# Repetitive transcranial magnetic stimulation induces protocol and layer specific transcriptomic plasticity in the human cortex

**DOI:** 10.64898/2026.04.23.720509

**Authors:** Rebecca C.S Ong, Alexander D Tang

## Abstract

Repetitive transcranial magnetic stimulation is a popular method of non-invasive brain stimulation used to study the human brain and treat neurological disorders. However, despite over three decades of use, the cellular mechanisms underlying its effects remain unknown which has limited its use. Using spatial transcriptomics on excised human cortical tissue, we characterized and mapped changes to gene expression following stimulation with two common stimulation protocols. We find that repetitive magnetic stimulation alters the expression of genes related to neuronal and glial plasticity mechanisms that were mostly cortical layer, cell type, and protocol dependent. These findings show that repetitive transcranial magnetic stimulation acts on multiple neural plasticity mechanisms simultaneously in the human brain, which to some extent, can be biased by the stimulation frequency used.

---

Brain stimulation has had longstanding use in neuroscience, both as a tool to study the nervous, but also as a treatment for several neurological disorders (*1*). Of the different techniques used today, repetitive transcranial magnetic stimulation (rTMS) has arguably gained the most popularity, expanding from an experimental treatment of depression in the 1990’s (*2*), to a staple tool in basic neuroscience to study cognitive processes, and as an approved treatment for several mental health conditions (*3*). During rTMS, electromagnetic pulses are delivered over the scalp to affect the underlying neural circuits, with the timing of the pulses (stimulation frequency) thought to be one of the most important stimulation parameters in determining the long-lasting effects on neural plasticity (*3, 4*). For example, it has been long thought that 10Hz stimulation and the complex pattern protocol of intermittent theta burst stimulation (iTBS – 2s trains of 3 pulses of 50Hz repeated at 5Hz) promotes neural activity, driving activity-dependent synaptic plasticity in the stimulated region (*5*). However, to date, support for synaptic plasticity as the cellular mechanism underlying the effects of rTMS in the human brain is limited to non-invasive neurophysiological or behavioral measures (*6, 7*). As such, there have been no direct measurements of the changes induced in human neural circuits following rTMS, limiting the interpretation of the effect of rTMS on cognitive processes, and the optimization of rTMS to improve its clinical efficacy.

Given that many neuronal and glial plasticity mechanisms are known to be activity-dependent (*8, 9*), and that rTMS acts non-discriminately with all cell types in the targeted brain region stimulated simultaneously, it is likely that rTMS acts on multiple neural plasticity mechanisms in the human brain that extend beyond the synapse. This is supported by fundamental research using laboratory animals where 10Hz and iTBS have been shown to induce synaptic (*10–13*), intrinsic (*14, 15*), and glial plasticity (*16–19*). Moreover, recent rodent studies suggest that the exact mechanisms of rTMS-induced plasticity depend not only on the stimulation frequency used, but on the layer of the cortex, and cell subtype (*20, 21*). For example, iTBS to the mouse sensorimotor cortex has been shown to act on multiple neuronal and glial plasticity mechanisms, but the specific changes varied between the different cortical layers of the sensory and motor cortex despite both brain regions receiving identical stimulation intensity. However, while animal and experimental models have provided the greatest insight into the cellular mechanisms of rTMS to date, the ultimate goal is to characterize the direct effect of rTMS on the human brain given the physiological and anatomical differences between species that may affect direct translation. To address this, excised neurosurgical tissue presents a novel method of studying the mechanisms of rTMS as excised cortical tissue retains physiological cortical layering, cellular function, and cellular heterogeneity (*22*) that can be combined with the advanced neuroscience techniques that are too invasive for human participants.

In this study, we used spatial transcriptomics on excised human neurosurgical tissue to characterize the direct effect of rTMS on the human cortex (Fig. 1A-C). Several acute brain slices were made from each tissue sample, allowing us to characterize and map changes to gene expression following the 10Hz, iTBS and sham for every tissue donor. In addition, a small number of whole cell patch clamp recordings were made from layer 2/3 pyramidal neurons in iTBS and sham stimulated.

**Fig. 1:**
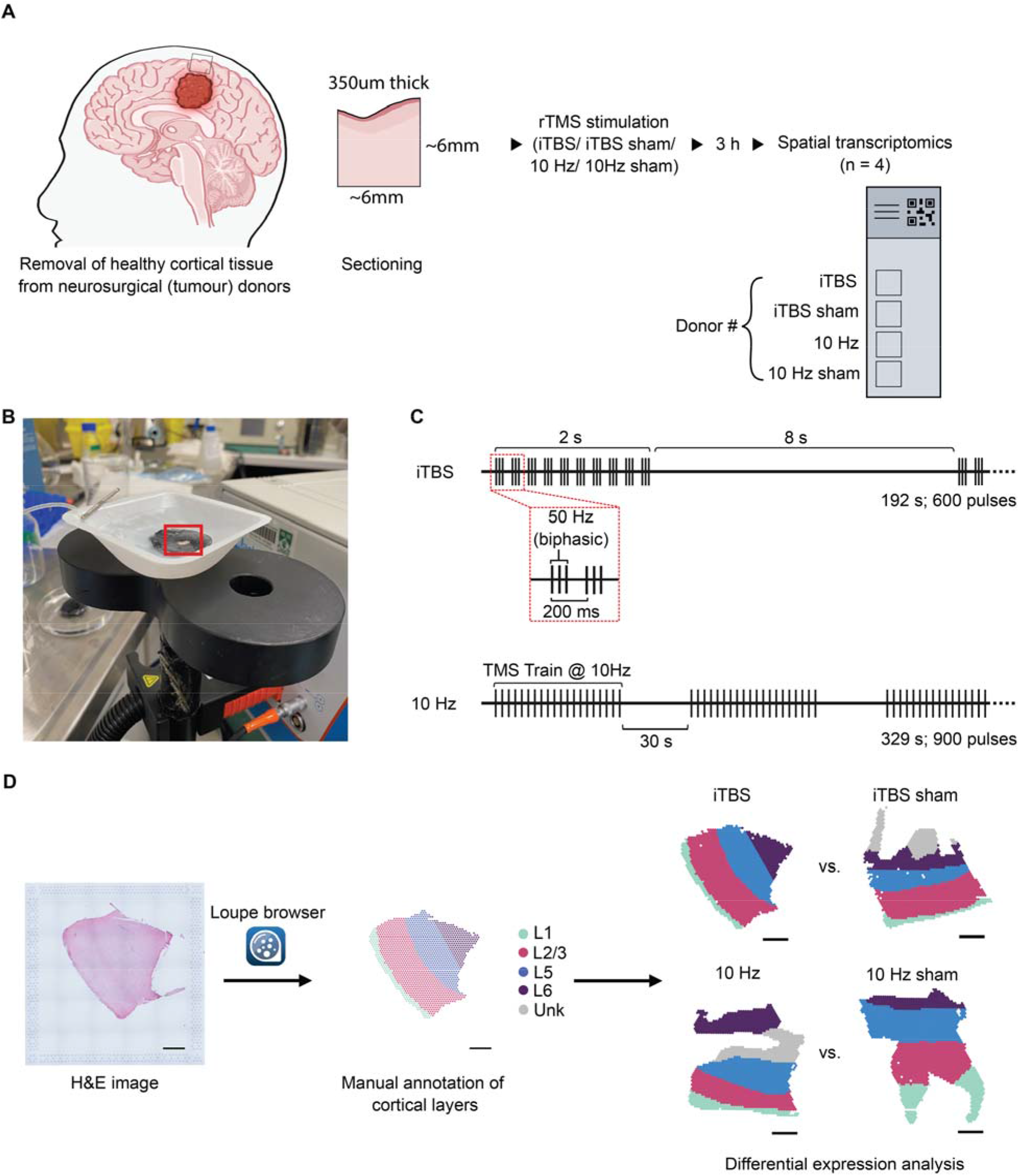
Overview of study design and repetitive transcranial magnetic stimulation (rTMS) protocols. **(A)** Experimental workflow to assess the effects of intermittent theta burst stimulation (iTBS) and 10 Hz rTMS on ‘live’ human cortical tissue. Cortical tissue sections obtained from each donor were used 3 hours following stimulation for spatial transcriptomics. **(B)** Photograph of stimulation setup. Cortical slices were stimulated using a standard 75-mm human TMS figure-of-eight coil. Slices were placed on a mesh surface situated in a weigh boat filled with 37°C artificial cerebral spinal fluid (ACSF) bubbled with carbogen and positioned directly on the center of the coil. **(C)** Schematic of the iTBS and 10 Hz rTMS protocols. **(D)** Manual subsetting of cortical layers from H&E images.

## Pseudobulk analysis reveals no uniform effect of rTMS on gene expression

We started by aggregating the transcriptomic profile of all tissue covered spots to form a pseudobulk analysis, similar to bulk RNA-Seq, to investigate if there were gross changes to gene expression irrespective of spatial location in the cortex. Principal component analysis showed no distinct clustering pattern amongst biological replicates of each stimulation group (Fig. S1A). Instead, samples obtained from the same tissue donor clustered together, indicative of the inherent inter-individual variability between donor samples (Fig. S1B). Accordingly, there were no significant differentially expressed genes (DEGs) following iTBS or 10Hz stimulation in the pseudobulk dataset.

## Spatial transcriptomics reveal cortical layer, and protocol dependent rTMS plasticity mechanisms

Given that each cortical layer has distinct structure, organization, and cell type composition, spatial transcriptomics was used to assess whether rTMS induced different changes in gene expression between the cortical layers, and if this varied between iTBS and 10Hz stimulation (Fig. 1D and Fig. S2). Due to the variability in the size of the samples resected and quality of tissue after cryosectioning, not all cortical layers were present in every sample. Therefore, our analysis was restricted to layers 1 and 2/3 for iTBS and iTBS sham, and layers 1, 2/3, 5, and 6 for 10Hz and 10Hz sham.

Due to the limited sample size (4 donors) and substantial variability in gene expression between each tissue donor (Fig. S2), we applied a conservative analysis approach by restricting our analysis to genes that showed large changes relative to sham from the same tissue donor (absolute log_2_FC ≥ 1) before computing the overlap between all tissue donors (Fig. 2A). Through this analysis, comparisons between iTBS and iTBS sham stimulated tissue identified 266 DEGs universally altered across three individuals in cortical layer 1, and 8 DEGs universally altered across four individuals in cortical layer 2/3. Interestingly, for the majority of genes identified, the directionality of expression change was not consistent between individuals, thus uncovering substantial inter-individual variability on the effect of rTMS at the transcriptomic level (Data S1 and S2), similar to measurements made at the network and behavioral level in human participants. Nevertheless, we used the National Center for Biotechnology Information (NCBI) human gene database as well as existing literature to determine the specific functional role of each DEG in the brain, if any. Specifically, genes could be manually assigned into one of nine different neural processes of interest based on their known biological function (Table 1; for the full list of DEGs with their corresponding annotations see Data S1 and S2). Within cortical layer 1, iTBS was found to regulate the greatest number of genes that had established roles in various immune and/or inflammatory responses within the brain. For example, genes encoding for antigens (e.g., *CD200* and *CD53)* known to often be involved in the activity of immune cells, and correspondingly, genes encoding for transcription factors (e.g., *NFATC2* and *NFATC3)* that regulate gene expression in human T cells were found to be differentially regulated in cortical layer 1 across all individuals (n = 3) following iTBS. Notably, whilst both *NFATC2* and *NFATC3* are classically known to be expressed in immune T cells, several rodent studies have further characterised their roles in regulating various glial cells in the brain, with *NFATC2* actively involved in oligodendrocyte differentiation (*23*) and microglial activation (*24*), and *NFATC3* in the regulation of astrocyte reactivity (*25*). Several genes with known functions in various components of the synapse and synaptic plasticity responses were also found to be altered following iTBS in cortical layer 1. Some examples (Fig. 2C, Fig S3) include *CACNG3* which encodes for the transmembrane AMPA receptor regulatory protein (TARP) involved in AMPA receptor trafficking (Fig. 4A), and genes involved in synaptic vesicle exocytosis (e.g., *STX1A* and *DOC2A)*. Additionally, we also identified genes involved in regulating various cellular mechanisms that mediate inhibitory neurotransmission, including alterations in the expression of two different GABA-A receptor subunits, *GABRA4* and *GABRG2* (Fig. 2C), and a notable marker of a subtype of inhibitory neurons, somatostatin *(SST)*. Within cortical layer 1, iTBS was also found to regulate the expression of genes involved in additional forms of plasticity including several encoding for various potassium channel components (e.g., *KCNH1, KCNIP1, KCNJ9*, and *KCNS2)* often involved in regulating membrane excitability, and genes involved in myelination (e.g., *PRRG1* and *MAG)*. Other neuronal processes that were also found to be altered following iTBS in cortical layer 1 included genes relating to neurogenesis (e.g., *ALKBH3* and *AHR)*, neurotrophic factors (e.g., *ADCYAP1)*, cell death (e.g., *BTBD10, PTPN4*, and *TRIM27)*, and mitochondrial function (e.g., *AKAP1)*. Across all individuals, iTBS was found to induce a lower number of significant gene expression changes in cortical layer 2/3 compared to cortical layer 1, with this reduced effect reflected in the total number of genes that were differentially regulated across all four tissue donors. Adopting the same approach to identify the known function of DEGs within cortical layer 2/3, only two out of the eight were found to have a well characterised role within the brain, with the gap junction protein gene *GJB1* expressed within myelin sheaths and transmembrane protein *WDR81* involved in regulating neurogenesis in the adult brain.

**Table 1:**
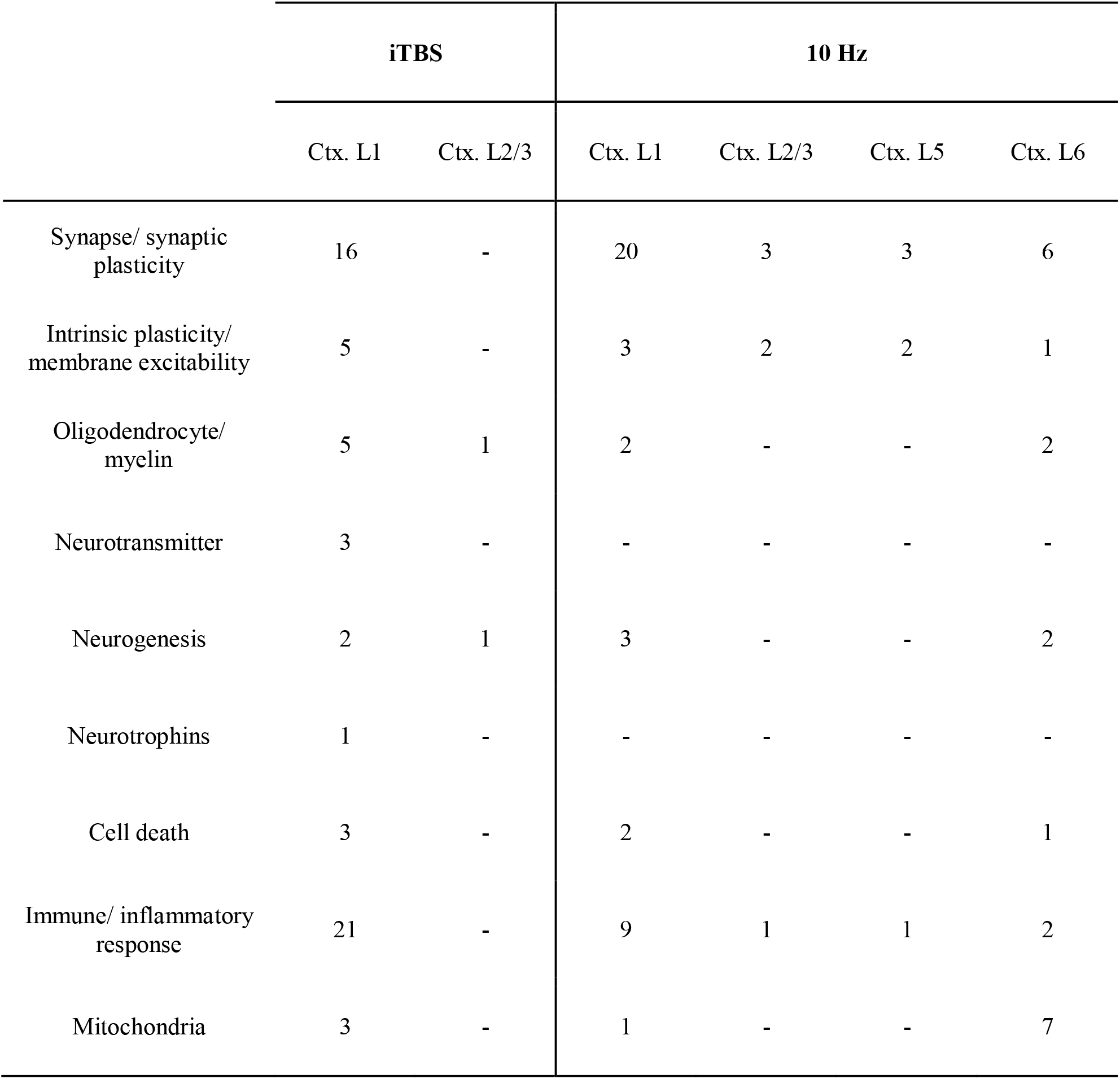
Higher-order functional category assignment for genes with a significant change in expression 3 h following iTBS, and 10Hz rTMS in different cortical layers of the human brain. Genes with a significant change in expression following iTBS, and 10 Hz stimulation in cortical layers 1-6 (Ctx. L1 – L6) were categorised into various neural processes, where possible, based on their known function in the literature. Significant genes were classified as having a log_2_ fold-change ≥ 1 within each individual and were also found to be differentially expressed across all individuals. Dashes indicate no identified differentially expressed genes in that category. Complete list of differentially expressed genes with their assigned functional classes can be found in Data S1 – S7.

**Fig. 2:**
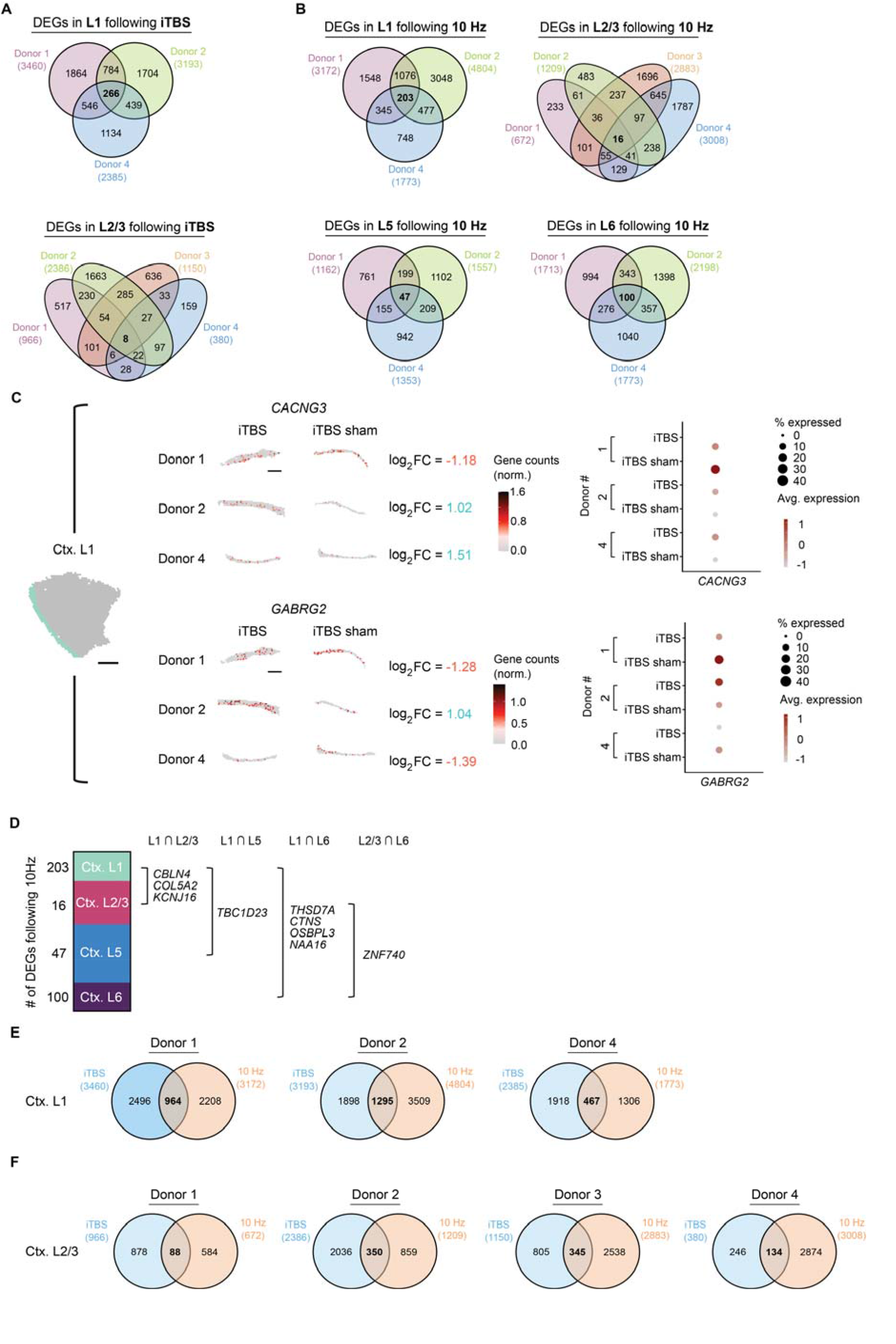
Identification of cortical layer-specific genes altered 3 hours following iTBS and 10 Hz rTMS. **(A)** Venn diagrams displaying the number of DEGs (absolute log_2_FC ≥ 1) altered for each individual in cortical layers 1 and 2/3 following iTBS and **(B)** in cortical layers 1, 2/3, 5 and 6 following 10 Hz rTMS. Numbers bolded in each venn diagram indicate the final number of DEGs considered to be altered by stimulation. **(C)** Example distribution of changes to gene expression relative to sham following iTBS in cortical layer 1, with *CACNG3* and *GABRG2* shown. **(D)** Of the significant DEGs following 10 Hz rTMS, some genes were found to be altered across two different cortical layers as visualised (∩ denoting overlap). No genes were found to be altered across more than two cortical layers following 10 Hz rTMS. No overlapping genes between cortical layers was found following iTBS. **(E)** Venn diagrams displaying the number of DEGs unique to 10Hz and iTBS, and their overlap for each tissue donor, in cortical layers 1 and **(F)** 2/3.

Similar to iTBS, we found the greatest number of genes universally altered by 10 Hz stimulation across three tissue donors was in cortical layer 1, with a total of 203 genes displaying a significant change in expression (absolute log_2_FC ≥ 1) compared to sham . Within the other cortical layers, 10 Hz stimulation induced a significant change in the expression of 16 genes common in cortical layer 2/3, and in cortical layers 5 and 6, we found 47 and 100 significant DEGs, respectively. Again, the directionality of expression change induced by 10 Hz stimulation varied between individuals, with approximately less than 40% of genes found to be universally up- or down-regulated within each cortical layer (i.e., 21/203 DEGs in cortical layer 1, 2/16 DEGs in cortical layer 2/3, 18/47 DEGs in cortical layer 5, and 10/100 DEGs in cortical layer 6 with the same regulation in expression) (Data S3-6). Assessing the functional implications of significant genes, we found alterations in the expression of genes relating to synaptic plasticity, intrinsic plasticity, and immune responses across all cortical layers following 10 Hz (Table 1; for the full list of DEGs with their corresponding annotations see Data S3-6). Surprisingly, there was little overlap amongst the specific genes expressed within each functional category across cortical layers, with only the cerebellin 4 gene *CBLN4*, and inwardly rectifying potassium channel gene *KCNJ16* found to be both differentially expressed in cortical layers 1 and 2/3 (Fig. S4). *CBLN4* encodes for a synaptic protein essential in the formation and maintenance of inhibitory GABAergic synapses (*26*), and in addition to *CBLN4*, several other synapse-specific genes within cortical layer 1 were also found to have a regulatory role in inhibitory interneuron synthesis, notably *DLX1* (*27*) and *ERBB4* (*28*). Alongside these inhibitory synapse-specific genes, parvalbumin *PVALB*, a notable gene marker for a subtype of GABAergic interneurons was also found to have a significant change in expression within cortical layer 1 following 10 Hz stimulation. Furthermore, 10 Hz stimulation modulated the expression of genes associated with various other synaptic functions including neurotransmitter release (e.g., *PI4K2A* and *DCAF12)*, long-term potentiation (e.g., *ACVR1C* and *ARHGEF6)*, dendritic branching (e.g., *PCDH11X*, and *KIFC3)*, and receptor regulation at the postsynaptic density (e.g., *NETO1*, and *ADAM22)* that were only affected in cortical layer 1. Beyond the synapse, 10 Hz stimulation was found to alter the expression of several genes encoding for several ion channels involved in regulating intrinsic neuronal excitability (e.g., *CACNA1C, KCNJ16*, and *NKAIN4)*, as well as genes expressed during active myelination (e.g., *GAB1* and *UGT8)*, and the microglial immune response (e.g., *CASP4* and *MOK)*. Within cortical layer 2/3, in addition to the modulation of *CBLN4* and *KCNJ16*, 10 Hz stimulation altered the expressed of synaptic (e.g., *FOXP2* and *GRIN3A)* and intrinsic (e.g., *GLRA3)* plasticity genes that were unique to the cortical layer. In both cortical layers 5, and 6, the few DEGs identified to play a role in regulating synaptic functions or intrinsic plasticity/ membrane excitability were unique to each cortical layer. However, in line with the aforementioned alterations of genes involved in different aspects of inhibitory cortical circuitry, we also identified changes in genes within both cortical layers 5 and 6 relating to several other inhibitory-related synaptic mechanisms. For example, the lysosomal-associated membrane protein gene *LAMP5*, expressed in a subtype of cortical GABAergic neurons, underwent a change in expression within cortical layer 5 whilst in cortical layer 6, 10 Hz stimulation altered the expression of the vasoactive intestinal peptide *VIP*, another known subtype marker of GABAergic neurons. Additionally, within cortical layer 6, we observed alterations in genes relating to oligodendrocytes (e.g., *MYO1D* and *ZBTB45)*, neurogenesis (e.g., *ALKBH3)*, apoptosis (e.g., *FAS)*, and mitochondrial functions (e.g., *GASP, METTL17*, etc.).

### iTBS and 10Hz show some overlap in DEGs at the individual level

In both basic and clinical neuroscience, iTBS and 10Hz stimulation are typically used for the same purpose, to increase neural activity and promote long-term potentiation like effects in the cortex. Therefore, we investigated the degree of overlap in DEGs between iTBS and 10Hz for each tissue donor in cortical layers 1 and 2/3 (i.e. layers where we had transcriptomic data for both protocols-Data S7). Whilst iTBS and 10Hz mostly altered the expression of different genes for each tissue donor, there was some degree of overlap (Fig. 1E and F). Interestingly, when looking at the overlapping DEGs for iTBS and 10Hz stimulation, there was a consistent trend in the proportion of DEGs related to specific neural plasticity mechanisms in all tissue donors, with ∼20% of the overlapping genes related to synaptic plasticity, ∼10% related to intrinsic plasticity, and ∼8 related to oligodendrocyte/myelin plasticity.

### Deconvolution provides insight into cell-specific effects of rTMS protocols across the human cortex

To gain insight into whether rTMS affects multiple cell types, and the specific genes in those cell types, we applied the robust cell-type decomposition (RCTD) model to our spatial transcriptomic dataset to enable the assignment of individual spots to cell-type groups using the cell-type specific enrichment percentage as a guide. Across all samples, we categorised spots as excitatory neurons (ExN), inhibitory neurons (InN), microglia, oligodendrocytes (Oligo), and/or astrocytes (Astro) if they contained at least 25% of the gene expression profile of that cell, defined using a published human prefrontal cortex snRNA-seq dataset (*29*) (Fig. 7A and S4). After excluding samples that had a poor representation of cell-type specific spots (i.e., less than 20% of the highest count for each cell-type category) and cell types that did not have a large number of cell specific spots, we were left with oligodendrocytes and astrocytes for the iTBS group, and excitatory neurons, microglia, oligodendrocytes, and astrocytes for the 10Hz group. We found that following iTBS, differential expression analysis identified 172 genes that had a significant change in expression across two individuals within oligodendrocyte-enriched spots, and 143 shared genes across two individuals within astrocyte-enriched spots (Fig 8A and B). To obtain a broad characterisation of the underlying effects of iTBS on each of the cell-types, significant DEGs identified were individually assessed for any potential functional relevance within their respective cell-type context. Through this, genes were manually assigned into higher-order functional categories where possible (Table 2; for the full list of DEGs with their corresponding annotations see Data S8 and S9). Within oligodendrocytes, iTBS was found to modulate the expression of several genes involved in regulating different aspects of the myelination process and/or cellular differentiation. For example, the myelin oligodendrocyte glycoprotein *MOG* encodes for notable protein involved in maintaining myelin structure and stability (*30*). In line with this, other genes identified within this functional category are known to have supporting roles both in controlling oligodendrocyte differentiation, thus resulting in the promotion of myelin sheaths by mature oligodendrocytes in the brain (e.g., *SMAD7, NPY*, and *BRCC3)* (Data S8). In addition to the oligodendrocyte-specific functional processes found to be altered following iTBS, we also found changes in genes primarily expressed in neural cells but have also been shown to impact oligodendrocyte function. An example of this is the postsynaptic density protein *SHANK3*, which displayed an up-regulation in expression across two individuals following iTBS. Other cellular processes identified amongst iTBS-induced DEGs in oligodendrocytes included genes involved in calcium signaling (e.g., *SLC8A3)*, mitochondrial function (e.g., *MTO1, MRPL39*, etc.), immune responses (e.g., *CD93* and *ABCC8)*, and cell death (e.g., *EPHB3* and *XAF1)* (Data S8). Within astrocytes, although 143 DEGs were identified to be altered following iTBS, many didn’t have a known astrocyte-specific function. Nonetheless, of the genes that were grouped into higher-order functional categories, several were found to be associated with astrocyte-specific immune responses (Table 2). In particular, the interleukin 1 alpha *IL1A*, and metallothionein 1H *MT1H* gene have both been implicated in mediating neuroprotective astrocytic responses (*31, 32*). Furthermore, another interleukin receptor gene *IL4R* was also identified to be differentially expressed and is known to play an immuno-protective role in inhibiting astrocyte activation and initiating the secretion of neurotrophic factors in astrocytes (*33*). Additionally, several genes encoding for ion channels (e.g., *TRPV2* and *KCNN2)* were also found to be altered in astrocytes following iTBS. The transient receptor potential channel *TRPV2* gene is involved in regulating intracellular calcium levels within astrocytes and has been thought to be a negative regulator of cell survival (*34*) whilst the potassium channel *KCNN2* specifically encodes for a small-conductance calcium-activated potassium channel, involved in modulating membrane excitability. Several mitochondrial-related genes were also found to be differentially regulated in astrocytes following iTBS.

Differential expression analysis of 10 Hz and 10 Hz-sham stimulated samples and comparisons across individuals identified a total of 203 DEGs in excitatory neurons that were shared across two individuals, 81 DEGs in microglia that were shared across four individuals, 93 DEGs in oligodendrocytes that were shared across two individuals, and 49 DEGs in astrocytes shared across two individuals (Fig 8C-F). Within excitatory neurons we identified the greatest number of DEGs with functional roles within the synapse and/or involved in regulating synaptic function (Table 2; for full list of DEGs and their corresponding annotations see Data S10). For example, we found 10 Hz-induced alterations in genes shown to encode for structural dendritic proteins (e.g., *AMOT*, C*UX2*, and *C1QL2)*, and various aspects of GABAergic inhibitory transmission (e.g., *LAMP5, NPAS4*, and *DGCR8)*. In addition to alterations at the synapse, genes encoding proteins associated with the regulation of neuronal excitability were also found to be altered within excitatory neurons following 10 Hz stimulation (Table 2 and Data S10). For example, the gene regucalcin *RGN* encodes for a protein involved in modulating the activity of membrane calcium pumps, thereby *influencing* the intracellular calcium concentrations within neurons (*35*) whilst similarly, the ring finger protein gene *RNF138* plays a role in regulating voltage-gated calcium 2.1 channel protein levels in neurons (*36*). We also identified changes to the expression of genes with known functions in adult neurogenesis (e.g., *ERBB3* and *FOXO4)*, myelination (e.g., *GPR17* and *CMTM5)*, immune responses (e.g., *GBP1, SOCS1*, and *ASCC3)*, cell survival (e.g., *DICER1, PLEKHO2, BIRC3*, etc.), and mitochondrial-related functions. Within microglia, 10 Hz stimulation was found to primarily induce alterations in the expression of immune response-related genes, several of which are known to contribute to the activation of microglia (e.g., *STAT1, STAT3* and *OGA)* (Table 2; for full list of DEGs and their corresponding annotations see Data S11). Alongside these gene changes, amongst the list of significant DEGs, we also identified alterations in the expression of a gene involved in the regulation of the GABA neurotransmitter uptake in microglia, syntaxin 1A *STX1A*, as well as a notable gene marker for activity-induced synaptic plasticity *ARC*. Within oligodendrocytes, we identified 10 Hz-induced changes in genes primarily involved in regulating oligodendrocyte differentiation (e.g., *YAP*1, *CLCN2*, and *LAMB1)* and several synapse-specific genes, including *LAMP5*, also found to be modulated in excitatory neurons following 10 Hz stimulation (Fig. 3G; for full list of DEGs and their corresponding annotations see Data S12). Whilst we also found genes associated with various other cellular processes including myelination, cell death, immune responses, and mitochondria amongst the list of 10 Hz-induced DEGs in oligodendrocytes, only a single gene was identified for each category. Within astrocytes, we predominantly identified changes in the expression of genes associated with immune and/or inflammatory-responses, particularly genes known to be regulators of astrocyte reactivity (e.g., *HAS1* and *KLRF11)* (Table 2; for full list of DEGs and their corresponding annotations see Data S13).

**Fig. 3:**
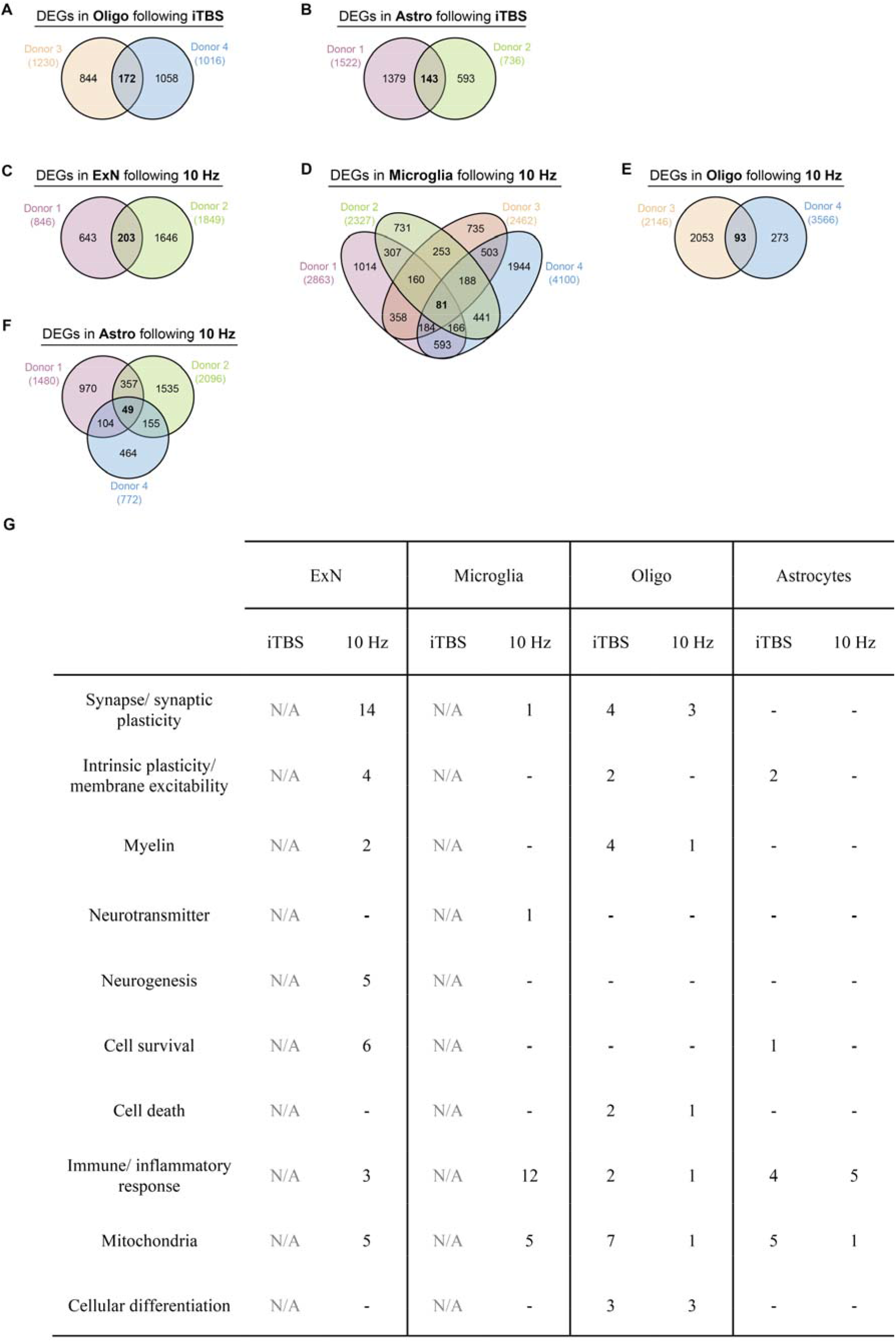
Number of cell-type specific differentially expressed genes (DEGs) following iTBS and 10 Hz rTMS following single cell deconvolution. For each individual that had sufficient cell-type specific spot numbers in both stimulated and sham stimulation control tissue following deconvolution, differential expression analysis was performed for each cell-type of interest within each individual. Cross-comparison of significant DEGs (absolute log_2_FC ≥ 1) was performed to identify the final list of cell-type specific DEGs considered to be altered by stimulation. Venn diagrams display the number of DEGs altered 3 h following iTBS specifically in **(A)** oligodendrocytes (Oligo), and **(B)** astrocytes (Astro). Following 10 Hz rTMS, venn diagrams display the number of DEGs altered specifically in **(C)** excitatory neurons (ExN), **(D)** microglia, **(E)** oligodendrocytes, and **(F)** astrocytes. Numbers bolded in each venn diagram indicate the number of DEGs common between all individuals used in analysis. **(G)** Higher-order functional category assignment of the DEGs in excitatory neurons (ExN), microglia, oligodendrocytes (Oligo), and astrocytes. Genes were categorized into different neural processes of interest based on their known function. Complete list of differentially expressed genes with their assigned functional classed can be found in Data S8 – S13. Note N/A reflects cell types that could not be resolved with the deconvolution algorithm.

**Fig. 4:**
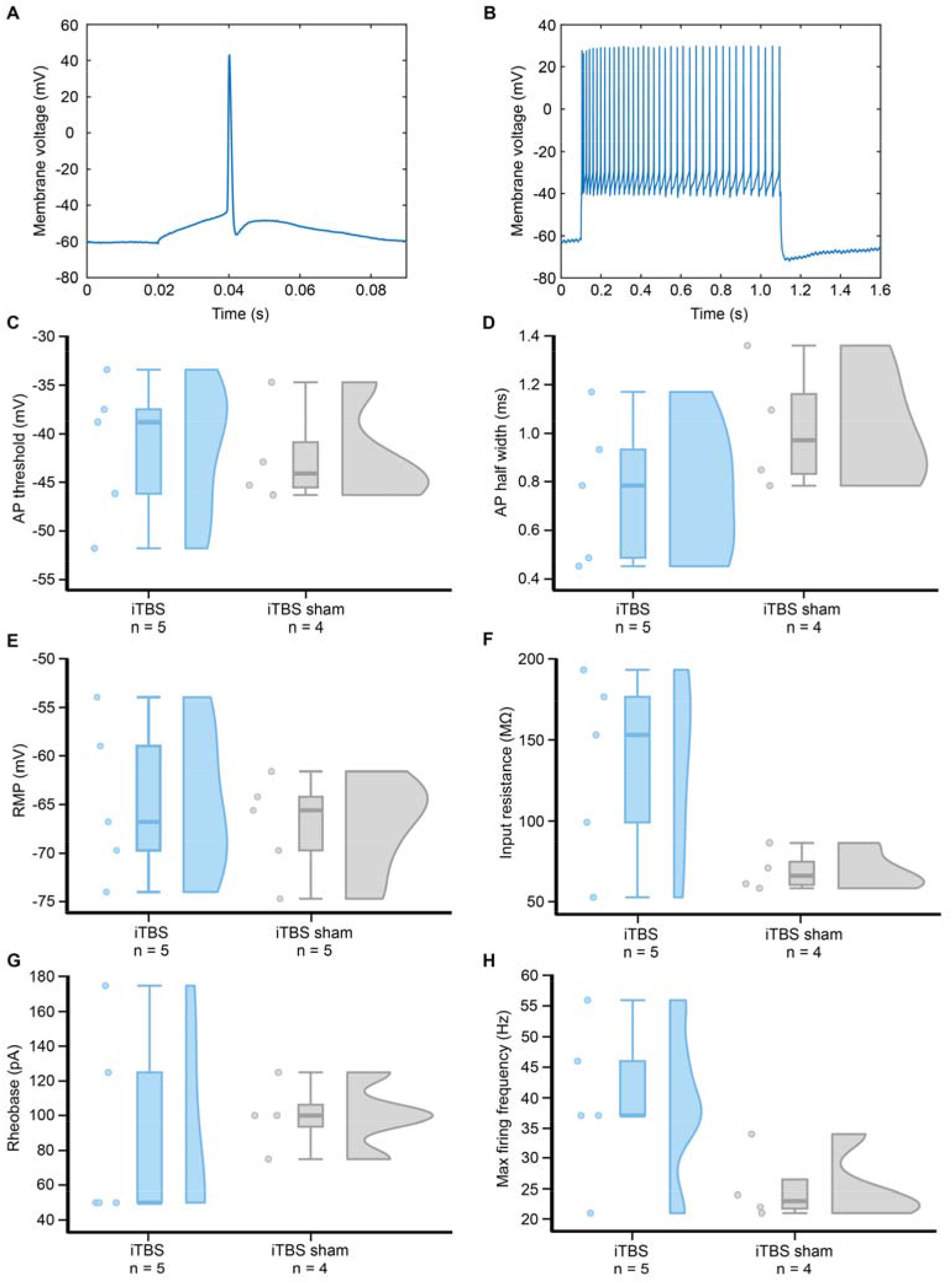
Whole-cell current clamp recording data from layer 2/3 pyramidal neurons following sham or iTBS. **A)** Representative trace of a single action potential (AP) of a human pyramidal layer 2/3 neuron in response to a 5 ms current injection. **(B)** Representative trace of the AP firing activity in response to a 1 s current injection. Plots depicting individual data points, box plots, and distributions for **(C)** AP threshold, **(D)** AP half-width, **(E)** resting membrane potential (RMP), **(F)** input resistance, **(G)** rheobase, and **(H)** max firing frequency of neurons treated with iTBS, and iTBS sham stimulation. For all measures, the number of cells analysed for each treatment group is presented below the treatment names. Cells were recorded from a total of n = number of individual cells, with at least one cell recorded from each tissue donor used for spatial transcriptomics.

## Patch clamp recordings provide preliminary evidence of iTBS induced intrinsic plasticity in human layer 2/3 pyramidal neurons

While the purpose of our study was to characterize and map the neural plasticity mechanisms of rTMS in the human cortex using spatial transcriptomics, an excess of iTBS stimulated slices allowed us to perform a small number of whole-cell current clamp recordings to assess changes to excitability and intrinsic properties in layer 2/3 pyramidal neurons. As we only recorded from 5 iTBS active and 5 iTBS sham cells (at least one cell per tissue donor used for spatial transcriptomics), we only report central tendency measures (Fig 4; for summary of raw patch-clamp data collected from each cell see Data S14).

In our data, we did not observe large changes to resting membrane potential, action potential threshold, rheobase, or action potential half-width. We did however observe large differences in input resistance (134.90 ± 57.96 MΩ iTBS active vs 69.24 ± 10.96 MΩ iTBS sham) and the evoked maximum firing frequencies (39.40 ± 12.93 Hz iTBS active vs 25.25 ± 5.96 Hz iTBS sham). Taken together, these results provide preliminary support for an increase in neuronal excitability through intrinsic plasticity three hours post-iTBS.

## Discussion

rTMS has been used widely since the 1990’s, yet the cellular mechanisms underlying its effects on the human brain, and how they are influenced by different stimulation protocols remain unclear. Through spatial transcriptomics, we found that rTMS alters acts on multiple neuronal and glial plasticity mechanisms simultaneously, which to a large degree, were cortical layer and stimulation frequency dependent. Supporting this, deconvolution for cell type specific analysis suggested that rTMS alters plasticity related genes in both neurons and glia. Furthermore, a small number of patch clamp recordings from iTBS stimulated layer 2/3 neurons provided preliminary support for increased input resistances and evoked maximum spike firing frequencies, providing further evidence of iTBS induced intrinsic plasticity three hours post-stimulation.

While the longstanding assumption within the field has been that rTMS acts on synaptic plasticity in the human brain (*6, 37*), our transcriptomic data extends this by showing rTMS also acts on intrinsic and glial plasticity mechanisms. These findings are consistent with the mechanistic data from laboratory animals (*10, 11, 14, 12, 15–17, 13, 19, 38*), both in the types of neural plasticity mechanisms implicated, but also that rTMS induced plasticity is cell type, cortical layer, and stimulation frequency dependent (*20, 21*). Interestingly, while 10Hz and iTBS protocols acted on multiple plasticity mechanisms, we found that iTBS had a greater effect on oligodendrocyte/myelin plasticity and immune/inflammatory related genes, whereas 10Hz was more biased to synaptic plasticity related genes. This may suggest that iTBS is more suitable for studying and modulating oligodendrocyte/myelin plasticity in the human brain than 10Hz, which is supported by data in mice with iTBS having a stronger effect on oligodendrocyte/myelin plasticity gene expression than cTBS (*20*), and the only protocol to promote the survival of adult born oligodendrocytes (*16*). Ten hertz on the other hand may be most suited when the primary purpose of rTMS is to bias the desired effect to synaptic plasticity with minimal action on other plasticity mechanisms. However, it should be noted that the different neural plasticity mechanisms observed in our study are unlikely to work in isolation, and to some extent, support each other. For example, we speculate that increases to neuronal activity via intrinsic plasticity supports synaptic and myelin plasticity which are well known to be activity-dependent (*39, 40*). Similarly, it is already known from organotypic rodent slice cultures that 10Hz induced synaptic plasticity require an increase in cytokine release from microglia (*19*). Therefore, while different rTMS protocols may bias towards specific neural plasticity mechanisms, it is most likely that a synergistic effect is required to some degree.

One of the interesting outcomes of our study was the variability in changes to gene expression following rTMS, not only between individual tissue donor samples, but within tissue donor samples to iTBS and 10Hz. While it is well established, and often a criticism, that rTMS has large inter-individual variability in its neurophysiological (e.g., motor evoked potentials) and behavioral outcomes (*41–43, 6*), it was unknown whether similar variability occurs at the molecular level. We find that the effect of rTMS at the transcriptomic level not only varies in the specific genes it affects between individuals, but that even for commonly altered genes, there was variability in the direction of the change. While we can only speculate on the functional and behavioral outcome of such gene transcriptomic variability following rTMS, it may explain in part why rTMS can have opposite effects on neurophysiological and behavioral measures from one individual to another. However, it is worth noting that within each tissue donor sample we found that 10Hz and iTBS consistently had a greater number of unique DEGs compared to the total number of overlapping DEGs, providing further support for the protocol-dependent mechanisms of rTMS we saw at the group level. Furthermore, despite the total number of DEGs altered by each protocol varying substantially between each tissue donor, characterization of the overlapping DEGs between 10Hz and iTBS for each tissue donor showed a similar proportion of genes related to the different neural plasticity mechanisms. This would suggest that while 10Hz and iTBS induce protocol-dependent effects, that show wide variability at the molecular level, the proportion of overlap between the two protocols within an individual is less variable.

While we have characterized and mapped the neural plasticity mechanisms induced by two common rTMS protocols in the human cortex, it is important to note the key limitations of our interpretation. As detailed elsewhere (*20*), it is highly likely that rTMS-induced plasticity varies with time post-stimulation, with an immediate effect of rTMS not mediated by changes to gene expression, whereas the longer lasting changes that develop over hours and days likely relying on changes to gene expression. Furthermore, neurophysiological measures in humans (*44*), and spatial transcriptomic data (*21*) have suggested that the mechanisms of rTMS are impacted by normal aging, with aged mice showing less changes to gene expression following rTMS, which to some extent is stimulation frequency dependent (*21*). Lastly, while our study provided insight into inter-individual variability at the transcriptomic level in response to 10Hz and iTBS, as well as the genes that showed large changes in expression (absolute log2FC>1) consistently across our tissue donors, much larger sample sizes are needed to perform more in depth statistical testing to determine true “group level” transcriptomic changes in human neural circuits following rTMS.

## Acknowledgments

The authors thank the Sir Charles Gairdner Neurosurgery Department for identifying suitable participant and tissue collection for the study. OpenAI (ChatGPT) was used to identify genes of interest related to synaptic, intrinsic, and myelin plasticity from the list of overlapping DEGs between 10Hz and iTBS for each individual tissue donor that was then interpreted by the authors.

## Funding

Provide complete funding information, including grant numbers, complete funding agency names, and recipient’s initials. Each funding source should be listed in a separate paragraph.

2022 NARSARD Young Investigator Grant - ADT

Future Health and Research Innovation Near Miss Award – ADT

Sarich Family Research Fellowship - ADT

Australian Rotary Health PhD Scholarship – RCSO

University of Western Australia - RCSO

## Author contributions

Conceptualization: RCSO, ADT

Methodology: RCSO, ADT

Investigation: RCSO, ADT

Visualization: RCSO

Funding acquisition: RCSO, ADT

Project administration: ADT

Supervision: ADT

Writing – original draft: RCSO, ADT

Writing – review & editing: RCSO, ADT

## Data, code, and materials availability

The analysis scripts used in this manuscript can be found at https://github.com/rebecca-ong/rTMS-spatial. The data for this study have been deposited in the Gene Expression Omnibus (GEO) database (GSE328945). All data needed to evaluate the conclusions in the paper are present in the paper and/or the Supplementary Materials.

