## Supplementary material for "Repetitive transcranial magnetic stimulation induces protocol and layer specific transcriptomic plasticity in the human cortex"

#### Materials and Methods

Participant recruitment and details

All human brain tissue experiments were approved by the Sir Charles Gairdner and Osborne Park Health Care Group Human Research Ethics Committee (RGS0000005422) and UWA Human Ethics Office (2022/ET000645). Human neocortical tissue was obtained from donors undergoing tumor resection surgery where excess 'healthy' cortical tissue was removed to access the lesion. Inclusion criteria included adults ≥ 18 years of age of any gender or race/ethnicity with prior arrangements with the Sir Charles Gairdner Hospital neurosurgery department to undergo brain resection surgery. Individuals were excluded if they were unable to provide consent themselves or had previous neurological conditions, or any current neurological conditions that were unrelated to their reason for surgery. All cases were confirmed to be suitable by the consultant neurosurgeon responsible for each participant and included a review of neuroimaging by the surgical team to rule out swelling or possible lesions in the tissue sample collection area. Written informed consent was obtained from all donors prior to the surgery. Following exclusion of samples with potential tumour infiltration or poor-quality following slicing, data from 4 male tissue donors were included for sequencing and analysis (see Data S15 for age and brain region details).

Human brain tissue collection

Tissue was sharp dissected with minimal to no coagulation and was placed in a 50 ml falcon tube filled with ice-cold, recently carbogenated cutting artificial cerebral spinal fluid (ACSF) solution composed of (in mM): 110 Choline chloride, 26 $NaHCO_3$, 0.5 $CaCl_2$, 7 $MgCl_2$, 2.5 KCl, 1.25 $NaH_2PO_4$, 10 Glucose, 11.6 Na-ascorbate, and 3.1 Na-pyruvate (*45*). To ensure that there was minimal delay between the carbogenation of the solution and collection of the sample, a disposable nitrile glove filled with carbogen (5% $CO_2$/ 95% $O_2$) connected to a 3-way valve and a perfusion line, as designed by Straehle et al. (*46*), was used to constantly bubble the ACSF during transportation from the laboratory to the surgical theatre. Immediately following sample collection, the sample was transported on ice to the laboratory (~ 10-15 mins) for all following experimental procedures.

Slice preparation

Brain tissue samples were sectioned on a vibratome (Campden Instruments 7000 SMZ-2) in ice-cold carbogenated cutting ACSF solution. Slices of 300 μm thickness were cut perpendicular to the pial surface to capture all cortical layers using a ceramic blade and immediately transferred to a storage chamber filled with recovery ACSF solution containing (in mM): 125 NaCl, 25 $NaHCO_3$, 5 HEPES, 1 $CaCl_2$, 6 $MgCl_2$, 3 KCl, 1.25 $NaH_2PO_4$, and 10 Glucose bubbled with carbogen (5% $CO_2$/ 95% $O_2$). Slices were incubated in recovery ACSF at 35°C for 30 mins. Following recovery, sections were transferred to a separate storage chamber filled with normal ACSF solution composed of (in mM): 125 NaCl, 3 KCl, 1.25 $NaH_2PO_4$, 25 $NaHCO_3$, 0.7 $MgCl_2$, 1.2 $CaCl_2$, 10 Glucose bubbled with carbogen (5% $CO_2$/ 95% $O_2$). Slices were held in normal ACSF at 35°C for the remainder of the experiment.

rTMS parameters and administration

For rTMS stimulation, tissue sections were first transferred to a synthetic membrane situated in a square weigh boat containing warmed (~35°C) normal ACSF constantly bubbled with carbogen (5% $CO_2$/ 95% $O_2$). Tissue sections were stimulated using a MagPro R30 Stimulator (MagVenture) connected to a flat figure-of-eight coil (75 mm outer diameter; MC-B65-HO-2 Butterfly Coil; MagVenture). Tissue sections were placed directly on the center of the coil (i.e., intersection between the two wings), where the electric field strength would be at its maximum. Stimulation was delivered as biphasic pulses and two different protocols of active rTMS was tested: high-frequency 10 Hz stimulation (9 trains of 100 pulses delivered at 10 Hz, with an intertrain interval of 30 s, equivalent to 900 pulses (*10*)) and iTBS (2 s train of TBS [3 pulses delivered at 50 Hz repeated in 5 Hz intervals] with an intertrain interval of 8 s, equivalent to 600 pulses (*47*)). Parameters for 10 Hz rTMS were set directly through the MagPro R30 Stimulator to deliver iTBS, an external waveform generator (33500B; Agilent Technologies) containing the pulse pattern was used to trigger the MagPro R30. Stimulation intensity was set to 43% of maximum stimulator output, which was determined based on the max intensity at which 10 Hz rTMS and iTBS could be delivered without the coil temperature exceeding 41°C. As controls, sham stimulation for both 10Hz rTMS and iTBS was performed on slices from the same participant for the equivalent duration for each stimulation protocol (i.e., 10Hz rTMS sham for 329 s, iTBS sham for 192 s) under the same conditions but received no active stimulation. Immediately following stimulation, slices were returned to the storage chamber containing normal ACSF at 35°C for at least 3 hours before subsequent experimental procedures.

Induced electric field values are the most accurate measure of rTMS intensity. Therefore, to estimate the intensity of the electric field induced in the human brain slices with the MagVenture B65 butterfly TMS coil, finite element modelling in MATLAB (2025a) was done on a human brain slice (6mm x 6mm x 0.35mm) immersed in a saline bath (30mm x 30mm x 10mm). The slice was modeled as a thin tetrahedral mesh with conductivities of 0.40 S/m (cortical tissue) and 1.65 S/m (saline). The coil was positioned 0.2 mm below the base of the saline bath and was represented as two circular windings (10 turns per wing) with inner and outer diameters of 27 mm and 97 mm, respectively, and a winding height of 6 mm. The Biot–Savart law was used to compute the E-field at each mesh node with the fields scaled to 41% of maximum stimulator output (dI/dt = 161 kA/s), yielding realistic electric field magnitudes within the slice (25.56 V/m – fig S9). As expected, the estimated induced electric field was lower than whole-head simulations due to reduced tissue volume, boundary effects of the model, and the reduced MSO% needed to maintain coil temperature for both stimulation protocols.

Sample preparation & sequencing

Individual brain tissue slices were embedded in optical cutting temperature (OCT) medium (Scigen) in a 10 mm x 10 mm mold, ensuring that each slice laid as flat as possible, and were snap-frozen using liquid nitrogen-cooled isopentane. From each donor, brain samples treated with 10 Hz rTMS, iTBS, and the corresponding sham stimulation for each protocol (i.e., iTBS sham and 10Hz sham) were frozen and stored at -80°C until required. Fresh frozen samples were cryosectioned (CryoStar NX70, Thermo Fisher Scientific) at 10 µm thickness for all Visium spatial transcriptomics experiments. Prior to mounting cryosections on Visium slides, approximately 10-15 sections were collected from samples to assess RNA quality. The RNeasy Mini Kit (Qiagen) and RNase-Free DNase Set (Qiagen) was used to extract RNA, according to manufacturer's instructions, and quantified using a LabChip GX Touch Nucleic Acid Analyzer. Only samples that had an RNA Integrity Number (RIN) > 7.5 were used in subsequent Visium workflows.

To determine the optimal permeabilisation time that would maximise the mRNA yield from tissues, several sections (10 µm) from additional, un-stimulated brain tissue from two of the four donors were mounted onto Visium Spatial Tissue Optimization (TO) slides and processed according to manufacturer's instructions. In brief, TO slides were incubated with a permeabilisation enzyme for various timeframes (6, 10, 12, 14, 15, 16, 18, 20, 25, 30 mins) before captured RNA was reverse transcribed to cDNA and tagged with fluorescently labelled nucleotides. Enzymatic removal of tissue was performed prior to visualization of fluorescent cDNA using a Nikon Eclipse Ti2-E inverted microscope. In our optimization samples, 18 mins produced the greatest fluorescent cDNA intensity with the lowest signal diffusion and thus, was used for all subsequent Visium Spatial Gene Expression experiments.

One 10 µm section from each of the different stimulation-treated brain samples was mounted onto a Visium Spatial Gene Expression slide (n = 4) and processed according to manufacturer's instructions. Stimulated and sham sections from each donor were placed on the same slide. H&E staining was performed, and sections were imaged using a Nikon Eclipse Ti2-E inverted microscope at 10x magnification (numerical aperture = 0.45, 2424-pixel by 2424-pixel resolution). Raw bright-field images were processed in FIJI Image J. Following H&E imaging, tissue samples were permeabilized for 18 mins, before reverse transcription was performed and the cDNA collected from the slide. Sequencing-ready libraries were constructed from each cDNA sample and indexed using the Dual Index Kit TT Set A (10X Genomics).

A total of 16 dual-indeed, paired-end cDNA libraries sequenced on a NovaSeq 6000 instrument (Illumina) at the Australian Genome Research Facility (AGRF; Melbourne, Australia). Samples were sequenced at a minimum depth of 50,000 read pairs for each spot covered by tissue using a special sequencing read configuration as outlined in the Visium protocol: read 1, 28 bp; i7 index, 10 bp; i5 index, 10 bp; read 2, 90 bp. An average depth of 427,469,701 reads was obtained per sample.

<u>Spatial transcriptomics data processing and QC</u>

Using the Visium Lope Browser 7 application, manual spot alignment was performed for each sample to ensure that all spots that overlaid the tissue were selected and the output .JSON file was used for subsequent analyses. Raw sequencing FASTQ data files were aligned to the *Homo sapiens* GRC38 reference genome and along with the corresponding H&E images, each sample was processed using the 10x *SpaceRanger* pipeline (v3.1.3; 10X Genomics). Outputs of *SpaceRanger* include QC metrics (available in Data S1) and a summarized count table of genes within each tissue-covered spot [i.e., unique molecular identified (UMI) counts]. For each sample, the summarized counts table and corresponding H&E images were processed in R (v4.5.1) and stored in a *Seurat* object using the *STUtility* package (v1.1.1) (fig. S10). The *scuttle* package (v1.18.0) was used to calculate additional quality control metrics of spots, including mitochondrial expression, low library size, and low gene detection. Spots that had low library sizes and low number of gene counts which fell beyond a 3x median absolute deviation (MAD) threshold were discarded. As rTMS has previously been shown to modulate various mitochondrial functions (*48*), mitochondrial expression rates were not used as a quality control metric to ensure that a comprehensive understanding of the effects of rTMS on mitochondria in human neural cells is also obtained. A total of 149 spots were removed, resulting in a filtered dataset of 25,733 genes across 18,985 spots. Counts were normalised to account for variability in sequencing depth/library size across samples and batch-corrected for the effect of different donor tissues using the SCTransform function in *Seurat* (v5.3.0) (fig S11).

Unsupervised clustering analysis

Unsupervised clustering of spatial transcriptomic data was performed using PRECAST (v1.7) (*49*), a package that takes into consideration the spatial embeddings [i.e., the reduction of high-dimensional spatial transcriptomics data into a lower 2-dimensional space that captures the spatial relationship and gene expression patterns of each spot (*50*)] of different tissue sections to achieve a better clustering accuracy across samples. To assess the optimal number of clusters that would best delineate different regions across the tissue, PRECAST was run using $k = 2$ through to $k = 15$. The Akaike information criterion (AIC) method was used to compare each of the clustering models and evaluate the best-fitting model for the data, with models that have lower AIC scores considered to be the better estimates of spatial clustering. As AIC scores from $k = 5$ onwards were considerably lower than the preceding clustering models (Fig. S_ ; AIC plot), only clustering groups from $k = 5$ through to $k = 15$ were plotted across all tissue samples. Each clustering resolution was visually assessed to identify the number of clusters that most accurately reflected different tissue regions with the lowest number of clusters.

Pseudobulking spatial transcriptomics data

To assess the global effects of stimulation (i.e., independent of cell type and cortical layer), all tissue-covered Visium spots were pseudobulked within each sample by totaling the raw gene expression counts across each spot. The DESeq2 package (v1.48.2) was subsequently used to perform differential expression analysis of the pseudobulked samples. To visualise the data, raw counts were normalised using the variance stabilising transformation (VST) and dimensionality reduction was performed using the principal component analysis (PCA). Differential expression analysis using the Wald test was performed between tissues treated with each stimulation protocol and the corresponding sham tissue samples (i.e., iTBS vs. iTBS sham, and 10Hz vs. 10Hz sham) via the *DESeq* function. *P*-values for each gene was adjusted for multiple comparisons using the Benjamini-Hochberg false discovery rate correction (*51*). Genes were determined to have undergone a significant change in expression if they had an adjusted *P*-value (p.adj) of $\leq 0.05$ and absolute $\log_2$-transformed fold change (log2FC) $\geq 1$. Data plots were generated using *ggplot2* (v4.0.0).

Manual annotation of cortical layers

To conduct a cortical layer-level analysis of the effects of stimulation, each sample was manually annotated, using the Loupe Browser application (v9.0) by 10X Genomics, to identify spots that overlaid different cortical layers based on each corresponding H&E image. Manual annotation of cortical layers was independently performed by both authors and cross checked to ensure an accurate identification of each region. Where possible, tissue samples were annotated to identify cortical layer 1 (Ctx1) , cortical layer 2/3 (Ctx2/3), cortical layer 5 (Ctx5), and cortical layer 6 (Ctx6). Following manual annotation, spots were pseudobulked into layer-level data by summing the raw gene expression counts across all spots within each cortical layer for each given donor.

Differential expression analysis was performed between stimulated vs. sham samples within the same donor to identify genes altered by iTBS and 10 Hz rTMS that are specific to each individual. As previously mentioned, the *FindMarkers* function was used for differential expression analysis. Similarly, genes were only considered to have undergone a significant change in expression if the absolute log2FC was ≥ 1. Once significant differentially expressed genes were identified for each layer within each donor, cortical layer-specific gene lists were compared to identify genes that overlapped across all four donors. Data plots were generated with *ggplot2* (v4.0.0) or visualization tools in *STUtility*.

<u>Single-cell RNA-seq deconvolution</u>

As individual Visium spots often span multiple of the same and/or different cell types, we performed spot deconvolution using a recently published single nuclei RNA-seq (snRNA-seq) dataset by Jeffries et al. (*29*) to predict the likely cellular composition of each spot. The dataset by Jeffries et al. (*29*) included isolated nuclei obtained from 19 fresh-frozen human prefrontal cortex (PFC) tissue donors ranging from the ages of 20 weeks to 104 years old. Specifically, we used the transcriptomic profiles of seven different cell type clusters annotated in the study: excitatory neurons (ExN), inhibitory neurons (InN), microglia, astrocytes (AST), oligodendrocytes, oligodendrocyte precursor cells (OPC) and endothelial cells. The robust cell type decomposition (RCTD) algorithm (*52*) was used to predict the cell-type proportions of each spot via the *spacexr* (v1.0.0) package. RCTD was set to run using a 'multi' model to allow for any number of cell types to be assigned per spot without constraint. The output table containing the proportions of each cell type identified per spot was used to subsequently categorize spots across all samples into different cell types of interest. Specifically, spots that expressed at least 25% of the gene expression profile of either excitatory neurons, inhibitory neurons, microglia, oligodendrocytes, or astrocytes were correspondingly categorised into one or more of each cell type group.

To assess the cell type specific effects of stimulation, differential expression analysis was performed on each cell type of interest. Only samples that contained a minimum of 20% of the highest spot count number for each specific cell type were included in the analysis. The *FindMarkers* function in Seurat was used to identify differentially expressed genes within each donor (i.e., comparing stimulation vs. stimulation control tissue from the same individual) for both iTBS and 10Hz rTMS. In brief, a Wilcoxon Rank Sum test was applied to generate *P* values that were adjusted for multiple comparisons using the Bonferroni correction method. As differential expression analysis was conducted within the same biological sample, we only defined genes that had an absolute log2FC ≥ 1 change in expression as significant. Subsequently, each list of significant differentially expressed genes for each cell type was compared across all four donors, and genes that were present in all donors were considered to be genes that had undergone a significant change in expression following stimulation.

Whole cell patch-clamp electrophysiology

For whole cell current-clamp recordings, slices were placed in a chamber constantly perfused with normal ACSF solution (~1.1 ml/mins) and maintained at 35 ± 2°C (Warner Instruments TC-324B). Borosilicate glass pipettes (1.5 mm outer diameter x 0.86 mm inner diameter; Harvard apparatus GC150F-15) were pulled using a micropipette puller (Sutter Instruments P1000) and had an open tip resistance of 4-7 MΩ. Recording pipettes were filled with an internal solution composed of (mM): 135 potassium gluconate, 10 HEPES, 7 NaCl, 2 $Na_2$-ATP, 0.3 $Na_3$-GTP, and 2 $MgCl_2$.

Somatic recordings were obtained from human layer 2/3 pyramidal neurons using an Olympus BX51-W1 microscope equipped with an Oxford Instruments Andor Zyla 4.2 camera and visualised at 60X magnification under bright field and infrared differential interference contrast video microscopy. Resting membrane potential (RMP) was immediately recorded upon access into the cell, in current-clamp mode with no holding current. Cells were excluded if the series resistance exceeded 25 MΩ or if the series resistance changed by more than 20% of the initial value

Several evoked measures were obtained from each cell, where possible. A series of 1 s current steps increasing by 25 pA increments, ranging from -100 pA to +500 pA was applied to the cell. Input resistance was calculated from the successive steady state voltage responses produced from the first four negative current steps (i.e., ranging from -100 pA to -25 pA). Rheobase was defined as the first current step that evoked at least one action potential (AP). For the duration of each current step, the number of APs was calculated by counting the number of peaks that passed a 10-mV membrane potential threshold. Maximum number of APs for each cell was obtained by determining the current step at which the most peaks were produced.

For single AP measures, a 5 ms depolarising current step with increasing 25 pA increments was applied until a single AP was induced. Measurements from two single AP were averaged and used to determine AP properties. AP threshold was defined as the membrane voltage where dV/dt exceeded 10 V/s during the rising phase of the AP. AP half-width was defined as the time between the point of AP threshold to the point where the AP reached half of its peak amplitude. AP amplitude was defined as the voltage difference between the AP threshold and AP peak. Patch clamp data was analysed in SutterPatch (V2.3.1) and MATLAB (2025a), with central tendency measures plotted using JASP (0.95.4).

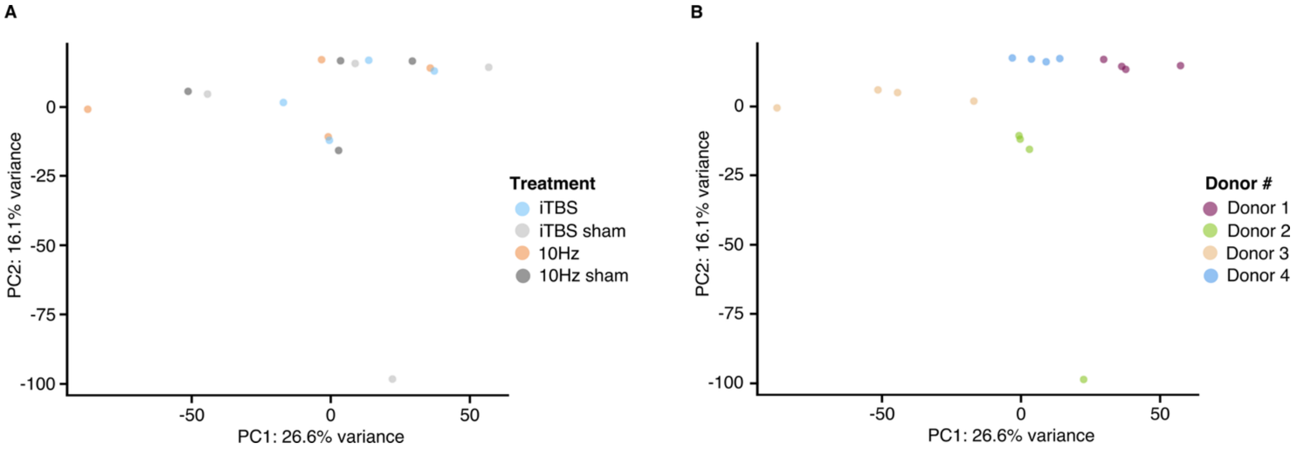


**Fig. S1: Principal component plots of pseudobulked spatial transcriptomics samples.** Spots from each Visium capture area were pseudobulked by summing the total UMIs from each sample. A two-dimensional principal component analysis (PCA) plot of the pseudobulked samples coloured by **(A)** stimulation treatment group and **(B)** donor captures the inter-individual variability present between samples, with no distinct clustering amongst stimulation groups.

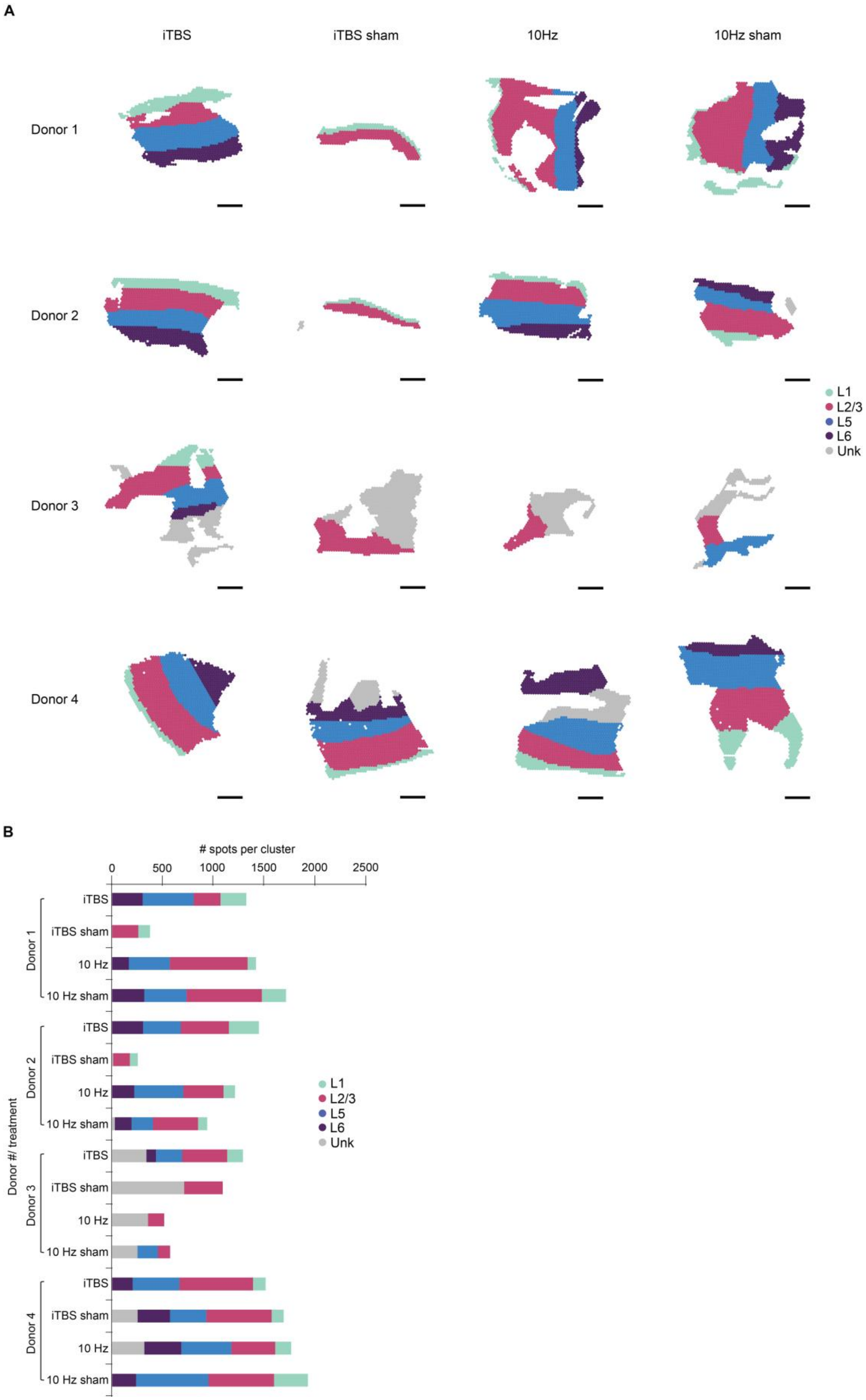
A
iTBS
iTBS sham
10Hz
10Hz sham
Donor 1
Donor 2
Donor 3
Donor 4
L1
L2/3
L5
L6
Unk
B
### spots per cluster
0
500
1000
1500
2000
2500
Donor #/ treatment
Donor 1
Donor 2
Donor 3
Donor 4
iTBS
iTBS sham
10 Hz
10 Hz sham
L1
L2/3
L5
L6
Unk

**Fig. S2: Manual annotation of cortical layers. (A)** All Visium tissue samples (n = 16) were manually annotated to identify each of the different cortical layers, guided by the H&E histology images. Each spot was annotated to either cortical layer 1 (L1), cortical layer 2/3 (L2/3), cortical layer 5 (L5), cortical layer 6 (L6), or as ‘unknown’ if the underlying tissue was tumour or there was a lack of information to accurately categorise spots. All scale bars represent 1 mm. **(B)** Number of visium spots for each cortical layer, split by stimulation group and tissue donor.

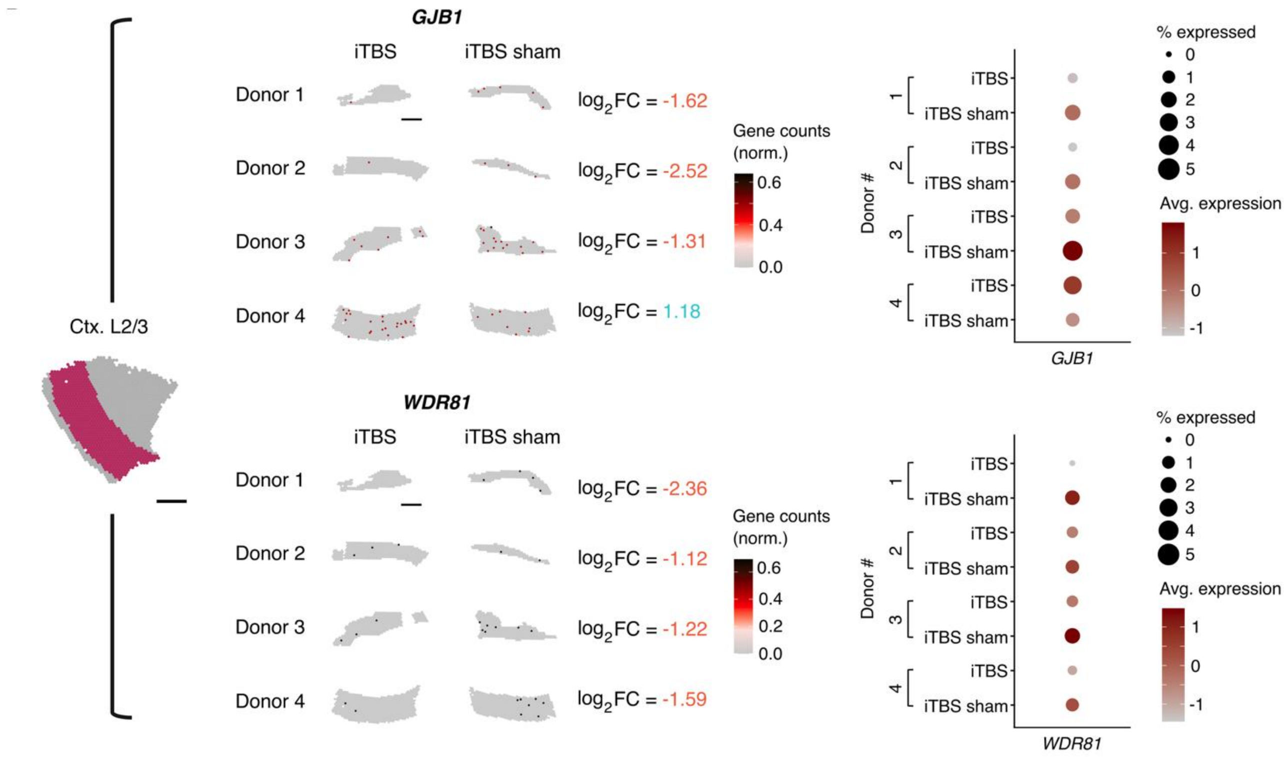


**Fig S3: Spatial gene expression plots of genes altered in cortical layer 2/3 following iTBS.** Significant genes were identified as having an absolute $\log_2$ fold-change ($\log_2$FC) $\geq 1$ in expression relative to sham within each individual and was also altered across all other individuals. Spatial gene expression plots (left) display the normalised counts for each gene within specific cortical layer spots for each individual and the corresponding $\log_2$FC following iTBS. Dot plots (right) summarise the expression pattern of each gene within the cortical layer across all individuals. Dot diameter indicates the proportion of spots expressing the gene whilst the colour gradient reflects the average expression levels of the gene. All scale bars represent 1 mm.

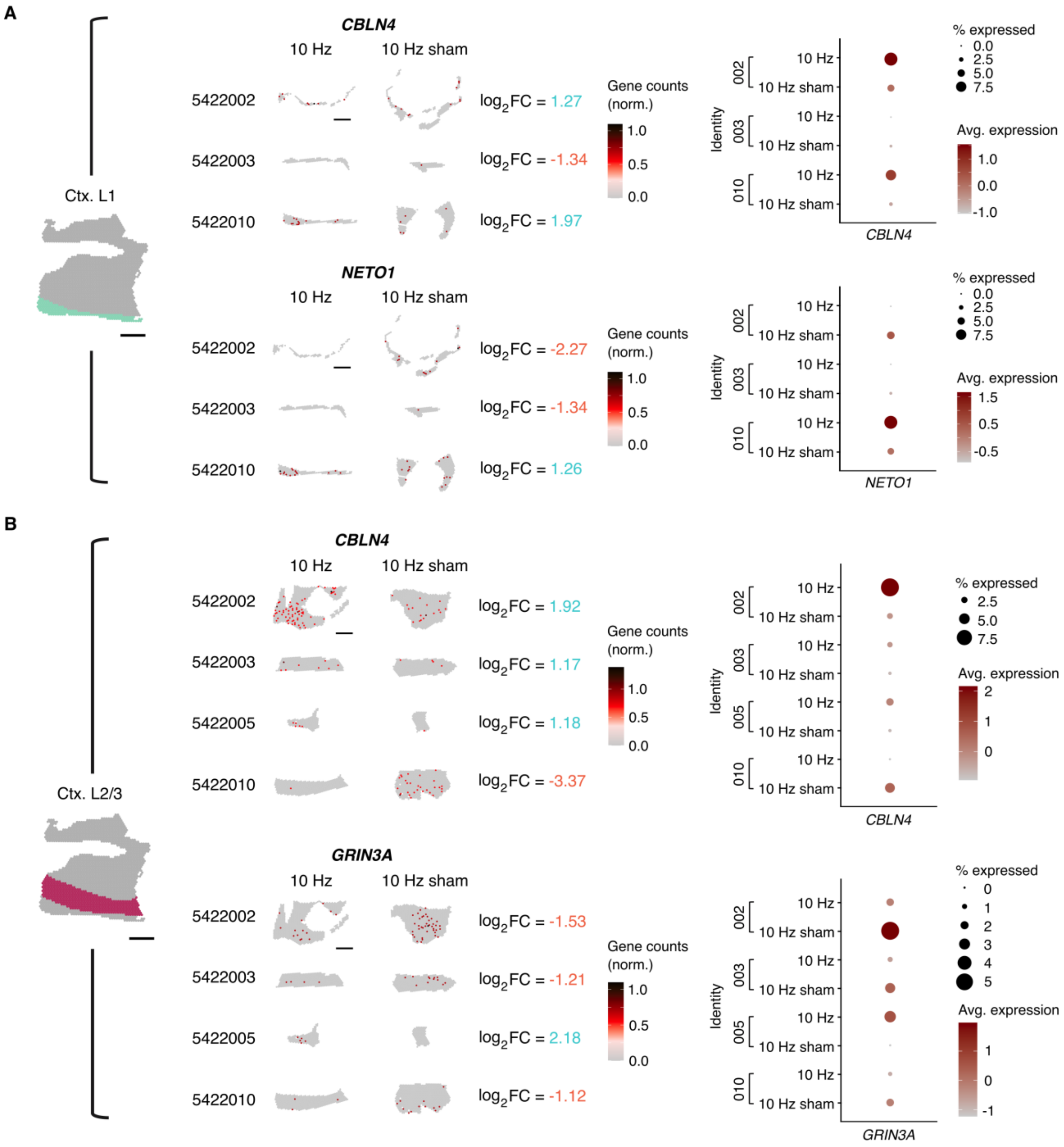


**Fig. S4: Spatial gene expression plots of genes altered in cortical layers 1, and 2/3 following 10 Hz.** Example plots of genes with a significant change (absolute $\log_2$ fold-change ($\log_2$FC) ≥ 1) in expression in **(A)** cortical layer 1 (Ctx. L1; n = 3) and **(B)** cortical layer 2/3 (Ctx. L2/3; n = 4), 3 h following 10 Hz stimulation. Significant genes were determined within each individual and cross-compared to identify the list of significant genes that were commonly altered across all individuals following stimulation. Spatial gene expression plots (left) display the normalised counts for each gene for spots specific to each cortical layer for each individual. The corresponding $\log_2$FC of the gene after 10 Hz stimulation is also displayed. Dot plots (right) summarise the expression pattern for each gene for each individual, with the dot diameter indicating the proportion of spots expressing the gene within the cortical layer and the colour gradient reflecting the average expression levels of the gene. All scale bars represent 1 mm.

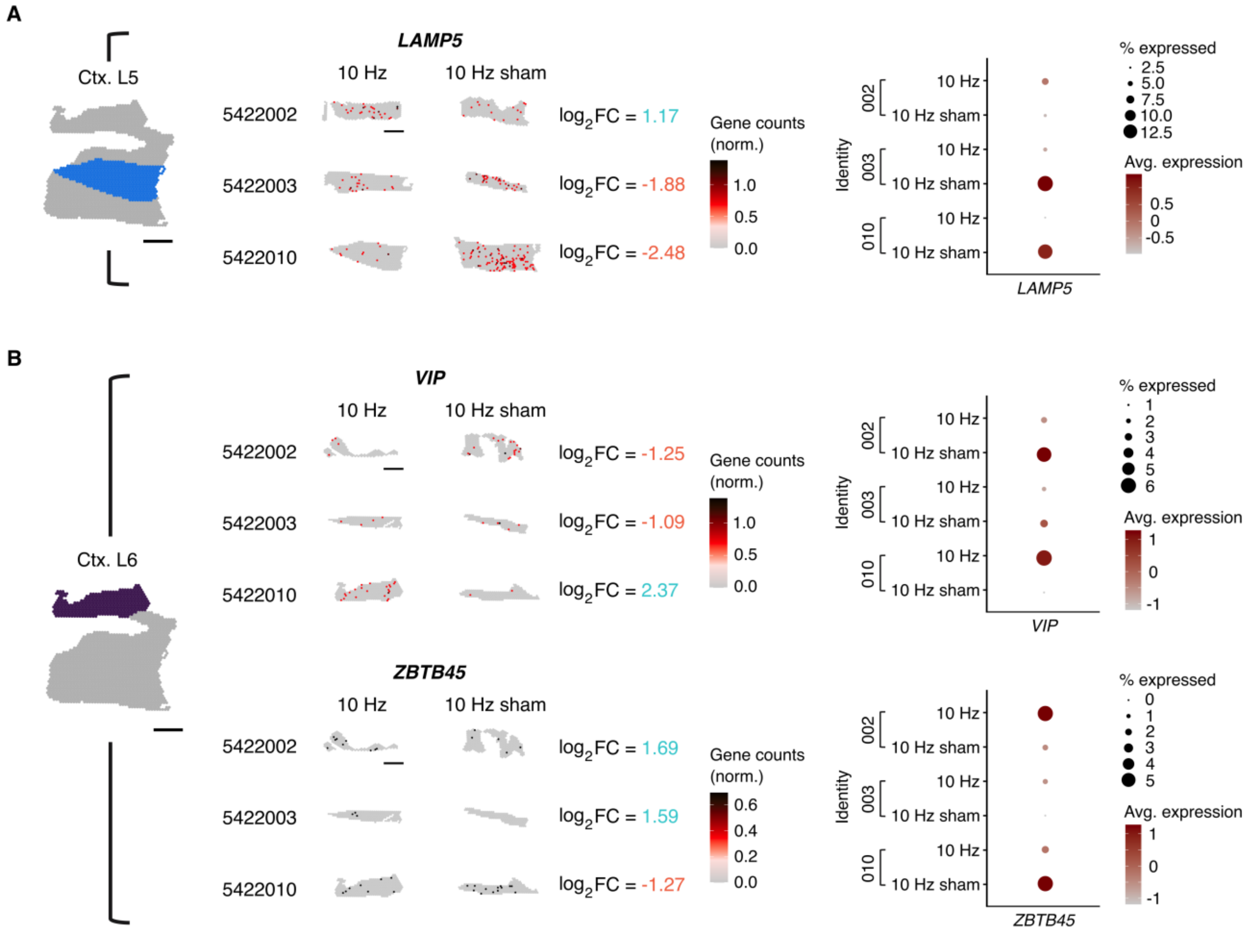


**Fig. S5: Spatial gene expression plots of example genes altered in cortical layers 5 and 6 following 10 Hz.** Example plots of genes with a significant change (absolute $\log_2$ fold-change ($\log_2$FC) ≥ 1) in expression within **(A)** cortical layer 5 (Ctx. L5; n = 3) and **(B)** cortical layer 6 (Ctx. L6; n = 3), 3 h following 10 Hz stimulation. Significant genes were determined within each individual and the overlap of genes commonly altered across all individuals were identified. Spatial gene expression plots (left) display the normalised counts for each gene for spots specific to each cortical layer. The corresponding $\log_2$FC of the gene for that individual following 10 Hz stimulation is shown. Dot plots (right) summarise the expression pattern for each gene for each individual, with the dot diameter indicating the proportion of cortical layer spots expressing the gene, and the colour gradient indicating the average expression levels of the gene. All scale bars represent 1 mm.

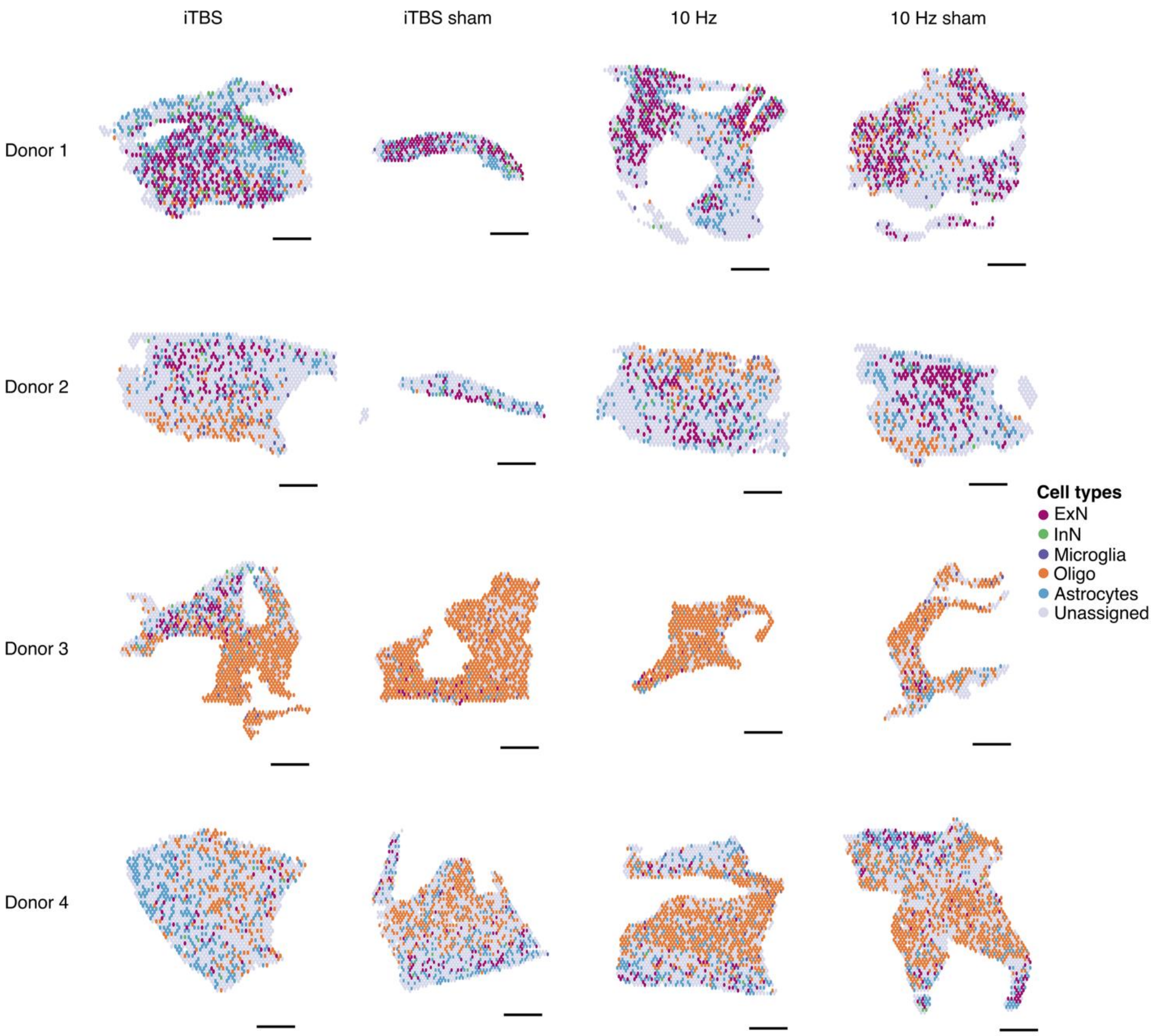


**Fig. S6: Cell-type enrichment annotation of Visium samples.** Robust cell type decomposition (RCTD) of Visium spots was performed using the transcriptomic profiles of various cell types of interest obtained from a published single-nuclei RNA-seq reference dataset. Individual spots were annotated as excitatory neurons (ExN), inhibitory neurons (InN), microglia, oligodendrocytes (Oligo), and/or astrocytes as determined by the cell-type enrichment percentage. Specifically, spots that contained at least 25% or more of a particular cell-type transcriptomic profile were categorised as that cell-type. Spots that didn't fall into any of the cell-type categories of interest were labelled as 'unassigned'. All scale bars represent 1 mm.

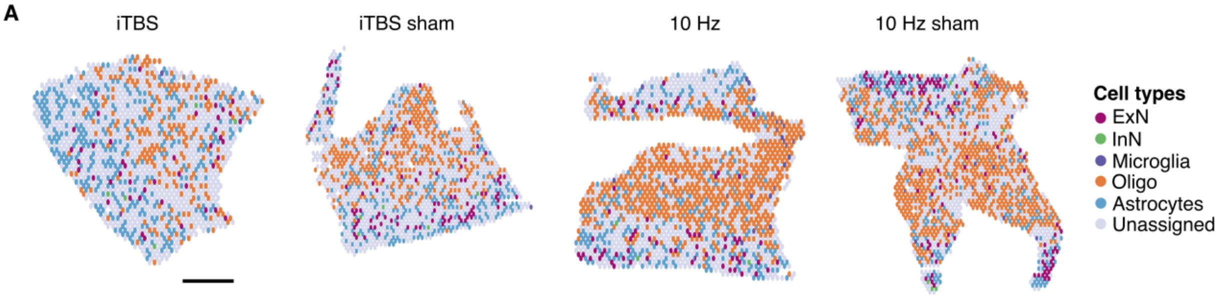


B

| | 5422002 | | | | 5422003 | | | | 5422005 | | | | 5422010 | | | |
|---|---|---|---|---|---|---|---|---|---|---|---|---|---|---|---|---|
| | iTBS | iTBS sham | 10 Hz | 10 Hz sham | iTBS | iTBS sham | 10 Hz | 10 Hz sham | iTBS | iTBS sham | 10 Hz | 10 Hz sham | iTBS | iTBS sham | 10 Hz | 10 Hz sham |
| ExN | 338 | 140 | 314 | 366 | 135 | 31 | 123 | 140 | 95 | 28 | 9 | 16 | 46 | 93 | 72 | 129 |
| InN | 64 | 21 | 37 | 33 | 10 | 5 | 14 | 5 | 12 | 1 | 0 | 2 | 5 | 5 | 3 | 6 |
| Microglia | 10 | 3 | 12 | 16 | 18 | 3 | 18 | 10 | 22 | 18 | 15 | 18 | 1 | 2 | 26 | 19 |
| Oligo | 38 | 5 | 33 | 126 | 149 | 1 | 125 | 52 | 674 | 849 | 418 | 288 | 222 | 390 | 695 | 750 |
| Astrocytes | 401 | 111 | 260 | 170 | 176 | 58 | 186 | 157 | 174 | 45 | 32 | 53 | 360 | 282 | 281 | 302 |

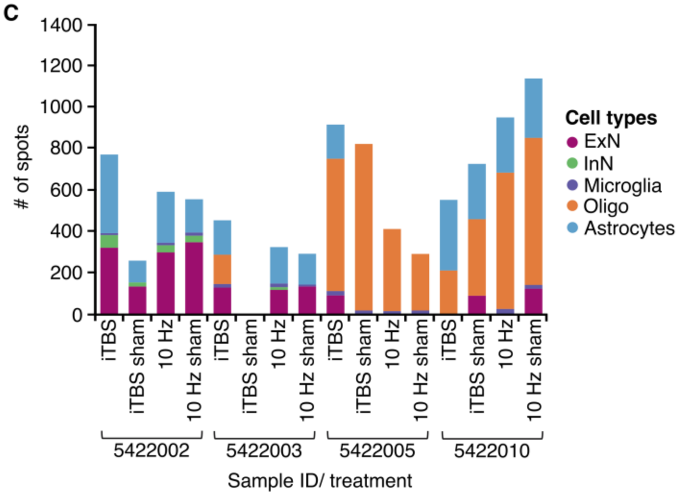


**Fig. S7: Single-cell guided deconvolution of neuronal and glial cell type enriched Visium spots. (A)** Representative spatial plots showcasing the distribution of spots enriched with a specific cell type (i.e., contains >25% of a specific cell-type's gene expression profile). Cell types that were identified include excitatory neurons (ExN), inhibitory neurons (InN), microglia, oligodendrocytes (Oligo), and astrocytes. Remaining spots that didn't contain at least 25% expression of any of the mentioned cell types were left unassigned. Scale bar represents 1 mm. **(B)** Summary table displaying the number of spots from each sample that were enriched with each cell type of interest. For each cell type, samples that contained a total number of cell-type enriched spots that was less than 20% of the highest number of spots were omitted from differential expression analysis. Samples that weren't used for further analysis are coloured in red. **(C)** Bar graph summarising the number of cell-type specific spots in each sample that was used for differential expression analysis.

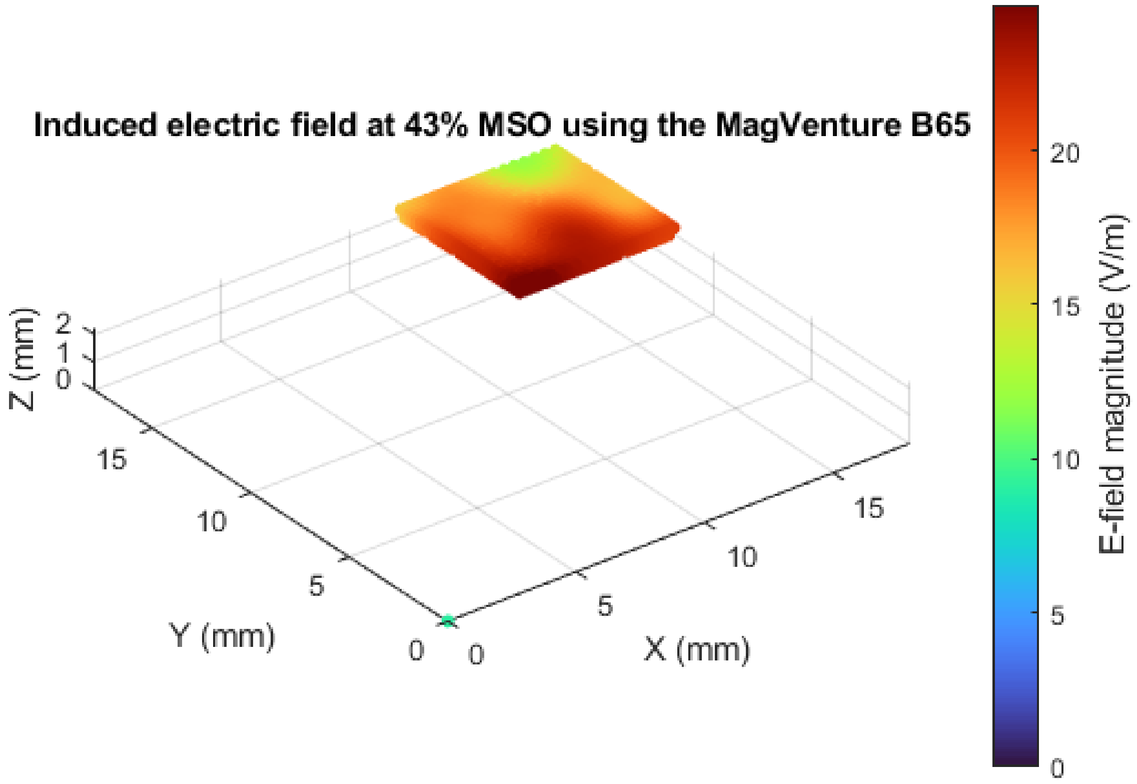


**Fig. S9: Modelling estimate of the induced electric field within the human brain slices.** Brain slice was modelled as a square piece of tissue 6mm x 6mmx 0.35mm sitting 2mm from the bottom of an ACSF bath (bath component removed from visualization to focus on slice). Brain slices were stimulated from below and induced an estimated peak of 25.56V/m within the cortical samples.

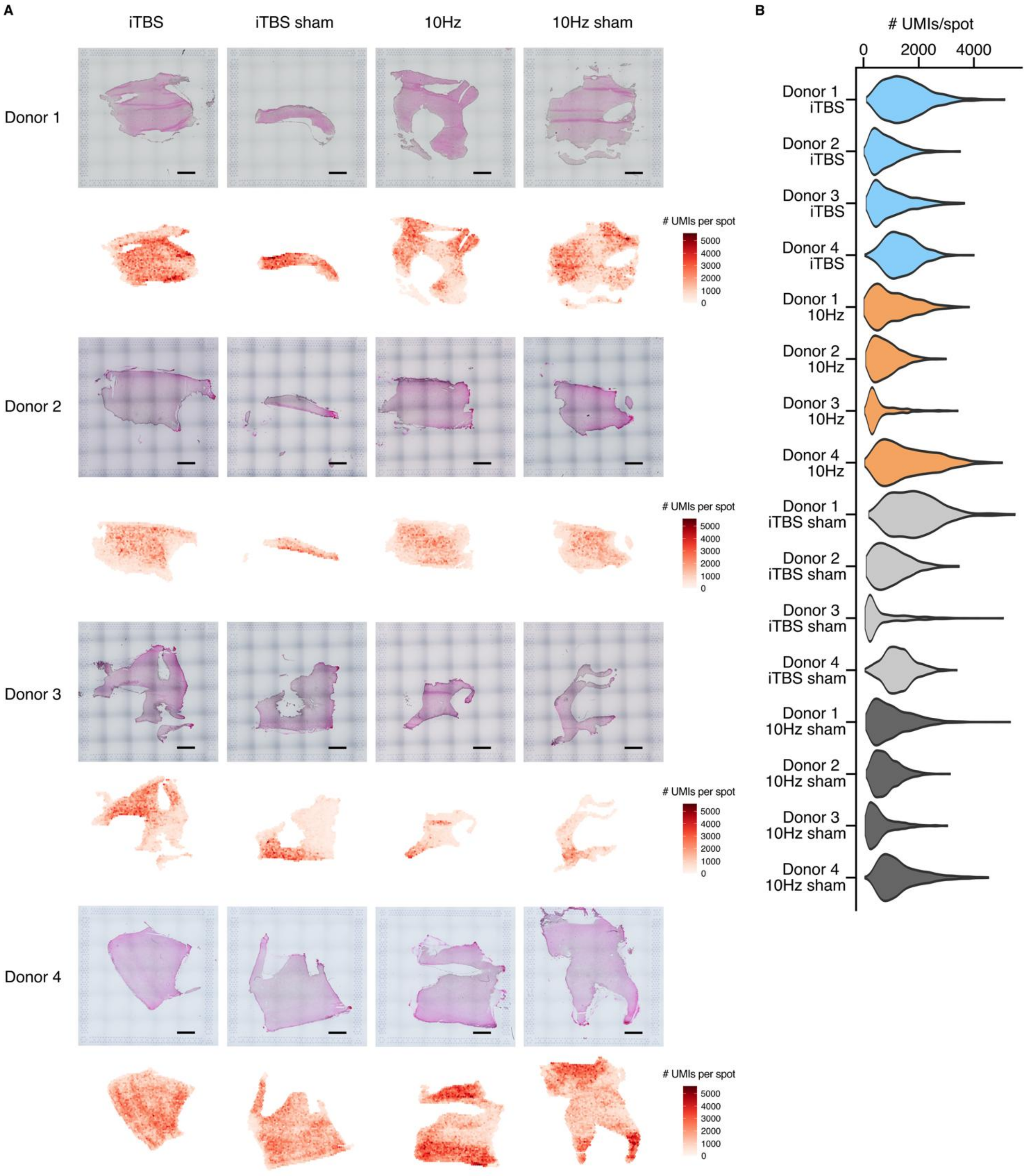
A
iTBS
iTBS sham
10Hz
10Hz sham
Donor 1
Donor 2
Donor 3
Donor 4
### UMIs per spot
5000
4000
3000
2000
1000
0
B
### UMIs/spot
0
2000
4000
Donor 1 iTBS
Donor 2 iTBS
Donor 3 iTBS
Donor 4 iTBS
Donor 1 10Hz
Donor 2 10Hz
Donor 3 10Hz
Donor 4 10Hz
Donor 1 iTBS sham
Donor 2 iTBS sham
Donor 3 iTBS sham
Donor 4 iTBS sham
Donor 1 10Hz sham
Donor 2 10Hz sham
Donor 3 10Hz sham
Donor 4 10Hz sham

**Fig. S10: Summary of mRNA expression across Visium human cortical sections. (A)** H&E histology images (top) and the accompanying unique molecular identifier (UMI) counts for all tissue-covered spots (bottom) from iTBS (n = 4), iTBS sham (n = 4), 10 Hz (n = 4), and 10 Hz sham (n = 4) stimulated cortical tissue sections, obtained from four different donors. Scale bars represent 1 mm. **(B)** Violin plot of the total number of UMIs (i.e., total number of mRNA reads per a single spot) for each Visium sample.

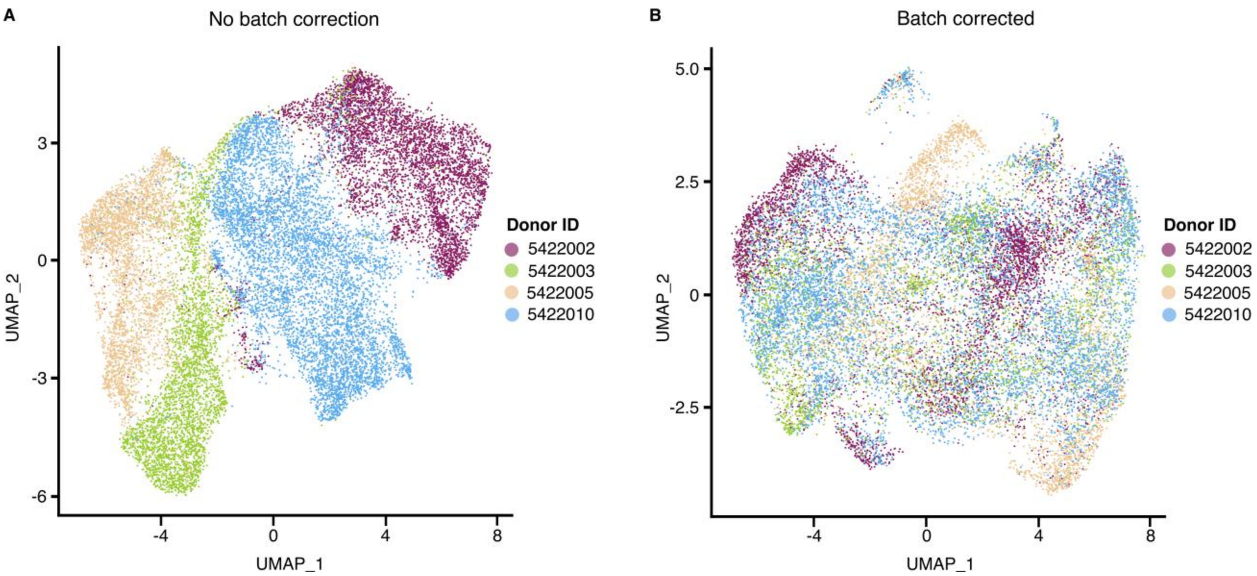


**Fig. S11: Batch correction of Visium spatial transcriptomics samples.** Uniform Manifold Approximation and Projection (UMAP) representation of spots from all human brain spatial transcriptomics samples (n =16) obtained from four donors. **(A)** Samples from the same donor (n = 4) were processed on the same Visium Gene Expression Slide and thus, prior to correction spatial gene expression patterns exhibited batch effects. **(B)** Thus, samples were batch corrected to reduce the technical variation between samples.

**Table S1: Summary of the number of spots in each cortical layer across all donor samples.** H&E histology-guided manual annotation of Visium spots was performed to identify cortical layer 1 (Ctx. L1), cortical layers 2/3 (Ctx. L2/3), cortical layer 5 (Ctx. L5), and cortical layer 6 (Ctx. L6).

| | Donor 1 | | | | Donor 2 | | | | Donor 3 | | | | Donor 4 | | | |
|---|---|---|---|---|---|---|---|---|---|---|---|---|---|---|---|---|
| | iTBS | iTBS sham | 10 Hz | 10 Hz sham | iTBS | iTBS sham | 10 Hz | 10 Hz sham | iTBS | iTBS sham | 10 Hz | 10 Hz sham | iTBS | iTBS sham | 10 Hz | 10 Hz sham |
| Ctx. L1 | 254 | 115 | 82 | 238 | 295 | 77 | 113 | 89 | 155 | 0 | 0 | 0 | 124 | 118 | 158 | 333 |
| Ctx. L2/3 | 264 | 257 | 765 | 742 | 474 | 165 | 394 | 444 | 442 | 379 | 158 | 119 | 723 | 642 | 430 | 645 |
| Ctx. L5 | 502 | 0 | 404 | 415 | 370 | 0 | 483 | 211 | 259 | 0 | 0 | 201 | 461 | 358 | 493 | 712 |
| Ctx. L6 | 299 | 0 | 163 | 315 | 304 | 0 | 218 | 164 | 93 | 0 | 0 | 0 | 201 | 318 | 364 | 235 |

**Table S2: Number of excitatory neuron (ExN)-enriched spots within each cortical layer for each Visium sample.** Spots were categorised as ExN if they expressed > 25% of the cell-type specific expression profile as calculated by the robust cell type decomposition algorithm using a single-nuclei RNA-sequencing human cell database as a reference.

| | | Ctx. L1 | Ctx. L2/3 | Ctx. L5 | Ctx. L6 |
|---|---|---|---|---|---|
| Donor 1 | iTBS | 17 | 78 | 157 | 86 |
| | iTBS sham | 51 | 89 | 0 | 0 |
| | 10 Hz | 10 | 204 | 53 | 47 |
| | 10 Hz sham | 38 | 167 | 80 | 81 |
| Donor 2 | iTBS | 29 | 66 | 39 | 1 |
| | iTBS sham | 0 | 31 | 0 | 0 |
| | 10 Hz | 0 | 38 | 50 | 55 |
| | 10 Hz sham | 0 | 50 | 52 | 38 |
| Donor 3 | iTBS | 18 | 66 | 11 | 0 |
| | iTBS sham | 0 | 27 | 0 | 0 |
| | 10 Hz | 0 | 9 | 0 | 0 |
| | 10 Hz sham | 0 | 7 | 9 | 0 |
| Donor 4 | iTBS | 1 | 26 | 8 | 11 |
| | iTBS sham | 0 | 63 | 11 | 4 |
| | 10 Hz | 22 | 30 | 4 | 13 |
| | 10 Hz sham | 40 | 15 | 27 | 47 |

**Table S3: Number of inhibitory neuron (InN)-enriched spots within each cortical layer for each Visium sample.** Spots were categorised as InN if they expressed > 25% of the cell-type specific expression profiles as calculated by the robust cell type decomposition algorithm, using a single-nuclei RNA-sequencing human cell database as a reference.

| | | Ctx. L1 | Ctx. L2/3 | Ctx. L5 | Ctx. L6 |
|---|---|---|---|---|---|
| Donor 1 | iTBS | 12 | 24 | 17 | 11 |
| | iTBS sham | 8 | 13 | 0 | 0 |
| | 10 Hz | 2 | 24 | 5 | 6 |
| | 10 Hz sham | 2 | 17 | 7 | 7 |
| Donor 2 | iTBS | 5 | 4 | 1 | 0 |
| | iTBS sham | 0 | 5 | 0 | 0 |
| | 10 Hz | 0 | 3 | 8 | 3 |
| | 10 Hz sham | 0 | 1 | 2 | 2 |
| Donor 3 | iTBS | 6 | 6 | 0 | 0 |
| | iTBS sham | 0 | 1 | 0 | 0 |
| | 10 Hz | 0 | 0 | 0 | 0 |
| | 10 Hz sham | 0 | 1 | 1 | 0 |
| Donor 4 | iTBS | 0 | 3 | 0 | 2 |
| | iTBS sham | 0 | 5 | 0 | 0 |
| | 10 Hz | 1 | 2 | 0 | 0 |
| | 10 Hz sham | 3 | 0 | 2 | 1 |

**Table S4: Number of microglia-enriched spots within each cortical layer for each Visium sample.** Spots were categorised as microglia if they expressed > 25% of the cell-type specific expression profiles as calculated by the robust cell type decomposition algorithm, using a single-nuclei RNA-sequencing human cell database as a reference.

| | | Ctx. L1 | Ctx. L2/3 | Ctx. L5 | Ctx. L6 |
|---|---|---|---|---|---|
| Donor 1 | iTBS | 1 | 0 | 6 | 3 |
| | iTBS sham | 1 | 2 | 0 | 0 |
| | 10 Hz | 2 | 6 | 3 | 1 |
| | 10 Hz sham | 5 | 3 | 5 | 3 |
| Donor 2 | iTBS | 2 | 2 | 1 | 13 |
| | iTBS sham | 2 | 1 | 0 | 0 |
| | 10 Hz | 8 | 8 | 2 | 0 |
| | 10 Hz sham | 4 | 4 | 0 | 2 |
| Donor 3 | iTBS | 0 | 1 | 3 | 1 |
| | iTBS sham | 0 | 3 | 0 | 0 |
| | 10 Hz | 0 | 0 | 0 | 0 |
| | 10 Hz sham | 0 | 1 | 0 | 0 |
| Donor 4 | iTBS | 0 | 0 | 1 | 0 |
| | iTBS sham | 1 | 0 | 0 | 0 |
| | 10 Hz | 0 | 1 | 9 | 4 |
| | 10 Hz sham | 2 | 9 | 6 | 2 |

**Table S5: Number of oligodendrocyte (Oligo)-enriched spots within each cortical layer for each Visium sample.** Spots were categorised as oligodendrocytes if they expressed > 25% of the cell-type specific expression profiles as calculated by the robust cell type decomposition algorithm, using a single-nuclei RNA-sequencing human cell database as a reference.

| | | Ctx. L1 | Ctx. L2/3 | Ctx. L5 | Ctx. L6 |
|---|---|---|---|---|---|
| Donor 1 | iTBS | 1 | 2 | 13 | 22 |
| | iTBS sham | 1 | 4 | 0 | 0 |
| | 10 Hz | 0 | 11 | 13 | 9 |
| | 10 Hz sham | 11 | 73 | 34 | 8 |
| Donor 2 | iTBS | 1 | 13 | 33 | 102 |
| | iTBS sham | 0 | 1 | 0 | 0 |
| | 10 Hz | 38 | 77 | 10 | 0 |
| | 10 Hz sham | 29 | 23 | 0 | 0 |
| Donor 3 | iTBS | 17 | 125 | 167 | 85 |
| | iTBS sham | 0 | 315 | 0 | 0 |
| | 10 Hz | 0 | 127 | 0 | 0 |
| | 10 Hz sham | 0 | 61 | 66 | 0 |
| Donor 4 | iTBS | 7 | 65 | 113 | 37 |
| | iTBS sham | 1 | 70 | 126 | 112 |
| | 10 Hz | 4 | 98 | 316 | 80 |
| | 10 Hz sham | 109 | 363 | 234 | 44 |

**Table S6: Number of astrocyte-enriched spots within each cortical layer for each Visium sample.** Spots were categorised as astrocytes if they expressed > 25% of the cell-type specific expression profiles as calculated by the robust cell type decomposition algorithm, using a single-nuclei RNA-sequencing human cell database as a reference.

| | | Ctx. L1 | Ctx. L2/3 | Ctx. L5 | Ctx. L6 |
|---|---|---|---|---|---|
| Donor 1 | iTBS | 82 | 78 | 182 | 59 |
| | iTBS sham | 28 | 83 | 0 | 0 |
| | 10 Hz | 4 | 146 | 89 | 21 |
| | 10 Hz sham | 4 | 76 | 51 | 39 |
| Donor 2 | iTBS44 | 44 | 63 | 51 | 18 |
| | iTBS sham | 20 | 38 | 0 | 0 |
| | 10 Hz | 6 | 66 | 82 | 32 |
| | 10 Hz sham | 5 | 84 | 40 | 28 |
| Donor 3 | iTBS | 36 | 102 | 28 | 5 |
| | iTBS sham | 0 | 41 | 0 | 0 |
| | 10 Hz | 0 | 23 | 0 | 0 |
| | 10 Hz sham | 0 | 12 | 41 | 0 |
| Donor 4 | iTBS | 55 | 197 | 78 | 30 |
| | iTBS sham | 40 | 127 | 43 | 30 |
| | 10 Hz | 52 | 86 | 60 | 65 |
| | 10 Hz sham | 77 | 51 | 109 | 65 |
