## Supplementary data for "Repetitive transcranial magnetic stimulation induces protocol and layer specific transcriptomic plasticity in the human cortex"

**Data S1. Differentiall**

| Gene symbol |
|---|
| A1BG-AS1 |
| BTBD10 |
| PTPN4 |
| TRIM27 |
| ARG2 |
| CD200 |
| CD53 |
| CENPN |
| CKLF |
| DUSP2 |
| IFIT5 |
| ILRUN |
| JCAD |
| KBTBD7 |
| LCP2 |
| MB21D2 |
| NFATC2 |
| NFATC3 |
| NFKBIZ |
| SRGAP1 |
| SUSD4 |
| TXNIP |
| USP19 |
| FRMPD2 |
| KCNH1 |
| KCNIP1 |
| KCNJ9 |
| KCNS2 |
| AKAP1 |
| CDK5RAP1 |
| CKMT1A |
| MAG |
| NTRK3 |
| PRRG1 |
| FBLN7 |
| ALKBH3 |
| AHR |
| SOCS6 |
| SST |
| GABRA4 |
| GABRG2 |
| ADCYAP1 |
| SLITRK5 |

STX1A
ACVR1C
AMH
CACNG3
CDH10
CPEB1
DCC
DMXL2
DNMT3A
DOC2A
NRGN
PIP4P1
POGZ
TIAM2
AARSD1
AASS
ABCD1
ACAD11
ADGRL4
AKR1C3
ALKBH6
ANKRD52
AOPEP
APTR
ARMC6
ASCC2
ASXL3
BBIP1
BBS7
BCL11A
BLOC1S3
BMS1
BORCS5
BRINP3
C1orf174
CARNMT1
CAVIN2
CCDC90B
CDK11A
CDK11B
CEP250
CEP63
CEP76
CHMP1A
CIR1
CLBA1

CLIP2
CLN5
COMMD1
COMMD3
COMMD5
COMTD1
COQ5
CRB1
CRYZ
CTBP1-DT
CYP4X1
DCTN1-AS1
DCUN1D5
DGUOK
DYRK4
ECD
EGR4
EIF5AL1
EMP3
EMSY-DT
ENTREP3
EPM2A-DT
ERCC5
EXOSC5
FAM171A2
FAM222B
FAM228B
FAM241B
FBXO6
FOXJ2
FUT1
GAA
GNB3
GPR161
GTF2IRD2B
H2AC25
HAR1A
HEY1
HNMT
HS6ST2
HYCC1
ID3
IDE
INO80
ITGA6
JKAMP

KBTBD4
KIFBP
KLF12
KMT2C
L2HGDH
LAMC1
LATS2
LCNL1
LIG3
LINC00342
LINC00662
LINC01007
LINC01511
LINC02609
LNX1
LPGAT1
LYRM1
MAGT1
MAN1B1
MARK1
MARS1
MCM6
MCPH1
MCU
MDM4
METTL2A
MEX3D
MGAT5
MIR7-3HG
MON2
MPPE1
MPV17L2
MRS2
MSANTD4
MTARC2
MTHFS
MYL9
NAPEPLD
NEU3
NEURL1B
NEUROD6
NOL8
NPAT
NT5DC1
NUBP1
NUDT14

OPN3
OSCP1
PAGR1
PCBP1-AS1
PCYOX1
PEPD
PIP4K2C
POLD4
PPARG
PPP2R3C
PRKAB2
PRPF3
PYROXD1
RALGAPA2
RASA2
RASL11A
RBMX2
RCE1
RNF144B
RNF25
RRP12
SALL3
SAMHD1
SDHAP4
SETMAR
SH3YL1
SHISAL1
SLC15A2
SLC25A19
SLC30A10
SLF2
SMARCD2
SMURF1
SPATA2L
SPATA33
SPEF1
SRPK1
SRRM4
ST6GALNAC5
SUPT6H
SYT13
TAF5L
TAMM41
TBL1X
TDRD9
TIMM21

TIMM22
TMEM117
TMEM121B
TMEM192
TMEM201
TMEM229B
TMEM243
TMEM63C
TMEM69
TRHDE
TRIM46
TRUB1
TTC9
TYW1
UBOX5
UBXN2B
UHMK1
UNK
USP2
UST
VPS13A
VXN
WDFY3
WDR55
WDR91
WRAP73
XYLT1
ZBED3
ZBTB20
ZBTB22
ZBTB8OS
ZDHHC7
ZFPM2
ZFY
ZFYVE9
ZNF48
ZNF568
ZNF584
ZNF689

y expressed genes (DEGs) in cortical layer 1 (L1) following iTBS.

| Gene name |
| --- |
| A1BG antisense RNA 1 |
| BTB domain containing 10 |
| protein tyrosine phosphatase non-receptor type 4 |
| tripartite motif containing 27 |
| arginase 2 |
| CD200 molecule |
| CD53 molecule |
| centromere protein N |
| chemokine like factor |
| dual specificity phosphatase 2 |
| interferon induced protein with tetratricopeptide repeats 5 |
| inflammation and lipid regulator with UBA-like and NBR1-like domains |
| junctional cadherin 5 associated |
| kelch repeat and BTB domain containing 7 |
| lymphocyte cytosolic protein 2 |
| Mab-21 domain containing 2 |
| nuclear factor of activated T cells 2 |
| nuclear factor of activated T cells 3 |
| NFKB inhibitor zeta |
| SLIT-ROBO Rho GTPase activating protein 1 |
| sushi domain containing 4 |
| thioredoxin interacting protein |
| ubiquitin specific peptidase 19 |
| FERM and PDZ domain containing 2 |
| potassium voltage-gated channel subfamily H member 1 |
| potassium voltage-gated channel interacting protein 1 |
| potassium inwardly rectifying channel subfamily J member 9 |
| potassium voltage-gated channel modifier subfamily S member 2 |
| A-kinase anchoring protein 1 |
| CDK5RAP1 mitochondrial tRNA methylthiotransferase |
| creatine kinase, mitochondrial 1A |
| myelin associated glycoprotein |
| neurotrophic receptor tyrosine kinase 3 |
| proline rich and Gla domain 1 |
| fibulin 7 |
| alkB homolog 3, alpha-ketoglutarate dependent dioxygenase |
| aryl hydrocarbon receptor |
| suppressor of cytokine signaling 6 |
| somatostatin |
| gamma-aminobutyric acid type A receptor subunit alpha4 |
| gamma-aminobutyric acid type A receptor subunit gamma2 |
| adenylate cyclase activating polypeptide 1 |
| SLIT and NTRK like family member 5 |

syntaxin 1A
activin A receptor type 1C
anti-Mullerian hormone
calcium voltage-gated channel auxiliary subunit gamma 3
cadherin 10
cytoplasmic polyadenylation element binding protein 1
DCC netrin 1 receptor
Dmx like 2
DNA methyltransferase 3 alpha
double C2 domain alpha
neurogranin
phosphatidylinositol-4,5-bisphosphate 4-phosphatase 1
pogo transposable element derived with ZNF domain
TIAM Rac1 associated GEF 2
alanyl-tRNA synthetase domain containing 1
aminoadipate-semialdehyde synthase
ATP binding cassette subfamily D member 1
acyl-CoA dehydrogenase family member 11
adhesion G protein-coupled receptor L4
aldo-keto reductase family 1 member C3
alkB homolog 6
ankyrin repeat domain 52
aminopeptidase O (putative)
Alu-mediated CDKN1A/p21 transcriptional regulator
armadillo repeat containing 6
activating signal cointegrator 1 complex subunit 2
ASXL transcriptional regulator 3
BBSome interacting protein 1
Bardet-Biedl syndrome 7
BCL11 transcription factor A
biogenesis of lysosomal organelles complex 1 subunit 3
BMS1 ribosome biogenesis factor
BLOC-1 related complex subunit 5
BMP/retinoic acid inducible neural specific 3
chromosome 1 open reading frame 174
carnosine N-methyltransferase 1
caveolae associated protein 2
coiled-coil domain containing 90B
cyclin dependent kinase 11A
cyclin dependent kinase 11B
centrosomal protein 250
centrosomal protein 63
centrosomal protein 76
charged multivesicular body protein 1A
corepressor interacting with RBPJ, CIR1
clathrin binding box of aftiphilin containing 1

CAP-Gly domain containing linker protein 2
CLN5 intracellular trafficking protein
copper metabolism domain containing 1
COMM domain containing 3
COMM domain containing 5
catechol-O-methyltransferase domain containing 1
coenzyme Q5, methyltransferase
crumbs cell polarity complex component 1
crystallin zeta
CTBP1 divergent transcript
cytochrome P450 family 4 subfamily X member 1
DCTN1 antisense RNA 1
defective in cullin neddylation 1 domain containing 5
deoxyguanosine kinase
dual specificity tyrosine phosphorylation regulated kinase 4
ecdysoneless cell cycle regulator
early growth response 4
eukaryotic translation initiation factor 5A like 1
epithelial membrane protein 3 (MAM blood group)
EMSY divergent transcript
endosomal transmembrane epsin interactor 3
EPM2A divergent transcript
ERCC excision repair 5, endonuclease
exosome component 5
family with sequence similarity 171 member A2
family with sequence similarity 222 member B
family with sequence similarity 228 member B
family with sequence similarity 241 member B
F-box protein 6
forkhead box J2
fucosyltransferase 1 (H blood group)
alpha glucosidase
G protein subunit beta 3
G protein-coupled receptor 161
GTF2I repeat domain containing 2B
H2A clustered histone 25
highly accelerated region 1A
hes related family bHLH transcription factor with YRPW motif 1
histamine N-methyltransferase
heparan sulfate 6-O-sulfotransferase 2
hyccin PI4KA lipid kinase complex subunit 1
inhibitor of DNA binding 3
insulin degrading enzyme
INO80 complex ATPase subunit
integrin subunit alpha 6
JNK1/MAPK8 associated membrane protein

kelch repeat and BTB domain containing 4
kinesin family binding protein
KLF transcription factor 12
lysine methyltransferase 2C
L-2-hydroxyglutarate dehydrogenase
laminin subunit gamma 1
large tumor suppressor kinase 2
lipocalin like 1
DNA ligase 3
long intergenic non-protein coding RNA 342
long intergenic non-protein coding RNA 662
long intergenic non-protein coding RNA 1007
long intergenic non-protein coding RNA 1511
long intergenic non-protein coding RNA 2609
ligand of numb-protein X 1
lysophosphatidylglycerol acyltransferase 1
LYR motif containing 1
magnesium transporter 1
mannosidase alpha class 1B member 1
microtubule affinity regulating kinase 1
methionyl-tRNA synthetase 1
minichromosome maintenance complex component 6
microcephalin 1
mitochondrial calcium uniporter
MDM4 regulator of p53
methyltransferase 2A, tRNA N3-cytidine
mex-3 RNA binding family member D
alpha-1,6-mannosylglycoprotein 6-beta-N-acetylglucosaminyltransferase
MIR7-3 host gene
MON2 homolog, regulator of endosome-to-Golgi trafficking
metallophosphoesterase 1
MPV17 mitochondrial inner membrane protein like 2
magnesium transporter MRS2
Myb/SANT DNA binding domain containing 4 with coiled-coils
mitochondrial amidoxime reducing component 2
methenyltetrahydrofolate synthetase
myosin light chain 9
N-acyl phosphatidylethanolamine phospholipase D
neuraminidase 3
neuralized E3 ubiquitin protein ligase 1B
neuronal differentiation 6
nucleolar protein 8
nuclear protein, coactivator of histone transcription
5'-nucleotidase domain containing 1
NUBP iron-sulfur cluster assembly factor 1, cytosolic
nudix hydrolase 14

opsin 3
organic solute carrier partner 1
PAXIP1 associated glutamate rich protein 1
PCBP1 antisense RNA 1
prenylcysteine oxidase 1
peptidase D
phosphatidylinositol-5-phosphate 4-kinase type 2 gamma
DNA polymerase delta 4, accessory subunit
peroxisome proliferator activated receptor gamma
protein phosphatase 2 regulatory subunit B"gamma
protein kinase AMP-activated non-catalytic subunit beta 2
pre-mRNA processing factor 3
pyridine nucleotide-disulphide oxidoreductase domain 1
Ral GTPase activating protein catalytic subunit alpha 2
RAS p21 protein activator 2
RAS like family 11 member A
RNA binding motif protein X-linked 2
Ras converting CAAX endopeptidase 1
ring finger protein 144B
ring finger protein 25
ribosomal RNA processing 12 homolog
spalt like transcription factor 3
SAM and HD domain containing deoxynucleoside triphosphate triphosphohydrolase 1
SDHA pseudogene 4
SET domain and mariner transposase fusion gene
SH3 and SYLF domain containing 1
shisa like 1
solute carrier family 15 member 2
solute carrier family 25 member 19
solute carrier family 30 member 10
SMC5-SMC6 complex localization factor 2
SWI/SNF related BAF chromatin remodeling complex subunit D2
SMAD specific E3 ubiquitin protein ligase 1
spermatogenesis associated 2 like
spermatogenesis associated 33
sperm flagellar 1
SRSF protein kinase 1
serine/arginine repetitive matrix 4
ST6 N-acetylgalactosaminide alpha-2,6-sialyltransferase 5
SPT6 homolog, histone chaperone and transcription elongation factor
synaptotagmin 13
TATA-box binding protein associated factor 5 like
TAM41 mitochondrial translocator assembly and maintenance homolog
transducin beta like 1 X-linked
tudor domain containing 9
translocase of inner mitochondrial membrane 21

translocase of inner mitochondrial membrane 22
transmembrane protein 117
transmembrane protein 121B
transmembrane protein 192
transmembrane protein 201
transmembrane protein 229B
transmembrane protein 243
transmembrane protein 63C
transmembrane protein 69
thyrotropin releasing hormone degrading enzyme
tripartite motif containing 46
TruB pseudouridine synthase family member 1
tetratricopeptide repeat domain 9
tRNA-yW synthesizing protein 1 homolog
U-box domain containing 5
UBX domain protein 2B
U2AF homology motif kinase 1
unk zinc finger
ubiquitin specific peptidase 2
uronyl 2-sulfotransferase
vacuolar protein sorting 13 homolog A
vexin
WD repeat and FYVE domain containing 3
WD repeat domain 55
WD repeat domain 91
WD repeat containing, antisense to TP73
xylosyltransferase 1
zinc finger BED-type containing 3
zinc finger and BTB domain containing 20
zinc finger and BTB domain containing 22
zinc finger and BTB domain containing 8 opposite strand
zinc finger DHHC-type palmitoyltransferase 7
zinc finger protein, FOG family member 2
zinc finger protein Y-linked
zinc finger FYVE-type containing 9
zinc finger protein 48
zinc finger protein 568
zinc finger protein 584
zinc finger protein 689

| EntrezID | Gene category | Donor 1 | |
|---|---|---|---|
| | | log2fc | p.adj |
| 503538 | | -1.558232135 | 1 |
| 84280 | cell death | -1.530217759 | 1 |
| 5775 | cell death | -1.323766881 | 1 |
| 5987 | cell death | -1.365587057 | 1 |
| 384 | immune response | -1.442754918 | 1 |
| 4345 | immune response | -1.579293751 | 1 |
| 963 | immune response | -1.673709353 | 1 |
| 55839 | immune response | -2.143194636 | 1 |
| 51192 | immune response | -1.295197729 | 1 |
| 1844 | immune response | -3.017663754 | 1 |
| 24138 | immune response | -1.602626254 | 1 |
| 64771 | immune response | -1.069194054 | 1 |
| 57608 | immune response | -2.88016023 | 1 |
| 84078 | immune response | -1.465122731 | 1 |
| 3937 | immune response | 1.359305705 | 1 |
| 151963 | immune response | -1.558232135 | 1 |
| 4773 | immune response/ oligodendrocytes | -1.143194636 | 1 |
| 4775 | immune response | -1.628621463 | 1 |
| 64332 | immune response | -1.728157137 | 1 |
| 57522 | immune response | 2.316236983 | 1 |
| 55061 | immune response | -1.465122731 | 1 |
| 10628 | immune response | 1.178733459 | 1 |
| 10869 | immune response | -1.657767809 | 1 |
| 143162 | intrinsinc plasticity | -2.365587057 | 1 |
| 3756 | intrinsinc plasticity | -2.521706259 | 0.749147845 |
| 30820 | intrinsinc plasticity | -1.005691112 | 1 |
| 3765 | intrinsinc plasticity | -1.221197148 | 1 |
| 3788 | intrinsinc plasticity | -2.465122731 | 1 |
| 8165 | mitochondria | -1.465122731 | 1 |
| 51654 | mitochondria | -1.950549558 | 1 |
| 548596 | mitochondria | -1.258671853 | 1 |
| 4099 | myelin | 1.026730366 | 1 |
| 4916 | myelin/ oligodendrocytes | -1.323766881 | 1 |
| 5638 | myelin/ oligodendrocytes | -2.143194636 | 1 |
| 129804 | myelin/ oligodendrocytes | -2.017663754 | 1 |
| 221120 | neurogenesis | -1.365587057 | 1 |
| 196 | neurogenesis/ immune response | 1.441767865 | 1 |
| 9306 | immune response | 1.249122787 | 1 |
| 6750 | neurotransmitter | -1.810619297 | 1 |
| 2557 | neurotransmitter/ synaptic plasticity | -1.143194636 | 1 |
| 2566 | neurotransmitter/ synaptic plasticity | -1.28069816 | 0.359158854 |
| 116 | neurotrophin | -1.88016023 | 1 |
| 26050 | synapse | -1.442754918 | 1 |

| | | | |
|---|---|---|---|
| 6804 | synapse | -1.440176374 | 0.021870903 |
| 130399 | synaptic plasticity | -1.821266541 | 1 |
| 268 | synaptic plasticity | -1.143194636 | 1 |
| 10368 | synaptic plasticity | -1.178818546 | 1 |
| 1008 | synaptic plasticity | -1.143194636 | 1 |
| 64506 | synaptic plasticity | -2.365587057 | 1 |
| 1630 | synaptic plasticity | -1.728157137 | 1 |
| 23312 | synaptic plasticity | -1.143194636 | 1 |
| 1788 | synaptic plasticity | -1.143194636 | 1 |
| 8448 | synaptic plasticity | -1.095888921 | 1 |
| 4900 | synaptic plasticity | -1.324801442 | 8.09E-17 |
| 90809 | | -1.230657477 | 1 |
| 23126 | synaptic plasticity | -1.221197148 | 1 |
| 26230 | synaptic plasticity | -1.628621463 | 1 |
| 80755 | | -1.728157137 | 1 |
| 10157 | | -1.465122731 | 1 |
| 215 | | -1.728157137 | 1 |
| 84129 | | -3.143194636 | 1 |
| 64123 | | -1.465122731 | 1 |
| 8644 | | 1.664160286 | 1 |
| 84964 | | -1.036279432 | 1 |
| 283373 | | 1.441767865 | 1 |
| 84909 | | -1.465122731 | 1 |
| 100505854 | | -1.143194636 | 1 |
| 93436 | | -1.143194636 | 1 |
| 84164 | | -1.143194636 | 1 |
| 80816 | | -1.143194636 | 1 |
| 92482 | | -1.017663754 | 1 |
| 55212 | | -1.365587057 | 1 |
| 53335 | | -1.295197729 | 1 |
| 388552 | | -1.143194636 | 1 |
| 9790 | | -1.143194636 | 1 |
| 118426 | | -1.143194636 | 1 |
| 339479 | | -1.143194636 | 1 |
| 339448 | | -1.143194636 | 1 |
| 138199 | | -2.143194636 | 1 |
| 8436 | | 1.856805364 | 1 |
| 60492 | | -1.143194636 | 1 |
| 728642 | | -1.558232135 | 1 |
| 984 | | -2.143194636 | 1 |
| 11190 | | -2.143194636 | 1 |
| 80254 | | -1.143194636 | 1 |
| 79959 | | -1.728157137 | 1 |
| 5119 | | -1.295197729 | 1 |
| 9541 | | -1.143194636 | 1 |
| 122616 | | -1.406229042 | 1 |

| | | |
|---|---|---|
| 7461 | -1.628621463 | 1 |
| 1203 | -1.950549558 | 1 |
| 150684 | -3.017663754 | 1 |
| 23412 | -1.335839714 | 1 |
| 28991 | -1.465122731 | 1 |
| 118881 | -1.143194636 | 1 |
| 84274 | 1.441767865 | 1 |
| 23418 | -2.728157137 | 1 |
| 1429 | -2.143194636 | 1 |
| 92070 | -1.143194636 | 1 |
| 260293 | -2.843634354 | 0.693932838 |
| 100189589 | -1.728157137 | 1 |
| 84259 | -1.230657477 | 1 |
| 1716 | -1.036279432 | 1 |
| 8798 | -1.628621463 | 1 |
| 11319 | -1.143194636 | 1 |
| 1961 | -2.094285035 | 0.000170976 |
| 143244 | -2.465122731 | 1 |
| 2014 | -1.505764715 | 1 |
| 124902718 | -1.143194636 | 1 |
| 10712 | -1.143194636 | 1 |
| 100507557 | -2.143194636 | 1 |
| 2073 | -2.628621463 | 0.826271773 |
| 56915 | -1.143194636 | 1 |
| 284069 | -1.060732476 | 1 |
| 55731 | -1.143194636 | 1 |
| 375190 | 1.664160286 | 1 |
| 219738 | -1.143194636 | 1 |
| 26270 | 1.664160286 | 1 |
| 55810 | -1.406229042 | 1 |
| 2523 | -1.143194636 | 1 |
| 2548 | -1.143194636 | 1 |
| 2784 | -1.143194636 | 1 |
| 23432 | -1.406229042 | 1 |
| 389524 | -1.728157137 | 1 |
| 92815 | 1.026730366 | 1 |
| 768096 | -1.628621463 | 1 |
| 23462 | -1.491117939 | 1 |
| 3176 | -1.143194636 | 1 |
| 90161 | -1.728157137 | 1 |
| 84668 | 1.664160286 | 1 |
| 3399 | 1.079197786 | 1 |
| 3416 | -1.143194636 | 1 |
| 54617 | -1.602626254 | 1 |
| 3655 | 1.079197786 | 1 |
| 51528 | -1.313119637 | 1 |

| | | |
|---|---|---|
| 55709 | -1.143194636 | 1 |
| 26128 | -1.143194636 | 1 |
| 11278 | -2.558232135 | 1 |
| 58508 | -1.406229042 | 1 |
| 79944 | -1.558232135 | 1 |
| 3915 | 2.178733459 | 1 |
| 26524 | 1.664160286 | 1 |
| 401562 | -3.728157137 | 1 |
| 3980 | -1.465122731 | 1 |
| 150759 | -1.143194636 | 1 |
| 148189 | -1.795271332 | 1 |
| 100506527 | -2.991191542 | 0.009110555 |
| 100506791 | -2.728157137 | 1 |
| 105378853 | -1.465122731 | 1 |
| 84708 | -2.465122731 | 0.032659198 |
| 9926 | -1.465122731 | 1 |
| 57149 | -1.88016023 | 1 |
| 84061 | 1.178733459 | 1 |
| 11253 | -1.027717418 | 1 |
| 4139 | -1.28069816 | 1 |
| 4141 | -1.558232135 | 1 |
| 4175 | -1.465122731 | 1 |
| 79648 | -1.406229042 | 1 |
| 90550 | -1.143194636 | 1 |
| 4194 | -1.558232135 | 1 |
| 339175 | -1.991191542 | 1 |
| 399664 | -1.558232135 | 1 |
| 4249 | -1.764683013 | 1 |
| 284424 | -1.28069816 | 1 |
| 23041 | -1.143194636 | 1 |
| 65258 | -1.335839714 | 1 |
| 84769 | -1.88016023 | 1 |
| 57380 | 1.441767865 | 1 |
| 84437 | -1.991191542 | 1 |
| 54996 | -1.728157137 | 1 |
| 10588 | -1.143194636 | 1 |
| 10398 | 1.856805364 | 1 |
| 222236 | -1.657767809 | 1 |
| 10825 | -1.728157137 | 1 |
| 54492 | -1.143194636 | 1 |
| 63974 | -2.558232135 | 1 |
| 55035 | -1.143194636 | 1 |
| 4863 | -1.143194636 | 1 |
| 221294 | -1.143194636 | 1 |
| 4682 | 1.178733459 | 1 |
| 256281 | -1.213583964 | 1 |

| | | |
|---|---|---|
| 23596 | -1.143194636 | 1 |
| 127700 | -1.313119637 | 1 |
| 79447 | -1.005691112 | 1 |
| 400960 | -1.465122731 | 1 |
| 51449 | -1.225656796 | 1 |
| 5184 | -1.060732476 | 1 |
| 79837 | -1.005691112 | 1 |
| 57804 | -1.991191542 | 1 |
| 5468 | -1.728157137 | 1 |
| 55012 | -1.465122731 | 1 |
| 5565 | -1.406229042 | 1 |
| 9129 | 1.026730366 | 1 |
| 79912 | -1.143194636 | 1 |
| 57186 | -1.558232135 | 1 |
| 5922 | -2.365587057 | 1 |
| 387496 | -1.558232135 | 1 |
| 51634 | -1.143194636 | 1 |
| 9986 | -1.143194636 | 1 |
| 255488 | -1.143194636 | 1 |
| 64320 | -1.821266541 | 1 |
| 23223 | -1.384202735 | 1 |
| 27164 | -1.143194636 | 1 |
| 25939 | -1.465122731 | 1 |
| 220729 | -2.143194636 | 1 |
| 6419 | -2.728157137 | 1 |
| 26751 | 1.316236983 | 1 |
| 85352 | -1.295197729 | 1 |
| 6565 | 1.557245082 | 1 |
| 60386 | -1.406229042 | 1 |
| 55532 | -1.143194636 | 1 |
| 55719 | -1.950549558 | 1 |
| 6603 | -1.143194636 | 1 |
| 57154 | -1.728157137 | 1 |
| 124044 | -1.143194636 | 1 |
| 124045 | -1.143194636 | 1 |
| 25876 | -1.521706259 | 1 |
| 6732 | -1.335839714 | 1 |
| 84530 | -1.973269634 | 1 |
| 81849 | -2.221197148 | 0.317991463 |
| 6830 | -1.25010984 | 1 |
| 57586 | -1.517590151 | 0.722058892 |
| 27097 | -1.821266541 | 1 |
| 132001 | -1.143194636 | 1 |
| 6907 | 1.441767865 | 1 |
| 122402 | -1.88016023 | 1 |
| 29090 | -1.28069816 | 1 |

| | | |
|---|---|---|
| 29928 | -1.143194636 | 1 |
| 84216 | -1.143194636 | 1 |
| 27439 | -1.406229042 | 1 |
| 201931 | 1.664160286 | 1 |
| 199953 | -1.88016023 | 1 |
| 161145 | -1.795271332 | 1 |
| 79161 | -3.143194636 | 1 |
| 57156 | -2.143194636 | 1 |
| 51249 | -1.143194636 | 1 |
| 29953 | -1.821266541 | 1 |
| 80128 | -1.88016023 | 1 |
| 142940 | -1.28069816 | 1 |
| 23508 | -1.295197729 | 1 |
| 55253 | -2.465122731 | 1 |
| 22888 | -1.143194636 | 1 |
| 137886 | -1.728157137 | 1 |
| 127933 | -1.143194636 | 1 |
| 85451 | -1.465122731 | 1 |
| 9099 | -1.491117939 | 1 |
| 10090 | -1.558232135 | 1 |
| 23230 | -1.143194636 | 1 |
| 254778 | -1.20459518 | 1 |
| 23001 | -1.491117939 | 1 |
| 54853 | -1.143194636 | 1 |
| 29062 | -1.143194636 | 1 |
| 49856 | -1.728157137 | 1 |
| 64131 | -1.143194636 | 1 |
| 84327 | -1.143194636 | 1 |
| 26137 | 2.178733459 | 1 |
| 9278 | -1.728157137 | 1 |
| 339487 | -2.143194636 | 1 |
| 55625 | -1.673709353 | 1 |
| 23414 | -1.143194636 | 1 |
| 7544 | 1.441767865 | 1 |
| 9372 | -1.602626254 | 1 |
| 197407 | 1.178733459 | 1 |
| 374900 | -1.406229042 | 1 |
| 201514 | -1.465122731 | 1 |
| 115509 | -1.465122731 | 1 |

| Donor 2 | | Donor 4 | | |
|---|---|---|---|---|
| log2fc | p.adj | log2fc | p.adj | |
| -1.937784604 | 1 | 1.51340924 | 1 | |
| 1.232140398 | 1 | 1.343484238 | 1 | |
| -1.200819009 | 1 | 1.09837174 | 1 | |
| 1.647177897 | 1 | -1.241478262 | 1 | |
| 1.232140398 | 1 | -1.071553261 | 1 | |
| 1.121109085 | 1 | 1.51340924 | 1 | |
| -1.160177025 | 1 | -1.187030478 | 1 | y |
| -1.352822103 | 1 | 2.250374834 | 1 | |
| 1.232140398 | 1 | 1.09837174 | 1 | |
| 1.335233891 | 1 | -1.171088935 | 1 | |
| -2.259712698 | 1 | -1.808518855 | 1 | y |
| 1.31014291 | 1 | -1.071553261 | 1 | |
| -2.522747104 | 1 | -2.071553261 | 1 | y |
| -2.937784604 | 1 | -1.656515762 | 1 | y |
| -1.044699807 | 1 | 1.250374834 | 1 | |
| -1.130429681 | 1 | 1.928446739 | 1 | |
| -1.937784604 | 1 | -1.071553261 | 1 | y |
| -1.937784604 | 1 | -1.656515762 | 1 | y |
| -1.130429681 | 1 | 1.735801661 | 1 | |
| -1.352822103 | 1 | 2.250374834 | 1 | |
| 1.521647015 | 1 | 1.09837174 | 1 | |
| -1.089787697 | 1 | 1.560714954 | 1 | |
| 1.062215396 | 1 | 1.51340924 | 1 | |
| -2.937784604 | 1 | 1.928446739 | 1 | |
| 1.521647015 | 1 | -1.656515762 | 1 | |
| -1.352822103 | 1 | 1.387878358 | 1 | |
| 1.647177897 | 1 | -1.241478262 | 1 | |
| -2.937784604 | 1 | 1.928446739 | 1 | |
| -1.259712698 | 1 | 2.51340924 | 1 | |
| 1.232140398 | 1 | 1.250374834 | 1 | |
| 2.062215396 | 1 | 1.043923956 | 1 | |
| 1.869570319 | 1 | 1.169454838 | 1 | y |
| 1.347617615 | 1 | -1.48659076 | 1 | |
| -2.259712698 | 1 | 1.250374834 | 1 | |
| -1.937784604 | 1 | -1.393481356 | 1 | y |
| -1.352822103 | 1 | 1.51340924 | 1 | |
| -1.937784604 | 1 | 2.250374834 | 1 | |
| -1.130429681 | 1 | 1.15083916 | 1 | |
| 1.140217908 | 1 | 1.1300806 | 1 | |
| -1.044699807 | 1 | 1.09837174 | 1 | |
| 1.042037515 | 1 | -1.393481356 | 1 | |
| 1.521647015 | 1 | -2.878908183 | 1 | |
| -1.285707907 | 1 | 1.51340924 | 1 | |

| | | | | |
|---|---|---|---|---|
| 1.499620709 | 1 | 1.572302929 | 1 | |
| -1.089787697 | 1 | -1.656515762 | 1 | y |
| -1.200819009 | 1 | -1.656515762 | 1 | y |
| 1.016411707 | 1 | 1.51340924 | 1 | |
| 1.384143491 | 1 | -1.656515762 | 1 | |
| 1.232140398 | 1 | 1.343484238 | 1 | |
| -2.522747104 | 1 | 1.51340924 | 1 | |
| -1.011785185 | 1 | 1.802915857 | 1 | |
| 1.647177897 | 1 | -1.393481356 | 1 | |
| 1.725180409 | 1 | 1.217953356 | 1 | |
| 1.030135425 | 3.96E-06 | -1.026465371 | 5.46E-06 | |
| 1.232140398 | 1 | -1.35166118 | 1 | |
| -1.107709605 | 1 | 1.15083916 | 1 | |
| 1.817102899 | 1 | -1.48659076 | 1 | |
| 1.647177897 | 1 | 1.735801661 | 1 | |
| -1.200819009 | 1 | -1.393481356 | 1 | |
| -1.937784604 | 1 | 1.928446739 | 1 | |
| -1.352822103 | 1 | 1.928446739 | 1 | |
| -1.352822103 | 1 | -2.071553261 | 1 | |
| -1.937784604 | 1 | 1.735801661 | 1 | |
| 1.149678238 | 1 | 1.928446739 | 1 | |
| -1.615856509 | 1 | -1.071553261 | 1 | |
| 2.232140398 | 1 | 1.51340924 | 1 | |
| 1.521647015 | 1 | -1.071553261 | 1 | |
| -1.937784604 | 1 | -2.656515762 | 1 | |
| 1.384143491 | 1 | 1.735801661 | 1 | |
| -1.352822103 | 1 | 1.928446739 | 1 | |
| 1.232140398 | 1 | 1.387878358 | 1 | |
| 1.869570319 | 1 | 2.387878358 | 1 | |
| 1.149678238 | 1 | 1.250374834 | 1 | |
| -1.937784604 | 1 | -1.656515762 | 1 | |
| 1.232140398 | 1 | -1.071553261 | 1 | |
| -1.522747104 | 1 | -1.293945682 | 1 | |
| 1.232140398 | 1 | 1.928446739 | 1 | |
| -1.522747104 | 1 | 1.51340924 | 1 | |
| -1.522747104 | 1 | -1.071553261 | 1 | |
| -2.937784604 | 1 | 1.250374834 | 1 | |
| -1.937784604 | 1 | -1.071553261 | 1 | |
| -1.200819009 | 1 | -1.656515762 | 1 | |
| -1.352822103 | 1 | -1.656515762 | 1 | |
| 1.762655115 | 1 | -1.071553261 | 1 | |
| -1.200819009 | 1 | -1.071553261 | 1 | |
| -1.937784604 | 1 | 1.51340924 | 1 | |
| -1.011785185 | 1 | 1.191481145 | 1 | |
| -1.130429681 | 1 | 1.250374834 | 1 | |
| -2.937784604 | 1 | 1.51340924 | 1 | |

| | | | |
|---|---|---|---|
| 1.016411707 | 1 | -1.071553261 | 1 |
| 1.062215396 | 1 | 1.51340924 | 1 |
| -1.130429681 | 1 | 1.043923956 | 1 |
| 1.062215396 | 1 | 1.735801661 | 1 |
| 1.062215396 | 1 | 1.15083916 | 1 |
| -2.937784604 | 1 | 1.51340924 | 1 |
| -1.352822103 | 1 | -1.393481356 | 1 |
| -1.615856509 | 1 | 1.928446739 | 1 |
| -1.352822103 | 1 | 1.51340924 | 1 |
| -1.937784604 | 1 | 1.51340924 | 1 |
| -1.522747104 | 1 | -1.656515762 | 1 |
| -1.352822103 | 1 | -1.656515762 | 1 |
| -1.352822103 | 1 | 1.928446739 | 1 |
| 1.31014291 | 1 | 1.867046194 | 1 |
| -1.259712698 | 1 | 1.250374834 | 1 |
| 1.062215396 | 1 | -3.53098488 | 1 |
| 1.117899529 | 1 | -1.282120247 | 1 |
| -1.452357776 | 1 | 1.51340924 | 1 |
| -1.274819591 | 1 | 1.250374834 | 1 |
| -1.937784604 | 1 | 1.928446739 | 1 |
| 1.232140398 | 1 | -1.293945682 | 1 |
| -1.522747104 | 1 | -1.071553261 | 1 |
| 1.762655115 | 1 | -1.071553261 | 1 |
| 1.062215396 | 1 | -1.334587667 | 1 |
| 2.384143491 | 1 | 1.15083916 | 1 |
| -1.937784604 | 1 | -1.071553261 | 1 |
| -1.352822103 | 1 | -1.071553261 | 1 |
| 1.454532819 | 1 | 1.928446739 | 1 |
| -1.352822103 | 1 | -2.071553261 | 1 |
| 1.647177897 | 1 | -1.241478262 | 1 |
| -1.937784604 | 1 | -1.656515762 | 1 |
| -1.200819009 | 1 | 1.250374834 | 1 |
| -1.937784604 | 1 | 1.928446739 | 1 |
| -2.352822103 | 1 | 1.250374834 | 1 |
| -1.352822103 | 1 | 1.928446739 | 1 |
| 1.177692614 | 1 | -1.071553261 | 1 |
| -1.200819009 | 1 | 1.51340924 | 1 |
| 1.384143491 | 1 | 1.665412333 | 1 |
| -1.937784604 | 1 | 1.191481145 | 1 |
| -1.937784604 | 1 | 1.51340924 | 1 |
| -1.937784604 | 1 | -1.808518855 | 1 |
| -2.300354683 | 1 | 1.11287131 | 1 |
| -2.522747104 | 1 | 1.51340924 | 1 |
| 1.647177897 | 1 | -1.656515762 | 1 |
| -1.200819009 | 1 | 1.250374834 | 1 |
| -1.044699807 | 1 | 1.250374834 | 1 |

| | | | |
|---|---|---|---|
| 1.762655115 | 1 | 1.343484238 | 1 |
| -1.160177025 | 1 | 1.09837174 | 1 |
| -1.937784604 | 1 | -1.071553261 | 1 |
| 1.062215396 | 1 | -1.293945682 | 1 |
| -2.352822103 | 1 | 2.250374834 | 1 |
| -1.674750198 | 1 | 1.735801661 | 1 |
| -1.937784604 | 1 | -1.293945682 | 1 |
| -1.522747104 | 1 | -1.293945682 | 1 |
| -1.200819009 | 1 | 1.51340924 | 1 |
| 1.521647015 | 1 | -1.393481356 | 1 |
| 1.062215396 | 1 | -1.071553261 | 1 |
| -1.352822103 | 1 | 1.928446739 | 1 |
| 1.384143491 | 1 | -1.878908183 | 1 |
| -1.200819009 | 1 | 1.51340924 | 1 |
| 1.000814852 | 1 | -1.071553261 | 1 |
| 1.062215396 | 1 | 1.250374834 | 1 |
| -2.130429681 | 1 | 1.387878358 | 1 |
| -1.200819009 | 1 | 2.928446739 | 1 |
| 2.31014291 | 1 | -1.071553261 | 1 |
| -1.352822103 | 1 | 1.387878358 | 1 |
| 1.762655115 | 1 | 1.065950263 | 1 |
| 1.062215396 | 1 | 1.15083916 | 1 |
| -1.937784604 | 1 | 1.250374834 | 1 |
| -1.352822103 | 1 | -1.393481356 | 1 |
| 1.454532819 | 1 | -1.393481356 | 1 |
| -1.937784604 | 1 | 2.250374834 | 1 |
| -3.937784604 | 1 | -1.656515762 | 1 |
| 2.454532819 | 1 | -1.656515762 | 1 |
| 1.647177897 | 1 | -1.209056785 | 1 |
| -1.407269887 | 1 | 1.065950263 | 1 |
| 1.647177897 | 1 | 1.665412333 | 1 |
| -1.937784604 | 1 | -1.393481356 | 1 |
| -1.522747104 | 1 | -2.071553261 | 1 |
| 1.521647015 | 1 | 1.250374834 | 1 |
| -1.352822103 | 1 | 1.51340924 | 1 |
| -1.352822103 | 1 | 3.250374834 | 1 |
| -2.352822103 | 1 | 1.51340924 | 1 |
| -1.937784604 | 1 | 1.51340924 | 1 |
| -1.937784604 | 1 | -1.071553261 | 1 |
| -1.522747104 | 1 | -1.071553261 | 1 |
| -1.352822103 | 1 | 1.51340924 | 1 |
| -1.937784604 | 1 | 1.51340924 | 1 |
| -1.200819009 | 1 | 1.250374834 | 1 |
| -1.200819009 | 1 | 2.51340924 | 1 |
| -1.352822103 | 1 | -2.393481356 | 1 |
| 1.585777353 | 1 | 1.006449251 | 1 |

| | | | |
|---|---|---|---|
| 1.817102899 | 1 | 1.191481145 | 1 |
| 1.232140398 | 1 | -1.071553261 | 1 |
| 1.969105992 | 1 | 1.15083916 | 1 |
| -1.352822103 | 1 | -1.071553261 | 1 |
| 1.121109085 | 1 | -1.171088935 | 1 |
| 1.521647015 | 1 | 1.006449251 | 1 |
| 1.564715737 | 1 | 1.15083916 | 1 |
| 1.521647015 | 1 | 1.854446158 | 1 |
| -1.937784604 | 1 | 1.928446739 | 1 |
| -1.615856509 | 1 | 1.928446739 | 1 |
| 1.062215396 | 1 | 1.51340924 | 1 |
| -1.089787697 | 1 | 1.735801661 | 1 |
| -1.352822103 | 1 | 1.928446739 | 1 |
| 1.232140398 | 1 | -1.656515762 | 1 |
| 1.062215396 | 1 | 1.735801661 | 1 |
| -1.352822103 | 1 | 2.51340924 | 1 |
| 1.647177897 | 1 | 1.15083916 | 1 |
| 1.062215396 | 1 | -1.071553261 | 1 |
| -1.200819009 | 1 | 1.51340924 | 1 |
| -2.522747104 | 1 | 2.928446739 | 1 |
| 1.299254594 | 1 | -1.602067978 | 1 |
| -1.352822103 | 1 | -1.071553261 | 1 |
| 1.521647015 | 1 | 1.15083916 | 1 |
| 1.384143491 | 1 | 1.250374834 | 1 |
| -1.352822103 | 1 | 1.51340924 | 1 |
| 1.969105992 | 1 | 1.09837174 | 1 |
| 1.062215396 | 1 | -1.241478262 | 1 |
| -1.044699807 | 1 | 1.043923956 | 1 |
| -1.352822103 | 1 | -1.878908183 | 1 |
| -1.352822103 | 1 | 1.928446739 | 1 |
| -1.615856509 | 1 | -1.293945682 | 1 |
| -1.522747104 | 1 | 2.250374834 | 1 |
| -1.130429681 | 1 | 3.09837174 | 1 |
| 2.384143491 | 1 | 1.09837174 | 1 |
| -1.715392182 | 1 | -1.293945682 | 1 |
| -1.522747104 | 1 | 1.09837174 | 1 |
| -1.089787697 | 1 | 1.250374834 | 1 |
| 1.097839306 | 1 | -1.071553261 | 1 |
| 1.762655115 | 1 | 3.387878358 | 1 |
| 1.384143491 | 1 | 1.15083916 | 1 |
| 1.585777353 | 1 | -1.071553261 | 1 |
| -1.674750198 | 1 | 1.250374834 | 1 |
| -1.352822103 | 1 | -2.393481356 | 1 |
| -1.937784604 | 1 | 1.928446739 | 1 |
| 1.647177897 | 1 | -1.656515762 | 1 |
| 1.232140398 | 1 | 1.735801661 | 1 |

| | | | |
|---|---|---|---|
| 1.521647015 | 1 | 1.928446739 | 1 |
| -2.937784604 | 1 | 2.51340924 | 1 |
| -1.352822103 | 1 | 1.51340924 | 1 |
| -1.352822103 | 1 | -1.48659076 | 1 |
| -1.937784604 | 1 | 2.735801661 | 1 |
| -2.937784604 | 1 | 1.51340924 | 1 |
| 1.062215396 | 1 | -2.393481356 | 1 |
| 1.762655115 | 1 | 1.51340924 | 1 |
| -1.615856509 | 1 | -1.071553261 | 1 |
| -1.200819009 | 1 | 2.51340924 | 1 |
| 1.762655115 | 1 | -2.071553261 | 1 |
| 1.869570319 | 1 | 1.250374834 | 1 |
| 1.521647015 | 1 | 1.387878358 | 1 |
| 1.232140398 | 1 | 1.51340924 | 1 |
| -1.937784604 | 1 | 1.51340924 | 1 |
| -1.352822103 | 1 | 1.250374834 | 1 |
| 2.31014291 | 1 | 1.191481145 | 1 |
| 1.062215396 | 1 | -1.48659076 | 1 |
| 2.521647015 | 1 | -1.071553261 | 1 |
| 1.062215396 | 1 | 1.343484238 | 1 |
| 2.062215396 | 1 | -2.656515762 | 1 |
| 2.258612609 | 1 | 1.201465233 | 1 |
| 1.869570319 | 1 | 1.693981485 | 1 |
| -1.200819009 | 1 | -2.071553261 | 1 |
| 1.521647015 | 1 | -1.071553261 | 1 |
| -1.937784604 | 1 | 2.250374834 | 1 |
| -1.423211431 | 1 | 2.51340924 | 1 |
| -2.937784604 | 1 | 1.250374834 | 1 |
| -1.937784604 | 1 | 1.735801661 | 1 |
| -1.937784604 | 1 | 2.09837174 | 1 |
| 1.454532819 | 1 | -1.071553261 | 1 |
| -1.200819009 | 1 | 2.09837174 | 1 |
| 1.232140398 | 1 | 1.51340924 | 1 |
| -1.352822103 | 1 | 1.928446739 | 1 |
| -1.200819009 | 1 | 1.250374834 | 1 |
| -2.522747104 | 1 | -2.071553261 | 1 |
| 1.969105992 | 1 | -1.393481356 | 1 |
| -2.522747104 | 1 | 1.51340924 | 1 |
| -1.522747104 | 1 | 2.928446739 | 1 |

**Data S2. Differentiall**

| Gene symbol |
| --- |
| DNAH17 |
| DOCK5 |
| GJB1 |
| LINC00595 |
| MX2 |
| TVP23A |
| WDR81 |
| ZNF587 |

**y expressed genes (DEGs) in cortical layer 2/3 (L2/3) following iTBS.**

| Gene name | EntrezID |
|---|---|
| dynein axonemal heavy chain 17 | 8632 |
| dedicator of cytokinesis 5 | 80005 |
| gap junction protein beta 1 | 2705 |
| long intergenic non-protein coding RNA 595 | 414243 |
| MX dynamin like GTPase 2 | 4600 |
| trans-golgi network vesicle protein 23 homolog A | 780776 |
| WD repeat domain 81 | 124997 |
| zinc finger protein 587 | 84914 |

my

| Gene category | Donor 1 | | Donor 2 | |
|---|---|---|---|---|
| | log2fc | p.adj | log2fc | p.adj |
| myelin | 1.283158525 | 1 | -2.84434913 | 1 |
| | 1.183622851 | 1 | 1.284933887 | 1 |
| | -1.623732071 | 1 | -2.522421035 | 1 |
| | 1.376267929 | 1 | -1.107383535 | 1 |
| | 1.183622851 | 1 | -1.744813456 | 1 |
| neurogenesis | -1.261161992 | 1 | -1.84434913 | 1 |
| | -2.360697665 | 1 | -1.107383535 | 1 |
| | 1.96123043 | 1 | -1.522421035 | 1 |

immune response
mitochondria

synaptic plasticity

synaptic plasticity

synaptic plasticity
synaptic plasticity
synaptic plasticity
immune response

yelin/ oligodendrocytes

intrinsinc plasticity

| Donor 3 | | Donor 4 | |
|---|---|---|---|
| log2fc | p.adj | log2fc | p.adj |
| -1.029203443 | 1 | 1.288009269 | 1 |
| -2.079829516 | 1 | -1.064507146 | 1 |
| -1.309311362 | 1 | 1.182214605 | 1 |
| -1.543776616 | 1 | 1.314004477 | 1 |
| 1.058259398 | 1 | 1.050970072 | 1 |
| 1.100079574 | 1 | 1.413540151 | 1 |
| -1.221848521 | 1 | -1.586459849 | 1 |
| 1.36311398 | 1 | 2.050970072 | 1 |

**Data S3. Differentially**

| Gene symbol |
|---|
| NFIA |
| CHAC1 |
| FADD |
| AKNA |
| BMP7 |
| CASP4 |
| FCGR1A |
| IL32 |
| MOK |
| NFKB2 |
| PTPN2 |
| CACNA1C |
| KCNJ16 |
| NKAIN4 |
| WNT2B |
| GAB1 |
| UGT8 |
| FZD1 |
| KAT8 |
| KCTD10 |
| DLX1 |
| EHD4 |
| ERBB4 |
| FCHO2 |
| GDNF |
| KIFC3 |
| LTA4H |
| NETO1 |
| PCDH11X |
| PDE7A |
| PI4K2A |
| PVALB |
| VTI1A |
| ACVR1C |
| ADAM22 |
| ARHGEF6 |
| DAB1 |
| CBLN4 |
| DCAF12 |

PPP3CC
ABCB8
ABHD5
ADK
AGAP4
ANKRD26
AP4E1
AP5B1
ATAD3A
ATF7IP2
ATL2
BAALC-AS1
BCAN
BTAF1
C8orf76
CCDC112
CCDC24
CCDC28A
CCDC71L
CDK12
CHD1L
COL5A2
CPSF3
CSNK1G3
CTNS
CYBA
DENND6A
DLL4
DYNC2H1
EML6
ENTPD1
EPHX4
EXOSC7
EXT1
EXT2
FAM120A
FAM151B-DT
FCF1
FES
FMN1
FNTB
GALNT13
GALNT7

GALT
GATAD1
GMPR2
GON4L
GTF2E2
GUCD1
HEATR3
HSF4
HTR5A-AS1
HUNK
INTS8
ITGA3
KCTD4
KIF22
KLF3
KLHL18
L3HYPDH
LATS1
LEPROTL1
LMF1
LPCAT3
MECR
MEGF10
MIEF1
MN1
MRPL48
MSANTD1
MT1F
NAA16
NOM1
NSUN5P1
NUTM2B-AS1
OSBPL3
P3H2
PARL
PAXBP1
PBX3
PCCB
PEX19
PHF6
PIK3IP1
PRELID3B
PRR18

PTPN9
RANBP3
RBMS3
RCL1
RDX
REC8
RFPL2
RFX1
RIOK1
RPUSD4
RUBCN
SALL1
SBNO1-AS1
SFXN4
SGCD
SGF29
SHISAL1
SLC25A16
SLC25A29
SLC25A42
SLC43A2
SLC9A7
SLF2
SMARCE1
SMG8
SMURF2
SNRNP48
SOS1
SPAG5
SPTLC1
SRGAP3
SSPN
STAG1
STRIP2
SUFU
SYCE1L
TAF5L
TAFA1
TARS1
TARS3
TASOR2
TBC1D22A
TBC1D22B

TBC1D23
TCP11L1
THSD7A
TIGD5
TIMM8A
TLCD1
TMBIM4
TMEM38A
TMEM42
TMEM64
TNPO3
TNRC6C
TPT1-AS1
TRABD
TRAF3IP2-AS1
TRMT11
TRPV2
TTC21B
TYMP
URGCP
USP36
VIPAS39
VSTM2A
WHAMM
WIF1
ZBTB33
ZC3H4
ZC3HAV1
ZFAND2A-DT
ZFHX3
ZNF181
ZNF254
ZNF3
ZNF629
ZSCAN26

**y expressed genes (DEGs) in cortical layer 1 (L1) following 10 Hz rTMS.**

| Gene name | EntrezID |
|---|---|
| nuclear factor I A | 4774 |
| ChaC glutathione specific gamma-glutamylcyclotransferase 1 | 79094 |
| Fas associated via death domain | 8772 |
| AT-hook transcription factor | 80709 |
| bone morphogenetic protein 7 | 655 |
| caspase 4 | 837 |
| Fc gamma receptor Ia | 2209 |
| interleukin 32 | 9235 |
| MOK protein kinase | 5891 |
| nuclear factor kappa B subunit 2 | 4791 |
| protein tyrosine phosphatase non-receptor type 2 | 5771 |
| calcium voltage-gated channel subunit alpha1 C | 775 |
| potassium inwardly rectifying channel subfamily J member 16 | 3773 |
| sodium/potassium transporting ATPase interacting 4 | 128414 |
| Wnt family member 2B | 7482 |
| GRB2 associated binding protein 1 | 2549 |
| UDP glycosyltransferase 8 | 7368 |
| frizzled class receptor 1 | 8321 |
| lysine acetyltransferase 8 | 84148 |
| potassium channel tetramerization domain containing 10 | 83892 |
| distal-less homeobox 1 | 1745 |
| EH domain containing 4 | 30844 |
| erb-b2 receptor tyrosine kinase 4 | 2066 |
| FCH and mu domain containing endocytic adaptor 2 | 115548 |
| glial cell derived neurotrophic factor | 2668 |
| kinesin family member C3 | 3801 |
| leukotriene A4 hydrolase | 4048 |
| neuropilin and tolloid like 1 | 81832 |
| protocadherin 11 X-linked | 27328 |
| phosphodiesterase 7A | 5150 |
| phosphatidylinositol 4-kinase type 2 alpha | 55361 |
| parvalbumin | 5816 |
| vesicle transport through interaction with t-SNAREs 1A | 143187 |
| activin A receptor type 1C | 130399 |
| ADAM metallopeptidase domain 22 | 53616 |
| Rac/Cdc42 guanine nucleotide exchange factor 6 | 9459 |
| DAB adaptor protein 1 | 1600 |
| cerebellin 4 precursor | 140689 |
| DDB1 and CUL4 associated factor 12 | 25853 |

| | |
|---|---|
| protein phosphatase 3 catalytic subunit gamma | 5533 |
| ATP binding cassette subfamily B member 8 | 11194 |
| abhydrolase domain containing 5, lysophosphatidic acid acyltransferase | 51099 |
| adenosine kinase | 132 |
| ArfGAP with GTPase domain, ankyrin repeat and PH domain 4 | 119016 |
| ankyrin repeat domain containing 26 | 22852 |
| adaptor related protein complex 4 subunit epsilon 1 | 23431 |
| adaptor related protein complex 5 subunit beta 1 | 91056 |
| ATPase family AAA domain containing 3A | 55210 |
| activating transcription factor 7 interacting protein 2 | 80063 |
| atlastin GTPase 2 | 64225 |
| BAALC antisense RNA 1 | 100499183 |
| brevican | 63827 |
| B-TFIID TATA-box binding protein associated factor 1 | 9044 |
| chromosome 8 open reading frame 76 | 84933 |
| coiled-coil domain containing 112 | 153733 |
| coiled-coil domain containing 24 | 149473 |
| coiled-coil domain containing 28A | 25901 |
| coiled-coil domain containing 71 like | 168455 |
| cyclin dependent kinase 12 | 51755 |
| chromodomain helicase DNA binding protein 1 like | 9557 |
| collagen type V alpha 2 chain | 1290 |
| cleavage and polyadenylation specific factor 3 | 51692 |
| casein kinase 1 gamma 3 | 1456 |
| cystinosin, lysosomal cystine transporter | 1497 |
| cytochrome b-245 alpha chain | 1535 |
| DENN domain containing 6A | 201627 |
| delta like canonical Notch ligand 4 | 54567 |
| dynein cytoplasmic 2 heavy chain 1 | 79659 |
| EMAP like 6 | 400954 |
| ectonucleoside triphosphate diphosphohydrolase 1 | 953 |
| epoxide hydrolase 4 | 253152 |
| exosome component 7 | 23016 |
| exostosin glycosyltransferase 1 | 2131 |
| exostosin glycosyltransferase 2 | 2132 |
| family with sequence similarity 120 member A | 23196 |
| FAM151B divergent transcript | 121232370 |
| FCF1 rRNA-processing protein | 51077 |
| FES proto-oncogene, tyrosine kinase | 2242 |
| formin 1 | 342184 |
| farnesyltransferase, CAAX box, subunit beta | 2342 |
| polypeptide N-acetylgalactosaminyltransferase 13 | 114805 |
| polypeptide N-acetylgalactosaminyltransferase 7 | 51809 |

| | |
|---|---|
| galactose-1-phosphate uridylyltransferase | 2592 |
| GATA zinc finger domain containing 1 | 57798 |
| guanosine monophosphate reductase 2 | 51292 |
| gon-4 like | 54856 |
| general transcription factor IIE subunit 2 | 2961 |
| guanylyl cyclase domain containing 1 | 83606 |
| HEAT repeat containing 3 | 55027 |
| heat shock transcription factor 4 | 3299 |
| HTR5A antisense RNA 1 | 100128264 |
| hormonally up-regulated Neu-associated kinase | 30811 |
| integrator complex subunit 8 | 55656 |
| integrin subunit alpha 3 | 3675 |
| potassium channel tetramerization domain containing 4 | 386618 |
| kinesin family member 22 | 3835 |
| KLF transcription factor 3 | 51274 |
| kelch like family member 18 | 23276 |
| trans-L-3-hydroxyproline dehydratase | 112849 |
| large tumor suppressor kinase 1 | 9113 |
| leptin receptor overlapping transcript like 1 | 23484 |
| lipase maturation factor 1 | 64788 |
| lysophosphatidylcholine acyltransferase 3 | 10162 |
| mitochondrial trans-2-enoyl-CoA reductase | 51102 |
| multiple EGF like domains 10 | 84466 |
| mitochondrial elongation factor 1 | 54471 |
| MN1 proto-oncogene, transcriptional regulator | 4330 |
| mitochondrial ribosomal protein L48 | 51642 |
| Myb/SANT DNA binding domain containing 1 | 345222 |
| metallothionein 1F | 4494 |
| N-alpha-acetyltransferase 16, NatA auxiliary subunit | 79612 |
| nucleolar protein with MIF4G domain 1 | 64434 |
| NSUN5 pseudogene 1 | 155400 |
| NUTM2B antisense RNA 1 | 101060691 |
| oxysterol binding protein like 3 | 26031 |
| prolyl 3-hydroxylase 2 | 55214 |
| presenilin associated rhomboid like | 55486 |
| PAX3 and PAX7 binding protein 1 | 94104 |
| PBX homeobox 3 | 5090 |
| propionyl-CoA carboxylase subunit beta | 5096 |
| peroxisomal biogenesis factor 19 | 5824 |
| PHD finger protein 6 | 84295 |
| phosphoinositide-3-kinase interacting protein 1 | 113791 |
| PRELI domain containing 3B | 51012 |
| proline rich 18 | 285800 |

| | |
|---|---|
| protein tyrosine phosphatase non-receptor type 9 | 5780 |
| RAN binding protein 3 | 8498 |
| RNA binding motif single stranded interacting protein 3 | 27303 |
| RNA terminal phosphate cyclase like 1 | 10171 |
| radixin | 5962 |
| REC8 meiotic recombination protein | 9985 |
| ret finger protein like 2 | 10739 |
| regulatory factor X1 | 5989 |
| RIO kinase 1 | 83732 |
| RNA pseudouridine synthase D4 | 84881 |
| rubicon autophagy regulator | 9711 |
| spalt like transcription factor 1 | 6299 |
| SBNO1 antisense RNA 1 | 112268105 |
| sideroflexin 4 | 119559 |
| sarcoglycan delta | 6444 |
| SAGA complex associated factor 29 | 112869 |
| shisa like 1 | 85352 |
| solute carrier family 25 member 16 | 8034 |
| solute carrier family 25 member 29 | 123096 |
| solute carrier family 25 member 42 | 284439 |
| solute carrier family 43 member 2 | 124935 |
| solute carrier family 9 member A7 | 84679 |
| SMC5-SMC6 complex localization factor 2 | 55719 |
| SWI/SNF related BAF chromatin remodeling complex subunit E1 | 6605 |
| SMG8 nonsense mediated mRNA decay factor | 55181 |
| SMAD specific E3 ubiquitin protein ligase 2 | 64750 |
| small nuclear ribonucleoprotein U11/U12 subunit 48 | 154007 |
| SOS Ras/Rac guanine nucleotide exchange factor 1 | 6654 |
| sperm associated antigen 5 | 10615 |
| serine palmitoyltransferase long chain base subunit 1 | 10558 |
| SLIT-ROBO Rho GTPase activating protein 3 | 9901 |
| sarcospan | 8082 |
| STAG1 cohesin complex component | 10274 |
| striatin interacting protein 2 | 57464 |
| SUFU negative regulator of hedgehog signaling | 51684 |
| synaptonemal complex central element protein 1 like | 100130958 |
| TATA-box binding protein associated factor 5 like | 27097 |
| TAFA chemokine like family member 1 | 407738 |
| threonyl-tRNA synthetase 1 | 6897 |
| threonyl-tRNA synthetase 3 | 123283 |
| transcription activation suppressor family member 2 | 54906 |
| TBC1 domain family member 22A | 25771 |
| TBC1 domain family member 22B | 55633 |

| | |
|---|---|
| TBC1 domain family member 23 | 55773 |
| t-complex 11 like 1 | 55346 |
| thrombospondin type 1 domain containing 7A | 221981 |
| tigger transposable element derived 5 | 84948 |
| translocase of inner mitochondrial membrane 8A | 1678 |
| TLC domain containing 1 | 116238 |
| transmembrane BAX inhibitor motif containing 4 | 51643 |
| transmembrane protein 38A | 79041 |
| transmembrane protein 42 | 131616 |
| transmembrane protein 64 | 169200 |
| transportin 3 | 23534 |
| trinucleotide repeat containing adaptor 6C | 57690 |
| TPT1 antisense RNA 1 | 100190939 |
| TraB domain containing | 80305 |
| TRAF3IP2 antisense RNA 1 | 643749 |
| tRNA methyltransferase 11 homolog | 60487 |
| transient receptor potential cation channel subfamily V member 2 | 51393 |
| tetratricopeptide repeat domain 21B | 79809 |
| thymidine phosphorylase | 1890 |
| upregulator of cell proliferation | 55665 |
| ubiquitin specific peptidase 36 | 57602 |
| VPS33B interacting protein, apical-basolateral polarity regulator, spe-39 homolog | 63894 |
| V-set and transmembrane domain containing 2A | 222008 |
| WASP homolog associated with actin, golgi membranes and microtubules | 123720 |
| WNT inhibitory factor 1 | 11197 |
| zinc finger and BTB domain containing 33 | 10009 |
| zinc finger CCCH-type containing 4 | 23211 |
| zinc finger CCCH-type containing, antiviral 1 | 56829 |
| ZFAND2A divergent transcript | 101927021 |
| zinc finger homeobox 3 | 463 |
| zinc finger protein 181 | 339318 |
| zinc finger protein 254 | 9534 |
| zinc finger protein 3 | 7551 |
| zinc finger protein 629 | 23361 |
| zinc finger and SCAN domain containing 26 | 7741 |

| Gene category | Donor 1 | | Don |
|---|---|---|---|
| | log2fc | p.adj | log2fc |
| | 1.537265759 | 1 | -1.081411126 |
| cell death | 1.537265759 | 1 | -1.344445531 |
| cell death | 2.537265759 | 1 | -1.344445531 |
| immune response | 2.537265759 | 1 | -1.929408032 |
| immune response | 1.122228259 | 1 | -1.929408032 |
| immune response | 2.537265759 | 1 | -1.344445531 |
| immune response | 2.537265759 | 1 | -1.666373626 |
| immune response | 1.537265759 | 1 | -1.929408032 |
| immune response | 1.122228259 | 1 | -1.344445531 |
| immune response | 3.122228259 | 1 | -1.344445531 |
| immune response | -1.047696742 | 1 | -3.344445531 |
| intrinsinc plasticity | 1.274231353 | 1 | -2.344445531 |
| intrinsinc plasticity | 1.537265759 | 1 | -1.344445531 |
| intrinsinc plasticity | 2.022692586 | 1 | 1.530023586 |
| mitochondria | 1.537265759 | 1 | -1.344445531 |
| myelin | -1.462734241 | 1 | -2.666373626 |
| myelin | 1.537265759 | 1 | -1.566837953 |
| neurogenesis | 1.215337664 | 1 | -1.344445531 |
| neurogenesis | -2.632659243 | 1 | -2.929408032 |
| neurogenesis | 1.952303258 | 1 | -2.929408032 |
| synapse | 2.122228259 | 1 | -1.344445531 |
| synapse | 1.537265759 | 1 | -1.666373626 |
| synapse | 1.122228259 | 1 | -1.344445531 |
| synapse | 1.537265759 | 1 | -1.344445531 |
| synapse | 1.537265759 | 1 | -1.929408032 |
| synapse | 1.215337664 | 1 | -1.344445531 |
| synapse | 1.051838932 | 1 | -2.344445531 |
| synapse | -2.270089163 | 1 | -1.344445531 |
| synapse | 1.537265759 | 1 | -1.344445531 |
| synapse | -1.047696742 | 1 | -1.344445531 |
| synapse | 2.022692586 | 1 | -1.759483031 |
| synapse | 2.952303258 | 1 | -1.929408032 |
| synapse | 1.537265759 | 1 | -2.344445531 |
| synapse/synaptic plasticity | 2.122228259 | 1 | -1.344445531 |
| synapse/synaptic plasticity | -1.369624837 | 1 | -1.344445531 |
| synapse/synaptic plasticity | 1.537265759 | 1 | -1.929408032 |
| synapse | 2.537265759 | 1 | -1.344445531 |
| synaptic plasticity | 1.274231353 | 1 | -1.344445531 |
| synaptic plasticity | -1.270089163 | 1 | -1.344445531 |

| | | | |
|---|---|---|---|
| synaptic plasticity/ immune response | 1.537265759 | 1 | 1.240516969 |
| | -1.047696742 | 1 | -1.344445531 |
| | -1.462734241 | 1 | -2.344445531 |
| | 1.537265759 | 1 | -1.344445531 |
| | 1.537265759 | 1 | -1.344445531 |
| | -1.047696742 | 1 | -2.666373626 |
| | 2.537265759 | 1 | -1.344445531 |
| | 1.122228259 | 1 | -1.344445531 |
| | 1.537265759 | 1 | -1.344445531 |
| | 2.344620681 | 1 | -2.666373626 |
| | 1.952303258 | 1 | -1.666373626 |
| | 3.122228259 | 1 | -1.344445531 |
| | 1.03030577 | 1 | 1.103013446 |
| | -1.270089163 | 1 | 1.655554469 |
| | 2.344620681 | 1 | -1.344445531 |
| | -1.632659243 | 1 | -1.344445531 |
| | 1.537265759 | 1 | -1.344445531 |
| | -1.784662336 | 1 | -1.344445531 |
| | 2.274231353 | 1 | -1.344445531 |
| | 1.952303258 | 1 | -1.344445531 |
| | 2.537265759 | 1 | -2.344445531 |
| | 1.537265759 | 1 | -1.344445531 |
| | 2.537265759 | 1 | -1.344445531 |
| | 1.122228259 | 1 | -1.929408032 |
| | 2.122228259 | 1 | -2.344445531 |
| | 1.051838932 | 1 | -1.344445531 |
| | -1.047696742 | 1 | -1.344445531 |
| | 2.537265759 | 1 | -1.929408032 |
| | 1.122228259 | 1 | -2.344445531 |
| | 1.122228259 | 1 | -1.344445531 |
| | 1.537265759 | 1 | -1.929408032 |
| | -1.047696742 | 1 | -1.344445531 |
| | 2.122228259 | 1 | 1.240516969 |
| | 1.537265759 | 1 | -1.344445531 |
| | 1.537265759 | 1 | -1.344445531 |
| | 1.122228259 | 1 | -1.666373626 |
| | 1.537265759 | 1 | -1.344445531 |
| | -1.047696742 | 1 | 1.655554469 |
| | 1.537265759 | 1 | -1.929408032 |
| | -1.270089163 | 1 | -1.344445531 |
| | 3.537265759 | 1 | -1.344445531 |
| | 1.859193854 | 1 | -1.344445531 |
| | 1.537265759 | 1 | -1.929408032 |

| | | |
|---|---|---|
| -1.270089163 | 1 | -1.344445531 |
| -1.270089163 | 1 | -1.344445531 |
| -1.270089163 | 1 | 1.655554469 |
| -1.047696742 | 1 | -1.344445531 |
| 1.537265759 | 1 | -1.344445531 |
| 3.122228259 | 1 | -1.929408032 |
| 1.122228259 | 1 | -1.344445531 |
| 1.537265759 | 1 | -1.929408032 |
| 1.122228259 | 1 | -2.344445531 |
| 1.537265759 | 1 | -1.929408032 |
| 1.215337664 | 1 | -1.344445531 |
| 2.537265759 | 1 | -1.344445531 |
| 1.314873337 | 1 | -1.344445531 |
| 1.537265759 | 1 | -2.344445531 |
| 1.537265759 | 1 | 1.655554469 |
| 1.537265759 | 1 | -1.344445531 |
| 1.537265759 | 1 | -1.344445531 |
| 1.952303258 | 1 | -1.344445531 |
| 1.051838932 | 1 | -2.666373626 |
| 1.537265759 | 1 | -1.344445531 |
| 1.537265759 | 1 | -1.666373626 |
| 1.215337664 | 1 | -1.344445531 |
| 1.122228259 | 1 | -1.929408032 |
| -1.047696742 | 1 | -1.929408032 |
| 1.122228259 | 1 | -1.344445531 |
| 1.537265759 | 1 | -1.929408032 |
| 1.537265759 | 1 | -1.344445531 |
| 1.122228259 | 1 | -1.132941426 |
| 1.537265759 | 1 | -1.344445531 |
| 1.537265759 | 1 | -1.929408032 |
| 1.537265759 | 1 | -1.929408032 |
| 3.122228259 | 1 | -1.344445531 |
| -1.047696742 | 1 | -1.344445531 |
| 1.537265759 | 1 | -2.344445531 |
| 1.122228259 | 1 | -2.344445531 |
| 1.122228259 | 1 | -1.344445531 |
| -1.270089163 | 1 | -1.929408032 |
| 1.537265759 | 1 | -1.929408032 |
| -1.462734241 | 1 | -2.344445531 |
| -1.92216586 | 1 | -1.929408032 |
| 1.215337664 | 1 | -1.344445531 |
| 1.274231353 | 1 | 1.070591968 |
| -1.047696742 | 1 | -1.514370533 |

| | | |
|---|---|---|
| 1.537265759 | 1 | 1.655554469 |
| -1.270089163 | 1 | -1.666373626 |
| 2.537265759 | 1 | -1.344445531 |
| 1.537265759 | 1 | -1.344445531 |
| 1.537265759 | 1 | -1.722957155 |
| 1.537265759 | 1 | -1.929408032 |
| 2.537265759 | 1 | -1.344445531 |
| 1.122228259 | 1 | -1.929408032 |
| -1.047696742 | 1 | -2.929408032 |
| 1.952303258 | 1 | -1.344445531 |
| -1.784662336 | 1 | -1.344445531 |
| 2.537265759 | 1 | -1.344445531 |
| 2.537265759 | 1 | -1.344445531 |
| 1.122228259 | 1 | -2.344445531 |
| 2.122228259 | 1 | -1.929408032 |
| 1.537265759 | 1 | -1.929408032 |
| 1.051838932 | 1 | -1.929408032 |
| 1.537265759 | 1 | -1.344445531 |
| 1.75965818 | 1 | 1.655554469 |
| -1.784662336 | 1 | -2.344445531 |
| 1.274231353 | 1 | -2.151800454 |
| 1.051838932 | 1 | -2.344445531 |
| -1.047696742 | 1 | -1.344445531 |
| 2.274231353 | 1 | -2.666373626 |
| 1.537265759 | 1 | -1.929408032 |
| 1.537265759 | 1 | -1.929408032 |
| 1.537265759 | 1 | -1.344445531 |
| 1.122228259 | 1 | -1.666373626 |
| 3.122228259 | 1 | -1.344445531 |
| -1.632659243 | 1 | -1.566837953 |
| -1.270089163 | 1 | -1.081411126 |
| 1.537265759 | 1 | -1.929408032 |
| 1.537265759 | 1 | -2.929408032 |
| 2.122228259 | 1 | -1.344445531 |
| 1.537265759 | 1 | -1.344445531 |
| -1.462734241 | 1 | -1.344445531 |
| 3.537265759 | 1 | -1.344445531 |
| 1.158754135 | 1 | -1.344445531 |
| 1.537265759 | 1 | -1.566837953 |
| 3.122228259 | 1 | -1.566837953 |
| 1.952303258 | 1 | -1.344445531 |
| -1.047696742 | 1 | -1.666373626 |
| 1.122228259 | 1 | -1.344445531 |

| | | |
|---|---|---|
| -1.462734241 | 1 | -1.344445531 |
| 1.537265759 | 1 | -1.344445531 |
| 2.274231353 | 1 | -1.929408032 |
| 1.537265759 | 1 | -1.344445531 |
| 1.122228259 | 1 | 1.240516969 |
| 2.537265759 | 1 | -1.344445531 |
| -1.047696742 | 1 | -1.344445531 |
| 1.022692586 | 1 | 1.240516969 |
| -1.784662336 | 1 | -1.929408032 |
| 1.122228259 | 1 | -1.929408032 |
| -1.270089163 | 1 | -1.344445531 |
| 2.859193854 | 1 | -1.344445531 |
| 2.122228259 | 1 | -1.344445531 |
| 2.122228259 | 1 | -1.344445531 |
| -1.047696742 | 1 | -1.929408032 |
| -1.047696742 | 1 | -1.344445531 |
| -1.047696742 | 1 | -1.344445531 |
| 1.537265759 | 1 | 1.240516969 |
| 2.537265759 | 1 | -2.151800454 |
| 2.122228259 | 1 | -1.344445531 |
| 1.800300165 | 1 | -1.344445531 |
| 3.537265759 | 1 | -1.344445531 |
| 1.006751042 | 1 | -1.344445531 |
| -1.270089163 | 1 | -1.081411126 |
| 2.952303258 | 1 | -1.344445531 |
| 2.537265759 | 1 | -2.344445531 |
| 1.537265759 | 1 | -1.929408032 |
| -1.270089163 | 1 | -1.344445531 |
| 1.537265759 | 1 | -1.344445531 |
| -1.047696742 | 1 | -1.929408032 |
| 1.537265759 | 1 | -1.344445531 |
| -1.047696742 | 1 | -1.344445531 |
| 1.537265759 | 1 | -1.344445531 |
| 1.537265759 | 1 | -1.344445531 |
| 2.122228259 | 1 | -1.929408032 |

| ior 2 | Donor 4 | |
|---|---|---|
| p.adj | log2fc | p.adj |
| 1 | -1.011865222 | 1 |
| 1 | 1.177477233 | 1 |
| 1 | -1.731757303 | 1 |
| 1 | 1.075597619 | 1 |
| 1 | 1.001597037 | 1 |
| 1 | -1.731757303 | 1 |
| 1 | -1.071243769 | 1 |
| 1 | -2.094327383 | 1 |
| 1 | 1.616166 | 1 |
| 1 | 1.993135459 | 1 |
| 1 | 1.29799004 | 1 |
| 1 | 1.158059779 | 1 |
| 1 | 1.075597619 | 1 |
| 1 | -1.246330476 | 1 |
| 1 | 1.075597619 | 1 |
| 1 | -1.061905905 | 1 |
| 1 | -1.039879599 | 1 |
| 1 | 1.075597619 | 1 |
| 1 | 1.168707023 | 1 |
| 1 | 1.1495982 | 1 |
| 1 | 1.753669524 | 1 |
| 1 | 1.268242697 | 1 |
| 1 | 1.268242697 | 1 |
| 1 | 1.075597619 | 1 |
| 1 | 1.66056012 | 1 |
| 1 | 1.397525714 | 1 |
| 1 | -1.094327383 | 1 |
| 1 | 1.256169865 | 1 |
| 1 | 1.490635118 | 1 |
| 1 | 1.590170792 | 1 |
| 1 | 1.371053502 | 1 |
| 1 | 1.905672617 | 1.14E-10 |
| 1 | -1.924402381 | 1 |
| 1 | 3.24552262 | 1 |
| 1 | 1.039973709 | 1 |
| 1 | -1.146794802 | 1 |
| 1 | 1.075597619 | 1 |
| 1 | 1.968682415 | 1 |
| 1 | 1.812563213 | 1 |

| | | |
|---|---|---|
| 1 | 2.075597619 | 1 |
| 1 | -1.924402381 | 1 |
| 1 | -1.316719804 | 1 |
| 1 | 1.753669524 | 1 |
| 1 | 1.950066737 | 1 |
| 1 | -1.246330476 | 1 |
| 1 | 1.075597619 | 1 |
| 1 | 1.075597619 | 1 |
| 1 | 1.075597619 | 1 |
| 1 | 1.423520922 | 1 |
| 1 | 1.462620742 | 1 |
| 1 | 1.075597619 | 1 |
| 1 | -1.25697772 | 1 |
| 1 | 1.24552262 | 1 |
| 1 | 1.213101143 | 1 |
| 1 | -1.509364882 | 1 |
| 1 | 1.268242697 | 1 |
| 1 | 1.66056012 | 1 |
| 1 | -2.383834 | 1 |
| 1 | 1.338632025 | 1 |
| 1 | 1.24552262 | 1 |
| 1 | 2.397525714 | 1 |
| 1 | -1.509364882 | 1 |
| 1 | 1.753669524 | 1 |
| 1 | 1.29799004 | 1 |
| 1 | -1.782383376 | 1 |
| 1 | 1.397525714 | 1 |
| 1 | 1.075597619 | 1 |
| 1 | 1.812563213 | 1 |
| 1 | 1.882952541 | 1 |
| 1 | -1.246330476 | 1 |
| 1 | 1.227600712 | 1 |
| 1 | 1.490635118 | 1 |
| 1 | 1.011467281 | 1 |
| 1 | 1.168707023 | 1 |
| 1 | -1.646868406 | 1 |
| 1 | 1.075597619 | 1 |
| 1 | 1.075597619 | 1 |
| 1 | 1.590170792 | 1 |
| 1 | 1.561024446 | 1 |
| 1 | 1.397525714 | 1 |
| 1 | 1.075597619 | 1 |
| 1 | 2.490635118 | 1 |

| | | |
|---|---|---|
| 1 | 1.982488215 | 1 |
| 1 | -1.246330476 | 1 |
| 1 | 1.134491308 | 1 |
| 1 | 1.71302754 | 1 |
| 1 | 1.227600712 | 1 |
| 1 | -1.146794802 | 1 |
| 1 | 1.66056012 | 1 |
| 1 | 1.075597619 | 1 |
| 1 | 1.365104236 | 1 |
| 1 | 2.24552262 | 1 |
| 1 | 1.490635118 | 1 |
| 1 | 1.075597619 | 1 |
| 1 | 2.397525714 | 1 |
| 1 | 1.126223692 | 1 |
| 1 | -1.98329607 | 1 |
| 1 | 1.005208291 | 1 |
| 1 | 1.490635118 | 1 |
| 1 | 1.338632025 | 1 |
| 1 | 1.001597037 | 1 |
| 1 | 1.950066737 | 1 |
| 1 | 1.29799004 | 1 |
| 1 | 1.075597619 | 1 |
| 1 | -1.039879599 | 1 |
| 1 | 2.29799004 | 1 |
| 1 | 1.075597619 | 1 |
| 1 | 1.075597619 | 1 |
| 1 | 1.535029238 | 1 |
| 1 | 1.528109824 | 0.075105415 |
| 1 | -1.509364882 | 1 |
| 1 | 2.66056012 | 1 |
| 1 | -2.094327383 | 1 |
| 1 | 1.075597619 | 1 |
| 1 | 1.075597619 | 1 |
| 1 | -1.924402381 | 1 |
| 1 | -1.246330476 | 1 |
| 1 | 1.227600712 | 1 |
| 1 | 1.075597619 | 1 |
| 1 | 1.075597619 | 1 |
| 1 | 1.24552262 | 1 |
| 1 | 1.075597619 | 1 |
| 1 | -1.094327383 | 1 |
| 1 | -1.094327383 | 1 |
| 1 | -1.117047459 | 1 |

| | | |
|---|---|---|
| 1 | 1.075597619 | 1 |
| 1 | 1.075597619 | 1 |
| 1 | -2.246330476 | 1 |
| 1 | 1.66056012 | 1 |
| 1 | -1.028739041 | 1 |
| 1 | 1.075597619 | 1 |
| 1 | 1.075597619 | 1 |
| 1 | 1.561024446 | 1 |
| 1 | 1.812563213 | 1 |
| 1 | 1.075597619 | 1 |
| 1 | 1.126223692 | 1 |
| 1 | 1.075597619 | 1 |
| 1 | -1.039879599 | 1 |
| 1 | 1.075597619 | 1 |
| 1 | 1.075597619 | 1 |
| 1 | 1.24552262 | 1 |
| 1 | 1.853205198 | 1 |
| 1 | 1.075597619 | 1 |
| 1 | 1.416634537 | 1 |
| 1 | 1.365104236 | 1 |
| 1 | 1.582557608 | 1 |
| 1 | 1.075597619 | 1 |
| 1 | -1.302914004 | 1 |
| 1 | 1.075597619 | 1 |
| 1 | 2.075597619 | 1 |
| 1 | 1.561024446 | 1 |
| 1 | 1.812563213 | 1 |
| 1 | 1.213101143 | 1 |
| 1 | 1.29799004 | 1 |
| 1 | -1.071243769 | 1 |
| 1 | 1.121401309 | 1 |
| 1 | -1.039879599 | 1 |
| 1 | 1.523056596 | 1 |
| 1 | 1.24552262 | 1 |
| 1 | 1.075597619 | 1 |
| 1 | 1.338632025 | 1 |
| 1 | 1.490635118 | 1 |
| 1 | 1.753669524 | 1 |
| 1 | 1.130045403 | 1 |
| 1 | -1.061905905 | 1 |
| 1 | 1.075597619 | 1 |
| 1 | 1.075597619 | 1 |
| 1 | -1.039879599 | 1 |

| | | |
|---|---|---|
| 1 | 1.703628842 | 1 |
| 1 | 1.882952541 | 1 |
| 1 | 1.075597619 | 1 |
| 1 | 1.882952541 | 1 |
| 1 | 1.561024446 | 1 |
| 1 | 1.075597619 | 1 |
| 1 | -1.28697246 | 1 |
| 1 | 1.021149835 | 1 |
| 1 | 1.075597619 | 1 |
| 1 | -1.509364882 | 1 |
| 1 | 1.075597619 | 1 |
| 1 | 1.1495982 | 1 |
| 1 | -2.246330476 | 1 |
| 1 | 1.075597619 | 1 |
| 1 | -2.094327383 | 1 |
| 1 | 1.438167698 | 1 |
| 1 | -1.246330476 | 1 |
| 1 | 1.075597619 | 1 |
| 1 | -1.831292977 | 1 |
| 1 | 1.075597619 | 1 |
| 1 | 1.019014091 | 1 |
| 1 | 1.075597619 | 1 |
| 1 | 1.389706209 | 0.012791322 |
| 1 | 1.66056012 | 1 |
| 1 | 1.898719857 | 1 |
| 1 | 1.29799004 | 1 |
| 1 | 1.438167698 | 1 |
| 1 | -1.246330476 | 1 |
| 1 | -1.509364882 | 1 |
| 1 | -2.094327383 | 1 |
| 1 | 1.24552262 | 1 |
| 1 | 1.075597619 | 1 |
| 1 | 1.812563213 | 1 |
| 1 | -1.509364882 | 1 |
| 1 | 1.075597619 | 1 |

**Data S4. Differentially expressed genes (DEGs) in cortical layer 2/3 (L2/3) following 10 Hz r[illegible]**

| Gene symbol | Gene name | EntrezID |
|---|---|---|
| CBLN4 | cerebellin 4 precursor | 140689 |
| COL5A2 | collagen type V alpha 2 chain | 1290 |
| FOXP2 | forkhead box P2 | 93986 |
| FOXP4 | forkhead box P4 | 116113 |
| GLRA3 | glycine receptor alpha 3 | 8001 |
| GRIN3A | glutamate ionotropic receptor NMDA type subunit 3A | 116443 |
| IL1RN | interleukin 1 receptor antagonist | 3557 |
| INO80D | INO80 complex subunit D | 54891 |
| KANSL1-AS1 | KANSL1 antisense RNA 1 | 644246 |
| KCNJ16 | potassium inwardly rectifying channel subfamily J member 16 | 3773 |
| LAMB1 | laminin subunit beta 1 | 3912 |
| SERPINH1 | serpin family H member 1 | 871 |
| SLIT2 | slit guidance ligand 2 | 9353 |
| SNURF | SNRPN upstream open reading frame | 8926 |
| TMEM125 | transmembrane protein 125 | 128218 |
| ZNF740 | zinc finger protein 740 | 283337 |

**ΓMS.**

| **Gene category** | Donor 1 | | Donor 2 | |
|---|---|---|---|---|
| | log2fc | p.adj | log2fc | p.adj |
| synaptic plasticity | 1.917485291 | 0.000407153 | 1.172364047 | 1 |
| | 2.910155749 | 2.99E-06 | 1.494292142 | 1 |
| synapse | -1.365968656 | 1 | -1.827635953 | 1 |
| | -1.266432982 | 1 | -1.149564048 | 1 |
| intrinsnc plasticity | 1.510548291 | 1 | 1.494292142 | 1 |
| synaptic plasticity | -1.529467388 | 1 | -1.206147576 | 1 |
| immune response | 1.033961951 | 1 | 1.662689673 | 1 |
| | 1.849044235 | 1 | -1.412598454 | 1 |
| | -1.044040561 | 1 | 1.172364047 | 1 |
| intrinsnc plasticity | 1.955959439 | 1 | 1.172364047 | 1 |
| | 1.228977933 | 1 | 1.020360953 | 1 |
| | 1.955959439 | 1 | 1.342289048 | 1 |
| | 1.318529518 | 1 | -1.753635372 | 1 |
| | -1.044040561 | 1 | 1.287841264 | 1 |
| | -1.629003062 | 1 | 1.757326548 | 1 |
| | -1.143576235 | 1 | -2.287067572 | 1 |

| Donor 3 | | Donor 4 | |
|---|---|---|---|
| log2fc | p.adj | log2fc | p.adj |
| 1.175999516 | 1 | -3.36923381 | 0.708843425 |
| 1.591037015 | 1 | -1.900464326 | 1 |
| -1.145928579 | 1 | 1.087462841 | 1 |
| 2.591037015 | 1 | 1.754887502 | 1 |
| 1.398391937 | 1 | -1.222392421 | 1 |
| 2.175999516 | 1 | -1.115477217 | 1 |
| 1.91296511 | 1 | -1.415037499 | 0.079337145 |
| 1.175999516 | 1 | 1.021061616 | 1 |
| -2.73089108 | 1 | 1.192645078 | 1 |
| 1.175999516 | 1 | 1.263034406 | 1 |
| 2.175999516 | 1 | -2.662965013 | 1 |
| 1.175999516 | 1 | 1.432959407 | 1 |
| 1.175999516 | 1 | 1.169925001 | 1 |
| -1.993925486 | 1 | 1.672425342 | 1 |
| 1.175999516 | 1 | -1.187627003 | 1 |
| 1.91296511 | 1 | 1.044394119 | 1 |

**Data S5. Differentially expressed genes (DEGs) in cortical layer 5 (L5) following 10 Hz rTMS.**

| Gene symbol | Gene name | EntrezID |
|---|---|---|
| SLC38A3 | solute carrier family 38 member 3 | 10991 |
| CCL8 | C-C motif chemokine ligand 8 | 6355 |
| KCNS1 | potassium voltage-gated channel modifier subfamily S member 1 | 3787 |
| RBFOX3 | RNA binding fox-1 homolog 3 | 146713 |
| LAMP5 | lysosomal associated membrane protein family member 5 | 24141 |
| ADCY8 | adenylate cyclase 8 | 114 |
| ASPA | aspartoacylase | 443 |
| ASPHD2 | aspartate beta-hydroxylase domain containing 2 | 57168 |
| CENPP | centromere protein P | 401541 |
| CSF3 | colony stimulating factor 3 | 1440 |
| EIF1AY | eukaryotic translation initiation factor 1A Y-linked | 9086 |
| FAM149B1 | family with sequence similarity 149 member B1 | 317662 |
| FBXO6 | F-box protein 6 | 26270 |
| FGGY | FGGY carbohydrate kinase domain containing | 55277 |
| FUT11 | fucosyltransferase 11 | 170384 |
| GARNL3 | GTPase activating Rap/RanGAP domain like 3 | 84253 |
| GJA4 | gap junction protein alpha 4 | 2701 |
| GPR26 | G protein-coupled receptor 26 | 2849 |
| HBA1 | hemoglobin subunit alpha 1 | 3039 |
| HBA2 | hemoglobin subunit alpha 2 | 3040 |
| HEXIM2 | HEXIM P-TEFb complex subunit 2 | 124790 |
| LIN54 | lin-54 DREAM MuvB core complex component | 132660 |
| LINC03025 | long intergenic non-protein coding RNA 3025 | 440896 |
| LNCBRM | lncRNA SMARCA2 (BRM) associated | 100996645 |
| MTMR7 | myotubularin related protein 7 | 9108 |
| NAA25 | N-alpha-acetyltransferase 25, NatB auxiliary subunit | 80018 |
| NAA30 | N-alpha-acetyltransferase 30, NatC catalytic subunit | 122830 |
| NKX3-1 | NK3 homeobox 1 | 4824 |
| PNOC | prepronociceptin | 5368 |
| PRKD2 | protein kinase D2 | 25865 |
| PRRG3 | proline rich and Gla domain 3 | 79057 |
| PTCHD1 | patched domain containing 1 | 139411 |
| RPP40 | ribonuclease P/MRP subunit p40 | 10799 |
| SCYL3 | SCY1 like pseudokinase 3 | 57147 |
| SH2B2 | SH2B adaptor protein 2 | 10603 |
| SH3D19 | SH3 domain containing 19 | 152503 |
| SLC12A8 | solute carrier family 12 member 8 | 84561 |
| SLC35B3 | solute carrier family 35 member B3 | 51000 |
| SNED1 | sushi, nidogen and EGF like domains 1 | 25992 |

| | | |
|---|---|---|
| SOX6 | SRY-box transcription factor 6 | 55553 |
| TBC1D23 | TBC1 domain family member 23 | 55773 |
| UTP20 | UTP20 small subunit processome component | 27340 |
| ZFYVE26 | zinc finger FYVE-type containing 26 | 23503 |
| ZNF212 | zinc finger protein 212 | 7988 |
| ZNF222 | zinc finger protein 222 | 7673 |
| ZNF567 | zinc finger protein 567 | 163081 |
| ZNF766 | zinc finger protein 766 | 90321 |

| **Gene category** | Donor 1 | | Don |
|---|---|---|---|
| | log2fc | p.adj | log2fc |
| neurotransmitter | 1.176259567 | 1 | -1.19478019 |
| immune response | 1.913225161 | 1 | 1.127147905 |
| intrinsic plasticity | 1.038756043 | 1 | -1.406284295 |
| intrinsinc plasticity/synaptic plasticity | -1.018577132 | 1 | 1.022450526 |
| synapse | 1.176259567 | 1 | -1.8806716 |
| synaptic plasticity | 1.623718544 | 1 | -1.872852095 |
| | 1.846110966 | 1 | 1.80521981 |
| | 1.126218885 | 1 | -1.697280531 |
| | -1.183636378 | 1 | 1.390182311 |
| | -1.048706798 | 1 | -1.320311072 |
| | -1.224278362 | 1 | 1.127147905 |
| | -1.546206457 | 1 | -1.19478019 |
| | -1.546206457 | 1 | -1.19478019 |
| | -1.283172051 | 1 | -1.417172611 |
| | 1.038756043 | 1 | 1.220257309 |
| | -1.020137646 | 1 | 1.027612231 |
| | 1.038756043 | 1 | -1.364705192 |
| | 1.089382117 | 1 | -1.542703494 |
| | -1.768598879 | 1 | -1.846856887 |
| | -1.633669298 | 1 | -1.109891293 |
| | 2.360684138 | 1 | -2.19478019 |
| | -1.039246469 | 1 | 1.220257309 |
| | -1.768598879 | 1 | -1.19478019 |
| | -1.484805913 | 1 | -1.609817689 |
| | -1.11324705 | 1 | -1.364705192 |
| | -1.283172051 | 1 | -1.002135112 |
| | -1.005638076 | 1 | -1.429245444 |
| | -2.283172051 | 1 | -1.387425268 |
| | -1.020137646 | 1 | -1.095244517 |
| | 1.846110966 | 1 | 1.612574732 |
| | -1.183636378 | 1 | -1.042777097 |
| | -1.131168958 | 1 | -1.19478019 |
| | -1.283172051 | 1 | -1.19478019 |
| | -1.076721174 | 1 | -1.609817689 |
| | -1.131168958 | 1 | -2.516708285 |
| | -1.546206457 | 1 | 1.390182311 |
| | 2.360684138 | 1 | -1.457814596 |
| | -1.131168958 | 1 | -2.364705192 |
| | 1.176259567 | 1 | -2.042777097 |

| | | |
|---|---|---|
| 1.453793543 | 1 | 1.127147905 |
| -1.202252056 | 1 | -1.294315864 |
| 1.261148465 | 1 | -1.931745784 |
| 1.453793543 | 1 | -1.002135112 |
| 1.261148465 | 1 | -1.609817689 |
| 1.453793543 | 1 | -1.609817689 |
| -1.183636378 | 1 | -1.19478019 |
| 2.208681045 | 1 | -1.19478019 |

| ...or 2 | Donor 4 | |
|---|---|---|
| p.adj | log2fc | p.adj |
| 1 | -1.054672906 | 1 |
| 1 | 1.230729313 | 1 |
| 1 | -3.014030922 | 0.262156518 |
| 1 | -1.315539475 | 0.002446832 |
| 0.103999173 | -2.483516205 | 1.02E-06 |
| 1 | -1.277065327 | 1 |
| 1 | 1.803308089 | 1 |
| 1 | -1.692102827 | 1 |
| 1 | 1.378286501 | 1 |
| 1 | -2.820207652 | 0.02394175 |
| 1 | -1.126822692 | 1 |
| 1 | -1.306211673 | 1 |
| 1 | -1.929142024 | 1 |
| 1 | -1.469710405 | 1 |
| 1 | -1.121787102 | 1 |
| 1 | -1.157766399 | 1 |
| 1 | -1.317707312 | 1 |
| 1 | -1.054672906 | 1 |
| 1 | 1.399045061 | 1 |
| 1 | 1.814592617 | 1 |
| 1 | -1.054672906 | 1 |
| 1 | -2.054672906 | 1 |
| 1 | 1.115252095 | 1 |
| 1 | -1.317707312 | 1 |
| 1 | -1.054672906 | 1 |
| 1 | -1.157766399 | 1 |
| 1 | -1.000225122 | 1 |
| 1 | -1.054672906 | 1 |
| 1 | -2.585187623 | 1 |
| 1 | -1.147782311 | 1 |
| 1 | -1.469710405 | 1 |
| 1 | 1.044862767 | 1 |
| 1 | -1.054672906 | 1 |
| 1 | 1.015716422 | 1 |
| 1 | -1.972210746 | 1 |
| 1 | -1.972210746 | 1 |
| 1 | -1.235245152 | 1 |
| 1 | -1.395709824 | 1 |
| 1 | -1.132675418 | 1 |

| | | |
|---|---|---|
| 1 | -2.277065327 | 1 |
| 1 | -1.101978621 | 1 |
| 1 | -1.054672906 | 1 |
| 1 | -1.344179523 | 1 |
| 1 | -1.929142024 | 1 |
| 1 | -1.344179523 | 1 |
| 1 | -1.469710405 | 1 |
| 1 | 1.530289595 | 1 |

**Data S6. Differentially expressed genes (DEGs) in cortical layer 6 (L6) following 10 Hz rTMS.**

| Gene symbol | Gene name | EntrezID |
|---|---|---|
| FAS | Fas cell surface death receptor | 355 |
| KCNQ1OT1 | KCNQ1 opposite strand/antisense transcript 1 | 10984 |
| SOCS5 | suppressor of cytokine signaling 5 | 9655 |
| KCNAB3 | potassium voltage-gated channel subfamily A regulatory beta subunit 3 | 9196 |
| COQ5 | coenzyme Q5, methyltransferase | 84274 |
| EXOG | exo/endonuclease G | 9941 |
| GSAP | gamma-secretase activating protein | 54103 |
| METTL17 | methyltransferase like 17 | 64745 |
| MRS2 | magnesium transporter MRS2 | 57380 |
| POLL | DNA polymerase lambda | 27343 |
| SLC25A40 | solute carrier family 25 member 40 | 55972 |
| MYO1D | myosin ID | 4642 |
| ZBTB45 | zinc finger and BTB domain containing 45 | 84878 |
| ALKBH3 | alkB homolog 3, alpha-ketoglutarate dependent dioxygenase | 221120 |
| NXPH3 | neurexophilin 3 | 11248 |
| RNF34 | ring finger protein 34 | 80196 |
| SYT2 | synaptotagmin 2 | 127833 |
| VIP | vasoactive intestinal peptide | 7432 |
| FBLN2 | fibulin 2 | 2199 |
| AMH | anti-Mullerian hormone | 268 |
| ADHFE1 | alcohol dehydrogenase iron containing 1 | 137872 |
| AGO1 | argonaute RISC component 1 | 26523 |
| AGO4 | argonaute RISC component 4 | 192670 |
| ATAD2 | ATPase family AAA domain containing 2 | 29028 |
| BBOX1 | gamma-butyrobetaine hydroxylase 1 | 8424 |
| C3orf62 | chromosome 3 open reading frame 62 | 375341 |
| CDH10 | cadherin 10 | 1008 |
| CDK6 | cyclin dependent kinase 6 | 1021 |
| COLEC12 | collectin subfamily member 12 | 81035 |
| CRYBG3 | crystallin beta-gamma domain containing 3 | 131544 |
| CTH | cystathionine gamma-lyase | 1491 |
| CTNS | cystinosin, lysosomal cystine transporter | 1497 |
| CYTH4 | cytohesin 4 | 27128 |
| DCAF1 | DDB1 and CUL4 associated factor 1 | 9730 |
| DCTN6-DT | DCTN6 divergent transcript | 118597835 |
| DDR2 | discoidin domain receptor tyrosine kinase 2 | 4921 |
| FAM118A | family with sequence similarity 118 member A | 55007 |
| FAM149A | family with sequence similarity 149 member A | 25854 |
| FLRT1 | fibronectin leucine rich transmembrane protein 1 | 23769 |

| | | |
|---|---|---|
| FN3K | fructosamine 3 kinase | 64122 |
| G2E3 | G2/M-phase specific E3 ubiquitin protein ligase | 55632 |
| GCDH | glutaryl-CoA dehydrogenase | 2639 |
| GLE1 | GLE1 RNA export mediator | 2733 |
| HILPDA | hypoxia inducible lipid droplet associated | 29923 |
| HMBOX1 | homeobox containing 1 | 79618 |
| ICAM2 | intercellular adhesion molecule 2 | 3384 |
| IPPK | inositol-pentakisphosphate 2-kinase | 64768 |
| ISYNA1 | inositol-3-phosphate synthase 1 | 51477 |
| KANSL2 | KAT8 regulatory NSL complex subunit 2 | 54934 |
| KLK7 | kallikrein related peptidase 7 | 5650 |
| KMT2E-AS1 | KMT2E antisense RNA 1 | 100216545 |
| LINC00507 | long intergenic non-protein coding RNA 507 | 100862680 |
| LINC00951 | long intergenic non-protein coding RNA 951 | 401260 |
| LMNB2 | lamin B2 | 84823 |
| LRCH3 | leucine rich repeats and calponin homology domain containing 3 | 84859 |
| MAN2B2 | mannosidase alpha class 2B member 2 | 23324 |
| MAPKAPK3 | MAPK activated protein kinase 3 | 7867 |
| METTL23 | methyltransferase 23, arginine | 124512 |
| MRI1 | methylthioribose-1-phosphate isomerase 1 | 84245 |
| MRPL44 | mitochondrial ribosomal protein L44 | 65080 |
| MUC12 | mucin 12, cell surface associated | 10071 |
| MYRIP | myosin VIIA and Rab interacting protein | 25924 |
| NAA16 | N-alpha-acetyltransferase 16, NatA auxiliary subunit | 79612 |
| NBEAL1 | neurobeachin like 1 | 65065 |
| NIPAL2 | NIPA like domain containing 2 | 79815 |
| NOTCH4 | notch receptor 4 | 4855 |
| OSBPL3 | oxysterol binding protein like 3 | 26031 |
| PAGR1 | PAXIP1 associated glutamate rich protein 1 | 79447 |
| PAK4 | p21 (RAC1) activated kinase 4 | 10298 |
| PIK3C2B | phosphatidylinositol-4-phosphate 3-kinase catalytic subunit type 2 beta | 5287 |
| PIPOX | pipecolic acid and sarcosine oxidase | 51268 |
| PRELP | proline and arginine rich end leucine rich repeat protein | 5549 |
| PTAR1 | protein prenyltransferase alpha subunit repeat containing 1 | 375743 |
| PTPRJ | protein tyrosine phosphatase receptor type J | 5795 |
| PWP2 | PWP2 small subunit processome component | 5822 |
| RAB9B | RAB9B, member RAS oncogene family | 51209 |
| RAD51D | RAD51 paralog D | 5892 |
| RAD9A | RAD9 checkpoint clamp component A | 5883 |
| RBM38 | RNA binding motif protein 38 | 55544 |
| RGP1 | RGP1 homolog, RAB6A GEF complex partner 1 | 9827 |
| SEC23IP | SEC23 interacting protein | 11196 |
| SH3D21 | SH3 domain containing 21 | 79729 |

| | | |
|---|---|---|
| SHC4 | SHC adaptor protein 4 | 399694 |
| SLC19A2 | solute carrier family 19 member 2 | 10560 |
| SOSTDC1 | sclerostin domain containing 1 | 25928 |
| SPATA7 | spermatogenesis associated 7 | 55812 |
| TAB3 | TGF-beta activated kinase 1 (MAP3K7) binding protein 3 | 257397 |
| TFDP2 | transcription factor Dp-2 | 7029 |
| THSD7A | thrombospondin type 1 domain containing 7A | 221981 |
| TNS1 | tensin 1 | 7145 |
| TRMT6 | tRNA methyltransferase 6 non-catalytic subunit | 51605 |
| TTLL1 | TTL family tubulin polyglutamylase complex subunit L1 | 25809 |
| XPO5 | exportin 5 | 57510 |
| ZFHX4 | zinc finger homeobox 4 | 79776 |
| ZNF274 | zinc finger protein 274 | 10782 |
| ZNF385B | zinc finger protein 385B | 151126 |
| ZNF436 | zinc finger protein 436 | 80818 |
| ZNF585B | zinc finger protein 585B | 92285 |
| ZNF740 | zinc finger protein 740 | 283337 |
| ZWILCH | zwilch kinetochore protein | 55055 |

| Gene category | Donor 1 | | Don |
|---|---|---|---|
| | log2fc | p.adj | log2fc |
| cell death | 1.535442365 | 1 | 1.58936768 |
| immune response | 1.535442365 | 1 | -1.41063232 |
| immune response | 1.298403168 | 1 | 1.004405179 |
| intrinsic plasticity | -1.049520136 | 1 | -1.41063232 |
| mitochondria | 1.120404866 | 1 | -1.732560415 |
| mitochondria | -1.049520136 | 1 | 2.396722602 |
| mitochondria | 2.272407959 | 1 | -1.41063232 |
| mitochondria | 1.950479864 | 1 | -1.995594821 |
| mitochondria | 1.950479864 | 1 | -1.732560415 |
| mitochondria | -1.371448231 | 1 | 1.911295775 |
| mitochondria | -2.136982977 | 1 | -1.147597914 |
| myelin/ oligodendrocytes | 1.950479864 | 1 | -1.732560415 |
| myelin/ oligodendrocytes | 1.687445458 | 1 | 1.58936768 |
| neurogenesis | 1.435906691 | 1 | 2.174330181 |
| synapse | 1.143124942 | 1 | -1.995594821 |
| synapse | 1.602556561 | 1 | 1.048799298 |
| synapse | -1.049520136 | 1 | 1.396722602 |
| synapse | -1.251153997 | 1 | -1.088704225 |
| synapse/ neurogenesis | 1.602556561 | 1 | -1.732560415 |
| synaptic plasticity | 2.272407959 | 1 | -1.995594821 |
| | -1.856875058 | 1 | 1.58936768 |
| | 1.272407959 | 1 | -1.41063232 |
| | 1.272407959 | 1 | -1.41063232 |
| | 1.535442365 | 1 | 1.174330181 |
| | 1.687445458 | 1 | -1.147597914 |
| | 2.272407959 | 1 | -1.41063232 |
| | 1.780554863 | 1 | -1.088704225 |
| | 1.21351427 | 1 | -2.41063232 |
| | 1.435906691 | 1 | -1.995594821 |
| | 1.687445458 | 1 | -1.995594821 |
| | -1.049520136 | 1 | 1.289807398 |
| | -1.634482637 | 1 | -1.41063232 |
| | 1.950479864 | 1 | -1.995594821 |
| | 1.172872285 | 1 | 1.759292681 |
| | 1.313049944 | 1 | -1.995594821 |
| | 1.365517363 | 1 | -1.41063232 |
| | 1.535442365 | 1 | -1.147597914 |
| | 1.535442365 | 1 | -1.147597914 |
| | 1.365517363 | 1 | -1.995594821 |

| | | |
|---|---|---|
| 1.435906691 | 1 | -1.41063232 |
| -1.371448231 | 1 | 1.58936768 |
| 1.272407959 | 1 | -1.217987242 |
| 2.757834786 | 1 | 1.174330181 |
| 2.120404866 | 1 | -1.995594821 |
| 1.687445458 | 1 | -2.732560415 |
| 1.435906691 | 1 | -2.41063232 |
| 1.409911483 | 1 | -1.088704225 |
| 1.435906691 | 1 | -1.732560415 |
| -1.552020476 | 1 | -1.580557322 |
| 1.298403168 | 1 | 2.174330181 |
| -2.049520136 | 1 | 1.004405179 |
| -1.693376326 | 1 | 1.396722602 |
| -1.049520136 | 1 | -1.732560415 |
| 3.272407959 | 1 | -1.41063232 |
| 1.950479864 | 1 | -1.732560415 |
| 1.120404866 | 1 | -1.41063232 |
| 1.272407959 | 1 | -1.995594821 |
| 1.365517363 | 1 | -1.633024741 |
| -1.634482637 | 1 | 1.58936768 |
| -1.049520136 | 1 | -1.303717116 |
| 2.272407959 | 1 | -1.995594821 |
| 1.028482376 | 1 | -1.303717116 |
| 1.687445458 | 1 | -1.825669819 |
| -1.371448231 | 1 | 1.174330181 |
| -1.428031759 | 1 | -1.088704225 |
| 1.757834786 | 1 | 1.174330181 |
| 1.535442365 | 1 | -1.147597914 |
| -1.049520136 | 1 | -1.41063232 |
| 1.365517363 | 1 | -2.732560415 |
| 1.365517363 | 1 | -1.995594821 |
| -1.049520136 | 1 | 1.58936768 |
| 1.757834786 | 1 | -2.870063939 |
| 1.328991487 | 1 | -1.147597914 |
| 1.065957082 | 1 | -2.111072038 |
| 1.365517363 | 1 | 1.396722602 |
| 1.120404866 | 1 | 1.174330181 |
| -1.371448231 | 1 | -2.995594821 |
| 1.365517363 | 1 | 1.58936768 |
| -1.371448231 | 1 | 1.58936768 |
| 1.535442365 | 1 | -1.995594821 |
| 1.120404866 | 1 | -1.732560415 |
| 1.535442365 | 1 | 1.911295775 |

| | | |
|---|---|---|
| 2.272407959 | 1 | -1.995594821 |
| 1.313049944 | 1 | -1.062709017 |
| -1.049520136 | 1 | -1.732560415 |
| -1.371448231 | 1 | 1.58936768 |
| 1.313049944 | 1 | 1.004405179 |
| 1.21351427 | 1 | -1.41063232 |
| -1.049520136 | 1 | -1.258629227 |
| 1.535442365 | 1 | -1.41063232 |
| 1.313049944 | 1 | 1.58936768 |
| 1.272407959 | 1 | -1.41063232 |
| 1.172872285 | 1 | -1.41063232 |
| 1.687445458 | 1 | -1.147597914 |
| 1.365517363 | 1 | -2.732560415 |
| 1.065957082 | 1 | 1.911295775 |
| 1.272407959 | 1 | 2.396722602 |
| 2.272407959 | 1 | 1.911295775 |
| 2.535442365 | 1 | 1.174330181 |
| -1.049520136 | 1 | -1.995594821 |

| ıor 2 | Donor 4 | |
|---|---|---|
| p.adj | log2fc | p.adj |
| 1 | -1.368243288 | 1 |
| 1 | -1.046315193 | 1 |
| 1 | -1.368243288 | 1 |
| 1 | -2.347484728 | 0.28371297 |
| 1 | -1.216240194 | 1 |
| 1 | -1.216240194 | 1 |
| 1 | -1.046315193 | 1 |
| 1 | -1.52436249 | 1 |
| 1 | -1.16179241 | 1 |
| 1 | -2.046315193 | 1 |
| 1 | -1.071850285 | 1 |
| 1 | -1.894312099 | 1 |
| 1 | -1.268707614 | 1 |
| 1 | 1.216719213 | 1 |
| 1 | -1.28335439 | 1 |
| 1 | -1.216240194 | 1 |
| 1 | -2.681903767 | 4.55E-09 |
| 1 | 2.368722306 | 1 |
| 1 | 1.953684807 | 1 |
| 1 | -1.331717412 | 1 |
| 1 | -1.268707614 | 1 |
| 1 | -1.557277112 | 1 |
| 1 | -1.4792746 | 1 |
| 1 | -2.046315193 | 1 |
| 1 | -1.768781217 | 1 |
| 1 | -1.046315193 | 1 |
| 1 | -2.009789317 | 1 |
| 1 | -1.046315193 | 1 |
| 1 | -1.953205789 | 1 |
| 1 | -1.631277694 | 1 |
| 1 | -1.911385613 | 1 |
| 1 | -1.768781217 | 1 |
| 1 | 2.538647308 | 1 |
| 1 | -1.145850866 | 1 |
| 1 | -1.16179241 | 1 |
| 1 | -1.4792746 | 1 |
| 1 | -1.009789317 | 1 |
| 1 | 1.105687901 | 1 |
| 1 | -1.438632616 | 1 |

| | | |
|---|---|---|
| 1 | -1.853670115 | 1 |
| 1 | 1.368722306 | 1 |
| 1 | 1.069162025 | 1 |
| 1 | -3.331717412 | 1 |
| 1 | 1.176077228 | 1 |
| 1 | -1.631277694 | 1 |
| 1 | -1.133778034 | 1 |
| 1 | -1.631277694 | 1 |
| 1 | 1.538647308 | 1 |
| 1 | -1.138237682 | 1 |
| 1 | -1.953205789 | 1 |
| 1 | 1.226703301 | 1 |
| 1 | 1.484199524 | 1 |
| 1 | 1.368722306 | 1 |
| 1 | -1.505746812 | 1 |
| 1 | -1.631277694 | 1 |
| 1 | -1.290240776 | 1 |
| 1 | 1.134257053 | 1 |
| 1 | -1.408885272 | 1 |
| 1 | 1.176077228 | 1 |
| 1 | -1.247949054 | 1 |
| 1 | 1.368722306 | 1 |
| 1 | -1.39681244 | 1 |
| 1 | -1.116704521 | 1 |
| 1 | 1.046794211 | 1 |
| 1 | -1.090709312 | 1 |
| 1 | -1.145850866 | 1 |
| 1 | -1.046315193 | 1 |
| 1 | -1.046315193 | 1 |
| 1 | -1.631277694 | 1 |
| 1 | -4.090709312 | 1 |
| 1 | -1.631277694 | 1 |
| 1 | -1.438632616 | 1 |
| 1 | -1.347484728 | 1 |
| 1 | -1.438632616 | 1 |
| 1 | -1.894312099 | 1 |
| 1 | -1.116704521 | 1 |
| 1 | -2.368243288 | 1 |
| 1 | -1.631277694 | 1 |
| 1 | -1.39681244 | 1 |
| 1 | -1.216240194 | 1 |
| 1 | -1.408885272 | 1 |
| 1 | -1.408885272 | 1 |

| | | |
|---|---|---|
| 1 | 1.953684807 | 1 |
| 1 | -1.116704521 | 1 |
| 1 | 1.134257053 | 1 |
| 1 | -1.090709312 | 1 |
| 1 | -1.39681244 | 1 |
| 1 | -1.046315193 | 1 |
| 1 | 1.368722306 | 1 |
| 1 | -1.145850866 | 1 |
| 1 | -1.046315193 | 1 |
| 1 | -1.216240194 | 1 |
| 1 | -1.116704521 | 1 |
| 1 | -1.046315193 | 1 |
| 1 | -1.090709312 | 1 |
| 1 | -1.438632616 | 1 |
| 1 | -1.461352692 | 1 |
| 1 | -1.631277694 | 1 |
| 1 | -2.331717412 | 1 |
| 1 | -1.046315193 | 1 |

**Data S7. List of unique, and overlapping significant DEGs (absolute log2FC >= 1) between i**

| Donor 1 | | | | | |
|---|---|---|---|---|---|
| Cortical layer 1 | | | Cortical layer 2/3 | | |
| iTBS | iTBS ∩ 10 Hz | 10 Hz | iTBS | iTBS ∩ 10 Hz | 10 Hz |
| NEFL | SV2B | PLAT | SPP1 | HSPA6 | MBP |
| NRGN | EGR4 | RAB31 | CCL4L2 | MT1H | HOPX |
| THY1 | SNCG | COP1 | CCL4 | CSF1 | PLP1 |
| ENC1 | PPP3R1 | DCXR | DOK6 | IRAK1 | KRT17 |
| TUBB2A | ZYG11B | GPSM2 | HSPA1A | DIRAS2 | SNCA |
| ARPP19 | RAB3A | HGF | HSPA1B | C9orf72 | LAMP5 |
| UCHL1 | TAC3 | FAM131B-AS2 | DNAJB1 | CRYAB | QDPR |
| CALM3 | CALB1 | PTCH1 | MAD2L2 | CRTAC1 | PCP4 |
| CHN1 | DGKE | EEF1AKMT2 | IL1B | PLOD2 | CLSTN2 |
| PPIA | TMEM38A | HPCAL1 | HERC4 | IL1R1 | CALB1 |
| VSNL1 | CDK5R2 | GATD1 | CHD6 | TENM1 | PCDH8 |
| MDH1 | FAM98A | PDGFRA | GFAP | DAAM2 | RAPGEF4 |
| CLSTN1 | C2orf80 | EMP1 | TFG | MAG | NPAS4 |
| BEX1 | PDK3 | TNRC6C | GLRX | DIS3L | MET |
| TUBA1B | PCMT1 | SHISA6 | GPSM1 | RIPK2 | CUX2 |
| CRYM | SH2D5 | BLMH | TMEM201 | ARHGAP4 | LINC00507 |
| YWHAH | NAP1L2 | FTX | SHISAL1 | NUDT18 | NTNG2 |
| PLD3 | PVALB | TPD52L1 | TRBC2 | GPR180 | TESPA1 |
| RGS4 | CPNE9 | WIF1 | NADK | LRRC28 | LINC01007 |
| YWHAZ | C1orf115 | SELENOS | TOMM70 | BET1 | MAP2K1 |
| SLC17A7 | ABLIM2 | ATP2B4 | MARK1 | TMCC3 | RXFP1 |
| RTN4 | RELN | USP10 | AFMID | SALL2 | VSTM2A |
| SCN3B | ACTR1B | ARID4B | CDR2L | CNDP1 | HBA2 |
| FAM131A | CPNE6 | LYST | RILPL1 | HELQ | MGP |
| MAP1B | CDIP1 | DVL1 | MED4 | B3GALT2 | RASGRF2 |
| PHYHIP | DLGAP1-AS4 | MIDEAS | MED12L | MFAP3 | CALB2 |
| RTN1 | FREM3 | ACSL3 | DNAJC9 | SYNC | SOWAHA |
| CREG2 | RTN4RL2 | ITPKB | CSNK1E | TFEB | COL5A2 |
| GAP43 | EPB41L3 | PHC1 | RTN4R | BOK | ITPKA |
| GDI1 | ZEB1 | SLC20A1-DT | ITGA6 | LPP | MOBP |
| TMEM59L | YOD1 | TFDP2 | SPRN | FSTL5 | TF |
| NSG2 | MFSD8 | PKD2 | TRAPPC3 | COG8 | TAFA2 |
| APP | DDX19A | POLR1C | UTRN | SGMS2 | RCN1 |
| PAK1 | MRPL17 | ASF1A | DENND6B | PRIMPOL | IPCEF1 |
| LINGO1 | SLC25A27 | GTPBP10 | VRK1 | CHN2 | NECAB2 |
| NELL2 | CCDC85A | ERICH1 | THAP11 | RBBP8 | EFHD2 |
| SYN2 | CERS6 | PRAG1 | DROSHA | CCDC80 | LZTS1 |
| ATP1B1 | STRIP2 | ZNF462 | PEX10 | ERMP1 | HS3ST4 |

| | | | | | |
|---|---|---|---|---|---|
| PRKAR1B | USP7 | LINC00595 | CBARP | ELF1 | RNASE1 |
| AP2M1 | TAFA1 | FNTB | NCLN | POLR1A | CBLN4 |
| STXBP1 | MRPS23 | VIPAS39 | CLOCK | RBM15 | PPP1R14A |
| RAB6B | TSPAN17 | KCNK10 | DOT1L | FEZF2 | DGKB |
| STMN2 | PPME1 | RFLNB | EIF3C | HHIP | GALNT18 |
| LMO4 | BCAP29 | SIRT7 | SLC2A11 | GBP1 | CNTN2 |
| GABBR2 | PTGS2 | ZNF426 | MKRN2 | POT1 | SEMA3E |
| TESPA1 | TMEM132D | LRFN1 | DCAF5 | TMEM260 | SMYD2 |
| PPP3CB | HR | TFIP11-DT | ACBD6 | PAWR | DTNBP1 |
| GNG3 | TMEM240 | LASP1 | UQCC3 | STAC2 | RPRM |
| CXCL14 | CLIP2 | PPP1R12C | MLH3 | CDC6 | IGFBP5 |
| TAGLN3 | PNMA6A | DNAJC3 | DYRK3 | FABP6 | TMEFF2 |
| ALDOA | SLC2A8 | FOXN3 | SLCO2B1 | GJB1 | SPON2 |
| MEF2C | GALNT18 | MRPL22 | PXDC1 | RCL1 | CPNE4 |
| TUBA1A | PRMT5 | NRN1 | MAPKAPK2 | HPS5 | FAM107B |
| PRKCG | CIART | SNRPB | FLNA | HBA1 | TMEM178B |
| PEG3 | GPRASP1 | FGF13 | KCNA3 | CA7 | HS6ST3 |
| HSPA12A | FABP3 | TBL1XR1 | CHIC1 | ITFG2 | GRIA4 |
| ATL1 | GABRD | ITGB1 | ANKHD1 | HS3ST2 | CD52 |
| SHISA4 | NEUROD2 | BRAF | SLC39A12 | CBLN1 | FLRT2 |
| KIAA1549L | GAK | TRIB2 | SCFD2 | SLC14A1 | ADCYAP1 |
| SLC26A4-AS1 | TMEM178B | C8orf76 | FAM98B | ZNRF2 | BTBD3 |
| TPBG | BTBD3 | GPATCH8 | DPP9 | LINC00499 | KLK6 |
| CDC42 | ISOC2 | MAFK | CSTF1 | CA13 | ERMN |
| HOPX | STMP1 | NSMCE1 | UBXN2A | PLD4 | FREM3 |
| BAIAP2 | DUSP2 | ATL2 | TRIP6 | ANGPT2 | TMEM144 |
| PI4KA | ZNF385B | ADAM10 | ADORA2B | CMTM5 | PRAG1 |
| ATP1A1 | TBC1D23 | ZNF503 | H4C3 | ZNF561-AS1 | NTM |
| STMN4 | THRB | PWP1 | COL9A2 | SOX6 | NTRK3 |
| PTPRN | USP6NL | TRIM23 | KBTBD4 | AMH | FBLN7 |
| SNCB | TYRO3 | SOX7 | ERBIN | ZNF471 | ABTB3 |
| ATP2A2 | NFKBIA | GTF3C4 | MMGT1 | SNHG28 | HS6ST1 |
| GFAP | ELP1 | LYPD1 | RELT | NOC3L | DAPK1 |
| LINC01007 | SLC25A39 | ARHGAP12 | MMP15 | NKX6-2 | PDZD2 |
| NUDT4 | CSE1L | BIRC3 | MZF1 | RBBP9 | ENPP2 |
| ENSA | CADPS2 | REXO2 | IFI44 | ISYNA1 | NPY |
| SYN1 | AUTS2 | SLC5A3 | ZCCHC7 | NKX3-1 | MYRF |
| APBB1 | LENG8 | ATF7IP2 | PCYT2 | PARP11 | NR4A2 |
| CAP2 | FIG4 | WFDC2 | NOPCHAP1 | DERA | CDH22 |
| STX1A | MSR1 | CNP | ARMH3 | TRIM22 | CDK18 |
| ITPKA | PAFAH1B2 | HRAS | TOPBP1 | DYSF | CXCL14 |
| CA10 | KCNK12 | CMC2 | NFKBID | CXCL3 | MAPK4 |
| PRRT1 | NPM2 | SGSM2 | ZCCHC14 | ARHGEF37 | NEUROD1 |

| | | | | | |
|---|---|---|---|---|---|
| NECAB1 | CELF2 | TPM3 | MRPL23 | TPPP3 | B3GAT2 |
| GRSF1 | SRGAP3 | MATK | CNPPD1 | TRAF5 | BRINP2 |
| PPP3CA | TMEM151A | NDUFA10 | EML6 | SPG11 | TSPAN14 |
| LNX1 | TMBIM4 | PCDH8 | NCKIPSD | MOG | PPP4R4 |
| GRIA3 | CDC123 | CHGB | ID3 | RELA-DT | CLDND1 |
| LINC00507 | CHRM3 | EHMT2 | ECSIT | NRG1 | KIF1C |
| BSCL2 | SPRED2 | KCNT1 | PRELP | LINC03048 | BCAS1 |
| GGCT | DDRGK1 | MGP | OBSCN | | LINC02762 |
| POLR2H | STRIP1 | MRTFA | MCF2L2 | | AKAP5 |
| MLLT11 | LINC00294 | DDX3Y | COL6A1 | | CD24 |
| ATP2B2 | EXTL1 | FOSL2 | UBXN1 | | TCAP |
| LINC00632 | ZNF629 | DTNA | TDRKH | | BEGAIN |
| NEFM | SMPD4 | PINK1 | KNOP1 | | CDKN1C |
| SHISA9 | CREB3 | CAPZA2 | THOC5 | | KCNG1 |
| LDOC1 | CDH8 | MPZL1 | ZNF318 | | VAT1L |
| TUBA4A | TAFA2 | BTBD2 | ELMO2 | | CARTPT |
| PPA1 | WDR7 | VPS26C | MSX1 | | PIP5K1B |
| KCNK1 | MAPKAPK5-AS | PGM1 | CASP9 | | FAM13A |
| MYL12B | TMUB2 | TSPAN31 | ALG3 | | VIP |
| PPM1L | ORC4 | KMT2B | BCL11B | | TKFC |
| CD47 | RNF141 | TNFRSF1A | C12orf43 | | MAN1A1 |
| PRRT2 | SHISAL1 | BZW2 | ATL3 | | STRIP2 |
| MET | SMIM10L1 | SLC29A4 | UBASH3B | | GLRA3 |
| HTR2A | SEC23B | NRDC | KCNAB1 | | FAT3 |
| ARPC5L | TMEM65 | SMYD2 | SMG1P5 | | ASS1 |
| FBXL2 | GSTM4 | CAMK4 | PHAF1 | | ICAM1 |
| USP14 | TET3 | MAPK8IP2 | TBC1D16 | | GPR26 |
| CLSTN3 | NEUROD6 | ZDBF2 | CELF6 | | HRK |
| ZBTB18 | WAC-AS1 | CHMP5 | LRP1B | | RAB3B |
| VIM | CCKBR | EP300 | CRTC3 | | KIRREL3 |
| NCKAP1 | DNAJC24 | MTA3 | MRPL36 | | SULF2 |
| MCHR2 | WSCD2 | SERPINB6 | ZNF268 | | ARL4A |
| SNCA | NOVA2 | PPT2 | ALDH1A3 | | GATB |
| NPDC1 | CHPF | URGCP | ZNF436 | | DHCR7 |
| ARPP21 | NLGN4X | PPP1R16A | SEC23IP | | ETV1 |
| KALRN | FEM1B | DICER1 | CHST10 | | GNAL |
| NRXN2 | CHRM1 | ANKRD13B | KIF9 | | IGFBP2 |
| DIRAS1 | KCND2 | RSPH1 | WDPCP | | OPALIN |
| CCDC167 | H2AC6 | RPS6KA3 | KCTD8 | | ITGA9 |
| CYP46A1 | CLASRP | PRDM16-DT | ADAMTS13 | | PDLIM7 |
| EGR1 | QARS1 | FAM131C | TSR1 | | NOL4 |
| PTPRF | CAPN10 | ZNF697 | MRPS30-DT | | MTUS2 |
| FRRS1L | MOCS2 | ANKRD35 | POLR3E | | THSD7A |

| | | | | |
|---|---|---|---|---|
| TOLLIP | SSH2 | GMCL1 | CXCL16 | ZBTB25 |
| ST6GALNAC5 | TBC1D16 | ARL6IP6 | TMOD3 | GRIN3A |
| NGEF | DMAC1 | PLEKHM3 | GTF2IRD1 | RRAD |
| GABRG2 | ACBD3 | NPPC | ZNF777 | UGT8 |
| AGPAT3 | OSBP2 | DGKD | NFYB | MFSD14B |
| RBFOX1 | MPP1 | SLC6A11 | NRG3 | NECTIN3 |
| BEX2 | MPHOSPH9 | HYAL2 | MPHOSPH8 | LRRC4 |
| S100B | CBLN4 | CBLB | PPWD1 | PITPNM2 |
| HMGCS1 | PNMA8B | ALDH1L1 | PAOX | KCNK3 |
| C2CD2L | DOLK | ZNF639 | MTMR14 | WFDC2 |
| ITFG1 | GON7 | LPP | MRPL10 | NLGN3 |
| CCL2 | TAFAZZIN | LCORL | GPR153 | TRERF1 |
| NFE2L1 | KAT7 | CMYA5 | NRP2 | NXPH3 |
| SPP1 | TUSC3 | SLC25A48 | OXTR | CNTN4 |
| NAPA | KCTD13 | IBTK | SPAG16 | SLC6A15 |
| CX3CL1 | AARSD1 | TMEM74 | DGKE | GNA15 |
| NRXN1 | ACTR6 | MED30 | GFRA1 | MAN1C1 |
| PSIP1 | TMEM8B | HAUS6 | DACH1 | GRB10 |
| CRYAB | NUTF2 | MLLT10 | ZNF628 | PENK |
| RUFY3 | NAPG | KIAA1217 | PRKAR2A | NAA60 |
| CADM3 | ZNRF1 | LINC00839 | GRIK3 | CRLF1 |
| VGF | NELL1 | NUTM2B-AS1 | WDR55 | SOHLH1 |
| WIPF3 | HSPA4 | CHUK | PRR5 | NFKB2 |
| CYP4X1 | PLK2 | CASP7 | UTP14A | ELOVL1 |
| RTN4RL1 | PIAS1 | AFAP1L2 | SLC22A15 | CPNE7 |
| TLE1 | CA4 | EDRF1 | RAD17 | TMEM65 |
| SYT13 | MAN1C1 | AMPD3 | RASL11A | CROT |
| TENM2 | FAM217B | PARVA | TAF1 | PYDC1 |
| KCNH1 | CS | SAAL1 | ZNF654 | FAM153A |
| GALNT8 | CALU | HPS5 | KCTD21 | MOB1B |
| CDK20 | EFR3B | FRMD8 | ARMC8 | KCTD3 |
| PNMA8C | SSX2IP | CTSC | PLEK | SNAPC2 |
| CDH18 | SEMA4C | GRIK4 | TRAM1 | ATF7IP2 |
| ERCC5 | AHNAK2 | RASSF8-AS1 | BBX | THEMIS |
| ATP6V1G2 | PCDH11X | ITGA7 | LINC02245 | MSANTD4 |
| FKBP1A | TTLL7 | MSI1 | IL17RA | RASGRP1 |
| SIRT3 | PRMT8 | SETD1B | ACVR1C | SLN |
| ARHGEF25 | SLC4A8 | DNAAF2 | TBC1D22B | ST6GAL2 |
| TUBB3 | SLITRK5 | PRKCH | LINC01089 | TMEM88B |
| SERPINF1 | TMEM42 | RLBP1 | FPR1 | C8orf90 |
| LRFN2 | FOXRED1 | TARS3 | LIFR | ISLR |
| DOCK3 | PTPRR | NEURL4 | KDM2B | MRPL47 |
| GOT1 | MRPL55 | SPAG5 | TBL2 | CASTOR1 |

| | | | | |
|---|---|---|---|---|
| SLC30A3 | HTR5A-AS1 | EMILIN2 | PRKAG2 | C1QL2 |
| GUCY1B1 | SYNJ2BP | DUS3L | PPID | FANCL |
| ELMO1 | FMNL1 | C5AR1 | RPS6KC1 | ASGR1 |
| TSR2 | RBM24 | MYT1 | SULT1C4 | PCDHGC3 |
| SH3BP5 | MRPS18B | TXLNGY | MAP3K20 | CRHR1 |
| EFNB3 | LYRM1 | SLC38A2 | SNX18 | TMEM63A |
| TGFBR3L | NHSL2 | PON2 | ZNF250 | SSTR2 |
| CCDC184 | GGNBP2 | REEP2 | SSTR3 | TOX3 |
| SCN2A | NPEPL1 | ARL4C | COQ5 | CA4 |
| DNAJB1 | EMC10 | PDS5A | CNFN | CLDN11 |
| SYNGR1 | ENTREP2 | CD74 | SNRNP27 | MKNK1 |
| DCP2 | TOMM40L | NSD2 | TNFRSF1B | SLC17A6 |
| B4GALT4 | MTMR3 | FRG1 | KDM3B | CACNA1C |
| COMMD1 | FASTKD1 | ODC1 | YES1 | CACNA2D3 |
| C11orf87 | NUDT6 | PNMA1 | C1orf54 | PYGO1 |
| HMGCR | UIMC1 | DYNLT2B | TPT1-AS1 | ARHGAP10 |
| ATP6V1D | ZCCHC7 | GABRA5 | NPR2 | ANLN |
| GRM5 | EDC4 | PUF60 | WWOX | SLC24A4 |
| DNM1L | RAB8A | TRIB1 | EHD1 | IGFBP4 |
| CDK5R1 | ZNF146 | NBEAL1 | ERGIC2 | LINC02607 |
| MICOS13 | CRNKL1 | RASA1 | SLC35F6 | CAPN3 |
| VXN | DUSP26 | MRPL24 | PSMA3-AS1 | SLC2A8 |
| AP3M2 | BRPF3 | PHC2 | C6orf120 | PTGR3 |
| PTPRO | ZNF134 | CBX5 | DCP1A | HMGB3 |
| RAP1GAP | CXCL8 | OPTN | GTPBP4 | SESN1 |
| DLGAP1 | TMX4 | AGO3 | SH3YL1 | DAB1 |
| RALYL | PRKN | GPATCH4 | MMAB | HAPLN2 |
| BRI3BP | RBM12B | EPB41L5 | LINC00844 | HSDL1 |
| NPM3 | LRTM2 | LZTFL1 | BAZ1B | CDH4 |
| NDUFAF3 | IKBKG | PDE4D | CACNA1I | SERTM1 |
| RGS7 | NCOR2 | THSD7A | TLK2 | RSPRY1 |
| GABRB2 | C1orf216 | AHCYL2 | RMND5B | PCGF1 |
| ARMCX2 | LYSMD2 | BICD2 | KAT6A | EP300 |
| CDC16 | PCDH19 | CRLF1 | SLC25A16 | INO80D |
| NAV1 | COA1 | PIP5K1A | THAP9-AS1 | GULP1 |
| SNAPC3 | GALNT1 | PDCD7 | RND3 | MPPED2 |
| GABRA1 | CRIM1-DT | CDC42EP4 | PIK3R4 | CCNO |
| TAC1 | PAQR3 | ZMYM2 | MFSD1 | ZFP64 |
| CACNA2D1 | RIOK3 | PCSK2 | IFRD2 | CPNE8 |
| CYRIA | LASP1NB | LONRF2 | PYROXD1 | HRH2 |
| FRMPD2 | SLC25A26 | EMC1 | YJU2B | DIAPH1 |
| CPEB1 | GPRIN1 | H3-3A | FAM120C | ARL14EP |
| BOLA2B | C1QBP | EMX2OS | PIK3C2B | SLC37A1 |

| | | | | |
|---|---|---|---|---|
| AP2S1 | SNAP47 | USPL1 | PDGFRB | PRKCQ-AS1 |
| CDKN2AIPNL | FNIP2 | RASA3 | LINC01503 | IRF2BP1 |
| PSMD3 | MAPK14 | IGSF3 | CLN3 | FNBP1L |
| ACTR3B | GPAT3 | FCGR1A | PKNOX1 | NR2F2 |
| B4GALNT1 | CDH12 | CTSS | PWP2 | GPCPD1 |
| AK5 | MAK16 | C1QL2 | INTS12 | SOCS5 |
| SLBP | PDZD7 | CCNT2 | JDP2 | C8orf76 |
| LUC7L | B3GNT4 | CC2D2A | USP39 | COQ10A |
| NUMBL | MRGBP | FRYL | MALT1 | SLIT2 |
| VDAC3 | ICOSLG | ELOVL6 | GALNT1 | COMMD2 |
| KIF3A | MDM4 | HUS1 | ADM | VLDLR |
| NRXN3 | CBLN2 | ZNF680 | COG4 | MPP7 |
| NEGR1 | LRP8 | AP4M1 | PDE12 | PLAC9 |
| ADGRB1 | MIGA1 | ARHGEF10 | MTX3 | HTR1A |
| PIK3R1 | RASSF5 | ZBTB5 | UQCC6 | TENM4 |
| PIH1D1 | FHL2 | ZNF248 | PLCXD3 | CNIH3 |
| PSMB3 | C3orf80 | CCNT1 | TRAF2 | ADGRF5 |
| USP46 | DYRK2 | HMG20A | RPAP1 | KIT |
| MAPRE3 | GSS | LINC01563 | S100A10 | TPBG |
| CACNG3 | SCN1A | NETO1-DT | ADAM10 | ERLIN1 |
| GNG2 | ZNF410 | ZNF211 | ABCG2 | GRM7 |
| CNKSR2 | SCAMP1 | DGCR8 | CCNB1 | DHRS4L2 |
| PNCK | AMFR | TRIM44 | SERAC1 | HSPA2 |
| ZDHHC8 | PRXL2B | FGFR3 | GDPD5 | JAG1 |
| DHCR24 | PHTF1 | NOL9 | DNAH10OS | EBF4 |
| CDH2 | ETV1 | SDC3 | ZNF766 | GPR37 |
| FAM131B | ABCB9 | CELF4 | CAPN10 | CNTNAP5 |
| NCALD | MTA1 | DYNLT3 | TNIP1 | KMT2B |
| RASGRF1 | PDCD6IP | AHDC1 | ZNF227 | FOXP2 |
| PRSS3 | LARP4B | WDR12 | LAMA4 | CRABP2 |
| CAB39 | DYRK3 | SRPRB | TMEM161B | NIPSNAP3A |
| OXR1 | MCM6 | ADCYAP1R1 | PDIK1L | SDR16C5 |
| NDRG3 | AKAP1 | MROH1 | ITGA2 | NPPA |
| KIAA0232 | MRPL19 | FKBP2 | TMEM54 | TMEM132D |
| SIVA1 | BAIAP2-DT | DISP2 | TMEM19 | RBMXL1 |
| AZI2 | MTMR6 | PTPRE | GGT7 | RFX4 |
| FBXW2 | HTRA2 | BCR | STOX1 | SGMS1 |
| ARSA | LINC01896 | OLR1 | BRD1 | PLEKHH1 |
| PPFIA2 | BTBD1 | COPS3 | NSMCE4A | SLC8B1 |
| SYT4 | BRD7 | SELENOF | PTPN12 | WASH2P |
| NDFIP2 | ANKRD40 | BCAN | MFSD3 | CTXN3 |
| ZNF597 | TRIM46 | DBNDD2 | FAM89A | SHISA8 |
| DHRS11 | EXOC6B | NRIP1 | STX17 | TRNT1 |

| | | | | |
|---|---|---|---|---|
| STS | COL5A2 | GFRA2 | PIERCE1 | PDE7B |
| PART1 | FAM151B-DT | ACVR1B | CEP164 | KCTD4 |
| GATB | CSMD1 | NUBP2 | PHF11 | SERPINH1 |
| SRRM4 | ADCYAP1 | SLC4A1AP | MED11 | KCNJ16 |
| ATP6AP1 | PDE1B | PYGB | CCL8 | PBX2 |
| SST | KCNB1 | MAP6 | INTS2 | MTRES1 |
| NISCH | CNTNAP4 | RAPGEF4 | AFF1 | KRBOX5 |
| MAPK9 | CCDC24 | ABCF2 | MUL1 | LAYN |
| ANO3 | LTA4H | RDH10 | IPMK | FNDC9 |
| DDX10 | MSMO1 | PSMD2 | CHCHD7 | MYBPHL |
| SPON2 | NAPEPLD | SLIRP | ODF2 | DHRS4-AS1 |
| DAPK1 | RCC2 | RHOC | RASSF1 | TNFSF9 |
| CORO2B | AGL | LHFPL3 | RPP25 | MTARC1 |
| ZBTB4 | LINC00882 | SLC24A3 | IL1A | SP4 |
| COLGALT2 | EEFSEC | PARP1 | NKAIN1 | SERPINE1 |
| ICA1L | NUP155 | SMOC1 | TNFAIP3 | NINJ2 |
| STAM | FCHO2 | NKAIN4 | PGAP2 | CDH20 |
| NOMO1 | TYW1 | COPS2 | PTPN6 | IRF7 |
| GLRA3 | RUNDC3B | IFI16 | TRPV2 | TBC1D10A |
| NDUFA8 | AKNA | GPD2 | ZNF844 | TCERG1L |
| IARS1 | EIF5AL1 | NECTIN3 | ZNF793 | WNT3 |
| DOCK4 | TVP23A | GNB4 | ZNF587 | PGPEP1 |
| CD200 | SRR | ST6GAL1 | A1BG-AS1 | HGF |
| MSX1 | CA5BP1 | MAN2B2 | RNF128 | ADARB2 |
| RFTN1 | SLC17A6 | SNX25 | GYG1 | ANKRD35 |
| DNAJC19 | SLC27A4 | VCAN | ABCB1 | REXO4 |
| MBD1 | MCHR1 | HGSNAT | PPP4R1 | ADAMTS8 |
| PGP | CDC42BPB | ENDOG | AIF1L | POLR3B |
| AP2A1 | H2AJ | GCN1 | OSBP2 | ZSCAN31 |
| TMEM132A | UBE2T | SPTLC2 | EEFSEC | GRIK1 |
| NTRK3 | ACVR1C | IFI27L1 | C1QTNF3 | RPP30 |
| GPRC5B | RNF25 | NAA60 | SLC36A1 | RAD54L2 |
| LRPPRC | COPS7B | USP36 | PHYKPL | KCNS1 |
| ATRNL1 | ARL6 | CD320 | RRP8 | SFI1 |
| FBXO44 | ZDHHC22 | SLC25A29 | CRISPLD2 | PALLD |
| PPP4R4 | GPR137 | IQSEC1 | MTHFR | WDR25 |
| PNMA8A | VPS13D | KLHL18 | PTPN18 | DEPDC5 |
| AP3S1 | CFAP97 | IRF1 | CCDC14 | GPR19 |
| ICAM5 | TRIQK | DLAT | MCHR2 | GAB2 |
| KLHDC10 | RNF112 | ARCN1 | RTN4IP1 | TYW1B |
| IFIT3 | GPR61 | AP1G1 | PDIA4 | HMGN5 |
| CGREF1 | PRKAA1 | NETO1 | DCAF12 | FAM43B |
| GSK3A | TENT2 | SELENOP | SETD7 | CNOT10 |

| | | | | |
|---|---|---|---|---|
| MRPS34 | ZNF282 | MANSC1 | PHF13 | C1R |
| TSPAN5 | SLF2 | TOMM40 | WDR75 | SOCS2 |
| MTCH2 | FGF9 | KAT8 | TMEM87A | STEEP1 |
| SERINC3 | FCF1 | BOD1L1 | PEX26 | REEP3 |
| CISD1 | CDK5RAP1 | TMEM167B | KLHL29 | TYMSOS |
| BRINP1 | SLC35A2 | RAB12 | PELO | SLC25A1 |
| SATB2 | APOOL | PRPS1 | PUS1 | CNN3-DT |
| ZNF800 | USP5 | ELAVL1 | TMEM163 | TRABD2A |
| RPGRIP1L | CDK11B | CHPT1 | TRPM3 | SLC5A11 |
| EIPR1 | MUL1 | PSMD6 | FSTL1 | RRP15 |
| UQCC5 | C1orf53 | SF3B5 | ZDHHC2 | TRPS1 |
| MZT1 | RASL11B | TMEM33 | ANXA1 | TFPI2 |
| NLK | ZKSCAN8 | KCND3 | SYNRG | TMEM125 |
| KIF21B | RSPH3 | FBXO41 | ID1 | H2BC21 |
| LMBRD2 | TRMT12 | MEGF9 | MIR222HG | CLN6 |
| CYTH2 | ALG9 | FAM200B | SLC25A38 | CABYR |
| IGSF21 | DPAGT1 | EGR3 | FCGR3A | ZNF184 |
| KCNV1 | FAR2 | CCSAP | ATP10B | LAMB1 |
| MFSD6 | ST20 | GPBP1 | BAALC-AS1 | SHROOM3 |
| SPNS2 | RAB3D | TRNP1 | APLNR | PXYLP1 |
| LSM4 | RSPH14 | MXI1 | WDR81 | EDEM1 |
| GADD45G | LINC02245 | ETFB | PTPRT-DT | SELE |
| PTK2B | LSP1P5 | ARRB1 | NSL1 | ASIC4 |
| CDK14 | HDAC3 | SPTAN1 | DAB2 | RNF216P1 |
| SEC23A | PPP1R10 | ZCCHC14 | OSTF1 | CCDC30 |
| CORT | CHST10 | RBM17 | STK4 | FN3K |
| SGIP1 | ANKS1A | SNHG15 | GON4L | PRDM16-DT |
| ACTN2 | SPIDR | GLRX3 | SOCS4 | SNAI3-AS1 |
| UBQLN1 | ACADSB | YAF2 | PARN | RNF25 |
| HS6ST3 | PLA2G6 | TMEM144 | CDC42EP1 | ZNF496-DT |
| KDM5B | TMEM150C | CYB5D2 | PIK3R3 | RGR |
| JCAD | NSFL1C | CREG1 | UBXN8 | NCAPD3 |
| UQCC4 | PI4K2A | GCA | MSI1 | PLCB4 |
| CLVS2 | MED4 | CMTR1 | TDRD9 | MYO1B |
| YIPF1 | GAS6 | CPSF6 | MOSPD2 | ZBTB10 |
| FBXO25 | FBXO33 | MED13 | ZC4H2 | RBM4B |
| LY6E-DT | IK | NFIA | B3GALNT2 | ANXA4 |
| HINFP | DMXL2 | MMS19 | VEGFA | CNTN5 |
| KCNH5 | CDH9 | ITGAX | CDKL2 | ABHD12B |
| EZH1 | MECR | ALDH3A2 | FZD8 | NAGLU |
| DYNC1I1 | TMEM170B | FHOD3 | NEIL1 | TGFBR2 |
| UBXN2B | MIR22HG | N4BP2L1 | PWWP3B | SALL3 |
| SPRN | SEMA4F | SH3GL2 | EIF2AK4 | EPHA6 |

| | | | | |
|---|---|---|---|---|
| VPS39 | IFIT5 | CCDC106 | TNNT2 | TRABD-AS1 |
| TMEM63B | ATAD1 | BPNT2 | HLA-L | JAKMIP3 |
| MARK2 | FGFR1OP2 | DLG2 | LINC02668 | MREG |
| TNFAIP1 | KIAA0319 | SPOCK1 | DHX35 | RHBDD3 |
| GOLGA8A | TOR1A | SETD5 | SLAIN2 | MRAP2 |
| ATP6V1H | ANKRD13C | TMEM129 | TVP23A | ENTPD3 |
| LINC01106 | OTUD7A | CCDC186 | MTR | TGFB1 |
| DCTN1 | TLK1 | DNAJC4 | PMS2P4 | NIBAN2 |
| SEPTIN11 | GABRA4 | MCM3AP | PEX1 | TUBGCP4 |
| MKRN2 | SLC22A23 | ASMTL | CARNMT1 | KLF12 |
| KATNAL1 | ERICH3 | CASK | VWC2 | HLA-DQB1 |
| DNAL4 | BDH1 | FKBP3 | FRMPD2 | DGKA |
| PPP2R5D | RAB11FIP2 | MFSD2A | ST8SIA1 | MYOM2 |
| PTPA | TMEM229B | ZNF326 | NBPF1 | TIMP1 |
| AMACR | PAQR6 | ARV1 | CBLL1-AS1 | RHBDL3 |
| FHL3 | LINC00662 | TBCE | ARHGAP22 | CD22 |
| ADGRF5 | TENM3 | CNTNAP5 | OLMALINC | KIAA1755 |
| LCNL1 | FNDC9 | GALNT13 | SETDB2 | TSPAN6 |
| BRICD5 | VPS52 | SLC25A38 | MSRB1 | NGB |
| INTS2 | ABRACL | CWC27 | IQCK | ZRSR2 |
| HOMER1 | SNX30 | CREBRF | ZNF23 | PIR |
| ZBTB8OS | KCTD10 | RMND5B | FTSJ3 | GREM2 |
| FDPS | EXD2 | CEP43 | HIF3A | USP21 |
| TTC1 | SMARCE1 | STEAP2 | ZNF446 | FOXP4 |
| UBFD1 | GTF2F2 | ITPRIP | DACT1 | FAM210B |
| ANXA6 | BCL7C | GPAM | PATZ1 | RAB20 |
| JADE2 | CRB1 | CHST11 | PLRG1 | NUDT15 |
| SPATS2 | DIS3L2 | CTDSPL2 | CCDC32 | ZUP1 |
| KITLG | SETMAR | KCTD5 | ASIC2 | TP53RK |
| RASAL1 | SOWAHB | TMX3 | HSPB6 | FAM124A |
| ARFRP1 | KIAA1586 | MIER2 | BBS5 | DLX6-AS1 |
| CALN1 | CBLL1-AS1 | FEM1A | HLA-DPB1 | MAP3K14 |
| SLC2A6 | TBXAS1 | LMF2 | PHTF2 | MKX |
| SNX32 | TCP11L1 | SYNGAP1 | GPR83 | PPIP5K1 |
| ESYT1 | TUT1 | ESYT2 | HDAC7 | INPP5J |
| CDADC1 | P2RX2 | EFHD2 | DCAF15 | LRRC2 |
| NANOS3 | AVEN | NXF1 | LENG1 | WIPF1 |
| MICU3 | SQOR | VEGFB | ZBED1 | PLEKHH3 |
| UBE2E2 | TPPP3 | NDEL1 | DENND11 | KRI1 |
| RPP30 | ELMO2 | MFSD12 | CAT | SLC35E4 |
| TCEA1 | SUPT6H | BLTP1 | EPS8 | FUOM |
| TIMM17A | RPF2 | MANF | RNPEPL1 | TMEM97 |
| CDK5 | PPIL2 | FAM229B | LRMDA | CXXC4 |

| | | | | |
|---|---|---|---|---|
| SACM1L | MOB1B | SLC18B1 | PLAAT4 | NUP214 |
| BTRC | LRRC4 | TECPR2 | PPL | NR3C2 |
| GRIN2A | SIAE | EPS15L1 | LMLN | POLR1C |
| NECAP1 | ERGIC2 | EHBP1 | PPARGC1B | POMK |
| LRFN5 | RNF19A | PAPSS1 | MANEA | CMC4 |
| KCTD16 | VIPR1 | TEX264 | CHRAC1 | WDR36 |
| SYT7 | LARS2 | EMC6 | SHC4 | GALNT10 |
| IGIP | BBX | POLR2G | FSTL3 | GALNT14 |
| ZSWIM8 | ST3GAL4 | KCNQ3 | PLCG1 | HIF1AN |
| ERVK3-1 | PUS1 | SPRYD7 | RNF182 | CENPC |
| RBP4 | AKAP13 | HSPD1 | RIOX2 | C2CD3 |
| CHTOP | FCHO1 | LAMP5 | GUCY1A1 | PRORP |
| PDP1 | AMIGO1 | ADNP | ARFIP1 | EID2B |
| FLOT2 | IL6 | CHP1 | PCDH1 | MPHOSPH9 |
| ST8SIA5 | STT3B | C19orf53 | PPP3CC | AGO4 |
| IPO5 | TNKS | MAGI2 | TCTN1 | TMEM41A |
| DTNBP1 | PPFIA4 | ZCRB1 | TBC1D4 | ASXL2 |
| GRIA4 | SECISBP2 | VSIR | GEMIN2 | RELN |
| OAZ2 | SLC35B1 | YIPF6 | CSNK1G1 | EML1 |
| GAS2L1 | ALKBH6 | SIK1 | TSSC4 | DPAGT1 |
| ACOT13 | DSTYK | MALSU1 | RIN1 | KCTD15 |
| PNMA3 | EPHX4 | UBAP2 | THBD | PTGER4 |
| GLG1 | MECP2 | MMP17 | DOK5 | BBOF1 |
| SCAP | TAF5L | SKIC8 | RFTN2 | KLHL17 |
| CYRIB | VPS13B-DT | COG4 | ORC3 | PTPRD-AS1 |
| DOC2A | PEX14 | CDYL2 | THEM6 | FA2H |
| MADD | LYPLAL1 | PHF6 | SNAP23 | GLDN |
| PLP1 | MYO1B | PPP2R5C | PLA2G4B | UNC5C |
| AASDHPPT | METTL21A | PPP1R14A | JADE1 | FZD6 |
| HIVEP2 | GUF1 | RCAN1 | UFSP2 | GXYLT1 |
| HRH2 | OSBPL3 | TCEAL1 | UGGT2 | TMEM235 |
| PRKDC | FAM135B | TRIP12 | TTC17 | PPM1J |
| L1CAM | MLLT3 | CIT | PLXDC1 | PCDHB4 |
| SLC30A9 | PTPN3 | SMIM10L2A | THEM4 | FAM47E |
| ME3 | GRID1 | PRR36 | METTL25B | MGST2 |
| PPA2 | CCDC34 | CAMTA2 | SYT14 | RAVER2 |
| TOMM34 | FAM76B | ZFP36L1 | VPS54 | CDKL3 |
| PAICS | DYRK4 | LUC7L3 | FAM114A2 | NFYA |
| THOC7 | NFATC3 | ENY2 | LY86 | IL11 |
| SLC12A8 | ZNF574 | YKT6 | NHSL1 | PPP2R1B |
| ARFGEF3 | RRP7BP | EI24 | NUP43 | IGLON5 |
| FBXO34 | MTMR1 | EFNB2 | ZNF398 | MAILR |
| OLFM3 | BRINP2 | HEXIM1 | DOLK | TP53I13 |

| | | | | |
|---|---|---|---|---|
| SPHKAP | UBE4B | PPP6R1 | IDE | ZNF780B |
| ARMC9 | KCNA1 | OGFR | GNG13 | ZCCHC18 |
| CCNO | OPRL1 | PGLS | TANGO6 | TMTC4 |
| ZUP1 | PARP8 | SEC14L2 | C17orf100 | SLC2A5 |
| CEP126 | CTTNBP2 | RNF157 | PEX12 | LSMEM1 |
| COG7 | PTPRJ | TOMM6 | MPP3 | R3HCC1L |
| TM9SF2 | CKLF | KIF3C | TMEM101 | UTP15 |
| MAGEF1 | USP25 | LRRTM1 | DSN1 | FNDC3B |
| NOC2L | USP15 | SEMA5A | LAMC1 | ZC3H8 |
| PLCL1 | MAML2 | FAM219B | PRKD3 | RGS7BP |
| PPP1R21 | MYL9 | PSMD4 | CEP19 | RERG |
| RGS14 | LSM12 | ANKRD65 | CHIC2 | B3GNT4 |
| CAPZB | PDRG1 | PTAR1 | TMA16 | SNHG20 |
| PLPPR4 | RBM4B | RNF185 | ZNF789 | ANKRD26 |
| DPM3 | HAPLN2 | KLHL5 | CREB3L2 | EFS |
| AMPH | PXN | PELO | CHD7 | C16orf95 |
| CELF3 | LATS1 | C1QL1 | SAAL1 | PTCHD1 |
| TEX2 | CIBAR1 | IFI30 | ZFHX2 | PRR3 |
| RBBP4 | LMAN2 | CERK | GID4 | NKD1 |
| TIAM2 | CAPRIN2 | INKA2 | OFD1 | RTKN |
| INTS11 | CYTH3 | CMSS1 | ST3GAL1 | MAP3K3 |
| YTHDF3 | LEPROTL1 | TTC14 | WDR12 | TYK2 |
| NLGN3 | ANKLE2 | ENPP5 | LY6E-DT | FAM110C |
| GPR26 | SIAH1 | NIBAN2 | SORCS1 | SPRY1 |
| AHCY | DNMT1 | MESD | MDK | TMEM209 |
| ARL2 | CSNK1G3 | PARN | IPO8 | MYO1E |
| MGAT4B | WIPI2 | SCAMP3 | RAB7B | GEMIN4 |
| BTBD10 | PDPN | PLRG1 | MRPS9 | PHKA1 |
| GNL1 | TMTC1 | WASHC5 | SVIL-AS1 | DNAJC1 |
| VAMP4 | NPPA | DIO2 | NEK9 | NARS2 |
| CARMIL2 | GDAP2 | HS3ST2 | SNX11 | TGIF1 |
| RTN4R | KCNN3 | RPA2 | PCIF1 | METTL15 |
| LMO7 | KLHDC8B | ALAS1 | SRARP | CD93 |
| HACD2 | PXYLP1 | RUBCN | SLC39A13 | BTN2A1 |
| CAMLG | FAM47E | CCDC28A | HOMER3 | SANBR |
| LNP1 | RWDD4 | DPM2 | PUSL1 | BCL3 |
| ELP3 | RIOK1 | BBS1 | LINC01750 | LIG1 |
| FAM216A | ZBTB22 | RASGRP1 | ENTPD3-AS1 | SLC31A2 |
| OXLD1 | PTCD1 | ASPDH | NAF1 | MTA2 |
| LINC01128 | LCOR | GPCPD1 | GPANK1 | TNFRSF14 |
| GRM2 | AKIP1 | DENND6B | GTF2IRD2B | RHNO1 |
| ORC5 | TTC5 | PSMB6 | TCIM | CCDC22 |
| NADSYN1 | WSCD1 | STK16 | ZNF397 | TMC6 |

| | | | | |
|---|---|---|---|---|
| MAGEL2 | SPRED3 | DLL3 | CEP250 | FRMD4B |
| CAP1 | L3MBTL1 | CHGA | ZBTB46 | TOE1 |
| VCP | DEPDC5 | SH3RF1 | GBE1 | PELI2 |
| MRPS9 | HS6ST2 | FAT1 | ARHGEF10L | NUDT7 |
| PIP4P1 | REPS1 | RBL2 | GLI4 | SPACA6 |
| FBLN7 | ZBTB20 | PIGA | PPTC7 | LIN52 |
| RIPPLY2 | RARRES2 | TBC1D25 | ZNF496 | MYO19 |
| RAB27B | LINC00869 | GAMT | MOGS | RRAS |
| U2AF1L4 | KHK | ALDH4A1 | SYNPO2 | TMEM273 |
| COL26A1 | STT3A | MIER1 | NR2E1 | ZNF595 |
| WBP11 | FKRP | CRTC2 | SNX16 | CC2D2A |
| PREP | TMEM18 | SNHG28 | DCAF4 | ART3 |
| EBAG9 | NIFK | TPR | SLC30A4 | THOC3 |
| TBL3 | EOLA2 | PTPN18 | RHPN2 | TMEM168 |
| DHX33 | CARF | SCRN3 | SLC35A3 | CLEC7A |
| SPIN3 | OXNAD1 | HIBCH | MDM4 | NXN |
| PLCB1 | PINX1 | ZNF654 | DNAJC27-AS1 | RAMP2 |
| POU3F1 | INTS8 | TMCC1-DT | P3H2 | CA5BP1 |
| ZNF335 | ADCY8 | DGKG | SRD5A3 | FBLN1 |
| SNAP29 | LINC03025 | AFAP1 | PMS2 | RGS8 |
| TAFA5 | MPPED2 | METAP1 | PNPLA7 | LYPD6 |
| GSK3B | C11orf91 | IQCE | CELF2-DT | ZNF141 |
| RXRG | DSCAML1 | INTS15 | MRPS31 | SLC12A4 |
| RNASEH2B | TNS2-AS1 | ALKBH4 | ARAF | SCARF2 |
| LHFPL4 | UTP20 | LYN | LYVE1 | LYSMD3 |
| KCNJ9 | ITGAM | MOB3B | CSTF2T | CAD |
| PANK3 | EDEM2 | PABIR1 | UBXN11 | TMEM177 |
| LRATD1 | ZNF542P | NRARP | POLR1B | SOX21 |
| NUDT14 | HSPB3 | SLC29A3 | GK5 | RAI2 |
| FLRT2 | TMEM201 | CAT | STK35 | SPR |
| GALNT17 | ISG20L2 | ATM | PAFAH2 | STAR |
| DDA1 | AADAT | RPUSD4 | IFI44L | PLP2 |
| KIFAP3 | NRG2 | ST8SIA1 | SLC27A3 | GLMN |
| NCKIPSD | TAP2 | DDX51 | ELN | POMC |
| AKAP11 | BICRAL | ABHD13 | CLDN15 | TBC1D8 |
| DAGLA | ELAPOR2 | SAV1 | RAB11FIP1 | RPP40 |
| KLHL2 | ZMYND19 | TRIP11 | CABCOCO1 | LRATD2 |
| SNX10 | ZNF37A | CSK | TMEM218 | TCP11L1 |
| SORBS2 | NANOS1 | SNTB2 | TNS2-AS1 | ROBO3 |
| DRD1 | CDAN1 | CDK12 | NAA15 | ATF7 |
| PPM1J | METTL22 | NKIRAS2 | SPON1 | TNFRSF10B |
| NBPF19 | MPV17L2 | LINC01476 | TMF1 | MEAK7 |
| UCN | RAB36 | DHX40 | MC1R | NBPF11 |

| | | | | |
|---|---|---|---|---|
| ANKZF1 | FAF1 | TRIM47 | NSUN4 | RPIA |
| SDR16C5 | KLHL23 | SNHG20 | NOTCH2NLA | ZDHHC11B |
| DNAJC1 | TRIM24 | CCDC57 | IL17RB | PDSS2 |
| COX15 | TEK | MTCL1 | ARHGEF26-AS1 | PUS7 |
| TOX2 | CFAP68 | ISYNA1 | CD2AP | IRAG2 |
| GRAMD1B | JAM3 | RELB | NSUN6 | ZNF740 |
| SELENON | ACAD8 | CHD6 | PANK1 | PRDM1 |
| NPEPPS | GPHN | SETD4 | KCNQ1OT1 | LAMA2 |
| EHBP1L1 | MRPL46 | ANOS1 | GPD1 | MIRLET7A1HG |
| ASB6 | NCBP3 | KRBOX4 | TMEM132C | POLD3 |
| VPS36 | NDUFAF5 | PABIR2 | ZNF507 | HEPN1 |
| RNF114 | ATG16L1 | AFF2 | ZNF677 | SLC1A5 |
| TMEM183A | STON2 | EP400 | ZNF217 | SALL1 |
| ACSL4 | PDLIM5 | HEBP2 | SDAD1 | CFAP210 |
| PEX5L | STARD10 | MEPCE | SEL1L3 | ZNF138 |
| ADISSP | PRKAG1 | TAF1D | ANKRD36B | FADD |
| PRKAG2-AS1 | MUS81 | USP34 | CNTN6 | ASXL3 |
| RAB11FIP3 | PDGFB | LRP3 | CPVL | POU2F2 |
| SEC61A2 | GOLGA4 | HNRNPH3 | FIGNL1 | FOXQ1 |
| IGF1R | KIRREL3 | SEPTIN4 | TAPBPL | NFATC1 |
| SORL1 | MPLKIP | RBM3 | TMEM117 | IGSF3 |
| NARF | AP4S1 | BNIP3L | NUP107 | CCDC74A |
| BCL11A | SPATA2L | LPIN1 | P2RX2 | SLC12A8 |
| SSTR1 | NUP62 | SNX13 | TMEM265 | GALNT7 |
| IPO13 | CYBA | DLD | NOS2 | ZBTB14 |
| IFITM3 | SLC15A2 | UXT | MYH7B | ZNF76 |
| PCYOX1 | MEGF10 | NOP10 | IL3RA | TMEM98 |
| KCTD1 | SSPN | BROX | SPATA33 | ZBTB40 |
| CNOT2 | FBLN1 | IGFBP5 | BLZF1 | C2orf88 |
| SLC6A15 | PPM1H | UBTF | STAT4 | TNFAIP6 |
| TM7SF2 | MTSS2 | HID1 | RRP9 | PDYN |
| GOLGA8B | MRPS16 | MYL12A | EMSY | SV2C |
| C1QA | CHAMP1 | CLIP1 | L2HGDH | KLF5 |
| DBNDD1 | EMP3 | YPEL5 | SOCS1 | ECE2 |
| CACNB2 | RNF41 | GSTA4 | METTL16 | WNT7B |
| PAM | WDR43 | BEGAIN | NKIRAS2 | EPB41 |
| DNAJB4 | PAPOLG | UBE2F | NUDT19 | SLC50A1 |
| SEC22B | JAK2 | SSR3 | BCLAF3 | SEPTIN10 |
| NDUFA9 | GALT | IPCEF1 | BRWD3 | LRRC10B |
| LARGE1 | CSTF3 | GCLM | ADRA2C | STYX |
| SYNM | UGGT2 | COPZ1 | TDRD3 | PCED1B-AS1 |
| ANTKMT | LIMD2 | DENND10 | B3GNT2 | SRGAP1 |
| SEZ6 | COQ8B | SLU7 | XIRP1 | CD44 |

| | | | | |
|---|---|---|---|---|
| PPP1R7 | ZMYM6 | CLCC1 | FSD1L | EIF4EBP1 |
| COQ8A | SANBR | SELENOH | EDRF1 | NIPSNAP3B |
| ABHD17B | BCAM | RSRP1 | TMEM179B | SPRYD4 |
| PMPCB | GPR176 | ACP1 | ARHGAP17 | HIC1 |
| KMT2C | PSME3 | RNASEH2C | PDGFB | CCDC157 |
| EGFL7 | ABCA5 | SNRPD1 | KIAA1217 | TTC38 |
| ARHGEF2 | TFCP2 | STK40 | CIAO2A | RAB35-AS1 |
| STAU2 | SMPD1 | MARK3 | LHX6 | SNURF |
| PGM2L1 | PPM1E | OPA1 | MAP4K3-DT | BNIP1 |
| COPS7A | GABRA3 | CSNK1D | SOX10 | HGH1 |
| PHF24 | NAB1 | VMP1 | IFIH1 | CEP78 |
| AP1M1 | LGALS3BP | UBP1 | ABCB7 | CTNS |
| EMC3 | YPEL1 | PRKCSH | THBS4 | TMEM132E |
| PITRM1 | TBC1D1 | TMEM39B | ANAPC2 | REL |
| PHAX | AP5M1 | ZC3H12A | UNC5B | LY6G5B |
| NAGPA | PNO1 | CPT2 | ABCC12 | ZNF783 |
| YBX1 | ADO | DAB1 | ENGASE | EPHX2 |
| ST8SIA3 | CABLES1 | FGGY | SLC13A3 | DLL4 |
| AKAP5 | UPP1 | SLC44A5 | GPRASP3 | SGSH |
| C6orf89 | MRPL49 | COL24A1 | BRCC3 | NT5C1A |
| VPS29 | CUL4A | SLC35A3 | ATG4C | TSTD3 |
| MGAT5 | ABCA1 | CHD1L | ZNF687 | CDC37L1-DT |
| MRPS7 | CSGALNACT2 | HCN3 | RASGRP3 | ZNF367 |
| ZFAND6 | BCL9 | FCGR2A | MAP3K2-DT | JUP |
| ZNF365 | TMUB1 | NME7 | SCAMP1-AS1 | IL1RN |
| ABTB1 | BIN3 | SEC16B | FAM131B-AS1 | EPS8L2 |
| RPS6KA4 | UNC5D | SERTAD4 | SLC7A2 | CSF3 |
| RALGAPA1 | DNAAF11 | INTS7 | ZNF251 | KANSL1-AS1 |
| HAGHL | INIP | PROX1 | POLE | IL6 |
| ZDHHC7 | DNAJC9 | RAB4A-AS1 | ABHD4 | ACKR1 |
| CDKN1A | ZDHHC16 | WDR35 | SNX33 | |
| FAM234B | FHIP1B | TTC27 | ASPA | |
| PHLDA1 | PGAP6 | HK2 | KCNAB3 | |
| PPT1 | ITGA3 | RPIA | BORCS6 | |
| PENK | NXT1 | ANKRD23 | ITGB4 | |
| B9D1 | COL20A1 | SPOPL | PRCD | |
| KLHL22 | MSN | NOSTRIN | ERCC8 | |
| CHMP2B | LNCBRM | NEUROD1 | SGCD | |
| RAB33A | DDX27 | PPP1R1C | CERS4 | |
| ATF3 | VARS1 | NUP35 | SLC16A14 | |
| MAP9 | PEX26 | WDR75 | SLITRK3 | |
| OTUD6B-AS1 | ELAC2 | CTDSP1 | SLC35A4 | |
| FARSB | NICN1 | CT75 | PTTG1 | |

| | | | |
|---|---|---|---|
| ADAM11 | RANBP6 | CCDC174 | MMP16 |
| TARDBP | ZPR1 | RBMS3 | GTF2H3 |
| COA6 | ERG28 | ZNF621 | OBI1 |
| ETV5 | GAA | SFMBT1 | SLCO1A2 |
| NAA20 | SETDB1 | MITF | LINC01141 |
| BDNF | HYCC1 | EPHA3 | BASP1-AS1 |
| LRRC47 | AKR1C3 | PROS1 | ADAM19 |
| MMADHC | CKAP4 | GRAMD1C | AP4M1 |
| WRNIP1 | BACE2 | SLC49A4 | ATAD2 |
| SNAPC5 | EFNB1 | ARHGEF26 | AKR1C2 |
| RAPGEFL1 | TSPAN6 | RNF212 | ERCC6 |
| CRAT | LCP2 | UVSSA | LINC00595 |
| FTO | L3MBTL4 | LGI2 | INTS13 |
| STXBP5L | ABHD5 | TBC1D19 | FBLN5 |
| CCNDBP1 | KCTD15 | RBM47 | RLBP1 |
| LAMB1 | MAPK11 | GABRB1 | MYO1C |
| SAMD12 | SEC24D | SGMS2 | MIR23AHG |
| CCN1 | SLC25A16 | CYP2U1-AS1 | EIF5B |
| VPS41 | NECAP2 | SPRY1 | B3GALT1 |
| CHID1 | PHRF1 | SAP30 | WDR3 |
| SEC14L1 | DSEL | TMEM267 | GIGYF2 |
| BCS1L | EIF2B3 | POLR3G | RYK |
| TMEM198 | GPR161 | FAM114A2 | TAX1BP3 |
| ASPHD2 | RPS6KC1 | CCNJL | MAP2K7 |
| ANKRD9 | C2orf42 | PHYKPL | CYP20A1 |
| RAB3GAP1 | IFT122 | ZFP62 | RFC4 |
| UGCG | NELFA | FARS2 | TLR2 |
| PSMG3 | CARTPT | TNXB | MSC |
| C6orf136 | SLF1 | HSD17B8 | ZNF169 |
| ATG4B | HCG11 | EFHC1 | ARRDC1-AS1 |
| RNF2 | SLC29A1 | COL9A1 | PACSIN3 |
| EMX1 | PDE10A | MED23 | GCH1 |
| LINC01511 | ZNF394 | USP42 | RHOJ |
| TMEM223 | ANGPTL2 | UMAD1 | FAM83D |
| GXYLT1 | ARMH3 | SOSTDC1 | GGT5 |
| OSER1-DT | MBD6 | CDCA7L | TCF7L2 |
| SNRPC | EME2 | YAE1 | LIN54 |
| MRFAP1L1 | CNEP1R1 | IGFBP3 | LHFPL3 |
| TP53BP1 | VPS25 | STAG3L4 | NCOA3 |
| DENND5B | SLCO4A1 | COL1A2 | TMEM60 |
| CD151 | TMEM121B | ING3 | WDR59 |
| EXTL3 | PLEKHM1 | KBTBD11-AS1 | ATAD3B |
| KCNG1 | DENND3 | SLC7A2 | ATR |

| | | | |
|---|---|---|---|
| SYNE1 | VPS16 | BMP1 | ZNF346 |
| KCNMB4 | DMXL1 | EIF4EBP1 | TJAP1 |
| CCNL2 | HAVCR2 | CLXN | CEP57L1 |
| CSNK1E | GOLT1B | ADHFE1 | ABHD11 |
| PSMA5 | CCDC107 | TAF2 | P4HA1 |
| COPS4 | TMEM115 | ATAD2 | USP54 |
| KLHL12 | ZBTB11 | NTAQ1 | NUFIP1 |
| UTP4 | RBM33 | KCNK9 | RPUSD2 |
| CAB39L | PTDSS2 | ZFTRAF1 | ZNF48 |
| MCTS1 | TBC1D15 | VLDLR-AS1 | ALOX12-AS1 |
| TPD52 | PAGR1 | IL33 | CHD1L |
| SNRNP25 | SLC38A10 | HRCT1 | CYP1B1 |
| MIR100HG | PRPF6 | ANKRD18A | C7orf25 |
| MARK1 | SMARCA1 | PTGS1 | NSMAF |
| PLXNA4 | ZBTB33 | GBGT1 | POLL |
| EMX2 | TNFRSF21 | CACFD1 | PITPNA-AS1 |
| INO80 | AFF3 | ARHGAP22 | ESCO1 |
| PTGR3 | WDFY1 | SLC16A9 | RALGAPA2 |
| LRRC41 | ANAPC4 | TSPAN15 | RFPL2 |
| KLHL21 | RAD1 | FAS | SHROOM2 |
| GOT2 | SUDS3 | DNMBP | GLMP |
| CHMP1A | GMPR2 | STN1 | SLC30A10 |
| VPS35 | LMF1 | ZDHHC6 | COG2 |
| LIN7B | TIMM22 | ZDHHC13 | TFPI |
| MGAT4A | ARRDC2 | LINC02696 | TYW5 |
| RAB4A | EPS8 | KCNK7 | SCLY |
| PTPN4 | CITED2 | FADD | SETMAR |
| SLITRK1 | LMCD1 | TPBGL-AS1 | MANBA |
| VIP | TMEM192 | CASP4 | AGA |
| EDEM3 | ARHGAP27 | VWA5A | CDKAL1 |
| PRPF40B | PRRX1 | ROBO4 | UST |
| WDR59 | LTBP1 | HYLS1 | MDFIC |
| CLDND1 | INO80D-AS1 | GSG1 | NUDCD1 |
| AACS | CFAP44 | RECQL | TAF2 |
| DYNC1I2 | NFYA | ETFBKMT | FUT11 |
| BRD3OS | PRDM1 | LINC00938 | HSPB2 |
| U2SURP | ACSL5 | MMP19 | TRAFD1 |
| PPP1R9A | COQ5 | NUP107 | TM9SF1 |
| LRWD1 | CAVIN1 | SBNO1-AS1 | ARMH4 |
| KLF12 | NAGLU | MTUS2 | FMN1 |
| DHRS7B | TGM2 | LACC1 | RANBP10 |
| WDFY3 | TBL1X | TGDS | LIMK2 |
| HEY1 | SPATA20 | COL4A2 | TMCC1 |

| | | | |
|---|---|---|---|
| RFX3 | FBXO30 | DHRS1 | LIF |
| CUL2 | PIGV | STYX | TRMT9B |
| TSPAN14 | ECHDC2 | SOCS4 | SLC15A4 |
| USP2 | ADGRL4 | PLEKHG3 | B3GNT5 |
| KPNA3 | LRIG2 | ZNF839 | SHPRH |
| DRG1 | ACKR1 | DLL4 | ZNF407 |
| LPGAT1 | C1orf35 | AP4E1 | SESN2 |
| MPND | RRP9 | PIGB | SEMA3F |
| UQCC1 | PBX2 | USP3-AS1 | CYB561D2 |
| EFCAB7 | PEX3 | JPT2 | SHROOM1 |
| D2HGDH | ZFAND2A-DT | ZNF205 | GTF2H4 |
| OSBPL11 | C7orf25 | NKD1 | DNAAF5 |
| ARL15 | RNF32-DT | SALL1 | SOSTDC1 |
| KLF5 | ZNF623 | CDH5 | ERI1 |
| CACNA1H | KCNK4 | TRADD | PARP10 |
| ABCC12 | PANX1 | RANBP10 | RGP1 |
| ZNF423 | RNF26 | FANCA | RESF1 |
| GAS8 | KMT2D | PLD6 | BAZ1A |
| YBEY | ZNF689 | TLCD1 | PLEKHG3 |
| SYN3 | BORCS6 | RAD51D | NPIPB13 |
| ARHGAP4 | SMG8 | CDC6 | COL6A2 |
| MAN1A1 | UNK | MAP3K14 | KDM6A |
| PCDH7 | GUCD1 | ERN1 | CELSR3 |
| ARHGAP23 | H2AC25 | METTL4 | BMPR1B |
| ECHDC1 | DHRS3 | CDH19 | LSM11 |
| CDC42EP3 | ZNRF3 | TINCR | RHOG |
| UXS1 | SMAD1 | CLEC4G | NSRP1 |
| LRRC14 | CACNA1C | ZNF136 | HSF4 |
| MTPAP | ASAP3 | DCAF15 | PROX1 |
| SEC23IP | ZNRF2 | ADGRE5 | GALNT15 |
| POLD4 | NUP188 | ANO8 | TRIM35 |
| METTL2A | SPSB3 | ZNF675 | HMBOX1 |
| GGA3 | LPL | ZNF461 | ADCY8 |
| SPHK2 | EIF2B1 | ZNF585B | GLE1 |
| ZSCAN5A | GLT8D1 | DHX34 | AMPD3 |
| RHOT1 | CDH10 | GRIN2D | CASP4 |
| SOCS7 | SLC12A2-DT | RRAS | C1S |
| RTBDN | PPP3CC | ZNF614 | TUNAR |
| H4C8 | VCPIP1 | ZNF432 | TRADD |
| TBRG1 | ZBTB10 | ZNF581 | C21orf91 |
| SDHAP4 | GLIPR1 | JAG1 | ZFY |
| CCNK | RBM23 | ZNF133 | DHFR |
| MORC2 | EMC9 | ACSS1 | ATP6AP1-DT |

| | | | |
|---|---|---|---|
| DNASE1L1 | GAN | IFT52 | DUSP12 |
| DDHD1 | SLC25A42 | TP53RK | SEC22A |
| PER1 | UBOX5 | RBM11 | SMC2 |
| FH | CBFA2T2 | CRYBB1 | SFMBT2 |
| LMBR1 | CEP104 | KREMEN1 | TEX30 |
| EPDR1 | FUCA1 | SHISA8 | SPPL2A |
| UNC119 | WDR3 | TYMP | HIGD1B |
| SLC2A13 | CPSF3 | SHROOM4 | LRFN3 |
| TRHDE | HS1BP3 | HMGN5 | FCHSD2 |
| SCN8A | SMYD5 | APLN | CEP126 |
| GLRB | ATR | SMIM10 | KTI12 |
| NOPCHAP1 | PIGG | TMEM187 | LRIG2 |
| PEF1 | ADAM19 | OLIG1 | LINC01138 |
| CSRNP3 | PGBD1 | AP2A2 | GORAB |
| CDC42BPA | KHDC1-AS1 | PCBP1 | KANSL1L |
| LINC00641 | GTF2IRD2 | KLHL24 | FAM86DP |
| AGRN | UPK3BL2 | MRPS25 | CLCN2 |
| GFM1 | PDE7A | SMIM20 | ABHD18 |
| RNF121 | PTPDC1 | NUP205 | ZNF608 |
| ZC3H15 | REXO4 | CORO1B | ZKSCAN8 |
| GSTO1 | MIR1915HG | RUVBL2 | LFNG |
| AJAP1 | MPP7 | RAP1A | DOCK5 |
| NKAIN2 | TCERG1L | ARMC8 | THAP1 |
| CCDC25 | RAB35-AS1 | PAQR8 | TMEM67 |
| TSFM | CLYBL | SSRP1 | TIGD5 |
| INTS6 | DCAF4 | SAMD14 | PUM3 |
| STOM | SULT1A3 | MYRF | MIR1915HG |
| RAB5B | FA2H | S100A10 | ACSL5 |
| MPG | DVL2 | CEP350 | CASP7 |
| SRA1 | ZNF615 | GNPDA2 | MCAM |
| SPATA2 | A1BG-AS1 | SRD5A3 | CPM |
| TUBG2 | BMP2 | CCDC112 | ZNF140 |
| PSMA3 | SLC2A4RG | BRAT1 | USP3 |
| GARS1 | RFPL2 | TRAPPC9 | TBC1D2B |
| CD53 | GCNA | SPTLC1 | SETD6 |
| CLEC2L | NDST2 | ARRDC1-AS1 | ZNF566 |
| RALA | TAGAP-AS1 | UNG | ZNF583 |
| DGKA | IFFO1 | TMEM179 | GART |
| KTN1 | SNHG16 | YJU2 | MX2 |
| NIPA1 | NIPBL-DT | OFD1 | SLC44A3 |
| DGKB | H1-2 | TSTD1 | OSBPL11 |
| ELK1 | FAM210A | NFIX | HYCC1 |
| HSPBP1 | CTBS | PSMB10 | CDK6 |

| | | | |
|---|---|---|---|
| RC3H2 | FSTL1 | RNASE1 | TNFRSF10D |
| STOML1 | UMPS | SRCIN1 | ITIH5 |
| SLC7A6OS | BCHE | POLR2I | FIBIN |
| CC2D1B | GDNF | GNPTAB | NAA40 |
| ZNF496-DT | QRSL1 | TSC22D3 | AP5B1 |
| LYPD8 | CLEC16A | MAPK10 | CWF19L2 |
| ALKAL2 | GID4 | AHCYL1 | DHRS1 |
| TMEM87B | CCL8 | ITGB5 | KIAA0586 |
| ACAD11 | ZNF175 | WWTR1 | IRF8 |
| SPTSSB | ZNF419 | LRRN3 | FAM222B |
| RFC4 | PI4KAP1 | EGLN3 | P2RY11 |
| TNIP2 | MRPL53 | CCDC88A | TRPC5 |
| CPVL | NFKB2 | ATAD3B | GSTM5 |
| TMEM243 | RHOT2 | FNDC10 | COL23A1 |
| VIPR2 | SYNC | PLA2G5 | MBIP |
| SPX | DCTN1-AS1 | SESN2 | YAE1 |
| MVK | POLK | DNALI1 | MED9 |
| MTRFR | SLC36A1 | EPHA10 | CDK11B |
| PCDH20 | FOXP4 | BGLAP | SH3D21 |
| ZWILCH | CHRM4 | DDR2 | ESRRG |
| ZNF778 | RELA-DT | BATF3 | GRASLND |
| MPP3 | ABCG4 | HEATR1 | CAPG |
| FDXR | ANKRD13A | NTSR2 | SOWAHC |
| BAIAP2L2 | HEATR3 | ALMS1 | SCRN3 |
| CPT1B | SNAI3-AS1 | ANKRD36C | COL7A1 |
| GPRASP3 | CDK10 | PAX8-AS1 | NEK11 |
| CARD19 | HEXD | CYBRD1 | UVSSA |
| ADGRA1 | DCC | DLX1 | NOP14 |
| TUSC2 | HIF3A | ACKR3 | SLC35B3 |
| KYAT3 | HAS1 | LRRC3B | HCG11 |
| PHF5A | ZSCAN1 | CCR5 | TRIM39 |
| ASS1 | GNB1L | CCDC51 | MTERF3 |
| DCUN1D5 | SPSB1 | ECE2 | NOL8 |
| EPHB6 | PARP14 | REST | FKBP15 |
| CACNA1B | EXOSC2 | USP38 | KIN |
| DNAJA2 | WDR20 | CTSO | RBMS2 |
| SLC24A2 | RETREG3 | CPLANE1 | FCF1 |
| KNDC1 | TANGO2 | SGCD | WDR24 |
| ZMAT3 | PRPF3 | ZSCAN26 | PAK4 |
| INF2 | BLZF1 | DXO | RDH13 |
| HNRNPH2 | SOAT1 | TAP1 | ZNF550 |
| PLEKHA1 | LBR | AK9 | HCN3 |
| SEMA4D | STAT4 | ERI1 | DUBR |

| | | | |
|---|---|---|---|
| RNMT | MBD4 | HMBOX1 | CEP44 |
| HM13 | CEP41 | TOX-DT | MMD2 |
| BAG1 | PARD3 | CHD7 | MBLAC1 |
| TSPAN13 | IFIT2 | ZFHX4 | KCTD9 |
| ABCC5 | NSMCE4A | MAILR | PSKH1 |
| OMG | METTL15 | MYC | MLYCD |
| ARL16 | TMOD3 | TNC | DNAH17 |
| ANGPTL4 | ZNF397 | REEP3 | APBA3 |
| HBS1L | LINGO3 | MACROH2A2 | TICAM1 |
| PJA1 | MAG | SGPL1 | GTPBP3 |
| ZFYVE9 | ALKBH3 | LDB1 | ZNF570 |
| MTERF4 | EXOSC7 | ATL3 | NKX2-2 |
| TBCC | SLC30A10 | PYGM | RPGR |
| LRRC73 | B3GALNT2 | UCP2 | MOSPD1 |
| ELOVL4 | NSUN2 | CNTN5 | NEK7 |
| AGPAT4 | SH3PXD2B | ADCY6 | NUP85 |
| TMEM25 | SDHAF4 | ATF1 | TOR3A |
| PPHLN1 | SLC25A32 | ZNF740 | ELOVL7 |
| GLCE | ZFPM2 | ATF7 | BMPER |
| DNAJB5 | B4GALT1 | THAP2 | CHRNB1 |
| HDAC2 | NPAT | CHFR | ZNF154 |
| BORCS8 | TPT1-AS1 | PAN3 | NUPR1 |
| PTPN5 | DMAC2L | ARHGEF40 | GALC |
| SYAP1 | SLC38A7 | LIN52 | RNASEH1-DT |
| ATRN | TRAPPC8 | ATXN3 | ATXN7L2 |
| SVBP | SETBP1 | CAPN3 | DRAM2 |
| NBDY | TK2 | SEMA6D | UCN |
| MYH10 | SDS | TRIP4 | LTBP1 |
| EEF1E1 | PAK1IP1 | BBS4 | GYPC |
| KIF21A | ATP2B1-AS1 | CBFB | SLC11A1 |
| GTF2A1 | HPS4 | CTNS | XPC |
| KCNQ5 | SNRNP40 | MAP3K3 | PPP2R3A |
| EIF3D | OBSCN | MYO15B | LCORL |
| TMEM123 | MAD1L1 | ESCO1 | ERAP1 |
| NLGN1 | PDE1C | ZNF561-AS1 | ZBTB9 |
| ATAT1 | ZNF568 | ZNF791 | SNX10-AS1 |
| STXBP5 | SLC25A17 | ZNF507 | ZDHHC12 |
| BABAM1 | MTF1 | KPTN | C17orf114 |
| HAT1 | BRINP3 | EBF4 | GJC1 |
| UBAP1 | EIF2D | SYS1 | DHRSX |
| FAM43B | GAREM2 | RGS19 | LIX1 |
| TMEM237 | MOGS | IL10RB | ZNF585B |
| ATXN1L | HNMT | MX1 | AP3B1 |

| | | | |
|---|---|---|---|
| RAB40C | LINC02610 | ABCG1 | SEC16B |
| NR2F1-AS1 | GAB1 | RTL10 | PDLIM4 |
| PEPD | ZNF827 | PIR | MVP |
| DPP10 | LY86-AS1 | ZRSR2 | VCAM1 |
| UBE2V2 | TBC1D22B | CHST7 | LINC01090 |
| PPM1A | BAALC-AS1 | EIF3I | H1-2 |
| KLHDC9 | PTGR1 | COX17 | STK3 |
| RARA | GSTO2 | PLOD2 | PIGO |
| MCAT | JAKMIP3 | H4C3 | TBC1D2 |
| ERCC1 | APIP | NABP2 | TOR2A |
| TRIM27 | HSPB2 | BECN1 | SYT9 |
| CCNQ | C3AR1 | CNTN2 | LINC02696 |
| SUPT7L | IKZF4 | SNRNP27 | FOLH1 |
| RPN1 | SOX21 | MRPL12 | THOC6 |
| SEC13 | SHF | EIF2B5 | DMKN |
| POP7 | CEMIP | BLVRA | ZFP14 |
| MFGE8 | ZFR2 | RBFOX3 | INSM1 |
| POGZ | PLA2G4C | SOCS3 | TOP3B |
| LPAR1 | AAR2 | ZCCHC24 | AFF2 |
| DTD1 | WWC3 | MBOAT2 | ZNF395 |
| PRPS2 | UPF3B | IMPDH1 | CTBS |
| CACNB1 | TNFRSF10B | TIMP2 | MASTL |
| AMZ2P1 | DGKH | SKIL | IFIT2 |
| NECTIN1 | GYPC | SPART | |
| PPM1M | PLXND1 | ZDHHC18 | |
| CPOX | IRAK1 | WDTC1 | |
| IGSF11 | RLF | NOTCH2 | |
| ZNF189 | MFSD1 | PCCB | |
| LRRTM3 | CASQ1 | VWA5B2 | |
| ZDHHC5 | TMEM177 | YTHDC2 | |
| DIXDC1 | DENND6A | MRNIP | |
| UBE3B | IQCB1 | PMS2 | |
| LHFPL6 | ZNF141 | GPR107 | |
| SLC12A6 | CTBP1-DT | ADK | |
| FAM174B | C1QTNF3 | CCND1 | |
| AMDHD2 | ATP10B | MRPL48 | |
| USP19 | THOC3 | RDX | |
| CPNE8 | MDGA1 | KANSL2 | |
| THUMPD3-AS1 | TRMT11 | RNF34 | |
| MOB2 | CCDC71L | SERTM1 | |
| SCAMP4 | ZNF767P | ELF1 | |
| SLK | GTF2E2 | TMEM87A | |
| DCAF11 | NUDCD1 | SMAD3 | |

| | | |
|---|---|---|
| KLHL42 | AGPAT2 | LITAF |
| NRG3 | TRAF2 | KIF22 |
| RUFY2 | ECD | RUNDC1 |
| SPEF1 | SEC31B | MCOLN1 |
| PIK3CD | MEN1 | YJU2B |
| P3H1 | GNB3 | AXL |
| SPRTN | BORCS5 | PNKP |
| DESI2 | STAT6 | RNF24 |
| BLTP3A | CAPS2 | RRBP1 |
| BAG2 | OGFOD2 | SRP72 |
| POT1 | GABPB1 | HYI |
| KCNS2 | DDX28 | SUCO |
| FAM78A | NIP7 | FEM1C |
| SLC36A4 | ZNF830 | KDM7A |
| ACSF3 | RSAD1 | MON1B |
| MIEF2 | ASXL3 | SDC4 |
| ELL | MOB3A | FAM110B |
| ZNF790-AS1 | JAK3 | MRPS5 |
| ZNF224 | ZNF260 | ATPAF1 |
| PXMP4 | CNFN | TSPOAP1 |
| UTP14A | FUT1 | ITGB1BP1 |
| MSANTD4 | ZGPAT | UBA1 |
| IWS1 | SCAF4 | PPOX |
| TENT4A | RBMX2 | HLA-DPA1 |
| FOCAD | GALE | SLC17A5 |
| POLE3 | DCAF17 | UNC13B |
| ZFYVE27 | TRIM62 | NELFB |
| TMED3 | LINC00622 | FBH1 |
| WDR18 | MSTO1 | NT5C2 |
| MISP3 | CXCR4 | AMN1 |
| LRRC7 | KANSL1L | SCAMP2 |
| KIAA2013 | USP37 | HBA1 |
| MYO9A | NUP210 | ARL5B |
| FLYWCH2 | NAA80 | VWA1 |
| MRPL50 | P3H2 | ZNF362 |
| NWD2 | GRK4 | RNPEP |
| UFL1 | ART3 | LRRN2 |
| NT5M | MCUB | CLIP4 |
| SMAD7 | NPY1R | ERICH2 |
| ZNF567 | LYSMD3 | EAF1 |
| SPATC1L | FSTL4 | MON1A |
| ZNF24 | RNF144B | SETD7 |
| CDC14B | RMND1 | GNPDA1 |

| | | |
|---|---|---|
| CNNM1 | RUNX1T1 | RAB24 |
| WDR26 | PYCR3 | HEIH |
| CTNNA1 | DKK1 | PLGRKT |
| MB21D2 | SUFU | MINDY3 |
| RNF150 | LDLRAD3 | SORCS1 |
| B4GALT7 | MS4A7 | DDX47 |
| ZCCHC2 | USP28 | HRK |
| RMCX5-GPRAS | TMEM218 | COG3 |
| TSNAX | ITGA5 | NUMB |
| SND1 | GPC5 | NPRL3 |
| MRPS2 | ANKDD1A | SETD1A |
| TMEM106B | KCNJ12 | SYCE1L |
| ATXN7L3B | NOL11 | PHF12 |
| CEP250 | TTC39C | ASIC2 |
| C2CD2 | SALL3 | PNMT |
| CACNB4 | ZNF415 | KRT17 |
| NTMT1 | XPNPEP3 | KAT2A |
| FEZ2 | PLP2 | APCDD1 |
| MORN2 | DCBLD2 | RIT2 |
| PFDN6 | NSMAF | RBCK1 |
| CACNA1G | TRABD | ZHX3 |
| UBE2O | RPS6KA6 | PCNT |
| NUDT11 | ZNF576 | SLC25A18 |
| GRM7 | | GTPBP6 |
| CRYBG3 | | FGD1 |
| PIK3R4 | | GLA |
| SMIM43 | | TTTY14 |
| GPX3 | | B3GAT3 |
| POLR3B | | CTTN |
| CLN5 | | HNRNPD |
| SNX11 | | ZDHHC14 |
| ZNF570 | | RAP2A |
| SARS2 | | SAR1A |
| RRP1 | | GLRX2 |
| RAB9B | | KCNJ3 |
| RASA2 | | NCK2 |
| CBY1 | | RND3 |
| DHPS | | QRICH1 |
| BTG3 | | TARS1 |
| EPHB2 | | UNC13C |
| CRYZ | | AHSA1 |
| PASK | | SENP6 |
| GK5 | | ARPC4 |

| | |
|---|---|
| NUDT12 | DR1 |
| SLC12A9 | LYPLA2 |
| FGF17 | RGS16 |
| HIF1AN | SLC20A1 |
| POC1B | CYFIP1 |
| ATP6V0A2 | GDPD1 |
| TMEM63C | PIAS2 |
| TUNAR | TIAM1 |
| SLC30A4 | WDR45 |
| CENPN | RTF1 |
| PFAS | TBC1D9B |
| MOCS3 | SNPH |
| ZC4H2 | PDE8B |
| CISD3 | PPP4C |
| PGD | CD9 |
| SRGAP2 | SPTBN4 |
| POM121 | SIAH2 |
| ARL1 | CLINT1 |
| MFAP1 | FGFR1 |
| MED31 | N4BP2L2 |
| PDGFA | MT1F |
| BEND6 | AK3 |
| NUTM2A-AS1 | BCL6 |
| MARS1 | SDF4 |
| SMPD3 | SOX2-OT |
| MANEAL | CA2 |
| EFNA3 | CPNE4 |
| AHCTF1 | CLSTN2 |
| TMEM131 | LINC02217 |
| DNAJC21 | ADAM22 |
| MRPL2 | UBE2Z |
| SLC45A4 | SLC1A1 |
| HIKESHI | LRP11 |
| PRICKLE1 | AKAP9 |
| NIPA2 | SCAF11 |
| MPI | CES4A |
| LRRC49 | UBE2QL1 |
| CKS2 | ZNF638 |
| GNG11 | DZIP1 |
| WDR5 | NAE1 |
| GFOD2 | XRN2 |
| UBE2Q1 | MRPL40 |
| ARPC1B | MIA3 |

| | |
|---|---|
| ITGB3 | ROBO2 |
| JPX | DAB2IP |
| TMEFF2 | GLOD4 |
| ZSWIM7 | DUSP16 |
| SLC7A1 | TMEM141 |
| FAM32A | ZBTB38 |
| PITPNA | EPB41L4B |
| NEK6 | PSMC2 |
| MAGI1 | NTM |
| ACAA1 | COX7A2L |
| HDAC9 | NARS1 |
| FARSA | FAM168A |
| UBL4A | PTPRZ1 |
| ATF6 | MCF2L |
| LAP3 | SLC20A2 |
| NDUFAF4 | CTDNEP1 |
| PELP1 | TMEM256 |
| MT1H | ZNHIT3 |
| LETMD1 | RAB4B |
| RNF126 | NNT |
| PLEKHA3 | TSR3 |
| NCEH1 | PLEKHM2 |
| NEURL1B | SSBP1 |
| DNAAF9 | CLK1 |
| CAMK1G | DHX9 |
| ERC2 | C1D |
| PALM2AKAP2 | ATG12 |
| RBM10 | WARS1 |
| NPAS4 | EIF4A3 |
| ATP8A2 | PEX19 |
| TOX4 | TOR1AIP2 |
| MED24 | TCAIM |
| AFG3L2 | RABL3 |
| ELP2 | C3orf70 |
| SACS | TRAPPC11 |
| ARID4A | ABLIM3 |
| MAP2K4 | F12 |
| GPBP1L1 | DMTF1 |
| SUSD4 | VPS50 |
| SIDT1 | LONRF1 |
| SLC9B2 | CARNS1 |
| RAB33B | DHCR7 |
| MARCHF1 | UNKL |

| | |
|---|---|
| TRAPPC13 | INO80C |
| SIRT5 | SMAD2 |
| CCNC | PKN1 |
| COMMD5 | TCFL5 |
| AOPEP | SCNM1 |
| VPS11 | LINC00844 |
| PPP2R3C | OAF |
| SPATA7 | PPIE |
| DERL2 | NRAS |
| KRT222 | UAP1 |
| BCAS3 | PSEN2 |
| SAMHD1 | JMJD4 |
| RASD2 | KLF7 |
| CDC73 | SLC16A14 |
| TRRAP | ZNF385D |
| ARMC1 | LRRFIP2 |
| PDE4A | NIT2 |
| NCAPH2 | RXFP1 |
| LAMP2 | HPF1 |
| ATN1 | FBXO38 |
| CCDC92 | FAF2 |
| RGS5 | KHDC1 |
| LGI3 | GSDME |
| ADAMTS1 | GATAD1 |
| ZRANB2 | CNOT4 |
| C6orf120 | ZNF746 |
| XKR4 | ARHGAP39 |
| CRELD2 | DCAF12 |
| ACAT1 | TLE4 |
| HTR1A | PBX3 |
| RRP12 | AIF1L |
| SRP54 | MARCHF8 |
| OSBPL6 | UBE2D1 |
| COMMD9 | BTAF1 |
| UBALD1 | MFSD13A |
| COX11 | BET1L |
| FBXL3 | ZC3H12C |
| VKORC1 | PPP2R1B |
| IKZF5 | SRPRA |
| COMMD4 | NINJ2 |
| PDAP1 | CSAD |
| CUX2 | SPATA13 |
| SRGAP1 | CINP |

| | |
|---|---|
| ZNF154 | FMN1 |
| QSOX1 | TTBK2 |
| MICAL2 | ZNF280D |
| INPP1 | WHAMM |
| ADNP2 | RHBDL1 |
| UHMK1 | ADCY9 |
| SCN3A | MED11 |
| RAPH1 | PHF23 |
| RRAGD | NAT9 |
| FZD3 | PGS1 |
| SS18L1 | RANBP3 |
| SIRT1 | CCDC9 |
| ZNF281 | NCOA5 |
| DROSHA | CHRNA4 |
| CCAR2 | SH3BGR |
| NUP214 | TXNRD2 |
| TNFAIP8L1 | TBC1D10A |
| KDM4B | TOM1 |
| CRLS1 | EBP |
| TAB3 | MRAS |
| SLC25A14 | CADM1 |
| NUCB2 | MRPL27 |
| RASGEF1B | DCTN4 |
| FURIN | IP6K2 |
| PDZD11 | PIP4P2 |
| ANKRD39 | GANAB |
| DAGLB | PWWP3A |
| PLPBP | TXNDC12 |
| LEMD3 | MCTP1 |
| GLT1D1 | TCERG1 |
| ILRUN | TERF1 |
| TBCD | PARP2 |
| LINC02693 | CDH11 |
| SC5D | HS2ST1 |
| DDX42 | PRPF38B |
| EZR | NR3C1 |
| ZBTB40 | ALDH5A1 |
| FAM76A | CCZ1 |
| TCEANC2 | SGCE |
| NBPF12 | FMC1 |
| SCYL3 | CALCOCO1 |
| MAP3K21 | SPTSSA |
| THNSL2 | HAPSTR1 |

| | |
|---|---|
| SHQ1 | EVI5L |
| LINC01990 | CDKL5 |
| KLHL6 | ATP2C1 |
| ACOX3 | CAD |
| FRAS1 | TPRA1 |
| LINC02433 | NEIL2 |
| HAPLN1 | PRXL2C |
| CTXN3 | LZTS2 |
| NKAPL | CHORDC1 |
| DUS4L | USP30 |
| NUDT18 | MLXIP |
| NPBWR1 | LEO1 |
| LINGO2 | IMPACT |
| ZBTB34 | STX10 |
| PNPLA7 | MAGED2 |
| MKX | PHLDB1 |
| LRMDA | KCNA3 |
| TRIM68 | TARS2 |
| DDB2 | SEMA6C |
| LIPT2-AS1 | MRPL9 |
| HTR3B | ALDH9A1 |
| APOA1 | RGS8 |
| PIK3C2G | EPHX1 |
| CABP1-DT | DNAJC27 |
| TRPC4 | FASTKD2 |
| DICER1-AS1 | CAPN7 |
| SHC4 | GORASP1 |
| CIB2 | TMEM175 |
| GLIS2 | ZNF622 |
| CBLN1 | RGS7BP |
| MMP2 | JMY |
| ZNF469 | THBS4 |
| SLC16A11 | CD14 |
| ZNF431 | TTBK1 |
| ZNF350 | FAM184A |
| ZNF549 | NHSL1-AS1 |
| TCP10L | ARID1B |
| CCDC117 | SLC35B4 |
| PPEF1 | SORBS3 |
| ATRAID | C8orf82 |
| CAST | FUBP3 |
| PCP4L1 | EXT2 |
| SLC2A5 | SYT12 |

| | |
|---|---|
| IRF7 | ARAP1 |
| CHI3L1 | ARHGDIB |
| ARG2 | PRKD1 |
| SMG7 | KIAA0586 |
| BACE1 | EIF2B2 |
| CORO1C | MEIS2 |
| CACNA1A | ISG20 |
| AIG1 | ENKD1 |
| CCNYL1 | NFAT5 |
| STOX1 | C16orf46 |
| BCKDK | PNPO |
| FUNDC1 | RNF152 |
| STARD3NL | TIMM29 |
| ULK1 | FCGRT |
| NFKBIZ | NOL4L |
| RER1 | DNTTIP1 |
| UTP3 | EMID1 |
| NACAD | BCLAF3 |
| TRUB1 | HDAC6 |
| CAPN5 | CPNE5 |
| JAG2 | TCEAL6 |
| TIMM21 | WTAP |
| MIR7-3HG | USP4 |
| L3MBTL2 | RHOBTB3 |
| SAMD4B | SH2B3 |
| DNAJC18 | GALNS |
| VPS18 | COASY |
| ATG101 | FOXK2 |
| RNF6 | PANX2 |
| PANK4 | NKRF |
| RAB3B | ICAM1 |
| FAM78B | CNOT7 |
| DPP10-AS1 | BAG3 |
| ACVR2A | RAB5A |
| ISY1 | FBXW5 |
| ZNF76 | SLC12A2 |
| TDRP | GDI2 |
| MMP16 | ECHS1 |
| STYK1 | RANBP9 |
| ABCD2 | RNF5 |
| SUOX | CSTPP1 |
| UQCC6 | NUMA1 |
| NUDT15 | FBLL1 |

| | |
|---|---|
| TBC1D4 | EIF2AK1 |
| RABGGTA | KXD1 |
| TUBG1 | RDH11 |
| ATP9B | SBF1 |
| ARL8A | CNDP2 |
| CELSR3 | TRIR |
| ASB8 | NRBP1 |
| RYR2 | IL17D |
| DGKI | RBM5 |
| VPS13A | FOXJ3 |
| KANSL1 | PLEK |
| TMEM242 | C2orf69 |
| CCPG1 | HMGXB3 |
| KCNIP3 | PPARD |
| FAM174A | FERMT2 |
| CBX4 | SEC11A |
| MBD3 | FMN2 |
| RPS6KA2 | PI4KB |
| ZNF25 | SLC41A1 |
| EGFR | MPHOSPH10 |
| ODF2 | PAIP2B |
| TOR3A | GABRG1 |
| MCM7 | MYLIP |
| EPHX2 | TP53TG1 |
| DNAH10OS | CBLL1 |
| TMEM126B | TNPO3 |
| FAM171A2 | ZC3HAV1 |
| GNAZ | PNPLA2 |
| GALNT11 | FCHSD2 |
| RGS3 | RAB2B |
| TRAPPC2 | ISCA2 |
| NSDHL | CLPX |
| GSN | ZNF710-AS1 |
| ASRGL1 | EVI2A |
| CCR1 | CNDP1 |
| MFSD10 | LINC00665 |
| ANKRD50 | NFKBIB |
| TM7SF3 | VRK3 |
| TLE3 | SPECC1L |
| PFDN4 | NNT-AS1 |
| TCEAL8 | RTL5 |
| NBPF1 | KCNQ4 |
| AKR1A1 | ELOVL1 |

| | |
|---|---|
| PRELP | FAAH |
| B3GNT2 | PRKAA2 |
| STAMBP | PIGK |
| ST6GAL2 | GOLPH3L |
| UGGT1 | PIK3C2B |
| B3GALT1 | ZNF692 |
| THUMPD3 | TMEM214 |
| ZNF518B | SPAST |
| SYNPO2 | RAB11FIP5 |
| UBTD2 | SH3RF3 |
| KLHL32 | SLC35F5 |
| ALG2 | POLR2D |
| ADAMTS8 | GTF3C3 |
| ERC1 | GMPPA |
| ZFYVE1 | GIGYF2 |
| CTXN2 | EPHA6 |
| MSRB1 | ACAP2 |
| TMEM231 | HS3ST1 |
| SP2-DT | CETN3 |
| CUEDC1 | SLC35A4 |
| ZNF846 | DIAPH1 |
| ZNF135 | PCDH1 |
| HAR1A | PDLIM7 |
| CACNA1I | RIPK1 |
| VKORC1L1 | MAPK13 |
| ALKBH2 | EPHA7 |
| SLC15A4 | TRAF3IP2-AS1 |
| DHDDS | PRR18 |
| PRKCD | ELFN1 |
| MFN1 | ZNF12 |
| LUZP2 | CHN2 |
| INPPL1 | PPP1R35 |
| TTC9 | THAP5 |
| JMJD8 | CHPF2 |
| USP32 | MCM4 |
| YIPF2 | NBN |
| DEDD | PLPPR1 |
| AFTPH | FKTN |
| CKMT1A | ANKRD26 |
| CPD | ZNF487 |
| MAGOH | STAMBPL1 |
| RIPOR2 | PWWP2B |
| HLA-DPB1 | ANO5 |

| | |
|---|---|
| MED21 | PRDM11 |
| UBA3 | TRPT1 |
| DLGAP2 | SYVN1 |
| TM2D2 | VWF |
| TAF9B | ZNF84 |
| PPM1G | VWA8 |
| SCAF8 | BIVM |
| LIN7A | OSGEP |
| CENPT | NGDN |
| ZNF506 | TM9SF1 |
| TBC1D20 | GPR137C |
| KAT14 | ACOT2 |
| GID8 | EIF2AK4 |
| XPR1 | PRMT7 |
| FAM241B | TXNL4B |
| HDGFL2 | ZFHX3 |
| RABEPK | SPIRE2 |
| SFXN3 | TRPV2 |
| SMARCC2 | STAT5B |
| DNAL1 | MAPT-AS1 |
| MICAL3 | TLK2 |
| ME2 | PRPSAP1 |
| FAM174C | PRELID3A |
| GGT7 | ZNF562 |
| POLR2J | ZNF254 |
| RFC1 | PCED1A |
| AP3B1 | PCNA |
| RBM7 | GZF1 |
| FAHD1 | NFS1 |
| GPN2 | LAMA5 |
| MTARC2 | DIP2A |
| NVL | ZNF74 |
| SH3BP5L | GAL3ST1 |
| RABL2A | MIEF1 |
| RNPEPL1 | TCF20 |
| PPARG | TBC1D22A |
| SEMA3F | ZNF711 |
| GTPBP8 | ENOX2 |
| PAQR9 | TM9SF3 |
| AIMP1 | C9orf78 |
| C5orf22 | TRIM2 |
| SREK1IP1 | SFXN1 |
| RASGEF1C | CHSY1 |

| | |
|---|---|
| RSPH9 | LDLR |
| TUBE1 | PQBP1 |
| PEX1 | PIK3CB |
| RBM48 | FAAP20 |
| SMURF1 | PMF1 |
| TTC39B | ZNF512 |
| RPP25L | MTHFD2 |
| DYNC2I2 | SP3 |
| FAM149B1 | CCDC50 |
| AP3M1 | ARL4A |
| CDC42EP2 | MIGA2 |
| UCHL3 | SOX6 |
| C16orf87 | FOSL1 |
| MYO19 | SCYL2 |
| SYNRG | POLR2M |
| PSPN | RORA |
| ZNF44 | TENT4B |
| THBD | PEMT |
| SOX18 | RFNG |
| THOC2 | ATG4D |
| KCMF1 | SMG9 |
| CCN2 | RAE1 |
| LAMC1 | ARFGAP3 |
| RNF166 | FRMPD4 |
| CYP7B1 | CITED1 |
| FOXJ2 | IFT57 |
| HIRIP3 | RNF145 |
| ZBTB17 | ATIC |
| CLK2 | NFKBIE |
| BACH2 | ORAI2 |
| CCDC126 | CEMIP2 |
| ZIC2 | CCAR1 |
| FRMD6 | PDHX |
| CTDP1 | TRAF7 |
| POLR3A | KARS1 |
| HBEGF | LSM14A |
| FKBP4 | G3BP1 |
| DDI2 | FBXO32 |
| LRRC4C | NOTCH1 |
| ZFP1 | TRAF3 |
| SLITRK4 | SOS1 |
| SLC66A1 | ERBB4 |
| LINC01141 | EIF2A |

| | |
|---|---|
| NT5C1A | ZNF703 |
| THBS3 | HSF1 |
| LCLAT1 | TMEM276 |
| LMAN2L | TRAPPC6B |
| ZNF197 | AP3S2 |
| C3orf62 | RNF40 |
| DNAH1 | MSI2 |
| C4orf33 | PLXNB1 |
| ATPSCKMT | ARID5B |
| BASP1-AS1 | ADIPOR1 |
| RIOK2 | CISD2 |
| FER | SNRPD3 |
| MDC1 | DPYSL4 |
| EGFL8 | STX12 |
| TSBP1-AS1 | RYBP |
| POPDC3 | CSF1R |
| DSE | MRPS33 |
| EPM2A-DT | PRKAG2 |
| RAMACL | CHURC1 |
| SMIM18 | EIF3J |
| CA13 | TELO2 |
| RSPO2 | TOMM70 |
| ZC3H3 | PCGF3 |
| CARNMT1 | TNS3 |
| WHRN | SH3PXD2A |
| KYAT1 | PCNX1 |
| CYP2E1 | BCAR1 |
| CLP1 | URI1 |
| SPINDOC | TULP4 |
| RHEBL1 | PITHD1 |
| MFSD5 | TNPO2 |
| DNAJC14 | SF3B2 |
| TMCC3 | COPA |
| SLC7A8 | CADPS |
| DPF3 | TMEFF1 |
| TDP1 | CRIM1 |
| C16orf95 | FMNL2 |
| BANP | HYOU1 |
| GTPBP3 | FAM177A1 |
| ZNF571 | ROCK2 |
| PRRG1 | COPE |
| F8 | PPP1R11 |
| RBM6 | DBNL |

| | |
|---|---|
| BCL7B | IDH3A |
| PEAK1 | NFASC |
| LIG1 | BDP1 |
| ZNF304 | VPS26B |
| COPS6 | KCNJ4 |
| IMMP2L | SH3BGRL3 |
| PRMT6 | C1orf43 |
| SLC27A3 | H2AZ2 |
| FCGR3A | PAK6 |
| ASXL2 | GON4L |
| FOXN2 | ADAMTS4 |
| NXPH2 | RETSAT |
| GULP1 | HACL1 |
| SATB2-AS1 | NMD3 |
| CHL1-AS2 | RNF168 |
| RPP14 | RAB28 |
| SCFD2 | RELL1 |
| SHROOM3 | FAM149A |
| SAP30-DT | MFAP3 |
| SHROOM1 | RNF8 |
| KLHL3 | GFRA1 |
| GM2A | ANKRD42 |
| LTV1 | CMTM5 |
| SHPRH | PLEKHH1 |
| SERAC1 | SNX29 |
| AGFG2 | TADA2A |
| GGH | CASKIN2 |
| FBXO10 | PTPN2 |
| R3HCC1L | CROCCP2 |
| P2RX4 | SHC1 |
| FAM124A | FNBP1 |
| TEX29 | TUBA1C |
| IPO4 | MCRS1 |
| TDRD9 | PXK |
| TMEM62 | FAM169A |
| PIGBOS1 | KHSRP |
| TANGO6 | CD2BP2 |
| TSEN54 | SLC25A36 |
| CCDC137 | RHOU |
| ILF3-DT | BOLA3 |
| PET117 | PSD2 |
| ZMAT1 | TMEM181 |
| NADK | NRIP3 |

| | |
|---|---|
| OSCP1 | ANKRD10 |
| HEATR5B | STX8 |
| SESTD1 | MRPS26 |
| NME5 | UBA5 |
| RNFT2 | ITGAE |
| IQGAP1 | SAE1 |
| FAM234A | CMPK1 |
| VAC14 | FADS2 |
| RPA1 | EGLN1 |
| TUBA8 | VOPP1 |
| PPP2R5A | CENPB |
| ZMYM3 | NR4A2 |
| RPL26L1 | ZFAND3 |
| HEXB | MIDN |
| ICA1 | RMDN1 |
| VWC2 | TASOR2 |
| VCPKMT | DTX1 |
| STX6 | STX2 |
| PPFIBP1 | CES2 |
| DAP3 | TGFB1 |
| CD55 | RGMB |
| PPIG | ZNF232-AS1 |
| PDCD10 | DUSP4 |
| ZFYVE28 | CXCL12 |
| RCHY1 | ATAD3A |
| ABCE1 | GPR153 |
| KIFBP | UBR4 |
| GGA2 | CSMD2 |
| RAC3 | ADPRS |
| EXOSC5 | DPH2 |
| U2AF2 | SRGAP2B |
| CFAP298 | KIAA0040 |
| FAM228B | RAB29 |
| LIX1 | PFKFB2 |
| GFPT2 | ERO1B |
| FAM167A | SOX11 |
| CCS | NOL10 |
| TMPO | KLHL29 |
| ZNRD2 | KANSL3 |
| RITA1 | SAP130 |
| SYT17 | TTC21B |
| TLCD3B | PMS1 |
| SOCS6 | TYW5 |

| | |
|---|---|
| MAGT1 | TNS1 |
| EXOSC10 | DOCK10 |
| TOPORS | SLC35A5 |
| WBP1L | DNAJC13 |
| PYGO2 | STAG1 |
| TAF13 | SLC35G2 |
| FBXO42 | IL1RAP |
| ATG7 | HES1 |
| SLIT2 | LSG1 |
| SRPK1 | SEL1L3 |
| VIRMA | STIM2 |
| MVB12B | KIT |
| COMMD3 | KLHL8 |
| PSTK | BDH2 |
| UVRAG | UGT8 |
| IFT46 | GUCY1A1 |
| HBQ1 | CBR4 |
| MPPE1 | CCDC127 |
| MTMR9 | LIFR |
| ZFPL1 | OSMR |
| CHERP | SMIM15 |
| TNS2 | SNRNP48 |
| MELTF | RNF182 |
| OSTF1 | BTN2A1 |
| CCDC7 | TRIM26 |
| ZNF768 | HCG18 |
| WFDC1 | NT5E |
| NXPH3 | REV3L |
| CYTH4 | PERP |
| PLEKHG5 | DYNLT1 |
| MAP6D1 | PLEKHA8 |
| UBE3C | KCTD7 |
| KIAA1671 | NSUN5 |
| SKIC2 | MOSPD3 |
| CCDC90B | MFHAS1 |
| KEAP1 | RGS20 |
| ARMC6 | CSPP1 |
| TMEM185A | PAG1 |
| RUSF1 | NDUFAF6 |
| MRPL4 | EXOSC4 |
| TAF11 | RCL1 |
| PPTC7 | CD274 |
| INAFM2 | CNTNAP3 |

| | |
|---|---|
| LINC00511 | FAM27C |
| CDK13 | TSTD2 |
| UBE2D4 | MSANTD3 |
| SHISA7 | APBB1IP |
| USP1 | ENTPD1 |
| YAP1 | HPS6 |
| FRA10AC1 | FNBP4 |
| PCMTD2 | LRRC10B |
| ZFAND2B | PCNX3 |
| RPUSD3 | FOLR2 |
| WDR55 | NKAPD1 |
| MAP7 | TLCD5 |
| NT5C3A | KDM5A |
| SMU1 | RPAP3 |
| EXOC6 | ASCL1 |
| PHLPP2 | GIT2 |
| GSE1 | DIS3 |
| HELZ | L3HYPDH |
| NOL4 | SLC8A3 |
| POLRMT | ZFYVE19 |
| ANKRD24 | EHD4 |
| ATP13A1 | CGNL1 |
| BAZ2A | MEGF11 |
| APPL2 | CELF6 |
| CCN3 | C15orf40 |
| KIF5B | SEMA4B |
| DAAM2 | FES |
| DMAP1 | ALDH1A3 |
| MDP1 | ORAI3 |
| RHOBTB2 | PYDC1 |
| ARL10 | FHOD1 |
| STK38L | WWOX |
| WASHC4 | CCDC43 |
| AHNAK | RPRML |
| TACC2 | TMEM100 |
| C1orf159 | MRC2 |
| SLC45A1 | CYBC1 |
| WASF2 | KDSR-DT |
| BBS7 | NR2C2AP |
| XPA | DPY19L3 |
| FERMT3 | ZNF527 |
| ARHGAP20 | PAK4 |
| SMUG1 | DYRK1B |

| | |
|---|---|
| UNC79 | ZNF780B |
| ANKFY1 | OPA3 |
| LRRC37B | ATF5 |
| EID2B | ZNF582 |
| ERCC2 | SMIM17 |
| BACH1 | TRIB3 |
| TMEM191C | SNHG11 |
| TTLL1 | FAM210B |
| TBC1D22A-DT | TRPM2 |
| MBTPS2 | CRYBB2P1 |
| HSCB | FOXRED2 |
| MON2 | LINC01278 |
| SART3 | CHRDL1 |
| KMT5A | DCX |
| PGAP1 | PCDH11Y |
| PLCL2 | USP9Y |
| NGLY1 | NENF |
| ABHD6 | TEAD1 |
| IPPK | DTWD1 |
| CERS4 | PCGF2 |
| HIRA | CNOT3 |
| PATZ1 | SEPTIN6 |
| ALS2 | WDR13 |
| NUDT16 | DOCK9 |
| SELENOO | ARHGEF7 |
| RABL2B | IRF2BP2 |
| MACO1 | SCG2 |
| PIGZ | HERC4 |
| TESC | PNKD |
| NANS | TIA1 |
| BBS5 | APPBP2 |
| CNTN4 | MOB1A |
| TRIM36 | JARID2 |
| STK17A | KCTD4 |
| CLN8 | ST6GALNAC6 |
| PMPCA | H2AX |
| C2CD5 | NFU1 |
| GATC | CHCHD5 |
| CALCOCO2 | MARCHF7 |
| ALYREF | GYG1 |
| SYNDIG1 | GOLIM4 |
| ANLN | MAP3K4-AS1 |
| LRIF1 | DIP2C |

| | |
|---|---|
| FBXO6 | CLDN10 |
| EVA1B | RYR3 |
| B3GALT2 | IL34 |
| LYPD6 | MLLT6 |
| CCRL2 | ORMDL1 |
| PLCXD2 | TOB2 |
| MASP1 | IFITM10 |
| CYP2U1 | SRSF4 |
| SMIM3 | ACSL1 |
| GEMIN5 | LRRTM2 |
| SNX16 | PDCD2 |
| NET1 | PLOD3 |
| DHTKD1 | MLC1 |
| NHLRC2 | NR1D2 |
| PARP11 | SPEN |
| PTPN6 | SNHG12 |
| LATS2 | RNF19B |
| ZBTB25 | GATAD2B |
| ARRDC4 | IGFBP2 |
| LPCAT2 | ZDHHC23 |
| TMEM107 | PIK3CA |
| PMAIP1 | TRIM7-AS2 |
| TICAM1 | NELFE |
| BCAT2 | AVL9 |
| CTU1 | MAPKAP1 |
| SMIM14 | RASGEF1A |
| CPEB2 | RB1 |
| ABTB3 | RGS6 |
| EIF1AY | TRAPPC10 |
| RFTN2 | SLC19A1 |
| SCFD1 | TMEM35A |
| ME1 | F8A1 |
| KBTBD8 | CUX1 |
| MATN2 | DNAJC16 |
| RGP1 | KPNA6 |
| CEP290 | CTPS1 |
| RCBTB2 | SFT2D2 |
| NEK9 | KCNK3 |
| ANKRD52 | PANTR1 |
| POLR3GL | SLC4A7 |
| C1orf54 | NKTR |
| SH3YL1 | ACAD9 |
| KLF15 | B3GNT5 |

| | |
|---|---|
| XRN1 | AP1AR |
| SRSF12 | CNOT8 |
| DNAJC30 | RARS2 |
| ELOA | PMS2P4 |
| PRKAB2 | FZD1 |
| NIT1 | CASD1 |
| C2orf76 | CPSF4 |
| CALCRL | CASTOR3P |
| TRAF3IP1 | SLC37A3 |
| NEK10 | LYPLA1 |
| SUCLG2 | RNF20 |
| FAIM | TMEM268 |
| NOP14 | USP20 |
| ADRA2C | ADARB2 |
| GAR1 | ILK |
| C1GALT1 | ADM |
| GTF2IP1 | NAT10 |
| TSPAN33 | MED19 |
| ZNF777 | MARCHF9 |
| LRRC61 | MAPKAPK5 |
| MCPH1 | MBIP |
| PLEKHF2 | EXOC5 |
| SMC5 | MED6 |
| NUDT5 | CD276 |
| WASHC2C | RGMA |
| CNNM2 | GFER |
| ANO4 | MOSMO |
| PRKAB1 | SRCAP |
| MIR4500HG | COQ9 |
| MEG8 | KIFC3 |
| CLBA1 | IER3IP1 |
| FRMD5 | POP4 |
| TCAP | ZNF529 |
| SLC25A19 | ETHE1 |
| SYNGR2 | UBE2G2 |
| SPACA6 | ARMCX1 |
| KCNS1 | RABGGTB |
| RWDD2B | TPP1 |
| SLC2A11 | TMED2 |
| CARHSP1 | INTS1 |
| HIVEP3 | PSMD13 |
| PCBD1 | TCEAL5 |
| ARHGAP29 | PACS2 |

| | |
|---|---|
| PACC1 | TDG |
| SERINC5 | DCUN1D3 |
| IGF2R | ATG3 |
| NOL6 | ABCB8 |
| SUPV3L1 | MOBP |
| ERLIN1 | TF |
| RASGRP2 | NOL7 |
| GABRG3 | PPP1R1B |
| CXXC1 | NUS1 |
| ZBTB45 | ADI1 |
| MYADM | SHFL |
| TFAM | SAR1B |
| AAGAB | HSPA6 |
| BBIP1 | RGS1 |
| THAP4 | CHST1 |
| FBXO45 | UBE2J2 |
| KCNIP1 | APOLD1 |
| KLHL7 | PRKAR2B |
| NYAP1 | QDPR |
| KCNH2 | SH3GL3 |
| HTR5A | PALLD |
| RBM18 | MT1X |
| NAV2 | PTBP1 |
| MED17 | GATM |
| TIGAR | GTF2F1 |
| PIP4K2C | KCNA2 |
| FGF1 | RERE |
| MATCAP2 | MAP4K4 |
| NCBP1 | MAFB |
| CENATAC | SLC1A4 |
| JKAMP | SLC39A14 |
| UFD1 | FAM120A |
| OPN3 | NAA30 |
| PRADC1 | MAL |
| MARCHF4 | WNK2 |
| TCTA | TBCK |
| RASSF1 | EXOC2 |
| ATP11B | MIR9-3HG |
| DELE1 | RBM38 |
| GRM1 | SOX2 |
| DNAJC25 | ALDH1A1 |
| PPP1R3E | GAPVD1 |
| RCOR1 | STAT2 |

| | |
|---|---|
| ANKS3 | SFSWAP |
| XYLT1 | GPR183 |
| HSDL1 | SLC43A2 |
| KSR1 | PRELID3B |
| SPPL2B | KLHL17 |
| NELFCD | SNIP1 |
| ALG13 | SCMH1 |
| TMEM134 | TUT4 |
| RALGAPB | HOOK1 |
| CLOCK | LRRC40 |
| ID3 | LRRC8C-DT |
| SLC27A1 | DPH5 |
| ZFP82 | NTNG1 |
| ITM2A | WNT2B |
| CTH | FCGR1BP |
| OTUD7B | NOTCH2NLA |
| WDFY3-AS2 | RAB13 |
| KANSL1-AS1 | ATP8B2 |
| CAVIN2 | THBS3-AS1 |
| IFI44 | METTL25B |
| LINC01750 | MIR29B2CHG |
| LINC01138 | ZNF496 |
| NUDT4B | E2F6 |
| ARNT | GREB1 |
| RAB7B | KCNS3 |
| SLC11A1 | IFT172 |
| ERCC8 | ELMOD3 |
| MED7 | FRZB |
| MRS2 | NABP1 |
| ATF6B | KCTD18 |
| B3GAT2 | SMARCAL1 |
| TNFAIP3 | STK36 |
| PLAGL1 | SP110 |
| DLL1 | COLQ |
| SCIN | ITGA9-AS1 |
| AHR | ENTPD3 |
| SNX10-AS1 | TMIE |
| PAXIP1 | SLC25A20 |
| CA8 | AMT |
| STK3 | RNF123 |
| PARP10 | GMPPB |
| KDM4C | STIMATE |
| RHOG | NEPRO |

| | |
|---|---|
| GSTZ1 | MYLK |
| SARM1 | CCDC14 |
| TMEM235 | DBR1 |
| OGFOD3 | ALG3 |
| ZNF585A | ZNF595 |
| GLRA2 | MSANTD1 |
| ZFY | ADGRA3 |
| C11orf68 | CENPC |
| MRPS18C | UNC5C |
| MDN1 | MANBA |
| RORB | ELF2 |
| TSC1 | INPP4B |
| MAN1B1 | LRBA |
| ACYP1 | GALNT7 |
| CFDP1 | SLC9A3-AS1 |
| NCLN | ZDHHC11B |
| THBS1 | ZNF131 |
| GMEB1 | ITGA2 |
| GRIK3 | UTP15 |
| LINC02609 | ZCCHC10 |
| NOS1AP | PCDHB14 |
| QPCT | SPRY4 |
| MCEE | PDGFRB |
| GCFC2 | TRIM52 |
| STK11IP | KDM1B |
| FILIP1L | TRIM39 |
| HERC6 | PRR3 |
| NDST3 | CCHCR1 |
| WDR70 | TENT5A |
| CRHBP | COQ3 |
| PCSK1 | NR2E1 |
| CEP120 | CEP85L |
| PWWP2A | FBXO5 |
| LINC00680 | MRM2 |
| MANEA | RBAK |
| DCBLD1 | FAM221A |
| HSF2 | BMPER |
| THBS2 | PSPH |
| ICA1-AS1 | RCC1L |
| AASS | NSUN5P1 |
| SQLE | PHTF2 |
| TEX10 | ABCB1 |
| LHX2-AS1 | ZNF3 |

| | |
|---|---|
| PDCD11 | TRIP6 |
| KCNQ1OT1 | RINT1 |
| TALAM1 | METTL2B |
| PUS7L | ZC3HC1 |
| KBTBD7 | NOM1 |
| COL4A1 | POLR3D |
| ADPRHL1 | IKBKB |
| WDR24 | CRH |
| CRAMP1 | TRPS1 |
| ZNF75A | EXT1 |
| MBTPS1-DT | TIGD5 |
| LIG3 | ZNF252P |
| HSD17B1-AS1 | WASHC1 |
| ZNF653 | FRMD3 |
| ZNF584 | TRIM14 |
| RBBP9 | TLR4 |
| LINC00320 | FPGS |
| ZCCHC18 | PHYH |
| PWWP3B | MSANTD7 |
| IL13RA1 | RET |
| RASSF8 | AGAP4 |
| SLC25A1 | TMEM273 |
| KLHL35 | KAT6B |
| SLCO3A1 | SHLD2 |
| PPAN | NOC3L |
| NSL1 | VTI1A |
| AIMP2 | C10orf88 |
| HAPLN4 | GATD1-DT |
| ALPL | PIDD1 |
| NR3C2 | SPTY2D1 |
| PSMB8 | GAL3ST3 |
| SLC39A4 | DPP3 |
| ZNF384 | SSH3 |
| ACADS | AAMDC |
| SNAP23 | GAB2 |
| EIF3C | PRSS23 |
| ZNF48 | CWF19L2 |
| NSRP1 | ZW10 |
| GOSR1 | MCAM |
| ZNF536 | NCAPD3 |
| CLDND2 | LTBR |
| PIKFYVE | LPCAT3 |
| CNTN3 | SLCO1A2 |

| | |
|---|---|
| COG1 | CNPY2-AS1 |
| TRMT1 | METTL1 |
| FAM117B | DPY19L2 |
| MRPL13 | SLC8B1 |
| CHST15 | DDX55 |
| RELT | ZNF268 |
| C1S | ZMYM5 |
| ENTREP3 | GPR12 |
| B4GALT3 | SUPT20H |
| ADORA1 | MRPS31 |
| PIGF | NAA16 |
| DUSP28 | OBI1 |
| DCP1A | HAUS4 |
| FAM193A | ZFHX2 |
| XPO5 | REC8 |
| RNGTT | NPAS3 |
| MTRES1 | BAZ1A |
| PPP1R14C | GCH1 |
| CREB5 | TMEM260 |
| HERPUD2 | CPSF2 |
| RIPK2 | HHIPL1 |
| UPF2 | CHAC1 |
| ZNF22 | CSNK1G1 |
| USP54 | DIS3L |
| PPIF | THAP10 |
| ALG5 | PTPN9 |
| IFT43 | EFL1 |
| WDR73 | LRRC28 |
| ARHGAP17 | IL32 |
| PHF20 | CLUAP1 |
| LIME1 | ITPRIPL2 |
| GRAMD4 | KNOP1 |
| SURF6 | SGF29 |
| LSM11 | CFAP119 |
| CDK11A | ZNF668 |
| PPIH | CPNE2 |
| NDC1 | SETD6 |
| GPR89B | HSF4 |
| THEM4 | SLC9A5 |
| GRHL1 | ST3GAL2 |
| SMC6 | OSGIN1 |
| UBXN2A | CTU2 |
| LINC01122 | TLCD3A |

| | |
|---|---|
| CFAP210 | PITPNA-AS1 |
| RBM45 | ALOX12-AS1 |
| CNOT10 | TP53 |
| ACY1 | ATPAF2 |
| RIOX2 | RFFL |
| CEP70 | AATF |
| XXYLT1 | DHX58 |
| OSTC | RND2 |
| DHFR | FAM117A |
| WDR36 | SMURF2 |
| SAYSD1 | ABCA8 |
| UST | KCNJ16 |
| BET1 | SLC39A11 |
| ZNF862 | SGSH |
| ASIC3 | RNF213 |
| SLC4A2 | THOC1 |
| TMEM67 | C18orf21 |
| C9orf40 | SERPINB8 |
| ZBTB6 | TSHZ1 |
| MIR600HG | TCF3 |
| RPP38 | ZNF554 |
| ENKUR | TLE2 |
| FGFBP3 | ARHGEF18 |
| HMBS | ZNF561 |
| PTHLH | TYK2 |
| RASSF3 | HOOK2 |
| FGD6 | MRI1 |
| RASL11A | MIR23AHG |
| L2HGDH | HOMER3 |
| MAP3K9-DT | TMEM161A |
| RFX7 | MAU2 |
| ZSCAN32 | RHPN2 |
| EARS2 | ZNF181 |
| PDP2 | HAMP |
| DDX19B | SIPA1L3 |
| WRAP53 | POU2F2 |
| CTC1 | ZC3H4 |
| ADORA2B | BICRA |
| LLGL1 | ZNF71 |
| TOP3A | ZNF264 |
| PGAP3 | ZNF530 |
| MEX3D | ZNF814 |
| CHAF1A | ZCCHC3 |

| | |
|---|---|
| MAP2K7 | SNAP25-AS1 |
| P2RY11 | CTNNBL1 |
| HSPB6 | RPRD1B |
| ZNF471 | FITM2 |
| RALGAPA2 | LINC01260 |
| ZBTB46 | DOK5 |
| RNF215 | MIR155HG |
| ZXDB | GABPA |
| RNF113A | HUNK |
| ITGA6 | URB1 |
| DDAH2 | HLCS |
| RGS10 | MX2 |
| CUL5 | TOP3B |
| ING5 | SMTN |
| RASL10B | APOL6 |
| BCL2L13 | HCCS |
| NOTCH4 | ABCB7 |
| LACTB2 | TRPC5 |
| TMEM52B | ARHGEF6 |
| LRP6 | MCF2 |
| ZCCHC8 | BGN |
| PLA2G2C | CHMP7 |
| BCL10 | E2F4 |
| MINDY1 | MTR |
| ABCB10 | KLF3 |
| SUMF1 | FAM193B |
| FHIT | FUCA2 |
| CD86 | EEPD1 |
| HPS3 | PSMD5 |
| CXCL3 | KDM2B |
| ABCG2 | FAM161B |
| SLC39A8 | PML |
| LY86 | E4F1 |
| AARS2 | CDH20 |
| STX11 | LIPE-AS1 |
| NKX3-1 | RGN |
| LRRCC1 | DCN |
| DEPTOR | ACAA2 |
| HGH1 | PUSL1 |
| CEP78 | CLCN6 |
| NAA35 | CCDC28B |
| GLE1 | UTP11 |
| ZEB1-AS1 | RPF1 |

| | |
|---|---|
| DERA | FNBP1L |
| ANGEL1 | SLC30A7 |
| SNX33 | PBXIP1 |
| NUBP1 | NCSTN |
| ASPA | UCK2 |
| TMEM98 | FLVCR1 |
| CDR2L | FBXO28 |
| TMC6 | SLC35F3 |
| KATNAL2 | MAP4K3-DT |
| RTTN | SOCS5 |
| ZNF577 | EML6 |
| ZNF582-DT | LGALSL |
| PLAGL2 | TGFA |
| DGUOK | MITD1 |
| GGA1 | TGFBRAP1 |
| MATCAP1 | IL1RN |
| GCNT4 | TMEM163 |
| GPR22 | CD302 |
| THAP12 | STRADB |
| RAVER2 | LRRN1 |
| WRAP73 | JAGN1 |
| LINC00339 | ELP6 |
| EPB41 | NCK1 |
| TIE1 | PARL |
| TOE1 | BMP2K |
| NSUN4 | SLC12A7 |
| AGBL4 | ANKRD33B |
| ATG4C | NIM1K |
| PRUNE1 | COMMD10 |
| PIGC | PTTG1 |
| GALNT15 | C6orf47 |
| ATG10 | HLA-DMA |
| MAN2A1-DT | RGL2 |
| P4HA2 | SLC35B2 |
| PCYOX1L | MOXD1 |
| GTF2H4 | TBRG4 |
| LINC00951 | KMT2E-AS1 |
| KHDRBS2 | COG5 |
| STXBP5-AS1 | GCC1 |
| WDR27 | RBM28 |
| GET4 | PODXL |
| GTF2IRD2B | KIAA1549 |
| STAR | PAXIP1-DT |

| | |
|---|---|
| CRISPLD1 | ZMAT4 |
| ZNF34 | KAT6A |
| CENPP | TOX |
| ANAPC2 | ARFGEF1 |
| ASAH2B | TMEM64 |
| AIFM2 | ZHX2 |
| FRAT2 | GLDC |
| CYB5R2 | ENG |
| BDNF-AS | PPP1R26 |
| NAA40 | SVIL-AS1 |
| RAD9A | CHST3 |
| NEU3 | AVPI1 |
| NAALAD2 | SORCS3 |
| USP2-AS1 | SFXN4 |
| MSANTD2 | SPI1 |
| P3H3 | AP5B1 |
| PRANCR | RPS6KB2 |
| TEDC1 | FAM181B |
| GALK2 | DYNC2H1 |
| TBC1D2B | BCL9L |
| ALDH1A3-AS1 | LOH12CR2 |
| RAB37 | ATF7IP |
| TYMSOS | LRRK2 |
| CEP76 | GPD1 |
| ZNF98 | GNS |
| ZNF8-DT | CRY1 |
| ZNF8 | PSMD9 |
| GK | LRP10 |
| RP2 | AP1G2 |
| CCDC120 | DACT1 |
| CCDC22 | PIGH |
| ABCD1 | MOK |
| GLTP | NUP93 |
| COMMD8 | CIAPIN1 |
| NUP42 | RILP |
| TRAF4 | NUP88 |
| CAPG | SMCR8 |
| RPRM | SNHG30 |
| NEK1 | TBKBP1 |
| TRERF1 | FTSJ3 |
| HMGCLL1 | MAP2K6 |
| FAM118B | SMCHD1 |
| LMBR1L | ZNF521 |

| | |
|---|---|
| PELI2 | ZNF516 |
| ARMH4 | CC2D1A |
| DCAKD | RFX1 |
| ENGASE | KLHL26 |
| ZNF329 | LRFN3 |
| BCL2A1 | FPR1 |
| IPO11 | PIGU |
| CHRNB2 | TTI1 |
| KHDC4 | TNFRSF6B |
| DUSP23 | USP16 |
| SLC19A2 | PAXBP1 |
| FAM110C | PPM1F |
| DNMT3A | MN1 |
| CYP1B1 | CHADL |
| PEX13 | POLDIP3 |
| CIR1 | TUBGCP6 |
| DALRD3 | BCOR |
| USP13 | MAOA |
| CDKL2 | TAF1 |
| ARHGAP10 | CHIC1 |
| MRPS30-DT | TIMM8A |
| PGGT1B | HMGB3 |
| STYXL1 | CALB2 |
| APTR | BRD4 |
| ACHE | TUB |
| PDIA4 | CLASP1 |
| WWP1 | BUD23 |
| LRP12 | LUZP1 |
| NIPSNAP3B | GNL3 |
| BMPR1A | CDS1 |
| ADRB1 | GNA12 |
| KBTBD4 | EBPL |
| PCSK7 | TRMT61A |
| TUBGCP3 | WIZ |
| KLHL28 | SLC9A7 |
| PIGQ | IGBP1 |
| MPV17L | NQO2 |
| PLXDC1 | CYB5R1 |
| HAP1 | LIMS1 |
| MEX3C | HMGB2 |
| MALT1 | WDR46 |
| PEDS1 | NKAIN3 |
| ADSL | EED |

| | |
|---|---|
| SHROOM2 | UBASH3B |
| FAM120C | ING1 |
| DIAPH2 | NR1D1 |
| LRCH2 | PIK3IP1 |
| AKT1S1 | CYP26B1 |
| KRR1 | SLC39A1 |
| PREB | MYNN |
| RNF175 | TFPI2 |
| ITSN2 | VSTM2A |
| DCK | FASTK |
| DIMT1 | HMOX1 |
| TMEM196 | VAMP5 |
| POU6F1 | FARP2 |
| MGAT2 | ADGRV1 |
| ANKRD27 | CALD1 |
| METTL13 | UHRF2 |
| VARS2 | WDFY2 |
| TCF7L2 | PHAF1 |
| SLC35E3 | NT5C |
| ERAL1 | BMP7 |
| TXNIP | |
| ADAMTS3 | |
| SH3D19 | |
| CPNE8-AS1 | |
| CDKN2C | |
| GPR88 | |
| GPR89A | |
| ZNF672 | |
| SLC35F6 | |
| POLR1B | |
| CWC22 | |
| MAIP1 | |
| INO80D | |
| LRRC2 | |
| IL17RB | |
| NXPE3 | |
| SMARCAD1 | |
| INTU | |
| ARHGEF28 | |
| GFM2 | |
| MCC | |
| RAPGEF6 | |
| TFEB | |

NT5DC1
RNF217
NIPAL2
NAPRT
PIGO
NPR2
PLPP7
ZNF33A
MCU
TBC1D12
TMEM80
INTS5
RCE1
ATG16L2
EMSY
FDX1
TDRD3
ADCK1
RPUSD2
MTHFS
UBN1
MIR762HG
SPATA33
MED9
C17orf58
RMC1
EPG5
XRCC1
BLOC1S3
PRR12
MZF1
NFATC2
PKNOX1
CCDC157
MTFP1
ZBED1
NDP
SLC10A3
PTRH1
HBB
WIPF1
ZNF445
KCTD8

DENND4C
PHYHD1
APH1B
HCST
SESN1
TSEN34
AMOTL1
TRIM32
COMTD1
TCTN3
VPS33A
GTF2H3
CBFA2T3
PCBP1-AS1
CXXC4
HLA-DQB1
CUTC
CPM
ZNF407
MIR4435-2HG
GPRC5A
TMEM234
S100PBP
DRAM2
POLR3C
USP21
ESRRG
PKDCC
TMEM17
POLR1A
LINC00342
BCL2L11
ZC3H8
NIFK-AS1
LINC02993
WDSUB1
PDK1
TFPI
EDEM1
TAMM41
PLCD1
EXOG
CACNA1D

SLC9A9
COMMD2
COX18
STBD1
GMDS
ZNF318
ASCC3
BBS9
HECW1
CROT
SH2B2
SMKR1
WDR91
NOL8
ZNF169
BMS1
IPMK
IDE
POLL
USP35
KBTBD3
C11orf71
PYROXD1
ABCC9
TMEM117
SLC16A7
NR2C1
TCHP
GPR180
KHNYN
DTD2
ZFYVE26
GOLGA5
PPIP5K1
SPG11
FBXO22
VPS33B
HS3ST4
XAF1
FAM222B
PRCD
FN3K
ZNF519

ANKRD29
FECH
GCDH
MED26
ZNF493
ZNF766
ITCH
MYH7B
MHENCR
SAMD10
SLC37A1
GGT5
ASCC2
CXorf38
MIR222HG
GPR173
RNF128
RAP2C
CACTIN
CIAO2A
LRRC8D
DBF4
WASHC2A
UNC45A
ASNSD1
PPARGC1A
TMEM60
UBTD1
OPALIN
PRDM16
C1orf174
AGTRAP
PHACTR4
SZT2
SLC6A9
TESK2
GBP2
CDC7
ECM1
CRABP2
TRMT1L
TGFB2
KRCC1

| |
|---|
| TBC1D8 |
| GPR17 |
| CYP27A1 |
| SCLY |
| IL17RC |
| SUSD5 |
| HYAL3 |
| DUBR |
| SEC22A |
| CEP63 |
| PPP2R3A |
| PLCH1 |
| PLAAT1 |
| NOP14-AS1 |
| EXOSC9 |
| SCLT1 |
| FASTKD3 |
| SETD9 |
| CCNB1 |
| ZBED3 |
| CDKL3 |
| MAML1 |
| E2F3 |
| GPLD1 |
| ABCC10 |
| COL12A1 |
| BCKDHB |
| FBXL4 |
| PDSS2 |
| FIGNL1 |
| MBLAC1 |
| MUC12 |
| COPG2 |
| FAM131B-AS1 |
| WDR86 |
| PEBP4 |
| ANK1 |
| MTERF3 |
| LINC01503 |
| ENTR1 |
| IDI2-AS1 |
| TAF3 |
| CELF2-DT |

| |
|---|
| AGAP6 |
| FLRT1 |
| XRRA1 |
| EMSY-DT |
| AQP11 |
| GPR83 |
| ITFG2 |
| RESF1 |
| FKBP11 |
| RACGAP1 |
| MSRB3 |
| BBS10 |
| PAWR |
| LRCH1 |
| DHRS12 |
| MRPS31P5 |
| CDKL1 |
| BBOF1 |
| FAM181A |
| CLMN |
| TM6SF1 |
| FCSK |
| GEMIN4 |
| SMG6 |
| TMEM101 |
| FMNL1-AS1 |
| TOM1L1 |
| SMARCD2 |
| ITGB4 |
| IZUMO4 |
| AMH |
| DNASE2 |
| PDE4C |
| FBXO17 |
| ZNF575 |
| LYPD5 |
| ZNF528 |
| AP5S1 |
| KIF16B |
| ACTR5 |
| ZNF217 |
| LKAAEAR1 |
| C21orf91 |

LINC01637
PABPC1L2B
KLHL4
C1GALT1C1
OTUD3
FKBP14
TMEM69

**ΓBS and 10 Hz rTMS within cortical layer 1, and 2/3 across all donors.**

| Donor 2 | | | | | |
|---|---|---|---|---|---|
| Cortical layer 1 | | | Cortical layer 2/3 | | |
| iTBS | iTBS ∩ 10 Hz | 10 Hz | iTBS | iTBS ∩ 10 Hz | 10 Hz |
| MT-ATP8 | VGF | HSPA1A | MT-ATP8 | CSKMT | CCL2 |
| UCHL1 | PHYHIP | EIF5 | MBP | TAFA1 | CCL4 |
| NRGN | THY1 | EEF1B2 | PLP1 | ATF7IP2 | CCL3 |
| APLP1 | FAM181B | SNHG14 | MALAT1 | FAM98C | CCL3L3 |
| BAIAP2 | SYN2 | NCL | HPCAL1 | IL1RN | TSC22D4 |
| TAGLN3 | CAMK2B | CDC42 | INHBA | CALB1 | CCL4L2 |
| SNCB | FKBP15 | C1orf122 | EGR2 | BATF3 | PPP1R14A |
| ARPP19 | YTHDF2 | MT1M | KCNIP2 | SDR39U1 | CRYM |
| MAP1B | CTSH | S100A13 | ATF3 | USP24 | NR4A3 |
| ATP1A3 | PI4KA | DDAH1 | ZFR | CTSZ | IGSF21 |
| ATP2B1 | STAT3 | NAMPT | CALB2 | CORO1A | THEMIS |
| ENO2 | PSMA1 | EIF3A | SOWAHA | CSF1R | ANKRD37 |
| NCDN | BAZ1A | NTRK2 | RASGRF2 | PIR | TP53INP2 |
| EGR1 | IRS1 | MEF2C | PCDH8 | PBRM1 | AQP1 |
| PRORP | LST1 | IFITM3 | DUSP2 | SPRY1 | SYTL2 |
| MTCH1 | GABRA1 | EDF1 | CALR | TF | SEMA3B |
| SIL1 | ATL1 | NDUFC2 | TRIB1 | ABCB1 | GAD2 |
| SNX22 | UPF3A | CD99 | CDC37L1 | OBSCN | DCXR |
| ARC | NPY | C11orf96 | HS3ST2 | GJA4 | CHCHD1 |
| PFDN2 | GRIN1 | NOP58 | LAMP5 | NCK1 | MRPL19 |
| EGR4 | THAP2 | HSPD1 | SLC12A8 | SEZ6 | ANXA1 |
| SYN1 | SRRD | FRRS1L | CHCHD6 | ICAM2 | TUFM |
| PLD3 | PLEKHB2 | EPS15 | POLR2J | H1-0 | EFCAB14 |
| FUS | BMERB1 | DCTN2 | RGS12 | EBPL | RNF38 |
| ID3 | ATP6V1A | MLLT11 | CNR1 | HS6ST3 | CLK4 |
| SERPINI1 | ATP6V0B | RPL36AL | MIA3 | TATDN3 | CCSAP |
| KIAA0930 | ACTR10 | ABI2 | DUSP6 | ZC3H18 | SPARC |
| PPP3CA | CELSR2 | MORF4L1 | MKNK2 | IGFBP5 | NTSR2 |
| LMO3 | CASP4 | CCNI | FNDC9 | TMEM144 | TMEM147 |
| SNCA | SLC14A1 | CALY | GNG2 | UIMC1 | ABRACL |
| WBP2 | JPT1 | ATP1A2 | CRLF1 | THEMIS2 | FSTL1 |
| CEBPB | CAMKV | PCDH9 | NUPR1 | CTNS | GALNT9 |
| TAF4B | NECAB1 | GSTP1 | NPY | RNF128 | CEP78 |
| MEX3D | ARHGDIG | PTPN11 | FAM89A | RPP25L | CCN2 |
| MAPRE3 | BLOC1S1 | MRPL41 | RPL7L1 | LSG1 | EID2 |
| NSF | STAG2 | EIF3H | ERRFI1 | NR2F1 | PVALB |
| C3orf70 | KPNB1 | DNAJC6 | C1QL2 | SUPT20H | ARX |
| TMEM120B | REV3L | UGP2 | RSRC2 | METTL3 | GEMIN5 |

| | | | | | |
|---|---|---|---|---|---|
| STYX | PRKAG2-AS1 | CAPRIN1 | RBBP7 | P2RY11 | FIBIN |
| PRMT2 | LYN | NR1D2 | EIF3M | PCBP1-AS1 | PCP4 |
| ISOC1 | KIAA1143 | ALDH6A1 | MAN1C1 | TMEM179B | CD164 |
| MYO16 | HERPUD1 | PNRC1 | SNHG12 | LNCBRM | ABCD3 |
| SEPTIN7 | SLC25A25 | DDR1 | DAGLB | IGSF3 | NMB |
| NDUFB6 | ARL6IP5 | SCAND1 | TXNL4B | SUMF1 | THRB |
| ARF1 | SRCIN1 | NDRG3 | PARP6 | MIR23AHG | DCAKD |
| CCT4 | TGFBR3L | HNRNPA3 | GSG1L | ZSCAN5A | POLR3H |
| ACAP3 | PPDPF | SSBP1 | SLC9A7 | ZNF141 | KCNN3 |
| BUD31 | KCNAB2 | PREPL | PIN4 | TSPAN17 | TP53RK |
| RAP1GAP | PITPNA | CUTA | MEST | SP4 | HBP1 |
| PTPRN | DUSP2 | SPCS1 | CYB561 | RNF169 | IGFBP2 |
| GPRC5B | VAMP5 | NAPB | GALC | RNF122 | RASD2 |
| PTN | PER1 | PPP2R1A | SRP68 | CARNS1 | MISP3 |
| ATP5PO | ZNRF3 | PPFIA1 | PHACTR3 | TMEM233 | CRYL1 |
| PRDX2 | SMAP2 | EXOC5 | PSMD13 | BCL7A | NSMAF |
| CXCL14 | MARCKS | SNPH | RPSA2 | CFLAR | FAM229B |
| CYFIP2 | CD24 | FYN | MMUT | PCCA | TCTA |
| P4HTM | SHLD2 | AQP1 | SLC11A2 | RCN1 | NKAIN4 |
| WASF1 | MRPL11 | MPDZ | CD200 | CLDN11 | NKX6-2 |
| LYRM1 | AMPH | THRA | UNC5B | PLXDC1 | TMEM245 |
| LEPROT | RAB11B | MAP2K1 | KSR2 | SIDT2 | DUSP7 |
| SEC22C | GABRG1 | PCGF5 | KCND3 | THOC3 | MCUB |
| NPTX2 | FAM171B | CAMK2A | PTPRR | FAM151B-DT | POP5 |
| UBE2S | PPP2R2C | ARID5B | ZNF184 | SETX | PTPRT |
| KLC1 | ZNF496 | MRPS24 | ORC4 | XKR4 | IFIT5 |
| MYBPC1 | HYAL2 | KLF10 | DUSP26 | ENTREP3 | ADGRB3 |
| MAP3K8 | COX10 | SORL1 | LBH | SH3RF1 | SORCS2 |
| EBPL | ZNF626 | SPIRE1 | ATAT1 | RAPGEF6 | SLC35F5 |
| VXN | DCTN1 | MAN1A2 | SNHG15 | CRHR1 | CYP2U1-AS1 |
| ZBTB47 | SYP | TCF25 | NCLN | COLQ | INTS2 |
| UBE2B | CTIF | MRPS21 | NCOA1 | PLVAP | C3 |
| TMEM59L | MYT1L | SLC25A5 | NTRK3 | EIF2B4 | RTF2 |
| COQ4 | UHMK1 | NDUFS6 | C12orf43 | MRPL36 | PRPF39 |
| CPNE6 | NPTX1 | DDX21 | RIPK1 | EXOSC7 | COMT |
| SLC25A22 | PDP1 | ADD1 | ITPKC | CHMP6 | IFI44 |
| BRD2 | WIPF3 | MMP24OS | CMIP | SYN3 | EGFL7 |
| DIRAS2 | ANXA4 | CAPNS1 | RSPH9 | TBC1D2B | TMEM165 |
| ANKS1B | ZNF354A | IER5 | DCAF5 | L3MBTL1 | GCSH |
| SCG5 | ITIH5 | POLR2A | CMTM4 | RWDD2A | ZNF576 |
| PIP5K1C | ORMDL2 | CLASP2 | EIF3E | ATG16L2 | NCOR1 |
| GTF2F2 | DOP1B | C6orf62 | ATPSCKMT | KCNJ16 | ZNF540 |
| BAG5 | SOX2 | MGLL | NFATC2 | QPRT | TMEM123 |

| | | | | | |
|---|---|---|---|---|---|
| ARPC1B | PRKCB | TPPP | ITGAV | TADA2A | FAM107B |
| FLCN | PLEKHM2 | HDLBP | PPP1R15B | SLC7A2 | ZNF789 |
| ANKH | PAK1 | TAX1BP1 | ABLIM1 | TNFRSF12A | MRPL10 |
| SPRYD3 | TMEM130 | SNX6 | GPR88 | ACAD11 | SNHG17 |
| ARHGAP31 | STRBP | FCER1G | ZNF697 | CCHCR1 | SPTBN4 |
| APP | MAGED2 | PTCD3 | NUTF2 | PTGR2 | ADARB1 |
| CAMKK1 | KLF9 | CCDC6 | CNN3-DT | DONSON | USP28 |
| GRAMD1C | ATP13A2 | KDSR | SIPA1L3 | GJB1 | PAPSS1 |
| ZFPM1 | CTXND1 | PANK2 | LRATD1 | OPALIN | PTTG1 |
| JOSD1 | PSMA6 | LRRC47 | HBB | SLC24A3 | FKBP9 |
| STMN2 | SUMO3 | EIF4G3 | UBE2D1 | ARMCX5 | BBS5 |
| ATP6AP1 | ARPC5L | SSR1 | DGKB | ZNF254 | ITGA5 |
| ARID5A | SIPA1L1 | TM2D2 | GJB6 | HLA-C | MTR |
| STPG1 | PNISR | LRRC59 | CD53 | KANSL3 | TET3 |
| CYP27A1 | CXCL2 | CCNY | SDHAF4 | CWF19L1 | USF3 |
| SHROOM1 | YIF1A | CELF4 | SNRNP25 | ATP11A | FAR1 |
| LY86 | MAPK9 | ACTR3 | SGMS2 | TMC6 | ACACA |
| GPLD1 | FNDC3A | HADHB | SSR1 | MVB12B | CASKIN1 |
| ZNF92 | CSF1 | DNER | LGI1 | MAP2K5 | TRAPPC3 |
| RRP8 | OCIAD1 | CALR | MIB1 | CNIH3 | TPBG |
| PCNX3 | BTBD8 | ARHGEF12 | RASA3 | FAXC | LINC01122 |
| CDAN1 | ADGRB2 | MORF4L2 | CYTH3 | CD302 | SLC12A7 |
| FBN1 | RAB26 | SSU72 | KIF1B | RGL2 | ALDH7A1 |
| WDR81 | FGF13 | MPC1 | HTT | UXS1 | RHOG |
| IGLON5 | ING1 | NCKAP1 | OTUD5 | MLXIPL | PDZD2 |
| TWF1 | OS9 | PGRMC1 | FOXO3 | TYW1B | RNFT2 |
| KCNK1 | UBE2QL1 | AFF4 | AIFM2 | GNG13 | EFNA5 |
| PRKCG | HCFC1R1 | PPP2CB | USP33 | SST | CYBRD1 |
| CTTN | PCNP | KMT2E | ANAPC1 | MCHR1 | LMTK2 |
| HS3ST2 | UBE2D2 | CREBBP | ENDOV | ARHGDIB | PGAP1 |
| BCAS1 | SMIM29 | RAD23A | DIP2C | XAF1 | ZCCHC24 |
| UBTF | UQCRC2 | FNBP1 | CRYZL1 | EBF4 | ALKBH2 |
| TRIB1 | RUNDC3A | RHOQ | SYT16 | PCDH1 | LPAR6 |
| SPINT2 | SYT5 | LINGO1 | ZSCAN16-AS1 | TAF8 | LINC03013 |
| EIF3F | KIF3A | GARS1 | AMFR | RNASEH2B | SPATA20 |
| MYL6B | PFKFB3 | SERPINE2 | HEPACAM | RPS6KA4 | SPRN |
| WDFY3 | GRIA3 | PRKCA | ADCYAP1R1 | PTP4A3 | ELP6 |
| CD14 | SLC23A2 | NDUFB8 | NFAT5 | MPST | BRD7 |
| CDKL3 | IFIT2 | SEPTIN2 | SSR2 | SCAMP2 | TBC1D22A-DT |
| TNFRSF10D | HPCA | EIF4B | CDC42EP3 | THUMPD3-AS1 | FBXL2 |
| PAOX | IFIT5 | ATP6V0A1 | HERC1 | DIPK1C | NAB2 |
| DDX47 | ACSL5 | ARHGDIA | SLC27A1 | STOX1 | WDR36 |
| MMP19 | TMEM201 | MLF2 | LRRC73 | TFPI | IL33 |

| | | | | | |
|---|---|---|---|---|---|
| TMEM229B | SSR2 | STMN4 | WFDC2 | CNEP1R1 | ZNF232-AS1 |
| TOP3A | GABBR1 | NDUFB4 | PPM1K | TNS2 | CCR1 |
| CD300A | SELENOM | RAB7A | RAB40B | CNTN3 | ANP32B |
| COLEC12 | MFSD2A | GRINA | PPFIA2 | FAM43B | ACVR1B |
| ZNF527 | NUDT4 | LY6H | CASC15 | CDC7 | POLR1B |
| ZNF628 | LARP1 | PFN1 | RRAS2 | PACC1 | FBXO38 |
| PCMT1 | PSMC5 | ZYX | TP53I11 | CDK11A | GALNT17 |
| TMEM100 | SPNS2 | RAPGEF4 | NCAM2 | MREG | DPH1 |
| ZNF483 | IFT20 | XRCC5 | LSM1 | MIR4500HG | CERS2 |
| DCLK1 | NETO2 | ATP6V1C1 | CBLN2 | ZNF227 | CEROX1 |
| BSN | RAB40C | WTAP | TESPA1 | CAPN3 | UBE4A |
| KIF3C | RPL7L1 | PRR13 | COBL | H2BC21 | TNFAIP1 |
| VCP | ELP1 | TRIM9 | SERPINE1 | HS1BP3 | GPR137B |
| DHPS | STX1A | MSX1 | SOX2 | TBC1D8 | TNFRSF21 |
| OTUB1 | NKAIN4 | AHNAK | GRK5 | SLC8A3 | CDC5L |
| PSMC2 | SIN3B | KDM6B | TNNT1 | DNAJC16 | SERTM1 |
| GCLM | ACTN2 | ALDH1A1 | DZIP1 | AGPS | BCKDHB |
| SNORC | YBX3 | SLC2A5 | MAP2K7 | WWC1 | NDUFAF1 |
| DUSP1 | PPP2R3C | TGFB1 | TBC1D30 | MDGA1 | KCNAB1 |
| HARS1 | CXXC5 | SDCBP | TMEM35A | TAF9B | CWC25 |
| POLR2M | ARHGEF9 | CSNK2B | CEP43 | ACVR2A | MAPRE1 |
| HAGH | GSTM5 | LAMTOR2 | MBNL1 | ALG2 | RNF185 |
| BCAT1 | PLPP3 | TIMM10B | MGAT3 | FAM110C | ARHGAP33 |
| RCN2 | CCDC32 | RAB15 | SOCS4 | GPAM | FHIP2B |
| CHST1 | IFNGR2 | AP1G1 | RGS8 | VWA1 | CMTM5 |
| LAT2 | SLC66A2 | PTPN2 | SPON2 | ZNRF3 | PANK2 |
| ERP44 | PFDN4 | ADNP | CDH9 | SNX2 | USP54 |
| SNRNP70 | TBC1D5 | PQBP1 | ZNF251 | MERTK | LDB3 |
| PDE4B | SPATA2L | DNAJC19 | MLYCD | ADCYAP1 | MFSD3 |
| POGK | FLRT2 | JPH4 | PPM1A | KMT2E-AS1 | ZBTB21 |
| MAN1C1 | PRDM8 | ZNF580 | CHKA | DCUN1D3 | MYBPHL |
| PJA2 | ZC3HAV1 | APOLD1 | NDUFAF4 | SNX29 | TTC21B |
| SORT1 | RNF216 | CIPC | ZFP62 | LPXN | RNF168 |
| PPP4R3B | RIPK2 | PCLO | ZNG1B | IRAG2 | SHROOM3 |
| IP6K2 | CACNB3 | C1QTNF4 | MYL11 | BCAM | HBQ1 |
| UROD | HAX1 | TUBG2 | MT1H | RP2 | RARS1 |
| ATG13 | MAN1B1 | CSNK1D | COMMD9 | BLOC1S4 | VIPAS39 |
| SH3GL2 | RAB5B | CYC1 | TAF4B | ADCY6 | ZNF304 |
| CCL4 | FBXO11 | ATF6 | NBPF1 | OSBPL5 | WWP2 |
| MLF1 | SUN2 | FKBP5 | TMEM87B | ATAD2 | MUL1 |
| ARPP21 | PDZD2 | BRAF | DENR | DPAGT1 | LRMDA |
| PDE2A | SRRT | TCFL5 | SLC24A4 | CT75 | PIK3AP1 |
| HPCAL4 | OPN3 | HNRNPF | ERBB4 | ZNF641 | SESTD1 |

| | | | | | |
|---|---|---|---|---|---|
| CELF2 | NFAT5 | IFI27L2 | MFSD4A | ZNF778 | RILPL1 |
| DIRAS1 | CFL2 | WAC | LRRC7 | ZNF626 | HDAC9 |
| MARF1 | ATXN10 | A2M | DNAJA2 | CD300A | ZNF740 |
| PLEKHJ1 | LRRC42 | GDE1 | MFSD13A | HGF | KLK7 |
| MAP1LC3A | CTH | KCNMB4 | ATF5 | FOXP4 | MAD2L2 |
| EIF6 | HDAC9 | ZNF655 | FAM50B | LINC00951 | TTLL4 |
| KCNIP4 | MRPL52 | FMR1 | IKZF5 | C8orf82 | ANKRD18A |
| MAPK10 | KLHL21 | UBE2L3 | CHMP3 | HAPLN2 | ARVCF |
| SV2B | ZNHIT6 | ATP5PF | MTX1 | BTN2A1 | ZNF74 |
| TAF7 | FCGR3A | CCT8 | MAP4K5 | STYK1 | MAPK8 |
| ZC3H15 | ERO1B | PHPT1 | ADNP2 | CARS2 | RORB |
| EPB41L4A-AS1 | PNPT1 | TCEAL2 | MTMR6 | GRM7 | DENND11 |
| SULT4A1 | KCTD18 | SUGP2 | UBE2G2 | ROBO3 | SOX8 |
| PSMD8 | GTF2H4 | TMEM9 | ZNF711 | SLC15A4 | NMRK1 |
| ETV5 | TMEM209 | MRPL51 | SNRK | HAR1A | ARID1A |
| DEDD2 | RRS1 | KCNJ10 | SGTB | THBD | NNMT |
| RRP12 | POLR1E | TKT | C1orf52 | SLC36A1 | CCDC74A |
| PLAAT4 | GAL3ST3 | ECHS1 | PABIR2 | ZFYVE26 | FUOM |
| ANXA6 | L2HGDH | ISG15 | MIR99AHG | ZNF524 | B3GALT6 |
| ID4 | NIP7 | APOC2 | HAPLN4 | HAT1 | WDR74 |
| HNRNPU | DNAJC11 | SEZ6L2 | ZNF428 | LMF1 | FAM149A |
| CAMK4 | CDC5L | TXNL4A | ITPRIPL2 | USP27X | PUM2 |
| ATP8B2 | CALB1 | BEX2 | CDS2 | SATB2-AS1 | SP100 |
| KRCC1 | MGAT5 | HSPB1 | MRPL23 | ZNF536 | LINC00320 |
| JCAD | MAP7 | SCIN | GRIN2A | CCDC107 | MCTP1 |
| POLR2E | ZCCHC3 | C1QBP | CYP1B1 | SLIT2 | GLS2 |
| RTN4RL2 | ANKRD9 | AURKAIP1 | SH3GL3 | ELK3 | MINCR |
| ELOVL1 | PHF20L1 | MYL12B | SIRT2 | GPR161 | LINC02762 |
| SP100 | MAN2C1 | ARMCX3 | INTS11 | CCL20 | EFEMP2 |
| LDB1 | GOLGA4 | COPS8 | CAMSAP3 | ZNF783 | TRIM46 |
| POLI | RGMB | MAP1LC3B | ZMYM2 | NOM1 | PDGFRA |
| STK11 | CORO1A | OGFRL1 | RRM1 | RBM12B | TMCO3 |
| SLC24A3 | EIF3I | UBL3 | RNF152 | MRE11 | PPM1D |
| UCKL1 | CRKL | IL6ST | WDR11 | SMAD9 | CDK5RAP1 |
| PDK4 | KCTD16 | SNHG5 | SERPINB9 | PRKD1 | PHKG2 |
| NTN4 | TIAM2 | UBXN1 | ACTL6B | ALOX12-AS1 | SMIM10L1 |
| TPD52L1 | MDN1 | PIM3 | CENPV | CCDC120 | CDK10 |
| AP1M1 | IGBP1 | LAMTOR1 | MYRF | PDYN | KPNA6 |
| CDIP1 | ATP13A3 | CHGA | LINC01346 | ZNF267 | BUB3 |
| MEAF6 | LAMTOR4 | DUSP6 | TRMT1L | CHCHD5 | STK4 |
| CABIN1 | STXBP5 | GYPC | CDC6 | CACNG4 | WAC-AS1 |
| CKMT1B | MEF2D | ADIRF | HES6 | HECTD3 | ST6GALNAC5 |
| SLC35E2B | KCNQ2 | TIMM8B | TRIQK | AGA | CORO6 |

| | | | | | |
|---|---|---|---|---|---|
| GMFB | KIAA1191 | HLA-DRB1 | AKAP9 | ZNF587 | VAT1 |
| COA5 | PANK3 | C12orf57 | SMARCA4 | C4orf46 | SCAF11 |
| TFRC | AGAP3 | PIP4K2A | ANXA2 | PAXIP1-DT | BDH2 |
| MAP2K3 | LINC-PINT | MAPK6 | TOMM70 | CNTN5 | ZNF593 |
| EPHB6 | RASGRF2 | CHMP4B | PHF3 | KHNYN | ANKFY1 |
| SCN1B | POU3F3 | SLC24A2 | TRIM52-AS1 | PITPNA-AS1 | TMEM131 |
| IMPDH2 | MRPL33 | POLR2L | DBNL | ASPA | RPRD2 |
| HIGD1A | BBOX1 | NRCAM | AIDA | KCNAB3 | GPD1 |
| SELENON | OSBPL2 | SPCS2 | RYBP | AGO3 | TP53I13 |
| SUPT7L | ERRFI1 | RPS4Y1 | SETD3 | SLC25A42 | KDM6A |
| FRMD4A | CLOCK | TMOD2 | DIRAS3 | MOBP | HMGN5 |
| MORN4 | GAPVD1 | BTF3L4 | LRRC3B | LINC00882 | GPRASP3 |
| RTL6 | DBP | NUDCD2 | DTX4 | SPG11 | LDLRAP1 |
| LONP1 | FNBP1L | RFK | TMEM135 | MYO1C | KLF15 |
| MICU1 | C1D | RC3H2 | RNF214 | ASXL2 | INTS12 |
| DEXI | IPCEF1 | SLC43A3 | BAZ1A | PPIL6 | SPACDR |
| IGSF9B | CCZ1B | HLF | MSRB1 | HPS5 | PRKCQ-AS1 |
| TSNAX | JAZF1 | FAM174C | HSDL1 | DHRS4-AS1 | C11orf71 |
| PRKAR2B | GRB10 | TMED1 | CCDC85A | ZNF423 | KIAA0753 |
| SLTM | DDX19A | CPNE1 | CASP3 | SLC46A1 | ZNF575 |
| PSMC6 | SYT13 | CISD2 | JOSD2 | ZNF571 | DENND3 |
| C19orf25 | ARG2 | CDC42SE2 | LINC00632 | TMEM106C | ZFYVE9 |
| EFR3A | CKAP5 | LARS1 | SLC2A13 | TUBGCP3 | TXNDC5 |
| FNTA | ELMO1 | BUD23 | KCNJ3 | CNNM1 | PLXDC2 |
| CD86 | KIAA0232 | DYNC1I1 | WFS1 | DLX6-AS1 | IRGQ |
| PLOD2 | PPP1R2 | MICU3 | GABRA3 | METTL13 | WSCD2 |
| LINC00320 | CAMTA1 | CNOT7 | C11orf87 | CDH10 | DZANK1 |
| PIGA | MCTP1 | TBC1D24 | PRDM16-DT | NCOA2 | BAMBI |
| HLA-DQB1 | PPP4C | TAOK1 | PTPRO | LENG1 | ARL14EP |
| TMEM123 | NMT1 | HSPA13 | NTM | CYP27A1 | CDC42EP1 |
| PRPF31 | LRRN2 | BAP1 | CHM | UGT8 | MRPL17 |
| TRIP12 | ARL13B | GPAA1 | MAG | PHYKPL | MIR7-3HG |
| DGCR5 | UBA6 | CHMP2A | BPTF | SLC25A13 | ETV3 |
| DAPK1 | IL13RA1 | NOMO1 | GGCT | ANKS6 | MRPL13 |
| HSPA4 | DPF1 | PPCS | EBAG9 | NAALAD2 | ZNF148 |
| SCN8A | TBC1D4 | ELAPOR1 | NPAT | MMP15 | KCNK12 |
| FIS1 | HECTD1 | GPATCH4 | NACC2 | PRR34-AS1 | DRD1 |
| TULP4 | SH3BP2 | SYT14 | C5orf15 | RUBCNL | OBSL1 |
| KAT7 | CD200 | COA6 | ERF | TACO1 | KIAA0319 |
| BPGM | CCL3L3 | VKORC1L1 | MSX1 | ZNF83 | CERT1 |
| ATP2B2 | EPHA4 | NCALD | MINDY3 | HRK | FKTN |
| RANBP9 | RALB | UBR5 | NKAPD1 | PMM2 | TNFRSF14 |
| SF1 | ENTPD4 | ADARB2 | KAT14 | ATG10 | FAM114A2 |

| | | | | | |
|---|---|---|---|---|---|
| TOMM34 | TRAF7 | ZDHHC5 | KRT17 | ZNF775 | IGF1R |
| EIF2AK2 | LSM12 | STX5 | THSD7A | PLEKHF2 | TUB |
| CACYBP | URB1 | LLPH | SULT1A3 | VSTM4 | PPIG |
| GFPT1 | PLEKHA6 | BLTP3B | MORC2 | TMEM273 | LINC01719 |
| KCTD5 | DCAF5 | KCTD10 | SAMD1 | RAB30-DT | INTS8 |
| PRPSAP2 | MOK | IGF1R | PPP1R12A | MEGF11 | DCHS1 |
| C15orf61 | ADGRL2 | UTP4 | TBPL1 | DSN1 | HYPK |
| FLYWCH1 | GAS6 | GRN | PLXNA3 | BCL2 | HSF4 |
| DCUN1D3 | USP2 | KANSL1 | CCDC57 | GREM2 | CLPP |
| CARNS1 | MAP4K3 | PSMG2 | FHOD3 | PAPSS2 | WWC2 |
| NIFK | ABCE1 | TOM1 | NSDHL | CABP1-DT | GALNT15 |
| FSTL3 | STOX2 | ITM2A | GLI3 | DACH1 | BCHE |
| MAFF | BLVRA | ADIPOR2 | TEN1 | SNURF | SEC31B |
| SCN2A | CASD1 | SDHA | FBXO32 | TMEM88 | MC1R |
| TPD52 | SFSWAP | AKR1B1 | CNDP1 | SAP130 | ZNF738 |
| FBXL15 | LMF2 | GNB2 | ACBD6 | GPR19 | URB1-AS1 |
| CKMT1A | IMMT | C1orf43 | THAP9-AS1 | FAM222A | ENTPD3 |
| MXRA7 | STARD3NL | PTTG1IP | CIART | RELN | LRCH1 |
| KLC2 | BAZ1B | APOL2 | MAST4 | RHOJ | FOXC1 |
| PSME4 | IGSF21 | ENAH | VPS37B | H2AJ | CCDC149 |
| NDUFAF3 | MAP7D2 | PNO1 | TIE1 | ECHDC2 | ZNF614 |
| LANCL1 | AP3B2 | EIF3J | SLC16A9 | KRBA1 | ZNF777 |
| EIF2AK3 | SPIDR | ERC2 | WRAP53 | CA13 | LIN7A |
| LARP7 | KLF16 | HMCES | DNAH17 | KLHDC4 | AGAP4 |
| SLC31A1 | HS6ST1 | FNIP1 | CD93 | ZNF382 | CCDC117 |
| MAGEH1 | PLPPR4 | MCMBP | SLC66A1 | EVA1C | SH3BP1 |
| SNHG3 | CELF3 | GTF2A1 | ZBTB22 | ZMAT4 | BACH1 |
| GPR161 | TRAPPC2L | SPECC1 | SLC6A9 | TMEM63A | SUGP1 |
| RFTN2 | G3BP1 | MRPL57 | DRAM2 | PRR5 | CYB5A |
| KIT | PIP4K2C | MAP1S | DIS3L2 | SEMA3C | MLLT6 |
| SLC25A48 | MIR4435-2HG | FGD5-AS1 | ART3 | RGS7BP | RSAD1 |
| RGS14 | CHAC1 | GSTM3 | CCNO | MS4A7 | MRPL43 |
| LSMEM1 | GABRG2 | PTPRA | MCPH1 | MBD6 | TLR4 |
| CREB3L2 | ISG20 | NOP53 | PIGO | ASCL1 | SMIM30 |
| POMK | NDN | CCN1 | TAF5 | PLGLB1 | APLNR |
| CACFD1 | ZNF331 | DNAJB14 | ABCC9 | CAND2 | RPA3 |
| NOP9 | TTC32 | RPN2 | RACGAP1 | TNFAIP3 | ATL3 |
| RYR1 | TTC7A | AAMP | YEATS4 | TMEM209 | RAB13 |
| MYL9 | MAP3K2-DT | COPS4 | HCFC2 | TGS1 | CYP26B1 |
| CNTN2 | ZNF277 | KIF2A | SALL1 | ANO4 | ZNF317 |
| PAK1IP1 | MAK16 | TMEM126B | RAB11B-AS1 | MRI1 | ELP3 |
| SMC1A | RGS3 | GLYR1 | ZNF507 | ARFGEF1 | TMTC1 |
| ANKRD46 | NARS2 | FGFR1OP2 | PDCD2L | CCDC61 | RPRD1B |

| | | | | | |
|---|---|---|---|---|---|
| SNN | EXOSC8 | PSMB4 | MYCBP2 | LTBP1 | NOL9 |
| AAK1 | PELI2 | GADD45A | RAB43 | ABHD14B | ATOSB |
| PIP4K2B | ARMH4 | TAGLN2 | TNRC6A | FAM241A | CDO1 |
| DOP1A | SHC4 | OPCML | CACNA1E | PTCD2 | PHIP |
| PCSK7 | FLYWCH2 | XPO6 | DIO2 | TMEM176A | ID3 |
| PDE4A | PITPNA-AS1 | XBP1 | ITPRID2 | EPHX2 | ANKS1A |
| KCNV1 | KIAA0753 | ENSA | RELT | NFATC3 | ATP11B |
| FAM171A2 | ZNF554 | LRP1 | RSAD2 | FMNL1-AS1 | DALRD3 |
| MMD | DHDH | PSME1 | NUMB | LIG1 | RANBP6 |
| DDN | TMEM47 | NCOA1 | MEX3C | GLRA3 | DTX1 |
| POLR3H | USP34 | STMN3 | DLGAP2 | CROT | VIPR1 |
| EIF2S2 | PUSL1 | DPYSL3 | TMEM65 | PTBP3 | DTNB |
| ZNF706 | AGTRAP | ANAPC11 | MED25 | AREL1 | AHCY |
| NUDT16L1 | SCYL3 | AZIN1 | ADD3 | ALG8 | ERBB3 |
| SUCLG1 | CAVIN2 | ARPC2 | CHMP1A | BBOF1 | ICE2 |
| EGLN3 | SMC4 | PRKAR1B | RGS5 | MDC1 | POLG2 |
| PNMA3 | CXCL11 | SCAMP5 | FOXQ1 | TMEM19 | ZNF579 |
| NDUFB5 | ETFDH | ANXA7 | SVIP | FASTKD1 | ARL15 |
| ASAP1 | MANEA | TALDO1 | ECT2 | POGLUT1 | THEM6 |
| PLEKHA5 | NOS3 | NFE2L2 | LRRC57 | PCDHGC3 | COG8 |
| GAS7 | ZNF251 | STX4 | SMAD3 | GLIPR2 | TTC14 |
| CD276 | COMTD1 | PDCD6 | FBN1 | ASCC1 | ZFPM2 |
| ECE1 | ENTPD1 | P4HB | DLGAP1-AS4 | TRIM68 | TMEM176B |
| CNPY2 | GDPD5 | BSCL2 | ASF1A | EDC3 | MYO1E |
| NUS1 | NNMT | RASSF8 | SLC7A8 | KANSL1-AS1 | RBM6 |
| SLC29A4 | RHEBL1 | VIM | ZFPM1 | KDSR-DT | GLT8D1 |
| SMNDC1 | CLBA1 | PAFAH1B1 | ZNF846 | PPARGC1A | LRRFIP1 |
| MED25 | VMO1 | RABGAP1L | PPP1R1A | NAAA | NQO1 |
| GGT7 | LINC01535 | SPATS2L | IL6 | CTSO | BOK |
| LSM5 | GMFG | TOMM5 | PGM3 | EXOSC3 | TBC1D1 |
| EXOC7 | PLCB1 | IFI44L | KANSL2 | SEC14L5 | PKD2 |
| ANP32A | AKR1A1 | TOR1AIP1 | BZW2 | LLGL1 | RFX5 |
| MFN1 | RIMS2 | CUL5 | EIF5B | CASKIN2 | SLN |
| BRD9 | ADSS2 | FBXO7 | AP1G2 | RGS1 | IFT20 |
| TERF2 | GYG1 | EIF3K | CFAP119 | USP1 | SLIT3 |
| MT1H | SMIM20 | RAB31 | ARL5A | WNT10B | E2F6 |
| SRSF4 | ZBED5 | ANKRD40 | KDM1A | XAB2 | EGFR |
| ANKRD6 | RIC8B | TPD52L2 | MPZL1 | CLEC16A | GNB1L |
| SCAMP2 | CDK10 | HDGF | ABHD6 | HOOK2 | GNPDA1 |
| STAT5B | KIZ | PSMD2 | UPF1 | DHRS9 | ATAD2B |
| ATP6V1D | ANKHD1 | ISCU | FBXO31 | SYNE2 | C2orf88 |
| TPGS2 | GDA | STIP1 | ZNF703 | ACBD4 | PPP1R9A-AS1 |
| MAST1 | CCT7 | NDUFV1 | FCGR1A | SUSD5 | NDUFA4L2 |

| | | | | | |
|---|---|---|---|---|---|
| RNF168 | SCYL1 | ATP5PD | TMEM167A | LINC00595 | C19orf25 |
| PCCA | H4C8 | GRPEL1 | C16orf89 | DPY19L2 | ZNF821 |
| SOCS4 | DNMT3A | PCDH10 | RAB5C | KCTD4 | CHRNA4 |
| CD93 | IMP3 | NR4A2 | ATXN3 | GABPA | SMIM17 |
| MGRN1 | AVL9 | FGFR3 | NUDT9 | FOXO4 | MAVS |
| TPRG1L | RNF6 | NFIB | EMP3 | MAML1 | PAK5 |
| HBS1L | TLN2 | COPS6 | SEC63 | KYAT1 | ZFAND1 |
| BCL2L1 | FOXP1 | USP11 | CCDC28A | PDSS1 | RNGTT |
| EP300 | NECAP1 | TRAPPC5 | CHD8 | IGLON5 | LSMEM1 |
| OAF | LBH | MCFD2 | KCTD1 | FKBP15 | FAM135B |
| ARHGEF25 | GNAI1 | MPC2 | RPS6KA3 | CXCL2 | ST8SIA1 |
| ST8SIA5 | G3BP2 | SIRPA | PTPRD | | PNO1 |
| CNNM1 | ABCF2 | ARHGAP32 | FAM204A | | SEMA3F |
| TGFB2 | CNTNAP1 | MACO1 | IPO13 | | ZNF487 |
| LYPD1 | NOP56 | STAMBP | NECAB2 | | ABCG4 |
| COLQ | NARF | PDE6D | MXI1 | | TICAM1 |
| CYGB | LAGE3 | ARL4C | FAM219B | | ZSCAN1 |
| ZNF263 | ATF2 | GUCY1B1 | NOTCH1 | | USE1 |
| LIX1 | PDZRN3 | FAM169A | CANT1 | | CPNE3 |
| HERC1 | XYLT1 | STARD4 | KHDC4 | | PRPF3 |
| RBFOX2 | ACADVL | SLU7 | LPIN2 | | ASTN2 |
| UBE3B | SEC31A | RIOK1 | CCDC28B | | CWC22 |
| C2CD2 | AOPEP | TSPYL4 | SORCS1 | | RIOX2 |
| SPATA2 | DNAJA3 | NEIL2 | OFD1 | | CXCL3 |
| EDNRB | DHX38 | ARFGEF1 | LINC00507 | | ANO3 |
| FN3KRP | VEZF1 | KCNT1 | METAP2 | | CEP164 |
| NPLOC4 | PENK | PAX6 | SH3D21 | | SOCS1 |
| CHD2 | PLEKHM3 | CSTF3 | SNX25 | | CIDEA |
| ARF5 | DMTF1 | APLNR | SLC31A1 | | CCDC9 |
| HAPSTR1 | FOCAD | N4BP2L2 | LDB1 | | PIK3IP1 |
| ELMOD1 | DIPK1B | FBXO33 | ARID2 | | RNF20 |
| TRIM2 | EXT2 | DPP8 | UGGT2 | | PSMG4 |
| HRAS | RELT | GOSR1 | ZNF419 | | SFMBT2 |
| PRRC2B | KMT2D | RBFOX3 | CHMP2B | | CCKBR |
| ILRUN | ANKRD13B | FOXK2 | SPAG16 | | PPTC7 |
| LMBR1 | AKT1S1 | MEX3C | NOP14 | | RASL11B |
| MAP4K5 | MIEF1 | ACOT8 | FAM118B | | LINC01896 |
| PBDC1 | DENND6B | PMM1 | RAB7B | | ACOT9 |
| LRIF1 | SHC3 | PRR5 | MXD1 | | SPATA2L |
| RPA3 | TMEM132B | NAP1L3 | TRIM24 | | TRPM7 |
| BRSK2 | RTN4R | LPIN1 | TCF20 | | GORAB |
| ABCF3 | LARP6 | MED10 | FBLL1 | | NEK4 |
| CGGBP1 | UBE2E1 | TMEM248 | UTP25 | | NAF1 |

| | | | | |
|---|---|---|---|---|
| PNMA6A | TIMM50 | NKAIN3 | GRIK1 | FOLH1 |
| NAGK | PSMD6 | RUSC2 | JMJD8 | C21orf91 |
| FAXDC2 | PPP2R5E | FAM234B | SPOP | MED30 |
| COA4 | PALM2AKAP2 | MYG1 | PMF1 | GPR83 |
| GSK3A | NAPG | HBA2 | FRS2 | TMEM98 |
| SATB1 | SLC25A38 | VSTM2L | TRIO | IFNGR2 |
| PAXX | LUC7L2 | LARGE1 | SLC39A9 | RBM4B |
| GLRB | TMEM181 | DDI2 | KCTD20 | TM9SF1 |
| OCIAD2 | DYNLT3 | ZMYM4 | HIPK3 | ZNHIT6 |
| KARS1 | SHOC2 | ST3GAL3 | ENG | EXOSC4 |
| ITPK1 | FOXJ3 | TOE1 | RFX3 | DHX33 |
| FAM168A | ATXN7L3 | STRIP1 | PNOC | PI4KAP1 |
| ZSWIM6 | JARID2 | MAGI3 | EDNRB | STX18 |
| BLTP1 | FBXW7 | UBAP2L | BDP1 | OSBPL2 |
| VASP | CELF1 | NCSTN | ATAD1 | SEC24D |
| TIMM10 | AKR7A2 | RGS5 | PDE8B | YJU2B |
| C1orf53 | PLK3 | SLC41A1 | DAAM1 | TMEM43 |
| MARCHF4 | PPP1R1A | DDX18 | ERC1 | CYP7B1 |
| RNF25 | LCMT1 | TMEM185B | TIPARP | SNIP1 |
| IFT80 | TST | ERCC3 | STX4 | TRIM36 |
| SNHG18 | DCAF1 | AGFG1 | ELOVL7 | SEC24A |
| EPHA7 | TMEM170B | ISY1 | CTXN3 | POGK |
| SPACDR | ST20 | VPS8 | PPP1R14C | UBXN7 |
| ATAD2 | SCOC | SLIT2 | RANBP3L | SIRT5 |
| DOLPP1 | KLHDC3 | GAB1 | PMPCA | RIPOR2 |
| SLC16A9 | SNTB2 | ICE1 | RO60 | IPPK |
| IDE | ASB8 | SERINC5 | LRRTM4 | CHRM1 |
| CNTN5 | CSF3 | CNR1 | SYNJ2BP | RYR3 |
| PUS7L | LRRFIP2 | CDK19 | TSC2 | PARVA |
| TTLL5 | COL9A2 | TMEM242 | BCAR3 | HIRIP3 |
| GABPB1-AS1 | CACNA2D3 | PHF14 | RASAL2 | XRCC1 |
| MEGF11 | SETD7 | LMTK2 | LST1 | FOXN2 |
| CRISPLD2 | KIAA0319 | ZNF398 | PDCD7 | BEST1 |
| DCC | CLN8 | LEPROTL1 | THBS3 | RELA |
| ZNF584 | GPATCH8 | VCPIP1 | EOGT | TRMT10C |
| COL6A2 | AFG3L2 | PLAA | CDHR1 | PARP9 |
| PPM1F | VPS45 | GLIPR2 | SCAF4 | MBLAC2 |
| GGT5 | TRIM46 | PABIR1 | UTP14A | ST8SIA4 |
| TTC28 | HECW2 | POLE3 | FAM131B | FOXP2 |
| TMEM268 | GFM1 | TAF3 | LRRC20 | ZNF202 |
| LMO2 | TCOF1 | ANKRD26 | TJP1 | GNB3 |
| SEMA6D | ABCB1 | HBB | RASSF5 | NUP107 |
| RPSA2 | BTRC | UQCC3 | COL23A1 | FGD6 |

| | | | | |
|---|---|---|---|---|
| BRPF3 | TMEM63C | WDR74 | NCKAP5L | TINCR |
| ZBTB22 | ZNF770 | CDC42EP2 | ATP6V0A2 | ZNF581 |
| FPGS | STARD3 | NABP2 | SLC25A38 | BRAT1 |
| FSTL1 | SF3A2 | FGD6 | MRGBP | BMPR1A |
| PIK3CD | ZNF317 | SH2B3 | NEDD9 | PATZ1 |
| EPHA8 | NFS1 | C12orf43 | MPHOSPH10 | C1GALT1 |
| BMP8A | ZNF365 | FOXO1 | RETSAT | INO80D |
| ECHDC2 | PPIL2 | NUFIP1 | CEP85L | RSU1 |
| COL11A1 | MICAL2 | LRRC49 | PPP3CC | MRM3 |
| HIPK1-AS1 | AIF1 | CRTC3 | FAM13C | TENT5A |
| NBPF26 | PITPNC1 | ATF7IP2 | MLC1 | KIT |
| MINDY1 | MTHFD1L | NFATC2IP | CTDSPL | BCL9L |
| ANKRD45 | FLOT1 | E2F4 | OSGEP | SFI1 |
| DYRK3-AS1 | TTL | VAT1L | DNPH1 | APOL6 |
| ESRRG | MUSTN1 | PELP1 | AIFM3 | DUSP11 |
| ALKAL2 | ARMH3 | MED31 | ZNF12 | CTBS |
| TP53I3 | TRPT1 | SMARCE1 | SRCIN1 | OGG1 |
| SLC30A6 | MSI1 | DHX58 | CFAP410 | SLC35B2 |
| LINC01159 | P2RX4 | DUSP3 | PLCB1 | UMAD1 |
| ST6GAL2 | MPHOSPH9 | DCAKD | UBE3A | SH2D3C |
| GLI2 | ICAM2 | CUEDC1 | CREBL2 | SOHLH1 |
| CHROMR | XAB2 | CANT1 | FAM20B | TOP3A |
| NUP35 | GGCT | SMAD4 | TLCD3B | POLI |
| FZD5 | DAGLB | SNAPC2 | STAM | CCDC159 |
| KCNE4 | INPP5A | MEIS3 | PRKCI | ZNF136 |
| ANKMY1 | SPHK1 | CTNNBL1 | KCNS1 | TMEM42 |
| FAM240C | NBEA | DIDO1 | RBM42 | EEIG2 |
| IL17RE | ITGA2 | PCMTD2 | DUS1L | TRMT61B |
| XPC | LRRC41 | LTN1 | PAQR8 | HEATR1 |
| CMTM8 | B4GALT3 | SLC25A18 | ANKRD28 | TCAIM |
| CMTM7 | TAF5L | UFD1 | SLITRK4 | LINC00499 |
| PLA1A | PDLIM3 | PACSIN2 | GPX3 | MAGOHB |
| GATA2 | UNC5D | SCO2 | PCNT | RILP |
| SPTSSB | MLLT10 | SCML1 | TMEM175 | RWDD2B |
| LMLN | VWA5A | UBE2K | GBP3 | AVPI1 |
| ZBTB49 | HMG20B | PNMA2 | FCRLB | TMEM145 |
| FAM184B | ZC3H4 | RIPOR1 | ZNF496 | TGFBRAP1 |
| LINC01088 | PAK5 | RCAN2 | TMEM44 | ZC3HAV1 |
| RAI14 | VPS35 | PDPN | COL25A1 | CRB1 |
| CARD6 | HDDC3 | CRIPT | GALNTL6 | MLIP |
| ITGA1 | HIGD2A | SMARCC1 | PCDHB16 | NR2E1 |
| OCLN | NDUFS3 | KPNA4 | PDSS2 | ZNF252P |
| FOXD1 | TMEM178A | CHST11 | FBXL13 | NPIPB13 |

| | | | | |
|---|---|---|---|---|
| ZBED3 | ACBD7 | SBNO1 | GARIN1B | RAB3D |
| PCDHB7 | TSPAN5 | CDC42BPA | GATD1-DT | FBXO17 |
| FGFR4 | MYH9 | NRP2 | PRR5L | MMP24 |
| GMPR | PRRT1 | NR3C1 | TECTA | SEMA6A |
| APOM | ST6GALNAC5 | CASTOR3P | LACC1 | SCNM1 |
| FKBPL | FAM50B | FGFR1 | CPB2-AS1 | B4GALT3 |
| ARMC2 | TMEM276 | SNX30 | TEX29 | GATB |
| LAMA2 | ERCC5 | MS4A7 | HAUS4 | DGLUCY |
| PARP12 | PRMT5 | NT5DC3 | KLHDC1 | CKAP4 |
| GIMAP6 | FAM117A | PIAS1 | C15orf48 | CYB5D2 |
| PAXIP1-AS2 | SRXN1 | MSI2 | GOLGA6B | HDAC3 |
| PAXIP1 | PEDS1 | ACAA2 | TSNAXIP1 | GPR27 |
| FUT10 | MCHR1 | ADGRE5 | ADORA2B | IPO11 |
| CRH | SMARCD1 | TYMP | GRIN2D | STX2 |
| KCNS2 | MTERF2 | PRELID1 | ALDH16A1 | ATG4D |
| FANCG | FAM98B | AGAP2 | SNAP25-AS1 | CEP63 |
| LINC03025 | SLC30A3 | FARSA | SPIN4 | LINC02482 |
| ZNF33B | URGCP | PNKD | VAT1L | ITGB4 |
| FRMPD2 | CRABP2 | CNIH1 | CDK18 | TDG |
| DNA2 | JMJD4 | CMIP | NIM1K | LAS1L |
| PDLIM1 | KCNAB1 | FDX2 | C14orf119 | TMEM150C |
| AS3MT | WDR19 | HNRNPAB | IPO9 | PIK3C2B |
| SFR1 | HTR5A-AS1 | AGPAT3 | LRRTM2 | RBMS3 |
| SYT9 | C8orf82 | MRPL55 | STAT2 | SH3TC2 |
| LINC02552 | TBC1D13 | DDHD2 | LRP3 | MIR219A2HG |
| TBCEL | VPS37C | PGM2L1 | SSRP1 | NUBPL |
| TMEM117 | NUDT22 | ARIH1 | ITM2A | CMTM3 |
| TNS2 | FOXJ2 | ZNHIT3 | ZBTB43 | LDHD |
| NDUFA4L2 | WDFY2 | SAP30BP | LRRC4 | ZBTB14 |
| TMEM263-DT | NEO1 | BTBD3 | ARID4A | CHAF1A |
| TMEM119 | NXT1 | REEP2 | COA8 | C3orf18 |
| SDSL | TEKT4P2 | WNK2 | PRKAA2 | TMEM191A |
| KBTBD7 | TNRC6B | RAB21 | SWI5 | FAM241B |
| MRPS31P5 | RBMX2 | PTMS | SLC25A1 | TTC32 |
| OLFM4 | CFDP1 | MAGOH | LINC01503 | POMK |
| PIBF1 | SNX10 | CDC37 | TOPORS | METTL17 |
| PCCA-DT | MIR7-3HG | NDUFS4 | FGGY | INSYN1 |
| LINC02328 | CCDC106 | LPL | DOLPP1 | AMDHD2 |
| FOXN3-AS1 | USP12 | CNIH2 | DDX18 | SPECC1L |
| MYH11 | CA10 | SYNJ1 | YIF1B | MOB3B |
| C17orf114 | SLC8A2 | MAST3 | BMP2K | DSTYK |
| NLRP1 | MKNK1 | ELOVL5 | FAM161A | CDADC1 |
| LINC01563 | MRPL24 | DNAJB9 | FRZB | ODF2 |

| | | | | |
|---|---|---|---|---|
| SLC46A1 | SELENOI | MIGA1 | TNIP2 | LSM2 |
| TCAP | REEP1 | MAGED1 | RNF180 | ARPC1B |
| TRIM65 | ABCC5 | OSBPL1A | EIF4EBP3 | ARHGEF18 |
| TYMSOS | EHBP1L1 | TRIM28 | PSMG3-AS1 | MOG |
| CFAP53 | ETFRF1 | CLTA | VIRMA | IGSF11 |
| ZNF30 | RAD51C | FABP7 | HMGXB4 | CLYBL |
| IGFLR1 | FAM177A1 | UBA1 | AFF3 | ZNF518B |
| ZNF225 | PIP4P1 | SORBS1 | PSMA3-AS1 | CCNB1IP1 |
| MAMSTR | ORMDL1 | CCND2 | MAPKAPK5 | CPNE8-AS1 |
| POLD1 | CHCHD1 | MFAP3L | FCGR1BP | NUDT6 |
| TMEM238 | ABHD17A | CAMLG | UCK2 | LY86 |
| ZNF582-DT | ADD2 | RNF114 | LYST | LRTM2 |
| ZNF776 | NTRK3 | PSMD11 | ERCC6L2 | CLCN2 |
| FERMT1 | JKAMP | H3-3A | DDHD1 | SAPCD2 |
| SLX4IP | NUCB1 | NONO | ELL | DGAT2 |
| CCM2L | FGF12 | FKBP1A | TJP2 | AMH |
| POFUT1 | TDP2 | PIN1 | POU3F3 | TMEM161A |
| SLC2A10 | TARDBP | PPP1R7 | BRI3BP | NR2C2AP |
| LINC01637 | SLC8A1 | GOLM2 | NCOA5 | ZNF528 |
| EPIC1 | BMP7 | MAFK | VPS13D | NOL3 |
| NEXMIF | OTUD3 | CTBP1 | FAM98A | JAG2 |
| FAM133A | TUT4 | SURF4 | FRY | FZD1 |
| EIF3E | ATP11B | BRI3 | LNX1 | WASHC1 |
| RTCA | TRIM27 | GOLGB1 | PREX1 | GRIN3A |
| ZNNT1 | NUDT2 | BSG | CALCOCO1 | CCNT1 |
| VPS13A | KBTBD4 | FNIP2 | TBC1D23 | TAF1C |
| CC2D1A | GAB2 | WDR26 | CAPZA1 | RARA |
| GMCL1 | RNF34 | S100A6 | NPRL3 | POLG |
| COPG1 | ARHGEF40 | CSNK1A1 | ANKRD6 | HNMT |
| TBC1D1 | FAN1 | FUBP3 | SLC39A14 | KLF12 |
| PPIL1 | INO80 | PDPR | IL17RC | LILRB4 |
| PAGR1 | CTDP1 | DCTN3 | P3H1 | NFX1 |
| STRADA | DOHH | SEPTIN5 | WDR3 | FAIM |
| KCNC3 | LRRC4B | VCAN | C2orf42 | AMPD3 |
| PRUNE2 | ZNF512B | NEDD9 | NABP1 | NOP9 |
| IFITM1 | FBXL2 | TNFAIP3 | DHX32 | PRKCH |
| TMEM14B | CACNG8 | ASAH1 | MGP | ARHGAP29 |
| ANKRD10 | MED15 | INF2 | BFAR | GABRG3 |
| LGI4 | CYP1B1 | GBP2 | ASGR1 | PLCH2 |
| PPP2R5D | OXSR1 | PI4K2A | CEP95 | LAMC1 |
| CHPF | CLK4 | B4GALT5 | HOMER3 | RUFY1 |
| SFT2D1 | PSMD5 | PTPA | RYR1 | CD52 |
| CREB5 | SEPHS1 | HSBP1 | ZNF134 | SNRNP48 |

| | | | | |
|---|---|---|---|---|
| NRIP3 | HIF1AN | STK40 | MACROD2 | ZNF432 |
| SOCS6 | CFAP68 | CTDNEP1 | CACNG2 | NHSL2 |
| PAFAH1B3 | PLBD2 | KXD1 | WNT7B | HDAC8 |
| PA2G4 | PRKAB1 | PDCD5 | C8orf76 | TRAF5 |
| SPOCK3 | EMC9 | EIF5B | GPR3 | NXPH2 |
| DOC2A | IFI27L1 | KLF4 | ALYREF | SOWAHB |
| PTPN5 | BCL11B | ZFP36L1 | ZNF542P | PRR3 |
| CHRD | ARHGEF18 | NDUFAB1 | WDR91 | USP49 |
| TUSC2 | ZNF254 | DNAJC7 | TALAM1 | MED23 |
| DLG3 | PTCHD1 | KHDRBS1 | TTC33 | CDK6 |
| FAM133B | SGTA | SEPTIN3 | ARHGAP39 | BHLHE22 |
| ZDHHC14 | LINC02693 | ASH1L | ATP13A1 | RSPO2 |
| HSPA12A | POR | CD47 | ADGRA3 | INTS4 |
| UBE2A | HGSNAT | FAM219A | MACROD1 | IKZF4 |
| NME3 | HRH3 | DIO2 | RASSF8-AS1 | MTRFR |
| VEGFA | TCEA1 | VEGFB | BIVM | PLD4 |
| AHI1 | LINC01128 | SLC4A4 | SETBP1 | TRADD |
| NOL7 | BCAR3 | ADH5 | OCRL | NKX2-2 |
| EFTUD2 | CD55 | SPTBN2 | NTAN1 | MN1 |
| CACNB1 | BRMS1L | COMT | SAP30 | ASCC2 |
| LAPTM4B | ADGRV1 | SNHG32 | DCUN1D5 | MCF2 |
| TMF1 | S100A10 | PSAT1 | MAP6D1 | TRBC2 |
| DBN1 | UBXN2B | TMEM33 | SCO1 | RBP1 |
| LGMN | PIANP | TMEM30A | FMNL1 | POLR1H |
| NEDD8 | ATAD1 | UBE2N | ORMDL1 | LACTB |
| TXNL4B | SERPING1 | CADM3 | FAM184A | GAREM2 |
| TAP1 | LRRTM2 | RAB3GAP1 | RND3 | LIX1 |
| GBP1 | PTGES2 | R3HDM1 | UBIAD1 | ANKRD13D |
| SMIM45 | LGALS3 | HABP4 | AGTRAP | LHX6 |
| SUCO | RYR2 | CDC42BPB | MARCHF4 | CNPY3 |
| SH3YL1 | SCN2B | APRT | GIGYF2 | ZNF266 |
| PIKFYVE | TMEM50B | CA11 | ELFN1 | SPTB |
| CMTR1 | RBP4 | OGT | ZC3HC1 | CMPK2 |
| SCARB1 | PCSK2 | NDUFC1 | MATN2 | ORC2 |
| KAT2A | EFNB3 | PTPRS | MACROH2A2 | MYNN |
| EZH1 | CERS1 | ZFAND3 | RAMAC | NDST3 |
| MISP3 | RPL26L1 | GNG10 | PRELID3A | F12 |
| ZNF568 | PRDM16-DT | NSA2 | SALL3 | KDM1B |
| TSEN34 | WASF2 | RNF187 | LRRC75B | PKIB |
| PRKCI | GNL2 | MCRIP1 | CCDC134 | PRORP |
| SPG21 | ANKRD35 | RAMP1 | RNPEP | ZNF768 |
| PICK1 | CGREF1 | RANBP2 | ITGA6 | ATPAF2 |
| CERCAM | ASB1 | MT1F | PPRC1 | RASIP1 |

| | | | | |
|---|---|---|---|---|
| PDK1 | UGT8 | NDUFB1 | AMBRA1 | NOTCH2NLA |
| ZNF385D | MTRF1L | DANCR | RTL8A | SOWAHC |
| PDCD6IP | VPS37A | PTK2 | ARHGAP12 | DCBLD2 |
| BBS7 | EML1 | BRSK1 | LRRC27 | ZNF318 |
| SKIC3 | TDRD9 | ACSL1 | SOCS7 | CENPP |
| PHIP | GSG1L | PSMD12 | SESN2 | CTNNA3 |
| CSMD1 | CDYL2 | MTLN | LPGAT1 | SERPINH1 |
| KIF13B | CEBPA | TBRG1 | PPP4R3A | BIRC3 |
| MCM4 | ALDH4A1 | DLG4 | RBCK1 | TBC1D16 |
| DNAJB5 | LHFPL4 | FUBP1 | LRRC58 | PLPP2 |
| MTMR2 | AGGF1 | MLLT6 | PRMT7 | DLG5 |
| MYO18A | SMAD3 | PRRC2C | WASF2 | POU6F1 |
| HP1BP3 | WFDC1 | ACBD3 | DNAJC10 | UNC5D |
| SGCB | SETD4 | FAM98A | ICA1L | MYH14 |
| HBP1 | SRGAP2C | UGDH | DEGS1 | ELP4 |
| ILK | SLC5A6 | LINC02217 | SMIM19 | CDC73 |
| CDK5 | SEMA4F | H4C3 | KRR1 | GLI4 |
| MAL | LRP1B | UCK1 | PITPNM2 | SNX30 |
| MARS1 | EFNA5 | NUMB | A1BG | SPI1 |
| UBE2G1 | PTP4A3 | IVD | ANGEL2 | EFNA3 |
| ECH1 | NOL6 | NDUFV2 | BCAS1 | RTKN |
| PDK3 | FKTN | CDC34 | SH3BP5L | IL17RB |
| KCNJ9 | XPNPEP1 | URI1 | GPR17 | DGKQ |
| SENP6 | ARL14EP | AP2B1 | UBA6-DT | ZNF608 |
| NUDT14 | AMBRA1 | MTHFD2 | ARSB | ADARB2 |
| BLVRB | RPS6KA4 | PRR18 | SIPA1 | ZNF574 |
| CLCN4 | LPCAT3 | CTSZ | CARMIL3 | TMEM63B |
| RGL1 | NAA16 | RNF115 | SNX33 | BCOR |
| SUSD4 | BIVM | CD84 | TANGO6 | KIAA1586 |
| SATB2 | CHD8 | GTDC1 | SNHG20 | C3AR1 |
| NUP98 | APH1B | MAP3K20 | KLHL7 | SPOUT1 |
| FDX1 | USP3 | PCBP4 | EPHB3 | EPHA5 |
| PFDN6 | BCKDK | LETM1 | TRAPPC13 | STAMBPL1 |
| ST3GAL1 | GHDC | SEPTIN11 | KANSL1 | TMEM109 |
| MMS19 | VPS25 | USP53 | THAP2 | MTCL1 |
| ANKRD42 | CASKIN2 | DNAJC21 | POLR2M | FMC1 |
| SUPT4H1 | TMC6 | SRF | CFDP1 | TNRC18 |
| CACNA1E | KLHL26 | SNX14 | TEAD1 | HSPB6 |
| DNTTIP2 | EID2B | TAB2 | TRNT1 | LINC01106 |
| DOCK3 | TMEM145 | ANKIB1 | RPAP2 | TMEM108 |
| SGK1 | IRF2BP1 | DHCR7 | NOL10 | OPA3 |
| PPM1A | PAXBP1 | THAP12 | PYDC2-AS1 | TBC1D22A |
| GDAP1 | IL17RA | VWF | ELF2 | AMY2B |

| | | | | |
|---|---|---|---|---|
| SLIT1 | ASCC2 | NAV3 | CGNL1 | MAPKAPK3 |
| AIDA | UBE2E2 | NFYB | NIP7 | CEP97 |
| PLCH2 | SLC25A46 | IAPKAPK5-AS | DMKN | FUCA2 |
| BABAM2 | ABCF1 | STARD13 | SDF2L1 | NCAPD3 |
| METTL5 | FAM241B | USP8 | B9D1 | KLK6 |
| ANKS1A | FAM153CP | ZFAND6 | FAM153CP | EOLA1 |
| ELP2 | TPST2 | SLC38A7 | KIF1C | SCAF8 |
| HDGFL2 | MED8 | SLC35B1 | LAMA4 | YEATS2 |
| GRIA1 | QSOX1 | RPS6KB1 | CENATAC | AGFG2 |
| CDC37L1 | MDM4 | MRC2 | AKTIP | C1S |
| RAP1GDS1 | ZCCHC12 | MARK4 | PPM1F | CYP4X1 |
| CHIC2 | CHRM3 | OLIG2 | NEFH | TMEM68 |
| TGOLN2 | TTC14 | CUL4B | MAT2B | CA4 |
| IRF2BP2 | DSE | RALY | FXR2 | NOTCH3 |
| PKN2 | CPSF1 | UBE2J2 | MID1IP1 | DHDDS |
| H2BC21 | CDK9 | HMGCL | MFSD6 | TAF6L |
| RDH14 | ZNF710-AS1 | RBMXL1 | PROX1 | SLC37A4 |
| TRUB1 | PRELID3B | ASAP2 | DNAL4 | ZFYVE1 |
| LRFN4 | RRP7A | MRPS25 | KCNQ4 | CRABP1 |
| NSRP1 | LYRM4 | ZDHHC3 | CLK2 | EPG5 |
| MAG | LRP11 | GRM2 | FCGR3A | PPARA |
| ASPHD2 | RANBP3 | TRMT10C | ZNF514 | WNT2B |
| HAPLN2 | RPRD1B | LPP | ANO10 | ABCB10 |
| FAM193A | SST | FRG1 | CFAP44 | USP40 |
| ERICH3 | SOCS2 | CRHBP | SRD5A3 | NUP210 |
| JAGN1 | PTRH2 | KDM3B | CRACD | EEFSEC |
| MRPL3 | TSTD1 | TBCC | ZNF623 | GSAP |
| YEATS2 | LIF | SLC25A37 | QSOX2 | DDX47 |
| SNX2 | SRRM4 | MAPKAP1 | ZNF488 | L2HGDH |
| NRP1 | EIF2AK1 | WDR5 | BDNF | HNRNPA1L3 |
| NUP160 | NDST1 | CPEB3 | CORO1B | BORCS6 |
| C16orf87 | ARHGEF10 | CACUL1 | CBFB | LYPD5 |
| PLA2G15 | CXCL12 | STIM1 | LINC00665 | GUCD1 |
| ZNF232-AS1 | CSTF2T | TTC9C | GPR26 | RAC2 |
| FKRP | ZNF432 | CLSTN3 | EBLN3P | CTNND1 |
| GCNA | PTPRF | CMAS | CLEC2B | CERS4 |
| CORO2B | KIFBP | SVOP | DGKE | ACTR6 |
| MAP2K2 | KLC4 | ADPRHL1 | SLC35C2 | TMEM163 |
| C1orf50 | MRPL16 | MEIS2 | FBXL7 | SPEG |
| PLCXD3 | ATF1 | TMED3 | RGS10 | PLEKHG1 |
| IGFBP3 | ERC1 | PDPK1 | TAFA2 | ZFHX4 |
| SMIM30 | NRN1 | CNOT1 | DISP3 | ABCA5 |
| SEL1L | PATL1 | RAI1 | CDK5RAP2 | BRD1 |

| | | | | |
|---|---|---|---|---|
| GPSM2 | RPAIN | MRPL10 | CDR2 | PREP |
| TIMM9 | TJP1 | SNX11 | HEBP1 | CD84 |
| MGAT4B | MLST8 | APPBP2 | RAB28 | KCTD3 |
| KLF13 | CCDC3 | GNA11 | NRAS | TRIT1 |
| PCP4 | ZBTB20 | CELF5 | ELK4 | CASQ1 |
| SS18L2 | OSER1-DT | SEMA6B | ALG14 | MTIF2 |
| DNAJC16 | ZBTB21 | PRR36 | THBS1 | IRS1 |
| RNPC3 | ZEB1 | MANBAL | CDK5RAP3 | METTL15 |
| HNMT | TMCO1 | GRAMD4 | PHLDA3 | RESF1 |
| WDR75 | ZNF207 | PLXNB2 | LCMT1 | COX10 |
| FRMD4B | ZBTB8OS | PNCK | NME5 | LIG3 |
| CFAP90 | TRNAU1AP | SKI | NT5C3A | SLC25A20 |
| AHR | POLR1A | TNFRSF14 | FBXO25 | LIAS |
| MCM7 | PDE12 | PER3 | SNED1 | TMEM243 |
| DCAF13 | PREP | MRTO4 | FEZF2 | AP4M1 |
| ZFYVE27 | HEY1 | DDOST | NUDT1 | TNPO3 |
| PPM1H | TGFBR1 | GPN2 | WHRN | ALG9 |
| SMAD9 | IL34 | PPP1R8 | CDH20 | PYM1 |
| COL4A1 | SMCR8 | GMEB1 | PRPF31 | TMTC2 |
| METTL3 | RABL2B | RNF19B | PRPF38A | ZBTB25 |
| INTS14 | FTX | COA7 | EHBP1L1 | DHRS12 |
| NKIRAS2 | B3GALT6 | ST6GALNAC3 | SMURF1 | NAPRT |
| SLC16A3 | PLA2G2C | DIPK1A | UBN2 | FGF9 |
| FECH | P3H1 | SYT6 | SLC19A2 | MTUS2 |
| CHST8 | EPHX4 | GLMP | KBTBD6 | PRMT6 |
| GRWD1 | SMYD5 | PEX19 | HINT3 | TBCE |
| MAVS | ERICH2 | SFT2D2 | MAP7 | ZNF273 |
| RUNX1 | CREB1 | G0S2 | RCC2 | MAILR |
| GSPT2 | SETD5 | KCTD3 | PHTF1 | KCNK9 |
| RBM6 | CDKL2 | ABCB10 | STXBP5L | MRRF |
| PSPC1 | ANAPC10 | NOL10 | SNX1 | TIMM23B |
| PTAR1 | MARCHF1 | E2F6 | ETS1 | COMTD1 |
| CSGALNACT2 | TRIM41 | MTA3 | FAM210B | KCNJ11 |
| IQSEC3 | MRPS18B | KCNK12 | XPA | INTS5 |
| SPSB3 | RRP36 | KDM3A | HECW1 | CEP83 |
| VSTM2B | MRAP2 | TGFBRAP1 | ATRAID | RPP25 |
| MRPL36 | TRAF3IP2-AS1 | MFSD6 | MOB4 | CTU2 |
| RLF | TUBE1 | HYCC2 | TRAPPC12 | GPR61 |
| PCNX2 | DLL1 | MTERF4 | ILRUN | SOX11 |
| CPD | NDRG1 | TADA3 | UBE4B | EPHA6 |
| HID1 | ENTREP1 | QRICH1 | TPST2 | ALKBH3 |
| CARM1 | AKNA | ABHD14A | VEZF1 | FNTB |
| CFAP298 | ANAPC2 | FEZF2 | PLPP5 | GABPB1-AS1 |

| | | | | |
|---|---|---|---|---|
| KDM1A | SFXN4 | NIT2 | LRIF1 | TRIP4 |
| TENT5A | EDRF1 | SENP7 | TOGARAM1 | CDH7 |
| GPNMB | ALG8 | CHCHD6 | ZNF330 | TMEM125 |
| COL18A1 | PRICKLE1 | ACAD9 | PLEKHM3 | PPM1J |
| EMC3 | LRP10 | PXYLP1 | NETO1 | COL5A2 |
| NECAP2 | SBK1 | FNDC3B | PI4K2A | P3H2 |
| PLPPR1 | CKLF | FAM131A | EIF3B | SMIM3 |
| KIAA0319L | TACO1 | P3H2 | QSOX1 | TRIM35 |
| TOM1L2 | CYB5A | TMEM271 | ARFGAP3 | ANAPC2 |
| CNKSR2 | ADAMTS10 | TNIP2 | RBMXL1 | ZNF33B |
| RCC2 | FIZ1 | TBC1D9 | RAVER1 | MARCHF8 |
| CUL3 | MHENCR | SMAD1 | PIGS | ZBED5-AS1 |
| PPP2R5B | CYTH4 | TMEM184C | SMCHD1 | ZWILCH |
| FBXW5 | CHM | LRBA | CNTNAP5 | PRCD |
| LUC7L3 | TMEM69 | DCTD | ADRA1B | PMAIP1 |
| SEC62 | KCNN3 | TTC33 | AHCTF1 | RDH13 |
| THAP9-AS1 | LGALSL | RIMOC1 | MED12L | FAM182A |
| PPP6C | KLHL23 | SNX18 | MANF | TXLNGY |
| BTBD10 | E2F3 | PDE4D | ZNF91 | SIGMAR1 |
| EMP3 | TRMT11 | ERAP1 | PSEN2 | PUS1 |
| SEPTIN4 | TRPS1 | PGGT1B | PPOX | TPT1-AS1 |
| SELENOP | TLR4 | CNOT8 | CFAP210 | CCNK |
| LINC01963 | SLF2 | RAB24 | TRMT11 | E4F1 |
| PLA2G12A | TMPO | C6orf47 | IGFBP3 | TMEM192 |
| HSPA4L | SNRNP35 | DAXX | ARHGEF10 | EIF3J-DT |
| NIM1K | BLTP2 | TJAP1 | RPUSD4 | SPPL2B |
| VPS26A | DDX52 | SMAP1 | CKAP2 | SLC2A5 |
| TP53I11 | YIPF2 | ELOVL4 | PHF12 | MTHFD1L |
| SPATS2 | PET117 | RPS6KA2 | SMAD7 | DHX57 |
| BORCS8 | ACSS2 | AFDN | LKAAEAR1 | DCAF17 |
| CEP250 | DDX46 | HECW1 | KAT5 | PPARG |
| PHF5A | NELFE | AEBP1 | ENPP2 | RASGEF1C |
| ZNF511 | CHERP | DDX56 | PNPT1 | EXOSC8 |
| AIMP1 | GP1BB | GUSB | RIGI | PPL |
| CPTP | TELO2 | TPST1 | SMAD1 | RABEP2 |
| ACKR1 | PRKAB2 | CACNA2D1 | SNX6 | MPP3 |
| WDFY1 | INPP1 | RBM28 | ANO8 | HAMP |
| CCDC66 | KMT2C | GSTK1 | PDGFRB | PRKD2 |
| ZFYVE28 | EXOSC4 | FASTK | THBS2 | ZNF274 |
| MAPK13 | REEP3 | DLGAP2 | CDS1 | SPIN2B |
| DENND4C | RAB30 | MTMR9 | ZNF704 | ARMCX5-GPRAS |
| LINC01503 | PDE1B | SLC39A14 | BAHCC1 | COL4A5 |
| ANKRD13D | MED4 | EXTL3 | F8 | SIDT1 |

| | | | | |
|---|---|---|---|---|
| GNS | POLG | ZC2HC1A | PIK3R3 | NTNG1 |
| VPS33A | ERAL1 | RALYL | KCNK4 | HINFP |
| TMEM179 | LMAN1 | CIBAR1 | USP35 | MARCHF9 |
| SLC12A6 | PNMA8C | SAMD12 | ANO6 | RRP15 |
| ARHGAP27 | GTPBP1 | KCNQ3 | ERBB2 | NTHL1 |
| MPPE1 | SHISAL1 | RIGI | KARS1 | CDKN2C |
| PIK3C3 | MYLIP | RORB | EIF3D | GUSB |
| JOSD2 | LINC00869 | XPA | RFTN1 | CBLN4 |
| TTPAL | PRELP | TEX10 | GPR176 | POFUT2 |
| PBXIP1 | CPSF3 | PTGS1 | USP3 | SMARCAL1 |
| HOOK3 | FBLN7 | DPM2 | TREM2 | ALKBH4 |
| C1orf216 | TTLL4 | SURF6 | XPO5 | ZNF7 |
| CCDC136 | PLXND1 | MED22 | DDX19B | DERA |
| STX7 | SPON2 | NOTCH1 | STX10 | RPAP1 |
| CRBN | HLA-F | UBTD1 | ARHGAP4 | CSTF1 |
| RNF167 | BLTP3A | NPM3 | POLR3GL | HSCB |
| CHST7 | USP45 | VTI1A | ALMS1 | TAF1 |
| CCDC85A | ABHD11 | BUB3 | MAPK13 | RAB9B |
| ZHX3 | RUNDC3B | CHST15 | CCDC126 | GNAL |
| STX16 | TRIP6 | FUOM | ABHD11 | SGPP1 |
| SOX4 | NAPEPLD | SIGIRR | STAR | RBBP8 |
| SEC13 | INTS8 | DEAF1 | RRS1 | CLDND2 |
| SUPT6H | PLEKHF2 | SERGEF | GLE1 | CENPC |
| ARHGAP33 | PHF19 | LINC00294 | VWA8 | TBCK |
| CDK16 | ST6GALNAC4 | TM7SF2 | TCF3 | NSMCE2 |
| SCAP | ZNF22 | KDM2A | MOB3A | KHDC1 |
| BMPR2 | FAM13C | DYNC2H1 | TIMM29 | SLC22A15 |
| GNA13 | GRK5 | ZBTB16 | DNASE2 | SMC6 |
| CDK2AP1 | MDK | GRAMD1B | ZNF420 | TGFA |
| GNG12 | EMSY-DT | CACNA1C | ZNF615 | ELMOD3 |
| MPZL1 | ESAM | CLEC7A | PAQR3 | SLC20A1-DT |
| MTMR10 | SP1 | CCDC59 | TOPBP1 | CCDC174 |
| SNX5 | YEATS4 | POC1B | APOC2 | CYB561D2 |
| SDHC | TMTC2 | LTA4H | PRDX3 | THBS4 |
| CDC42EP4 | KLF12 | PWP1 | RNF25 | HSD17B8 |
| FDFT1 | RNF31 | DDX55 | TMOD1 | PEX3 |
| SH3GL3 | KHNYN | GALNT9 | GABPB1-IT1 | RNF216P1 |
| AUP1 | VPS18 | GPR12 | ANOS1 | CBLL1-AS1 |
| KIF21A | NKD1 | TDRD3 | ST3GAL1 | DENND2A |
| ARL8A | ZNF778 | RBM26 | ASH2L | FAM131B-AS2 |
| TM4SF1 | MRM3 | DNAJC3 | DOCK10 | FZD6 |
| TM2D3 | TMEM101 | GCH1 | CHIC2 | SSH3 |
| BDP1 | ABHD3 | COX16 | ORC3 | PTPN21 |

| | | | | |
|---|---|---|---|---|
| PNMA8B | MPV17L2 | ACYP1 | FKBP14 | TDP1 |
| SAFB2 | BMP2 | MTA1 | LIF | TNFAIP2 |
| YRDC | INTS6L | SNAP23 | MED14 | PLCB2 |
| NR2F1-AS1 | FBXO30 | RORA | PIGK | CDAN1 |
| HACL1 | HIBCH | BBS4 | TRIB3 | HIC1 |
| MTREX | ZNF213-AS1 | NMB | CDC42SE1 | PFAS |
| ENTR1 | SYNRG | MRPL28 | ABCD4 | PGPEP1 |
| ZFPL1 | LARP4 | RHBDL1 | BRSK2 | KCNG1 |
| BICDL1 | PDLIM4 | UBALD1 | TASOR2 | P2RX6 |
| YLPM1 | ANGPTL4 | EIF3C | UBL4A | TOP3B |
| ZNF592 | PVR | KIF22 | BCL7C | EPHB1 |
| USB1 | ZNF622 | FHOD1 | PPCS | CUTC |
| RANGRF | TARS3 | SF3B3 | SERAC1 | CACHD1 |
| AARSD1 | ACVR1 | IST1 | TRHDE | USP20 |
| ELF2 | IFIH1 | JPH3 | PML | TRIM32 |
| SUGP1 | SERPINB9 | TUBB3 | CBFA2T2 | EIF2AK4 |
| LAMA5 | TRIM26 | RAP1GAP2 | RCC1L | GOLPH3L |
| GPR153 | TMEM8B | CTNS | RRP1 | HCN3 |
| TRAPPC12 | AIFM3 | SPAG7 | CEP41 | LMAN2L |
| LINC00342 | HSF2 | DRG2 | TOX | ZNF346 |
| RABL2A | GPBP1 | MED1 | KLHDC9 | BLTP3A |
| TATDN2 | HDHD2 | PLCD3 | LSM5 | GET4 |
| MRPS22 | NETO1 | COX11 | C6orf120 | ZKSCAN5 |
| PPARGC1A | LGALS8 | DHX40 | NUBP2 | MET |
| PPID | AKAP8L | USP32 | PPT2 | MSANTD2 |
| DROSHA | ASMTL | ALYREF | STAG3L4 | SLC38A6 |
| KDM1B | CLSTN2 | PTPRM | PDCL | MAPK7 |
| NKAIN2 | LNX1 | CXXC1 | DLST | A1BG-AS1 |
| CRCP | NR1D1 | RELCH | UNC50 | IFNAR2 |
| SBDSP1 | FABP3 | KLF2 | DPCD | PABPC1L2B |
| APTR | EPAS1 | SLC25A42 | DRG2 | LINC01003 |
| VPS50 | TPM4 | SNRPA | ZNF639 | CASP9 |
| KIAA1549 | PRKAG2 | ZNF470 | FAT1 | STK36 |
| KAT6A | HAUS2 | ADISSP | STX8 | MIER2 |
| RNF170 | CLIP4 | GPCPD1 | CBR3 | GNE |
| TMEM65 | BCL11A | CBFA2T2 | TXNDC15 | LAMB1 |
| PDCL | PLXDC2 | ITCH | ARL4A | SORCS3 |
| DDX51 | DNAJC12 | PTPRT | LHFPL4 | TAFAZZIN |
| TTC9 | AIP | VPS26C | TUBGCP2 | ACKR3 |
| THAP11 | RALGAPA1 | TUBA8 | AKR7A2 | SLC4A2 |
| ST3GAL2 | HEXA | PATZ1 | AJM1 | CD24 |
| EPS15L1 | FSD1 | HMGXB4 | GLA | PRMT9 |
| POU2F2 | TRNP1 | PARVB | TBC1D14 | NINJ2 |

| | | | | | |
|---|---|---|---|---|---|
| SAMHD1 | AHDC1 | SLC9A7 | PEX11B | | ZNF677 |
| RALGAPB | ANP32E | FTSJ1 | GATAD2A | | TFCP2 |
| IFT27 | STX6 | GRIPAP1 | EPS8 | | GPR180 |
| DRP2 | TARS1 | RBM41 | ZDHHC20 | | TMTC4 |
| DPP10 | TNFRSF21 | PSMD10 | CALN1 | | |
| NUMBL | HTR2A | NSDHL | ZBTB1 | | |
| TMEM41B | MINDY2 | ATP2B3 | EMC9 | | |
| SH3PXD2A | RNF40 | VAMP7 | C2orf69 | | |
| PEPD | SPATA20 | USP9Y | LRRN3 | | |
| NECAB3 | SSTR2 | EIF1AY | SBK1 | | |
| NCKIPSD | UQCC1 | EPRS1 | CPTP | | |
| WIPF2 | TNIP1 | COX17 | EPHB2 | | |
| DRG1 | PLPP7 | SELENOF | ZBTB47-AS1 | | |
| HIPK3 | BBIP1 | TSPAN13 | TP53TG1 | | |
| SF3B4 | POLR3E | SARNP | ERCC6 | | |
| KHDC4 | ENDOV | DGCR6L | SRGAP1 | | |
| LMBRD2 | RRM2B | HTATSF1 | COG7 | | |
| TCAF1 | C2CD3 | LDOC1 | ZNF668 | | |
| RASGEF1A | SYNGR2 | IL17D | B9D2 | | |
| RUVBL2 | TPRKB | TRIM37 | PNPLA4 | | |
| SNHG17 | STAM2 | GNG2 | MARK2 | | |
| ABTB1 | RBMS3 | GPRASP1 | EP400 | | |
| LONRF1 | SLIT3 | PAK3 | VPS45 | | |
| ZBTB43 | LEMD2 | NEGR1 | LINC01511 | | |
| PLS3 | SLC17A5 | PSD3 | PLEKHG5 | | |
| SEC11C | UST | NSFL1C | BTAF1 | | |
| AP3S1 | TMEM243 | RNF7 | RNF111 | | |
| FAR1 | KRIT1 | HNRNPUL2 | B3GALNT2 | | |
| SHISA5 | NBN | SMARCA2 | ABHD5 | | |
| DMXL2 | EBAG9 | NRSN1 | FRMD6 | | |
| EEA1 | ZHX2 | HIVEP2 | SNX13 | | |
| WASHC2A | TUT7 | KCNC1 | DNAJC25 | | |
| IER3 | ASB6 | SAFB | DDX49 | | |
| FAM217B | ECD | RFPL1S | FUNDC2 | | |
| DCUN1D5 | CCKBR | C1orf52 | PSRC1 | | |
| PITPNM2 | PHF21A | FDPS | HINT2 | | |
| SMIM10L2B | SNX15 | LONRF2 | UPF3A | | |
| UBL4A | SLC36A4 | CCSER2 | CRYBB2P1 | | |
| GNAI2 | DSTNP2 | MYCBP2 | LRRC4C | | |
| PTRHD1 | NR2C1 | PSMA4 | CKLF | | |
| FAM163B | ZMYM5 | NUDT21 | SMC1A | | |
| PSMD14 | FMN1 | CDH13 | WDR35 | | |
| VPS26B | MSRB1 | NHERF1 | WDR43 | | |

| | | | |
|---|---|---|---|
| JPX | NOB1 | ODC1 | CUL7 |
| CHD3 | SUZ12 | MATK | BAX |
| RIF1 | UNK | RUFY3 | UBFD1 |
| HYI | ANKRD29 | SLC44A2 | CDK11B |
| UTP18 | NFATC1 | PGRMC2 | POU3F1 |
| GRM3 | LINC00662 | SPCS3 | TMEM198 |
| DDX54 | IL4I1 | PACSIN1 | NEK10 |
| DLGAP3 | IL11 | PPP2R2A | UFSP2 |
| PCYOX1 | ESF1 | CHPT1 | PGGT1B |
| CREB3 | PI4KAP1 | PSMA3 | CEP120 |
| ISG20L2 | PIK3IP1 | LDLR | ZKSCAN8 |
| KCNH1 | LCP2 | ZSCAN18 | THAP1 |
| TPRA1 | OGFR | PPP3R1 | MIRLET7A1HG |
| THBS2 | PTPRG | MYO10 | ABHD13 |
| WDR91 | PPIB | GPC1 | HEATR5A |
| NDST2 | PLXNA1 | PLXNB1 | DTD2 |
| C10orf95-AS1 | ENDOD1 | RPN1 | MAP3K3 |
| CHRM4 | RBP1 | RNF4 | NEXMIF |
| MSANTD4 | TREM2 | NEU1 | MTHFD2 |
| HINFP | TFPT | DOCK9 | PRKAB2 |
| RGS6 | PHACTR3 | TMEM147 | FASTK |
| RRN3 | METTL9 | RBM42 | MIIP |
| SPNS1 | FUNDC2 | PTPRN2 | ZNF680 |
| TIMM22 | C2CD2L | APBA2 | IQCK |
| ERN1 | FAM131B | SYNGR1 | RER1 |
| THOC1 | ZNF703 | PKIG | CDH2 |
| INSR | ZNF281 | PSMB7 | DAGLA |
| HPS4 | CARMIL1 | HMGN1 | MTMR14 |
| H1-0 | TERF1 | CAPZA2 | MX1 |
| STEEP1 | SNHG10 | GFOD1 | RMDN3 |
| ATG5 | TNFAIP8L3 | GLRX | TYW3 |
| FNDC5 | WWP2 | TJP2 | NT5E |
| MANEAL | AKAP1 | CTNNB1 | RBAK |
| ANKRD36 | HCST | SSBP3 | TMEM132C |
| CNPPD1 | MRPL40 | TMED5 | SHISA8 |
| ARL6 | ARSA | CHMP3 | SLC1A1 |
| RIOX2 | TXNIP | LMBRD1 | SYNPO2 |
| IFT122 | STAG1 | RANGAP1 | HLA-DQB1 |
| SDHAP4 | DGLUCY | LHFPL2 | MTA2 |
| TAPT1 | AGO1 | RAB18 | SARS2 |
| RNF150 | MBOAT2 | MOB1A | GPATCH3 |
| LINC01511 | WBP1 | MSN | GSTM2 |
| SPEF2 | ZNF771 | RAB5C | SRGAP2B |

| | | | |
|---|---|---|---|
| GFPT2 | FAM47E | GRIA2 | CDH12 |
| SLC35B2 | TRAPPC14 | SV2A | CASP8AP2 |
| MTO1 | RSPO2 | OPALIN | IL11RA |
| FAM229B | ABCC8 | RABEP1 | MIR1915HG |
| BRAT1 | FBXO27 | LPAR1 | IDE |
| TMEM196 | PVALB | RGS1 | WEE1 |
| SLC25A40 | HAUS7 | NBL1 | LIPT2-AS1 |
| CALD1 | TMEM11 | RAB5A | C11orf54 |
| SLC1A1 | FBLN1 | ACTN4 | ALOX5AP |
| CD274 | TUBB6 | CLIP1 | TVP23A |
| GAS1 | ZCCHC2 | CCNL1 | NKD1 |
| URM1 | TFIP11 | NDUFA7 | CCL8 |
| SPOUT1 | PIM2 | SEC23B | HEXIM2 |
| RPP30 | VKORC1 | SND1 | ZNF154 |
| TSSC4 | PNPO | TRAM1 | EDEM2 |
| FOXRED1 | RING1 | ABCA1 | ZBTB46 |
| GPR19 | LAMTOR3 | HSDL2 | EEPD1 |
| CSAD | SMIM7 | CTR9 | F8A1 |
| GLS2 | PCSK1 | FADS1 | PTRHD1 |
| AP5M1 | MRPS7 | ATM | DHX36 |
| PCNX4 | WDR7 | ARFRP1 | KLC4 |
| ZNF423 | TMEM131 | NCS1 | MFSD5 |
| OSGIN1 | FOXN3 | SLC25A11 | DPH2 |
| IGFBP4 | CAMSAP1 | PPT1 | ARNT |
| CCDC159 | BST2 | ARID1A | PDIA4 |
| ZNF420 | NABP1 | SERTAD2 | NHLRC2 |
| GMEB2 | FAHD1 | UQCC5 | ZNHIT2 |
| NUP50-DT | SIKE1 | GNA12 | TCHP |
| RTCB | TFPI | SPTLC1 | TRAFD1 |
| ERGIC3 | ATXN7 | PTBP3 | RTBDN |
| ITPRID2 | KCNIP1 | PDHX | CUL4A |
| ACTR3B | WHRN | TBC1D15 | LYN |
| DDX27 | KATNAL1 | STON2 | MLLT10 |
| UBP1 | CPNE7 | PDXDC1 | NUCB2 |
| HSF1 | ZNF540 | SNHG16 | ASPHD2 |
| MED17 | ZNF542P | RPS6KA3 | BRCC3 |
| SYNDIG1 | MX2 | SLCO1A2 | DDHD2 |
| MTRR | PKNOX1 | ARHGAP26 | PPP1R35 |
| ICMT | DGCR6 | KDELR2 | WDR83 |
| GMPS | STRIP2 | CIAO2B | HERC3 |
| RXRB | DLG5 | RGS16 | IQCB1 |
| CAPN1 | SYT16 | UGGT1 | TENM3 |
| HDAC7 | NTSR2 | SUPV3L1 | PENK |

| | | | |
|---|---|---|---|
| CDC16 | CPNE2 | PTPRK | FZD8 |
| DTX3 | TNFRSF1B | SGCE | PCNX3 |
| TIMMDC1 | IDH1 | PLOD3 | AKAP5 |
| SLC11A2 | NKTR | ERBB3 | OSBP2 |
| ARRDC4 | TAFA1 | ZBTB1 | STAT6 |
| PPP1R14A | RCHY1 | RPA1 | KDELR1 |
| GSK3B | MAN2A1 | GPS1 | ECE1 |
| ATP5F1C | CDH11 | CIT | MAN1B1 |
| EMC10 | PTPN4 | NLK | WIZ |
| SMARCD3 | STX10 | NUFIP2 | DMTF1 |
| NIBAN2 | NTPCR | CADM4 | EMX1 |
| MYO1E | RNF38 | SPART | DLGAP3 |
| USP16 | PADI2 | SSR4 | MRPL53 |
| BAIAP3 | ACVR1C | DAB2 | MSANTD3 |
| GDAP1L1 | CNPY3 | COA8 | CEP57 |
| WDR43 | C9orf72 | DYRK1A | DMAP1 |
| APBB1IP | BICD2 | USP33 | BCL10 |
| EIF5AL1 | SDHAF2 | CLNS1A | DDAH2 |
| RASSF8-AS1 | ATF7IP | ADAR | CLPX |
| CGRRF1 | SYMPK | C1orf198 | CALCOCO2 |
| TBC1D20 | FAT3 | PUF60 | LARP1B |
| HCCS | SLITRK4 | PRPF8 | AGO4 |
| MFSD4A | IDNK | CERS2 | CHST2 |
| CACNG3 | LUZP1 | DNMT1 | MTX3 |
| SERPINB8 | RASA3 | NUP62 | NUDT2 |
| SLC48A1 | STIM2 | NCOA7 | EIF5AL1 |
| NTM | SSPN | NANS | EEA1 |
| SPOCK1 | SLC66A1 | NDUFS1 | MED1 |
| RBM25 | ZNF362 | RNF5 | RNASEH1 |
| RWDD1 | LRRC40 | ZER1 | TM9SF4 |
| GLDN | GRM7 | FGFR2 | PTAR1 |
| FAM13B | TTLL3 | CRIP2 | KCTD5 |
| H2AC25 | TCAIM | TPM1 | HLA-L |
| TMEM158 | NCK1 | ATP2B4 | ZDHHC8 |
| CDC42EP3 | HSD17B11 | DCLK2 | KLHDC8B |
| CCT3 | RARS1 | PMVK | FILIP1L |
| PDZD8 | CASC15 | NAA50 | GPNMB |
| ZIC2 | CYREN | DLD | RCE1 |
| LIMS1 | FAM131B-AS1 | ALKBH5 | TDRD9 |
| C1QC | TGS1 | GAMT | CYB5D1 |
| SCN3A | ZNG1C | AP3D1 | SHISA6 |
| ADAP1 | VAV2 | SMARCA4 | SPAG5 |
| AP3M2 | TMEM132C | DAD1 | ERP44 |

| | | | |
|---|---|---|---|
| TXLNG | ZBTB25 | ARL2 | NKAIN3 |
| PYGO2 | RNF112 | IGFBP7 | PTPRU |
| GTF3C2 | SMARCD2 | DNM1L | CUX2 |
| TMEM198 | ESCO1 | WASF3 | VCPKMT |
| DUSP7 | RNF152 | CCT6A | PCMTD2 |
| TRPC1 | RBFA | SF3B5 | RAB39B |
| IL1RAP | LTBP4 | UBE2V2 | OSBPL6 |
| SH3D19 | SPEF1 | ACTN1 | BBX |
| CDH10 | ZXDB | CAST | PDLIM4 |
| GNE | TRIL | TNS3 | TUSC1 |
| RNASEH2B | SLC35A5 | AKAP9 | RAB35 |
| FAM124A | TSPAN33 | ADAM9 | ATXN7 |
| MEX3B | CLIP2 | NEK6 | MCM7 |
| PDIA2 | PRDM2 | DDT | PEX2 |
| ZNF136 | MESP1 | MACROH2A1 | PIGA |
| TBC1D10A | WDR24 | TSN | EIF2B5 |
| RNF113A | RCE1 | NFKB1 | NNT-AS1 |
| NEPRO | GPATCH3 | NIPA2 | UQCC6 |
| POLD4 | FAM229A | DNM2 | BCL9 |
| NEFH | SLC22A15 | CHL1 | CRHBP |
| SCNN1D | LMAN2L | LMNA | ACHE |
| OSCP1 | LIMS2 | POLR2I | NUP160 |
| LINC02607 | PARP3 | PHGDH | STARD13 |
| CYB561D1 | RUVBL1 | OGA | ZNF436 |
| SMG5 | PAQR9 | MFF | ZDHHC23 |
| RXRG | CTBP1-DT | ENHO | GKAP1 |
| MOGS | SV2C | INPP5F | PSMD5 |
| ZNF385B | PCYOX1L | STAT1 | KLHL35 |
| RBSN | PI16 | CMPK1 | IGFBP4 |
| MLH1 | AARS2 | RETREG2 | RIPK2 |
| VIPR1 | EPM2A | FOXG1 | PTGR1 |
| RWDD4 | MCPH1 | DLGAP4 | SLC39A12 |
| BCKDHB | UBXN8 | MECP2 | HELZ |
| RNGTT | RBM12B | HES4 | ADISSP |
| GARS1-DT | SPAG1 | TMEM222 | SYMPK |
| TYW1 | ZNF34 | EFNA1 | SEPTIN6 |
| COL26A1 | NOL8 | CRACDL | CPNE4 |
| GPR37 | TBC1D2 | SRPRB | ANKRD29 |
| ADCK2 | ZEB1-AS1 | GAK | EPM2A |
| ZFPM2 | DEPP1 | PLEC | PTPN14 |
| JAK2 | NUTM2B-AS1 | NRXN2 | RNPEPL1 |
| SIGMAR1 | LIPT2-AS1 | PPP1R12A | NUP155 |
| SOHLH1 | AQP11 | ITFG1 | BAK1 |

| | | | |
|---|---|---|---|
| MASTL | RNF26 | BCAR1 | NHSL1 |
| BMS1 | GNB3 | CXCL16 | GTF2IRD2 |
| LZTS2 | MDM1 | RAB34 | TRIM22 |
| YPEL4 | VIPAS39 | JUP | RFX7 |
| ARAP1 | DICER1-AS1 | ZFR2 | NUBP1 |
| STX2 | CLMN | SNHG12 | PEX12 |
| PRKCH | KNOP1 | RBBP4 | BMI1 |
| CPEB1 | RPGRIP1L | IFT25 | MICU2 |
| NPIPB13 | FAM222B | ANKRD39 | HRH3 |
| ASPA | TBC1D16 | DHX36 | CAVIN1 |
| LRRC75A | ENOSF1 | NSD2 | PPA2 |
| GRIN2C | PLVAP | BOD1L1 | ATMIN |
| TIMM21 | ARMC6 | SLAIN2 | KCNJ10 |
| OPA3 | ZNF570 | LIFR | TMEM196 |
| POLR3F | ZNF224 | GRAMD2B | MX2 |
| RALGAPA2 | A1BG-AS1 | TMEM151B | SLC43A2 |
| CDK5RAP1 | NFATC2 | ST7 | PLIN2 |
| SOX10 | TBL1X | C8orf33 | SEMA5B |
| TAB1 | HS6ST2 | MEGF9 | SFXN4 |
| COL4A5 | ABCD1 | NET1 | KRCC1 |
| CRB1 | SLC25A33 | EIF4EBP2 | TMEM161B |
| BOK | KCTD2 | FRA10AC1 | PPP1R3C |
| WWTR1 | FBXO45 | GLRX3 | MTG1 |
| TMEM41A | FGF2 | IRAG1 | TMEM223 |
| ZNF827 | BMPR1B | ESRRA | NEU3 |
| MTX3 | FZR1 | PPP6R3 | RECQL |
| ILRUN-AS1 | GNG11 | CLEC2B | ANAPC7 |
| ARHGAP12 | FAM151B-DT | NOC4L | PNMT |
| MTRFR | ZDHHC24 | LINC00641 | SLC25A19 |
| LINC00507 | NDUFAF2 | NAA30 | DUS3L |
| NRDE2 | GMPR2 | SNX1 | TGM2 |
| PLA2G4B | DFFA | TBCD | RGS19 |
| TMOD3 | COMMD1 | CYBA | COL11A1 |
| HMG20A | NFU1 | TRAP1 | GPRIN3 |
| CIB2 | ZC3H6 | SESN2 | PLAC9 |
| MYL11 | CIR1 | PGM1 | MORN1 |
| ZNF768 | MB21D2 | MIER1 | CSMD2 |
| RNF166 | SDHAF4 | SEC22B | EFCAB7 |
| DNASE2 | ASF1A | FCGR1A | C2orf76 |
| ZNF579 | FAM167A | ATL2 | TAMM41 |
| CHRNA4 | DDX25 | SOS1 | COL7A1 |
| RRP1B | FASTKD5 | RBMS1 | HYAL2 |
| LRCH2 | FUNDC1 | DNAJC10 | CBLB |

| | | | |
|---|---|---|---|
| TMEM185A | TUBA1C | MSL2 | ANAPC4 |
| RIMKLA | CHMP4A | COMMD2 | ZNF138 |
| DOT1L | SOBP | PCYT1A | FAM131B-AS1 |
| FBL | YTHDF1 | FGFRL1 | DEPTOR |
| GTF3C3 | EXOC3 | TMEM161B-DT | CHRAC1 |
| EGFR | PPRC1 | UBE2J1 | PACSIN3 |
| PTRH1 | SLC15A2 | REPS1 | RASSF3 |
| BAMBI | TAF15 | TMED4 | DHRS2 |
| SAC3D1 | ITPR1 | CHCHD3 | CENPN |
| GALK1 | CHMP1A | LSM1 | CEP76 |
| MED14 | DAG1 | PLEKHA2 | ZNF582 |
| PSMC1 | KAT8 | ARMC1 | ZFP64 |
| CCSAP | DOC2B | SDC2 | SLC38A5 |
| SVBP | ZFP82 | FAM120A | TIMM8A |
| ICA1L | SERPINB1 | GTF3C5 | LRFN5 |
| CCR1 | OGDHL | DDIT4 | FKRP |
| WDR6 | DERL1 | RAB11FIP2 | PAICS |
| MAP9 | GUCD1 | ATE1 | ADAM11 |
| ANKRD33B | ZFYVE9 | NAV2 | PXMP2 |
| POLR1C | CMPK2 | PELI3 | FLAD1 |
| KLHDC10 | ASXL2 | SLCO2B1 | TRIB2 |
| RELA | INO80B | CBL | EARS2 |
| EED | KANSL3 | SRPRA | NFKBIZ |
| TRAF3 | BCL2L11 | ERGIC2 | FAM234A |
| EXOSC5 | ITGA6 | IPO8 | LIN37 |
| SRC | NELFA | SLC4A8 | SLIT1 |
| WRAP73 | MAN2B2 | CDK4 | TSPAN31 |
| DDX20 | GALNT7 | ZC3H13 | SERPINB1 |
| IGSF3 | SLF1 | DIS3 | LRFN4 |
| MSTO1 | BTN2A1 | NGDN | FIP1L1 |
| INTS7 | MCM3 | BRF1 | USP42 |
| CNNM3 | NT5DC1 | GREM1 | NDUFAF6 |
| LINC01102 | AASS | RMDN3 | NCBP3 |
| TANC1 | TEK | EHD4 | TLN1 |
| PPARG | DCAF12 | PML | RPL23AP7 |
| SEMA3F | SLC25A16 | MESD | LRRC49 |
| ACTR8 | GFRA1 | BOLA2B | SPSB3 |
| CEP70 | TMEM216 | N4BP1 | MAP3K4-AS1 |
| ALG3 | MAP4K2 | EDC4 | TSPAN33 |
| INPP4B | CEP295 | FA2H | TOR1B |
| SNX25 | MCAM | MAF | PAMR1 |
| SLC12A7 | GRIK4 | ZCCHC14 | TRIP11 |
| PTGER4 | LMBR1L | STX8 | SYVN1 |

| | | | |
|---|---|---|---|
| TBC1D7 | ASCL1 | MED24 | LEPROTL1 |
| FGL2 | USP30 | MPP2 | DDX27 |
| LRRC61 | TNFAIP2 | SLC25A39 | CHADL |
| ADHFE1 | GABRG3 | RTTN | USP21 |
| MRRF | MTFMT | GET3 | SLC11A1 |
| SAPCD2 | FAM234A | HNRNPUL1 | GRPEL2 |
| MINPP1 | JMJD8 | MYH14 | ISCA2 |
| EIF1AD | DDX19B | XRN2 | SARM1 |
| FOLR2 | LIG3 | TRPC4AP | RALGAPA2 |
| CCDC90B | FKBP10 | MIR99AHG | CYTH4 |
| NCAPD3 | MRPL58 | TIAM1 | IL13RA1 |
| PAWR | PPAN | ADA2 | SLC6A7 |
| ZFYVE19 | KCTD15 | KREMEN1 | TRAF7 |
| UBAP1L | HAMP | NUP50 | LNPK |
| CIAO3 | SYS1 | TBC1D22A | WDR19 |
| MVP-DT | SNX21 | PLCXD1 | DMPK |
| COG8 | SCAF4 | PPM1L | MMACHC |
| IRF8 | ARMCX4 | GABRA2 | LNP1 |
| PFAS | TAGLN | PHF24 | MCF2L2 |
| IFI35 | GDPD1 | KCNMA1 | MASP1 |
| ACBD4 | MRPS15 | CCPG1 | MB21D2 |
| MFSD11 | CPNE3 | RNF103 | CXXC4 |
| CEP76 | AJM1 | CACNB4 | C7orf25 |
| GTPBP3 | SLITRK5 | KIDINS220 | MMP16 |
| ZNF575 | PHKG2 | PIM1 | GFRA1 |
| BLOC1S3 | ATOSB | SLC35F1 | HHIPL1 |
| UBOX5 | ABL1 | WASL | SLC5A11 |
| BGN | SLC27A1 | TXN2 | C17orf58 |
| ARHGAP4 | SLC9B2 | HIPK1 | CNBD2 |
| AJAP1 | LSM8 | PGBD5 | KIAA1755 |
| PYROXD1 | SMC3 | NRBP2 | ZNF217 |
| ETFB | HEBP1 | C11orf68 | DYNC2LI1 |
| RRAGC | ENG | ARRB1 | ARAP1 |
| MADD | ZCCHC17 | AARS1 | DNAJC27 |
| PHC2 | CHD9 | SRP68 | CHCHD7 |
| PLPP5 | GOLIM4 | PALM | GRM5 |
| FN1 | SRGAP2 | PSENEN | NR3C2 |
| HSPA6 | GABRA4 | GNAZ | PLOD3 |
| CSNK2A1 | PFKFB2 | PRPS1 | FGD4 |
| TWSG1 | ZIC1 | TSC22D3 | ROCK1 |
| RMND5B | SRPK1 | ARHGAP35 | PRPF18 |
| PRPSAP1 | P2RY1 | PSMA2 | TAOK3 |
| MAOA | RGS10 | SMOX | ZFPL1 |

| | | | |
|---|---|---|---|
| HCFC1 | CDK11B | SDHB | COLGALT1 |
| SMAD5 | NDC1 | ZRANB2 | SDE2 |
| AIG1 | CRYZ | CSRNP3 | NT5DC2 |
| PDE1A | ALG14 | CERS6 | INSR |
| MRPS5 | SEMA6C | RGS12 | DISP1 |
| BAZ2B | PIK3C2B | ACSL6 | GRAMD1C |
| MMGT1 | SH3BP5L | MRPL14 | PLXND1 |
| THUMPD1 | PRKD3 | TRA2A | TADA2B |
| RDH10 | PCBP1-AS1 | NSD3 | KLHL32 |
| FGF14 | SLC20A1-DT | STAU2 | CSMD1 |
| DGUOK | WDSUB1 | BRINP1 | LRRC10B |
| EHD1 | CALCRL | DPP7 | MEIS2 |
| STRN4 | TAMM41 | CARHSP1 | SLCO4A1 |
| OXA1L | XIRP1 | USP7 | MIGA2 |
| RGS11 | GBE1 | GGNBP2 | ID1 |
| USP25 | ACAD11 | MBOAT7 | NRSN1 |
| PER2 | HAUS3 | EPB41L1 | HECW2 |
| SNAP47 | LCORL | SREBF1 | MBD2 |
| PLPBP | SEL1L3 | FBXL17 | PIK3CA |
| TOR1A | PGM2 | CD2BP2 | GRIK2 |
| MED19 | CRACD | SERTAD3 | NUP62 |
| GPR176 | TIFA | NAT14 | TST |
| THOP1 | C4orf46 | IDH3B | TASOR |
| MAU2 | DHFR | TMEM230 | MAMLD1 |
| WDR45 | BAG2 | U2AF1 | ZDHHC9 |
| LAS1L | BMPER | SMIM10L1 | NECTIN3 |
| EPHB2 | NSUN5P1 | RAB11A | CSPP1 |
| TAF12 | SEMA3E | GIT1 | SLCO1C1 |
| SNRNP40 | PRKRIP1 | SLC25A36 | TMPO |
| ENTREP3 | NKX3-1 | AGRN | CMBL |
| BRINP3 | GTF2E2 | ANKRD65 | CCZ1B |
| CNTNAP5 | WRN | ATAD3B | TM6SF1 |
| SMIM8 | PDE7A | THAP3 | CYBA |
| CCDC126 | MAILR | UBIAD1 | ALDH4A1 |
| ING3 | LURAP1L | PTPRU | EIF2B3 |
| NCBP1 | PCSK5 | PPIE | GBP2 |
| PHLDB1 | SMC2 | TIE1 | PRELP |
| CDK8 | CDHR1 | BEND5 | ACAP2 |
| FAM161B | TAF5 | RAB3B | CCDC85C |
| INO80E | KCNQ1OT1 | CC2D1B | RTL10 |
| TMEM91 | ATL3 | LRP8 | SNX11 |
| BBC3 | MAP3K11 | USP24 | ASPSCR1 |
| OPRL1 | FADD | PRKAA2 | PCED1A |

| | | | |
|---|---|---|---|
| CHPF2 | MSANTD2 | USP1 | GCAT |
| MRPL46 | RBMS2 | SSX2IP | EXOC8 |
| ADCYAP1 | BBS10 | ARHGAP29 | SH3YL1 |
| PEX10 | COQ5 | GPR88 | CFAP90 |
| ZC3H12A | ACADS | OLFM3 | DXO |
| EVI5 | POLE | PRMT6 | GSDME |
| SEMA4A | MIR4500HG | AMIGO1 | GFUS |
| FLVCR1 | KLHL28 | DRAM2 | PITPNM1 |
| MTARC1 | ADCK1 | CD58 | GTF2F2 |
| QPCT | DIS3L | GDAP2 | SNAP29 |
| MSH2 | SGF29 | POLR3C | MTMR3 |
| VPS54 | ZNF688 | PIAS3 | SMIM20 |
| CYP26B1 | CENPN | NBPF12 | EIF2B1 |
| MCM6 | SNHG20 | CHD1L | FARP2 |
| WDR12 | ASXL3 | GPR89B | NSMCE4A |
| USP19 | SLC7A10 | SETDB1 | HS3ST4 |
| CACNA2D2 | ZNF565 | RFX5 | MMADHC |
| RASA2 | SPRED3 | RIIAD1 | MMP17 |
| STX18 | EHD2 | SNAPIN | RASSF2 |
| NR3C2 | BCAT2 | SLC27A3 | MINDY1 |
| F12 | ZNF577 | CKS1B | SNHG28 |
| KIF13A | PIGP | EFNA3 | PIGM |
| TRIM39 | LRRC75B | RUSC1 | TTC13 |
| FAXC | XPNPEP3 | DUSP23 | KCNG3 |
| YAE1 | ARAF | KLHDC9 | MCEE |
| CROT | EFNB1 | CCDC181 | FBLN2 |
| LRCH4 | TMEM192 | VAMP4 | TBCCD1 |
| UBN2 | AQR | COP1 | LGI2 |
| KCTD9 | RDX | NMNAT2 | SHROOM1 |
| TOX-DT | ADM | TRMT1L | HLA-DQA1 |
| PLGRKT | RB1 | ELK4 | ERMARD |
| ANKS6 | VIP | RCOR3 | PSPH |
| ADAMTSL2 | BEGAIN | ANGEL2 | TFPI2 |
| COMMD3 | RTF2 | BPNT1 | FBXO16 |
| MPP7 | THBS1 | MIA3 | TOR2A |
| TCERG1L | FNTB | NVL | BDNF-AS |
| RASSF7 | NFKBIZ | CNIH3 | LINC02716 |
| NELL1 | KDM7A | TMEM18 | RCOR2 |
| ANO5 | ANO6 | KLF11 | FLRT1 |
| UNC93B1 | NBR1 | UBXN2A | KCNK10 |
| CAPN5 | SAMD4A | SLC35F6 | FES |
| NUDT15 | IL6 | KHK | ADCY7 |
| CKAP2 | MARK1 | PREB | SLC7A6OS |

| | | | |
|---|---|---|---|
| CLN5 | MTHFS | FNDC4 | TMEM101 |
| C14orf119 | TFDP2 | SANBR | LINC01315 |
| DCAF4 | LCNL1 | GGCX | RTL8B |
| RPS6KL1 | RGN | MRPL35 | PCID2 |
| NGB | CD46 | CNNM4 | CARMIL2 |
| CCDC85C | MYRF | SOWAHC | ARHGAP20 |
| TM6SF1 | SOCS5 | ZC3H8 | ADAMTS4 |
| CFAP119 | TSPO | MAP3K2 | IQCE |
| ZZEF1 | LARS2 | PTPN18 | CTPS1 |
| KSR1 | BCAS3 | AMER3 | DLX1 |
| TRAF4 | | RND3 | IRAK2 |
| PNMT | | TNFAIP6 | CDKAL1 |
| HAP1 | | KCNJ3 | HR |
| VAT1 | | OSBPL6 | SLC25A16 |
| CCDC43 | | PRKRA | CUEDC1 |
| MTCL1 | | ZNF804A | SMTN |
| RFX2 | | KLF7 | BCS1L |
| ZNF653 | | UNC80 | HERC4 |
| SDHAF1 | | RPE | HS2ST1 |
| XRCC1 | | ATIC | NSUN5P1 |
| SH3BGR | | SLC4A3 | SNAI3-AS1 |
| ZC3H12B | | CT75 | FUCA1 |
| UTP14A | | DIS3L2 | RPS6KA1 |
| CBX1 | | EIF4E2 | KTI12 |
| WSB1 | | HDAC4 | NDC1 |
| PTTG1 | | THUMPD3 | COLGALT2 |
| POLR2K | | MTMR14 | KCNS3 |
| RNASE1 | | CAMK1 | CNNM3 |
| CREG1 | | RPUSD3 | VILL |
| TMEM151A | | RAF1 | SEL1L3 |
| DOCK10 | | CHCHD4 | TLR2 |
| SNRPF | | METTL6 | GDNF |
| PPP1R9B | | WDR48 | EPHA7 |
| PPP1R18 | | MYRIP | MBLAC1 |
| SEMA3B | | ABHD5 | ZNF169 |
| DTD1 | | TMEM42 | CCDC34 |
| NAP1L2 | | SACM1L | PRDM11 |
| ZCCHC10 | | RAD54L2 | FEN1 |
| SOD3 | | ACY1 | IGHMBP2 |
| YAP1 | | CHDH | CASP4 |
| MGP | | ABHD6 | DCAF4 |
| KPNA3 | | EOGT | AP4E1 |
| LEO1 | | ROBO2 | METTL16 |

| | | |
|---|---|---|
| NCBP3 | TMEM39A | KCNJ12 |
| ZNF217 | ZNF148 | MYO1D |
| TUBGCP6 | UBA5 | ADGRE5 |
| SGSM2 | PCCB | ZNF8 |
| PHLPP1 | DBR1 | TTI1 |
| WDR41 | MFSD1 | PTPRT-DT |
| ACOT2 | B3GALNT1 | XPNPEP3 |
| TP53RK | PHC3 | IFITM1 |
| SH3KBP1 | UBXN7 | LINC01007 |
| ALKBH6 | LRCH3 | KLHL17 |
| PPP1R21 | MFSD10 | DHRS3 |
| TBR1 | SORCS2 | TMEM69 |
| CCM2 | ABLIM2 | ITGB3BP |
| DHRS4L2 | CC2D2A | ADGRL4 |
| PLXNA3 | PCDH7 | TARS2 |
| DYRK3 | SMIM14 | HK2 |
| LPGAT1 | NOA1 | GALNT13 |
| LRP3 | DCK | KANSL1L |
| PLCG1 | SDAD1 | SLC9A9 |
| SLC1A4 | CXCL10 | C4orf33 |
| PCM1 | NUP54 | ERCC8 |
| LINC00511 | CNOT6L | SLC25A48 |
| ANKRD37 | PTPN13 | UST |
| CCDC74A | HERC6 | PAXIP1 |
| MON1A | PPA2 | PINX1 |
| LRRC58 | SEC24B | CNTNAP3B |
| CBR4 | ELOVL6 | MINPP1 |
| COMMD5 | TENM3 | ZNF408 |
| LRTM2 | CASP3 | PCED1B-AS1 |
| TRPM7 | CMBL | MDM1 |
| CORO7 | NUP155 | TMCC3 |
| SEC14L5 | GDNF | MSI1 |
| SSH2 | OSMR | CIAO2A |
| ZNF211 | RICTOR | CLN3 |
| WFDC2 | MRPS30-DT | C16orf95 |
| SLC25A14 | PARP8 | WNT3 |
| TMEM97 | HSPB3 | RBBP9 |
| SLC37A3 | DHX29 | SLC10A3 |
| SNRPC | PLPP1 | PIGC |
| TCEAL8 | LNCBRM | CPSF3 |
| POLK | DIMT1 | MCM3 |
| SEC23A | SLC30A5 | GID4 |
| KEAP1 | MRPS27 | TSPAN6 |

| | | |
|---|---|---|
| COBL | GCNT4 | FAM13A |
| CRIM1 | JMY | ZNF770 |
| MYH10 | SSBP2 | FGFRL1 |
| ATXN2L | MACIR | TXNDC9 |
| ADRA1B | TRIM36 | ZNF592 |
| DUS1L | CSNK1G3 | NOL11 |
| DDR2 | MEGF10 | TMEM214 |
| LAMC1 | CDKN2AIPNL | MYO16 |
| SGTB | JADE2 | FAT3 |
| PLLP | APBB3 | HLA-DPA1 |
| CXCL1 | GRPEL2 | SNF8 |
| FBXO6 | HMGXB3 | CHST10 |
| ADGRL4 | MRPL22 | SLC39A13 |
| SLC30A10 | SGCD | LRRTM3 |
| ITGA9 | FNDC9 | NRG3 |
| FLNB | FABP6 | MAPKBP1 |
| DNAJC30 | WWC1 | TMED3 |
| THAP5 | SFXN1 | FRMD4B |
| METTL2B | GPRIN1 | NUTM2B-AS1 |
| SLC4A2 | TSPAN17 | KLHL29 |
| MRPL50 | TMED9 | RRP9 |
| AKR1C2 | B4GALT7 | LSM11 |
| TLCD5 | CDYL | COL12A1 |
| LRRC57 | SIRT5 | COL26A1 |
| QPRT | MCUR1 | LRP6 |
| NAT9 | GMNN | TP53 |
| DMKN | H2AC6 | PIGL |
| TGM2 | ZKSCAN8 | L3MBTL4 |
| ZFY | ZSCAN26 | IRF2BP1 |
| PDCD2 | DHX16 | JADE2 |
| GLDC | VARS2 | HMGCL |
| DDX10 | EHMT2 | NGLY1 |
| PKNOX2 | PSMB8 | ASCC3 |
| PHF12 | MAPK14 | DYNC2I2 |
| LPCAT1 | UBR2 | ASXL3 |
| IMPDH1 | ZNF318 | KBTBD8 |
| SEC24C | SLC25A27 | ZMYM6 |
| RBM38 | CD2AP | LINC01750 |
| SNAPC5 | LINC00680 | LIX1L |
| ZNF24 | B3GAT2 | ANKRD36C |
| CLIC4 | CASP8AP2 | SETMAR |
| SPEN | FUT9 | CNOT10 |
| FAM228B | CCNC | CEP70 |

| | | |
|---|---|---|
| CHD6 | GRIK2 | LMLN |
| PCAT19 | RNF217 | INPP4B |
| MED13L | PERP | KCNIP1 |
| TMBIM1 | LTV1 | FRS3 |
| TECPR1 | MYCT1 | KCNH2 |
| VPS13B-DT | GTF2H5 | ZC3H3 |
| NRG3 | MAP3K4-AS1 | MTAP |
| HPS1 | FOXK1 | NPR2 |
| BEST1 | ZDHHC4 | ZNG1C |
| ACER3 | ZNF12 | STN1 |
| LEMD3 | THSD7A | LRRK2 |
| GLIPR1 | ANKMY2 | DNAH10OS |
| RCBTB1 | SKAP2 | NUFIP1 |
| JDP2 | BBS9 | KATNAL2 |
| SULT1A1 | DPY19L1 | LMAN1 |
| RBL2 | MPLKIP | RFX2 |
| FTSJ3 | C7orf25 | EHD2 |
| MON2 | TBRG4 | ESF1 |
| USP10 | UPP1 | SLC5A3 |
| GTPBP8 | STAG3L4 | ADA2 |
| TRAF3IP2 | POMZP3 | NHS |
| LINC01551 | STEAP2 | PIKFYVE |
| PRRG1 | NYAP1 | IL34 |
| ERMN | GIGYF1 | KCTD8 |
| TIMM44 | IFT22 | PAN2 |
| ANKRD52 | SLC35B4 | NEIL1 |
| C1orf174 | DGKI | TRAF4 |
| CYP4X1 | ZNF746 | JUP |
| THADA | REPIN1 | RNF138 |
| TGFBR2 | KCNH2 | YIPF2 |
| ENTPD3 | NOM1 | FRMD5 |
| POGLUT1 | XPO7 | GTF2H3 |
| IQCB1 | NPM2 | DHRS4L2 |
| NEURL1B | CCAR2 | ADGRV1 |
| LTC4S | DOCK5 | HCST |
| NFKBIL1 | RGS20 | SRRD |
| MCHR2 | XKR4 | PIH1D1 |
| NHSL1-AS1 | PAG1 | SPON1 |
| EPM2A-DT | TMEM64 | BOP1 |
| HIBADH | TSPYL5 | ZNF326 |
| MET | MAL2 | GPALPP1 |
| C8orf90 | SQLE | SLC25A39 |
| PIGO | TRAPPC9 | MOB3C |

| | | |
|---|---|---|
| CARNMT1 | GLI4 | CTSS |
| PTPDC1 | SCRIB | VPS26C |
| ASCC1 | PARP10 | VPS18 |
| TMEM254 | ADCK5 | PPP1R8 |
| ZDHHC16 | LRRC14 | HRH2 |
| CYB561A3 | WASHC1 | SLC25A17 |
| BORCS5 | ZNG1A | DYNC2H1 |
| ST8SIA1 | ZDHHC21 | LCP1 |
| RITA1 | MOB3B | GSTM5 |
| RBM19 | EBLN3P | RNPC3 |
| XPO4 | ANXA1 | CACNA2D2 |
| PARP4 | AUH | SNX24 |
| DPF3 | TTLL11 | PRRC1 |
| PAPLN | PBX3 | ING3 |
| PTGR2 | ZDHHC12 | UAP1L1 |
| UNC79 | TSC1 | EML3 |
| EIF3J-DT | BRD3OS | LTBR |
| LRRC28 | MRPS2 | P2RX7 |
| PMM2 | GTPBP4 | PDP2 |
| ZNF689 | SFMBT2 | SPATA33 |
| METTL16 | GAD2 | PANX1 |
| GID4 | EPC1 | NUDT10 |
| PLEKHH3 | DDX50 | ILDR2 |
| CCDC137 | BMPR1A | ZNF385D |
| ZNF426 | EXOC6 | MSH3 |
| ELL | LGI1 | TRIAP1 |
| LIN37 | PDCD11 | PRRG3 |
| YDJC | ADRA2A | DNLZ |
| YPEL1 | ADRB1 | THAP7 |
| SFI1 | CCDC186 | C1orf159 |
| NUDT10 | FHIP2A | TMEM51 |
| TAF1 | WDR11 | PDIK1L |
| ARMCX5 | IKZF5 | THBS3-AS1 |
| AFF2 | CYB5R2 | RXRG |
| POGZ | GTF2H1 | INTS7 |
| KLHDC8B | BDNF-AS | RNASEH1-DT |
| CEP63 | PRR5L | GALNT14 |
| DTX4 | CCDC86 | PCGF1 |
| APC2 | MTA2 | CCDC51 |
| GJA4 | ROM1 | BOC |
| DGAT2 | DNAJC4 | ELMOD2 |
| RFX4 | DPF2 | MARCHF11 |
| POLR3K | BRMS1 | PGBD1 |

| | | |
|---|---|---|
| ZNF48 | TMEM134 | DNAAF5 |
| TANGO6 | CCND1 | ZNF736 |
| ZNF397 | RNF121 | LINC01004 |
| SOX18 | MRPL48 | DOCK5 |
| MIIP | PAAF1 | EIF4EBP1 |
| SPOCD1 | AAMDC | LRATD2 |
| NT5C1A | INTS4 | CNTNAP3 |
| CDKN2C | ME3 | NOL8 |
| CPT2 | TMEM135 | ANKRD16 |
| MOV10 | GRM5 | PLPP4 |
| NBPF9 | ROBO3 | EMSY-DT |
| SELE | NFRKB | VRK1 |
| METTL13 | ITFG2 | LINC02352 |
| HS1BP3 | RHNO1 | EFL1 |
| VAMP8 | C1R | BRICD5 |
| ARL6IP6 | TMEM52B | CNTROB |
| ANKRD44 | PTPRO | ZNF577 |
| METTL21A | IRAG2 | MYH7B |
| SH3BP4 | MED21 | SNAI1 |
| CPNE9 | GXYLT1 | RNF215 |
| RARB | POU6F1 | RFPL2 |
| ZBTB47-AS1 | GIHCG | CPT1B |
| PARP9 | SLC16A7 | TMEM187 |
| KLHL6 | TBK1 | PTGFRN |
| NOP14-AS1 | DYRK2 | PPP1R26 |
| ABCG2 | KRR1 | VPS13C |
| ABHD18 | SLC6A15 | WASH2P |
| PTCD2 | GPN3 | CACUL1 |
| PCDHGC3 | FAM216A | GGA3 |
| MFAP3 | RNFT2 | ZHX2 |
| DRD1 | SUDS3 | MED9 |
| ZBTB9 | DNAH10OS | PLCXD2 |
| RTN4IP1 | GLT1D1 | HILPDA |
| PDSS2 | RIMBP2 | ATG14 |
| GTPBP10 | EP400 | EVI2A |
| TFR2 | MTMR6 | FAM210A |
| LINC01004 | RFXAP | SPATC1L |
| ZC3HC1 | LACC1 | GRASLND |
| ZNF212 | UGGT2 | LRBA |
| GIMAP5 | UBAC2 | PRIMPOL |
| IKBKB | PCID2 | VCAN |
| NSMCE2 | PARP2 | BNIP1 |
| LRATD2 | OSGEP | VARS2 |

| | | |
|---|---|---|
| ZNF623 | REC8 | RP9 |
| CDC37L1-DT | NPAS3 | PUS7 |
| CNTNAP3B | PYGL | FLNC |
| GRIN3A | SGPP1 | DCAF10 |
| ADAMTS13 | EXD2 | TBC1D12 |
| AKR1C3 | ENTPD5 | LGR4 |
| VSTM4 | SPTLC2 | TEX30 |
| TSPAN15 | FAM181A | VASH1 |
| GATD1-DT | PPP4R4 | WDR90 |
| UEVLD | CINP | RFFL |
| PRDM11 | ADSS1 | OLIG2 |
| SLC15A3 | MAGEL2 | TMEM191C |
| NUDT8 | RYR3 | TTC28 |
| NEU3 | DISP2 | FAM118A |
| CEP164 | DLL4 | TYW1 |
| USP2-AS1 | SPG11 | SLC25A29 |
| PUS3 | GNB5 | ZNF175 |
| PTPRB | FAM81A | GPR89B |
| CABP1-DT | C2CD4B | METTL25B |
| LATS2 | COMMD4 | RBM45 |
| PGF | SIN3A | NEUROD1 |
| ALKBH1 | SNUPN | ITGA9-AS1 |
| TEDC1 | PEAK1 | LINC02035 |
| PDCD7 | TLNRD1 | CC2D2A |
| LETR1 | EFL1 | ADAMTS2 |
| PCSK6 | BTBD1 | RGS17 |
| MIR193BHG | FES | MAS1 |
| SLX1A | HBA1 | PARP10 |
| ORAI3 | CRAMP1 | TEK |
| CMTM3 | NUBP2 | PCSK5 |
| NFATC3 | GFER | ENKUR |
| ADAT1 | NHERF2 | PAOX |
| MYO1C | ABCA3 | TCP11L1 |
| PIGW | IL32 | GRIK4 |
| HEXIM2 | ZSCAN32 | P3H3 |
| METTL2A | ADCY9 | C2CD5-AS1 |
| MIF4GD | UBN1 | TBC1D4 |
| TINCR | SHISA9 | DPF3 |
| RAB11B-AS1 | BFAR | TUNAR |
| CCNE1 | EARS2 | MEAK7 |
| PDCD2L | HS3ST4 | MAPT-AS1 |
| DHX34 | NUP93 | TEPSIN |
| FUT1 | CMTM4 | IZUMO4 |

| | | |
|---|---|---|
| ZNF473 | PARD6A | PDE4C |
| ZNF320 | GFOD2 | ZNF766 |
| ZNF274 | ZFHX3 | SOX18 |
| PXMP4 | MBTPS1 | TBL1X |
| PTPRT-DT | PIEZO1 | PTPRJ |
| FITM2 | DHX33 | THAP12 |
| CSTF1 | PITPNM3 | TAGLN |
| EVA1C | XAF1 | MFAP1 |
| RNF215 | PHF23 | INO80B |
| SSTR3 | GPS2 | ADCK2 |
| LINC01315 | NEURL4 | GHDC |
| CA5B | ACAP1 | GRWD1 |
| CLDN11 | TP53 | LYRM4 |
| SPATA33 | TRIM16 | SLC35A1 |
| RAB39B | LLGL1 | RASGRP1 |
| EGLN1 | B9D1 | MRPS16 |
| ANGPTL2 | RSKR | PCIF1 |
| UQCC4 | SEZ6 | PLA2G2C |
| SHARPIN | CORO6 | TMEM234 |
| SPIRE2 | C17orf75 | CPT2 |
| PRPF3 | STAC2 | SULT1C4 |
| IRF2BPL | ORMDL3 | TANC1 |
| RGL2 | RETREG3 | LRP2 |
| KYAT3 | C1QL1 | PMS1 |
| FCGR1BP | SMG8 | ACVR2B |
| RIT1 | TANC2 | SLC38A3 |
| PRRX1 | SOX9-AS1 | PLAAT1 |
| ARV1 | NT5C | HARS2 |
| PIGF | MGAT5B | LINC01202 |
| RASSF1 | BAIAP2-DT | MICA |
| NPRL2 | ARL16 | AZGP1 |
| ARHGEF26-AS1 | TYMS | STK3 |
| BACH2 | GNAL | SLA |
| SMURF1 | KCTD1 | RPP38 |
| FRMD3 | RNF138 | ZNF143 |
| PHYHD1 | PMAIP1 | ATF7 |
| FHIP1B | VPS4B | POLR3B |
| ATG2A | TMX3 | MRPS31P5 |
| ABCC9 | DOK6 | EFS |
| CCNT1 | R3HDM4 | TMEM260 |
| C16orf89 | CSNK1G2 | SLX1A |
| PRMT7 | MOB3A | ZFHX3 |
| FAM43A | SIRT6 | MED11 |

| | | |
|---|---|---|
| FAM20B | PSPN | CHRNB1 |
| SPOP | ZNF846 | TRPV2 |
| MPV17 | RAVER1 | NXPH3 |
| RRAGD | ZNF844 | WIPI1 |
| OSTM1 | MIR23AHG | ARMC7 |
| TMEM141 | RFX1 | CD22 |
| VSIR | NR2F6 | C19orf47 |
| CD53 | ANO8 | ZNF71 |
| UBXN4 | PGLS | PDE9A |
| PPIF | GDF15 | BAIAP2L2 |
| APCDD1 | TSHZ3 | SHROOM4 |
| SRP19 | GRAMD1A | CRTC2 |
| NALF1 | ZNF461 | GPATCH2 |
| ARCN1 | ZNF569 | SLC25A40 |
| R3HDM2 | DLL3 | SPATS2 |
| LYRM7 | MYPOP | NCOA3 |
| CEP41 | ZNF766 | ABL1 |
| RAC3 | ZNF787 | BRD3 |
| MTM1 | ZNF264 | PRKAA1 |
| SAP30 | ZNF544 | CEBPZ |
| CYP7B1 | ZNF324 | ITGB5 |
| CDK11A | SLC27A5 | PAK1IP1 |
| LINC02609 | ZBTB45 | ABCB8 |
| CNN3-DT | NRSN2 | ZFC3H1 |
| KCNK3 | STK35 | RABGGTA |
| ZNF514 | VPS16 | SIL1 |
| MITD1 | TRMT6 | SOCS2 |
| CCDC93 | KIF3B | GYS1 |
| DTYMK | TTI1 | GARIN5A |
| PLCXD2 | KIAA1755 | HES1 |
| ARHGEF26 | DHX35 | LATS1 |
| GNPDA2 | PLTP | SPA17 |
| NEK1 | PCIF1 | NR2F2 |
| WDR55 | SS18L1 | SCMH1 |
| LINC00847 | MRPL39 | MTF2 |
| RNF144B | HUNK | NUDT4B |
| LY6G5B | MORC3 | C1orf54 |
| ZNRF2 | KCNJ6 | GLMP |
| RP9 | ABCG1 | LBR |
| ZNF736 | PDE9A | PRKD3 |
| DLX6-AS1 | PEX26 | DCTN1-AS1 |
| ZMAT4 | KLHL22 | BTD |
| FCHSD2 | SDF2L1 | TGFBR2 |

| | | |
|---|---|---|
| NPAT | BCR | UVSSA |
| LIMA1 | GAS2L1 | TFEB |
| PRANCR | MTMR3 | BICRAL |
| CMTM5 | SEC14L2 | UFL1 |
| PRPF39 | PES1 | LFNG |
| COQ6 | RNF185 | STYXL1 |
| GOLGA5 | PRR14L | LHX2-AS1 |
| CPPED1 | MPPED1 | REXO4 |
| ZDHHC7 | GTPBP6 | DHTKD1 |
| BLMH | PUDP | DNAJC24 |
| GAA | FRMPD4 | TLCD5 |
| FAM210A | PRPS2 | CNPY2-AS1 |
| PTGR3 | MOSPD2 | GIT2 |
| AMH | PCYT1B | XPO4 |
| TICAM1 | RPGR | RNF31 |
| CERS4 | NUDT11 | DNAAF2 |
| GYS1 | IQSEC2 | COQ6 |
| SHISA7 | FAM120C | C15orf40 |
| HAR1A | RRAGB | COQ7 |
| SPECC1L | CHIC1 | ZNF747 |
| SHISA8 | TMEM35A | DHRS7B |
| HDAC10 | ARMCX2 | HAP1 |
| MAGT1 | FAM199X | CEP192 |
| MRS2 | ZBTB33 | ZNF181 |
| SOX9 | PHF6 | ZNF160 |
| GLRX2 | ARHGEF6 | ZNF583 |
| TMUB1 | EOLA1 | TASP1 |
| SEC23IP | FAM3A | CDH4 |
| UBE4A | DKC1 | TRIOBP |
| BRI3BP | TXLNGY | LRCH2 |
| IFT43 | ANLN | WDR44 |
| CLPX | RNH1 | C1GALT1C1 |
| NR1H2 | POLR2J | NEUROD6 |
| SMIM17 | GAS5 | MRPS9 |
| SLC30A7 | TBPL1 | TMEM121B |
| GAREM2 | GUCY1A2 | SLC27A3 |
| ANO10 | FAM210B | ALG5 |
| KBTBD8 | SLC35A3 | TUBGCP4 |
| LSM2 | RAPH1 | CEP250 |
| RPF2 | ZFAND2B | ELOA |
| AKAP7 | C4orf50 | ELOVL1 |
| DCHS1 | TBC1D14 | ITGA2 |
| DYRK4 | KLF3 | POC1B |

| | | |
|---|---|---|
| ESYT1 | SMARCAD1 | ABTB3 |
| TMEM19 | DNAJC2 | DMAC2L |
| AMDHD2 | CSGALNACT1 | GCHFR |
| RC3H1 | PHF2 | GDAP2 |
| EMG1 | BTAF1 | CNNM4 |
| SPPL2B | FIBIN | CASP7 |
| ZNF664 | PACSIN3 | FAM120C |
| POLDIP3 | SNX32 | LRCH3 |
| ZRANB1 | ABCD2 | UBA6 |
| SLC39A1 | GDF11 | INTU |
| RIMS4 | SOX21 | KCTD9 |
| GMPPA | C14orf132 | SMC5 |
| RFX3 | SLC25A29 | ZNF280D |
| TMTC1 | TECPR2 | CCDC43 |
| SLC41A2 | PTPN9 | TIMM21 |
| TTC4 | PDE8A | PCYT1B |
| CCL20 | PAQR4 | C12orf75 |
| LIAS | BBS2 | ZNF830 |
| HYCC1 | SLC7A6 | SLC45A1 |
| ASB13 | ZFP90 | TXNIP |
| RHBDL3 | COG4 | SLITRK3 |
| TMEM51 | ZNF358 | METTL14 |
| ARHGEF10L | PRKACA | CCNB1 |
| FUCA1 | OXR1 | HLA-DMA |
| TMEM234 | CCT5 | TBP |
| AGO4 | SNCG | RFC2 |
| AGBL4 | SLBP | ZNF23 |
| KTI12 | NCAN | IFI35 |
| RWDD3 | THRAP3 | CCDC137 |
| NTNG1 | MAP3K13 | ZNF653 |
| LINC00623 | FRYL | MPV17L2 |
| GOLPH3L | FAT1 | HDAC10 |
| ZNF687 | SEC24A | XK |
| F11R | ZNF25 | NLGN4Y |
| USP21 | DOCK1 | PPIH |
| TADA1 | TMEM263 | STEAP2 |
| MTARC2 | ANKLE2 | RPRM |
| ATAD2B | MPDU1 | DNAJC9 |
| RASGRP3 | DCAF7 | YIPF5 |
| THUMPD2 | MBD2 | CCNG1 |
| REL | MFSD12 | PLRG1 |
| DCTN1-AS1 | ATF5 | MGAT2 |
| GPR17 | ZNFX1 | NKRF |

| | | |
|---|---|---|
| FRZB | UBE2G2 | MED28 |
| CYP20A1 | RANBP1 | HAVCR2 |
| CTDSP1 | STOML2 | USO1 |
| SPHKAP | POLR2B | ITPR2 |
| SETMAR | CENPX | LEMD3 |
| CCDC174 | PFDN1 | TESK2 |
| BOC | HSD17B12 | AGBL4 |
| IQCG | LTBP3 | DNAJC27-AS1 |
| MYL5 | BCL6 | UCN |
| ADRA2C | OGDH | TTC7A |
| COX18 | ADAMTS4 | CYP20A1 |
| NAAA | YIPF4 | PPM1M |
| CXXC4 | TLK1 | CTBP1-DT |
| TET2 | PARP14 | PTGER4 |
| CYP2U1-AS1 | LINC03013 | CMYA5 |
| LINC00499 | CUX1 | GALNT10 |
| PRIMPOL | SETX | AARS2 |
| BASP1-AS1 | CTNNA3 | MANEA |
| SH3TC2 | TRIP11 | NHSL1-AS1 |
| SMIM3 | DICER1 | ICA1-AS1 |
| TSBP1-AS1 | HERC2 | CAV2 |
| PSMB9 | VPS39 | CHD7 |
| TREM1 | CLUAP1 | LINC02668 |
| XPO5 | LONP2 | ITIH5 |
| KIAA1586 | CBFB | SUFU |
| PLEKHA8 | MYBBP1A | DCPS |
| NEUROD6 | ARL4D | DBX2 |
| EEPD1 | DDX39A | FKBP11 |
| ZNF138 | C5AR1 | STYX |
| GTF2IRD2B | YKT6 | CLBA1 |
| ZNF394 | TMEFF1 | CRIP1 |
| AP4M1 | FAM32A | ZNF205 |
| MOSPD3 | SIK1 | ZNF629 |
| LINC01007 | RLIM | HIGD1B |
| ORC5 | RPS19BP1 | ZNF44 |
| PUS7 | SHISA4 | ZNF430 |
| FAM131B-AS2 | RYBP | ZNF585B |
| EIF4EBP1 | GNL1 | ZNF570 |
| CPQ | CYP46A1 | SHKBP1 |
| FZD6 | EPHX1 | DHX34 |
| EXOSC3 | VSTM2A | IL11 |
| SUSD1 | ANXA11 | ZNF471 |
| TOR2A | PGP | LINC01260 |

| | | |
|---|---|---|
| MCU | LRRC8B | GGT5 |
| FGFBP3 | PRKAR2A | TFIP11-DT |
| TCTN3 | OTUD4 | LIMK2 |
| C10orf88 | GABRB3 | DNAJC2 |
| SLC17A6 | MGA | HYCC1 |
| ALKBH3 | BLOC1S6 | TNFSF13 |
| DDB2 | VOPP1 | MRPL42 |
| CLP1 | SPTBN4 | EIF1AD |
| FZD4 | SLC20A1 | CNTNAP4 |
| TSPAN9 | CSPG5 | PAQR4 |
| LTBR | SYNE1 | UBR4 |
| DERA | NPEPPS | RICTOR |
| PCED1B-AS1 | AGAP1 | ZCCHC10 |
| AAAS | YME1L1 | NSMCE1 |
| SRGAP1 | PBX1 | ERN1 |
| HCFC2 | TAF9 | DLL3 |
| ACACB | GLO1 | PES1 |
| HRK | ZFAND2A | INTS3 |
| B3GNT4 | SYT7 | IFNGR1 |
| FGF9 | MRPL21 | WDR77 |
| RASL11A | NDUFV3 | OLFML3 |
| LNX2 | DGCR2 | WDR41 |
| DACH1 | MPHOSPH8 | RUNX1T1 |
| PLD4 | CCN2 | COX15 |
| ZWILCH | LSAMP | VTI1A |
| NUBP1 | IFITM10 | RBM19 |
| ABCC12 | SDC3 | OTUD7A |
| C16orf46 | ACTR1A | TMEM231 |
| CTC1 | UFM1 | TRMT6 |
| ADORA2B | HEXIM1 | OSER1-DT |
| HIGD1B | PRPF40A | TBC1D10A |
| KANSL1-AS1 | LRRFIP1 | RPL39L |
| MAP2K6 | PDS5A | LRP8 |
| SLC25A19 | ITGB8 | PLCB4 |
| MYO15B | USP47 | DUBR |
| CEP192 | ITGB3 | KLHL8 |
| SALL3 | SSNA1 | TIFA |
| ZNF507 | PPM1K | SLC41A2 |
| ZNF383 | FBXO9 | CLN6 |
| ZNF780B | BAG1 | CCDC22 |
| HAS1 | TMX4 | PUSL1 |
| ZNF528 | MAPK8IP3 | MGST2 |
| ZNF865 | BAD | LYSMD3 |

| | | |
|---|---|---|
| KIF16B | SRA1 | DUSP22 |
| TSHZ2 | DENND11 | NSUN5 |
| TAF4 | RPS27L | VPS13B |
| TMEM121B | SNRPD3 | WDR25 |
| CCDC117 | SLC30A1 | ZNF875 |
| CHADL | LRIG1 | FBXO6 |
| TTC38 | NAPA | CHD1L |
| OFD1 | VMA21 | ZNF687 |
| HMGN5 | SRSF8 | ATP8B2 |
| TMEM164 | SCN3B | ESRRG |
| SNED1 | PDHA1 | CAMKMT |
| SRD5A3 | RRBP1 | ANKRD36B |
| MICAL1 | APH1A | LINC01102 |
| AGFG2 | ABL2 | MAP3K2-DT |
| TMCC3 | CWC27 | ZNF197 |
| P2RX7 | MYO9B | NAA80 |
| TNFRSF25 | COPRS | DTX3L |
| PPP1R16A | ST6GAL1 | IQCG |
| SDF2 | KALRN | ACOX3 |
| LAMA4 | NNT | PCDHB14 |
| POLRMT | DBNL | ZNF354A |
| HTRA2 | RTF1 | HCG11 |
| SGPP2 | MEF2A | PSMB8 |
| PPT2 | HUWE1 | LCA5 |
| TATDN1 | WDR83OS | DCBLD1 |
| AIF1L | ZSCAN16-AS1 | DPY19L4 |
| PRPF18 | MTUS1 | ADCY8 |
| OLMALINC | CREG2 | ABITRAM |
| PSTK | FBXW4 | DNAJC1 |
| CPSF2 | FERMT3 | PARG |
| STS | CDC27 | TSPAN15 |
| MMP28 | LILRB4 | NANOS1 |
| NIPA1 | EIF3G | CYB561A3 |
| MRPL9 | LCP1 | NADSYN1 |
| PCYT2 | CYB5R3 | DNAJC14 |
| ZFHX4 | KLHDC2 | SLC16A7 |
| STC1 | SFXN5 | COQ5 |
| LIN52 | TPGS1 | TMEM120B |
| ANKFY1 | SDF4 | ABHD4 |
| DHRS3 | CCNL2 | GCH1 |
| SGMS2 | USP48 | SQOR |
| ALG13 | ATPAF1 | DVL2 |
| LMAN2 | SNRPE | MYO19 |

| | | |
|---|---|---|
| SCN4B | IARS2 | OSBPL7 |
| DCP1A | FOXN2 | SLC16A3 |
| AMN1 | UNC50 | ZNF561-AS1 |
| MTHFR | MBD5 | ZNF776 |
| ANTXR1 | TUBA4A | SLC13A3 |
| CAPN10 | EDEM1 | RBM11 |
| PHLDA2 | ZMAT3 | RAB36 |
| TRHDE | AFF1 | ALG12 |
| NOL3 | HHIP | EOLA2 |
| VRK3 | NADK2 | NOA1 |
| CMC4 | CPLANE1 | ZNF362 |
| | ERGIC1 | CITED4 |
| | GRK6 | SOX13 |
| | ACOT13 | LYPLAL1 |
| | PHF3 | KLF7 |
| | MICALL2 | NPPC |
| | GSR | HACL1 |
| | GOLGA2 | LARS2 |
| | RIMKLB | AMT |
| | TFCP2 | ARL13B |
| | RAB20 | CRYBG3 |
| | DHRS7 | OSBPL11 |
| | INAFM2 | PPAT |
| | TMEM87A | ANAPC10 |
| | IQGAP1 | ERAP1 |
| | ECI1 | PCYOX1L |
| | SLC43A2 | FARS2 |
| | TAX1BP3 | LURAP1L |
| | FLII | SLC31A2 |
| | TMEM98 | PNPLA7 |
| | TSPOAP1 | C11orf98 |
| | PLIN3 | SPINDOC |
| | ICOSLG | TSPAN9 |
| | LSS | PTPRB |
| | AMER2 | PHF11 |
| | C1orf159 | LYSET |
| | PHF13 | ZNF213 |
| | DISP3 | CBFA2T3 |
| | LINC01141 | TMEM235 |
| | CSMD2 | TLE2 |
| | EVA1B | ISYNA1 |
| | LSM10 | TSHZ3 |
| | SCMH1 | ZNF544 |

| | |
|---|---|
| NSUN4 | ZNF8-DT |
| MOB3C | GART |
| PSRC1 | SLC7A4 |
| MYBPHL | PLP2 |
| PMF1 | MPPED2 |
| MRPS14 | DLC1 |
| BRINP2 | STON2 |
| MIR29B2CHG | CLN8-AS1 |
| TBCE | DTNBP1 |
| RAB10 | C4orf50 |
| PKDCC | CD14 |
| SPR | MASTL |
| POLR1B | SMCR8 |
| WASH2P | ZMAT5 |
| NIFK-AS1 | TNFRSF1B |
| TTC21B | CYB5R1 |
| SCN1A | PIAS3 |
| PMS1 | ACY1 |
| FASTKD2 | MED7 |
| ZNF621 | SLC35B3 |
| EXOSC7 | ZBTB2 |
| CCRL2 | KDM4C |
| TCTA | TSTD2 |
| PBRM1 | LATS2 |
| TASOR | DECR2 |
| DZIP3 | PPIL2 |
| SNX4 | CTH |
| MBD4 | ZC3H8 |
| NLGN1 | CASP2 |
| MAGEF1 | NDRG1 |
| DCAF16 | PLD2 |
| NUDT9 | ADA |
| SLC39A8 | CSK |
| SMIM43 | MRPS17 |
| RNF175 | STPG1 |
| SAP30-DT | SERTAD4-AS1 |
| ING2 | TMEM17 |
| CYP4V2 | IFT88 |
| IPO11 | DHDH |
| SNX24 | TAF12 |
| DNAJC18 | ATXN7L2 |
| DIAPH1 | HSD11B1 |
| DELE1 | MTARC1 |

| | |
|---|---|
| ARHGEF37 | CAD |
| GALNT10 | WNT7A |
| PPP1R11 | XPC |
| VPS52 | CCDC80 |
| KCTD20 | QTRT2 |
| TFEB | LYAR |
| MMUT | SETD9 |
| PRDM1 | PCDHB4 |
| MUC12-AS1 | ADAM19 |
| RINT1 | ZUP1 |
| PDIA4 | SOSTDC1 |
| LINC01003 | FIGNL1 |
| HR | SEMA3E |
| PPP3CC | BET1 |
| DECR1 | TFR2 |
| VIRMA | ZNF277 |
| VPS13B | LRRCC1 |
| MTAP | LY6E-DT |
| CAAP1 | PTPRD-AS1 |
| MIR600HG | TBC1D2 |
| RALGPS1 | PTGS1 |
| RNF208 | KBTBD7 |
| UPF2 | SETDB2 |
| PAPSS2 | ARMH4 |
| WBP1L | PAPLN |
| CNNM2 | DICER1-AS1 |
| ZDHHC6 | ZNF75A |
| CARS1 | ZSCAN32 |
| ZNF408 | MIR762HG |
| OSBP | DDX28 |
| ZNRD2 | ACSF3 |
| C11orf54 | WDR81 |
| FAM76B | SLC13A5 |
| FAM118B | PIPOX |
| NOP2 | FKBP10 |
| PTPN6 | RECQL5 |
| EPS8 | MALT1 |
| TESPA1 | ZNF554 |
| METTL1 | RBM38 |
| PPFIA2 | PPP1R3D |
| SCYL2 | PUDP |
| WASHC3 | EFNB1 |
| PXN | ENOX2 |

| | |
|---|---|
| GTF2H3 | RNF216 |
| ATP11A | ACAD9 |
| PPP1R3E | MTMR1 |
| DMAC2L | PLPPR3 |
| TMEM260 | THNSL2 |
| FCF1 | C11orf24 |
| VASH1 | PCDHB10 |
| TRMT61A | TATDN1 |
| TYRO3 | MLLT3 |
| TUBGCP4 | TRPM3 |
| KLHL25 | SMUG1 |
| HAGHL | RASL11A |
| EME2 | AIFM1 |
| DCTPP1 | EPOP |
| PRR14 | RAMACL |
| MIR762HG | KIAA1143 |
| ZNF267 | CHCHD3 |
| GAS8 | CCDC92B |
| SMG6 | MPND |
| ELAC2 | NKAIN1 |
| PEMT | |
| ATPAF2 | |
| UTP6 | |
| PEX12 | |
| AATF | |
| COASY | |
| UBE2Z | |
| ITGB4 | |
| DNAH17 | |
| ENGASE | |
| RPTOR | |
| TEPSIN | |
| FN3K | |
| CABLES1 | |
| B4GALT6 | |
| SCAMP4 | |
| CAMSAP3 | |
| ZNF266 | |
| YJU2B | |
| TMEM38A | |
| NUDT19 | |
| LIPE | |
| ERCC2 | |

| |
|---|
| SPACA6 |
| CACNG7 |
| TRAPPC2B |
| TASP1 |
| SCARF2 |
| SGSM1 |
| GRK3 |
| TTC28-AS1 |
| SF3A1 |
| TCN2 |
| FOXRED2 |
| MPST |
| SLC25A17 |
| TEF |
| KLHL15 |
| GJB1 |
| TIMM8A |
| TCEAL1 |
| PABIR2 |
| PHB2 |
| NADK |
| NOL9 |
| UBR4 |
| ZNF593 |
| FAM76A |
| TXLNA |
| TMEM54 |
| SMIM12 |
| ADPRS |
| GPBP1L1 |
| PRPF38A |
| ZYG11B |
| PLPPR5 |
| AGL |
| C1orf54 |
| VPS72 |
| EIF2D |
| GGPS1 |
| MTR |
| RNF144A |
| PPM1G |
| RETSAT |
| PANTR1 |

| |
|---|
| ZNG1B |
| SAP130 |
| STK39 |
| FAM117B |
| CNOT9 |
| VHL |
| TMEM43 |
| DPH3 |
| LRRC3B |
| PTPN23 |
| TBC1D23 |
| STXBP5L |
| LINC01322 |
| HES1 |
| BDH1 |
| RUBCN |
| DCUN1D4 |
| ATPSCKMT |
| ELOVL7 |
| CERT1 |
| ATG12 |
| TCERG1 |
| JAKMIP2 |
| MGAT1 |
| SNRNP48 |
| TXNDC5 |
| C6orf136 |
| PPARD |
| FAM135A |
| MAP3K7 |
| POU3F2 |
| HECA |
| AP5Z1 |
| USP42 |
| ARL4A |
| POLR1F |
| KLHL7 |
| FKBP9 |
| CDK13 |
| LINC03011 |
| SEMA3C |
| RHOBTB2 |
| MRPL15 |

| |
|---|
| C8orf76 |
| MYC |
| AGO2 |
| RANBP6 |
| ABHD17B |
| IPPK |
| FAM120AOS |
| TMEM38B |
| DAB2IP |
| RBM18 |
| NELFB |
| WAC-AS1 |
| IPMK |
| TSPAN14 |
| EMX2 |
| ACADSB |
| MRPL23 |
| SBF2 |
| SOX6 |
| TSG101 |
| PRMT3 |
| KIAA1549L |
| LDLRAD3 |
| FNBP4 |
| UBE2L6 |
| FADS3 |
| TMEM179B |
| PYGM |
| CLPB |
| RSF1 |
| KCTD21 |
| UBASH3B |
| ETS1 |
| TNFRSF1A |
| COPS7A |
| ETV6 |
| SPX |
| KRAS |
| RESF1 |
| LETMD1 |
| ACVR1B |
| CTDSP2 |
| TMBIM4 |

| |
|---|
| VEZT<br>N4BP2L1<br>LRCH1<br>CAB39L<br>LMO7<br>TPP2<br>CUL4A<br>RAB2B<br>SALL2<br>RBM23<br>RABGGTA<br>EIF2B2<br>MFAP1<br>PLEKHO2<br>RAB11FIP3<br>PRRT2<br>FBRS<br>OGFOD1<br>CSNK2A2<br>CDH8<br>CTCF<br>RILP<br>DERL2<br>MBTD1<br>TNRC6C<br>USP36<br>ASPSCR1<br>SMCHD1<br>DLGAP1<br>GAREM1<br>IER3IP1<br>MRO<br>ZNF532<br>PLPPR3<br>ILVBL<br>WIZ<br>RAB8A<br>ZNF506<br>ZNF302<br>ZNF260<br>NFKBIB<br>AXL<br>PIH1D1 |

| |
|---|
| FPR1 |
| HSPBP1 |
| SNHG11 |
| ELMO2 |
| MIR155HG |
| PSMG1 |
| ZMAT5 |
| TRIOBP |
| RBBP7 |
| PRDX4 |
| SNX12 |
| PIN4 |
| GLA |
| PFKM |
| ATXN7L3B |
| CLASP1 |
| CASTOR2 |
| PGAP4 |
| PTEN |
| COX14 |
| CAMKK2 |
| TSR3 |
| PAM16 |
| CBARP |
| UNC13A |
| IDH3G |
| SRGAP3 |
| THOC7 |
| PODXL2 |
| CPLX1 |
| SORBS2 |
| BAG6 |
| YTHDF3 |
| KIFC2 |
| SNHG1 |
| SIK2 |
| ARL1 |
| PPP1CC |
| SPPL3 |
| EMC6 |
| ZBTB4 |
| HSD11B1L |
| OLFM2 |

| | |
|---|---|
| | SSBP4 |
| | SNTA1 |
| | CBX7 |
| | TCEAL6 |
| | BOD1 |
| | CCL4L2 |
| | TAF1D |
| | CORO1C |
| | DCUN1D1 |
| | RAPGEF1 |
| | HOMER1 |
| | UTP11 |
| | WDR82 |
| | ANAPC13 |
| | PAICS |
| | MTMR12 |
| | SAP30L |
| | ADCYAP1R1 |
| | STMP1 |
| | AGPAT5 |
| | CEMIP2 |
| | DNLZ |
| | MRPS16 |
| | LIPA |
| | PTPRE |
| | SPON1 |
| | TAOK3 |
| | SRP54 |
| | RHOT1 |
| | NOVA2 |
| | ISOC2 |
| | PTPN1 |
| | ELK1 |
| | SRSF11 |
| | ACAT1 |
| | DDX3Y |
| | NASP |
| | GALNT8 |
| | UBE2E3 |
| | KLHL2 |
| | WIPI2 |
| | TMEM9B |
| | PKIA |

| | |
|---|---|
| | CRYM |
| | AK2 |
| | NFIA |
| | UBE2Q1 |
| | UBQLN4 |
| | NSL1 |
| | GALNT2 |
| | SMPD4 |
| | PLCL1 |
| | TMEM237 |
| | AZI2 |
| | EPM2AIP1 |
| | KIF9 |
| | HS3ST1 |
| | RBPJ |
| | ARAP2 |
| | ZFYVE16 |
| | CHD1 |
| | GM2A |
| | UBLCP1 |
| | NUB1 |
| | UBE3C |
| | PRAG1 |
| | GPAT4 |
| | MRPS28 |
| | ZHX1 |
| | MTSS1 |
| | MROH1 |
| | KANK1 |
| | TLN1 |
| | ZNG1E |
| | GPR107 |
| | RUFY2 |
| | PIK3C2A |
| | LPXN |
| | STARD10 |
| | ZC3H12C |
| | TIGAR |
| | GPR162 |
| | DENND5B |
| | DENR |
| | HIP1R |
| | MIA2 |

| |
|---|
| ZDHHC22 |
| OTUD7A |
| RNF111 |
| MRPS11 |
| MCRIP2 |
| SMG1 |
| LGALS9 |
| PLEKHM1 |
| SAMD1 |
| APOL6 |
| ARFGAP3 |
| CERK |
| CIAO1 |
| POLR3GL |
| ROCK2 |
| PHACTR1 |
| UBE2I |
| PSMD4 |
| CEP170 |
| RAB3C |
| CWC15 |
| TMED2 |
| NOC2L |
| ATAD3A |
| PLEKHG5 |
| SLC45A1 |
| CORT |
| MAD2L2 |
| NPPA |
| PLOD1 |
| CASP9 |
| NBPF1 |
| CROCCP2 |
| IFFO2 |
| MUL1 |
| FAM43B |
| KIF17 |
| ZNF436 |
| PAQR7 |
| CEP85 |
| UBXN11 |
| RPS6KA1 |
| PIGV |

| |
|---|
| SLC9A1 |
| GPR3 |
| LINC02574 |
| RCC1 |
| TMEM39B |
| CCDC28B |
| AZIN2 |
| SH3D21 |
| SNIP1 |
| POU3F1 |
| CFAP57 |
| CCDC24 |
| TMEM53 |
| PTCH2 |
| TESK2 |
| FAAH |
| OSBPL9 |
| YIPF1 |
| DAB1 |
| ATG4C |
| CACHD1 |
| TYW3 |
| PIGK |
| SPATA1 |
| RPAP2 |
| SNX7 |
| DBT |
| CDC14A |
| GSTM4 |
| CEPT1 |
| WNT2B |
| ST7L |
| PTGFRN |
| ZNF697 |
| SRGAP2B |
| PEX11B |
| NBPF11 |
| NOTCH2NLC |
| PRUNE1 |
| S100A9 |
| THBS3-AS1 |
| GON4L |
| SLC25A44 |

| |
|---|
| PEAR1 |
| NIT1 |
| PCP4L1 |
| SLC19A2 |
| CENPL |
| B3GALT2 |
| KCNT2 |
| UBE2T |
| SYT2 |
| KDM5B |
| ETNK2 |
| RBBP5 |
| CAMK1G |
| SERTAD4-AS1 |
| GPATCH2 |
| DUSP10 |
| LBR |
| SDE2 |
| B3GALNT2 |
| CHML |
| DESI2 |
| KIF26B |
| ZNF496-DT |
| RNASEH1-DT |
| SOX11 |
| IAH1 |
| SLC66A3 |
| KCNS3 |
| DNAJC27 |
| EFR3B |
| POMC |
| TMEM214 |
| EIF2B4 |
| ZNF513 |
| TRMT61B |
| LCLAT1 |
| MEMO1 |
| LTBP1 |
| HEATR5B |
| CAMKMT |
| EML6 |
| PEX13 |
| FAM161A |

| |
|---|
| AFTPH |
| C2orf42 |
| TGFA |
| RAB11FIP5 |
| HK2 |
| TACR1 |
| ELMOD3 |
| USP39 |
| KCNIP3 |
| ANKRD36B |
| C2orf92 |
| INPP4A |
| REV1 |
| TBC1D8 |
| CNOT11 |
| LINC01106 |
| TMEM177 |
| IWS1 |
| AMMECR1L |
| TMEM163 |
| ACVR2A |
| EPC2 |
| GALNT13 |
| COBLL1 |
| LRP2 |
| BBS5 |
| GORASP2 |
| DLX1 |
| GPR155 |
| LNPK |
| COL5A2 |
| TYW5 |
| ALS2 |
| INO80D |
| ADAM23 |
| ERBB4 |
| MREG |
| IGFBP2 |
| ZNF142 |
| DNPEP |
| OBSL1 |
| IQCA1 |
| DUSP28 |

| |
|---|
| THAP4 |
| LMCD1 |
| OGG1 |
| IL17RC |
| HRH1 |
| NUP210 |
| FBLN2 |
| NKIRAS1 |
| THRB |
| ENTPD3-AS1 |
| LIMD1 |
| LZTFL1 |
| KLHL18 |
| COL7A1 |
| CELSR3 |
| LAMB2 |
| AMT |
| RNF123 |
| SLC38A3 |
| PPM1M |
| DNAH1 |
| STAB1 |
| STIMATE |
| RFT1 |
| PRKCD |
| DENND6A |
| RPP14 |
| PRICKLE2 |
| MITF |
| GPR27 |
| SHQ1 |
| FAM86DP |
| ROBO1 |
| C3orf38 |
| CRYBG3 |
| FILIP1L |
| LINC00882 |
| DUBR |
| ABHD10 |
| CFAP44 |
| SIDT1 |
| DTX3L |
| SLC49A4 |

| |
|---|
| ADCY5<br>OSBPL11<br>TMCC1<br>TMCC1-DT<br>CPNE4<br>PPP2R3A<br>CHST2<br>HPS3<br>C3orf80<br>ACTL6A<br>USP13<br>PARL<br>CLCN2<br>EPHB3<br>RPL39L<br>MASP1<br>ATP13A5<br>LSG1<br>SDHAP1<br>CEP19<br>MELTF<br>PIGG<br>TMEM129<br>NOP14<br>GRK4<br>MSANTD1<br>JAKMIP1<br>TADA2B<br>ANAPC4<br>NWD2<br>GUF1<br>EPHA5<br>UBA6-DT<br>TMEM150C<br>HERC5<br>BDH2<br>C4orf33<br>PCDH10-DT<br>MGARP<br>NAA15<br>ZNF330<br>TLR2<br>DDX60 |

| | |
|---|---|
| | SH3RF1<br>CEP44<br>WWC2<br>EXOC3-AS1<br>SLC9A3-AS1<br>ZDHHC11B<br>ZDHHC11<br>FASTKD3<br>OTULIN<br>RETREG1<br>CDH18<br>CDH6<br>C5orf22<br>AMACR<br>TMEM267<br>C5orf34<br>CCNO<br>ERCC8<br>SMIM15<br>RGS7BP<br>RAD17<br>FCHO2<br>UTP15<br>ARHGEF28<br>MSH3<br>ATG10<br>TMEM161B<br>LNPEP<br>WDR36<br>CCDC112<br>CEP120<br>MARCHF3<br>SLC12A2-DT<br>RAPGEF6<br>SOWAHA<br>C5orf15<br>CDC23<br>PCDHB10<br>HDAC3<br>PCDH1<br>SPRY4<br>YIPF5<br>GEMIN5 |

| |
|---|
| MAT2B |
| TENM2 |
| SPDL1 |
| SIMC1 |
| UNC5A |
| UIMC1 |
| PRR7 |
| RASGEF1C |
| SCGB3A1 |
| BPHL |
| PSMG4 |
| RPP40 |
| HIVEP1 |
| RNF182 |
| H1-2 |
| ZNF322 |
| ZNF184 |
| ZSCAN31 |
| DXO |
| ATF6B |
| HLA-DRB5 |
| TAP2 |
| WDR46 |
| ZNF76 |
| TBC1D22B |
| SAYSD1 |
| LINC00951 |
| TRERF1 |
| BICRAL |
| ABCC10 |
| EFHC1 |
| BEND6 |
| RAB23 |
| KHDC1-AS1 |
| LCA5 |
| SLC35A1 |
| COQ3 |
| ASCC3 |
| SESN1 |
| PKIB |
| PEX7 |
| CCDC28A |
| STXBP5-AS1 |

| |
|---|
| LATS1 |
| NUP43 |
| PPP1R14C |
| PLEKHG1 |
| RMND1 |
| MRPL18 |
| CAHM |
| C6orf120 |
| TBP |
| DNAAF5 |
| ZFAND2A-DT |
| INTS1 |
| PSMG3 |
| CHST12 |
| LFNG |
| MMD2 |
| TNRC18 |
| MIOS |
| UMAD1 |
| ICA1 |
| NUP42 |
| OSBPL3 |
| CPVL |
| MATCAP2 |
| EPDR1 |
| GLI3 |
| POLD2 |
| FIGNL1 |
| MRPS17 |
| PSPH |
| KCTD7 |
| NSUN5P2 |
| MLXIPL |
| GTF2IRD1 |
| GTF2IP1 |
| SRRM3 |
| ADAM22 |
| CDK14 |
| FZD1 |
| GATAD1 |
| RBM48 |
| TFPI2 |
| COL1A2 |

| | |
|---|---|
| | ZNF789<br>ZKSCAN5<br>TRIM4<br>ZNF3<br>PILRB<br>PCOLCE<br>ACHE<br>LRWD1<br>CCDC71L<br>COG5<br>GPR22<br>IMMP2L<br>WASL-DT<br>POT1<br>TNPO3<br>COPG2<br>CLEC2L<br>TRBC2<br>CUL1<br>ZNF282<br>ZNF777<br>ZNF775<br>ABCB8<br>FBXO25<br>MYOM2<br>MSRA<br>MTMR7<br>ASAH1-AS1<br>LZTS1<br>GFRA2<br>POLR3D<br>PDLIM2<br>BIN3<br>ZNF395<br>RNF122<br>BRF2<br>ANK1<br>PCMTD1<br>FAM110B<br>SGK3<br>LY96<br>JPH1<br>CRISPLD1 |

| |
|---|
| LRRCC1 |
| MMP16 |
| MATN2 |
| STK3 |
| BAALC-AS1 |
| EXT1 |
| CCN3 |
| DEPTOR |
| NTAQ1 |
| FAM91A1 |
| KCNK9 |
| PEG13 |
| CHRAC1 |
| ZC3H3 |
| NAPRT |
| TIGD5 |
| GFUS |
| SCRT1 |
| SLC39A4 |
| LRRC24 |
| ZNF7 |
| GLIS3 |
| RCL1 |
| HAUS6 |
| ELAVL2 |
| TUSC1 |
| TOPORS |
| GALT |
| IL11RA |
| SLC25A51 |
| ALDH1B1 |
| LERFS |
| ANKRD20A1 |
| PIP5K1B |
| TRPM3 |
| SECISBP2 |
| ZNF510 |
| MFSD14CP |
| NIPSNAP3B |
| ABITRAM |
| ASTN2 |
| TRIM32 |
| CNTRL |

| |
|---|
| DENND1A |
| SCAI |
| SWI5 |
| DOLK |
| CRAT |
| TOR1B |
| GTF3C4 |
| PPP1R26 |
| INPP5E |
| NOXA1 |
| DPH7 |
| ARRDC1 |
| EHMT1 |
| LINC02668 |
| ANKRD16 |
| SFTA1P |
| SEC61A2 |
| MSANTD7 |
| NMT2 |
| C1QL3 |
| SLC39A12 |
| NSUN6 |
| NEBL-AS1 |
| KIAA1217 |
| ENKUR |
| GPR158 |
| PDSS1 |
| SVIL-AS1 |
| PARD3 |
| LINC00839 |
| AGAP4 |
| ARHGAP22 |
| TMEM273 |
| ERCC6 |
| PARG |
| PCDH15 |
| LRRTM3 |
| SGPL1 |
| VCL |
| ADK |
| LRMDA |
| POLR3A |
| KIF20B |

| | |
|---|---|
| | PLCE1 |
| | ALDH18A1 |
| | ZNF518A |
| | R3HCC1L |
| | ERLIN1 |
| | DPCD |
| | POLL |
| | KCNIP2 |
| | NFKB2 |
| | SUFU |
| | SORCS1 |
| | NHLRC2 |
| | GPR26 |
| | FAM53B |
| | ABRAXAS2 |
| | C10orf143 |
| | ADGRA1 |
| | TUBGCP2 |
| | PTDSS2 |
| | PIDD1 |
| | RIC3 |
| | DENND2B |
| | AMPD3 |
| | SAAL1 |
| | ZDHHC13 |
| | HTATIP2 |
| | LUZP2 |
| | CCDC34 |
| | TCP11L1 |
| | FBXO3 |
| | PAMR1 |
| | LINC02696 |
| | LRRC10B |
| | TUT1 |
| | TMEM223 |
| | FLRT1 |
| | MEN1 |
| | SIPA1 |
| | AP5B1 |
| | EFEMP2 |
| | SYT12 |
| | SSH3 |
| | KMT5B |

| |
|---|
| C11orf24 |
| TPCN2 |
| LRRC51 |
| ATG16L2 |
| LRRC32 |
| CEP57 |
| LAYN |
| C11orf71 |
| APOA1 |
| TMEM25 |
| ZNF202 |
| STT3A |
| HYLS1 |
| RPUSD4 |
| SLC6A13 |
| PARP11 |
| KCNA1 |
| ZNF384 |
| LRRC23 |
| STYK1 |
| CREBL2 |
| SLCO1C1 |
| RECQL |
| C2CD5-AS1 |
| INTS13 |
| FGD4 |
| YARS2 |
| CPNE8 |
| LRRK2 |
| DBX2 |
| KANSL2 |
| DDX23 |
| PRKAG1 |
| MFSD5 |
| DGKA |
| PAN2 |
| SHMT2 |
| TSFM |
| WIF1 |
| MSRB3 |
| GRIP1 |
| FRS2 |
| PTPRR |

| |
|---|
| KCNC2 |
| CAPS2 |
| MGAT4C |
| KITLG |
| CEP83 |
| NOPCHAP1 |
| PRDM4 |
| SART3 |
| IFT81 |
| CUX2 |
| BRAP |
| TRIAP1 |
| POP5 |
| BCL7A |
| ZCCHC8 |
| SBNO1-AS1 |
| PXMP2 |
| NHLRC3 |
| VWA8 |
| ENOX1 |
| KCTD4 |
| TPT1-AS1 |
| PHF11 |
| PCDH20 |
| MZT1 |
| KLF5 |
| UCHL3 |
| COL4A2 |
| NAXD |
| TUBGCP3 |
| CCNB1IP1 |
| METTL17 |
| HAUS4 |
| ZFHX2 |
| IPO4 |
| PRKD1 |
| DTD2 |
| NUBPL |
| MBIP |
| TRAPPC6B |
| LRFN5 |
| FRMD6 |
| GNPNAT1 |

| |
|---|
| L3HYPDH |
| TRMT5 |
| PLEKHG3 |
| SPTB |
| GPHN |
| RIOX1 |
| ISCA2 |
| GPATCH2L |
| PTPN21 |
| LYSET |
| SIVA1 |
| JAG2 |
| TMEM121 |
| HERC2P9 |
| ENTREP2 |
| LINC02352 |
| KATNBL1 |
| PLCB2 |
| BAHD1 |
| GCHFR |
| NDUFAF1 |
| TMEM62 |
| HYPK |
| FRMD5 |
| CTDSPL2 |
| SLC28A2-AS1 |
| SQOR |
| COPS2 |
| AP4E1 |
| LACTB |
| AAGAB |
| CLN6 |
| PAQR5 |
| TLE3 |
| CELF6 |
| INSYN1 |
| ISLR2 |
| ISLR |
| EDC3 |
| ARNT2-DT |
| SNHG21 |
| HOMER2 |
| AEN |

| |
|---|
| FAM174B |
| ALDH1A3 |
| ALDH1A3-AS1 |
| LMF1 |
| NTHL1 |
| BRICD5 |
| TIGD7 |
| PPL |
| SNX29 |
| NPIPA1 |
| MPV17L |
| SYT17 |
| IGSF6 |
| COG7 |
| ARHGAP17 |
| SMG1P5 |
| SEPTIN1 |
| ZNF747 |
| ZNF629 |
| SETD1A |
| CNEP1R1 |
| HEATR3 |
| TENT4B |
| SALL1 |
| HNRNPA1L3 |
| MT1A |
| KIFC3 |
| TK2 |
| CA7 |
| HSF4 |
| SLC9A5 |
| CARMIL2 |
| SMPD3 |
| FCSK |
| HYDIN |
| ATXN1L |
| LDHD |
| SYCE1L |
| NECAB2 |
| MEAK7 |
| GSE1 |
| FBXO31 |
| ZC3H18 |

| |
|---|
| GALNS |
| DPH1 |
| MED11 |
| PLD2 |
| MIS12 |
| PLSCR3 |
| TNFSF12 |
| SOX15 |
| WRAP53 |
| TMEM88 |
| KCNAB3 |
| TMEM107 |
| PIK3R5 |
| TRPV2 |
| MED9 |
| AKAP10 |
| DHRS7B |
| PYY2 |
| UNC119 |
| SPAG5 |
| TLCD1 |
| DHRS13 |
| ZNF830 |
| RASL10B |
| CCL5 |
| CWC25 |
| FBXL20 |
| CDK12 |
| P3H4 |
| RUNDC1 |
| G6PC3 |
| FMNL1-AS1 |
| SP2 |
| COPZ2 |
| ITGA3 |
| LINC01476 |
| INTS2 |
| TLK2 |
| CEP95 |
| SMURF2 |
| HELZ |
| C17orf58 |
| KCNJ16 |

| |
|---|
| SLC39A11 |
| GGA3 |
| TSEN54 |
| RECQL5 |
| UBE2O |
| PRCD |
| METTL23 |
| BAHCC1 |
| HEXD |
| METTL4 |
| ZBTB14 |
| ARHGAP28-AS1 |
| RNMT |
| TTC39C |
| TRAPPC8 |
| ZSCAN30 |
| PSTPIP2 |
| ZNF407-AS1 |
| ZNF407 |
| CFD |
| TCF3 |
| IZUMO4 |
| LMNB2 |
| APBA3 |
| TNFAIP8L1 |
| KDM4B |
| DUS3L |
| CAPS |
| MAP2K7 |
| ZNF414 |
| ZNF561 |
| COL5A3 |
| SHFL |
| TYK2 |
| ECSIT |
| ZNF440 |
| ZNF791 |
| RTBDN |
| NANOS3 |
| ZNF333 |
| MVB12A |
| FCHO1 |
| JAK3 |

| |
|---|
| CRLF1 |
| HAPLN4 |
| ZNF430 |
| ZNF738 |
| ZNF493 |
| ZNF98 |
| C19orf12 |
| GARRE1 |
| ZNF181 |
| ZNF875 |
| SARS2 |
| PAK4 |
| LRFN1 |
| PLEKHG2 |
| DYRK1B |
| C19orf47 |
| SHKBP1 |
| BCKDHA |
| LIPE-AS1 |
| IRGQ |
| ZNF226 |
| BCL3 |
| PPP1R37 |
| TRAPPC6A |
| RTN2 |
| LIG1 |
| LMTK3 |
| LIN7B |
| BCL2L12 |
| CPT1C |
| FUZ |
| ASPDH |
| ZNF160 |
| RDH13 |
| ZNF524 |
| ZNF444 |
| ZFP28 |
| ZNF548 |
| ZNF419 |
| ZNF550 |
| ZIK1 |
| ZNF530 |
| ZNF154 |

| |
|---|
| ZNF8 |
| ZNF446 |
| TRIB3 |
| PLCB4 |
| NDUFAF5 |
| MACROD2 |
| CRNKL1 |
| GZF1 |
| ACSS1 |
| HCK |
| PLAGL2 |
| CNBD2 |
| ACTR5 |
| STK4 |
| ZNF335 |
| CDH22 |
| SLC35C2 |
| ARFGEF2 |
| CBLN4 |
| MTG2 |
| ARFGAP1 |
| LIME1 |
| SLC2A4RG |
| SAMD10 |
| RBM11 |
| RWDD2B |
| IFNAR2 |
| IL10RB |
| GART |
| SMIM11 |
| HLCS |
| POFUT2 |
| SPATC1L |
| BCL2L13 |
| MICAL3 |
| ESS2 |
| GNB1L |
| ARVCF |
| TRMT2A |
| P2RX6 |
| SLC7A4 |
| TMEM191C |
| SLC2A11 |

| |
|---|
| ADORA2A |
| KIAA1671 |
| TFIP11-DT |
| MN1 |
| HSCB |
| RHBDD3 |
| NIPSNAP1 |
| OSBP2 |
| MORC2 |
| RFPL2 |
| SYN3 |
| RASD2 |
| RAC2 |
| KCNJ4 |
| CBY1 |
| DNAL4 |
| CACNA1I |
| L3MBTL2 |
| TCF20 |
| RRP7BP |
| WNT7B |
| PPARA |
| TBC1D22A-DT |
| BRD1 |
| TRABD |
| NCAPH2 |
| CPT1B |
| SHANK3 |
| RPL23AP7 |
| AKAP17A |
| DHRSX |
| SHROOM2 |
| RAI2 |
| BCLAF3 |
| MBTPS2 |
| XK |
| NDP |
| EBP |
| GPKOW |
| GPR173 |
| SPIN2B |
| ZC4H2 |
| OPHN1 |

| | |
|---|---|
| | ZMYM3 |
| | CITED1 |
| | SLC16A2 |
| | DACH2 |
| | PCDH11X |
| | DIAPH2 |
| | ZMAT1 |
| | TMSB15B |
| | ZCCHC18 |
| | RADX |
| | MID2 |
| | CHRDL1 |
| | NKRF |
| | TENM1 |
| | OCRL |
| | RAP2C |
| | SLITRK2 |
| | EOLA2 |
| | MAMLD1 |
| | GABRA3 |
| | ZNF275 |
| | IRAK1 |
| | ATP6AP1-DT |
| | F8 |
| | KDM5D |
| | FCGRT |
| | GLRX5 |
| | NFIX |
| | HINT3 |
| | USP46 |
| | ERF |
| | LRATD1 |
| | RAB13 |
| | ACIN1 |
| | PPP4R1 |
| | ROCK1 |
| | GLS |
| | ALDH7A1 |
| | RBM17 |
| | CDIPT |
| | HM13 |
| | GCC2 |
| | HGS |

| | |
|---|---|
| | KTN1 |
| | VWA1 |
| | SIPA1L2 |
| | DARS1 |
| | ANXA2 |
| | DPY30 |
| | TIMP3 |
| | ACBD5 |
| | RERE |
| | SEMA4D |
| | SESN3 |
| | SCAF11 |
| | CYSTM1 |
| | IRF1 |
| | RAB22A |
| | MAPKAPK2 |
| | GOLGA7 |
| | RBM3 |
| | PDHB |
| | FAM171A1 |
| | NRXN3 |
| | PRDX3 |
| | AKAP6 |
| | ARGLU1 |
| | HAVCR2 |
| | ERBIN |
| | MGAT4A |
| | UBTD2 |
| | RPA2 |
| | MECR |
| | PPIH |
| | ARHGEF11 |
| | SOAT1 |
| | TSEN15 |
| | ADORA1 |
| | AHCTF1 |
| | CRIM1-DT |
| | ST3GAL5 |
| | MKRN2 |
| | GLB1 |
| | FIP1L1 |
| | RAD50 |
| | FBLL1 |

| |
|---|
| NSD1 |
| HEIH |
| ECHDC1 |
| SNX8 |
| NT5C3A |
| EXOC4 |
| DCAF10 |
| TMEM250 |
| TASOR2 |
| MGMT |
| COPB1 |
| PTPRJ |
| MARK2 |
| FRMD8 |
| ZPR1 |
| H2AJ |
| SMUG1 |
| ZNF84 |
| INTS6 |
| MGAT2 |
| FBXO34 |
| MEG8 |
| CEMIP |
| GRIN2A |
| USP31 |
| LPCAT2 |
| COQ9 |
| VPS9D1 |
| FMNL1 |
| SEPTIN9 |
| TNPO2 |
| PKN1 |
| HSD17B14 |
| ARHGEF17 |
| DSCAML1 |
| SUCLA2 |
| SNRNP25 |
| TRAPPC1 |
| PDK2 |
| PPFIA3 |
| CBX5 |
| BSDC1 |
| NR2F1 |

| |
|---|
| PLEKHA1 |
| KMT2A |
| ZNF385A |
| MARK3 |
| TXNDC17 |
| EPN2 |
| SH3GL1 |
| ICAM5 |
| RAB33A |
| VPS13D |
| LRRC7 |
| AKT1 |
| PSMD13 |
| CYTOR |
| GTF2I |
| SBNO2 |
| KPNA6 |
| SEMA4C |
| KCTD12 |
| LZIC |
| CDC73 |
| CCNYL1 |
| MYLK |
| AHCYL2 |
| RASSF4 |
| CCDC82 |
| GNPTAB |
| SDS |
| MICU2 |
| DAAM1 |
| FURIN |
| RPUSD1 |
| GAD1 |
| CLPTM1L |
| C6orf89 |
| MFN2 |
| EMC4 |
| CMTM6 |
| HCLS1 |
| EIF2S1 |
| NCOR1 |
| MAOB |
| FLNA |

| | |
|---|---|
| | VTI1B |
| | BPNT2 |
| | NRBP1 |
| | PSMB6 |
| | RPH3A |
| | VPS29 |
| | CD164 |
| | MSRB2 |
| | NECTIN1 |
| | OSER1 |
| | NAA10 |
| | C22orf39 |
| | MDH2 |
| | DCTN6 |
| | ADI1 |
| | PAK2 |
| | ABLIM1 |
| | MAFG |
| | AATK |
| | TLCD4 |
| | PIGX |
| | ANAPC15 |
| | FBXL3 |
| | WHAMM |
| | HSPA1B |
| | USP14 |
| | UNC13B |
| | EGLN2 |
| | H2AZ2 |
| | BCAN |
| | BTBD2 |

| Donor 3 | | | Donor 4 | | |
|---|---|---|---|---|---|
| Cortical layer 2/3 | | | Cortical layer 1 | | |
| iTBS | iTBS ∩ 10 Hz | 10 Hz | iTBS | iTBS ∩ 10 Hz | 10 Hz |
| MBP | MT1G | ADAMTS1 | NRGN | LINC00632 | MBP |
| PLP1 | PCDH8 | UQCR11 | RPL39 | LAMP5 | GFAP |
| CRYAB | CARTPT | CD74 | RPL27A | SELENOP | PLP1 |
| GFAP | HS6ST3 | PFKFB3 | CSRP1 | RRP12 | MT1X |
| MOBP | NPY | NTRK2 | PPP2CB | TTYH2 | CNP |
| EGR4 | LINC00507 | BLVRA | MALAT1 | CBR1 | PVALB |
| APOD | HS3ST4 | PFDN5 | DNAJB1 | RNASE1 | MT1E |
| LAMP5 | C14orf132 | HINT1 | SNORC | PPP1R14A | APOD |
| MAG | LPAR1 | CLIC4 | ARC | PRPSAP1 | SCD |
| HSPH1 | NSG1 | HEPACAM | HSPA1B | ISLR | CCL3 |
| CAMK2A | ITPKA | CERCAM | YPEL3 | MLF1 | MT1G |
| HOPX | TMEM144 | ADGRG1 | DEDD2 | GAB1 | DBNDD2 |
| VGF | CALB1 | NCOA7 | SERP2 | MTHFS | QKI |
| CKS2 | RSBN1L | VPS26B | HSPA6 | ADGRL2 | CRYAB |
| PMP22 | ETV1 | DDX3Y | MYO9B | ZNF689 | TUBB4A |
| NECAB2 | TMIGD3 | C9orf78 | HSPA1A | BAG3 | INA |
| HS3ST2 | GRIA4 | GLS | KCNF1 | HPS1 | MT1M |
| SLC12A5 | RASD2 | BCAN | SPRING1 | CRY1 | TUBA1A |
| CALB2 | SCIN | AIF1L | SNHG25 | CHAC1 | MOBP |
| PTGS2 | PICK1 | COPS5 | KMT5A | FAM136A | TF |
| PSD | PLPP5 | SF1 | ANKRD33B | PCGF1 | CARTPT |
| SOWAHA | PDZD2 | NASP | ECD | SLC30A4 | CKS2 |
| BCAS1 | CBLN4 | DARS1 | PRMT2 | EIF3C | CAMK2N2 |
| ANKRD37 | TSHZ2 | MFGE8 | SET | NUDT7 | VIM |
| GABRD | PRAG1 | SORL1 | DHCR7 | SHROOM2 | CCL3L3 |
| MET | MLXIP | TLE4 | PAM16 | AEBP1 | CCL4L2 |
| CARNS1 | FECH | ETNPPL | RASSF2 | EMP3 | VSTM2A |
| RASGRF2 | PCSK6 | PHLDA1 | C14orf119 | MYL9 | BCAS1 |
| FAM107B | ANKRD18A | CLK1 | CHD9 | DCTN6-DT | LINC00844 |
| HAPLN4 | TMEM88B | SEPTIN8 | AKT3 | PLPPR1 | HIVEP2 |
| VSTM2A | DLGAP2 | GIT2 | MOB4 | PPP2R5A | HAPLN4 |
| NOP58 | ZFYVE28 | ELP3 | TCEAL4 | PAMR1 | PLEKHB1 |
| SELENOP | TAFA1 | CLASP2 | GAP43 | ADAM9 | STRBP |
| CCN1 | MORC3 | SH3TC2 | ST6GALNAC5 | MATN2 | SYT2 |
| EFTUD2 | RBP4 | NUCKS1 | LARP7 | JCAD | MT1F |
| TTYH2 | KIF13B | HERC4 | TUBB | SLC9A7 | GAD1 |
| DIRAS2 | PTPRO | CAMSAP2 | GASK1B | CARNS1 | CCL4 |
| LIPA | RFX3 | CAST | HSBP1 | CCDC24 | HSBP1L1 |

| | | | | | |
|---|---|---|---|---|---|
| FBXL7 | MVP | GOLGA8A | EGR2 | SEC22A | MIAT |
| SLC17A6 | DZIP1 | TMX4 | KCTD12 | KLF3 | KCNIP2 |
| CYP46A1 | OSBP2 | PLIN2 | G3BP1 | SYNPO2 | KLF9 |
| DUSP16 | TOM1 | CORO2B | FAM3A | PRR18 | PDZRN3 |
| OPALIN | ERBB4 | PDE4B | STX1A | FAM120B | TAC3 |
| CUX2 | TMTC4 | KIF1C | HSPD1 | BET1 | AQP1 |
| PRKACA | ACKR1 | ELL2 | CMC1 | RASL11A | CLDND1 |
| WDR26 | HHIP | PPP3R1 | KCNN2 | VIPAS39 | SCRT1 |
| RNASE1 | ZC3H12A | ATP13A1 | GABRA5 | ZNF540 | PRKAG2-AS1 |
| LINC00844 | ABCC8 | C11orf71 | SYNPR | RBSN | MAPK6 |
| MAP3K3 | ZFYVE16 | YIPF4 | TSC22D3 | MAG | SLC43A2 |
| CPSF6 | FAM153A | CEROX1 | PMEPA1 | ABCD4 | BCAN |
| CCL3 | SSH3 | PTPN12 | APBA2 | TMEM38A | CLDN11 |
| SSTR2 | KRT17 | SF3B2 | NDUFV3 | WASHC3 | RBM48 |
| DDN | STX10 | FHL2 | SUMO2 | LINC00641 | MARCKSL1 |
| PRDM8 | RABL2A | SEPHS1 | POLR2K | SNHG10 | MGAT5B |
| SPTB | ZBTB5 | ATP5IF1 | PHF20L1 | STUM | MFSD4A |
| RASGRP3 | ZNF780B | MAP3K7 | SRP54 | SLC41A2 | TMEM144 |
| MAN1C1 | GRM7 | MRPL34 | PRPS1 | HSD17B7 | MT1A |
| LRATD1 | METTL25B | PTN | MAN1A2 | BTAF1 | EPHA4 |
| PIANP | CDH10 | RGS12 | CAP2 | SF3A3 | SNAPC2 |
| AMPD2 | CDS1 | DGKZ | MAPKAP1 | ENTPD3-AS1 | SPATS2L |
| ADCYAP1 | NOL4 | TSR1 | SMIM13 | LINC02482 | SOX2-OT |
| FNBP1 | BBC3 | FAF1 | DMD | BRAT1 | C1orf35 |
| YJEFN3 | ASS1 | PREX2 | POMGNT1 | ZBED5 | DOP1A |
| AZGP1 | UBR4 | MICAL2 | ABL2 | FIBIN | LHPP |
| DOCK5 | SQLE | SNRNP200 | BMPR2 | RPP25 | NFKB2 |
| GLDN | AQP1 | LSM7 | FAM3C | ZNF512B | MIEF1 |
| SDC4 | KCNQ4 | ANAPC5 | NIF3L1 | RNASEH1 | AQP4 |
| MAP3K11 | FLYWCH1 | FMNL2 | PDCD10 | CDKN1C | MAL |
| THEMIS | POMGNT2 | NOP2 | ZNF853 | SRXN1 | PAWR |
| RCN1 | CD14 | PHC1 | SMURF1 | SMC5 | P2RX6 |
| ZNF184 | SDS | NELFB | AHCYL2 | SEMA5B | ADAM22 |
| HSPA4L | PPFIA4 | LENG8 | TRIM37 | STAG1 | DAPK1 |
| RIPK1 | RAB26 | CDH13 | MARCHF9 | TMEM243 | ANKRD10 |
| XK | SRGAP1 | XIAP | EIF4EBP2 | CDKL1 | ESYT1 |
| RAB3B | MPP7 | DENND4C | SEC14L2 | UBAP1L | AIFM3 |
| MAL | PHLDA2 | ODF2L | AP1G1 | FES | CASKIN1 |
| FAM234B | NKX6-2 | THAP12 | PTPRN2 | NUBP1 | NRIP3 |
| GPR88 | FBXW4 | TFCP2 | ICAM5 | THOC1 | MAP3K20 |
| HECTD2 | PPP1R16A | NDUFA9 | TRIB1 | CYBA | HTT |
| BUD23 | CDH19 | SYNCRIP | POLD4 | VPS11 | ENPP2 |
| RXFP1 | SCYL1 | GOLGA4 | NFYB | CACNA1C | CCSAP |

| | | | | | |
|---|---|---|---|---|---|
| CACNG8 | COL5A2 | ABCA1 | AKAP1 | ATF7IP2 | PAQR6 |
| ZNF711 | GPR180 | RAB6A | RAVER2 | FBXO25 | ANKRD37 |
| SNHG3 | KCNH1 | REEP1 | UNC13C | MIR124-2HG | TAFA1 |
| TMEM65 | B4GALT6 | LINC-PINT | LIPE-AS1 | PTPN2 | UFM1 |
| NTNG1 | COL12A1 | CAPN3 | ITGAE | SOX8 | ZFAND2A |
| PDE1B | SPACDR | SNHG8 | FMR1 | UBTD2 | TRRAP |
| B4GALNT1 | SPTSSB | SMG7 | KNDC1 | TRMT61B | PPP3CC |
| TYW5 | PYDC2-AS1 | ODC1 | DCLK2 | GALNT13 | SNHG1 |
| ANKRD42 | BAG2 | TMEM151B | BOLA3 | EPHA5 | POMT1 |
| FGD1 | STK17B | NANS | VPS29 | UBA6-DT | EFNA3 |
| CHORDC1 | MED23 | SNAPC2 | FUT8 | ADCY8 | RTL5 |
| LINC02610 | GPR83 | NPC2 | ARL2BP | ZNF488 | HGF |
| PKNOX1 | TKFC | COG1 | RMND5A | CASP7 | PDZD8 |
| DKK1 | NAXD | SLC39A6 | FMC1 | CNTN5 | CDR2L |
| RASSF2 | MRPL53 | LRIG1 | CHORDC1 | TMEM132D | PI4K2A |
| GPHN | RIMBP2 | CRIPT | MEX3C | NOP9 | MISP3 |
| TAFA5 | GLRA3 | UBE2N | VPS13D | L2HGDH | WIF1 |
| GRIN2B | RAB9B | HABP4 | ZBTB44 | ZBTB14 | TRMT44 |
| GREM1 | MBIP | TIAM1 | MED13 | SERPINB8 | INTS6 |
| NKAIN2 | SH3BP1 | ACADVL | TPM2 | SIPA1L3 | HMGB2 |
| OSBPL8 | SNRPA | DNAJC12 | DPM3 | C1GALT1C1 | ACSL6 |
| SPTBN4 | LZTS1 | STX1A | NRDC | ITGB5 | NT5C2 |
| RGMB | MIER2 | PKN1 | FADS2 | CTTNBP2 | APLNR |
| MIR29B2CHG | CDC42EP3 | SLC12A2-DT | KCNQ2 | NACAD | PISD |
| PNCK | KCNH8 | INPP5K | SEC63 | LATS1 | CTSA |
| TMEM74B | IFI30 | LIF | RGS2 | ITGA7 | PLAAT3 |
| BAMBI | HIBCH | DBP | SLC11A2 | FAM241B | PEX19 |
| IVD | GSTM5 | COX7A1 | TJP1 | COL5A2 | TBC1D23 |
| PCDH19 | HAS1 | SPATA13 | BCS1L | RASA2 | SCN4B |
| XXYLT1 | CRLF1 | CCDC50 | RNF25 | CSNK1G3 | WDR37 |
| CHST15 | ENGASE | FRMD4B | SETD9 | KYAT1 | ALMS1 |
| NT5DC3 | SERPINE1 | TMEM50A | SMIM8 | TBK1 | EEFSEC |
| BOK | CRH | CYTH1 | MTAP | HOMER2 | C3orf18 |
| MYO1B | TAGAP-AS1 | GRIA1 | COL9A3 | LMF1 | ADK |
| GRIK1 | CACNA1G | RMDN3 | MAGT1 | ZNF585A | NDUFAF6 |
| FAM32A | CFAP298 | FASTKD2 | DGUOK | USP16 | ST3GAL5 |
| SHANK3 | OBSCN | GNPDA2 | ACBD7 | CLEC11A | CAMK1G |
| PREX1 | KCNN2 | RPP25L | SNTB2 | PRDM16 | KIFC3 |
| ABLIM2 | AFF2 | COQ6 | VCAN | FBXO6 | MPDU1 |
| NPM3 | DIPK1B | MIR7-3HG | ZCCHC24 | ADGRL4 | SGSM1 |
| STAG1 | FKBP15 | MED7 | ASH1L | HCN3 | RALGAPA1 |
| SCPEP1 | SHC4 | KCTD21 | YTHDC1 | WNT7A | GALT |
| TXN2 | SLC24A4 | CUL5 | SNHG3 | TMIE | VIPR2 |

| | | | | | |
|---|---|---|---|---|---|
| LINC01202 | VPS37D | BRF1 | RAPGEF4 | RIOX1 | ZNF195 |
| GRHL1 | MACROD1 | HDDC3 | MED28 | AVEN | HNRNPUL1 |
| YIF1A | RANBP10 | THAP7 | EMC1 | TRIP4 | GMPS |
| TMEM179B | ZRSR2 | PRXL2A | ZNF687 | ISLR2 | CACNB3 |
| EPHB1 | CACNA1C | VPS72 | SLC35F3 | TMEM186 | VSTM2L |
| NRG1 | TRIP4 | SEC24A | ARL13B | ZNF48 | TBC1D7 |
| AMPD3 | LMAN1 | DPF2 | WWC2 | DOK4 | CBLN4 |
| KLK6 | LIMS1 | VTI1A | NGB | ALYREF | NPHP4 |
| ATF7IP2 | PTPDC1 | TXNDC11 | SHISA9 | PGPEP1 | SLC17A6 |
| MRPS11 | LACC1 | DCTN4 | SRCAP | TNFRSF25 | XPOT |
| POU3F1 | NOTCH1 | SRSF9 | ADCYAP1 | MSANTD1 | BEND6 |
| IKBKB | FN1 | EMC6 | WDFY3 | KMT2B | NPY |
| KLF4 | LRRK1 | ARIH1 | CHMP4A | SPATC1L | CNST |
| CAMK1G | CNR1 | CDS2 | SEC22B | GRK3 | SMPD1 |
| KIF9 | FABP6 | LTBP3 | FKBP1B | ZNF184 | ACSL4 |
| SLC14A1 | LYST | CCAR2 | BBS7 | TMEM126A | RUBCN |
| ZNF736 | TAB3 | PDXDC1 | CLDND2 | FAM216A | ARHGAP39 |
| SEMA6A | LAMB1 | SLC7A1 | ADRB1 | MIR7-3HG | CPT1C |
| PRKAG1 | URI1 | LHFPL6 | ZEB2 | TASOR2 | ZMYND8 |
| SPRTN | GALE | DMAP1 | OSTF1 | KCNH1 | KIF22 |
| DRG1 | ABCG4 | CDKN1A | POLR3A | LNPK | RAB11FIP3 |
| RNF144A | DTYMK | SCN2A | MMS19 | IRF2 | FAM217B |
| CHN2 | SMC2 | LSM5 | CUL5 | ABCC10 | SMG1 |
| KLHDC8B | ITGB4 | ZBTB38 | PDS5B | UBE2D4 | ZNF146 |
| PKDCC | GALNT13 | PRSS3 | RBL2 | BDNF | IL1B |
| CA4 | JAGN1 | TAX1BP3 | RAB11B-AS1 | LTBR | SPARC |
| NINJ2 | ANK1 | CEBPZ | ADA2 | TTLL5 | CDC37L1 |
| GPR22 | TAC3 | B3GALT1 | NKAP | ZNF865 | PRKAB1 |
| LATS2 | PLAC9 | ZNF746 | KIF1B | BACH1 | MOK |
| MRPL58 | ERO1B | CDK5RAP2 | RGS1 | HOOK1 | PIGQ |
| ARRDC3 | CEP57 | LMBR1L | FBXL5 | ARHGAP29 | POU2F1 |
| YIPF2 | RBP7 | MED13L | MMD | CRB1 | TSTD2 |
| MTREX | VSIR | SPATA33 | SGMS1 | GJC2 | ABCG4 |
| CBLN2 | S100PBP | SHC2 | PPP1R3C | CYP26B1 | ZNF570 |
| RPRML | NSMCE4A | FBXO34 | EIF2AK2 | ZNF514 | CNBD2 |
| LRIG2 | TMEM98 | HMGN2 | GCN1 | RBMS3 | PITPNM1 |
| KCTD16 | TCFL5 | SNX13 | CRIP1 | RPP14 | ETV5 |
| SH3RF1 | MOB3A | LMBR1 | CDKN1B | FAIM | ULK1 |
| SCN1A | UBE2T | RIF1 | C3orf70 | MB21D2 | NKX6-2 |
| ZNF830 | CYP7B1 | CDC16 | ARFGEF2 | C4orf33 | TBC1D9 |
| SLC16A1 | AAMDC | NMB | SUOX | LINC01007 | SLC45A4 |
| MYH7B | RP2 | C19orf12 | ADAM10 | GCC1 | TARS1 |
| MGP | BRCC3 | RAB14 | CRLS1 | THAP1 | TMEM158 |

| | | | | | |
|---|---|---|---|---|---|
| VIPR2 | IFT172 | DDRGK1 | LARGE1 | BHLHE22 | GNS |
| SRPK1 | ANKRD44 | WASHC3 | MAFG | KCNS2 | WHAMM |
| INAFM1 | TRERF1 | PIP5K1C | CADM2 | FZD6 | EPS15L1 |
| NCEH1 | RSPH1 | RAB11FIP4 | IFRD2 | TBC1D2 | ADARB1 |
| NF2 | GNAL | RBM22 | MTLN | TSPAN15 | CLCN6 |
| SULF1 | NCBP1 | HSDL1 | SVIP | ZNF518A | RCN1 |
| EML1 | HMGXB4 | RTKN | FAM162A | HPS5 | TAFA5 |
| PIM2 | SCRN2 | HAPLN2 | NTRK3 | NTM-AS1 | TAC1 |
| ZDHHC12 | TNFRSF25 | SS18L2 | GRIK5 | STYK1 | GON4L |
| VIP | NUP210 | VAC14 | PCM1 | CEP83 | VIRMA |
| POLK | NME7 | CREBZF | RAB22A | GPN3 | ZNF121 |
| ZC3H6 | TTI1 | CD47 | FADS1 | LRRC28 | MSL2 |
| LRP12 | CCRL2 | PDZRN3 | RTF1 | ASB7 | HTR5A-AS1 |
| DPP10-AS1 | RASGEF1C | SH3GL3 | IPCEF1 | FN3K | SNRPA1 |
| DBT | MPP3 | CBR1 | CDK9 | ZNF236 | DAAM2 |
| BRIX1 | MHENCR | PHB2 | TMEM201 | CNFN | STXBP5L |
| LINC02607 | SFXN4 | BOD1 | CD84 | KLK6 | ZNF445 |
| DHRS1 | NUP50-DT | LGMN | NRBF2 | ZNF415 | GALNT7 |
| PLTP | NBAS | TAF15 | C2CD5-AS1 | ZNF337 | ZBTB5 |
| SMURF2 | HDHD5 | LAPTM5 | RCBTB2 | BAIAP2L2 | PATZ1 |
| TUG1 | LPCAT3 | RNF123 | SNUPN | DACH2 | SLC16A2 |
| PDCD4 | MFSD11 | GRK2 | FLOT1 | STEEP1 | GRAMD1C |
| HTR1A | PAK4 | MICOS10 | RAB11A | PPOX | LRRC73 |
| ATAD2 | PNKP | FBXO44 | CRIPT | BBIP1 | CEP43 |
| HIP1 | AP3M1 | ADCY1 | CFL2 | LYRM1 | KLHL42 |
| SCAPER | DPY19L2 | EDF1 | CACNG3 | HS2ST1 | CCNB1IP1 |
| FBLN7 | SPRYD4 | ALDH9A1 | EMC7 | TTC21B | CD99 |
| SYT17 | IFT74 | SDF4 | PTDSS1 | GDA | PCYT1A |
| SNUPN | MGST2 | LUC7L | ATRAID | C16orf89 | CEP95 |
| LNPK | UTP15 | MARK2 | TULP4 | YES1 | TMOD1 |
| NEUROD1 | TNFRSF10D | MED4 | AAK1 | WFDC2 | PEX5 |
| FAHD1 | PYDC1 | PACSIN2 | DLD | SRRM4 | CDH7 |
| QSOX2 | TMEM106C | ABAT | EGR4 | ARHGEF6 | SLC16A14 |
| CSPP1 | SLC19A1 | RBM42 | MGAT5 | RAPH1 | NUDT12 |
| CTDP1 | BCLAF3 | GOLGA7 | STIM1 | CACHD1 | ABTB2 |
| ZNHIT6 | LSM10 | INPP5E | SACS | ZMAT4 | HUNK |
| SUSD4 | CAT | CDC42EP4 | GRAMD2B | PUS1 | LINC01637 |
| ASCL1 | AARS2 | DGCR6L | LARS1 | SHISAL1 | PLPPR5 |
| RSPRY1 | PIGC | PELI1 | ITGB1BP1 | RIIAD1 | ATL2 |
| NTRK3 | TGFBR2 | TUSC3 | TEX264 | MASP1 | TENM2 |
| NECTIN3 | SOX6 | OLMALINC | DNAJA4 | RUFY1 | ATG2B |
| JAG1 | KCNJ12 | OLFM2 | VXN | RPP25L | NCAN |
| PAK6 | VLDLR | SLTM | BORCS7 | PIGU | DHDDS |

| | | | | | |
|---|---|---|---|---|---|
| C4orf33 | LRIF1 | EIF3I | NINJ1 | SDE2 | AGAP4 |
| IL13RA1 | EPHB3 | TMCO1 | PPM1K | INO80D-AS1 | KDM3A |
| RBMXL1 | CUL2 | NUP153 | FAM200B | HEG1 | DDX18 |
| NRDE2 | ZNF140 | BOLA3 | SYNJ1 | GUCY1A1 | CIAPIN1 |
| BMP2 | PAQR9 | POR | PDHB | NAF1 | GCC2 |
| PDE7B | STX8 | PPP2R5E | SUCLG1 | CDK7 | ISG20L2 |
| CLVS1 | DHX33 | NUP50 | WDR3 | GEMIN5 | C5orf22 |
| LINC01007 | ATP10B | MRPL57 | UTP25 | NSUN5P2 | SPIRE2 |
| CD93 | ADAMTS2 | SIRPA | USP37 | GRIN3A | WDR33 |
| HERPUD2 | MAMLD1 | MYCT1 | SLC25A27 | MIGA2 | MET |
| TMEM120B | CD302 | ATRN | VPS13A | HSPA14 | TNRC6C |
| ST6GALNAC3 | PTCHD1 | TBP | TTLL11 | FAM161B | SIDT2 |
| ANGPTL2 | ZNF767P | CD44 | ZFR2 | CLBA1 | RUSF1 |
| RCOR1 | PTPN13 | SLC30A7 | ARMC6 | TMEM121 | TPST1 |
| LARP1B | JDP2 | ANKRD35 | OGFR | NAGLU | SNAPC5 |
| ST3GAL6 | ADGRF5 | SEC16B | ANKRD40 | ITGA3 | NANS |
| NSUN4 | KLHL28 | RALGPS2 | NFKBIZ | YJU2 | CORO6 |
| LHFPL2 | ECHDC2 | RDH14 | MTCH2 | CC2D1A | SRGAP3 |
| IL17RB | RPGR | HNMT | AARSD1 | TBC1D22A | SNRPB2 |
| LIMK2 | UBIAD1 | ATG16L1 | TMEM80 | PLLP | SIAH1 |
| KLHDC9 | TMEM62 | CCDC127 | CLCC1 | LRRC7 | TJP2 |
| ARHGEF6 | COG6 | CEP78 | ERC2 | TMEM191A | PPP6R3 |
| KLHL25 | HAMP | HEBP1 | JADE1 | AUH | CCPG1 |
| ZXDB | CTXN3 | LRRK2 | TPMT | MCC | ENTPD4 |
| SLITRK5 | GALNT14 | WDFY2 | ZBTB22 | BTD | ZMYM4 |
| SHB | MIA2 | MEX3B | TTBK1 | SLF2 | KIF9 |
| TBL2 | MBD5 | PDP2 | PIERCE1 | NNMT | RASAL2 |
| THSD7A | PLXDC1 | SPAG5 | SLC39A9 | RPTOR | ILKAP |
| PCMTD1 | PLPPR1 | GYS1 | PARD6A | CCDC3 | MCRS1 |
| ECE2 | LHX2-AS1 | IL17RA | ZDHHC7 | JKAMP | CALCOCO1 |
| BHLHE22 | TSPAN15 | ARAF | KLHDC4 | DDOST | ZNF410 |
| NR2F2 | MAP2K6 | CITED1 | MYH14 | TNS1 | ZBTB21 |
| MIR1915HG | TTLL11 | RWDD1 | RABGGTB | POLR1C | PRKCG |
| GMPR2 | ARHGAP29 | PIP4P2 | TMEM176B | EXT2 | MPZL1 |
| POLR1H | HIF1AN | SRARP | ASPDH | ACAD8 | CMTM5 |
| DMKN | BCL9L | SNIP1 | SHANK3 | DHRS11 | MAF |
| BLMH | ZKSCAN8 | GPAT4 | SESN2 | ERMN | HTATSF1 |
| SMARCD2 | DBF4 | ACTN1 | MIR497HG | LINC01141 | PAIP2B |
| LINC01141 | CLEC16A | TMEM245 | VTI1B | PTPRU | ACKR1 |
| ENOX2 | USP30 | MSRB2 | CORT | LRRC8C-DT | C2orf72 |
| CCN2 | KLK7 | PRKAA2 | IDH1 | PPM1J | TMEM165 |
| PRMT8 | SLC25A19 | NPY1R | PHF2 | ACVR1C | DOLPP1 |
| CORO6 | GGTA1 | TBC1D2B | NETO2 | NPY1R | AP5M1 |

| | | | | | |
|---|---|---|---|---|---|
| CDC42EP2 | PRMT9 | ZNF668 | DPH3 | FASTKD3 | TMED5 |
| ATE1 | ABCB9 | SERPINB8 | MRPL47 | CDH9 | LINC01003 |
| CDK20 | KIAA0586 | PEX10 | RIMS1 | CCNO | IL1RN |
| NUFIP1 | LSM11 | LRRC41 | CHN2 | EXOC2 | ADGRB3 |
| CXCL2 | ZBTB24 | SEMA6C | TMEM168 | FOXP4 | SCYL3 |
| PIK3C3 | AGBL4 | CNOT10 | DMXL2 | RAMACL | NKX3-1 |
| SPPL2A | SH2B2 | MELTF-AS1 | IRGQ | SEMA3E | PPFIBP2 |
| BRMS1 | ZNF628 | C5orf22 | PCIF1 | MBLAC1 | P3H4 |
| THBS1 | DACT1 | MIOS | GABBR2 | TMEM67 | ZNF512 |
| ZDHHC6 | HSD17B6 | THAP1 | UBE2A | NDRG1 | OTUD3 |
| EPHA10 | ELP5 | DYNC2H1 | PVR | LINC02977 | RNF44 |
| RBM12B | C3orf33 | RPUSD4 | PDK4 | VTI1A | EPG5 |
| JAK3 | ESCO1 | SUPT20H | RNF114 | GXYLT1 | SLC19A2 |
| BAIAP2L2 | CFAP44 | TRPM7 | COBL | TMTC3 | PIGZ |
| KCNA3 | PSMG3-AS1 | NUDT7 | KCTD17 | PROSER1 | GSR |
| TNFAIP3 | ZBTB25 | KANSL1-AS1 | GABRG2 | MAGEL2 | EMX1 |
| PROX1 | GRK5 | DUS3L | ALCAM | AP4E1 | MRPL17 |
| CXCR4 | HIF3A | SAMM50 | GRIA4 | ALOX12-AS1 | DAGLB |
| AGRN | RAB13 | PPP1R9B | PANK2 | RBBP8 | ZNF655 |
| TMEM19 | ELF1 | BDP1 | GPC4 | AMH | RANBP3 |
| BICRAL | FRS3 | FCHO2 | FBRS | PDE4C | MTMR9 |
| AJM1 | NAGLU | ARL4A | CCT3 | PRRG3 | GNPDA1 |
| TPCN1 | CEP20 | SUPV3L1 | CDKN2AIPNL | PTGFRN | TBP |
| ZCCHC24 | GMEB2 | MGA | MAD1L1 | FAM110C | FSD1L |
| NUDCD2 | OLFML3 | ULK2 | HYCC1 | DNMT3A | RRM1 |
| VPS11 | CCDC85C | NKAIN4 | IFIT5 | SLC5A6 | SCNM1 |
| COPG2 | SEMA4F | PMF1 | NOP2 | DLX1 | HSPA2 |
| CDKL1 | BANP | MAD1L1 | KDM4B | SDHAP1 | LAMA4 |
| PRKD2 | ZNF362 | TNFSF12 | POFUT2 | CEP44 | MYPOP |
| RBM11 | LYPD1 | RGS19 | AMER2 | PPWD1 | GPX3 |
| WNK3 | EPHX4 | FLOT2 | MEIS3 | MDC1 | ADGRB3-DT |
| MAPK7 | IL1RN | NME3 | BBS5 | AASS | IQCK |
| TRAPPC12 | ALDH1A3 | SETD5 | WDR55 | BMP1 | SLAIN2 |
| DHRS3 | BBS9 | HSPA14 | VEGFA | HMBOX1 | ANGPT2 |
| PRDX4 | ZNF641 | LTA4H | TRAPPC9 | TIGD5 | DCAF15 |
| GTF2H3 | FBXW9 | TMCO3 | SLC9A5 | RCL1 | NHSL2 |
| ATF5 | B3GLCT | ARHGEF40 | TEKT4P2 | MCU | COL7A1 |
| PKNOX2 | BIRC3 | STRADA | G6PC3 | RRP8 | RIOK1 |
| MRPS14 | CHST8 | SOX8 | TRAPPC1 | PRR5L | DCAF12 |
| RAB37 | PLCB4 | TOMM6 | ANO10 | IGHMBP2 | ATP2B1-AS1 |
| LRRN1 | CYP2E1 | BPTF | GPR27 | RPAP3 | TEDC1 |
| LRRC37B | SLC13A5 | KIAA0319L | IGF2 | ABCB9 | TIAL1 |
| ALG5 | DHX35 | PRCC | PLOD1 | XPO4 | RASGEF1A |

| | | | | | |
|---|---|---|---|---|---|
| RASGRP1 | CNTNAP3 | HS1BP3 | MIER1 | MTUS2 | ME3 |
| RIOX1 | BDNF-AS | ZNF330 | ZNF672 | CLN3 | KCTD10 |
| CDAN1 | RAB3D | PIGH | IFT172 | NLE1 | AMFR |
| HRH2 | RARB | PALM | ANXA4 | DMKN | NRF1 |
| TOX2 | SLC5A11 | RELT | WDR54 | CSDC2 | RASSF7 |
| ZNF32 | RHOJ | PTBP2 | SCRN3 | FAM118A | CCS |
| TIMM21 | ANKRD36C | WBP1 | BOK | PRKX | USP2-AS1 |
| PITPNM2 | ELFN1 | SDC3 | IGSF11 | YPEL4 | DTWD1 |
| CDKN2C | CAD | UBE2J1 | PPT2 | C1orf174 | SLC24A4 |
| B3GAT2 | TTC7A | FAM221A | NT5DC1 | RBP7 | FGD1 |
| NNT-AS1 | ORC2 | KCTD2 | RNF217 | EYA3 | SLC25A29 |
| UQCC4 | TPM2 | USP24 | SP4 | HECTD3 | FAM229B |
| EML4 | ZBTB6 | EXOSC6 | SLC25A32 | TESK2 | LRRC57 |
| CUL9 | TMEM53 | RCOR3 | EXOSC3 | CRYZ | CRB2 |
| FAM184A | NDST3 | CPSF1 | PRMT3 | RPAP2 | INPPL1 |
| SOX11 | SLC8B1 | HMOX1 | CEP57 | LINC01750 | LIN7C |
| TMEM74 | ALG12 | TBC1D15 | TMEM117 | MTARC2 | FAM120A |
| ZNF528 | MDFIC | VPS13C | MFSD5 | CHML | MGST2 |
| NELFE | PAPLN | SCN2B | TRHDE | CMPK2 | LARP4 |
| TRPM2 | WDR81 | SP2-DT | GPR180 | KLHL29 | NSDHL |
| PTPRG | PLPP4 | CHTOP | TMEM260 | FOXN2 | CD14 |
| C1QL1 | ENTPD5 | N4BP2L1 | TUBGCP4 | EML6 | ANK1 |
| KBTBD8 | ADCY7 | ASRGL1 | THOC6 | TBC1D8 | RSF1 |
| GPX3 | MYT1 | MIGA1 | XYLT1 | RBM45 | FLYWCH2 |
| PIK3R3 | CTU2 | SEMA3B | ORAI3 | MREG | AKT1 |
| CARNMT1 | SNX29 | PDPK1 | ABCA5 | NPHP3 | TADA3 |
| COL4A2 | RGS8 | RAB31 | ENGASE | HPS3 | CREB5 |
| TMCC3 | EFL1 | RAB2A | TIMM29 | OTULIN | GPN1 |
| RNPEP | SNX7 | MAGI2 | ADM5 | CDH12 | SETDB2 |
| RHPN1 | OXNAD1 | ANGPTL4 | EBF4 | RNF180 | ZSCAN32 |
| DNAJC25 | KRI1 | HMGA1 | KCNG1 | ZNF354A | FMNL1-AS1 |
| HGF | CYB561D2 | SLIRP | LIMK2 | RASGEF1C | LIPA |
| RBM24 | CCDC80 | RBM25 | MRPS33 | CDKAL1 | NKAIN4 |
| NWD2 | FIBIN | SF3B5 | CREBZF | SHPRH | CYTH3 |
| STYK1 | GOLGA5 | SELENOM | PIP4P1 | ZNF398 | NUP58 |
| P3H1 | PMAIP1 | HLA-DRB5 | TMEM33 | KRBA1 | IPO9 |
| ACOT2 | F8 | YAP1 | ACAA2 | UBXN8 | HTR2A |
| TRMT9B | THEMIS2 | C2CD2L | ZNF335 | FBXL6 | SNHG17 |
| KIAA1755 | CPT2 | THUMPD1 | BZW2 | FAM27C | TMBIM4 |
| CHFR | DPH1 | UNC80 | MAST1 | NIPSNAP3A | RTKN |
| MRPL48 | PRCD | CDC40 | NREP | LHX2-AS1 | CAPN1 |
| TK2 | KNSTRN | SLC25A1 | FAM181B | FAM78A | U2AF1L4 |
| ZNF320 | NR2C2AP | ZDHHC9 | N4BP2L1 | CELF2-DT | ABHD16A |

| | | | | | |
|---|---|---|---|---|---|
| ANKRD65 | RNF215 | SEC61A2 | SOX9 | SFTA1P | SPTSSB |
| CELSR3 | PLP2 | ASPDH | GNAQ | LRMDA | SIRT5 |
| AMT | IL17RC | BPGM | PCP4 | GPR26 | WBP1L |
| BCL3 | SHQ1 | ZFHX4 | SDC2 | PAOX | PCED1A |
| CTNNA3 | COMTD1 | PEX5 | TRIM44 | ALKBH3 | UTP14A |
| DSCAML1 | HBQ1 | DDX56 | SCARB1 | SAC3D1 | PEX11B |
| PRPF3 | | WLS | MPHOSPH8 | C2CD3 | EVI5L |
| DDX23 | | FOXK1 | BLVRB | INTS13 | RRN3 |
| GLS2 | | JMJD8 | RAB33A | TDG | DDX42 |
| ATG4D | | OXSR1 | MAZ | POLR3B | LARP4B |
| CLK3 | | ASMTL | TMEM222 | KCTD4 | CAPN3 |
| TRIM47 | | PPP6R1 | HNRNPF | AKAP5 | PDCD2 |
| DNAJC11 | | ECI1 | SESN1 | MAP3K9-DT | NBPF19 |
| HSPB6 | | TARS3 | MYRF | VRK1 | HSCB |
| CACHD1 | | CDC27 | ANKIB1 | ARPIN | MYNN |
| CEP126 | | ALDH5A1 | CEP170B | LRRK1 | BACH2 |
| HEPN1 | | WTAP | CDKL5 | WDR90 | RANBP10 |
| MTFP1 | | GMFB | RPS27L | GNG13 | YPEL2 |
| EPHA6 | | POLR2E | ARHGEF2 | UNKL | SPIN3 |
| ADPRS | | TMED7 | ACAA1 | SLC5A11 | TTC33 |
| RASSF5 | | CNKSR2 | COPS4 | HEATR3 | USP47 |
| CIART | | GLRX5 | TXNDC17 | TMEM100 | DPYSL3 |
| MFSD13A | | NOL9 | CNTN2 | SMG8 | DOCK4 |
| NEURL4 | | NRAS | SMPD4 | DCC | DNAJC12 |
| MAU2 | | C1D | PRSS3 | PIGN | RHOG |
| SLC35C1 | | LMBRD2 | SIK2 | ZNF780B | CERS5 |
| HEBP2 | | ACLY | YAF2 | MMP24 | ERBB4 |
| SDSL | | SFXN5 | ACTR6 | HAR1A | FRS2 |
| FAM131B-AS1 | | MRPL47 | EXD2 | SLC7A4 | RSAD1 |
| FAM27C | | TUBGCP2 | HSD17B14 | CPT1B | FBXW4 |
| AATF | | FAM199X | MED25 | ANKRD13D | LINC03013 |
| UNC50 | | ABCD3 | HP1BP3 | AQR | THEMIS |
| ZBTB47-AS1 | | GCA | MTSS2 | CPEB1 | AFF2 |
| ZNF708 | | G3BP1 | SLC18B1 | RRAGB | MTMR4 |
| MTUS2 | | KCNJ10 | KIF2A | HAUS2 | FAM229A |
| TUBG1 | | UBTF | RNF2 | GSG1L | KCNA3 |
| RABL2B | | NFIB | BBX | NETO1 | ZDBF2 |
| RPS6KB2 | | FUBP3 | BBOX1 | CKS1B | ZKSCAN5 |
| RPP38 | | GPSM2 | SLCO1A2 | MCM6 | PPP4R1 |
| MYO16 | | RBM12 | WBP4 | C8orf76 | GSKIP |
| LRFN1 | | KDM5D | LRRC4 | TSPAN4 | EXT1 |
| NHSL2 | | ZYG11B | MOB2 | PAGR1 | PRKRIP1 |
| MTRFR | | ZNF24 | SLCO1C1 | PLXDC1 | DPF2 |

| | | | | |
|---|---|---|---|---|
| TMEM175 | MPG | KATNB1 | SAMHD1 | MIR99AHG |
| ARL6 | DLL1 | TMIGD3 | LINC02609 | VHL |
| VPS13A | MAPKAPK2 | SELENOI | INO80D | TRAPPC14 |
| NAB1 | ESRRA | TMEFF2 | MOB3C | CDS1 |
| F12 | PGLS | NPRL2 | ELK4 | APBA1 |
| LIPE-AS1 | PPP1R1B | PXK | PACC1 | LYRM7 |
| ACVR1C | THAP5 | MRPL3 | TAF5L | PJA1 |
| CCNO | BCL7C | RAB28 | SPRTN | LEPROTL1 |
| IMMP2L | CHDH | AGA | GCFC2 | EHBP1L1 |
| NRF1 | FOXP4 | SMIM15 | SHQ1 | LIN7A |
| FAM120C | OTUD1 | USP42 | SDHAP4 | RIT2 |
| ARL16 | WDR75 | CCAR1 | RCC1L | TRIM36 |
| ENOX1 | RBMS3 | RIC3 | PTCD1 | PBDC1 |
| VWC2 | ATXN7 | DCUN1D5 | LRRCC1 | GNA13 |
| NUP155 | SLC33A1 | TMTC2 | DNMBP | EPB41 |
| GUSB | GMNN | RBM26 | NPAT | RBMS1 |
| FOLH1 | TUBE1 | PPP2R3C | GPD1 | RFC2 |
| PTCH2 | FLNC | COA8 | ARHGAP5-AS1 | PDE7A |
| GOLPH3L | VPS37A | SEMA7A | FCF1 | MRM3 |
| RIOX2 | ZNF33A | CPLX3 | DLL4 | TP53RK |
| FILIP1L | WASHC2C | SEC14L5 | ATOSA | MORC2 |
| PGM2 | PARG | PHLPP2 | CSPG4 | DRP2 |
| TCERG1L | LDB3 | DHX38 | LRRC37B | HAUS7 |
| EML3 | DNAJC14 | ZNF91 | RARA | FAM174A |
| SEPTIN1 | HTR2A | HSPB6 | KRT17 | KMT2E-AS1 |
| FAAP20 | EMC9 | ALKBH6 | SMURF2 | POLR3E |
| ZNF383 | SALL1 | ANOS1 | RHBDF2 | KAT8 |
| MARK1 | IMPACT | MRPS27 | RFX2 | TBC1D12 |
| EPHA2 | VSTM2B | SDCCAG8 | ZNF626 | CEP290 |
| TRABD2A | ZNF566 | MCCC2 | ZNF567 | MRPL48 |
| ADRA2C | PRR14L | C2orf74 | RBBP9 | KLHL18 |
| VSTM4 | BCOR | CEP170 | CDK5RAP1 | ANKRD36 |
| PRELP | MED14 | TLK1 | RGS19 | LYPD6 |
| BLOC1S4 | KMT2C | BDH1 | MRPL39 | ACOX3 |
| ZNF518A | SLC25A26 | PHYKPL | MIR222HG | PCDHB14 |
| PEX2 | CCDC86 | CSGALNACT1 | TRIAP1 | TRMT12 |
| TMEM63A | CMTM5 | HEY1 | ENTREP3 | DHRS13 |
| TPGS1 | TXLNG | NTNG2 | IGF2R | DZANK1 |
| FAM181B | HMGB2 | DHTKD1 | FBXO42 | TENM1 |
| VASN | GNG5 | IFTAP | MDFIC | RBM15 |
| APBB1IP | TTC1 | SIAE | UVRAG | H2BC21 |
| PRRG1 | MTF1 | SP1 | KCTD21 | MRPL30 |
| PREB | TMEM250 | UNC119B | TSPAN9 | JADE2 |

| | | | | |
|---|---|---|---|---|
| ELOVL7 | PSRC1 | MPPE1 | TAFA2 | JAG1 |
| MID1 | ASF1A | SMAD2 | FAAP100 | DNAJC16 |
| JAK2 | AGPAT5 | CXADR | SALL3 | CCDC28A |
| ASIC2 | SLC25A32 | CHRDL1 | CACNA1B | ZNHIT2 |
| MCHR1 | ARAP1 | GET1 | EIF2B3 | KCNH5 |
| RPRM | NBR1 | EIF4G1 | USPL1 | KIAA0753 |
| TRIM26 | SLC27A1 | GPR3 | IL1RL1 | PRELID3A |
| UTP25 | SLC25A17 | PRMT6 | GNL2 | USP36 |
| ZNF354A | NR4A3 | ETV3 | TFB2M | COMMD10 |
| H4C8 | RNF208 | TAMM41 | CAPN7 | C1orf159 |
| ACAD9 | SEMA4D | COLQ | UMPS | PLGLB1 |
| NGB | WIPF3 | WRN | SV2C | SUMF1 |
| ALMS1 | TWF1 | PSMD5 | NEURL1B | PRIMPOL |
| DTNB | DCAKD | NCAPD3 | THSD7A | CASP2 |
| LNX2 | JAKMIP1 | PTPRR | MLXIPL | CWF19L1 |
| CHADL | HSPBP1 | ACSS3 | WDR91 | ADCY6 |
| ST7 | SPACA9 | DCUN1D2 | VPS13B | PRANCR |
| WDR54 | SYMPK | ZNF23 | RGP1 | HBQ1 |
| ZNF582 | NCBP2 | ZNF333 | CARNMT1 | CA4 |
| SRSF12 | RTRAF | NEFH | PTPN3 | B9D2 |
| TLCD3A | BAG1 | CMPK1 | SIPA1 | BACE2 |
| ACOX3 | RAB5IF | DHX15 | DYNC2H1 | SLC37A1 |
| KNOP1 | ROBO1 | DNAJC8 | ALG9 | CCDC157 |
| ZNF653 | DENND4A | NRBP2 | PTHLH | HADH |
| GEMIN5 | AHNAK | SLC9A6 | ANKRD52 | SLIT3 |
| C9orf40 | MTA2 | MFSD3 | MEG9 | ABCB8 |
| SLC7A8 | VPS41 | CSRNP2 | PPIP5K1 | UBE2W |
| GON7 | PIP5K1A | GABRA2 | THAP10 | RDX |
| ASCC2 | SERPINB1 | CCSER2 | EEF2K | MPHOSPH6 |
| YIPF1 | TMEM33 | PAQR4 | WSCD1 | APBB2 |
| MLXIPL | GLG1 | GRN | MAPK7 | PPIG |
| AMDHD2 | ARPC5L | ZNF503 | ZNF708 | MORN1 |
| SDR16C5 | PBRM1 | AKR1B1 | NUDT19 | SYNC |
| RWDD2A | MSL2 | TCEAL9 | CEP250 | EVA1B |
| TRIT1 | BDH2 | MRPL37 | PWP2 | C3orf62 |
| DLX6-AS1 | ZNF292 | RHOT2 | EOLA1 | PGM2 |
| SLC16A14 | GPR107 | SERPING1 | CCZ1B | PCDHB4 |
| BNIP1 | BRI3BP | CLIP2 | SNAPC3 | EXD3 |
| TYW1B | POLG | FAAH | IL1A | NEBL-AS1 |
| PYCR3 | WDR59 | ABLIM2 | GLT8D1 | P3H3 |
| MPPED2 | MPHOSPH6 | ST7 | HBA2 | KCNK3 |
| BBS10 | UBE2A | VAMP1 | HBB | ITGA9 |
| KCNN3 | KLC2 | B4GALT2 | | LMLN |

| | | | |
|---|---|---|---|
| CA7 | UBE2G1 | GTF3A | SOWAHB |
| VPS13B-DT | AKTIP | SLC44A3 | TCOF1 |
| CACNA1I | RHOC | RBBP5 | NR2E1 |
| UPK3BL2 | SACM1L | LPCAT1 | DHX32 |
| FAM76B | TRIO | H2AC6 | TCP11L1 |
| TCOF1 | IFT25 | SYNGAP1 | TMEM132C |
| PTPRR | CIAO3 | OSTM1 | POLE |
| PDCL | CCT2 | MIR124-1HG | CABYR |
| TBC1D8 | SLC44A2 | TCF7L2 | TASP1 |
| ANO6 | GET1 | EPS8 | PRKAR2A |
| INPP5B | MCUR1 | PAF1 | CHD8 |
| DCTN6-DT | GPT2 | GRIK1 | TINF2 |
| APOA1 | FARP1 | RTL10 | STK35 |
| DACH1 | WDR45B | ID3 | EPHB1 |
| ERI2 | DOCK9 | UBXN4 | SLIT2 |
| CYP2U1-AS1 | AP1S2 | CD68 | USP53 |
| TMEM233 | CKS1B | SCYL2 | PLPP5 |
| ALOX12-AS1 | CIAPIN1 | L1CAM | CNNM2 |
| ISLR | NF1 | ADAM15 | PKNOX2 |
| FAR2 | EMC3 | DCAF16 | UBE3B |
| TTC9 | FOXRED2 | MRPL11 | PPTC7 |
| ANKRD26 | UBXN11 | HSPB2 | INAFM2 |
| ZNF615 | PHF21A | USP46 | SMPD3 |
| ATP2B1-AS1 | AMOTL1 | NNT | NLGN4X |
| LRSAM1 | FGD6 | CCDC6 | MAP3K13 |
| AKAP5 | KATNAL1 | BGN | SF3B3 |
| RAC3 | LMO7 | SLC45A1 | BMP7 |
| ZNG1A | TUBGCP3 | PRKD3 | NEK1 |
| RRP1 | TMEM121B | TMEM108 | SLC38A3 |
| AGL | RHBDD3 | LINC01511 | ACY1 |
| FPGS | ARID1A | SLC22A23 | CCDC71L |
| RIOK2 | PRXL2B | POMZP3 | TAF3 |
| PIP5K1B | FGF2 | CDC37L1-DT | LINC00667 |
| FBXL12 | SLC25A37 | STRN3 | FCER1G |
| PRDM1 | KAT6B | SLC16A3 | MMAB |
| GNB1L | TEAD1 | XAB2 | RABL2A |
| CRHR1 | BNIP2 | DPY30 | NIPAL2 |
| REEP6 | RTCA | GLDN | IGFBP5 |
| NOTCH2 | FYTTD1 | JAKMIP1 | PRPSAP2 |
| CTSC | TYROBP | TUBA1C | TRIM47 |
| LRRC75B | NT5DC2 | LENG8 | PPP1R3E |
| PYGO2 | ZDHHC23 | CDC42EP2 | PPM1F |
| ABCC12 | PLXNA4 | LIF | TAGLN2 |

| | | | |
|---|---|---|---|
| SPG11 | LINC01003 | ANKRD65 | HMGB3 |
| GJB1 | TRAPPC9 | TOMM70 | PLCXD2 |
| BGN | SLC25A16 | SEC31A | SLC35B3 |
| TGFB2 | ANO3 | CCNG1 | RFX3 |
| BAK1 | CWF19L2 | NOPCHAP1 | SLC25A42 |
| IL4R | POU6F1 | DNAJA2 | UCKL1 |
| ZFP82 | VRK1 | RICTOR | MED12 |
| ZNF470 | PARD6A | RAB20 | TCEA1 |
| POLR1E | ZSWIM9 | PNKP | RPUSD4 |
| EXOC6 | CLDND2 | WRAP73 | SLC6A15 |
| CHRM4 | CD55 | MUL1 | WDR20 |
| ZNF384 | PRPF38A | PIGV | GMPR2 |
| MCF2L2 | C6orf136 | GOLPH3L | ZNF268 |
| SPIRE2 | MAGI2-AS3 | DYSF | TYMP |
| MTX1 | GPR137 | SPR | THBS2 |
| LNCBRM | HEPH | CNNM4 | C1QC |
| DCBLD1 | PDE8A | TSGA10 | FN1 |
| DHX58 | HNRNPH2 | GULP1 | NET1 |
| EVA1B | MAEA | ZNF148 | UGGT1 |
| PCBP3 | CCDC124 | P2RY12 | RIPK2 |
| SCMH1 | SOX2 | HPF1 | UQCC4 |
| UCN | GAD2 | TENM3 | TRAPPC2B |
| PRKCD | ITGB1 | LINC02217 | PPP1R14B |
| SPRY1 | MLEC | FARS2 | AGO3 |
| ERCC8 | GSKIP | POLR1H | TMEM198 |
| PENK | UBE2G2 | ZNF292 | SMIM30 |
| NAA40 | ZBTB47 | PPIL4 | CSMD1 |
| RIN1 | IFFO1 | AHR | NAT10 |
| SCARF1 | SYT2 | HUS1 | FNTB |
| PTPRT-DT | MFSD1 | VPS50 | CRYBB2P1 |
| ARHGEF26 | TRIB3 | TMEM74 | SYN3 |
| TENM1 | C1orf174 | WASHC1 | RAB9B |
| CMYA5 | BOLA1 | TRPM3 | LITAF |
| ANAPC7 | POU2F1 | TMEM250 | ZC2HC1A |
| FBXL6 | GNG4 | PDZD7 | LRRC8C |
| TAF5 | GREB1 | ANKRD42 | CEBPZ |
| RTTN | SLC35F6 | LAYN | NPAS4 |
| B9D1 | THADA | MAGOHB | CCNC |
| TUBA1C | USP4 | SLC16A7 | RALGAPB |
| IGFBP4 | SLC15A2 | SRGAP1 | ELOVL1 |
| LTBP1 | ZFP62 | SLC8B1 | AIF1L |
| ZNF143 | LY86-AS1 | LMO7 | CELSR3 |
| LPCAT2 | MRS2 | KNOP1 | SLX9 |

| | | | |
|---|---|---|---|
| HGH1 | UFL1 | MATCAP1 | SYT17 |
| MYBPHL | PEX3 | CENPN | TPT1-AS1 |
| LINC00595 | ZNF277 | MYO19 | SMAD7 |
| PSKH1 | STK3 | METTL2A | EDEM2 |
| CCDC61 | LURAP1L | SMARCD2 | THOC5 |
| SOX18 | ZNG1C | TCF3 | GAL3ST1 |
| PARP3 | ANKS6 | JAK3 | PLCH2 |
| CCDC137 | DNAJC1 | RTEL1 | PCCB |
| ZNF419 | TIMM23B | CACNG2 | CUX2 |
| NBPF11 | MCU | CCDC22 | SLCO3A1 |
| C2orf42 | R3HCC1L | PLK3 | VPS9D1 |
| CYB5R2 | BET1L | ADAM23 | ATG4D |
| FEN1 | SP1 | ICE1 | SPTLC1 |
| LRRCC1 | HCFC2 | FNDC9 | SFN |
| CAPS2 | CHAMP1 | THOP1 | VAV3 |
| PRR5 | DHRS4-AS1 | EXOSC5 | PCAT6 |
| E2F3 | NUBPL | CMTM6 | CBLB |
| ICAM2 | DCAF4 | CXCL12 | EMB |
| ZNF432 | CRIP1 | NRXN1 | PROB1 |
| CNPPD1 | IFT140 | KIFAP3 | CLDN15 |
| GLIPR2 | LDHD | DDAH2 | NOM1 |
| RNF113A | C16orf46 | ZBTB7A | C10orf143 |
| NLN | LINGO3 | MSR1 | SLC6A13 |
| LRRN2 | ZNF554 | TMEM43 | CA12 |
| SDHAP4 | ZNF585A | FAM204A | HAPLN3 |
| INTS12 | ZNF540 | METTL5 | PLD2 |
| AIMP2 | B9D2 | OTUD4 | MIS18A |
| XAF1 | SMIM17 | TXNIP | SLC14A1 |
| REL | CTNNBL1 | ARIH2 | TENT4A |
| WNT7A | THOC5 | SPG7 | GNPDA2 |
| LINC01511 | TRAPPC2 | ATPAF1 | TNFAIP3 |
| CEP295 | PDIA6 | NUDT14 | NIBAN2 |
| TMEM186 | TUFT1 | FAM50A | SLC15A3 |
| SMAD9 | STOML1 | SEL1L | DDI2 |
| RAB11B-AS1 | ERBB3 | SREBF1 | DDX20 |
| TRPM4 | TARS1 | MKKS | FAM78B |
| STPG1 | VSTM2L | ICMT | LINC01106 |
| PAOX | WFS1 | PSMA5 | NUP54 |
| ERBB2 | MED13 | TMED8 | ARHGAP10 |
| IFI44L | CTNNAL1 | KDM5C | SLC30A5 |
| DRD1 | TMEM256 | EFR3B | TRMT11 |
| HSD17B8 | SUCO | EIF4E2 | LINC03011 |
| EHD2 | CHD3 | ACKR3 | TUT7 |

| | | | |
|---|---|---|---|
| CSTF1 | ATG4B | HAVCR2 | PTBP3 |
| NPEPL1 | KCNB1 | MAN1A1 | LINC01503 |
| ATP6V0A2 | CNOT8 | ASH2L | MAP3K8 |
| RGS7BP | MAPK1IP1L | SBNO1 | UEVLD |
| FOXQ1 | CORO1A | PXMP2 | LYSET |
| TENT5A | PHLPP1 | TTC9 | ZC3H4 |
| TLR4 | PHGDH | SNW1 | RGS14 |
| DOLK | STX5 | SPPL2A | RNF216 |
| TRADD | PARP2 | RAMP2 | IFT20 |
| TINCR | SRXN1 | EFTUD2 | ARSA |
| ZSCAN1 | POP4 | NMT1 | RABL2B |
| CT75 | NUP54 | USP32 | GPR108 |
| ZNF585B | RNPC3 | RAB4A | OPRD1 |
| ZNF530 | BRINP2 | CAMTA2 | PTPN12 |
| COMMD8 | PRADC1 | UHMK1 | ABHD5 |
| ABCC10 | CYP4V2 | LYRM4 | KATNAL1 |
| P3H3 | B4GALT7 | NR1D2 | HHIP |
| ADPRHL1 | FBXO30 | NLGN3 | CHI3L1 |
| GCHFR | TECPR1 | PRKACA | GLOD4 |
| ZNF548 | HGSNAT | DNAJB11 | TMEM11 |
| ZNF587 | ENTREP1 | PDE4D | TPRA1 |
| GLMP | SAV1 | RGL2 | CDK13 |
| PPFIBP2 | REXO1 | PHIP | ZNF189 |
| GEMIN7 | BACH1 | KCNS1 | GLS2 |
| SNHG11 | PDPN | ESS2 | DGKE |
| HYI | USP1 | CDK5R2 | HIRA |
| RAD17 | KYAT3 | SEC16A | GPSM2 |
| FAM151B-DT | GTF3C2 | SCAMP4 | VAMP8 |
| RERG | PPAT | ZDHHC21 | NUDT6 |
| KCNAB3 | SIRT5 | PPP2R1B | ATP10B |
| IL6 | LEMD2 | SLC35E2B | TRAF3IP2-AS1 |
| APLNR | IQCE | OSMR | NSUN5P1 |
| MARCHF4 | OSGIN2 | BAG4 | MTMR7 |
| DENND2A | SNAPC4 | FGD4 | DICER1-AS1 |
| CDC37L1-DT | ZNF37A | CASKIN2 | IL32 |
| NEBL-AS1 | TAF1C | VAPB | ZFHX3 |
| MMP19 | RAPGEFL1 | SDF2L1 | DDX52 |
| ORMDL2 | POLRMT | TFIP11 | HSD17B1-AS1 |
| C19orf81 | ZNF426 | RFPL1S | TGM2 |
| SH3RF3 | GPCPD1 | SLITRK2 | YBEY |
| CD24 | MCM3AP | BSDC1 | EXOSC7 |
| CASP9 | PRKX | PBX1 | NRG2 |
| TMEM69 | CASP3 | PUM2 | MRM2 |

| | | | |
|---|---|---|---|
| SOCS1 | ITGB8 | C8orf33 | GLIDR |
| MX2 | ZMAT1 | SGCB | CDHR1 |
| DNASE1L1 | ELAPOR1 | BOLA1 | OSBPL5 |
| SLC13A3 | BMPR1B | CTSS | CSNK1G1 |
| RFC4 | SMAD1 | ILDR2 | DHX8 |
| ENOSF1 | KCNIP1 | ADSS2 | MALT1 |
| FZD5 | ARMT1 | PGAP1 | MIER2 |
| DHX37 | POMK | SH3BP4 | BICRA |
| TDP1 | DCAF13 | RBM15B | GPRASP3 |
| GNA15 | LRMDA | CD200 | CDK12 |
| ZNF219 | ALDH18A1 | ARMC8 | SOS1 |
| GON4L | LRFN4 | ZFPM2 | SLC33A1 |
| KCNK4 | COQ10A | AOPEP | GAS8 |
| MRE11 | SH2B3 | USP20 | SAMD14 |
| DLL4 | MEG8 | CTBP2 | PICK1 |
| NLGN4Y | EHD4 | MS4A7 | PPIH |
| TSPAN9 | SOCS7 | TRPT1 | MDN1 |
| GPR19 | APCDD1 | CLPB | URGCP |
| TVP23A | LRCH2 | SUDS3 | MFSD14CP |
| MRPL45 | DGAT1 | PHAF1 | ZMYND19 |
| EYA3 | RIMS4 | ATP13A1 | MINDY3 |
| GRASLND | FNDC5 | MAPK12 | GLMP |
| TMEM17 | CC2D1B | ALG14 | CYP27A1 |
| FRAS1 | LINC01719 | SELE | ELOVL7 |
| TAP2 | HEATR5B | TFDP2 | DOCK5 |
| MAPK13 | SP100 | TRAPPC11 | NRG1 |
| PKIB | MON1A | MEN1 | ZNF496-DT |
| ZNF550 | ACTR8 | ADPRHL1 | ZNG1B |
| UTY | GPR27 | SERTAD3 | BLTP3A |
| ENTPD3-AS1 | HCLS1 | CHMP3 | LRRC61 |
| NOP14-AS1 | PPARGC1A | PSMC6 | KLHL9 |
| PDZD7 | MANBA | CNN3-DT | LIMA1 |
| RACGAP1 | TRAPPC11 | PRPF3 | WDR25 |
| LINC03048 | RUNDC3B | LAMC1 | FMN1 |
| GAL3ST1 | PPP1R9A-AS1 | LYST | ZNF75A |
| THBS3-AS1 | UBXN8 | ZNF692 | RFX1 |
| CDH12 | ADCY8 | EFEMP1 | KIF16B |
| COL23A1 | PTCH1 | EIF4E3 | PHF8 |
| TMEM100 | TDRD7 | ZBTB20 | TIMM8A |
| SAMD10 | GRIN3A | VWA5B2 | ZBTB33 |
| P2RX6 | AFAP1L2 | DGKG | FCGR1A |
| C1GALT1C1 | CCDC34 | LHFPL3 | OAF |
| BACE2 | TIGAR | GPR37 | NFRKB |

| | | | |
|---|---|---|---|
| RGS14 | DYRK4 | INSIG1-DT | ID1 |
| AK9 | KCNK10 | PIP5K1B | CRYZL1 |
| SBNO1-AS1 | DOK4 | DPH7 | LIMS1 |
| VIPAS39 | THAP11 | AKR1C3 | SESTD1 |
| ISYNA1 | GAS8 | COMMD3 | IPO8 |
| ZNF446 | TBKBP1 | WASHC2C | EPHX4 |
| FAM131C | AXIN2 | BCAT1 | ITSN2 |
| TARS2 | FN3K | TWF1 | GTF3C2 |
| HK2 | MPPE1 | PCCA | MYO1B |
| C1QTNF3 | PSPN | METTL17 | GTF2E2 |
| CPVL | TMED1 | RPUSD1 | TTC39B |
| ZNF117 | KPTN | SLC13A5 | CDC14B |
| SEC31B | SPEF1 | ROCK1 | TCTN3 |
| AQP11 | MYL9 | TIMM21 | LEMD3 |
| LINC02762 | L3MBTL1 | ADGRE5 | RPS6KA5 |
| RNF214 | MIR99AHG | MYADM | ATXN3 |
| PELI2 | VPS26C | ZNF329 | ZNF266 |
| RWDD2B | YBEY | NDUFAF5 | RFXANK |
| TFIP11-DT | CYTH4 | PEX26 | PAXBP1 |
| GNL3L | WNT7B | ZMAT5 | ASPHD2 |
| KIF26B | FTX | ASCC2 | ZBED1 |
| LINC00951 | CHRDL1 | OSBP2 | ZBTB48 |
| RNF216P1 | NKAP | DNASE1L1 | ABCB10 |
| FKBP11 | GABRA3 | KDM1A | TTC13 |
| DCX | ACO2 | POGZ | KDM1B |
| XIRP1 | DUSP2 | ANTXR1 | RWDD2A |
| C3orf62 | CCNT2 | BAIAP3 | SNX10-AS1 |
| TEK | NRXN3 | SUPT6H | TAF5 |
| ASAH2B | FANCL | JMJD6 | COL4A1 |
| CLEC7A | STIM2 | RIOK3 | METTL22 |
| PDYN | ITPR2 | MEGF8 | TINCR |
| CCDC120 | LMF2 | VPS39 | ZNF562 |
| FEZF2 | PEF1 | CBARP | ZNF583 |
| SEMA3E | TM9SF1 | APC2 | CHST7 |
| GUF1 | PLAUR | PCLO | RASSF8 |
| CBR4 | LPIN1 | TFCP2 | SLC66A1 |
| ANKRA2 | C2orf74 | SMARCC2 | CTBS |
| POLR3B | ZNF316 | LRRN3 | LINC01719 |
| TRMT1L | GOLGA7B | ENTREP1 | ANKRD44 |
| STK36 | PLEKHM2 | ZFYVE21 | PPP2R3A |
| SEC22A | CNDP2 | CD44 | LYSMD3 |
| HSPB3 | CACNB3 | PCYOX1 | AZGP1 |
| ZFY | TNPO2 | DUSP2 | MYO15B |

| | | | |
|---|---|---|---|
| FKBP14 | TMEM208 | DCN | COLEC12 |
| RAVER2 | AMZ2 | ADH5 | ZNF382 |
| IRS1 | CADM1 | TOB1 | PCDH11Y |
| TRMT44 | TMUB1 | PEPD | FBXL19 |
| ZEB1-AS1 | RC3H2 | SMIM12 | CCN2 |
| TALAM1 | DHCR7 | ILRUN | CNIH3 |
| KLHL35 | RUNDC1 | PPM1E | RBKS |
| SOX21 | DHX40 | STPG1 | PPIL3 |
| PIGBOS1 | ARHGEF18 | AMY2B | P3H2 |
| ZSCAN5A | CC2D1A | TRIM46 | TBXAS1 |
| TBC1D22A-DT | RSAD2 | PIGM | NSUN6 |
| HMGN5 | MERTK | ZBTB47-AS1 | FAS |
| MACROH2A2 | GRK4 | DUBR | BIRC3 |
| AGTRAP | AHR | C4orf50 | PTPRB |
| ZNF561-AS1 | SLC27A4 | NAAA | RBM19 |
| LKAAEAR1 | MTMR6 | MFSD8 | DHX37 |
| ANAPC10 | MAN2C1 | DHX29 | MIR193BHG |
| WWOX | ABHD17C | EFNA5 | C17orf114 |
| TMEM107 | LYRM1 | LINC01574 | BCL10 |
| TRIOBP | LTN1 | MRS2 | SLC35F6 |
| ART3 | LAS1L | PBX2 | DAP |
| ZC3HC1 | TERF2 | SLC12A9 | CDK20 |
| RELA-DT | ILF3 | CBLL1-AS1 | DALRD3 |
| NAALAD2 | HBA2 | PAXIP1-DT | CDH18 |
| ABCD2 | SMIM12 | DNAAF11 | ATG5 |
| SLC38A6 | TOB2 | LY6E-DT | MRPL50 |
| ST20 | SENP2 | TTF1 | SLC39A13 |
| SRR | PDE1A | REXO4 | TM9SF1 |
| CDC6 | LRRC4 | TPBGL | SPHKAP |
| CIDEA | ZNF706 | CHAMP1 | RARB |
| KCTD15 | WDR3 | SETD6 | DNAH1 |
| TRPC5 | ATG12 | ADAT1 | SNRNP48 |
| XAB2 | LINC02609 | P2RX5 | IER3-AS1 |
| CDK11A | DUSP12 | ZNF461 | ZNF3 |
| TMEM177 | CACNA2D3 | IGLON5 | SNX16 |
| FAM78A | EIF5A2 | ZNF432 | ITFG2 |
| NNMT | NUCB2 | FIZ1 | GRIP1 |
| PEX12 | DNAL1 | KAT14 | ARMH4 |
| COPZ2 | ITSN1 | GMEB2 | TRIM16 |
| TTC32 | TTC9B | PNPLA4 | ZNF8 |
| SUFU | SAP130 | NBAS | STT3B |
| B3GNT4 | DECR1 | TMEM185B | TP53TG1 |
| WNT3 | SLC25A28 | IFT80 | CPSF3 |

| | | | |
|---|---|---|---|
| CENPP | C11orf87 | KCNQ5 | SH3D19 |
| HMBS | LIN52 | ZNF614 | NT5E |
| NUDT4B | SNX27 | TNPO1 | ZFAND2A-DT |
| RNASEH1-DT | WNK2 | CALD1 | SNX8 |
| P2RY12 | OCRL | SURF2 | NAA16 |
| CDK6 | LPL | ITGB4 | ZNF569 |
| CTNS | KAT6A | ETFA | RYR1 |
| G0S2 | HCFC1 | FAM241A | ARHGEF1 |
| RUVBL1 | RPP21 | TMEM192 | PPP2R3B |
| RFXAP | SIKE1 | NNT-AS1 | LINC00963 |
| ALG1 | B4GALT3 | GPR158 | MANF |
| RASIP1 | ARL13B | SUPV3L1 | ZNF436-AS1 |
| ARMC6 | GTF3C4 | METTL3 | AZIN2 |
| PLA2G2C | AGAP6 | IFI35 | CHD1L |
| LTC4S | WASHC2A | TBKBP1 | EDEM1 |
| ST6GALNAC4 | TSFM | UNK | NEK4 |
| TMEM52B | PLXNC1 | ZNF532 | B4GALT7 |
| RAMP2 | MPI | NPEPL1 | PAK1IP1 |
| CDH7 | CLUH | YIPF6 | STRIP2 |
| ZNF430 | TSHZ1 | LCP2 | LYPLA1 |
| IL4I1 | ACSL4 | AK2 | GOLGA1 |
| MSC | MAP3K2 | PREB | DPAGT1 |
| PCSK5 | ARFGEF1 | HNMT | RERG |
| DNAH17 | NRBF2 | TIMMDC1 | SGF29 |
| ATXN7L2 | ENDOD1 | TBL1XR1 | ZNF768 |
| ITGA9 | USP8 | CAMK4 | KCNAB3 |
| TBXAS1 | GCC2 | TTC1 | MED9 |
| TRAF2 | PWWP3A | NUDCD2 | TUBG1 |
| MS4A7 | PCNT | DNAJB12 | RAB37 |
| MSRB3 | NUDT16L1 | MARCHF5 | RAB3D |
| LINC01089 | NUP42 | N4BP2L2 | PDCD2L |
| ZNF213-AS1 | PRPF40A | CHMP1A | ZNF181 |
| ZNF747 | WDR17 | APPBP2 | ARX |
| FMNL1-AS1 | RHOBTB2 | PRDX4 | TARS3 |
| ZNF865 | UHRF2 | LSM2 | COL26A1 |
| SSTR3 | ZBTB43 | DAB2IP | INSIG2 |
| | SMC3 | PRDM8 | SMARCE1 |
| | IGFBP5 | PTDSS2 | ATAD3A |
| | PPM1L | SYTL2 | MECR |
| | TCERG1 | GRIN2B | MKNK1 |
| | NAE1 | SLC15A2 | RNPEP |
| | ATF2 | JAM3 | D2HGDH |
| | EXOC6B | SGTB | BRF2 |

| | | |
|---|---|---|
| THBS2 | SRP19 | CBL |
| FAM174C | PSMG3 | RUBCNL |
| STAG2 | MRPS17 | SGPP1 |
| SUPT16H | SLC52A2 | TMED3 |
| APOLD1 | MARK4 | CHKB |
| EDNRB | SMIM45 | SERPINA3 |
| WDR13 | SAMD8 | NFIA |
| LAMTOR2 | C11orf91 | MTX1 |
| NEO1 | USP2 | RASSF5 |
| GAD1 | GPR89A | PARL |
| PITRM1 | DGKD | GATAD1 |
| CTTN | MCM3 | ZC3HAV1 |
| DENND5A | TDRP | DCAKD |
| BTBD3 | SLC25A19 | SYT14 |
| LMTK3 | ZNF362 | FAM161A |
| SLC7A11 | RIMKLA | PLAC9P1 |
| BRI3 | ELAPOR1 | MCUB |
| NDUFA7 | SMARCAL1 | SLC12A7 |
| TAMALIN | FHIT | AK9 |
| NIPSNAP2 | MARCHF11 | PEBP4 |
| ANKH | CDH10 | FADD |
| ERH | CDC23 | CASP4 |
| HDAC11 | TRIM39 | SPATA7 |
| UBE2H | DYNC2I1 | KANTR |
| ZMAT3 | LINC02696 | MTM1 |
| C1QB | LPXN | SLC2A5 |
| SH3BGR | EED | TARS2 |
| PLEKHG5 | AEBP2 | TATDN3 |
| USP15 | RNFT2 | FANCL |
| FCER1G | GLT1D1 | METTL21A |
| SH3BP5L | KBTBD7 | THUMPD3 |
| GABRA4 | TDRD9 | GTF2H4 |
| CFAP97 | INO80 | MAD2L1BP |
| POLR1C | HIRIP3 | SLC29A1 |
| SLC17A5 | SETD1A | ZNRF2 |
| SLC20A2 | NFATC3 | SDHAF3 |
| DENND2B | SCARF1 | ZNF37A |
| TMEM109 | XAF1 | ZNF22 |
| MEIS2 | ZNF264 | RNF31 |
| RGS11 | ZBTB45 | EHD4 |
| CLUAP1 | MCAT | ENKD1 |
| ELAC2 | BRWD3 | DDX28 |
| LINC00511 | SEPTIN6 | SFI1 |

| | | | |
|---|---|---|---|
| | NXT1 | SLC10A3 | ZNF654 |
| | RRP1B | DPY19L4 | RUNX1 |
| | KRBOX4 | CASP9 | ACBD6 |
| | LRP1B | NBPF1 | HECW2 |
| | TADA2B | PDIK1L | FAM50B |
| | SERPING1 | GMEB1 | MIR219A2HG |
| | CCNB1IP1 | EPHA10 | EFS |
| | SCAMP2 | POU3F1 | PIK3IP1 |
| | ANKRD29 | DPH2 | BRI3BP |
| | FBXO42 | LRRC40 | DYRK3 |
| | BCL9 | OLFM3 | MCF2L2 |
| | TMCC2 | LINC00623 | CTSO |
| | EML6 | NBPF11 | CFAP97 |
| | GPD2 | THEM4 | FKBP14 |
| | GORASP1 | METTL25B | CDK6 |
| | IGSF11 | BRINP3 | TNFRSF10D |
| | ATP11B | ETNK2 | CPB2-AS1 |
| | DGKQ | HSD11B1 | MEX3B |
| | HERC3 | SLC30A10 | PLPPR3 |
| | CMTR1 | ANKRD36C | BBC3 |
| | PHF3 | FASTKD1 | KPTN |
| | MOXD1 | CT75 | C21orf91 |
| | CREB3L2 | PPARG | CBY1 |
| | HR | OXNAD1 | MOSPD2 |
| | CNTRL | KCNH8 | INTS6L |
| | NTNG2 | ZNF852 | CNPY3 |
| | CSTF2T | CNTN3 | TRIM52-AS1 |
| | C11orf91 | ZDHHC23 | PHTF2 |
| | PYROXD1 | NEK11 | ACHE |
| | SLC11A2 | ACAD11 | SOHLH1 |
| | THAP2 | ZNF595 | TACO1 |
| | ZBTB1 | TMEM128 | SLC1A6 |
| | AREL1 | NOA1 | MHENCR |
| | PARN | HELQ | TMEM185A |
| | BFAR | SAP30-DT | KDM5D |
| | BOLA2B | CFAP90 | LRRTM4 |
| | LLGL1 | PLCXD3 | KCNE4 |
| | AKAP10 | CRHBP | SEMA3F |
| | SYNRG | ATG10 | PAQR9 |
| | RAD51C | KLHL3 | FOXQ1 |
| | GGA3 | ZFP62 | TNXB |
| | TMC6 | HACE1 | KIAA1586 |
| | FPR1 | GTF2IRD2B | FAM131B-AS1 |

| | | |
|---|---|---|
| ZNF444 | COL1A2 | ATAD2 |
| FASTKD5 | ZNF277 | IDI2-AS1 |
| SLC2A11 | DGKI | FZD8 |
| SHROOM2 | NTAQ1 | SEC31B |
| WWC3 | NAA35 | SYCE1 |
| ARMCX4 | TRIM32 | EPS8L2 |
| CYB561D1 | SH2D3C | TMEM138 |
| SIN3A | ADAMTS13 | LPCAT3 |
| ELK1 | FRMPD2 | CDK8 |
| KTN1 | PWWP2B | KLHL28 |
| MAP2K4 | FEN1 | MTFMT |
| EMC8 | PCNX3 | CTNS |
| RBL2 | MUS81 | SPAG5 |
| ARHGAP33 | NUDT8 | TGIF1 |
| PVR | XRRA1 | ZNF426 |
| RABAC1 | EMSY-DT | HOOK2 |
| ALG14 | GNB3 | ZNF446 |
| ECM1 | PYROXD1 | PLEKHH3 |
| DDR2 | BMAL2 | EPB41L5 |
| GPR17 | FMNL3 | NDST1 |
| TMEM108 | TNS2-AS1 | CYREN |
| UVSSA | SLC15A4 | RASSF8-AS1 |
| RNF150 | SOX21 | NUP85 |
| TTC39B | DNAAF2 | GUCD1 |
| GLE1 | MLH3 | CENATAC |
| TXNDC16 | ALG1 | MEGF10 |
| CIB2 | COG8 | IFITM1 |
| NIP7 | FA2H | B3GAT2 |
| TAF12 | LDHD | PRELID3B |
| MED8 | TIMM22 | OTUD6B |
| MTF2 | MYO1D | FOLH1 |
| GPR61 | HEXIM2 | ORMDL2 |
| LINC01138 | ZNF652 | RILPL2 |
| NBPF19 | SOX9-AS1 | MRPL45 |
| MINDY1 | BTBD17 | TRIT1 |
| ZNF687 | BAIAP2-DT | STAT4 |
| RIIAD1 | ASXL3 | DUSP28 |
| PIGM | ZNF506 | CNTN4 |
| ZBTB37 | ZSWIM9 | NGLY1 |
| SYT14 | DSCAM | KLHL8 |
| VPS54 | CACNA1I | SEC24B |
| MOGS | LINC01315 | CARMIL1 |
| LINC01106 | NUP50-DT | HLA-DRB5 |

| | | |
|---|---|---|
| ZDBF2 | TBL1X | SCX |
| STK16 | ABCD1 | SLC25A16 |
| EDEM1 | ZFY | TMEM109 |
| MAPKAPK3 | INTS3 | PHF12 |
| ACY1 | KLHDC9 | NAT9 |
| PPM1M | COP1 | TANGO2 |
| ADAMTS9 | MARK1 | ZXDB |
| ZIC1 | MERTK | THOC2 |
| ARHGEF26-AS | KCNIP1 | FBLN2 |
| UBA6-DT | PRXL2C | CCDC112 |
| MCUB | SPTLC2 | TSTD3 |
| ABHD18 | NAA20 | TMEM64 |
| LRBA | LYPLA2 | MIRLET7A1HG |
| PDLIM3 | CABIN1 | ZBTB34 |
| ZDHHC11B | DCAF6 | PRKCQ-AS1 |
| DAB2 | LPGAT1 | VSTM4 |
| SPRY4 | BCL11A | CYP2E1 |
| ARL10 | NAB1 | WEE1 |
| CNOT6 | CNOT9 | FZD4 |
| ATF6B | SLMAP | PCED1B-AS1 |
| PDSS2 | MRPS18A | FLRT2-AS1 |
| CEP57L1 | PLXDC2 | MYL11 |
| MTHFD1L | AFAP1L2 | ZNF629 |
| ERMARD | CTSC | PIGL |
| LFNG | PTPRO | POLI |
| LINC03011 | ZDHHC17 | ZNF844 |
| RINT1 | CIAO2A | ZNF677 |
| MEST | GAA | PXMP4 |
| ZNF398 | DLGAP1-AS4 | TOP3B |
| ZNF777 | CLASRP | MICALL1 |
| PPP3CC | FAM210B | CEP85 |
| PHF20L1 | CXCR4 | ZMPSTE24-DT |
| ARHGAP39 | RAB30 | HS1BP3 |
| ZNF251 | REPIN1 | PIGF |
| NOL6 | TBC1D17 | SLC25A20 |
| SMC5 | EHD3 | GNB4 |
| ANXA1 | ARID1B | INTS12 |
| CHST3 | DOCK9 | HDAC9 |
| GRID1 | IFI27L1 | STX17 |
| ABRAXAS2 | LRP3 | ANKRD26 |
| SYCE1 | COMMD7 | FGFBP3 |
| ZBED5-AS1 | MAGEH1 | DGKA |
| SPINDOC | NOC2L | TMEM19 |

| | | | |
|---|---|---|---|
| | NUDT8 | DFFA | ZNF821 |
| | ZW10 | BCAP29 | DLGAP1-AS1 |
| | ZNF202 | MAN1B1 | PLP2 |
| | LTBR | NOL12 | CYB561D2 |
| | ZNF740 | PGBD5 | USP13 |
| | SUOX | COMMD1 | TNIP2 |
| | MARCHF9 | GYPC | ART3 |
| | ABTB3 | TMEM129 | CYP2U1 |
| | DTD2 | PPA2 | EXOC3-AS1 |
| | ZFYVE26 | ARRDC3 | ERAP1 |
| | TEDC1 | ATXN2 | RBM24 |
| | NUBP2 | CKMT1A | ZSCAN31 |
| | ZNF213 | KDELR1 | PLAGL1 |
| | NTAN1 | RUVBL2 | GSDME |
| | SNHG30 | STX16 | PUS7 |
| | PCGF2 | EPM2AIP1 | LAMB1 |
| | TACO1 | C6orf136 | ZNF212 |
| | COLEC12 | LRRC24 | ERBB2 |
| | PRELID3A | ODR4 | SCARF2 |
| | ZNF561 | SOD3 | WNT7B |
| | QTRT1 | DMXL1 | UPRT |
| | PGPEP1 | ATP11A | MID2 |
| | ZNF260 | ZNF846 | PIR |
| | TMEM91 | SLC9A1 | CXCL1 |
| | ZC3H4 | SLC39A7 | CCDC127 |
| | SEC23B | GTF3C4 | ELF2 |
| | GUCD1 | TSPAN14 | ITGB3 |
| | INPP5J | DDX25 | ZNF519 |
| | TBC1D22A | FOXJ2 | AAR2 |
| | GEMIN8 | RHBDL3 | SALL1 |
| | PIR | HRH3 | RSKR |
| | RBM41 | MRTFA | FTSJ3 |
| | ADD2 | SNIP1 | RPS6KA1 |
| | SLC37A3 | PHTF1 | TMEM125 |
| | PPP2R2D | NPL | LIX1L |
| | PWP2 | BIRC6 | TBC1D22B |
| | MXI1 | REV1 | PGAP2 |
| | WAPL | C2orf88 | BDNF-AS |
| | ZRANB2 | SLC6A11 | LTO1 |
| | PODXL2 | METTL6 | SSPN |
| | CLMN | PLCD1 | SBNO1-AS1 |
| | PHB1 | TMF1 | TSEN54 |
| | PYGB | LRRC58 | ZNF576 |

| | | | |
|---|---|---|---|
| | P4HB | SLC35G2 | JPX |
| | HPCA | MYL5 | COL4A5 |
| | TXNDC15 | NLN | DAB1 |
| | BACE1 | MARCHF3 | GREM2 |
| | RPN1 | RIPK1 | ENTPD3 |
| | SLC2A13 | SLC35A1 | TMEM42 |
| | VDAC1 | ASF1A | PCYOX1L |
| | UNC5B | UST | OSBPL3 |
| | PAXX | EEPD1 | ZNF273 |
| | COPZ1 | ARMC1 | PENK |
| | CTSA | SNX30 | ARL14EP |
| | PEX11B | GGTA1 | TTC9C |
| | BBX | NRP1 | CEP126 |
| | SLC7A14 | PLEKHA1 | CLEC2B |
| | ZDHHC4 | PHRF1 | HEATR5A |
| | SUPT6H | BTBD10 | CCDC92B |
| | NSRP1 | KBTBD4 | DHRS7B |
| | SOCS2 | ZDHHC5 | FBXL12 |
| | MATK | CCDC91 | PPAN |
| | PPP1R14B | KITLG | ZNF579 |
| | MSH6 | CINP | GALE |
| | LZTFL1 | CLPX | WIPF1 |
| | C9orf72 | NR2F2 | LTA4H |
| | FRMD4A | BANP | LINC00320 |
| | MYBBP1A | TMEM107 | DPH5 |
| | ZNF652 | TMEM98 | OBSCN |
| | SUPT4H1 | MMP28 | EFCAB2 |
| | SPHK1 | SKA2 | AGBL5 |
| | GSE1 | TWSG1 | DENND6A |
| | ZMYM6 | DOHH | C3orf33 |
| | RAPH1 | SMG9 | RFC4 |
| | MALSU1 | AKAP17A | PLAAT1 |
| | MARCHF5 | SCML1 | SCFD2 |
| | CHRM1 | SYT13 | METTL14 |
| | LRRC49 | POLR3K | HTR1A |
| | AEN | RNF185 | MSH3 |
| | CD2BP2 | CEBPA | RIC1 |
| | MAPK12 | DIPK1A | PTPRD-AS1 |
| | CXADR | STAMBP | ZDHHC16 |
| | BAZ2A | PLCL1 | ABRAXAS2 |
| | TMEM240 | BRPF1 | DCHS1 |
| | COL16A1 | NXPE3 | LOH12CR2 |
| | PPP1R12B | NEPRO | WNT10B |

| | | |
|---|---|---|
| FH | TMEM223 | CDAN1 |
| TMF1 | CORO1C | MBTD1 |
| DHX29 | POMT2 | ZNF222 |
| PANK3 | ARHGAP17 | HIF3A |
| DENND1A | ZNF304 | ZNF766 |
| STAM | ARG2 | SMIM17 |
| PDCD11 | HIVEP3 | ZNF550 |
| QSER1 | CYP2J2 | PRR14L |
| TMEM135 | EEIG2 | SLC38A6 |
| FOXO1 | SIKE1 | IFI44 |
| TPT1-AS1 | THBS3-AS1 | KRCC1 |
| PIAS2 | SERTAD4-AS1 | LMAN2L |
| FAM83D | FLVCR1 | LINC01102 |
| POFUT2 | IARS2 | NEK10 |
| MINDY2 | TRIM11 | ZNF608 |
| FBLL1 | EIF2B4 | FBXO16 |
| CLDN10 | GPATCH11 | NUDCD1 |
| ZNHIT1 | PNPT1 | ENTPD1 |
| H2AC25 | SMARCC1 | LRP4-AS1 |
| PABIR1 | USP19 | GEMIN2 |
| TMEM38B | BMP2K | STYX |
| DPCD | METAP1 | GSTZ1 |
| MGAT2 | FSTL5 | ZWILCH |
| TADA2A | AADAT | ALDH1A3-AS1 |
| FAM117A | DNAJC18 | ZNF423 |
| RPS6KA1 | REV3L | TRPV2 |
| CTBS | KPNA5 | ZFP82 |
| GJC2 | TECPR1 | ZNF528 |
| ZNF852 | NAPEPLD | ZFP64 |
| PCGF3 | COG5 | FAM43B |
| NOTCH4 | KAT6A | ATG4C |
| RIPPLY2 | TGS1 | RNPEPL1 |
| UAP1L1 | ZNF704 | ADAMTS9 |
| ECD | ZNNT1 | HCLS1 |
| FRAT2 | RIMS2 | ARHGEF26 |
| MTG1 | NMRK1 | ATPSCKMT |
| SIPA1 | NOL8 | HARS2 |
| MYBPC1 | MSANTD3 | MYCT1 |
| RILPL2 | RNF20 | SPAG1 |
| TGFB3 | TLR4 | ZDHHC12 |
| ALDH1A3-AS1 | ARRDC1 | SLC29A3 |
| EEF2K | COMTD1 | P4HA1 |
| ASGR1 | ATE1 | VPS33B |

| | | | |
|---|---|---|---|
| | PLEKHH3 | CHST15 | COPZ2 |
| | TLK2 | NKAPD1 | CCDC117 |
| | ZNF224 | LINC02762 | CXorf38 |
| | ZNF473 | SNRNP35 | FUT11 |
| | ZMAT5 | ELF1 | QTRT2 |
| | PPARA | CLN5 | RAB43 |
| | MIR222HG | SLITRK5 | LYAR |
| | RGS7 | MNAT1 | CTXN3 |
| | CIAO1 | TMEM63C | USP49 |
| | CNOT9 | GTF2A1 | INTS8 |
| | DPP8 | RAMAC | CNTNAP3 |
| | EGLN1 | KRBOX5 | CENPP |
| | SNRPA1 | ATXN1L | ZEB1-AS1 |
| | RBSN | CPNE7 | ERCC6 |
| | SNTB2 | NXPH3 | FUOM |
| | PDCD6 | PECAM1 | PHLDA2 |
| | ZBTB18 | TMX3 | RAD9A |
| | RASA1 | PIAS4 | LPAR6 |
| | CCAR1 | WDR83 | L3HYPDH |
| | EIF4EBP2 | SLC35E1 | ZNF628 |
| | FNDC3A | KLHL26 | OSER1-DT |
| | FAM89B | TSHZ3 | PPP1R3D |
| | ZMYM4 | DMAC2 | PCDH11X |
| | KDM5B | GEMIN7 | RBM41 |
| | TNS3 | KIAA1755 | AGTRAP |
| | EXOC5 | TOX2 | SZT2 |
| | PALS1 | IFNAR2 | NTNG1 |
| | CTSH | ACOT13 | RDH14 |
| | SLC25A39 | BMAL1 | CAD |
| | SPATA20 | MGP | SENP7 |
| | HGS | RFK | POGLUT1 |
| | SMARCA1 | MTMR1 | RYK |
| | PTTG1IP | CIAO1 | SNX25 |
| | NIFK | DEK | TENT2 |
| | TMEM178B | RWDD1 | HMGN4 |
| | TNKS2 | IDH2 | ABHD11 |
| | RASA3 | MBTPS1 | LSMEM1 |
| | CPNE1 | NBR1 | HILPDA |
| | FAAH | UFD1 | RNF122 |
| | DPYSL5 | OPN3 | BAALC-AS1 |
| | MAP9 | YIPF4 | ADAMTSL2 |
| | APBB3 | DPY19L1 | RET |
| | HECW1 | TCAF1 | TRAF6 |

| | | |
|---|---|---|
| ZHX1 | NELFB | EMSY |
| AMBRA1 | ORMDL3 | PAN2 |
| SLCO2B1 | GPD2 | PAPLN |
| CCDC82 | ZFYVE16 | CRABP1 |
| PEDS1 | DIAPH1 | SNHG21 |
| SHISAL1 | DELE1 | PIEZO1 |
| DAD1 | SGCE | FAM210A |
| ARMC10 | CHPF2 | ZNF254 |
| DOCK1 | CIBAR1 | FITM2 |
| MYL12A | ABL1 | SHISA8 |
| HIGD2A | KCNC1 | TRABD |
| SYPL1 | MARS1 | ATP6AP1-DT |
| RANBP6 | MON2 | UGT8 |
| POGK | PCDH8 | LINC01138 |
| PIP4K2C | TAOK2 | SHROOM3 |
| DNAJC24 | NELFCD | GDNF |
| PPHLN1 | RILPL1 | SRSF12 |
| PLEKHM1 | TGOLN2 | MTERF3 |
| AGPAT1 | PLXNA2 | TOR2A |
| ADAMTS4 | MFSD1 | RPUSD2 |
| ZNF667-AS1 | PEX2 | APBA3 |
| IDH3B | ZNF516 | KRI1 |
| PABPN1 | LNX1 | ADGRE2 |
| POLR2C | CCDC167 | ENOX2 |
| ST3GAL5 | BRAF | ITCH |
| PTPN23 | RIGI | TMEM51 |
| PRDM16-DT | CUL2 | MTF1 |
| ASAP3 | FNBP4 | AGBL4 |
| DAB1 | CD27-AS1 | LINC02607 |
| RWDD3 | MGAT2 | WNT2B |
| CNIH3 | ACOT1 | BGLAP |
| RBKS | ZNF267 | WDR75 |
| MCEE | COX11 | MARCHF4 |
| PLGLB1 | TIAM1 | NAA80 |
| METTL8 | NCAPH2 | CBR4 |
| NPPC | SLC4A7 | GUSBP1 |
| CCDC174 | ZNF131 | FAM151B-DT |
| MUSTN1 | TIAM2 | EIF4EBP3 |
| EEFSEC | IKZF5 | TRIM52 |
| RAB43 | MAML2 | BYSL |
| NAAA | VPS45 | QRSL1 |
| HELQ | ECM1 | ICA1-AS1 |
| KLHL8 | VPS72 | FGL2 |

| | | | |
|---|---|---|---|
| | OTULIN | BATF3 | EPHX2 |
| | RNF144B | B3GALT1 | PTCH1 |
| | ZSCAN9 | IL17RB | ERCC6L2 |
| | IER3-AS1 | SLC35A5 | SFXN4 |
| | PSMB8 | SENP2 | ANO3 |
| | PPIL6 | ZNF330 | GPR83 |
| | GET4 | JMY | ARID2 |
| | MUC12 | HEBP2 | LINC00507 |
| | CLDN15 | GRB10 | IFT88 |
| | MTMR7 | KMT2C | REC8 |
| | FBXO16 | LCNL1 | IPO4 |
| | UNC5D | UPF2 | RFX7 |
| | TRMT12 | BORCS5 | NEIL1 |
| | ADCK5 | LATS2 | PTPN9 |
| | PLGRKT | DNAL1 | TVP23A |
| | GALT | MFAP1 | XRCC1 |
| | TOR1B | SOCS1 | COL6A2 |
| | CELF2-DT | SPATA33 | MN1 |
| | NUTM2B-AS1 | SLC26A11 | PHF6 |
| | DNMBP | FZR1 | ANKRD50 |
| | POLL | ZNF317 | MPDZ |
| | PCGF6 | ADA | SYCE1L |
| | ELP4 | BRD1 | NKAIN1 |
| | POLD3 | LDOC1 | SLC30A7 |
| | USP2-AS1 | DPY19L3 | PLCH1 |
| | DGKA | SH3BP2 | BCHE |
| | CEP83 | DOC2A | TMEM161B |
| | NHLRC3 | ITGA6 | CETN3 |
| | SOX1 | CIR1 | SGCD |
| | WDHD1 | HIBCH | TRERF1 |
| | TDRD9 | TRIM26 | PACRG |
| | CLBA1 | GNAI1 | TMEM60 |
| | SNURF | GTF3C5 | TRIM35 |
| | THAP10 | ARHGAP12 | STAR |
| | CRABP1 | ARMC5 | SLC39A4 |
| | FAH | TRMT1 | ANKRD18A |
| | HOMER2 | ZNF536 | TCERG1L |
| | RAMAC | TRIM7-AS2 | LRRC27 |
| | WDR73 | MRPL2 | METTL15 |
| | TTC39C | ESRRA | NAALAD2 |
| | ST8SIA5-DT | NT5DC3 | SLC37A4 |
| | S1PR5 | EIF2S1 | ZNF140 |
| | SARS2 | ZNF232-AS1 | SORD |

| | | | |
|---|---|---|---|
| | ZNF814 | SH2D5 | WDR73 |
| | A1BG-AS1 | DOCK7 | TLCD1 |
| | RBBP9 | PRPF38B | WIPI1 |
| | DSN1 | DRAM2 | CDH19 |
| | ADA | KIAA0040 | ZSCAN1 |
| | TAF4 | NUP133 | ACTR5 |
| | RTEL1 | SH3YL1 | RPS6KA6 |
| | MN1 | PTPN4 | HSF4 |
| | XPNPEP3 | MARCHF7 | ZNF571 |
| | SMIM45 | CTDSPL | AMER3 |
| | ADCY3 | GBE1 | NXPH2 |
| | UFC1 | NMD3 | RUVBL1 |
| | COTL1 | GABRA4 | PRMT9 |
| | PCYT1A | PHAX | MRPS30-DT |
| | MICALL2 | BRD8 | SIMC1 |
| | NOMO1 | TDP2 | ZSCAN26 |
| | LAP3 | ARMC10 | TNPO3 |
| | PLEKHA3 | EMC2 | CNTRL |
| | MAPK8 | MRPL13 | PBX3 |
| | MED19 | DERL1 | NUTM2B-AS1 |
| | AKAP17A | SCRIB | AP5B1 |
| | PVALB | GLIPR2 | UNC93B1 |
| | MYDGF | KIFBP | ABCC9 |
| | EPHB6 | BMPR1A | ACACB |
| | KLF6 | BUB3 | ALKBH1 |
| | PSMC2 | PIK3C2A | C16orf87 |
| | EIF2S3 | TSG101 | KCNJ16 |
| | AP3M2 | NAB2 | ARMC7 |
| | NPDC1 | GPHN | CEP131 |
| | P4HTM | NSMCE1 | TAF4B |
| | DOC2A | ZNF771 | PRKD2 |
| | USP36 | CKLF | ZNF71 |
| | TRIM69 | MON1B | SAMD10 |
| | ITGAX | SPATA2L | MRPL20-AS1 |
| | HM13 | GGA3 | CCDC30 |
| | PLPPR5 | TEN1 | PIGC |
| | CNOT6L | PTPRM | CYP1B1 |
| | RGL2 | DOK6 | IFIH1 |
| | AGFG2 | NACC1 | SFMBT1 |
| | NSMAF | SPEF1 | ECE2 |
| | GFRA1 | OFD1 | CLCN2 |
| | EPS8 | COL18A1 | ADRA2C |
| | MAPKAPK5 | DARS1 | ARL15 |

| | | | |
|---|---|---|---|
| | FGF14 | ARAP2 | FCHO2 |
| | FXR2 | ANKRD12 | CCHCR1 |
| | DCC | CDK11B | PMS2P4 |
| | CACTIN | CDK11A | FZD1 |
| | EMID1 | CYP4X1 | CRISPLD1 |
| | BOP1 | SLC27A3 | AKNA |
| | TMEM179 | TOMM40L | SUFU |
| | CHIC1 | TADA1 | AS3MT |
| | MRTO4 | GALNT14 | FLRT1 |
| | ZBTB40 | MAP4K3-DT | CLEC7A |
| | MIER1 | DCTN1-AS1 | RHEBL1 |
| | RPAP2 | RPIA | CRADD |
| | B3GALNT2 | NIFK-AS1 | RAB35-AS1 |
| | PTRHD1 | FZD5 | DPF3 |
| | SEPTIN10 | NUP210 | NPIPB13 |
| | AMMECR1L | CFAP44 | PYDC1 |
| | ACKR3 | LINC01094 | MAPT-AS1 |
| | EOGT | SPEF2 | SCRN2 |
| | KLF15 | NUP43 | RFPL2 |
| | IFT122 | ZNF138 | RPGR |
| | CTSO | NSUN5 | LRCH2 |
| | SETD9 | WWP1 | |
| | DAXX | IFT74 | |
| | BYSL | ANKRD16 | |
| | RRP36 | C10orf88 | |
| | LRRC73 | ZNF143 | |
| | UTRN | ZBED5-AS1 | |
| | PACRG | VWCE | |
| | POLR1F | POLD3 | |
| | C7orf25 | NARS2 | |
| | GNG11 | RESF1 | |
| | PDIA4 | DBX2 | |
| | LYPLA1 | DPY19L2 | |
| | VPS13B | ZNF26 | |
| | IDNK | OBI1 | |
| | TNC | MAP3K9 | |
| | SPOUT1 | ADSS1 | |
| | NRARP | LINC02352 | |
| | GPR158 | TMEM62 | |
| | TSPAN14 | ALDH1A2 | |
| | LDB1 | SNX22 | |
| | NFKB2 | SLX1A | |
| | CD151 | CNEP1R1 | |

| | |
|---|---|
| KCNQ1OT1 | SLC46A1 |
| IGHMBP2 | CRLF3 |
| ARID2 | PEX12 |
| STAT6 | TOM1L1 |
| SNRNP35 | TTC39C |
| DDX55 | ZNF521 |
| PGAM5 | MEX3D |
| SLITRK1 | ZNF554 |
| VCPKMT | KANK3 |
| L2HGDH | CD22 |
| BBOF1 | ZNF566 |
| PYGO1 | LIPE |
| ARHGAP17 | BLOC1S3 |
| CDH8 | FUT1 |
| CPNE7 | RALGAPA2 |
| CCDC43 | LKAAEAR1 |
| BAHCC1 | WDR4 |
| DUS1L | TFIP11-DT |
| TNFAIP8L1 | PTCHD1 |
| KLF2 | RAP2C |
| LIN37 | KIF3B |
| AXL | CD53 |
| BMP7 | ADAM17 |
| TANGO2 | PRKDC |
| STEEP1 | SRRM3 |
| PLXNA3 | CEPT1 |
| CMC4 | POLR3C |
| CAPN7 | ATP8B2 |
| USP13 | CEP350 |
| DCUN1D4 | SOX13 |
| NAA15 | MDM4 |
| SLIT3 | LINC00342 |
| MRM2 | AFF3 |
| RNF216 | FBLN7 |
| OSBPL3 | ACVR1 |
| SLC7A2 | ZNF142 |
| PTGR1 | CNOT10 |
| NET1 | SIAH2 |
| TTC9C | C3orf80 |
| ING4 | GUF1 |
| PGAP6 | SERINC5 |
| SDF2L1 | MAML1 |
| PLA2G6 | FBXL4 |

| | |
|---|---|
| BCORL1 | NUP205 |
| SUN1 | CDC26 |
| SMG1 | SORCS1 |
| TDG | LGR4 |
| COP1 | FJX1 |
| MTR | DTX4 |
| TNFAIP6 | EIF1AD |
| HDAC3 | SSH3 |
| NUB1 | RNF214 |
| KCTD9 | ST3GAL4 |
| HEY1 | COQ5 |
| NAPRT | CAB39L |
| C8orf82 | TRPM7 |
| CDC26 | CNTROB |
| ODF2 | C17orf58 |
| ARHGAP12 | RNF152 |
| MSANTD4 | MPV17L2 |
| RXYLT1 | CEP89 |
| KHNYN | SLC7A10 |
| TMEM63C | ZNF568 |
| TRIP11 | ZNF581 |
| RNF111 | ZNF134 |
| RPTOR | ZNF274 |
| RAB8A | YDJC |
| DZANK1 | TTC28-AS1 |
| IFNAR1 | INPP5J |
| CHM | TAF1 |
| PRRT3 | TMEM35A |
| SETD1B | NLGN4Y |
| TAF9B | RAB13 |
| RRM2B | FAM228B |
| USP5 | ZDHHC24 |
| BHLHE41 | KIF17 |
| GAREM1 | PTCH2 |
| GAB1 | MMACHC |
| TM2D3 | ITGB3BP |
| NPIPA1 | PRKAB2 |
| CHD6 | TDRKH |
| GRIPAP1 | THBS3 |
| SH3GLB1 | EDEM3 |
| PLEKHA6 | ZNF496 |
| ACER3 | LYPD8 |
| ASXL1 | GNLY |

| | |
|---|---|
| SEZ6L | C2orf49 |
| GPATCH2 | SEPTIN10 |
| DNMT3A | POLR1B |
| MYNN | TMEM177 |
| MYOM2 | DCAF17 |
| XKR4 | TTLL4 |
| INTS8 | ARMC9 |
| ERLIN1 | SETMAR |
| SNX32 | XIRP1 |
| GPN3 | CCDC51 |
| PPL | ZNF141 |
| CASTOR1 | CTBP1-DT |
| MPST | GPRIN3 |
| TMEM51 | CYP2U1-AS1 |
| MOB3C | SCLT1 |
| CRYZ | RXFP1 |
| C1orf162 | TMA16 |
| LINC01750 | C7 |
| POLR3C | CMYA5 |
| PRUNE1 | RNF144B |
| MTARC1 | MOG |
| JMJD4 | CUL7 |
| LMAN2L | CYB5R4 |
| BCL2L11 | CEP57L1 |
| CFAP210 | ZUP1 |
| MAP3K20 | MOXD1 |
| NUP35 | STXBP5-AS1 |
| PMS1 | RGS17 |
| SMARCAL1 | PRKN |
| SUMF1 | DNAAF5 |
| OSBPL10 | ELFN1 |
| EXOSC7 | CPVL |
| TMEM39A | NEUROD6 |
| HEG1 | C7orf25 |
| NCK1 | TYW1 |
| ECT2 | TBL2 |
| B3GNT5 | FUT10 |
| CLCN2 | SFRP1 |
| SCFD2 | STK3 |
| MOB1B | HAUS6 |
| SLC39A8 | ZNF510 |
| LINC00499 | MRRF |
| ETFDH | CACFD1 |

| | |
|---|---|
| PARP8 | LINC02668 |
| RNF180 | CCDC7 |
| MTX3 | RHOBTB1 |
| MSH3 | PAPSS2 |
| CDKL3 | IDE |
| GALNT10 | GSTO2 |
| SNRNP48 | TRIM68 |
| ELOVL2 | SPINDOC |
| PRR3 | KCNK4 |
| EGFL8 | KBTBD3 |
| SAYSD1 | HRK |
| CILK1 | TRIM13 |
| CAHM | TMEM229B |
| STYXL1 | MEG8 |
| MBLAC1 | GABPB1 |
| ALKBH4 | ZNF609 |
| KIAA1549 | BRICD5 |
| KRBA1 | CPPED1 |
| PAXIP1-DT | BFAR |
| PARP10 | ITGAM |
| GLIS3 | KCNJ12 |
| CD274 | LIG3 |
| STX17 | XYLT2 |
| MSANTD3 | CEP76 |
| SUSD1 | CEP192 |
| PRKCQ-AS1 | LINGO3 |
| LINC00839 | CHAF1A |
| SLC16A9 | RTBDN |
| RHOBTB1 | PLVAP |
| ASCC1 | NFKBID |
| DLG5 | C19orf47 |
| EXOSC1 | CCDC97 |
| UBTD1 | ZSCAN5A |
| C10orf95-AS1 | ZNF471 |
| OSBPL5 | A1BG-AS1 |
| LRRC51 | ZNF584 |
| SOX5 | ZNF324 |
| PUS7L | UBOX5 |
| ACSS3 | PLCB4 |
| MSI1 | FAM182A |
| LRCH1 | CSTF1 |
| PHF11 | MTG2 |
| OBI1 | CHRNA4 |

| | |
|---|---|
| UBAC2 | TMEM121B |
| AP4S1 | TRABD-AS1 |
| SSTR1 | SLC38A5 |
| GSTZ1 | PORCN |
| AP4E1 | NUDT10 |
| MYO1E | GPR173 |
| MTFMT | PABPC1L2B |
| SNX33 | KLHL4 |
| PRC1 | RNF128 |
| MSRB1 | DCX |
| ZNF23 | TRPC5 |
| GEMIN4 | WDR44 |
| P2RX5 | HS6ST2 |
| RAB34 | KCNJ9 |
| LIG3 | KCNAB1 |
| MMP28 | TRIM27 |
| RECQL5 | HAGHL |
| CEP131 | ROCK2 |
| APBA3 | HMGCL |
| ADAMTS10 | CNNM3 |
| CCDC159 | ATG7 |
| ZNF738 | RARS1 |
| ZNF254 | SFMBT2 |
| LYPD5 | BRMS1L |
| ALDH16A1 | MIR155HG |
| ZNF471 | KLHL15 |
| ZNF337 | ATAD3B |
| NFATC2 | THAP3 |
| IL10RB | TTC4 |
| SLX9 | CACNA1E |
| DGCR8 | H2AC25 |
| CCDC157 | ASB3 |
| PHKA2 | KCNIP3 |
| SPIN2B | PER2 |
| TSPAN6 | ABCB1 |
| MOSPD1 | TAF6 |
| HAUS7 | PRAG1 |
| ALDH1A1 | TMEM254 |
| HES4 | SYVN1 |
| PTPRE | CCDC90B |
| ABCA5 | SFSWAP |
| THAP3 | COL4A2 |
| PHC3 | GABPB1-IT1 |

| | |
|---|---|
| SMARCC2 | CD276 |
| PAF1 | RAB26 |
| HDAC5 | NAE1 |
| DHDDS | KREMEN1 |
| BRPF1 | RASD2 |
| DIPK2A | EXOSC10 |
| TNFRSF21 | ALPL |
| FAM20C | ST6GALNAC3 |
| EEPD1 | LSP1P5 |
| SLC45A4 | BCL9 |
| ARMC5 | C1orf54 |
| LARGE1 | KLHL12 |
| MRPL2 | ACBD3 |
| NAPEPLD | AFTPH |
| SLC31A2 | ALS2 |
| PLCH2 | NBEAL1 |
| TMEM125 | CTDSP1 |
| MPHOSPH10 | SP100 |
| MGAT4A | ZNF385D |
| ALDH1L1 | MELTF-AS1 |
| MBD4 | PDS5A |
| PIGZ | FRYL |
| CCNG2 | AP1AR |
| ABCE1 | SPRY1 |
| ZCCHC10 | FAM153CP |
| LYRM4 | CLK4 |
| KCTD20 | HIVEP1 |
| HYCC1 | RIPOR2 |
| TRRAP | ADGRF5 |
| DGKI | KLHL7 |
| NBN | KCND2 |
| ERC1 | PNOC |
| NALCN | ZFHX4 |
| PCID2 | CCN3 |
| DCAF5 | TATDN1 |
| ZNF710-AS1 | COMMD5 |
| IGF1R | PRPF4 |
| SRCAP | BCCIP |
| ATXN1L | TMEM134 |
| MTMR4 | AAMDC |
| ZNF461 | AMOTL1 |
| C21orf91 | DLAT |
| JAM2 | PIP4K2C |

| | | |
|---|---|---|
| | MOSPD2 | GTF2H3 |
| | TIMP3 | EBPL |
| | SMG5 | SALL2 |
| | ICE1 | GCH1 |
| | ULK1 | LPCAT4 |
| | CYB5B | CGNL1 |
| | CXCL16 | PHKB |
| | JOSD2 | NOL3 |
| | EAF1 | TMEM231 |
| | CINP | HIC1 |
| | COMMD2 | GHDC |
| | TLN1 | RETREG3 |
| | SPTY2D1 | FAM171A2 |
| | JKAMP | SOCS6 |
| | EMP3 | CDC25B |
| | OSBPL2 | URB1-AS1 |
| | SRRD | ANKRD54 |
| | PBDC1 | RBMX2 |
| | BRSK1 | EOLA2 |
| | SAMD1 | KIF21B |
| | STRAP | MXD1 |
| | PLPP2 | LDB2 |
| | PSMD6 | SPON1 |
| | LRP11 | DIXDC1 |
| | AKIP1 | XIAP |
| | GDAP1 | POLR3H |
| | ATAD3A | GPATCH4 |
| | ZBTB41 | TUT4 |
| | MICAL1 | STK17B |
| | UBE2W | UBA3 |
| | ACO1 | DZIP3 |
| | ZNF511 | MCCC1 |
| | INPPL1 | EIF5AL1 |
| | BCAT1 | GPR137C |
| | GPATCH2L | LGALS9 |
| | LACTB | CHD6 |
| | TMEM38A | DIP2A |
| | NIPSNAP1 | TAF9B |
| | FOXK2 | FHL3 |
| | C1orf35 | ZFYVE9 |
| | DELE1 | FNBP1L |
| | TOPORS | GPR161 |
| | ZSWIM8 | KCNT2 |

| | |
|---|---|
| NDST2 | TBCE |
| LGI1 | CNOT11 |
| MED17 | IWS1 |
| SHMT2 | CAVIN2 |
| RGS6 | KANSL1L |
| SETD1A | DIS3L2 |
| RHBDL3 | THRB |
| MED24 | SUCLG2 |
| HDAC10 | NOP14 |
| SCO2 | STX18 |
| SFRP1 | CC2D2A |
| C3 | SMARCAD1 |
| TMEM63B | SLC39A8 |
| SMG1P5 | BDP1 |
| PIAS4 | GFM2 |
| APH1A | ZBED3 |
| ANGEL2 | KIF13A |
| MORN2 | H4C3 |
| TBC1D23 | SRPK1 |
| SMIM43 | RAB23 |
| RASSF8 | COQ3 |
| CAPRIN2 | POLR1F |
| ABHD13 | PDE1C |
| VAT1L | ASL |
| MRPL12 | AP4M1 |
| YTHDF1 | MOSPD3 |
| CCDC6 | MCPH1 |
| PIH1D1 | POMK |
| LANCL2 | UBXN2B |
| INPP5A | CRH |
| CCPG1 | RUNX1T1 |
| DNAJC10 | GLIS3 |
| MSX1 | GKAP1 |
| PDLIM2 | SECISBP2 |
| RUFY2 | GAPVD1 |
| PCBD1 | DOLK |
| EFS | DDX50 |
| BRD7 | RGR |
| ZDHHC7 | GPAM |
| SOCS6 | TRUB1 |
| MARK4 | SPRN |
| DENND6B | TRIM22 |
| MTMR1 | AKIP1 |

| | | |
|---|---|---|
| | ADO | SBF2 |
| | NELL1 | BEST1 |
| | ORMDL1 | SERPINH1 |
| | LSM2 | MSANTD4 |
| | SHFL | NINJ2 |
| | RAB4A | DYRK4 |
| | TSEN15 | ZNF384 |
| | MKRN2 | EMG1 |
| | LSM8 | DDX47 |
| | PDK2 | ABCD2 |
| | SLC39A3 | USP15 |
| | RUVBL2 | DYRK2 |
| | ELP1 | MGAT4C |
| | TLN2 | DNAH10OS |
| | TMEM126A | PRPF39 |
| | WHAMM | COQ6 |
| | RNF40 | TUBGCP5 |
| | BCAS3 | JMJD7 |
| | GGCT | USP3 |
| | DPYSL4 | ADPGK |
| | BCL7A | ZNF592 |
| | MRPS35 | RLBP1 |
| | DICER1 | CLUAP1 |
| | MARS1 | COG7 |
| | ABLIM1 | SLC38A7 |
| | CEBPZOS | ZC3H18 |
| | UBP1 | TADA2A |
| | ADH5 | MIF4GD |
| | ACSL1 | TRAPPC8 |
| | LONRF1 | PIK3C3 |
| | AP1M1 | SLC1A5 |
| | RPSA2 | ZNF667 |
| | TRIM9 | CRNKL1 |
| | CPEB4 | NFS1 |
| | SOX4 | COL20A1 |
| | FBXO41 | SH3BGR |
| | LIN7C | SLC25A17 |
| | REEP3 | TRAPPC2 |
| | SLC2A5 | PRRG1 |
| | EMC1 | APLN |
| | SLC1A4 | TXLNGY |
| | GIT1 | ZBTB8OS |
| | ARAP2 | TNFAIP6 |

| | |
|---|---|
| ZNF592 | MYOM2 |
| CARMIL2 | SH3PXD2A |
| FLCN | SENP6 |
| LRRC4B | TBC1D13 |
| UBE2R2 | NF2 |
| GPR137B | FBXO28 |
| POLR1B | STRN |
| EMG1 | CNPPD1 |
| ZKSCAN1 | SMARCD1 |
| DUSP4 | FBLL1 |
| CCDC3 | ZMYM6 |
| NFYC | SUSD4 |
| OPN3 | BOD1L1 |
| TTC14 | FIP1L1 |
| GCNT4 | BMPR1B |
| ILRUN-AS1 | MCTP1 |
| ORC3 | NRSN1 |
| PEX1 | COX19 |
| CSMD1 | ASIC1 |
| ZNF395 | TSPAN31 |
| SPON1 | NPRL3 |
| PAMR1 | DNAL4 |
| SALL2 | LZIC |
| GNPNAT1 | CROCC |
| PDCD7 | ZDHHC18 |
| SLC39A11 | WDTC1 |
| ZNF121 | GPBP1L1 |
| FCGRT | MRPL9 |
| HPS4 | CRTC2 |
| B3GALT6 | TOR3A |
| TMEM201 | MRPL53 |
| PIK3CD | ANKRD39 |
| EPHB2 | LYPD1 |
| FAM76A | UNC80 |
| ST6GALNAC5 | DNPEP |
| TGFBR3 | DTYMK |
| LIX1L | OGG1 |
| CHD1L | GPAT3 |
| NVL | FAM169A |
| CTDSP1 | RIOK2 |
| DGKD | SLC17A5 |
| FARP2 | ANKRD6 |
| TTLL3 | APTR |

| | |
|---|---|
| FSTL1 | KCNH2 |
| WWTR1 | RBM33 |
| SLITRK3 | NSMAF |
| STX18 | WDR5 |
| SLIT2 | BMS1 |
| PAICS | GRID1 |
| RIMOC1 | LZTS2 |
| MDC1 | LINC00294 |
| ZNF76 | RCE1 |
| NHSL1 | CNPY2-AS1 |
| FUCA2 | BICDL1 |
| PPP1R14C | ERCC5 |
| MATCAP2 | SLC7A8 |
| SMURF1 | LIN52 |
| PPP1R35 | EML1 |
| TMEM176B | TNFAIP2 |
| FAM167A | ZFYVE19 |
| SNX16 | RGS11 |
| RIGI | TMEM132E |
| ZNF510 | KANSL1 |
| PTPN3 | PIAS2 |
| PRPF4 | ISYNA1 |
| AKNA | ZNF98 |
| AGPAT2 | BCL3 |
| NOXA1 | ZNF83 |
| DHTKD1 | PTPRT |
| MTPAP | NFATC2 |
| ZNF22 | GART |
| TACC2 | CCDC134 |
| RRAS2 | RAB39B |
| PRMT3 | ARL5A |
| IFTAP | BCKDHA |
| SLC39A13 | ITGA2 |
| SYT12 | TCERG1 |
| EED | KLC4 |
| GXYLT1 | AP5Z1 |
| CDK4 | POLB |
| TMTC3 | SPTY2D1 |
| KITLG | RAPGEF3 |
| KLF5 | ATP9B |
| NOP9 | CORO1B |
| RPS6KA5 | XPR1 |
| PLD4 | GGPS1 |

| | |
|---|---|
| GALK2 | KANSL3 |
| USP3 | CCNT2 |
| RABEP2 | LNPEP |
| ZNF689 | IQCE |
| CES2 | CHCHD3 |
| ZFPM1 | CPQ |
| PLD2 | COG3 |
| LRRC75A | TIMM9 |
| FMNL1 | INPP5K |
| SLC38A10 | LZTR1 |
| CCDC57 | RTL6 |
| SMCHD1 | SEC24A |
| PTPRM | ATF7 |
| SMAD7 | AHNAK2 |
| TIMM29 | ZBTB17 |
| ZNF844 | ARHGEF10L |
| ZNF791 | ZBTB40 |
| NOTCH3 | GPN2 |
| SIPA1L3 | OSCP1 |
| NECTIN2 | KDM4A |
| KCNC3 | TMEM69 |
| TGM2 | CDC7 |
| SLC2A4RG | SLC35A3 |
| CCDC117 | NUDT4B |
| GSPT2 | LRRN2 |
| EIF1AY | INTS7 |
| ADCK1 | KCNS3 |
| DMPK | VPS54 |
| TNFRSF14 | LINC02245 |
| HCN3 | PCBP1-AS1 |
| FHIT | PAX8-AS1 |
| NEK11 | CCDC93 |
| RNF168 | IRAK2 |
| SMAD5 | XPC |
| FOXC1 | CEP63 |
| MTRES1 | NCK1 |
| ZNF273 | CHST2 |
| STMP1 | SLC7A14 |
| ARRDC1 | FNDC3B |
| MINPP1 | BLOC1S4 |
| INTS5 | FAM13A |
| POLE | TMEM184C |
| CBR3 | TMEM267 |

| | |
|---|---|
| MTMR3 | SLC6A7 |
| PIGA | FAM153A |
| FUNDC1 | BPHL |
| WDR44 | NOTCH4 |
| CENPV | TJAP1 |
| FBXW5 | ASCC3 |
| ACAT1 | EPM2A-DT |
| DDI2 | HIP1 |
| GMEB1 | SEMA3C |
| GBP2 | SH2B2 |
| DUSP23 | IMMP2L |
| UCK2 | TMEM209 |
| SCYL3 | VPS37A |
| SMC6 | CD274 |
| UBXN2A | TDRD7 |
| ASXL2 | VAV2 |
| RNF25 | ARRDC1-AS1 |
| COPS7B | FHIP2A |
| IQCA1 | ZNF511 |
| RNPEPL1 | BET1L |
| CAPN10 | LRRC4C |
| MTMR14 | CYB561A3 |
| GLT8D1 | RELA-DT |
| PLCXD2 | NEU3 |
| PLXND1 | FAR2 |
| TMCC1-DT | ATF1 |
| LRCH3 | GPALPP1 |
| ZNF518B | CKAP2 |
| PTGER4 | KLF12 |
| MRPS30-DT | SYNJ2BP |
| MCCC2 | ENTPD5 |
| GFM2 | GPR68 |
| SCAMP1-AS1 | FAM98B |
| ABLIM3 | STRC |
| PDGFRB | LEO1 |
| LINC01574 | FAM174B |
| UNC5A | SNX29 |
| GMDS | ZNF668 |
| RIOK1 | CDH5 |
| PAK1IP1 | WWP2 |
| KLC4 | NXN |
| SLC29A1 | BORCS6 |
| EFHC1 | FAM222B |

| | |
|---|---|
| HEY2 | MTCL1 |
| RSPH3 | LMNB2 |
| BMPER | LINC00662 |
| MUC12-AS1 | CCNE1 |
| PRKRIP1 | KCNC3 |
| TRBC2 | ZNF211 |
| PEBP4 | ZNF814 |
| GTF2E2 | DSN1 |
| PDE7A | ZGPAT |
| ZNF7 | PCBP3 |
| GNE | MTFP1 |
| FBXO10 | H1-0 |
| PBX3 | PRR34-AS1 |
| ZBTB34 | LINC01278 |
| CACFD1 | OCRL |
| SAPCD2 | F8A1 |
| PRPF18 | TIMP1 |
| FGFBP3 | SST |
| SORCS3 | GMPPA |
| NANOS1 | HERC3 |
| PSTK | CLDN12 |
| ADCY6 | EEA1 |
| SMUG1 | AGL |
| RASSF3 | SH3PXD2B |
| EEF1AKMT1 | FBXL20 |
| RCBTB2 | TTC28 |
| LINC02352 | FTX |
| JMJD7 | HBA1 |
| SQOR | TEX30 |
| ANKDD1A | MAP3K3 |
| ZWILCH | TNFRSF6B |
| ACSF3 | |
| CTC1 | |
| INTS2 | |
| NETO1 | |
| FXYD5 | |
| ZNF568 | |
| ZNF571 | |
| CLEC11A | |
| HAR1A | |
| HUNK | |
| YPEL1 | |
| APOL6 | |

| |
|---|
| FBLN1 |
| CPT1B |
| RAI2 |
| ACOT9 |
| NEXMIF |
| APOOL |
| RNF128 |
| ABCD1 |
| SRP54 |
| NEIL2 |
| XPA |
| KCTD13 |
| POLDIP2 |
| ARRDC2 |
| CCDC90B |
| GPBP1L1 |
| CEP350 |
| KIAA1143 |
| PARP14 |
| KLF3 |
| LPCAT1 |
| MIER3 |
| RPAP3 |
| RAB35 |
| RILPL1 |
| COG4 |
| SPATA2L |
| FAM171A2 |
| COX11 |
| ZNF302 |
| TBC1D17 |
| SLC35E2B |
| BMPR1A |
| FAM216A |
| CDH11 |
| TSPO |
| MEAF6 |
| OTUD3 |
| SF3B4 |
| GMPS |
| VPS8 |
| FAM153CP |
| VEGFA |

| |
|---|
| RCC1L |
| CHCHD3 |
| RARRES2 |
| PTAR1 |
| PRXL2C |
| HTATIP2 |
| CCNK |
| ZNF597 |
| ZC3H7A |
| VPS16 |
| ITCH |
| TXNDC9 |
| FAM117B |
| OBSL1 |
| SNRK |
| NEK4 |
| HLTF |
| FNDC3B |
| ATP13A4 |
| SDAD1 |
| TENM3 |
| SPEF2 |
| NAIP |
| SLC6A7 |
| VARS1 |
| MRAP2 |
| REV3L |
| GSTK1 |
| MMP16 |
| MATN2 |
| UNC13B |
| IPPK |
| ZMYND19 |
| NUDT5 |
| EIF5AL1 |
| PPP1R3C |
| FHIP1B |
| PRR5L |
| TP53I11 |
| DTX4 |
| CASP4 |
| UBR1 |
| PML |

| |
|---|
| AXIN1 |
| INO80E |
| NFATC3 |
| GHDC |
| POLG2 |
| MYO15B |
| ME2 |
| MIR23AHG |
| NR2F6 |
| KAT14 |
| ACSS2 |
| SEC14L2 |
| AK2 |
| LUC7L2 |
| CDK11B |
| NPPA |
| NECAP2 |
| LINC00339 |
| ZNF436-AS1 |
| TRIM62 |
| TTC4 |
| SLC35D1 |
| SLC44A3 |
| PRMT6 |
| DRAM2 |
| SYT6 |
| NBPF9 |
| CD84 |
| RXRG |
| RGS1 |
| B3GALT2 |
| SOX13 |
| TBCE |
| WDR35 |
| KLHL29 |
| ATAD2B |
| SLC5A6 |
| TRMT61B |
| PRKD3 |
| SANBR |
| WDPCP |
| ELMOD3 |
| ZNF514 |

| |
|---|
| ANKRD36B |
| LINC01102 |
| TMEM185B |
| GYPC |
| ACVR2A |
| CARF |
| INPP5D |
| CNTN4 |
| FBLN2 |
| VIPR1 |
| CACNA2D2 |
| CRYBG3 |
| BOC |
| SIDT1 |
| POGLUT1 |
| DTX3L |
| FAIM |
| P2RY1 |
| CEP19 |
| MYL5 |
| RBM47 |
| SLAIN2 |
| RASL11B |
| CXCL3 |
| SOWAHB |
| MFSD8 |
| CEP44 |
| ATPSCKMT |
| ZCCHC9 |
| CETN3 |
| GIN1 |
| PPIP5K2 |
| CEP120 |
| KLHL3 |
| PCDHB4 |
| PCDHB14 |
| LINC00847 |
| LY86 |
| GPLD1 |
| GPANK1 |
| TNXB |
| PPT2 |
| MDGA1 |

| |
|---|
| OARD1 |
| CUL7 |
| CD2AP |
| FBXL4 |
| SMPDL3A |
| NHSL1-AS1 |
| DNAAF5 |
| HERPUD2-AS1 |
| GLI3 |
| GTF2IRD2 |
| GTPBP10 |
| MOSPD3 |
| NOM1 |
| PINX1 |
| NUDT18 |
| PURG |
| MAILR |
| RSPO2 |
| NUDCD1 |
| FAM135B |
| PTP4A3 |
| GLDC |
| PIGO |
| OGN |
| MFSD14CP |
| LINC02977 |
| NIBAN2 |
| TOR2A |
| LINC00963 |
| PLPP7 |
| TTF1 |
| ADAMTS13 |
| AKR1C3 |
| PIK3AP1 |
| C10orf88 |
| LUZP2 |
| ABTB2 |
| ZFTA |
| GDPD5 |
| SERPINH1 |
| ALG8 |
| C11orf54 |
| ALKBH8 |

| |
|---|
| DDX47 |
| PTHLH |
| UBE3B |
| CCNA1 |
| DHRS12 |
| BIVM |
| RAB20 |
| ABHD4 |
| CARMIL3 |
| PRORP |
| FLRT2-AS1 |
| FAM181A |
| BCL11B |
| HHIPL1 |
| SRP14-DT |
| PLA2G4B |
| CD276 |
| CSPG4 |
| GOLGA6L9 |
| FES |
| IL32 |
| ZSCAN32 |
| CLN3 |
| CBLN1 |
| NKD1 |
| SMPD3 |
| MLYCD |
| ZNF778 |
| MRM3 |
| MED11 |
| DVL2 |
| DHRS13 |
| CCL8 |
| MLX |
| ITGA3 |
| KCNJ16 |
| BTBD17 |
| FADS6 |
| TYMS |
| DLGAP1-AS1 |
| CEP76 |
| ASXL3 |
| ZNF397 |

| |
|---|
| SALL3 |
| FGF22 |
| IZUMO4 |
| MAN2B1 |
| MVB12A |
| TMEM161A |
| PLEKHF1 |
| SLC7A10 |
| ZNF567 |
| ERCC2 |
| ZNF135 |
| EDEM2 |
| COL20A1 |
| DSCAM |
| ZNF74 |
| LINC01637 |
| TMEM191C |
| TTLL1 |
| PRR34-AS1 |
| ABCB7 |
| UPRT |
| DRP2 |
| PWWP3B |
| TMEM164 |
| EOLA2 |
| TMEM187 |
| TBCA |
| DANCR |
| VEGFB |
| KANK1 |
| DXO |
| DHRS4L2 |
| GAS2L1 |
| MRAS |
| SESTD1 |
| PTPMT1 |
| RPUSD3 |
| TRAPPC14 |
| IRF7 |
| STIM1 |
| NPAT |
| TMOD3 |
| QPRT |

| |
|---|
| CEP89 |
| SLC5A3 |
| GCNA |
| SCARB2 |
| SDR39U1 |
| SNORC |
| TRMT6 |
| AZIN2 |
| MANEAL |
| HECTD3 |
| RBM15 |
| SH3YL1 |
| DHX57 |
| KANSL3 |
| STAT4 |
| METTL21A |
| TMEM198 |
| USP19 |
| PLCH1 |
| PDGFRA |
| GAR1 |
| FER |
| KHDC1 |
| KRIT1 |
| FOXP2 |
| HTR5A |
| MFHAS1 |
| RGS20 |
| JPH1 |
| DNAAF11 |
| MINCR |
| NAA35 |
| ENTR1 |
| ERCC6 |
| GSTO2 |
| WDR11 |
| SIGIRR |
| ALKBH3 |
| PCNX3 |
| FMNL3 |
| SART3 |
| COQ5 |
| RNF34 |

| |
|---|
| TMEM132C |
| INSYN1 |
| METTL22 |
| BORCS6 |
| SLC46A1 |
| PTRH2 |
| EPG5 |
| ZNF333 |
| TRAPPC6A |
| BCAT2 |
| ZBTB45 |
| SLC24A3 |
| SLC35A2 |
| BRWD3 |
| UTP14A |
| VGLL4 |
| IER5L |
| MGST1 |
| ZER1 |
| IRF9 |
| HCN1 |
| MCRS1 |
| PORCN |
| SMARCA4 |
| AHDC1 |
| KCMF1 |
| SIVA1 |
| PRKAB2 |
| TRIM46 |
| PAPOLG |
| STAM2 |
| PHAX |
| RAB24 |
| MED27 |
| SFMBT2 |
| MORN4 |
| TIMM10B |
| NOPCHAP1 |
| PHAF1 |
| OPRL1 |
| PNMA8C |
| CHD9 |
| RAE1 |

| |
|---|
| IFT20 |
| CENPX |
| TOB1 |
| CARS1 |
| ZDHHC3 |
| NR2F1-AS1 |
| DDX46 |
| TUBGCP6 |
| ARL1 |
| MRPS10 |
| RALGAPA1 |
| ITGB5 |
| MAN2B2 |
| NETO2 |
| TIMM44 |
| DNAJB5 |
| PHLDA3 |
| RNASEH1 |
| MRPL35 |
| PANTR1 |
| TMCC1 |
| TTC33 |
| GRPEL2 |
| ANKMY2 |
| KIAA1217 |
| GDF11 |
| PLA2G4C |
| EBP |
| TNKS |
| SYNGR2 |
| ACSBG1 |
| IMPA1 |
| VCAN |
| MRPL22 |
| IRGQ |
| GLYR1 |
| ANKHD1 |
| UPK3BL1 |
| RAB5C |
| SATB2-AS1 |
| MRPL19 |
| GGA2 |
| KCNK3 |

| |
|---|
| INO80D |
| LSM6 |
| IBTK |
| PPIL4 |
| SHARPIN |
| MTAP |
| MLLT10 |
| TENM4 |
| SOCS4 |
| TRMT1 |
| GARRE1 |
| ZNF570 |
| DLL3 |
| FUZ |
| ZNF134 |
| LIME1 |
| DKC1 |
| KPNA6 |
| YRDC |
| FCGR1A |
| DYRK3 |
| PAX8-AS1 |
| BCS1L |
| CCDC149 |
| RICTOR |
| HEXB |
| KCND2 |
| ACTR3B |
| MOB3B |
| FAM120AOS |
| TGFBR1 |
| TSSC4 |
| TMEM134 |
| PRR14 |
| FHOD3 |
| TYK2 |
| LINC00662 |
| C16orf87 |
| OSBPL1A |
| C19orf53 |
| NUDT14 |
| ARHGEF2 |
| KLHDC10 |

| |
|---|
| BLVRB |
| SNAP91 |
| KLF16 |
| COA1 |
| SNAP47 |
| SDHAF2 |
| SNF8 |
| KIF21B |
| MED28 |
| PITPNM1 |
| DNAJC3 |
| AGTPBP1 |
| PEX16 |
| GCN1 |
| EAPP |
| YES1 |
| MECR |
| THEM4 |
| CAMKMT |
| FSTL5 |
| TCIM |
| CUTC |
| RGS10 |
| FZD4 |
| TAGLN |
| AVEN |
| EME2 |
| MIR497HG |
| DHRS7B |
| TBC1D16 |
| PDE4C |
| VASP |
| DNAL4 |
| BCKDK |
| SNRNP27 |
| SHISA9 |
| MAP3K4-AS1 |
| ADARB2 |
| ZDHHC5 |
| TNFAIP8L3 |
| TMED3 |
| CDK5RAP1 |
| BIN3 |

| |
|---|
| KSR2 |
| RFX5 |
| SAP30L |
| MMP17 |
| MCM4 |
| SNTG1 |
| ZNF704 |
| STARD3 |
| EXTL2 |
| C3orf80 |
| RASGEF1A |
| SNRNP40 |
| ODR4 |
| GPAT3 |
| SDHAF4 |
| SHPRH |
| PNOC |
| ADRA2A |
| PRKAB1 |
| STON2 |
| GABPB1-IT1 |
| MPV17L |
| COG7 |
| SOX9-AS1 |
| C2CD4C |
| LINC00665 |
| MACROD2 |
| RALGAPB |
| LINC01278 |
| F8A1 |
| MED12L |
| SLC35A1 |
| SMIM30 |
| TNKS1BP1 |
| BRAP |
| RABL3 |
| EID2B |
| VPS26A |
| RNF7 |
| BDNF |
| YDJC |
| PRDM11 |
| NUDT19 |

| |
|---|
| SCAF4 |
| UPF3B |
| DUSP6 |
| ZNF148 |
| CD53 |
| RBM7 |
| CREB3 |
| ARRB2 |
| DCTN5 |
| TNS2 |
| ERP29 |
| ME1 |
| CSRP2 |
| ITPA |
| DDX41 |
| HTRA2 |
| ABCG1 |
| VPS33B |
| WDR74 |
| STT3B |
| COQ8A |
| EBPL |
| MRPL38 |
| VIRMA |
| NCKAP5L |
| TRIAP1 |
| DNAJC9 |
| RANBP3 |
| ZCRB1 |
| POP7 |
| MYO10 |
| CRB1 |
| ZC3H12C |
| SPG21 |
| MAOA |
| HLCS |
| SGSM3 |
| PRR18 |
| DMWD |
| CKAP4 |
| COQ7 |
| SAMD14 |
| ADSL |

MZF1
GCAT
SFXN1
PDCL3
BBS7
TXLNA

| Cortical layer 2/3 | | |
|---|---|---|
| iTBS | iTBS ∩ 10 Hz | 10 Hz |
| CXCL8 | CCL4L2 | NRGN |
| GPR183 | CCL4 | SPARCL1 |
| IFI27 | CCL3 | SNAP25 |
| CXCL2 | CCL3L3 | GNAS |
| FLT1 | FBXO33 | SCN1B |
| PLEK | IL6 | EEF1A2 |
| TUBA1C | IL1RN | VSNL1 |
| RILPL2 | RBP4 | HS3ST2 |
| CHI3L1 | FRMPD2 | GFAP |
| RGR | CRH | CHGA |
| LRRC61 | PLAC9 | PLP1 |
| STUM | GPATCH4 | SNCB |
| CYBA | CHST15 | EFHD2 |
| RDH10 | FCGR1BP | CAMK2G |
| GPNMB | PEG13 | NDRG4 |
| KPNA3 | NEURL1B | NCDN |
| FZD1 | MYADM | NEFL |
| CEBPA | RAMP3 | CABP1 |
| IL1A | TREM1 | ALDOA |
| KLHL15 | ZNF865 | CCL2 |
| POLR3GL | LMNB2 | EGR3 |
| G0S2 | PAX8-AS1 | ZFP36 |
| FGFRL1 | P2RY12 | CHCHD10 |
| CEP104 | PDIK1L | GPI |
| HCST | RCL1 | STXBP1 |
| RO60 | PLCXD2 | GRIN1 |
| SMOC1 | LINC01007 | NEFM |
| CALB1 | EFEMP2 | ATP1A3 |
| LILRB4 | ZNF496-DT | NPTX1 |
| DGCR6 | BTBD1 | RAB3A |
| CPLX3 | CAPS2 | DKK3 |
| FRS2 | CBY1 | MAP2K1 |
| LRFN1 | C2CD4C | MAP1A |
| CXXC4 | NEUROD1 | HSPA5 |
| TUSC1 | AMMECR1L | DNM1 |
| CTXND1 | NOM1 | NGRN |
| ST6GAL1 | PIERCE1 | COX7A1 |
| MGP | TBP | SYN1 |

| | | |
|---|---|---|
| NAA16 | LRRC37A2 | ATP1B1 |
| BCAS4 | KTI12 | SNHG29 |
| LIX1L | PARG | BHLHE40 |
| HNMT | UTP25 | HERPUD1 |
| L2HGDH | POLR3C | VAMP1 |
| LY86 | NBPF12 | PDP1 |
| RGS19 | RDH13 | LZTS3 |
| MSR1 | SNX22 | DDIT3 |
| UQCC6 | GAS1 | SH3GL2 |
| DDX19A | RPAP3 | PNMA2 |
| LRRC3B | COX10 | RASD1 |
| DGKH | SEMA6C | NEFH |
| TCF3 | AGAP6 | SPRY2 |
| ITPR2 | LEMD3 | YPEL3 |
| ROCK1 | DHX8 | KIAA0513 |
| JPX | FRMD8 | NDFIP1 |
| NXPH2 | PUM3 | MALAT1 |
| SLC9A9 | ISLR2 | NCALD |
| SOS2 | MRPS31 | STX1B |
| NPRL3 | LRMDA | DIRAS2 |
| SPEF1 | RLBP1 | KCNAB2 |
| PLOD3 | EPB41 | PCSK1 |
| GSTO2 | LURAP1L | SCN4B |
| HVCN1 | FBLN2 | CLSTN2 |
| CDK5RAP1 | PARD3 | SSBP3 |
| BBC3 | AKR1C3 | CHAC1 |
| ANXA2 | CRIP1 | BMERB1 |
| CLDND2 | THEMIS | FAIM2 |
| LINC00595 | HSD11B1 | PVALB |
| P3H2 | ITGA9-AS1 | MOBP |
| MT1H | RBM24 | PNMA1 |
| RAD1 | LINC00507 | PREPL |
| MMP28 | THAP2 | DMTN |
| POLR1F | PRRC1 | CPLX1 |
| RFPL2 | GOLGA5 | NPTX2 |
| DPH5 | PYDC2-AS1 | KLF9 |
| ZNF488 | TUBGCP4 | RTN4RL2 |
| ZNF587 | GRIP1 | PHYHIP |
| MYL11 | BBS9 | TBR1 |
| GTPBP10 | FOLR2 | CEND1 |
| PIK3C2B | TMEM125 | CAMKK1 |
| ZBTB47-AS1 | GJB1 | SNAP91 |
| NUP210 | CCL20 | PFKP |

| | | |
|---|---|---|
| TEK | RIPOR2 | SSTR2 |
| HAVCR2 | HARS2 | PYGB |
| ARSB | EMSY-DT | GABRG2 |
| BLOC1S3 | BCL9L | UBE2QL1 |
| MTARC1 | LCP1 | PCBP2 |
| TP53I13 | STAR | CSRNP1 |
| COL26A1 | ARMC7 | ATP6V1G2 |
| CNOT4 | SNURF | NGEF |
| CCDC137 | COQ5 | HLF |
| OLMALINC | FREM3 | CPLX2 |
| TMCC1-DT | TCAP | TSC22D3 |
| APTR | GJC1 | SLC20A1 |
| CEP126 | FAM117B | NPTN |
| CCR1 | ERMP1 | ISCU |
| PPP1R3E | LINC01202 | ID3 |
| TNIP2 | P2RY11 | SCAMP5 |
| ANOS1 | ARHGEF1 | CADPS |
| NES | ZNF274 | NAPB |
| MARCHF9 | GPR68 | FAM43A |
| ERICH2 | CCDC93 | SCRT1 |
| PMAIP1 | PHACTR4 | KCNC1 |
| TCHP | C10orf88 | SERTAD1 |
| TVP23A | DDX28 | GDAP1L1 |
| CTNNA3 | SOSTDC1 | SST |
| C17orf100 | EIF5AL1 | SLC25A22 |
| FOXC1 | FRS3 | PEG3 |
| LRIG2 | CD300A | HEXIM1 |
| ZNF419 | PSPH | ZNF365 |
| GAL3ST1 | KCTD4 | SLC3A2 |
| ELF2 | GPLD1 | KLC2 |
| SLC43A3 | ZNF22 | ERRFI1 |
| SLC35B3 | PDSS2 | EPDR1 |
| GLDC | TMEM74 | KCNA2 |
| MFSD5 | PCDHGC3 | CLEC2L |
| COX18 | DCAF17 | CHD2 |
| SOX21 | XIRP1 | STAC2 |
| DNAH17 | POGLUT1 | SESN2 |
| PFAS | ITIH5 | LPCAT4 |
| TSPAN15 | CD52 | PCP4 |
| CITED4 | PRDM1 | ZDHHC22 |
| ELF1 | GRIN3A | IGSF9B |
| SPATA7 | ABCB10 | CAMK2A |
| LINC00623 | SETD9 | GADD45A |

| | | |
|---|---|---|
| CFAP97 | MSANTD2 | BAIAP2 |
| TMEM168 | MX2 | CSRNP2 |
| ARRDC4 | BCLAF3 | CNTNAP1 |
| GPR88 | LINC02668 | NAT8L |
| HELQ | MMACHC | GEM |
| CPVL | ALMS1 | FBXO32 |
| P3H3 | FGL2 | ANXA6 |
| KLF11 | PAPSS2 | PIP5K1C |
| CENPP | FAM161A | ETS2 |
| BTBD17 | ZFP64 | EVL |
| SLC36A1 | | GPR162 |
| ABHD12B | | AP2A1 |
| RRAS | | SLC2A3 |
| NUTM2B-AS1 | | CKS2 |
| IER3-AS1 | | SEPHS2 |
| ANKRD6 | | ISLR |
| SLC16A9 | | IFITM10 |
| OXNAD1 | | FAM3C |
| P2RX7 | | SLC25A12 |
| MINPP1 | | PFKM |
| DDB2 | | FAM163B |
| CD24 | | DEDD2 |
| WDR25 | | THRA |
| SLC19A1 | | CADPS2 |
| HDAC1 | | ADCY1 |
| ABTB2 | | ADRB1 |
| B4GALT4 | | RCAN2 |
| STX17 | | RRAD |
| SNX24 | | DLL1 |
| HYAL2 | | SDHA |
| CAV2 | | XBP1 |
| DGAT2 | | PIM3 |
| ZNF738 | | SLU7 |
| MYO19 | | USP2 |
| SATB2-AS1 | | MICAL2 |
| CCDC159 | | INA |
| ZSWIM9 | | COX7A2L |
| FBXL4 | | CDC37 |
| ZNF92 | | KHDRBS3 |
| C16orf95 | | MDK |
| NKAIN1 | | IQSEC3 |
| CDH9 | | RHBDL3 |
| SH3TC2 | | BSN |

| | |
|---|---|
| PSMG3-AS1 | EPHB6 |
| RUBCNL | KCNK1 |
| CHRAC1 | PPP1R10 |
| PCED1B-AS1 | FOXK1 |
| GPAT3 | PHF24 |
| WNT7B | TRIB1 |
| MRPL20-AS1 | KRT17 |
| TUNAR | COX20 |
| RPS6KA1 | FH |
| SNX18 | NPEPPS |
| TMEM161B | NDRG3 |
| SNX16 | SF1 |
| SYCE1L | RBFOX1 |
| CCL8 | PFKFB3 |
| TTC38 | CHST1 |
| NUP50-DT | CRABP1 |
| GBP2 | BCL2L2 |
| PLSCR3 | SPRYD3 |
| ITGB3BP | TRA2B |
| THBS4 | FBXW7 |
| TAF8 | IL1RL1 |
| ARHGEF26 | TOB1 |
| BCHE | PRMT7 |
| FKBP10 | BRD2 |
| RPIA | NDUFS2 |
| RASL11B | FBXO21 |
| FIGNL1 | EPB41L1 |
| LRRC2 | LRRC8A |
| CTXN3 | AOPEP |
| RP2 | SV2B |
| LGALS9 | PNMA8A |
| CD84 | UGP2 |
| PDCD11 | BLOC1S2 |
| UPRT | CHGB |
| DYSF | NFE2L1 |
| HPS3 | MYT1L |
| PPIL4 | RABGEF1 |
| MARCHF8 | TET3 |
| CASP4 | HIVEP2 |
| KLK7 | DAGLB |
| WDR81 | TMEM14A |
| MASP1 | IP6K1 |
| IFT74 | POLR2A |

| | |
|---|---|
| WDR90 | ITM2A |
| ABCD1 | HNRNPA0 |
| ACSL5 | HSPA12A |
| JPT2 | NAP1L5 |
| ARHGAP4 | RPRML |
| AP5S1 | CLPTM1 |
| FLNC | IER3 |
| CDK11B | AHDC1 |
| FCGR3A | DUSP16 |
| RSPH9 | ECM1 |
| MAS1 | LGALS3 |
| ADCY7 | TFRC |
| ETNK2 | CRY2 |
| RXRG | HSPA1A |
| SH2B2 | RNPS1 |
| TMEM209 | INPP5A |
| MAILR | AMPH |
| TSPAN4 | FOXG1 |
| ZNF561 | ZNF25 |
| SLC6A9 | SLC6A17 |
| GALNT10 | INPP5F |
| DOCK5 | RAB40C |
| PLAC9P1 | APMAP |
| FRZB | CRKL |
| IL33 | PRKAG2 |
| FZD4 | HAPLN4 |
| THEMIS2 | DAPK3 |
| LINC00499 | MAPK9 |
| RHBDF2 | RRP12 |
| LINC02977 | SYT7 |
| SRP14-DT | DAGLA |
| LACC1 | SIK2 |
| RAC2 | KIFAP3 |
| ANKRD18A | SYNDIG1 |
| SALL3 | ACOT9 |
| TNC | MIAT |
| | PPP1R14C |
| | ITPR1 |
| | SYNJ1 |
| | PRRC2B |
| | CAMTA2 |
| | AMZ2 |
| | COPS3 |

| |
|---|
| SLC35F1 |
| CLDN11 |
| CAMK1D |
| LRP4 |
| PRPF19 |
| CNIH1 |
| PRKAG2-AS1 |
| PSME4 |
| FOXJ2 |
| SEMA3E |
| ARHGEF25 |
| NTNG2 |
| SMYD2 |
| ATPAF1 |
| NAP1L3 |
| ABLIM1 |
| PI4KB |
| GALNT9 |
| SYNPO2 |
| TMEM248 |
| FBXO44 |
| SRRM4 |
| IP6K2 |
| ST8SIA5 |
| FOXO3 |
| PITPNA |
| ATXN2L |
| RELL2 |
| ZMYND8 |
| PAFAH1B2 |
| SYT2 |
| HS3ST4 |
| CITED2 |
| PTK2 |
| ALKBH5 |
| ZFYVE21 |
| PHYH |
| IFFO1 |
| CFAP20 |
| EDC4 |
| RERE |
| EIF3L |
| PAF1 |

| |
|---|
| GFOD1 |
| SPTBN2 |
| MAP7D2 |
| RGS16 |
| LINC01963 |
| IDH3A |
| TNFAIP6 |
| IARS1 |
| HR |
| TAMALIN |
| SCN1A |
| NECAP1 |
| RPRM |
| SLC39A14 |
| SDF2 |
| NFIC |
| HERC1 |
| ARRB1 |
| MAP2K3 |
| TSPYL4 |
| N4BP2L1 |
| KCNQ3 |
| ERP29 |
| SNHG1 |
| FOSL2 |
| DDX39B |
| RYR2 |
| LAMTOR3 |
| IL17D |
| DUSP14 |
| VAPB |
| BEND6 |
| H1-0 |
| RPS6KA2 |
| OPN3 |
| GPCPD1 |
| POLR2C |
| FKBP2 |
| LHX2 |
| KLHL21 |
| WDR47 |
| PER2 |
| SYBU |

| |
|---|
| PHLPP1 |
| OPCML |
| SCN2B |
| NFYC |
| MYOM2 |
| BICDL1 |
| DUSP5 |
| IMPA1 |
| HEBP2 |
| RASD2 |
| SHANK3 |
| ATN1 |
| MCF2L |
| TSHZ1 |
| DLC1 |
| MEPCE |
| PLEKHM2 |
| BEND5 |
| GPRASP1 |
| BRSK2 |
| NABP2 |
| SRPK2 |
| FAM131B |
| GAS6 |
| HPRT1 |
| KCTD10 |
| RNF41 |
| SBDSP1 |
| NR3C1 |
| SOX2 |
| FEZF2 |
| TSC2 |
| GLRX |
| KCNA1 |
| ARHGEF3 |
| RALYL |
| RIPPLY2 |
| PTPRT |
| BCOR |
| TMEM268 |
| STAU1 |
| NECAB3 |
| ZNF331 |

| |
|---|
| MIR22HG |
| TNFSF9 |
| CCDC6 |
| SRP68 |
| DUSP7 |
| SGSM1 |
| LINC01551 |
| PNMA8C |
| BLCAP |
| PDHA1 |
| KCNT1 |
| KIAA0408 |
| TMEM178A |
| CHST8 |
| ETNPPL |
| ANK1 |
| ABCF2 |
| NEIL2 |
| KIF3A |
| KIF21A |
| ANXA7 |
| PHF13 |
| NSG1 |
| PIM1 |
| LINC00869 |
| RAB24 |
| ALAS1 |
| ADI1 |
| ZFAND2A |
| CPNE5 |
| PCP4L1 |
| RAP1GAP2 |
| H2AX |
| EWSR1 |
| SNTB2 |
| TUBB3 |
| LINC01535 |
| FURIN |
| ARL4D |
| PDIA6 |
| PPL |
| SIK3 |
| FBXO34 |

| |
|---|
| LRRC4B |
| ODC1 |
| MRO |
| HMGB2 |
| HECW1 |
| ZNF667-AS1 |
| GSTM5 |
| CACNB1 |
| CLIP1 |
| IL34 |
| TAF7 |
| PJA1 |
| PDZD8 |
| KCNC2 |
| LUZP1 |
| ANKRD9 |
| ADGRA1 |
| RASGRP1 |
| RNF114 |
| ANAPC7 |
| CMIP |
| APLNR |
| CIZ1 |
| PRDM16-DT |
| PDPR |
| GFPT2 |
| MYBPHL |
| GFRA1 |
| ACOT1 |
| MYH7B |
| B3GAT3 |
| DCAF6 |
| DAB2IP |
| TOX4 |
| SPEN |
| SLC25A33 |
| EIF4A3 |
| KLF10 |
| EHD3 |
| KLF2 |
| MIR124-2HG |
| SMIM43 |
| SS18L2 |

| |
|---|
| NCAPD3 |
| CTSA |
| CASD1 |
| SLC4A10 |
| ACAA1 |
| SDS |
| RYBP |
| HYOU1 |
| ADM |
| PALM2AKAP2 |
| CTNNBIP1 |
| NONO |
| VSTM2B |
| RAB15 |
| EFNB2 |
| IQCK |
| BBS1 |
| PASK |
| ATG16L1 |
| DLG4 |
| NR4A2 |
| HRH2 |
| PTGR1 |
| FAM216A |
| CARMIL2 |
| OGDHL |
| SH3BP5 |
| KCNN1 |
| SLC32A1 |
| APBA1 |
| LSAMP |
| UNC5A |
| IMMT |
| MGST1 |
| JAK1 |
| RANBP9 |
| SNPH |
| RORA |
| ESRRA |
| CA7 |
| PCNT |
| BLTP2 |
| PIK3R3 |

| |
|---|
| CPEB2 |
| LMTK2 |
| PANK4 |
| SATB2 |
| UBE2W |
| RNF208 |
| SH3BGRL2 |
| CDK16 |
| NOL3 |
| WARS1 |
| PCDH7 |
| SNX27 |
| PPP1R9A |
| C2orf74 |
| CACNB4 |
| LCMT1 |
| PLEKHA1 |
| ZMYM2 |
| SHD |
| GOLGA7B |
| TLE3 |
| COA6 |
| ASRGL1 |
| SOX9 |
| LRRC75A |
| POLDIP2 |
| TEX2 |
| BTBD10 |
| BAIAP3 |
| MTRR |
| RPTOR |
| PRKCE |
| GLA |
| GPR155 |
| PLEKHG1 |
| TIAM1 |
| SUDS3 |
| NCK2 |
| ADD2 |
| ELOF1 |
| ELK1 |
| SSH1 |
| PDPK1 |

| |
|---|
| FOXN3 |
| RNF182 |
| OSGIN2 |
| PPP1R3F |
| PITRM1 |
| KANSL1 |
| MTR |
| MEAF6 |
| RIPOR1 |
| KCNS1 |
| FBXO9 |
| TMEM94 |
| KDM6B |
| MOB4 |
| DCAF8 |
| KMT2A |
| ADAM23 |
| EMD |
| PSPC1 |
| HDGFL2 |
| GRIK3 |
| PELI3 |
| LONRF1 |
| GLI4 |
| DSCAML1 |
| EHD4 |
| YY1AP1 |
| ZRANB1 |
| TSC22D2 |
| NXF1 |
| CCSAP |
| RESF1 |
| TECPR2 |
| ZNF530 |
| YTHDF2 |
| PDE4A |
| FAM13B |
| UFD1 |
| OGDH |
| PPP2R2A |
| BCL7A |
| PPP1R13B |
| AKR1C1 |

| |
|---|
| NOVA2 |
| USP7 |
| NETO1-DT |
| CACNA1B |
| UFM1 |
| CDC42BPB |
| PLEKHM1 |
| NR1D1 |
| THAP11 |
| DPF1 |
| FTO |
| TOMM22 |
| BACH2 |
| USP27X |
| SLC4A4 |
| KCNJ4 |
| ZYG11B |
| ACAD8 |
| DGLUCY |
| USP19 |
| KCTD8 |
| PDZD7 |
| CRTC2 |
| CCNB1IP1 |
| MRPL58 |
| DMKN |
| ENPP5 |
| MIDEAS |
| UBE2Q1 |
| PPP1R16B |
| MKX |
| ADCY8 |
| ZC3H7B |
| LRP8 |
| C1orf35 |
| PANX2 |
| SPNS2 |
| AFTPH |
| CAPN15 |
| EPHB1 |
| FGF13 |
| SIRT3 |
| DENND5B |

| |
|---|
| RIMKLA |
| CYTH2 |
| PPFIBP1 |
| PRKCA |
| TNPO2 |
| TOMM34 |
| STK40 |
| CERK |
| WDR74 |
| CHKB |
| DYRK1A |
| DECR1 |
| SIK1 |
| MAD2L1BP |
| STK35 |
| LAMC1 |
| SPRY4 |
| DENND3 |
| GPRC5A |
| NPLOC4 |
| SNAP47 |
| AQP1 |
| NCKIPSD |
| LMTK3 |
| BLVRA |
| R3HDM2 |
| ZCCHC2 |
| FBXL17 |
| KCND3 |
| TMCO3 |
| MSRA |
| SUSD6 |
| PLEKHG5 |
| PRICKLE2 |
| FBXO28 |
| ANKRD33B |
| COL12A1 |
| MIEF2 |
| REPS2 |
| PDRG1 |
| ESYT2 |
| HDAC4 |
| KDM3A |

| |
|---|
| FRMPD4 |
| NPPC |
| VILL |
| NXPH3 |
| SMG1P5 |
| NR1D2 |
| EAF1 |
| AMBRA1 |
| CCN1 |
| SPRED2 |
| STXBP5 |
| STX8 |
| BMS1 |
| OSBP |
| NT5C1A |
| CCDC3 |
| COQ10B |
| GRIK5 |
| CTH |
| ZBTB16 |
| KCNH3 |
| SSBP2 |
| RING1 |
| TRA2A |
| NUDT14 |
| SYNJ2 |
| REPIN1 |
| RTL5 |
| TXNDC15 |
| SORBS3 |
| CHM |
| TRIM8 |
| VCPKMT |
| ZNF622 |
| SRRT |
| SLC9A1 |
| TNIP1 |
| SOWAHA |
| FNDC9 |
| TLNRD1 |
| VGLL4 |
| TOMM40L |
| LRRC32 |

| |
|---|
| UAP1 |
| GDAP1 |
| ARFGAP3 |
| GATAD2B |
| GLS2 |
| PIAS4 |
| SPIRE1 |
| S1PR1 |
| GPBP1L1 |
| FAF1 |
| ZNF580 |
| DYNLT2B |
| KIF22 |
| UGT8 |
| TCTA |
| LDLR |
| FBXO3 |
| RTCA |
| ATG101 |
| IMPDH2 |
| BAZ1B |
| CBLN4 |
| PRICKLE1 |
| PRCP |
| CCDC91 |
| ARFIP2 |
| SLC7A1 |
| MTSS2 |
| THOC3 |
| ZBTB4 |
| MRPL23 |
| UNC119 |
| TIMM23 |
| METTL2A |
| CACNA1A |
| EIF2A |
| TOM1 |
| PSMG1 |
| SHOC2 |
| DNAJB14 |
| CYTH3 |
| TRUB2 |
| PARP3 |

| |
|---|
| KDELR1 |
| CDC42EP3 |
| EXOSC6 |
| PARVB |
| PLCXD3 |
| EIF5A2 |
| EIF2AK3 |
| PARVA |
| DCUN1D2 |
| COMMD2 |
| SRRM3 |
| BCL6 |
| WIPI1 |
| NANS |
| RPRD2 |
| GYG1 |
| SRPRA |
| GSTA4 |
| TRMT6 |
| PELP1 |
| VKORC1 |
| STOX1 |
| ADNP2 |
| GSTM2 |
| CIDEA |
| PCDH11Y |
| BCAR1 |
| SDSL |
| COG3 |
| TMEM18 |
| ZNF142 |
| RAC3 |
| CRTAC1 |
| ARIH2 |
| GGPS1 |
| SNRNP200 |
| PBXIP1 |
| RIN2 |
| LRRC57 |
| MIR124-1HG |
| MAGEE1 |
| SUPT6H |
| ZBTB21 |

| |
|---|
| AHNAK |
| MAP4K5 |
| HCN1 |
| ELAVL1 |
| ZSWIM7 |
| SEMA7A |
| ALDH4A1 |
| RB1 |
| PANK2 |
| PHKG2 |
| TMX3 |
| RRAGC |
| ABCC5 |
| SLC39A7 |
| KLHL18 |
| KCNIP2 |
| RAB11FIP3 |
| GPR137B |
| CREB5 |
| CARS1 |
| MCAM |
| UQCC4 |
| USP31 |
| AKAP17A |
| GEMIN7 |
| GAK |
| GTF2F1 |
| EGFL8 |
| LINC01503 |
| ADPRHL1 |
| SHB |
| RAI1 |
| ELP3 |
| RABL2B |
| CELF1 |
| ARHGEF11 |
| MDFIC |
| GRB10 |
| B4GALT6 |
| SIN3A |
| METTL23 |
| ANKMY2 |
| TMEM242 |

| |
|---|
| FBXO42 |
| RFX5 |
| DAPK1 |
| MIR9-3HG |
| CLUH |
| AACS |
| AJAP1 |
| ICE1 |
| FGFR1 |
| TMEM97 |
| GNA12 |
| SPATA2 |
| SEC23A |
| TAOK2 |
| LIMK1 |
| WNK2 |
| ULK3 |
| ANKRD17 |
| FAM81A |
| MKRN2 |
| WDR48 |
| SMG5 |
| CSGALNACT1 |
| ANKS1A |
| CLIP4 |
| MCRS1 |
| CNIH4 |
| OSBPL6 |
| CDH13 |
| FEM1A |
| MRPL22 |
| GAS2L1 |
| CRIM1 |
| TMEM145 |
| SEC16A |
| SHF |
| YTHDF1 |
| TOMM70 |
| CALN1 |
| ZNF367 |
| DDI2 |
| SLC30A7 |
| TNNT2 |

| |
|---|
| POMZP3 |
| ZNF140 |
| NRG3 |
| VIPR1 |
| BACE2 |
| KCNE4 |
| RAMACL |
| ZNF668 |
| KANSL2 |
| PKD2 |
| GFOD2 |
| TBC1D25 |
| RNF157 |
| FAT1 |
| BSDC1 |
| USP39 |
| G6PD |
| PAIP2B |
| VPS26A |
| MCTP1 |
| PPARD |
| HBP1 |
| CDK7 |
| FAM153CP |
| PTGR3 |
| MAT2B |
| PPP2R2D |
| DTD1 |
| MUS81 |
| GABPB1-IT1 |
| EFNA3 |
| CACNG8 |
| KCNK12 |
| JAKMIP1 |
| TIAM2 |
| FBXO31 |
| INPP5K |
| PACS1 |
| UQCC5 |
| TNS3 |
| ZNG1A |
| RAPGEFL1 |
| UTP11 |

| |
|---|
| ORMDL3 |
| IFFO2 |
| TIPRL |
| SRD5A1 |
| ZNG1E |
| CCZ1 |
| USP9Y |
| PDK3 |
| TERF2 |
| DMXL2 |
| GZF1 |
| ANKS6 |
| ANAPC2 |
| GABRG3 |
| GPR173 |
| PAXIP1 |
| ZNF250 |
| P2RX6 |
| PDS5B |
| GORASP2 |
| ADRA1B |
| BMAL1 |
| CSNK2A2 |
| CDS1 |
| THAP3 |
| EXOC3 |
| WDR36 |
| ZNF24 |
| PIK3CB |
| KIAA2013 |
| BOD1 |
| KCNC3 |
| UBE2D1 |
| SNRK |
| POLR2M |
| MARF1 |
| TNRC18 |
| NELL1 |
| CSNK1G3 |
| SLC9A7 |
| CNTFR |
| POPDC3 |
| LRP12 |

| |
|---|
| MEG9 |
| RANBP10 |
| MTMR1 |
| C6orf89 |
| SIPA1L2 |
| PEX26 |
| LAMP5 |
| TRIT1 |
| LINC01574 |
| NNMT |
| BCL11B |
| ZNF224 |
| TCF7L2 |
| KCTD13 |
| GALNT16 |
| WASHC2A |
| IQCE |
| VPS13D |
| DDX23 |
| KLF13 |
| PCYT1B |
| GPR89B |
| LMLN |
| ABTB3 |
| SEC11A |
| DPM1 |
| MARCHF7 |
| RABIF |
| MTA1 |
| AMFR |
| CRYBB2P1 |
| TMEM115 |
| DNAJC3 |
| OTUD3 |
| RNF26 |
| ZBTB38 |
| MED27 |
| EID2 |
| SLC35A1 |
| BRD8 |
| RPA1 |
| ARMC8 |
| NIN |

| |
|---|
| TMCC2 |
| BAHCC1 |
| ZBTB8OS |
| KLHL5 |
| PRKAA2 |
| DHX16 |
| FAM131B-AS2 |
| PRAG1 |
| DCUN1D3 |
| MAPK11 |
| TNRC6B |
| SLC25A28 |
| MTMR12 |
| ELP1 |
| NIFK |
| ILKAP |
| SLC25A38 |
| LMBR1 |
| C2CD4B |
| YRDC |
| STK11 |
| NEURL1 |
| MAIP1 |
| APPL2 |
| UNC45A |
| WDFY3-AS2 |
| RNF103 |
| NSUN2 |
| CACNA1G |
| CXCL1 |
| MYCT1 |
| ESRRG |
| TTC17 |
| ZNF592 |
| STRADA |
| KLHL12 |
| ORC3 |
| SOWAHB |
| KIAA1586 |
| OTUD6B-AS1 |
| ZFYVE1 |
| ST3GAL1 |
| MRPS30 |

| |
|---|
| POU3F2 |
| MAP3K8 |
| ZBTB22 |
| NFATC2 |
| HYCC2 |
| SPTY2D1 |
| MYC |
| RELT |
| SLC15A4 |
| FMN2 |
| FYTTD1 |
| ASNSD1 |
| RAB11FIP5 |
| RHOT1 |
| ICA1L |
| FAM193B |
| RELB |
| IFNGR1 |
| CDH10 |
| ZNF436 |
| EPM2A-DT |
| TAF3 |
| IFTAP |
| WDR46 |
| UNC80 |
| TUBA8 |
| UBE4B |
| NOS1AP |
| E2F6 |
| ZZEF1 |
| ARHGAP27 |
| ZNF324 |
| TBPL1 |
| SART1 |
| AAGAB |
| IGF1R |
| ACOX1 |
| GFPT1 |
| RASA3 |
| BAG4 |
| ENTREP3 |
| FAM98A |
| SERTAD2 |

| |
|---|
| VCPIP1 |
| MED19 |
| LMF2 |
| SNRNP27 |
| EXOSC4 |
| TRIM26 |
| OTUD1 |
| ZFYVE28 |
| FZR1 |
| CCDC82 |
| FLII |
| AZI2 |
| URGCP |
| STARD10 |
| STARD13 |
| THUMPD1 |
| POGZ |
| OLIG2 |
| SENP2 |
| IKBKG |
| IGFBP6 |
| NANOS3 |
| HCFC1 |
| ZNF84 |
| CRABP2 |
| C2CD5-AS1 |
| ZNF136 |
| ZNF571 |
| SLC35E2B |
| TBC1D7 |
| WHAMM |
| CACNG7 |
| CORO7 |
| CDK11A |
| FAM76A |
| GDPD5 |
| ALDH1A3 |
| CHL1-AS2 |
| ZNF318 |
| INPP5E |
| RNF6 |
| YOD1 |
| PTPN18 |

| |
|---|
| PHACTR2 |
| ZNF282 |
| HSPB6 |
| ELL |
| CBLN1 |
| UCHL5 |
| DLST |
| HIVEP1 |
| DENND2B |
| TMEM106C |
| GJB6 |
| COG5 |
| YEATS2 |
| THAP9-AS1 |
| NTSR2 |
| LARP1B |
| CASC15 |
| TNFRSF25 |
| MSANTD7 |
| MECP2 |
| DPP8 |
| PEX19 |
| CDH6 |
| UIMC1 |
| FAM161B |
| FBXL19 |
| BRCC3 |
| TK2 |
| NELFB |
| SLC43A2 |
| PPM1E |
| ETV3 |
| ENTPD3 |
| KATNA1 |
| PNPLA7 |
| TOP3A |
| ZNF304 |
| RPS6KC1 |
| IFT122 |
| GFRA2 |
| LINC00294 |
| UBP1 |
| PCNX2 |

| |
|---|
| UTP4 |
| CORT |
| CETN2 |
| IL1RAP |
| ZCCHC14 |
| DAZAP1 |
| SKI |
| TCF12 |
| MPHOSPH10 |
| HECW2 |
| MYRIP |
| HSPA4L |
| GAPVD1 |
| LINC00511 |
| SELENOO |
| SEMA4C |
| CLVS2 |
| KCNAB1 |
| GPR158 |
| PTPRM |
| ZNF787 |
| ACBD3 |
| MED12L |
| ESAM |
| TMTC1 |
| ARX |
| STX16 |
| MXD1 |
| SACM1L |
| UBQLN4 |
| SPIDR |
| KDM2A |
| SHISA6 |
| GRAMD2B |
| RPS6KA4 |
| KANSL1-AS1 |
| ZDHHC3 |
| MTF1 |
| ARNT |
| SEC22A |
| PM20D2 |
| AVEN |
| MIR193BHG |

| |
|---|
| SULT1A3 |
| ZNF565 |
| NEURL4 |
| NUP58 |
| CACNA2D3 |
| FAM13A |
| SART3 |
| PCIF1 |
| PLPP5 |
| MAP4K3 |
| AMIGO1 |
| TNRC6C |
| UBE2D4 |
| VPS11 |
| CRAT |
| COQ9 |
| PRKCI |
| AFF1 |
| TRAPPC10 |
| MRPS2 |
| SCAF1 |
| GPR3 |
| LHFPL4 |
| SS18L1 |
| STK17B |
| TOR1AIP1 |
| SPHK1 |
| KSR2 |
| B3GALT1 |
| FAM83D |
| DOP1B |
| CTDSP1 |
| CTIF |
| NT5DC2 |
| CEP70 |
| CEP41 |
| NIP7 |
| SMIM11 |
| BRPF1 |
| PPP2R1B |
| SGPL1 |
| NR2C1 |
| KLHL26 |

| |
|---|
| APH1A |
| LRRC49 |
| SUPT7L |
| DLGAP3 |
| GCNT4 |
| LONP2 |
| PDK1 |
| FXN |
| KRI1 |
| ZNF446 |
| C4orf33 |
| BRWD3 |
| NIT1 |
| LRWD1 |
| SERTAD3 |
| PTPN1 |
| RIOK3 |
| SPX |
| NIPA2 |
| KLHL29 |
| CAMKMT |
| DRD1 |
| SCX |
| DACH1 |
| PTGR2 |
| SETD5 |
| SLC12A8 |
| PLEKHM3 |
| PHC3 |
| SVBP |
| TINF2 |
| CAMSAP3 |
| COQ8B |
| UST |
| LRRC8C |
| INSR |
| TNFSF12 |
| CD320 |
| FCHSD2 |
| CCDC50 |
| EXOC6B |
| KANSL3 |
| ARHGEF10 |

| |
|---|
| CYB5R2 |
| RPS6KL1 |
| NUP133 |
| ITPRIP |
| HIRA |
| EZH1 |
| RNF8 |
| ADARB1 |
| TRIB3 |
| NAT9 |
| GAB2 |
| CWC25 |
| DNAJA3 |
| NKTR |
| FHIP1B |
| GPATCH8 |
| SUPV3L1 |
| PHKB |
| JAKMIP2 |
| SPON1 |
| DCUN1D5 |
| MZT1 |
| EXT1 |
| CERS5 |
| MED25 |
| RALGAPB |
| NECAB2 |
| SULT1A1 |
| MCM3AP |
| PIGK |
| SLC45A4 |
| CYB561 |
| RANBP6 |
| SBK1 |
| NFS1 |
| IRF3 |
| USP36 |
| STK38L |
| CIART |
| DAP3 |
| C1QB |
| MAP3K7 |
| TTBK2 |

| |
|---|
| SOD3 |
| ZNF384 |
| SLC35A2 |
| DHX57 |
| SHLD2 |
| BTBD7 |
| MBTD1 |
| SIPA1L3 |
| INTS12 |
| KRBA1 |
| GPR12 |
| FIZ1 |
| GTPBP6 |
| NDUFAF7 |
| SOAT1 |
| SLC38A3 |
| SETD1B |
| CDYL2 |
| SETBP1 |
| EPM2AIP1 |
| MED26 |
| NCBP3 |
| TSTD3 |
| SLC8B1 |
| COLGALT1 |
| TGFBRAP1 |
| CARMIL1 |
| RMND1 |
| BDNF |
| CBLN2 |
| INO80E |
| RYK |
| PGBD1 |
| LRFN2 |
| TRMT9B |
| RSPO2 |
| GMEB2 |
| GCNA |
| SETD7 |
| ARRDC2 |
| RBM4B |
| IRF9 |
| LRRC40 |

| |
|---|
| ST3GAL6 |
| PXYLP1 |
| ZMAT4 |
| NPIPA1 |
| PDCD2 |
| ZNNT1 |
| TLCD3B |
| ZDHHC6 |
| SLC9A3-AS1 |
| NME5 |
| CNOT8 |
| MACO1 |
| SVIL-AS1 |
| TMEM63C |
| LIN7A |
| ROM1 |
| RGS20 |
| DGKI |
| BIN3 |
| TSHZ3 |
| PAQR3 |
| POU6F1 |
| CLEC16A |
| ACAA2 |
| MARCHF5 |
| RNF216 |
| RRM1 |
| YAP1 |
| PLCG1 |
| CSDC2 |
| DNAJC9 |
| COL5A3 |
| LAMB1 |
| TOR3A |
| CELSR3 |
| FOXP4 |
| NPR2 |
| PLEKHH3 |
| TTC39C |
| MED8 |
| MIGA2 |
| FA2H |
| RETSAT |

| |
|---|
| DIPK2A |
| TMOD1 |
| H4C8 |
| ZNF76 |
| MTHFD1 |
| GMPS |
| BOD1L1 |
| FAM193A |
| LRATD2 |
| CLCC1 |
| VPS37A |
| ASTN2 |
| TEX261 |
| SMC3 |
| VWF |
| NOP2 |
| DHX40 |
| ZFYVE9 |
| SETD2 |
| ERCC5 |
| ZMYND19 |
| PIKFYVE |
| TRIM3 |
| ARMC5 |
| MAOA |
| AFDN |
| PALLD |
| BLTP3B |
| VAT1L |
| EMX2 |
| MTMR3 |
| STXBP3 |
| CDK13 |
| ASB13 |
| PPRC1 |
| PPIL2 |
| SAP30L |
| TPRA1 |
| TRAF3IP1 |
| TMEM267 |
| BMAL2 |
| TMEM233 |
| RIOX1 |

| |
|---|
| TRIM16 |
| PLXDC2 |
| HSD17B7 |
| HLA-F |
| SCARF2 |
| PAPOLG |
| H1-2 |
| DLG5 |
| LRRC23 |
| TUBGCP3 |
| WNT3 |
| SLC9A8 |
| MORC2 |
| TMEM88B |
| SGF29 |
| OSBPL9 |
| MELTF |
| RP9 |
| COX15 |
| SLC41A2 |
| FITM2 |
| TMEM185A |
| ASMTL |
| NFYB |
| CEP95 |
| COL5A2 |
| GPR89A |
| SEC16B |
| INPP4B |
| ZNF346 |
| KCNAB3 |
| RAB37 |
| CHAF1A |
| C2CD2 |
| PRKX |
| ZXDB |
| GPRASP3 |
| SLC7A8 |
| CNEP1R1 |
| MAP2K6 |
| TAFAZZIN |
| UGDH |
| CARTPT |

| |
|---|
| URB1 |
| CLDN15 |
| RUBCN |
| ELAPOR1 |
| GNPDA2 |
| TAF2 |
| SLITRK1 |
| TXLNA |
| ETHE1 |
| LINC01896 |
| TNFAIP3 |
| GFER |
| NPY |
| LATS1 |
| KCNG1 |
| ZNF451 |
| CNOT10 |
| MAD1L1 |
| GRID1 |
| RMC1 |
| ST6GALNAC5 |
| ZNF33A |
| RGMA |
| EXOC8 |
| SMIM8 |
| ADAM9 |
| B3GNT2 |
| CPNE9 |
| NIF3L1 |
| USP53 |
| RXFP1 |
| RRM2B |
| HNRNPLL |
| RSRC1 |
| ADRA2A |
| AP1AR |
| ASF1A |
| CRLF1 |
| NSDHL |
| HNRNPA1L3 |
| ASIC2 |
| JAK3 |
| THOC5 |

| |
|---|
| RNF38 |
| LARP4B |
| ANKFY1 |
| KCTD1 |
| IGFBP2 |
| DXO |
| PPT2 |
| UBR2 |
| MOSPD3 |
| MAP3K9 |
| DHRS7B |
| ZBTB45 |
| ECT2 |
| CRACD |
| ADAMTS2 |
| GTPBP2 |
| CHST12 |
| CXorf38 |
| FNBP1L |
| PPIL3 |
| CDC14B |
| TFCP2 |
| NDST1 |
| SMNDC1 |
| PEX11B |
| PHF23 |
| CLINT1 |
| ZMYM6 |
| RSBN1 |
| SLC6A6 |
| MVK |
| TTPAL |
| BRF2 |
| SRF |
| TMEM120A |
| CD14 |
| ANKRD54 |
| TMEM150C |
| RAVER1 |
| MEGF10 |
| RND3 |
| AGK |
| THAP12 |

| |
|---|
| SPECC1L |
| NETO2 |
| ASS1 |
| DSTYK |
| KIAA1671 |
| TRMT10C |
| CACNA1C |
| RNFT2 |
| SEC24B |
| REV1 |
| DNPEP |
| PDZRN3 |
| ARMH3 |
| YES1 |
| DDX27 |
| BBS5 |
| RAPGEF6 |
| CSGALNACT2 |
| GPN1 |
| GSE1 |
| ICAM2 |
| GALE |
| ZNF770 |
| CCNT2 |
| MSL3 |
| TFAM |
| RELCH |
| TLCD4 |
| CASQ1 |
| PDE4D |
| DDX51 |
| NPRL2 |
| CTNNAL1 |
| DNMT3A |
| SLITRK3 |
| MCM3 |
| IGF2R |
| GTF2E2 |
| TMEM120B |
| CRAMP1 |
| TAF1C |
| PUS1 |
| SLC44A3 |

| |
|---|
| WDR3 |
| IDE |
| TCTN1 |
| PROSER1 |
| CNBD2 |
| SLC35E4 |
| SH3BP1 |
| CA4 |
| ZNF134 |
| ECHDC2 |
| UBXN7 |
| TNXB |
| RET |
| ZNF423 |
| NUDT7 |
| FBXL20 |
| ZC3H12B |
| LNPEP |
| C15orf40 |
| ARHGAP29 |
| GMCL1 |
| LERFS |
| GRIK4 |
| GSTM4 |
| RABEPK |
| NSL1 |
| NDUFAF6 |
| PLA2G5 |
| ABHD14B |
| RFX7 |
| TAF4B |
| RBM15 |
| LINC01106 |
| GUF1 |
| DTWD1 |
| ALG1 |
| TBKBP1 |
| CCDC127 |
| SOHLH1 |
| ESYT1 |
| MTMR2 |
| CKS1B |
| EML6 |

| |
|---|
| ZNF385D |
| SLC35G2 |
| CCDC186 |
| SELENON |
| YPEL4 |
| LRFN5 |
| MAF |
| CDADC1 |
| MRTFA |
| TENM4 |
| BABAM2 |
| GTPBP4 |
| SLC18B1 |
| SLC33A1 |
| CHUK |
| NAF1 |
| CDKAL1 |
| NYAP1 |
| USP3 |
| TRPV2 |
| PCGF2 |
| CEP131 |
| XAB2 |
| HDAC3 |
| ST7 |
| TOX |
| DUSP11 |
| EXOSC8 |
| NUB1 |
| COP1 |
| KDM5B |
| USB1 |
| KREMEN1 |
| CBLL1 |
| LENG1 |
| UTP14A |
| KHK |
| TLCD5 |
| WDTC1 |
| ME3 |
| IFT46 |
| PDCL3 |
| ACADSB |

| |
|---|
| ZDHHC5 |
| STAM2 |
| IL11 |
| ZNF317 |
| MCCC1 |
| SURF6 |
| SLC39A13 |
| FRMD5 |
| OCRL |
| PRKAB2 |
| ZZZ3 |
| RPE |
| WDR55 |
| POLR1H |
| SAMD12 |
| WDR20 |
| CACNA1I |
| HRH3 |
| NTRK3 |
| PLA2G15 |
| SLC9B2 |
| NIPAL2 |
| C2CD5 |
| WDFY2 |
| FLCN |
| CHIC1 |
| NUP42 |
| C12orf43 |
| TRMU |
| CASP3 |
| KPNA5 |
| MAP3K5 |
| PRKRIP1 |
| HELZ |
| SLC25A40 |
| ECD |
| GALNT18 |
| UNC79 |
| GPKOW |
| E4F1 |
| IRAG1 |
| ZBTB43 |
| ROBO3 |

| |
|---|
| DSEL |
| TRAPPC14 |
| KMT2B |
| MRS2 |
| ZNF184 |
| RFX4 |
| FAM210B |
| RASA2 |
| MTHFD2L |
| DUSP4 |
| TRPM2 |
| NUP188 |
| WDR5 |
| MLX |
| ENTREP2 |
| LIX1 |
| CSMD2 |
| BOLA3-DT |
| DCBLD1 |
| RFXAP |
| SMG8 |
| RBM11 |
| KDM6A |
| UTY |
| PTHLH |
| SPRED3 |
| CHAF1B |
| UBLCP1 |
| CROCC |
| TFB2M |
| TTC7A |
| CT75 |
| MIER3 |
| ASAH2B |
| GCH1 |
| UBAP1L |
| INTS2 |
| ASXL3 |
| MTO1 |
| LINC00622 |
| BRINP3 |
| KLHL8 |
| FBXO10 |

| |
|---|
| CKAP2 |
| METTL17 |
| AXIN2 |
| TICAM1 |
| MTG2 |
| MINDY1 |
| COASY |
| ATOSA |
| TRNAU1AP |
| CNNM4 |
| MSTO1 |
| TTC13 |
| EXOG |
| SUCLG2 |
| SNX25 |
| SV2C |
| EFNA5 |
| BMPER |
| KCNK9 |
| ZNF252P |
| MPHOSPH9 |
| TRIOBP |
| TP53I11 |
| ERLIN1 |
| DMAC2 |
| LRFN3 |
| NRBF2 |
| MTHFR |
| AGO4 |
| VPS54 |
| ZFPM2 |
| IPMK |
| ABCD2 |
| RPS6KA5 |
| FAH |
| LRRC37B |
| LINC03048 |
| TMEM91 |
| FKRP |
| SPACA6 |
| HTRA2 |
| WBP1L |
| DHRS11 |

| |
|---|
| MYPOP |
| EDEM3 |
| HAUS2 |
| KMT5B |
| WIPF3 |
| LZTS2 |
| TOB2 |
| KDM7A |
| ASAP3 |
| ELK4 |
| USP42 |
| ZNF395 |
| KYAT1 |
| KLF12 |
| OGFOD3 |
| RBM33 |
| RPAP2 |
| OXTR |
| RAB43 |
| SAP30-DT |
| ZSCAN31 |
| TRIM39 |
| BAALC-AS1 |
| NIPSNAP3B |
| CYB561A3 |
| ASCL1 |
| GTPBP3 |
| ZNF615 |
| ING5 |
| BBOX1 |
| CORO1B |
| TRPM7 |
| MCMBP |
| TRO |
| APIP |
| CCDC28B |
| SH3BP2 |
| SLC30A5 |
| HERPUD2 |
| DNAJC25 |
| FMN1 |
| TGIF1 |
| HMGN4 |

| |
|---|
| FBXO30 |
| NAA25 |
| FGF9 |
| TM6SF1 |
| XRCC1 |
| BCL3 |
| UBOX5 |
| BMP2 |
| PKNOX1 |
| DGCR8 |
| SMG6 |
| GPR85 |
| SLC25A37 |
| C9orf72 |
| PSTK |
| POMT2 |
| HMGXB3 |
| CCDC28A |
| MLXIP |
| NCAPH2 |
| CDKN2C |
| ABT1 |
| KIAA1217 |
| CKAP4 |
| CTU2 |
| ZNF653 |
| ZNF791 |
| ZNF746 |
| RNASEH2B |
| DHRS3 |
| LRRTM2 |
| RCBTB2 |
| ODF2L |
| LGALSL |
| SUCO |
| RBBP5 |
| CHST2 |
| CMBL |
| UBTD1 |
| CENPC |
| KCTD5 |
| TRAPPC8 |
| MSANTD3 |

| |
|---|
| ABHD17C |
| SMYD3 |
| DIMT1 |
| SEC24A |
| CBL |
| TRAF4 |
| AAR2 |
| LRIF1 |
| MCM6 |
| C4orf50 |
| CCDC149 |
| HSPA14 |
| WASHC2C |
| BRD1 |
| ZNF330 |
| DTD2 |
| MBIP |
| LENG8 |
| MORC3 |
| SHANK1 |
| FSTL1 |
| ARAP1 |
| MARK1 |
| ABCB6 |
| PPP3CC |
| ZFAND1 |
| SS18 |
| ZBTB1 |
| WDR35 |
| GTF2IRD2B |
| FAM135B |
| CDC37L1-DT |
| MIR600HG |
| IDI2-AS1 |
| CDK8 |
| DLL4 |
| SOCS1 |
| ERBB2 |
| DCAF15 |
| CCDC9 |
| TAF4 |
| ZNRF3 |
| GABRA3 |

| |
|---|
| FBXO6 |
| PIGV |
| EOGT |
| BOC |
| TET2 |
| PSMB9 |
| AGAP4 |
| POLD3 |
| EMILIN2 |
| ZNF561-AS1 |
| FBH1 |
| NDC1 |
| CNOT11 |
| PAQR9 |
| MAML1 |
| ZC3HC1 |
| KMT2D |
| DNAH10OS |
| CCDC92B |
| ANKRD29 |
| RBFA |
| FLRT3 |
| AZIN2 |
| MAP3K12 |
| RAB9A |
| FUCA1 |
| FLVCR1 |
| LGI2 |
| SH3D19 |
| CCNB1 |
| PDE7A |
| SUSD1 |
| GLE1 |
| AKR1C2 |
| BEST1 |
| BORCS5 |
| ZFP82 |
| DISP3 |
| RNASEH1 |
| ZNF316 |
| MATN2 |
| CCDC181 |
| ARL6IP6 |

| |
|---|
| PARP8 |
| MED23 |
| LOH12CR2 |
| GPC5 |
| ANKDD1A |
| PMM2 |
| SEPTIN1 |
| CDH20 |
| EHD2 |
| MAVS |
| H2AC6 |
| ANXA1 |
| GDPD1 |
| SLC27A3 |
| ZNF354A |
| MASTL |
| FAS |
| RNF214 |
| MSRB3 |
| DEDD |
| ABHD5 |
| F8A1 |
| WAPL |
| GPR108 |
| PRUNE1 |
| RRP9 |
| ILRUN-AS1 |
| HMGCLL1 |
| KCTD21 |
| HMBS |
| ENDOV |
| CACTIN |
| GK |
| LRRC8D |
| RNF149 |
| ATG7 |
| C3orf80 |
| LARP4 |
| GGA3 |
| FTSJ1 |
| FGF17 |
| NOC3L |
| CEP295 |

| |
|---|
| PPP1R37 |
| LRRC42 |
| KMT2E-AS1 |
| ITGA7 |
| PPM1H |
| N4BP1 |
| PCED1A |
| STK4 |
| LSM11 |
| ZNF394 |
| GIGYF1 |
| FSD1L |
| PCGF6 |
| SORCS3 |
| RCE1 |
| JDP2 |
| INTS14 |
| CCDC57 |
| ZNF579 |
| HEPH |
| F8 |
| THBD |
| FUT11 |
| ANGEL2 |
| ZNF513 |
| NFKBIL1 |
| DIS3L |
| PPM1D |
| EIF4ENIF1 |
| ZNF777 |
| GPN2 |
| RNF113A |
| TMEM241 |
| AGO3 |
| EDEM1 |
| LIN54 |
| CDC23 |
| SH3PXD2B |
| IPPK |
| UEVLD |
| ST8SIA1 |
| MBD6 |
| UNG |

| |
|---|
| FBXL12 |
| PEX14 |
| MIER1 |
| SH3RF3 |
| NAA80 |
| NEK11 |
| WDR17 |
| ASCC1 |
| BICD1 |
| ZNF821 |
| ZGPAT |
| SLC25A14 |
| TSPYL5 |
| CCNT1 |
| GAS8 |
| SNIP1 |
| ALG14 |
| EPHB3 |
| PIGZ |
| MTHFD1L |
| DMTF1 |
| LINC01089 |
| ZNF26 |
| ZFP1 |
| ASCC2 |
| EPB41L4B |
| NGLY1 |
| PIGG |
| NET1 |
| MLLT10 |
| EED |
| PAN3 |
| NCOA3 |
| PEX13 |
| TUBE1 |
| SAP130 |
| TMEM39B |
| ZNF544 |
| NUP54 |
| LSM6 |
| STAMBPL1 |
| MID1 |
| CHRNB2 |

| |
|---|
| ZNF875 |
| MOSPD2 |
| TBC1D8 |
| XXYLT1 |
| NRG2 |
| ZNF767P |
| FZD6 |
| ZNF202 |
| ABCC9 |
| MEAK7 |
| GID4 |
| PDYN |
| SFI1 |
| PABPC1L2B |
| RASGRF2 |
| WIPF1 |
| WDPCP |
| MREG |
| LINC01322 |
| WDR19 |
| CMYA5 |
| EFHC1 |
| ZNF487 |
| DPP3 |
| CARMIL3 |
| MAGEL2 |
| HERC2P9 |
| SPAG5 |
| RBBP8 |
| ZNF566 |
| ALDH16A1 |
| ACTR5 |
| HMGN5 |
| DUSP10 |
| ZCCHC3 |
| VARS2 |
| TRRAP |
| PABIR2 |
| GLMN |
| THUMPD2 |
| MDC1 |
| ICA1-AS1 |
| CCDC126 |

| |
|---|
| AGFG2 |
| PRPF18 |
| TIMM23B |
| MTA2 |
| B3GNT4 |
| OPHN1 |
| GPR161 |
| NKX3-1 |
| PARP11 |
| CEP76 |
| ST3GAL3 |
| FAM181A |
| PUDP |
| VEGFA |
| KHDC4 |
| LYSMD4 |
| THEM4 |
| ATG10 |
| FES |
| ASGR1 |
| SIRT6 |
| CEP89 |
| FBXO17 |
| NKX2-2 |
| TENT4B |
| MEGF11 |
| MAP4K2 |
| LINC00882 |
| HACE1 |
| REV3L |
| EIF1AD |
| RABGGTA |
| MOK |
| TNFSF13 |
| GPD1 |
| DCAF1 |
| FRAS1 |
| KDM4C |
| SLC38A7 |
| ERCC2 |
| WDR44 |
| SEMA4A |
| B3GALT2 |

| |
|---|
| CUL9 |
| PABIR1 |
| EMSY |
| PIAS2 |
| INO80D |
| TMEM200C |
| ABCG1 |
| MAP3K20 |
| VASH1 |
| DPYSL5 |
| ZNF436-AS1 |
| NSD1 |
| SYNJ2BP |
| PPP4R1 |
| UBE2T |
| NIM1K |
| SMCR8 |
| TARS2 |
| SANBR |
| LNP1 |
| SORCS2 |
| TLR2 |
| RIOK1 |
| RBM28 |
| WASHC1 |
| MTAP |
| METTL3 |
| MRM3 |
| MIS12 |
| TSEN54 |
| CEP250 |
| NHS |
| NEIL1 |
| FAM91A1 |
| TAB3 |
| PPP1R8 |
| ACAD11 |
| SMARCAD1 |
| FAM221A |
| ACO1 |
| VPS37C |
| TDG |
| HEXD |

TIMM29
CROCCP2
SERTAD4-AS1
LRCH3
COMTD1
R3HCC1L
AMPD3
ANKRD42
MTERF2
SLC9A5
DHX34
TEKT4P2
KRBOX4
RICTOR
SLC25A27
CDK9
MTPAP
CWF19L1
TRIM13
GSPT2
PTPRU
DDR2
C2orf49
MERTK
XRN1
LPP
SLIT2
BNIP1
TTBK1
CYB5R4
TRAF3IP2-AS1
NSMCE2
PHRF1
GDF11
NUDT15
EBPL
PHAF1
SNAI3-AS1
COQ10A
CRISPLD1
SLC5A11
DOC2B
SOCS5

| |
|---|
| MRAP2 |
| EEF1AKMT1 |
| IFIH1 |
| CCDC66 |
| KLF15 |
| SLN |
| SLC38A6 |
| SGSH |
| IZUMO4 |
| RNF150 |
| PNMT |
| LYPD6 |
| ACVR2B |
| CRYZ |
| LINC01719 |
| SYT14 |
| INO80D-AS1 |
| ADRA2C |
| FBXO8 |
| HLA-DQA1 |
| THSD7A |
| IMMP2L |
| MCPH1 |
| CNTRL |
| ZBTB34 |
| FZD8 |
| LIPT2-AS1 |
| USP2-AS1 |
| ARID2 |
| ZWILCH |
| ORAI3 |
| SLC46A1 |
| ZNF44 |
| ZNF628 |
| ZNF776 |
| TAF1 |
| MID2 |
| PPP4R4 |
| TRABD2A |
| AMER3 |
| TAMM41 |
| ABHD18 |
| FASTKD3 |

| |
|---|
| ERMARD |
| WASL-DT |
| ZNF212 |
| MSC |
| TMEM218 |
| ZNF641 |
| RPAP1 |
| HYPK |
| ZNF609 |
| EIF3C |
| MIR762HG |
| NFATC3 |
| DDX19B |
| ADORA2B |
| RAB3D |
| DACH2 |
| GPR137C |
| MS4A7 |
| FAM174B |
| PYDC1 |
| PCDH19 |
| FBXL7 |
| H2BC21 |
| ALYREF |
| PTPN14 |
| GRK4 |
| TRIM52 |
| RTN4IP1 |
| ZNF789 |
| CAAP1 |
| OSBPL5 |
| IRAG2 |
| RACGAP1 |
| ADSS1 |
| MTFMT |
| POLG2 |
| ZNF260 |
| ZNF382 |
| ZNF227 |
| RSPH14 |
| SYN3 |
| SHISA8 |
| ZFY |

| |
|---|
| SMAD3 |
| TCOF1 |
| SLC19A2 |
| TBCE |
| BTD |
| ATP11B |
| CDKL2 |
| TMEM68 |
| NAA35 |
| C18orf21 |
| ARSA |
| ZNF623 |
| WDR43 |
| STPG1 |
| SH3D21 |
| TUFT1 |
| BCL2L11 |
| WDSUB1 |
| CACNA2D2 |
| FAM241A |
| CTSO |
| PEX7 |
| AHR |
| LACTB2 |
| UTP23 |
| PAWR |
| MSI1 |
| ZMYM5 |
| MEG8 |
| MAP3K14 |
| ZNF264 |
| RBMX2 |
| LINC02245 |
| STAT4 |
| LIAS |
| KIT |
| BPHL |
| ADHFE1 |
| PCID2 |
| LYSET |
| CLN6 |
| RSPRY1 |
| ITGB3 |

| |
|---|
| REEP6 |
| RHPN2 |
| MHENCR |
| TYMP |
| EPG5 |
| SLC6A11 |
| NFKBIZ |
| TNFAIP8L1 |
| ZFX |
| ZC3H12A |
| IL17RC |
| SPTSSB |
| PPARGC1B |
| KCNS2 |
| MED30 |
| ARRDC1 |
| WAC-AS1 |
| PYROXD1 |
| SNRNP35 |
| CHAMP1 |
| TMEM229B |
| FBXO22 |
| GLIS2 |
| EMP2 |
| CDH5 |
| CBFA2T3 |
| PDE9A |
| HTR5A |
| STK16 |
| VPS37D |
| LRCH4 |
| FOXP2 |
| PNOC |
| SUOX |
| MTUS2 |
| ADCY9 |
| C16orf46 |
| GYS1 |
| RAB9B |
| METTL13 |
| GPATCH2 |
| SETMAR |
| TMEM41A |

| |
|---|
| MSANTD1 |
| TMEM192 |
| FBXO38 |
| LTC4S |
| ZSCAN26 |
| TBL2 |
| SPACDR |
| ZNF510 |
| CUTC |
| SPA17 |
| VWA8 |
| PHF11 |
| THAP10 |
| PHF12 |
| SNHG30 |
| MAP3K3 |
| C19orf81 |
| ARHGAP10 |
| GPR26 |
| IFT172 |
| MRPS9 |
| CHDH |
| RIOX2 |
| GRAMD1C |
| ELOVL7 |
| PLAGL1 |
| PRKN |
| OSBPL3 |
| RBM48 |
| LSMEM1 |
| ADAMTSL2 |
| LRRK2 |
| SLC35E3 |
| MGA |
| CLUAP1 |
| KPTN |
| TMEM74B |
| AKAP7 |
| SALL2 |
| CDC7 |
| SLX1A |
| KCNJ12 |
| ZNF793 |

| |
|---|
| ZNF320 |
| BATF3 |
| SPON2 |
| ZNF608 |
| WNT7A |
| CTBP1-DT |
| TFR2 |
| ENTPD1 |
| ZNF471 |
| ZNF548 |
| SSTR3 |
| CCDC120 |
| GRM2 |
| NCBP1 |
| NIBAN2 |
| BCL2 |
| RNF32-DT |
| ERI1 |
| AGBL4 |
| INTS7 |
| LINC00342 |
| ANKRD36 |
| KCTD18 |
| TTLL4 |
| FCHO2 |
| HLA-L |
| SRSF12 |
| NAA40 |
| SERPINH1 |
| ZSCAN32 |
| PIGN |
| CCNE1 |
| MAP3K4 |
| C11orf91 |
| FAR2 |
| GMEB1 |
| GDAP2 |
| SOX13 |
| MCEE |
| TMEM184C |
| SPEF2 |
| BLTP3A |
| CEP57L1 |

| |
|---|
| GET4 |
| ANGPT2 |
| HMBOX1 |
| CDKL1 |
| WRAP53 |
| PRELID3A |
| SNAI1 |
| IFNAR2 |
| PCDH11X |
| CCN2 |
| CEP164 |
| WDR77 |
| NOTCH2NLA |
| PIGC |
| SOWAHC |
| TBC1D22B |
| TNPO3 |
| KCTD9 |
| PPP1R3C |
| PAN2 |
| REC8 |
| RCOR1 |
| ZNF48 |
| RFX1 |
| RAI2 |
| ARMCX5-GPRASP2 |
| BCL10 |
| AMY2B |
| PMS1 |
| ANKRD44 |
| DIS3L2 |
| COLQ |
| MCF2L2 |
| KCNN2 |
| DNAAF5 |
| PEX1 |
| PTCD1 |
| METTL2B |
| TCIM |
| LRRCC1 |
| INTS8 |
| POLR3B |
| ATP6V0A2 |

| |
|---|
| SLC30A4 |
| EARS2 |
| FAM182A |
| HMGXB4 |
| PTCHD1 |
| CCDC22 |
| NXPH1 |
| NBPF1 |
| ACVR2A |
| NEK1 |
| AIMP2 |
| PTGS1 |
| ENKUR |
| CPNE8-AS1 |
| SMAD9 |
| TMEM62 |
| LRRK1 |
| BANP |
| NUDT19 |
| GABPA |
| DONSON |
| METTL15 |
| CASP9 |
| PKDCC |
| PLGLB1 |
| LINC01102 |
| ANAPC10 |
| MARCHF11 |
| FAM114A2 |
| RNF144B |
| USP49 |
| MAK16 |
| ZNF34 |
| PUS7L |
| TMEM132C |
| HHIPL1 |
| C17orf114 |
| MAPK7 |
| ZNF519 |
| ZNF585B |
| MRPL39 |
| CPT2 |
| PPM1J |

| |
|---|
| NBPF19 |
| RNASEH1-DT |
| GRASLND |
| GMPPB |
| NOP14 |
| GUSBP1 |
| PCDHB14 |
| KDM1B |
| C6orf47 |
| CAHM |
| GOLGA1 |
| NARS2 |
| AKAP5 |
| TTLL5 |
| MPV17L2 |
| TMEM161A |
| PHKA2 |
| USP35 |
| C1QTNF3 |
| DLX6-AS1 |
| CAVIN1 |
| ACOX3 |
| ABCG2 |
| GLRA3 |
| ATPSCKMT |
| DCC |
| MYT1 |
| TRAF5 |
| ZNF736 |
| FLRT2-AS1 |
| ZNF205 |
| KIAA0753 |
| DCX |
| RNPC3 |
| KCTD7 |
| TMEM186 |
| SPRYD4 |
| TMEM101 |
| NBPF11 |
| TFPI |
| GK5 |
| HAUS3 |
| LINC01511 |

| |
|---|
| PCDHB4 |
| RNF216P1 |
| PPP1R9A-AS1 |
| PUS7 |
| KIAA0586 |
| ZNF23 |
| MC1R |
| MPP3 |
| CDH7 |
| TGFB2 |
| MCC |
| ELP4 |
| POU3F1 |
| TMEM87B |
| METTL14 |
| ABCC10 |
| MMD2 |
| ATAD2 |
| ZNF169 |
| AKNA |
| TBC1D30 |
| LATS2 |
| CIB2 |
| MPV17L |
| KCNJ16 |
| ZBTB14 |
| MEX3D |
| ZNF383 |
| MICALL1 |
| ZNF567 |
| PIK3CD |
| TMEM51 |
| P3H1 |
| ATXN7L2 |
| WNT2B |
| RARB |
| SEL1L3 |
| NAIP |
| BICRAL |
| COPG2 |
| SPAG1 |
| RASSF7 |
| ZNF740 |

| |
|---|
| CEP290 |
| KBTBD7 |
| MRPS31P5 |
| BIVM |
| DHX58 |
| ZNF790-AS1 |
| ZNF540 |
| BICRA |
| TTC28 |
| DNASE1L1 |
| TDRKH |
| USP21 |
| PRKD3 |
| DCTN1-AS1 |
| IL1R1 |
| NAAA |
| SGMS2 |
| SCAMP1-AS1 |
| ZNF251 |
| ZDHHC13 |
| ABCG4 |
| TINCR |
| GEMIN8 |
| FLRT1 |
| PCDH20 |
| C19orf47 |
| SLC38A5 |
| RBKS |
| LMAN2L |
| ECE2 |
| FAM153A |
| ZNF322 |
| GPANK1 |
| RNGTT |
| LAMA2 |
| ZNF398 |
| TNFRSF10D |
| WRN |
| ZNF462 |
| ZNF143 |
| DHRS1 |
| PYGL |
| ASB7 |

| |
|---|
| PAK4 |
| OPA3 |
| PLAGL2 |
| MOCS3 |
| PRRG3 |
| CYS1 |
| AGPAT2 |
| SLC24A4 |
| TOX3 |
| KCNQ4 |
| ST8SIA4 |
| SMIM18 |
| IRF8 |
| RAMP2 |
| PENK |
| DEPTOR |
| ZBTB40 |
| SMC4 |
| TIFA |
| LY6G5B |
| RSPH3 |
| ANKRD26 |
| PANK1 |
| OBI1 |
| LTBP1 |
| COL7A1 |
| FAM86DP |
| TRMT44 |
| ADAM19 |
| DENND2A |
| BMP1 |
| CNTNAP3 |
| CCDC7 |
| TMEM216 |
| LAYN |
| APOA1 |
| NPIPB13 |
| CDC6 |
| ZNF461 |
| TBL1X |
| NHSL2 |

**Data S8. Differe**

| Gene symbol |
|---|
| ABCB9 |
| ABCC8 |
| ABCG2 |
| ABCG4 |
| ADAM9 |
| AEBP2 |
| AGPS |
| ALDH1A3 |
| ANKRD36B |
| APTR |
| AQP11 |
| ARHGAP29 |
| ART3 |
| ASS1 |
| ATAD2 |
| BAIAP2-DT |
| BANP |
| BBC3 |
| BBS10 |
| BCAS4 |
| BRCC3 |
| CACNG8 |
| CAHM |
| CALB2 |
| CAPN15 |
| CCNO |
| CD276 |
| CD93 |
| CDKL1 |
| CEP295 |
| CGNL1 |
| CHRDL1 |
| COPB1 |
| COX15 |
| CUX2 |
| DCAF13 |
| DEPP1 |
| DESI2 |
| DHX32 |

DHX35
DLX1
DMKN
DOK4
DUS3L
ECHDC2
EDEM1
ELF1
EMC1
EPHB3
ESCO1
F8
FABP6
FAM78A
FAM89A
FKTN
GLMP
GPN2
GPR88
GRHL1
GRM7
HSD11B1
INPP5J
IPO4
JAK3
KLHDC8B
LINC01511
LINC01563
LMAN1
LRRC37B
LRRC8C
LYSMD3
LZTS1
MCAT
MCU
METTL25B
MICAL1
MID1
MINDY3
MOG
MPP3
MRPL39
MSH6

MTMR1
MTO1
MYO16
NAA40
NAALAD2
NIPSNAP3B
NLGN4Y
NOP14
NPY
NR2E1
NSUN4
PCBP3
PCMTD1
PDCD2L
PGAP3
PHLDA2
PKNOX2
PLPPR3
PMS2P4
POLR1E
POMZP3
PPTC7
PRKCQ-AS1
PRKD1
PRPF38A
PRR3
PTCH2
PTPN21
RAB28
RAB37
RANBP10
RBKS
RBM24
RFX3
RHNO1
RINT1
RNPEP
RPP25L
RPP38
RXFP1
SCN3A
SEPTIN1
SHANK3

SLC13A5
SLC1A6
SLC25A48
SLC50A1
SLC8A3
SLCO4A1
SMAD7
SNHG16
SOCS1
SPPL2A
STK36
SULF1
TAF6L
TBXAS1
THBD
THBS3-AS1
THSD7A
TMEM128
TMEM161B
TMEM65
TNFAIP8L1
TNFSF13
TNS2-AS1
TRIM3
TRIOBP
TTC28
TTC32
TYW5
UBXN2A
UTP15
VIP
VIPAS39
VPS13B-DT
WAC-AS1
XAF1
ZBTB25
ZEB1-AS1
ZNF10
ZNF135
ZNF252P
ZNF407
ZNF432
ZNF581

ZNF585B
ZNF621
ZNF747
ZXDB

**ntially expressed genes (DEGs) in oligodendrocytes following iTBS.**

| Gene name | EntrezID |
|---|---|
| ATP binding cassette subfamily B member 9 | 23457 |
| ATP binding cassette subfamily C member 8 | 6833 |
| ATP binding cassette subfamily G member 2 (JR blood group) | 9429 |
| ATP binding cassette subfamily G member 4 | 64137 |
| ADAM metallopeptidase domain 9 | 8754 |
| AE binding protein 2 | 121536 |
| alkylglycerone phosphate synthase | 8540 |
| aldehyde dehydrogenase 1 family member A3 | 220 |
| ankyrin repeat domain 36B | 57730 |
| Alu-mediated CDKN1A/p21 transcriptional regulator | 100505854 |
| aquaporin 11 | 282679 |
| Rho GTPase activating protein 29 | 9411 |
| ADP-ribosyltransferase 3 (inactive) | 419 |
| argininosuccinate synthase 1 | 445 |
| ATPase family AAA domain containing 2 | 29028 |
| BAIAP2 divergent transcript | 440465 |
| BTG3 associated nuclear protein | 54971 |
| BCL2 binding component 3 | 27113 |
| Bardet-Biedl syndrome 10 | 79738 |
| breast carcinoma amplified sequence 4 | 55653 |
| BRCA1/BRCA2-containing complex subunit 3 | 79184 |
| calcium voltage-gated channel auxiliary subunit gamma 8 | 59283 |
| colon adenocarcinoma hypermethylated | 100526820 |
| calbindin 2 | 794 |
| calpain 15 | 6650 |
| cyclin O | 10309 |
| CD276 molecule | 80381 |
| CD93 molecule | 22918 |
| cyclin dependent kinase like 1 | 8814 |
| centrosomal protein 295 | 85459 |
| cingulin like 1 | 84952 |
| chordin like 1 | 91851 |
| COPI coat complex subunit beta 1 | 1315 |
| cytochrome c oxidase assembly homolog COX15 | 1355 |
| cut like homeobox 2 | 23316 |
| DDB1 and CUL4 associated factor 13 | 25879 |
| DEPP autophagy regulator 1 | 11067 |
| desumoylating isopeptidase 2 | 51029 |
| DEAH-box helicase 32 (putative) | 55760 |

| | |
|---|---|
| DEAH-box helicase 35 | 60625 |
| distal-less homeobox 1 | 1745 |
| dermokine | 93099 |
| docking protein 4 | 55715 |
| dihydrouridine synthase 3 like | 56931 |
| enoyl-CoA hydratase domain containing 2 | 55268 |
| ER degradation enhancing alpha-mannosidase like protein 1 | 9695 |
| E74 like ETS transcription factor 1 | 1997 |
| ER membrane protein complex subunit 1 | 23065 |
| EPH receptor B3 | 2049 |
| establishment of sister chromatid cohesion N-acetyltransferase 1 | 114799 |
| coagulation factor VIII | 2157 |
| fatty acid binding protein 6 | 2172 |
| family with sequence similarity 78 member A | 286336 |
| family with sequence similarity 89 member A | 375061 |
| fukutin | 2218 |
| glycosylated lysosomal membrane protein | 112770 |
| GPN-loop GTPase 2 | 54707 |
| G protein-coupled receptor 88 | 54112 |
| grainyhead like transcription factor 1 | 29841 |
| glutamate metabotropic receptor 7 | 2917 |
| hydroxysteroid 11-beta dehydrogenase 1 | 3290 |
| inositol polyphosphate-5-phosphatase J | 27124 |
| importin 4 | 79711 |
| Janus kinase 3 | 3718 |
| kelch domain containing 8B | 200942 |
| long intergenic non-protein coding RNA 1511 | 100506791 |
| long intergenic non-protein coding RNA 1563 | 101060544 |
| lectin, mannose binding 1 | 3998 |
| leucine rich repeat containing 37B | 114659 |
| leucine rich repeat containing 8 VRAC subunit C | 84230 |
| LysM domain containing 3 | 116068 |
| leucine zipper tumor suppressor 1 | 11178 |
| malonyl-CoA-acyl carrier protein transacylase | 27349 |
| mitochondrial calcium uniporter | 90550 |
| methyltransferase like 25B | 51093 |
| microtubule associated monooxygenase, calponin and LIM domain containing 1 | 64780 |
| midline 1 | 4281 |
| MINDY lysine 48 deubiquitinase 3 | 80013 |
| myelin oligodendrocyte glycoprotein | 4340 |
| MAGUK p55 scaffold protein 3 | 4356 |
| mitochondrial ribosomal protein L39 | 54148 |
| mutS homolog 6 | 2956 |

| | |
|---|---|
| myotubularin related protein 1 | 8776 |
| mitochondrial tRNA translation optimization 1 | 25821 |
| myosin XVI | 23026 |
| N-alpha-acetyltransferase 40, NatD catalytic subunit | 79829 |
| N-acetylated alpha-linked acidic dipeptidase 2 | 10003 |
| nipsnap homolog 3B | 55335 |
| neuroligin 4 Y-linked | 22829 |
| NOP14 nucleolar protein | 8602 |
| neuropeptide Y | 4852 |
| nuclear receptor subfamily 2 group E member 1 | 7101 |
| NOP2/Sun RNA methyltransferase 4 | 387338 |
| poly(rC) binding protein 3 | 54039 |
| protein-L-isoaspartate (D-aspartate) O-methyltransferase domain containing 1 | 115294 |
| programmed cell death 2 like | 84306 |
| post-GPI attachment to proteins phospholipase 3 | 93210 |
| pleckstrin homology like domain family A member 2 | 7262 |
| PBX/knotted 1 homeobox 2 | 63876 |
| phospholipid phosphatase related 3 | 79948 |
| PMS1 homolog 2, mismatch repair system component pseudogene 4 | 5382 |
| RNA polymerase I subunit E | 64425 |
| POM121 and ZP3 fusion | 22932 |
| protein phosphatase targeting COQ7 | 160760 |
| PRKCQ antisense RNA 1 | 439949 |
| protein kinase D1 | 5587 |
| pre-mRNA processing factor 38A | 84950 |
| proline rich 3 | 80742 |
| patched 2 | 8643 |
| protein tyrosine phosphatase non-receptor type 21 | 11099 |
| RAB28, member RAS oncogene family | 9364 |
| RAB37, member RAS oncogene family | 326624 |
| RAN binding protein 10 | 57610 |
| ribokinase | 64080 |
| RNA binding motif protein 24 | 221662 |
| regulatory factor X3 | 5991 |
| RAD9-HUS1-RAD1 interacting nuclear orphan 1 | 83695 |
| RAD50 interactor 1 | 60561 |
| arginyl aminopeptidase | 6051 |
| ribonuclease P/MRP subunit p25 like | 138716 |
| ribonuclease P/MRP subunit p38 | 10557 |
| relaxin family peptide receptor 1 | 59350 |
| sodium voltage-gated channel alpha subunit 3 | 6328 |
| septin 1 | 1731 |
| SH3 and multiple ankyrin repeat domains 3 | 85358 |

| | |
|---|---|
| solute carrier family 13 member 5 | 284111 |
| solute carrier family 1 member 6 | 6511 |
| solute carrier family 25 member 48 | 153328 |
| solute carrier family 50 member 1 | 55974 |
| solute carrier family 8 member A3 | 6547 |
| solute carrier organic anion transporter family member 4A1 | 28231 |
| SMAD family member 7 | 4092 |
| small nucleolar RNA host gene 16 | 100507246 |
| suppressor of cytokine signaling 1 | 8651 |
| signal peptide peptidase like 2A | 84888 |
| serine/threonine kinase 36 | 27148 |
| sulfatase 1 | 23213 |
| TATA-box binding protein associated factor 6 like | 10629 |
| thromboxane A synthase 1 | 6916 |
| thrombomodulin | 7056 |
| THBS3 antisense RNA 1 | 105371450 |
| thrombospondin type 1 domain containing 7A | 221981 |
| transmembrane protein 128 | 85013 |
| transmembrane protein 161B | 153396 |
| transmembrane protein 65 | 157378 |
| TNF alpha induced protein 8 like 1 | 126282 |
| TNF superfamily member 13 | 8741 |
| TNS2 antisense RNA 1 | 283335 |
| tripartite motif containing 3 | 10612 |
| TRIO and F-actin binding protein | 11078 |
| tetratricopeptide repeat domain 28 | 23331 |
| tetratricopeptide repeat domain 32 | 130502 |
| tRNA-yW synthesizing protein 5 | 129450 |
| UBX domain protein 2A | 165324 |
| UTP15 small subunit processome component | 84135 |
| vasoactive intestinal peptide | 7432 |
| VPS33B interacting protein, apical-basolateral polarity regulator, spe-39 homolog | 63894 |
| VPS13B divergent transcript | 105375666 |
| WAC antisense RNA 1 (head to head) | 220906 |
| XIAP associated factor 1 | 54739 |
| zinc finger and BTB domain containing 25 | 7597 |
| ZEB1 antisense RNA 1 | 220930 |
| zinc finger protein 10 | 7556 |
| zinc finger protein 135 | 7694 |
| zinc finger protein 252, pseudogene | 286101 |
| zinc finger protein 407 | 55628 |
| zinc finger protein 432 | 9668 |
| zinc finger protein 581 | 51545 |

| | |
|---|---|
| zinc finger protein 585B | 92285 |
| zinc finger protein 621 | 285268 |
| zinc finger protein 747 | 65988 |
| zinc finger X-linked duplicated B | 158586 |

| Gene category | Donor 3 | | Don |
|---|---|---|---|
| | log2fc | p.adj | log2fc |
| | 1.81844279 | 1 | 1.812914447 |
| immune response | 1.163090961 | 1 | -1.187085553 |
| | 1.011087867 | 1 | 1.549880042 |
| | 2.44849318 | 1 | 1.075948853 |
| | 1.081477195 | 1 | 1.060841961 |
| | 1.268885625 | 1 | -1.509013647 |
| | 1.049222996 | 1 | 1.549880042 |
| | 2.333015962 | 1 | -1.187085553 |
| | 2.20748508 | 1 | 1.549880042 |
| | 1.163090961 | 1 | 1.272346066 |
| | -2.988912133 | 1 | 1.620269369 |
| | 1.420478804 | 1 | 1.005559525 |
| | 1.654944057 | 1 | 2.035306869 |
| | 1.140370884 | 1 | -1.187085553 |
| | 1.835516303 | 1 | 1.035306869 |
| | 1.1561382 | 1 | 1.260373424 |
| | 1.140370884 | 1 | 1.175484527 |
| | 1.333015962 | 1 | 1.397876948 |
| | 1.654944057 | 1 | -1.509013647 |
| | 1.44849318 | 1 | 1.175484527 |
| cellular differentiation | 1.110623541 | 1 | 1.928391665 |
| synapse | 1.121511857 | 1 | 1.024418553 |
| | 1.333015962 | 1 | 1.191426071 |
| synapse | 3.333015962 | 0.014633191 | 1.583432601 |
| | 1.603105126 | 1 | -1.113084971 |
| | 1.81844279 | 1 | -1.772048053 |
| | 1.152443717 | 1 | 1.134842542 |
| immune response | 2.069981557 | 1 | 1.549880042 |
| | 1.44849318 | 1 | 1.549880042 |
| | 1.502940964 | 1 | 2.620269369 |
| | 1.140370884 | 1 | 1.660911354 |
| | 1.163090961 | 1 | -1.850050565 |
| | 1.181012869 | 0.091109024 | 1.047379701 |
| | 1.333015962 | 1 | 1.343429164 |
| | 2.44849318 | 1 | 1.272346066 |
| | 1.110623541 | 1 | -1.689585893 |
| | 2.140370884 | 1 | 1.005559525 |
| | 1.033455681 | 1 | 1.660911354 |
| | -1.045495661 | 1 | 1.075948853 |

| | | | |
|---|---|---|---|
| | 1.011087867 | 1 | 1.620269369 |
| | 1.835516303 | 1 | 1.660911354 |
| | 1.596050368 | 1 | 1.982839449 |
| | 1.011087867 | 1 | -1.509013647 |
| | 1.773588554 | 1 | 1.175484527 |
| | 1.654944057 | 1 | 1.298341275 |
| | 1.333015962 | 1 | 1.549880042 |
| | 1.485019056 | 1 | 1.982839449 |
| | 1.150151905 | 1 | -1.274548394 |
| cell death | 1.239906558 | 1 | 1.397876948 |
| | 1.654944057 | 1 | 1.075948853 |
| | 1.711527586 | 1 | 1.227951947 |
| | -1.126415656 | 1 | -1.187085553 |
| | 2.502940964 | 1 | -1.357010554 |
| | -1.541453156 | 1 | -1.602123052 |
| | -1.319060734 | 1 | 1.397876948 |
| | -1.319060734 | 1 | 1.812914447 |
| | 1.391909651 | 1 | 1.053922547 |
| | 4.140370884 | 1 | 1.227951947 |
| | 1.917978463 | 1 | 1.134842542 |
| | 2.011087867 | 1 | 1.035306869 |
| | 1.271615418 | 1 | 1.32748762 |
| | 1.20748508 | 1 | -1.125685008 |
| | 1.239906558 | 1 | -2.187085553 |
| cellular differentiation | 2.069981557 | 1 | 1.660911354 |
| | 2.069981557 | 1 | 1.397876948 |
| | 2.333015962 | 1 | -1.994440475 |
| | 1.792447581 | 1 | 1.227951947 |
| | 1.220541233 | 1 | 1.191426071 |
| | 1.856577918 | 1 | -1.357010554 |
| | 1.284106362 | 1 | 1.053922547 |
| | 1.110623541 | 1 | 1.397876948 |
| | 1.695586042 | 1 | 1.093022367 |
| | 1.043509345 | 1 | -1.113084971 |
| | 1.033455681 | 1 | 1.134842542 |
| | -1.444591616 | 1 | 2.620269369 |
| | 1.069981557 | 1 | -1.187085553 |
| | 1.049222996 | 1 | 1.075948853 |
| | 1.214371466 | 1 | -1.357010554 |
| myelin | 1.232489086 | 0.264175053 | 1.028643138 |
| | -1.082021537 | 1 | 1.298341275 |
| mitochondria | 1.711527586 | 1 | 1.812914447 |
| | 1.058840999 | 1 | -1.602123052 |

| | | | |
|---|---|---|---|
| | -1.161748729 | 1 | 2.035306869 |
| mitochondria | 1.011087867 | 1 | 1.298341275 |
| | 1.792447581 | 1 | -1.509013647 |
| | 2.069981557 | 1 | 3.134842542 |
| | 1.917978463 | 1 | 2.035306869 |
| mitochondria | 1.748053462 | 1 | -2.509013647 |
| | 1.181012869 | 1 | 1.620269369 |
| | 1.239906558 | 1 | 1.32748762 |
| myelin | -1.020620992 | 1 | -1.894038578 |
| | 1.333015962 | 1 | -1.646517171 |
| mitochondria | 1.333015962 | 1 | 1.549880042 |
| | 1.654944057 | 1 | -1.994440475 |
| | 1.069981557 | 1 | 1.191426071 |
| | -1.126415656 | 1 | 1.035306869 |
| | 1.44849318 | 1 | -1.187085553 |
| | 1.502940964 | 1 | 1.227951947 |
| | 1.195512439 | 1 | 1.298341275 |
| | -1.251946538 | 1 | 1.053922547 |
| | -1.169484378 | 1 | 1.227951947 |
| | 1.163090961 | 1 | -2.772048053 |
| | 1.098550709 | 1 | -1.602123052 |
| mitochondria | 1.011087867 | 1 | 1.134842542 |
| | 2.069981557 | 1 | -1.409477974 |
| | 1.502940964 | 1 | -1.357010554 |
| | 1.098550709 | 1 | 1.199937571 |
| | 1.711527586 | 1 | 1.134842542 |
| | 1.333015962 | 1 | 1.397876948 |
| | 1.140370884 | 1 | 1.035306869 |
| | 1.16951723 | 1 | 1.075948853 |
| | 1.069981557 | 1 | 1.075948853 |
| | 1.695586042 | 1 | -1.187085553 |
| | 1.470519486 | 1 | 1.175484527 |
| | 1.333015962 | 1 | 1.397876948 |
| | 2.140370884 | 1 | 1.134842542 |
| | 1.917978463 | 1 | 1.397876948 |
| | 1.333015962 | 1 | 1.075948853 |
| | 1.502940964 | 1 | 2.397876948 |
| | 1.44849318 | 1 | 1.343429164 |
| | 2.917978463 | 1 | -1.509013647 |
| | -2.666984038 | 1 | 1.075948853 |
| intrinsic plasticity | 1.226100758 | 1 | -1.093976148 |
| | 1.069981557 | 1 | 1.227951947 |
| synapse | 1.470519486 | 1 | 1.982839449 |

| | | | |
|---|---|---|---|
| | 1.181012869 | 1 | 1.490986353 |
| synapse | 1.333015962 | 1 | 1.549880042 |
| mitochondria | 1.069981557 | 1 | 1.397876948 |
| | 1.44849318 | 1 | -1.061554671 |
| intrinsic plasticity | 1.250553802 | 1 | 1.075948853 |
| | 1.098550709 | 1 | -2.772048053 |
| myelin/ cellular differentiation | 1.555408384 | 1 | 1.175484527 |
| | 1.121511857 | 1 | -1.028387807 |
| | 1.140370884 | 1 | 1.982839449 |
| | 1.190996957 | 1 | 1.035306869 |
| | 1.110623541 | 1 | 1.397876948 |
| myelin/ cellular differentiation | -2.05930146 | 1 | 1.549880042 |
| | 1.470519486 | 1 | 1.005559525 |
| | 1.333015962 | 1 | -2.509013647 |
| | 1.20748508 | 1 | 1.298341275 |
| | 1.333015962 | 1 | 1.035306869 |
| | 1.654944057 | 1 | 2.134842542 |
| | 1.333015962 | 1 | 1.175484527 |
| | 1.110623541 | 1 | 1.620269369 |
| mitochondria | 1.479857351 | 1 | 1.053922547 |
| | -1.251946538 | 1 | 1.464991144 |
| | 1.333015962 | 1 | 1.075948853 |
| | 1.470519486 | 1 | 1.298341275 |
| | 1.087903465 | 1 | 1.060841961 |
| | 1.043509345 | 1 | 1.005559525 |
| | -1.782461255 | 1 | 1.812914447 |
| | 1.250553802 | 1 | 1.134842542 |
| | 1.011087867 | 1 | 2.397876948 |
| | -1.045495661 | 1 | -2.357010554 |
| | 1.470519486 | 1 | 1.134842542 |
| | 1.333015962 | 1 | 1.035306869 |
| | 1.792447581 | 1 | -1.187085553 |
| | 1.1561382 | 1 | 1.134842542 |
| | -1.107556629 | 1 | -1.093976148 |
| cell death | 1.226100758 | 1 | -1.509013647 |
| | 2.580943476 | 1 | 1.549880042 |
| | 1.011087867 | 1 | 1.397876948 |
| | 1.502940964 | 1 | 1.102421065 |
| | 1.069981557 | 1 | -2.887525271 |
| | -1.152410865 | 1 | 1.134842542 |
| | 1.181012869 | 1 | 1.812914447 |
| | 1.011087867 | 1 | 1.812914447 |
| | 1.110623541 | 1 | -1.187085553 |

| | | |
|---|---|---|
| 2.333015962 | 1 | 1.490986353 |
| 1.3971463 | 1 | -1.187085553 |
| 1.917978463 | 1 | 1.298341275 |
| 1.654944057 | 1 | 1.227951947 |

or 4

| p.adj |
| --- |
| 1 |
| 1 |
| 1 |
| 1 |
| 1 |
| 1 |
| 1 |
| 1 |
| 1 |
| 1 |
| 1 |
| 1 |
| 1 |
| 1 |
| 1 |
| 1 |
| 1 |
| 1 |
| 1 |
| 1 |
| 1 |
| 1 |
| 1 |
| 1 |
| 1 |
| 1 |
| 1 |
| 1 |
| 1 |
| 1 |
| 1 |
| 1 |
| 1 |
| 1 |
| 1 |
| 1 |
| 1 |
| 1 |
| 1 |

1
1
1
1
1
1
1
1
1
1
1
1
1
1
1
1
1
1
1
1
1
1
1
1
1
1
1
1
1
1
1
1
1
1
1
1
1
1
1
1
1
1
1

1
1
1
1

**Data S9. Differentially expressed genes (DEGs) in astrocytes following iTBS.**

| Gene symbol | Gene name |
|---|---|
| ADORA2B | adenosine A2b receptor |
| AGA | aspartylglucosaminidase |
| AP3M1 | adaptor related protein complex 3 subunit mu 1 |
| ARL15 | ARF like GTPase 15 |
| ATP10B | ATPase phospholipid transporting 10B (putative) |
| BACE2 | beta-secretase 2 |
| BMP1 | bone morphogenetic protein 1 |
| C1GALT1C1 | C1GALT1 specific chaperone 1 |
| CA7 | carbonic anhydrase 7 |
| CACNG2 | calcium voltage-gated channel auxiliary subunit gamma 2 |
| CALCRL | calcitonin receptor like receptor |
| CALD1 | caldesmon 1 |
| CCL4L2 | C-C motif chemokine ligand 4 like 2 |
| CD86 | CD86 molecule |
| CDH6 | cadherin 6 |
| CDK11B | cyclin dependent kinase 11B |
| CHAMP1 | chromosome alignment maintaining phosphoprotein 1 |
| COX10 | cytochrome c oxidase assembly factor heme A:farnesyltransferase COX10 |
| CRB1 | crumbs cell polarity complex component 1 |
| CRYBG3 | crystallin beta-gamma domain containing 3 |
| CTC1 | CST telomere replication complex component 1 |
| CTU2 | cytosolic thiouridylase subunit 2 |
| CXCL8 | C-X-C motif chemokine ligand 8 |
| DCAF15 | DDB1 and CUL4 associated factor 15 |
| DCPS | decapping enzyme, scavenger |
| DDI2 | DNA damage inducible 1 homolog 2 |
| DESI2 | desumoylating isopeptidase 2 |
| DNAJC1 | DnaJ heat shock protein family (Hsp40) member C1 |
| DNAL4 | dynein axonemal light chain 4 |
| DNMBP | dynamin binding protein |
| DYNC2I2 | dynein 2 intermediate chain 2 |
| DYSF | dysferlin |
| EBF4 | EBF family member 4 |
| EEF1AKMT2 | EEF1A lysine methyltransferase 2 |
| EEFSEC | eukaryotic elongation factor, selenocysteine-tRNA specific |
| EGFL8 | EGF like domain multiple 8 |
| EIF1AD | eukaryotic translation initiation factor 1A domain containing |
| FAM161A | FAM161 centrosomal protein A |
| FCSK | fucose kinase |

| | |
|---|---|
| FNDC3B | fibronectin type III domain containing 3B |
| GABPA | GA binding protein transcription factor subunit alpha |
| GLIDR | glioblastoma down-regulated RNA |
| GPRASP3 | G protein-coupled receptor associated sorting protein family member 3 |
| GREM2 | gremlin 2, DAN family BMP antagonist |
| HCG11 | HLA complex group 11 |
| HERC6 | HECT and RLD domain containing E3 ubiquitin protein ligase family member 6 |
| HYPK | huntingtin interacting protein K |
| ICA1-AS1 | ICA1 antisense RNA 1 |
| IL1A | interleukin 1 alpha |
| IL4R | interleukin 4 receptor |
| KCNN2 | potassium calcium-activated channel subfamily N member 2 |
| L2HGDH | L-2-hydroxyglutarate dehydrogenase |
| LGR4 | leucine rich repeat containing G protein-coupled receptor 4 |
| LINC00951 | long intergenic non-protein coding RNA 951 |
| LINC01089 | long intergenic non-protein coding RNA 1089 |
| LINC01122 | long intergenic non-protein coding RNA 1122 |
| LINC03025 | long intergenic non-protein coding RNA 3025 |
| LRMDA | leucine rich melanocyte differentiation associated |
| LY6G5B | lymphocyte antigen 6 family member G5B |
| MCU | mitochondrial calcium uniporter |
| METTL25B | methyltransferase like 25B |
| MMACHC | metabolism of cobalamin associated C |
| MOGS | mannosyl-oligosaccharide glucosidase |
| MPHOSPH9 | M-phase phosphoprotein 9 |
| MRPL39 | mitochondrial ribosomal protein L39 |
| MRPS31P5 | mitochondrial ribosomal protein S31 pseudogene 5 |
| MSRB3 | methionine sulfoxide reductase B3 |
| MT1H | metallothionein 1H |
| MTERF3 | mitochondrial transcription termination factor 3 |
| MYCT1 | MYC target 1 |
| NACAD | NAC alpha domain containing |
| NIFK-AS1 | NIFK antisense RNA 1 |
| NOP14 | NOP14 nucleolar protein |
| NOPCHAP1 | NOP protein chaperone 1 |
| NSUN5 | NOP2/Sun RNA methyltransferase 5 |
| NUDT15 | nudix hydrolase 15 |
| NUP155 | nucleoporin 155 |
| PACSIN3 | protein kinase C and casein kinase substrate in neurons 3 |
| PBX2 | PBX homeobox 2 |
| PCGF2 | polycomb group ring finger 2 |
| PENK | proenkephalin |
| PFAS | phosphoribosylformylglycinamidine synthase |

| | |
|---|---|
| PHACTR4 | phosphatase and actin regulator 4 |
| PIK3CD | phosphatidylinositol-4,5-bisphosphate 3-kinase catalytic subunit delta |
| PLEK | pleckstrin |
| PLEKHH3 | pleckstrin homology, MyTH4 and FERM domain containing H3 |
| PLOD3 | procollagen-lysine,2-oxoglutarate 5-dioxygenase 3 |
| PLXDC1 | plexin domain containing 1 |
| PNOC | prepronociceptin |
| PRELP | proline and arginine rich end leucine rich repeat protein |
| PRKCH | protein kinase C eta |
| PYDC2-AS1 | PYDC2 antisense RNA 1 |
| RAB35-AS1 | RAB35 antisense RNA 1 |
| RAB7B | RAB7B, member RAS oncogene family |
| RASGRP3 | RAS guanyl releasing protein 3 |
| RIGI | RNA sensor RIG-I |
| RSPH3 | radial spoke head 3 |
| SCLT1 | sodium channel and clathrin linker 1 |
| SCPEP1 | serine carboxypeptidase 1 |
| SEC22A | SEC22 homolog A, vesicle trafficking protein |
| SETD6 | SET domain containing 6, protein lysine methyltransferase |
| SHC4 | SHC adaptor protein 4 |
| SLC12A8 | solute carrier family 12 member 8 |
| SLC25A19 | solute carrier family 25 member 19 |
| SLC52A2 | solute carrier family 52 member 2 |
| SMKR1 | small lysine rich protein 1 |
| SORD | sorbitol dehydrogenase |
| SPRTN | SprT-like N-terminal domain |
| STX17 | syntaxin 17 |
| TCHP | trichoplein keratin filament binding |
| THUMPD2 | THUMP domain 2 tRNA and snRNA guanosine methyltransferase |
| TMEM168 | transmembrane protein 168 |
| TMEM19 | transmembrane protein 19 |
| TMEM216 | transmembrane protein 216 |
| TMEM234 | transmembrane protein 234 |
| TMEM254 | transmembrane protein 254 |
| TMEM62 | transmembrane protein 62 |
| TPM2 | tropomyosin 2 |
| TRIM25 | tripartite motif containing 25 |
| TRIM62 | tripartite motif containing 62 |
| TRPV2 | transient receptor potential cation channel subfamily V member 2 |
| TTC32 | tetratricopeptide repeat domain 32 |
| TTC38 | tetratricopeptide repeat domain 38 |
| UGGT2 | UDP-glucose glycoprotein glucosyltransferase 2 |
| USP37 | ubiquitin specific peptidase 37 |

| | |
|---|---|
| XXYLT1 | xyloside xylosyltransferase 1 |
| ZBED3 | zinc finger BED-type containing 3 |
| ZFYVE26 | zinc finger FYVE-type containing 26 |
| ZNF135 | zinc finger protein 135 |
| ZNF224 | zinc finger protein 224 |
| ZNF236 | zinc finger protein 236 |
| ZNF252P | zinc finger protein 252, pseudogene |
| ZNF3 | zinc finger protein 3 |
| ZNF367 | zinc finger protein 367 |
| ZNF383 | zinc finger protein 383 |
| ZNF507 | zinc finger protein 507 |
| ZNF510 | zinc finger protein 510 |
| ZNF554 | zinc finger protein 554 |
| ZNF626 | zinc finger protein 626 |
| ZNF677 | zinc finger protein 677 |
| ZNF790-AS1 | ZNF790 antisense RNA 1 |
| ZNF793 | zinc finger protein 793 |
| ZWILCH | zwilch kinetochore protein |

| EntrezID | Gene category | Donor 1 | |
|---|---|---|---|
| | | log2fc | p.adj |
| 136 | | -1.393610941 | 1 |
| 175 | | -2.075434981 | 1 |
| 26985 | | -1.022967562 | 1 |
| 54622 | | 1.468885535 | 1 |
| 23120 | | -1.367615733 | 1 |
| 25825 | | -1.630650139 | 1 |
| 649 | | -1.85304256 | 1 |
| 29071 | | 1.468885535 | 1 |
| 766 | | 1.731919941 | 1 |
| 10369 | | -1.005045654 | 1 |
| 10203 | | -1.268080059 | 1 |
| 800 | | 1.670519396 | 1 |
| 9560 | | 1.595715557 | 0.044422563 |
| 942 | | 1.14695744 | 1 |
| 1004 | | -1.683117559 | 1 |
| 984 | | -1.85304256 | 1 |
| 283489 | | -1.116076966 | 1 |
| 1352 | | -1.116076966 | 1 |
| 23418 | | 1.606389059 | 1 |
| 131544 | | -1.116076966 | 1 |
| 80169 | | -1.174970655 | 1 |
| 348180 | | -1.005045654 | 1 |
| 3576 | immune response | 1.439739189 | 1 |
| 90379 | | 1.14695744 | 1 |
| 28960 | | 1.14695744 | 1 |
| 84301 | | -2.531114465 | 1 |
| 51029 | | 1.316882441 | 1 |
| 64215 | | -1.045687638 | 1 |
| 10126 | | -1.512005642 | 1 |
| 23268 | | 1.14695744 | 1 |
| 89891 | | -1.116076966 | 1 |
| 8291 | | 1.731919941 | 1 |
| 57593 | | -1.005045654 | 1 |
| 399818 | | 2.053848036 | 1 |
| 60678 | | 1.316882441 | 1 |
| 80864 | | -1.438005061 | 1 |
| 84285 | | 1.539274863 | 1 |
| 84140 | | -1.438005061 | 1 |
| 197258 | | -1.367615733 | 1 |

| | | | |
|---|---|---|---|
| 64778 | | -1.045687638 | 1 |
| 2551 | | -1.438005061 | 1 |
| 389741 | | -1.005045654 | 1 |
| 80823 | | -1.630650139 | 1 |
| 64388 | | 1.731919941 | 1 |
| 493812 | | -1.116076966 | 1 |
| 55008 | | 1.731919941 | 1 |
| 25764 | | -1.531114465 | 1 |
| 100505938 | | -1.590008154 | 1 |
| 3552 | immune response | 2.053848036 | 1 |
| 3566 | immune response/ neurotrophin | 1.670519396 | 1 |
| 3781 | instrinsic plasticity | -1.152602842 | 1 |
| 79944 | | 1.731919941 | 1 |
| 55366 | | -1.85304256 | 1 |
| 401260 | | -2.590008154 | 1 |
| 338799 | | 2.606389059 | 1 |
| 400955 | | -1.531114465 | 1 |
| 440896 | | -1.116076966 | 1 |
| 83938 | | -2.590008154 | 1 |
| 58496 | | 1.14695744 | 1 |
| 90550 | mitochondria | -1.268080059 | 1 |
| 51093 | | 1.847397158 | 1 |
| 25974 | | -1.85304256 | 1 |
| 7841 | | -1.85304256 | 1 |
| 10198 | | -1.200965864 | 1 |
| 54148 | mitochondria | -1.005045654 | 1 |
| 100887750 | mitochondria | -1.438005061 | 1 |
| 253827 | | -1.531114465 | 1 |
| 4496 | immune response | -1.531114465 | 1 |
| 51001 | mitochondria | -1.85304256 | 1 |
| 80177 | | -2.268080059 | 1 |
| 23148 | | 1.110431564 | 1 |
| 254128 | | -1.85304256 | 1 |
| 8602 | | -1.85304256 | 1 |
| 121053 | | 1.847397158 | 1 |
| 55695 | | -1.005045654 | 1 |
| 55270 | | -1.322527843 | 1 |
| 9631 | | -1.045687638 | 1 |
| 29763 | | 1.053848036 | 1 |
| 5089 | | -1.338469387 | 1 |
| 7703 | | -1.152602842 | 1 |
| 5179 | | -1.660397482 | 1 |
| 5198 | | -1.045687638 | 1 |

| | | | |
|---|---|---|---|
| 65979 | | -1.683117559 | 1 |
| 5293 | | -1.116076966 | 1 |
| 5341 | | 1.234420281 | 1 |
| 79990 | | -1.045687638 | 1 |
| 8985 | | -1.152602842 | 1 |
| 57125 | | 1.394884953 | 1 |
| 5368 | | -1.305554765 | 1 |
| 5549 | | -1.85304256 | 1 |
| 5583 | | -1.045687638 | 1 |
| 103344926 | | 1.14695744 | 1 |
| 127138862 | | -1.268080059 | 1 |
| 338382 | | 1.205851129 | 1 |
| 25780 | | -1.85304256 | 1 |
| 23586 | | -1.200965864 | 1 |
| 83861 | | -1.116076966 | 1 |
| 132320 | | 2.14695744 | 1 |
| 59342 | | 1.561994939 | 1 |
| 26984 | | -1.490472481 | 1 |
| 79918 | | 1.468885535 | 1 |
| 399694 | | -2.268080059 | 1 |
| 84561 | | -1.85304256 | 1 |
| 60386 | mitochondria | -1.116076966 | 1 |
| 79581 | | 1.316882441 | 1 |
| 100287482 | | -1.367615733 | 1 |
| 6652 | | -1.174970655 | 1 |
| 83932 | | -1.630650139 | 1 |
| 55014 | | 1.394884953 | 1 |
| 84260 | | 1.14695744 | 1 |
| 80745 | | -1.116076966 | 1 |
| 64418 | | -1.045687638 | 1 |
| 55266 | | -1.045687638 | 1 |
| 51259 | | -1.116076966 | 1 |
| 56063 | | 1.468885535 | 1 |
| 80195 | | -1.531114465 | 1 |
| 80021 | | 1.14695744 | 1 |
| 7169 | | -1.715539036 | 1 |
| 7706 | | 1.085556895 | 1 |
| 55223 | | -1.045687638 | 1 |
| 51393 | cell survival | 1.14695744 | 1 |
| 130502 | | -1.531114465 | 1 |
| 55020 | | 1.606389059 | 1 |
| 55757 | | -1.660397482 | 1 |
| 57695 | | 1.053848036 | 1 |

| | | |
|---|---|---|
| 152002 | -1.268080059 | 1 |
| 84327 | -2.116076966 | 1 |
| 23503 | -1.85304256 | 1 |
| 7694 | -1.116076966 | 1 |
| 7767 | -1.85304256 | 1 |
| 7776 | -1.174970655 | 1 |
| 286101 | -1.116076966 | 1 |
| 7551 | 1.468885535 | 1 |
| 195828 | 1.14695744 | 1 |
| 163087 | 1.468885535 | 1 |
| 22847 | -1.268080059 | 1 |
| 22869 | 1.731919941 | 1 |
| 115196 | -1.045687638 | 1 |
| 199777 | -1.174970655 | 1 |
| 342926 | -1.590008154 | 1 |
| 284408 | -1.174970655 | 1 |
| 390927 | 1.606389059 | 1 |
| 55055 | -1.116076966 | 1 |

| Donor 4 | |
| --- | --- |
| log2fc | p.adj |
| 1.162271429 | 1 |
| 1.647698256 | 1 |
| -1.044179449 | 1 |
| 2.647698256 | 1 |
| -1.159656666 | 1 |
| -1.937264245 | 1 |
| -1.767339243 | 1 |
| 2.647698256 | 1 |
| -1.004378441 | 1 |
| 1.107129875 | 1 |
| 1.38466385 | 1 |
| 1.522167374 | 1 |
| 1.005808687 | 0.000953168 |
| 1.817623258 | 1 |
| 1.107129875 | 1 |
| 1.232660757 | 1 |
| -1.522226745 | 1 |
| -1.522226745 | 1 |
| 1.133125083 | 1 |
| 1.455053178 | 1 |
| 1.554588852 | 1 |
| -1.522226745 | 1 |
| 1.254355828 | 0.01426372 |
| 1.817623258 | 1 |
| -1.20029865 | 1 |
| -1.811733363 | 1 |
| 1.969626351 | 1 |
| -1.767339243 | 1 |
| 1.095157233 | 1 |
| -1.20029865 | 1 |
| 1.107129875 | 1 |
| -1.574694165 | 1 |
| -1.20029865 | 1 |
| -1.352301744 | 1 |
| 1.150198597 | 1 |
| -1.052741462 | 1 |
| -1.226770862 | 1 |
| -1.574694165 | 1 |
| -1.489805268 | 1 |

| | |
|---|---|
| -1.352301744 | 1 |
| 1.026209879 | 1 |
| -1.352301744 | 1 |
| -1.674229839 | 1 |
| 2.455053178 | 1 |
| 1.232660757 | 1 |
| 1.010268335 | 1 |
| 1.969626351 | 1 |
| 1.232660757 | 1 |
| 1.455053178 | 1 |
| 1.348137974 | 1 |
| 1.647698256 | 1 |
| 1.325770161 | 1 |
| -1.25276607 | 1 |
| -1.767339243 | 1 |
| -2.352301744 | 1 |
| 1.232660757 | 1 |
| -1.352301744 | 1 |
| 1.38466385 | 1 |
| -1.352301744 | 1 |
| -1.439764585 | 1 |
| 1.232660757 | 1 |
| 1.107129875 | 1 |
| 1.455053178 | 1 |
| -1.352301744 | 1 |
| 1.232660757 | 1 |
| -1.352301744 | 1 |
| 1.522167374 | 1 |
| 1.969626351 | 1 |
| -1.030373649 | 1 |
| -1.352301744 | 1 |
| -1.058570541 | 1 |
| -1.352301744 | 1 |
| -1.052741462 | 1 |
| 1.183751156 | 1 |
| -1.089267338 | 1 |
| -1.352301744 | 1 |
| 1.062735755 | 1 |
| -1.20029865 | 1 |
| -1.159656666 | 1 |
| -1.352301744 | 1 |
| -1.352301744 | 1 |
| -1.159656666 | 1 |

| | |
|---|---|
| -1.574694165 | 1 |
| -1.030373649 | 1 |
| 1.232660757 | 1 |
| -1.489805268 | 1 |
| 1.817623258 | 1 |
| -1.25919234 | 1 |
| -1.004378441 | 1 |
| 1.232660757 | 1 |
| 1.647698256 | 1 |
| -1.089267338 | 1 |
| -1.674229839 | 1 |
| 1.062735755 | 1 |
| 1.162271429 | 1 |
| -1.811733363 | 1 |
| 1.232660757 | 1 |
| 1.026209879 | 1 |
| 3.455053178 | 0.375187112 |
| 1.647698256 | 1 |
| -1.674229839 | 1 |
| -1.226770862 | 1 |
| -1.467778961 | 1 |
| -1.937264245 | 1 |
| -1.21479822 | 1 |
| -1.522226745 | 1 |
| -1.20029865 | 1 |
| 1.062735755 | 1 |
| 1.425305835 | 1 |
| 1.232660757 | 1 |
| 1.062735755 | 1 |
| 1.232660757 | 1 |
| 1.121629444 | 1 |
| 1.969626351 | 1 |
| -1.352301744 | 1 |
| -1.030373649 | 1 |
| -1.004378441 | 1 |
| 1.413233002 | 1 |
| -1.11783649 | 1 |
| 1.107129875 | 1 |
| -1.030373649 | 1 |
| -1.574694165 | 1 |
| 1.107129875 | 1 |
| 1.026209879 | 1 |
| -1.052741462 | 1 |

| | |
|---|---|
| -1.352301744 | 1 |
| 1.870090677 | 1 |
| -1.767339243 | 1 |
| -1.226770862 | 1 |
| 1.817623258 | 1 |
| -1.352301744 | 1 |
| -1.226770862 | 1 |
| 1.232660757 | 1 |
| 1.232660757 | 1 |
| 1.062735755 | 1 |
| 1.232660757 | 1 |
| -1.226770862 | 1 |
| -1.574694165 | 1 |
| 1.026209879 | 1 |
| -1.937264245 | 1 |
| -1.159656666 | 1 |
| -1.159656666 | 1 |
| -1.20029865 | 1 |

**Data S10. Differentially expressed genes (DEGs) in excitatory neurons following 10 Hz rTM**

| Gene symbol | Gene name | EntrezID |
|---|---|---|
| ADAMTS10 | ADAM metallopeptidase with thrombospondin type 1 motif 10 | 81794 |
| ADSS1 | adenylosuccinate synthase 1 | 122622 |
| AGO1 | argonaute RISC component 1 | 26523 |
| AMH | anti-Mullerian hormone | 268 |
| AMOT | angiomotin | 154796 |
| ANK1 | ankyrin 1 | 286 |
| ARHGAP10 | Rho GTPase activating protein 10 | 79658 |
| ARHGEF26 | Rho guanine nucleotide exchange factor 26 | 26084 |
| ASCC3 | activating signal cointegrator 1 complex subunit 3 | 10973 |
| ATF7IP | activating transcription factor 7 interacting protein | 55729 |
| ATP10B | ATPase phospholipid transporting 10B (putative) | 23120 |
| BACE2 | beta-secretase 2 | 25825 |
| BDNF-AS | BDNF antisense RNA | 497258 |
| BIRC3 | baculoviral IAP repeat containing 3 | 330 |
| BMP2 | bone morphogenetic protein 2 | 650 |
| BRWD3 | bromodomain and WD repeat domain containing 3 | 254065 |
| C17orf75 | chromosome 17 open reading frame 75 | 64149 |
| C1QL2 | complement C1q like 2 | 165257 |
| CA13 | carbonic anhydrase 13 | 377677 |
| CCDC134 | coiled-coil domain containing 134 | 79879 |
| CDK5RAP1 | CDK5RAP1 mitochondrial tRNA methylthiotransferase | 51654 |
| CHRNB1 | cholinergic receptor nicotinic beta 1 subunit | 1140 |
| CMTM5 | CKLF like MARVEL transmembrane domain containing 5 | 116173 |
| CNTN5 | contactin 5 | 53942 |
| COL5A2 | collagen type V alpha 2 chain | 1290 |
| COQ7 | coenzyme Q7, hydroxylase | 10229 |
| CPQ | carboxypeptidase Q | 10404 |
| CRACD | capping protein inhibiting regulator of actin dynamics | 57482 |
| CREB3L2 | cAMP responsive element binding protein 3 like 2 | 64764 |
| CROCC | ciliary rootlet coiled-coil, rootletin | 9696 |
| CSF1R | colony stimulating factor 1 receptor | 1436 |
| CTSC | cathepsin C | 1075 |
| CUX2 | cut like homeobox 2 | 23316 |
| CYFIP1 | cytoplasmic FMR1 interacting protein 1 | 23191 |
| DBF4 | DBF4-CDC7 kinase regulatory subunit | 10926 |
| DGCR8 | DGCR8 microprocessor complex subunit | 54487 |
| DHTKD1 | dehydrogenase E1 and transketolase domain containing 1 | 55526 |
| DICER1 | dicer 1, ribonuclease III | 23405 |
| DISP1 | dispatched RND transporter family member 1 | 84976 |

| | | |
|---|---|---|
| DNAJC1 | DnaJ heat shock protein family (Hsp40) member C1 | 64215 |
| DUS3L | dihydrouridine synthase 3 like | 56931 |
| EGFL8 | EGF like domain multiple 8 | 80864 |
| EHD2 | EH domain containing 2 | 30846 |
| EMX2 | empty spiracles homeobox 2 | 2018 |
| ERAP1 | endoplasmic reticulum aminopeptidase 1 | 51752 |
| ERBB3 | erb-b2 receptor tyrosine kinase 3 | 2065 |
| FBLN5 | fibulin 5 | 10516 |
| FHL3 | four and a half LIM domains 3 | 2275 |
| FOXO4 | forkhead box O4 | 4303 |
| FRZB | frizzled related protein | 2487 |
| FZD6 | frizzled class receptor 6 | 8323 |
| G2E3 | G2/M-phase specific E3 ubiquitin protein ligase | 55632 |
| GABPB1 | GA binding protein transcription factor subunit beta 1 | 2553 |
| GBP1 | guanylate binding protein 1 | 2633 |
| GFRA1 | GDNF family receptor alpha 1 | 2674 |
| GLE1 | GLE1 RNA export mediator | 2733 |
| GMPPA | GDP-mannose pyrophosphorylase A | 29926 |
| GPAT3 | glycerol-3-phosphate acyltransferase 3 | 84803 |
| GPN3 | GPN-loop GTPase 3 | 51184 |
| GPR17 | G protein-coupled receptor 17 | 2840 |
| GPRC5A | G protein-coupled receptor class C group 5 member A | 9052 |
| GUCD1 | guanylyl cyclase domain containing 1 | 83606 |
| HBB | hemoglobin subunit beta | 3043 |
| HCLS1 | hematopoietic cell-specific Lyn substrate 1 | 3059 |
| HELQ | helicase, POLQ like | 113510 |
| HILPDA | hypoxia inducible lipid droplet associated | 29923 |
| HPS4 | HPS4 biogenesis of lysosomal organelles complex 3 subunit 2 | 89781 |
| HS1BP3 | HCLS1 binding protein 3 | 64342 |
| IFI44 | interferon induced protein 44 | 10561 |
| IFIT5 | interferon induced protein with tetratricopeptide repeats 5 | 24138 |
| IGSF3 | immunoglobulin superfamily member 3 | 3321 |
| IL1RAP | interleukin 1 receptor accessory protein | 3556 |
| INTS12 | integrator complex subunit 12 | 57117 |
| INTS5 | integrator complex subunit 5 | 80789 |
| INTS6L | integrator complex subunit 6 like | 203522 |
| IQCE | IQ motif containing E | 23288 |
| ISYNA1 | inositol-3-phosphate synthase 1 | 51477 |
| JPT2 | Jupiter microtubule associated homolog 2 | 90861 |
| JUP | junction plakoglobin | 3728 |
| KCTD18 | potassium channel tetramerization domain containing 18 | 130535 |
| KCTD5 | potassium channel tetramerization domain containing 5 | 54442 |
| KRT17 | keratin 17 | 3872 |

| | | |
|---|---|---|
| LAMP5 | lysosomal associated membrane protein family member 5 | 24141 |
| LCLAT1 | lysocardiolipin acyltransferase 1 | 253558 |
| LIMA1 | LIM domain and actin binding 1 | 51474 |
| LINC00507 | long intergenic non-protein coding RNA 507 | 100862680 |
| LINC01007 | long intergenic non-protein coding RNA 1007 | 100506527 |
| LINC01896 | long intergenic non-protein coding RNA 1896 | 645321 |
| LINC02668 | long intergenic non-protein coding RNA 2668 | 105376354 |
| LINC03013 | long intergenic non-protein coding RNA 3013 | 100506725 |
| LNX2 | ligand of numb-protein X 2 | 222484 |
| LPAR6 | lysophosphatidic acid receptor 6 | 10161 |
| LRP2 | LDL receptor related protein 2 | 4036 |
| LRRC3B | leucine rich repeat containing 3B | 116135 |
| MACROH2A2 | macroH2A.2 histone | 55506 |
| MATCAP2 | microtubule associated tyrosine carboxypeptidase 2 | 23366 |
| MIRLET7A1H( | miRlet-7a-1/let-7f-1/let-7d cluster host gene | 112903833 |
| MMP19 | matrix metallopeptidase 19 | 4327 |
| MRC2 | mannose receptor C-type 2 | 9902 |
| MSI1 | musashi RNA binding protein 1 | 4440 |
| MSRB1 | methionine sulfoxide reductase B1 | 51734 |
| MYO15B | myosin XVB | 80022 |
| NAALAD2 | N-acetylated alpha-linked acidic dipeptidase 2 | 10003 |
| NECTIN3 | nectin cell adhesion molecule 3 | 25945 |
| NGB | neuroglobin | 58157 |
| NHS | NHS actin remodeling regulator | 4810 |
| NKAIN1 | sodium/potassium transporting ATPase interacting 1 | 79570 |
| NPAS4 | neuronal PAS domain protein 4 | 266743 |
| NPPC | natriuretic peptide C | 4880 |
| NRG2 | neuregulin 2 | 9542 |
| NSUN4 | NOP2/Sun RNA methyltransferase 4 | 387338 |
| NT5C1A | 5'-nucleotidase, cytosolic IA | 84618 |
| NUDT7 | nudix hydrolase 7 | 283927 |
| NUDT8 | nudix hydrolase 8 | 254552 |
| NUTM2B-AS1 | NUTM2B antisense RNA 1 | 101060691 |
| OLFML3 | olfactomedin like 3 | 56944 |
| PAK4 | p21 (RAC1) activated kinase 4 | 10298 |
| PARP14 | poly(ADP-ribose) polymerase family member 14 | 54625 |
| PAX8-AS1 | PAX8 antisense RNA 1 | 654433 |
| PHYKPL | 5-phosphohydroxy-L-lysine phospho-lyase | 85007 |
| PIGN | phosphatidylinositol glycan anchor biosynthesis class N | 23556 |
| PIGV | phosphatidylinositol glycan anchor biosynthesis class V | 55650 |
| PLEKHO2 | pleckstrin homology domain containing O2 | 80301 |
| PLIN3 | perilipin 3 | 10226 |
| PLOD2 | procollagen-lysine,2-oxoglutarate 5-dioxygenase 2 | 5352 |

| | | |
|---|---|---|
| PLOD3 | procollagen-lysine,2-oxoglutarate 5-dioxygenase 3 | 8985 |
| POU2F2 | POU class 2 homeobox 2 | 5452 |
| PPWD1 | peptidylprolyl isomerase domain and WD repeat containing 1 | 23398 |
| PRDM10 | PR/SET domain 10 | 56980 |
| PRKCD | protein kinase C delta | 5580 |
| PRRC1 | proline rich coiled-coil 1 | 133619 |
| PTAR1 | protein prenyltransferase alpha subunit repeat containing 1 | 375743 |
| PXYLP1 | 2-phosphoxylose phosphatase 1 | 92370 |
| RALGAPA2 | Ral GTPase activating protein catalytic subunit alpha 2 | 57186 |
| RELT | RELT TNF receptor | 84957 |
| RFT1 | RFT1 glycolipid translocator homolog | 91869 |
| RFX4 | regulatory factor X4 | 5992 |
| RGN | regucalcin | 9104 |
| RGR | retinal G protein coupled receptor | 5995 |
| RNASEH1-DT | RNASEH1 divergent transcript | 100506054 |
| RNF138 | ring finger protein 138 | 51444 |
| ROBO3 | roundabout guidance receptor 3 | 64221 |
| RSPRY1 | ring finger and SPRY domain containing 1 | 89970 |
| RXFP1 | relaxin family peptide receptor 1 | 59350 |
| SBF2 | SET binding factor 2 | 81846 |
| SCAMP1-AS1 | SCAMP1 antisense RNA 1 | 728769 |
| SCARF1 | scavenger receptor class F member 1 | 8578 |
| SCARF2 | scavenger receptor class F member 2 | 91179 |
| SCLY | selenocysteine lyase | 51540 |
| SELENOP | selenoprotein P | 6414 |
| SFMBT2 | Scm like with four mbt domains 2 | 57713 |
| SHROOM3 | shroom family member 3 | 57619 |
| SLC25A48 | solute carrier family 25 member 48 | 153328 |
| SLC2A5 | solute carrier family 2 member 5 | 6518 |
| SLC35A3 | solute carrier family 35 member A3 | 23443 |
| SMAD1 | SMAD family member 1 | 4086 |
| SOCS1 | suppressor of cytokine signaling 1 | 8651 |
| SOX6 | SRY-box transcription factor 6 | 55553 |
| SOX9-AS1 | SOX9 antisense RNA 1 | 400618 |
| SPON2 | spondin 2 | 10417 |
| SPRYD4 | SPRY domain containing 4 | 283377 |
| SPTB | spectrin beta, erythrocytic | 6710 |
| SSH3 | slingshot protein phosphatase 3 | 54961 |
| STRC | stereocilin | 161497 |
| STYX | serine/threonine/tyrosine interacting protein | 6815 |
| TAF5L | TATA-box binding protein associated factor 5 like | 27097 |
| TDP1 | tyrosyl-DNA phosphodiesterase 1 | 55775 |
| TDRD7 | tudor domain containing 7 | 23424 |

| | | |
|---|---|---|
| TGFA | transforming growth factor alpha | 7039 |
| TGFB2 | transforming growth factor beta 2 | 7042 |
| TGIF1 | TGFB induced factor homeobox 1 | 7050 |
| THEMIS | thymocyte selection associated | 387357 |
| THSD7A | thrombospondin type 1 domain containing 7A | 221981 |
| TMEM241 | transmembrane protein 241 | 85019 |
| TMEM88B | transmembrane protein 88B | 643965 |
| TMX1 | thioredoxin related transmembrane protein 1 | 81542 |
| TNNT2 | troponin T2, cardiac type | 7139 |
| TPBG | trophoblast glycoprotein | 7162 |
| TRIM68 | tripartite motif containing 68 | 55128 |
| TTC38 | tetratricopeptide repeat domain 38 | 55020 |
| TTC39C | tetratricopeptide repeat domain 39C | 125488 |
| UBXN11 | UBX domain protein 11 | 91544 |
| UNC93B1 | unc-93 homolog B1, TLR signaling regulator | 81622 |
| USP42 | ubiquitin specific peptidase 42 | 84132 |
| UTP15 | UTP15 small subunit processome component | 84135 |
| VPS33A | VPS33A core subunit of CORVET and HOPS complexes | 65082 |
| ZBTB46 | zinc finger and BTB domain containing 46 | 140685 |
| ZDHHC12 | zinc finger DHHC-type palmitoyltransferase 12 | 84885 |
| ZFYVE26 | zinc finger FYVE-type containing 26 | 23503 |
| ZNF136 | zinc finger protein 136 | 7695 |
| ZNF280D | zinc finger protein 280D | 54816 |
| ZNF330 | zinc finger protein 330 | 27309 |
| ZNF394 | zinc finger protein 394 | 84124 |
| ZNF44 | zinc finger protein 44 | 51710 |
| ZNF524 | zinc finger protein 524 | 147807 |
| ZNF562 | zinc finger protein 562 | 54811 |
| ZNF574 | zinc finger protein 574 | 64763 |
| ZNF605 | zinc finger protein 605 | 100289635 |
| ZNF672 | zinc finger protein 672 | 79894 |
| ZNF708 | zinc finger protein 708 | 7562 |
| ZNF766 | zinc finger protein 766 | 90321 |
| ZNF8 | zinc finger protein 8 | 7554 |
| ZNF844 | zinc finger protein 844 | 284391 |

[S.

| Gene category | Donor 1 | | Don |
|---|---|---|---|
| | log2fc | p.adj | log2fc |
| | -1.363883411 | 1 | 1.771731012 |
| | 2.543007184 | 1 | -1.813231488 |
| | 1.221079089 | 1 | -1.228268988 |
| synaptic plasticity | 2.221079089 | 1 | -2.135159583 |
| synapse | 1.599590713 | 1 | 1.06123763 |
| | 1.028434011 | 1 | -1.398193989 |
| | 2.147078508 | 1 | 1.771731012 |
| | -1.778920911 | 1 | -1.228268988 |
| immune response | 1.899150995 | 1 | -1.813231488 |
| | -1.586275833 | 1 | -1.398193989 |
| | -1.100849005 | 1 | 1.771731012 |
| | 2.336556307 | 1 | -1.398193989 |
| synaptic plasticity/ neurogenesis | -2.100849005 | 1 | -1.135159583 |
| cell survival | 1.80604159 | 1 | 1.771731012 |
| cell survival | 1.221079089 | 1 | 1.186768512 |
| | 1.221079089 | 1 | 1.771731012 |
| | -1.132557865 | 1 | 1.186768512 |
| synapse | 2.391004091 | 1 | -1.457087678 |
| synapse | 2.095548207 | 1 | 2.186768512 |
| | -1.19395841 | 1 | 2.186768512 |
| | 1.346609971 | 1 | 1.508696606 |
| | -1.041955316 | 1 | -2.398193989 |
| myelin/ oligodendrocytes | -1.363883411 | 1 | 1.186768512 |
| | 1.921518808 | 1 | -1.398193989 |
| | 1.472617856 | 0.633399827 | -1.03562391 |
| mitochondria | 1.028434011 | 1 | -1.398193989 |
| | -1.778920911 | 1 | 1.508696606 |
| | -1.001313332 | 1 | -1.135159583 |
| | -1.363883411 | 1 | -1.135159583 |
| | -1.026848424 | 1 | 1.186768512 |
| | -1.264347738 | 1 | -1.03562391 |
| | 1.221079089 | 1 | 1.508696606 |
| synapse | 1.221079089 | 1 | -1.813231488 |
| synapse | 1.474835682 | 1 | -1.093339408 |
| | -1.363883411 | 1 | -1.398193989 |
| synapse | 1.028434011 | 1 | -1.62058641 |
| mitochondria | 1.543007184 | 1 | 2.508696606 |
| cell survival | 1.028434011 | 1 | -2.135159583 |
| | 1.391004091 | 1 | 1.771731012 |

| | | | |
|---|---|---|---|
| | -1.456992816 | 1 | -1.398193989 |
| | 1.543007184 | 1 | -2.813231488 |
| neurogenesis | -1.363883411 | 1 | 1.508696606 |
| | 1.543007184 | 1 | 1.034765418 |
| | 1.127969685 | 1 | -1.228268988 |
| | -1.456992816 | 1 | 1.186768512 |
| neurogenesis | 1.221079089 | 1 | 1.186768512 |
| | 1.80604159 | 1 | 1.409160933 |
| | 1.543007184 | 1 | 2.508696606 |
| neurogenesis | 1.069075996 | 1 | 1.508696606 |
| | 1.221079089 | 1 | -2.135159583 |
| | -1.001313332 | 1 | -1.398193989 |
| | -1.14149099 | 1 | 1.186768512 |
| | 1.80604159 | 1 | -1.813231488 |
| immune response | 1.80604159 | 1 | -1.398193989 |
| | -1.653390029 | 1 | 1.466876431 |
| | 1.127969685 | 1 | 1.409160933 |
| | -1.314973811 | 1 | 2.994123434 |
| | 1.221079089 | 1 | 1.186768512 |
| | -1.293494083 | 1 | 1.994123434 |
| myelin/ oligodendrocytes | -2.586275833 | 1 | 1.186768512 |
| | 1.958044684 | 1 | 1.409160933 |
| | 1.028434011 | 1 | 1.186768512 |
| | -1.894398128 | 1 | -1.135159583 |
| | -1.363883411 | 1 | -1.398193989 |
| | -1.363883411 | 1 | 1.771731012 |
| | 1.028434011 | 1 | -2.398193989 |
| | 1.744641045 | 1 | -1.398193989 |
| | -1.264347738 | 1 | 1.771731012 |
| | -1.586275833 | 1 | 1.508696606 |
| | 1.80604159 | 1 | -1.90069433 |
| intrinsic plasticity | 1.221079089 | 1 | -1.135159583 |
| synapse | 1.105601872 | 1 | 1.409160933 |
| | -1.19395841 | 1 | 1.186768512 |
| | 1.373082183 | 1 | -1.398193989 |
| | 1.028434011 | 1 | -2.135159583 |
| | -1.219493502 | 1 | -1.491303394 |
| | 1.346609971 | 1 | -3.272663107 |
| | 1.543007184 | 1 | 2.508696606 |
| | -1.363883411 | 1 | 1.771731012 |
| | 1.221079089 | 1 | -1.398193989 |
| | 1.221079089 | 1 | 1.601806011 |
| | -2.62327004 | 1.03E-05 | 1.245662201 |

| | | | |
|---|---|---|---|
| synaptic plasticity | 1.786676265 | 0.000102797 | -1.205548911 |
| | 1.114163885 | 1 | 1.186768512 |
| | -1.363883411 | 1 | 1.771731012 |
| | 1.127969685 | 1 | 1.508696606 |
| | 1.282479634 | 1 | -1.813231488 |
| | 1.159678545 | 1 | 1.449802917 |
| | -1.041955316 | 1 | -1.398193989 |
| | 1.391004091 | 1 | 1.508696606 |
| | -1.586275833 | 1 | -1.398193989 |
| | -1.001313332 | 1 | -1.398193989 |
| | -2.100849005 | 1 | -1.398193989 |
| | 1.009574984 | 1 | -2.398193989 |
| | 1.095548207 | 1 | -1.813231488 |
| | 1.057580357 | 1 | 1.64620013 |
| | 1.028434011 | 1 | -1.135159583 |
| | -1.948845912 | 1 | -1.398193989 |
| | -1.515886505 | 1 | -1.398193989 |
| | 1.028434011 | 1 | -1.228268988 |
| cell survival | -1.100849005 | 1 | 1.508696606 |
| | 1.221079089 | 1 | -2.398193989 |
| | -1.19395841 | 1 | 2.186768512 |
| neurogenesis/ synapse | 1.102434593 | 1 | -1.135159583 |
| | 1.80604159 | 1 | -2.228268988 |
| | 1.028434011 | 1 | 1.508696606 |
| | 1.221079089 | 1 | -1.813231488 |
| synapse | -1.18920488 | 0.305186006 | 1.104306351 |
| | -1.19395841 | 1 | 2.508696606 |
| synaptic plasticity | 1.221079089 | 1 | 1.186768512 |
| mitochondria | -1.100849005 | 1 | -1.398193989 |
| | -1.778920911 | 1 | 1.771731012 |
| | -1.100849005 | 1 | -1.813231488 |
| mitochondria | 1.391004091 | 1 | -1.398193989 |
| | 1.221079089 | 1 | 1.508696606 |
| | -1.363883411 | 1 | -2.135159583 |
| | 1.543007184 | 1 | -1.03562391 |
| | 1.636116589 | 1 | -1.298658316 |
| | 1.391004091 | 1 | 1.771731012 |
| | 1.028434011 | 1 | -1.135159583 |
| | 1.028434011 | 1 | 1.186768512 |
| | 1.028434011 | 1 | -2.398193989 |
| cell survival | 1.221079089 | 1 | 1.186768512 |
| | 1.069075996 | 1 | -1.813231488 |
| | -1.05902883 | 1 | -1.03562391 |

| | | | |
|---|---|---|---|
| | -1.001313332 | 1 | -1.228268988 |
| | 1.636116589 | 1 | -1.98315649 |
| | 1.114163885 | 1 | 2.508696606 |
| | -1.100849005 | 1 | -1.398193989 |
| | 1.114163885 | 1 | -1.398193989 |
| | 1.221079089 | 1 | 1.186768512 |
| | 1.405503661 | 1 | 1.409160933 |
| | 1.80604159 | 1 | -1.166868443 |
| | 1.114163885 | 1 | -1.398193989 |
| | 1.291468417 | 1 | -1.135159583 |
| | 1.114163885 | 1 | -1.398193989 |
| | -2.100849005 | 1 | 2.186768512 |
| intrinsic plasticity | 1.735652262 | 1 | -1.398193989 |
| | 1.221079089 | 1 | 1.771731012 |
| | 1.069075996 | 1 | 1.771731012 |
| intrinsic plasticity | 1.308541931 | 1 | 1.923734106 |
| | -1.001313332 | 1 | -1.135159583 |
| | -1.026848424 | 1 | -1.398193989 |
| | -1.083775492 | 1 | -1.03562391 |
| | 1.221079089 | 1 | 1.771731012 |
| | 2.543007184 | 1 | -1.135159583 |
| | -1.586275833 | 1 | -1.398193989 |
| | 2.680510708 | 1 | -2.398193989 |
| | -1.041955316 | 1 | 1.771731012 |
| cell survival | -1.01992901 | 1 | -1.005876566 |
| | -1.157432534 | 1 | -3.135159583 |
| | 1.373082183 | 1 | 2.356693513 |
| mitochondria | 2.028434011 | 1 | 1.186768512 |
| | 1.221079089 | 1 | -2.398193989 |
| | -1.479360629 | 1 | -1.135159583 |
| intrinsic plasticity/ synaptic plasticity | 1.083575566 | 1 | 1.409160933 |
| immune response | 1.095548207 | 1 | 2.771731012 |
| | 1.221079089 | 1 | 1.508696606 |
| | -2.100849005 | 1 | -1.813231488 |
| | 1.443471511 | 1 | -1.398193989 |
| | 1.028434011 | 1 | 1.186768512 |
| | 1.164495561 | 1 | -1.491303394 |
| | -1.100849005 | 1 | 2.508696606 |
| | -2.100849005 | 1 | -1.135159583 |
| | 1.221079089 | 1 | -1.398193989 |
| | 1.320614763 | 1 | 1.994123434 |
| | -1.778920911 | 1 | -1.135159583 |
| | 1.127969685 | 1 | 1.186768512 |

| | | |
|---|---|---|
| -1.001313332 | 1 | 1.771731012 |
| 1.958044684 | 1 | -1.398193989 |
| -1.041955316 | 1 | -1.03562391 |
| -1.866383752 | 1 | 3.356693513 |
| 2.028434011 | 1 | -1.813231488 |
| -1.363883411 | 1 | 1.771731012 |
| -1.001313332 | 1 | 1.771731012 |
| -1.026848424 | 1 | -1.228268988 |
| 1.028434011 | 1 | -1.03562391 |
| 1.715843781 | 1 | -2.398193989 |
| -1.001313332 | 1 | -1.398193989 |
| 2.028434011 | 1 | -1.398193989 |
| -1.041955316 | 1 | -1.398193989 |
| -1.252852099 | 1 | 1.771731012 |
| 1.543007184 | 1 | -1.813231488 |
| 1.221079089 | 1 | -1.398193989 |
| -1.778920911 | 1 | 2.508696606 |
| -1.015960108 | 1 | -1.491303394 |
| -1.001313332 | 1 | -1.03562391 |
| 2.028434011 | 1 | -1.813231488 |
| 1.543007184 | 1 | -1.398193989 |
| 1.543007184 | 1 | 1.186768512 |
| 1.510585707 | 1 | 1.186768512 |
| 1.095548207 | 1 | -1.135159583 |
| -1.219493502 | 1 | 1.508696606 |
| -1.238352529 | 1 | 2.508696606 |
| 1.221079089 | 1 | 1.771731012 |
| -1.479360629 | 1 | -1.03562391 |
| 1.853347305 | 1 | 1.186768512 |
| 1.484113495 | 1 | -1.398193989 |
| -1.001313332 | 1 | -1.272663107 |
| 2.028434011 | 1 | 1.186768512 |
| 1.636116589 | 1 | -2.135159583 |
| 1.069075996 | 1 | 1.771731012 |
| -1.363883411 | 1 | 1.508696606 |

or 2

| p.adj |
| --- |
| 1 |
| 1 |
| 1 |
| 1 |
| 1 |
| 1 |
| 1 |
| 1 |
| 1 |
| 1 |
| 1 |
| 1 |
| 1 |
| 1 |
| 1 |
| 1 |
| 1 |
| 1 |
| 1 |
| 1 |
| 1 |
| 1 |
| 1 |
| 1 |
| 1 |
| 1 |
| 1 |
| 1 |
| 1 |
| 1 |
| 1 |
| 1 |
| 1 |
| 1 |
| 1 |
| 1 |
| 1 |
| 1 |
| 1 |

1
1
1
1
1
1
1
1
1
1
1
1
1
1
1
1
1
1
1
1
1
1
1
1
1
1
1
1
1
1
1
1
1
1
1
1
1
1
1
1
1
1
1

1
1
1
1
1
1
1
1
1
1
1
1
1
1
1
1
1
1
1
1
1
1
1
1
1
1
1
1
1
1
1
1
1
1
1
1
1
1
1
1
1
1
1

1
1
1
1
1
1
1
1
1
1
1
1
1
1
1
1
1
1
1
1
1
1
1
1
1
1
1
1
1
1
1
1
1
1
1

**Data S11. Differentially expressed genes (DEGs) in microglia following 10 Hz rTMS.**

| Gene symbol | Gene name | EntrezID |
|---|---|---|
| ACBD7 | acyl-CoA binding domain containing 7 | 414149 |
| AGFG1 | ArfGAP with FG repeats 1 | 3267 |
| APLP1 | amyloid beta precursor like protein 1 | 333 |
| ARC | activity regulated cytoskeleton associated protein | 23237 |
| ARL1 | ARF like GTPase 1 | 400 |
| ATG4B | autophagy related 4B cysteine peptidase | 23192 |
| CAMLG | calcium modulating ligand | 819 |
| CAND1 | cullin associated and neddylation dissociated 1 | 55832 |
| CPEB4 | cytoplasmic polyadenylation element binding protein 4 | 80315 |
| CS | citrate synthase | 1431 |
| CWC15 | CWC15 spliceosome associated protein homolog | 51503 |
| DCXR | dicarbonyl and L-xylulose reductase | 51181 |
| DGUOK | deoxyguanosine kinase | 1716 |
| DPP8 | dipeptidyl peptidase 8 | 54878 |
| ELMO1 | engulfment and cell motility 1 | 9844 |
| ENY2 | ENY2 transcription and export complex 2 subunit | 56943 |
| HSPA1B | heat shock protein family A (Hsp70) member 1B | 3304 |
| IFIT1 | interferon induced protein with tetratricopeptide repeats 1 | 3434 |
| IRF9 | interferon regulatory factor 9 | 10379 |
| ITFG1 | integrin alpha FG-GAP repeat containing 1 | 81533 |
| KANSL3 | KAT8 regulatory NSL complex subunit 3 | 55683 |
| KCNIP1 | potassium voltage-gated channel interacting protein 1 | 30820 |
| KRR1 | KRR1 small subunit processome component homolog | 11103 |
| LAMTOR4 | late endosomal/lysosomal adaptor, MAPK and MTOR activator 4 | 389541 |
| LPIN1 | lipin 1 | 23175 |
| LRRTM4 | leucine rich repeat transmembrane neuronal 4 | 80059 |
| MAPK1 | mitogen-activated protein kinase 1 | 5594 |
| MAPK10 | mitogen-activated protein kinase 10 | 5602 |
| MAPK9 | mitogen-activated protein kinase 9 | 5601 |
| MAX | MYC associated factor X | 4149 |
| MVB12B | multivesicular body subunit 12B | 89853 |
| NAP1L4 | nucleosome assembly protein 1 like 4 | 4676 |
| NCBP2 | nuclear cap binding protein subunit 2 | 22916 |
| NDUFS2 | NADH:ubiquinone oxidoreductase core subunit S2 | 4720 |
| NDUFS5 | NADH:ubiquinone oxidoreductase subunit S5 | 4725 |
| NECTIN2 | nectin cell adhesion molecule 2 | 5819 |
| NSF | N-ethylmaleimide sensitive factor, vesicle fusing ATPase | 4905 |
| NTM | neurotrimin | 50863 |
| NUDC | nuclear distribution C, dynein complex regulator | 10726 |

| | | |
|---|---|---|
| OFD1 | OFD1 centriole and centriolar satellite protein | 8481 |
| OGA | O-GlcNAcase | 10724 |
| P4HTM | prolyl 4-hydroxylase, transmembrane | 54681 |
| PDCD6 | programmed cell death 6 | 10016 |
| PHF6 | PHD finger protein 6 | 84295 |
| PMP22 | peripheral myelin protein 22 | 5376 |
| PPIB | peptidylprolyl isomerase B | 5479 |
| PPP1R1A | protein phosphatase 1 regulatory inhibitor subunit 1A | 5502 |
| PRKCA | protein kinase C alpha | 5578 |
| PSMD4 | proteasome 26S subunit ubiquitin receptor, non-ATPase 4 | 5710 |
| PTGS2 | prostaglandin-endoperoxide synthase 2 | 5743 |
| RAB40B | RAB40B, member RAS oncogene family | 10966 |
| RAB5B | RAB5B, member RAS oncogene family | 5869 |
| RBX1 | ring-box 1 | 9978 |
| RNF11 | ring finger protein 11 | 26994 |
| RPH3A | rabphilin 3A | 22895 |
| SCN3B | sodium voltage-gated channel beta subunit 3 | 55800 |
| SDHB | succinate dehydrogenase complex iron sulfur subunit B | 6390 |
| SEC14L1 | SEC14 like lipid binding 1 | 6397 |
| SEPTIN3 | septin 3 | 55964 |
| SERP1 | stress associated endoplasmic reticulum protein 1 | 27230 |
| SFXN3 | sideroflexin 3 | 81855 |
| SLC7A5 | solute carrier family 7 member 5 | 8140 |
| SPART | spartin | 23111 |
| STAT1 | signal transducer and activator of transcription 1 | 6772 |
| STAT3 | signal transducer and activator of transcription 3 | 6774 |
| STIP1 | stress induced phosphoprotein 1 | 10963 |
| STX1A | syntaxin 1A | 6804 |
| STX8 | syntaxin 8 | 9482 |
| SYT4 | synaptotagmin 4 | 6860 |
| TAPBP | TAP binding protein | 6892 |
| TCF25 | TCF25 ribosome quality control complex subunit | 22980 |
| TECR | trans-2,3-enoyl-CoA reductase | 9524 |
| TGOLN2 | trans-golgi network protein 2 | 10618 |
| TMEM126B | transmembrane protein 126B | 55863 |
| TNFRSF21 | TNF receptor superfamily member 21 | 27242 |
| TRIB2 | tribbles pseudokinase 2 | 28951 |
| TRIR | telomerase RNA component interacting RNase | 79002 |
| UBASH3B | ubiquitin associated and SH3 domain containing B | 84959 |
| UBE2D2 | ubiquitin conjugating enzyme E2 D2 | 7322 |
| WDR6 | WD repeat domain 6 | 11180 |
| ZSCAN16-AS1 | ZSCAN16 antisense RNA 1 | 100129195 |

| Gene category | Donor 1 | | Donor 2 | |
|---|---|---|---|---|
| | log2fc | p.adj | log2fc | p.adj |
| | 1.415037499 | 1 | -1.847996907 | 1 |
| | -1.169925001 | 1 | 1.152003093 | 1 |
| | -1.392317423 | 1 | 1.736965594 | 1 |
| synaptic plasticity | -1.169925001 | 1 | -1.432959407 | 1 |
| | 1 | 1 | 1.473931188 | 1 |
| | 1.415037499 | 1 | -1.847996907 | 1 |
| | 1.415037499 | 1 | -3.655351829 | 1 |
| | 2 | 1 | -1.847996907 | 1 |
| | -1.584962501 | 1 | -1.432959407 | 1 |
| | -1.169925001 | 1 | 1.152003093 | 1 |
| | -1.169925001 | 1 | 1.152003093 | 1 |
| | -1.584962501 | 1 | 2.611434712 | 1 |
| mitochondria | -1.584962501 | 1 | -1.847996907 | 1 |
| | -1.169925001 | 1 | -2.847996907 | 1 |
| | 2 | 1 | -2.432959407 | 1 |
| | 2.415037499 | 1 | -1.847996907 | 1 |
| | 2.584962501 | 1 | -1.263034406 | 1 |
| immune response | 1.415037499 | 1 | -1.847996907 | 1 |
| immune response | -1.169925001 | 1 | -1.847996907 | 1 |
| | -2.392317423 | 1 | 1.152003093 | 1 |
| | 1.415037499 | 1 | -2.432959407 | 1 |
| | 1.415037499 | 1 | -1.847996907 | 1 |
| | -1.169925001 | 1 | -1.432959407 | 1 |
| | -2.169925001 | 1 | -2.432959407 | 1 |
| | 1.415037499 | 1 | -1.432959407 | 1 |
| | 1.415037499 | 1 | -1.847996907 | 1 |
| immune response | -2.169925001 | 1 | -1.263034406 | 1 |
| immune response | -1.169925001 | 1 | -2.070389328 | 1 |
| immune response | 1.415037499 | 1 | -2.847996907 | 1 |
| | 2 | 1 | -1.169925001 | 1 |
| | 1.415037499 | 1 | -1.432959407 | 1 |
| | 2 | 1 | 1.152003093 | 1 |
| | 1.415037499 | 1 | -1.847996907 | 1 |
| mitochondria | 2.736965594 | 1 | -2.432959407 | 1 |
| mitochondria | -1.459431619 | 1 | -1.625604485 | 1 |
| | 2 | 1 | -1.847996907 | 1 |
| | -1.169925001 | 1 | 1.736965594 | 1 |
| | 1.415037499 | 1 | -2.432959407 | 1 |
| | -1.169925001 | 1 | -1.432959407 | 1 |

| | | | | |
|---|---|---|---|---|
| | 1.415037499 | 1 | -1.432959407 | 1 |
| immune response | 1.415037499 | 1 | 1.473931188 | 1 |
| | -1.584962501 | 1 | 1.473931188 | 1 |
| | -2.392317423 | 1 | -1.432959407 | 1 |
| | -1.169925001 | 1 | -1.847996907 | 1 |
| | -1.906890596 | 1 | 1.736965594 | 1 |
| | -1.906890596 | 1 | -2.263034406 | 1 |
| | 2 | 1 | 1.473931188 | 1 |
| | 2.415037499 | 1 | -3.655351829 | 1 |
| | -1.169925001 | 1 | -2.847996907 | 1 |
| | -1.169925001 | 1 | 2.152003093 | 1 |
| | -2.169925001 | 1 | 1.473931188 | 1 |
| | 1.830074999 | 1 | -1.847996907 | 1 |
| | -2.169925001 | 1 | -2.655351829 | 1 |
| immune response | -1.321928095 | 1 | -1.847996907 | 1 |
| | -1.169925001 | 1 | 1.152003093 | 1 |
| | -2.392317423 | 1 | -1.847996907 | 1 |
| | 2 | 1 | -2.432959407 | 1 |
| | -1.169925001 | 1 | -1.847996907 | 1 |
| | -1.169925001 | 1 | 1.959358016 | 1 |
| | -1.169925001 | 1 | 1.152003093 | 1 |
| mitochondria | 1.415037499 | 1 | -2.432959407 | 1 |
| immune response | 2.415037499 | 1 | -3.169925001 | 1 |
| immune response | 1 | 1 | -1.847996907 | 1 |
| immune response | 1.415037499 | 1 | -2.847996907 | 1 |
| immune response | 1.415037499 | 1 | -2.432959407 | 1 |
| | 1.152003093 | 1 | 1.152003093 | 1 |
| neurotransmitter | -1.584962501 | 1 | -1.847996907 | 1 |
| | 1.415037499 | 1 | -1.847996907 | 1 |
| | -1.169925001 | 1 | 1.152003093 | 1 |
| | -2.169925001 | 1 | -1.847996907 | 1 |
| | -1.906890596 | 1 | -1.432959407 | 1 |
| | 2 | 1 | 1.152003093 | 1 |
| | -1.169925001 | 1 | 2.152003093 | 1 |
| mitochondria | 2 | 1 | 1.152003093 | 1 |
| immune response | 1.415037499 | 1 | -1.847996907 | 1 |
| | 2 | 1 | -2.432959407 | 1 |
| | -1.169925001 | 1 | -1.169925001 | 1 |
| | 1.415037499 | 1 | -1.847996907 | 1 |
| | -1.584962501 | 1 | -1.263034406 | 1 |
| | -1.584962501 | 1 | -1.847996907 | 1 |
| | 1.415037499 | 1 | 1.473931188 | 1 |

| Donor 3 | | Donor 4 | |
|---|---|---|---|
| log2fc | p.adj | log2fc | p.adj |
| -1.321928095 | 1 | -2.7744403 | 1 |
| -2.321928095 | 1 | -1.7744403 | 1 |
| -2 | 1 | -1.163005588 | 1 |
| -1.115477217 | 1 | -1.452512205 | 1 |
| -1.321928095 | 1 | -1.037474705 | 1 |
| -2.058893689 | 1 | 1.132450296 | 1 |
| -1.736965594 | 1 | -1.452512205 | 1 |
| -1.321928095 | 1 | -2.259867127 | 1 |
| 3.432959407 | 1 | -2.259867127 | 1 |
| -2.058893689 | 1 | -2.037474705 | 1 |
| 1.847996907 | 1 | -2.7744403 | 1 |
| -1.321928095 | 1 | 1.547487795 | 1 |
| 1.847996907 | 1 | -1.452512205 | 1 |
| -1.321928095 | 1 | 1.86941589 | 1 |
| 1.847996907 | 1 | -1.189477799 | 1 |
| 1.847996907 | 1 | -2.037474705 | 1 |
| 1.263034406 | 1 | -1.13058411 | 1 |
| -1.321928095 | 1 | -1.037474705 | 1 |
| 1.847996907 | 1 | 1.132450296 | 1 |
| 1.847996907 | 1 | -1.037474705 | 1 |
| 1.263034406 | 1 | -1.452512205 | 1 |
| -1.321928095 | 1 | 1.547487795 | 1 |
| -1.321928095 | 1 | -3.037474705 | 1 |
| 1.678071905 | 1 | -1.370050045 | 1 |
| 1.847996907 | 1 | -2.452512205 | 1 |
| 1.263034406 | 1 | -1.452512205 | 1 |
| -1.736965594 | 1 | -1.452512205 | 1 |
| -1.321928095 | 1 | -2.452512205 | 1 |
| -2.058893689 | 1 | -1.7744403 | 1 |
| 3.070389328 | 1 | -1.037474705 | 1 |
| -1.058893689 | 1 | -1.452512205 | 1 |
| 1.847996907 | 1 | -2.7744403 | 1 |
| 1.847996907 | 1 | -2.037474705 | 1 |
| 3.070389328 | 1 | -2.452512205 | 1 |
| 1.432959407 | 1 | -1.189477799 | 1 |
| -1.321928095 | 1 | 1.132450296 | 1 |
| -1.736965594 | 1 | -1.259867127 | 1 |
| -1.321928095 | 1 | -2.037474705 | 1 |
| 1.847996907 | 1 | 2.132450296 | 1 |

| | | | |
|---|---|---|---|
| 1.263034406 | 1 | 1.132450296 | 1 |
| -1.321928095 | 1 | -1.259867127 | 1 |
| -1.321928095 | 1 | -1.674904626 | 1 |
| -2.906890596 | 1 | -1.037474705 | 1 |
| 1.263034406 | 1 | -1.452512205 | 1 |
| -2.058893689 | 1 | -1.452512205 | 1 |
| -1.321928095 | 1 | -1.300509111 | 1 |
| 1.847996907 | 1 | -1.13058411 | 1 |
| -1.321928095 | 1 | -1.189477799 | 1 |
| -1.321928095 | 1 | -1.037474705 | 1 |
| -1.321928095 | 1 | -2.037474705 | 1 |
| 3.070389328 | 1 | -1.7744403 | 1 |
| 1.847996907 | 1 | -1.037474705 | 1 |
| -2.736965594 | 1 | -2.259867127 | 1 |
| 1.485426827 | 1 | 1.547487795 | 1 |
| 1.847996907 | 1 | -2.452512205 | 1 |
| -2.058893689 | 1 | -2.7744403 | 1 |
| 1.847996907 | 1 | -2.037474705 | 1 |
| 1.847996907 | 1 | -1.037474705 | 1 |
| -1.321928095 | 1 | -1.452512205 | 1 |
| -2.321928095 | 1 | 1.132450296 | 1 |
| 1.847996907 | 1 | -1.037474705 | 1 |
| -2.321928095 | 1 | -1.452512205 | 1 |
| 2.263034406 | 1 | -2.452512205 | 1 |
| 1.263034406 | 1 | -1.452512205 | 1 |
| 1.263034406 | 1 | -1.7744403 | 1 |
| -1.321928095 | 1 | 2.354842717 | 1 |
| -2.321928095 | 1 | -2.259867127 | 1 |
| 2.263034406 | 1 | -1.452512205 | 1 |
| -1.736965594 | 1 | -2.037474705 | 1 |
| 1.847996907 | 1 | -2.7744403 | 1 |
| -1.736965594 | 1 | -1.452512205 | 1 |
| -1.321928095 | 1 | -2.911943823 | 1 |
| -1.736965594 | 1 | -1.622437206 | 1 |
| -1.321928095 | 1 | -1.452512205 | 1 |
| 1.263034406 | 1 | 1.86941589 | 1 |
| 1.263034406 | 1 | -2.037474705 | 1 |
| -1.152003093 | 1 | -1.326981323 | 1 |
| -1.321928095 | 1 | -2.037474705 | 1 |
| -2.544320516 | 1 | 1.547487795 | 1 |
| 1.263034406 | 1 | -1.452512205 | 1 |
| 2.584962501 | 1 | -1.037474705 | 1 |

**Data S12. Differentially expressed genes (DEGs) in oligodendrocytes following 10 Hz rTMS.**

| Gene symbol | Gene name |
|---|---|
| AFF2 | ALF transcription elongation factor 2 |
| ARL13B | ARF like GTPase 13B |
| ATXN3 | ataxin 3 |
| BCL10 | BCL10 immune signaling adaptor |
| BCL7A | BAF chromatin remodeling complex subunit BCL7A |
| BCOR | BCL6 corepressor |
| BHLHE22 | basic helix-loop-helix family member e22 |
| BMP2 | bone morphogenetic protein 2 |
| BOC | BOC cell adhesion associated, oncogene regulated |
| C11orf91 | chromosome 11 open reading frame 91 |
| CA4 | carbonic anhydrase 4 |
| CACNA1H | calcium voltage-gated channel subunit alpha1 H |
| CAPN5 | calpain 5 |
| CARNMT1 | carnosine N-methyltransferase 1 |
| CCKBR | cholecystokinin B receptor |
| CDKL1 | cyclin dependent kinase like 1 |
| CHMP7 | charged multivesicular body protein 7 |
| CLCN2 | chloride voltage-gated channel 2 |
| CMPK2 | cytidine/uridine monophosphate kinase 2 |
| CNTNAP3 | contactin associated protein family member 3 |
| CSTF1 | cleavage stimulation factor subunit 1 |
| DNAJC14 | DnaJ heat shock protein family (Hsp40) member C14 |
| DYRK2 | dual specificity tyrosine phosphorylation regulated kinase 2 |
| EPB41L4B | erythrocyte membrane protein band 4.1 like 4B |
| ERMARD | ER membrane associated RNA degradation |
| FAM241A | family with sequence similarity 241 member A |
| FAS | Fas cell surface death receptor |
| GMNN | geminin DNA replication inhibitor |
| GORAB | golgin, RAB6 interacting |
| HBA2 | hemoglobin subunit alpha 2 |
| HBB | hemoglobin subunit beta |
| HTR5A | 5-hydroxytryptamine receptor 5A |
| IFT81 | intraflagellar transport 81 |
| IPMK | inositol polyphosphate multikinase |
| IQCK | IQ motif containing K |
| KATNAL1 | katanin catalytic subunit A1 like 1 |
| KBTBD3 | kelch repeat and BTB domain containing 3 |
| KIF26B | kinesin family member 26B |
| KLHL23 | kelch like family member 23 |

| | |
|---|---|
| KRI1 | KRI1 homolog |
| L3MBTL1 | L3MBTL histone methyl-lysine binding protein 1 |
| LAMB1 | laminin subunit beta 1 |
| LAMP5 | lysosomal associated membrane protein family member 5 |
| LINC01003 | long intergenic non-protein coding RNA 1003 |
| LINGO3 | leucine rich repeat and Ig domain containing 3 |
| MCM6 | minichromosome maintenance complex component 6 |
| MGAT2 | alpha-1,6-mannosyl-glycoprotein 2-beta-N-acetylglucosaminyltransferase |
| MGAT4C | MGAT4 family member C |
| MIR155HG | MIR155 host gene |
| MTF2 | metal response element binding transcription factor 2 |
| MTFMT | mitochondrial methionyl-tRNA formyltransferase |
| NEUROD1 | neuronal differentiation 1 |
| NVL | nuclear VCP like |
| PACRGL | parkin coregulated like |
| PANX1 | pannexin 1 |
| PAXIP1-DT | PAXIP1 divergent transcript |
| PBX2 | PBX homeobox 2 |
| PIGC | phosphatidylinositol glycan anchor biosynthesis class C |
| POLD3 | DNA polymerase delta 3, accessory subunit |
| PPP3CC | protein phosphatase 3 catalytic subunit gamma |
| PRKCQ-AS1 | PRKCQ antisense RNA 1 |
| PTCH2 | patched 2 |
| RAB9B | RAB9B, member RAS oncogene family |
| RAMP2 | receptor activity modifying protein 2 |
| RBMS2 | RNA binding motif single stranded interacting protein 2 |
| RERG | RAS like estrogen regulated growth inhibitor |
| RIGI | RNA sensor RIG-I |
| RIOK1 | RIO kinase 1 |
| RNF215 | ring finger protein 215 |
| RPIA | ribose 5-phosphate isomerase A |
| RTL10 | retrotransposon Gag like 10 |
| SH2B2 | SH2B adaptor protein 2 |
| SLC24A4 | solute carrier family 24 member 4 |
| SMC6 | structural maintenance of chromosomes 6 |
| SMIM17 | small integral membrane protein 17 |
| SNED1 | sushi, nidogen and EGF like domains 1 |
| SPAG5 | sperm associated antigen 5 |
| SSH3 | slingshot protein phosphatase 3 |
| TMEM132D | transmembrane protein 132D |
| TMEM64 | transmembrane protein 64 |
| TNNT2 | troponin T2, cardiac type |
| TSPAN6 | tetraspanin 6 |

| | |
|---|---|
| TSTD3 | thiosulfate sulfurtransferase like domain containing 3 |
| VRK1 | VRK serine/threonine kinase 1 |
| WIPI1 | WD repeat domain, phosphoinositide interacting 1 |
| YAP1 | Yes1 associated transcriptional regulator |
| ZBTB14 | zinc finger and BTB domain containing 14 |
| ZBTB6 | zinc finger and BTB domain containing 6 |
| ZIC1 | Zic family member 1 |
| ZNF250 | zinc finger protein 250 |
| ZNF273 | zinc finger protein 273 |
| ZNF514 | zinc finger protein 514 |
| ZNG1B | Zn regulated GTPase metalloprotein activator 1B |

| EntrezID | Gene category | Donor 3 | | Don |
|---|---|---|---|---|
| | | log2fc | p.adj | log2fc |
| 2334 | | 2.269920791 | 1 | -2.212050477 |
| 200894 | | -1.759826552 | 1 | -1.375549209 |
| 4287 | | 1.578043087 | 1 | -1.060047384 |
| 8915 | | 1.784493964 | 1 | -1.185578266 |
| 605 | | -1.537434131 | 1 | 1.151697793 |
| 54880 | | 1.04752837 | 1 | 2.016768213 |
| 27319 | | -2.859362226 | 1 | -1.153156788 |
| 650 | | 1.04752837 | 1 | 1.247381142 |
| 91653 | myelin | 1.269920791 | 1 | 1.984346736 |
| 100131378 | | 2.269920791 | 1 | 1.109877618 |
| 762 | | 2.04752837 | 1 | -1.153156788 |
| 8912 | | -1.274399725 | 1 | -1.212050477 |
| 726 | | -3.344789053 | 1 | -1.0055996 |
| 138199 | | -1.759826552 | 1 | -1.243759337 |
| 887 | | -1.122396631 | 1 | -1.015653264 |
| 8814 | | -1.537434131 | 1 | -2.349554001 |
| 91782 | | 1.04752837 | 1 | -1.381975479 |
| 1181 | cellular differentiation | 1.04752837 | 1 | 1.372912024 |
| 129607 | | -1.122396631 | 1 | -1.5905621 |
| 79937 | | -1.759826552 | 1 | 1.694840118 |
| 1477 | | -3.122396631 | 1 | -1.08276746 |
| 85406 | | 1.269920791 | 1 | -1.027625906 |
| 8445 | | 1.269920791 | 1 | -1.305159882 |
| 54566 | | -1.537434131 | 1 | -1.0055996 |
| 55780 | | 1.04752837 | 1 | -1.112514804 |
| 132720 | | -1.759826552 | 1 | 1.332270039 |
| 355 | cell death | -1.274399725 | 1 | 1.431805713 |
| 51053 | | 1.04752837 | 1 | -1.112514804 |
| 92344 | | 2.04752837 | 1 | -1.268634005 |
| 3040 | | -1.047628863 | 1 | 1.485244972 |
| 3043 | | -1.601564468 | 1 | 1.31846424 |
| 3361 | | -2.122396631 | 1 | -1.027625906 |
| 28981 | | -2.537434131 | 1 | 1.694840118 |
| 253430 | | -2.122396631 | 1 | 1.197340459 |
| 124152 | | -1.707359132 | 1 | 1.846843212 |
| 84056 | | -1.385431037 | 1 | -1.08276746 |
| 143879 | | 1.199531464 | 1 | -1.060047384 |
| 55083 | | 1.04752837 | 1 | -1.027625906 |
| 151230 | | -1.274399725 | 1 | -1.091756243 |

| | | | | |
|---|---|---|---|---|
| 65095 | | -2.537434131 | 1 | 1.016768213 |
| 26013 | synaptic plasticity | 1.784493964 | 1 | -1.349554001 |
| 3912 | cellular differentiation | 2.269920791 | 1 | -1.697477304 |
| 24141 | synapse | -1.730079209 | 1 | -1.428542297 |
| 100128822 | | -1.859362226 | 1 | -1.189682664 |
| 645191 | | -1.859362226 | 1 | -2.060047384 |
| 4175 | | -1.344789053 | 1 | 1.18788013 |
| 4247 | | -1.915945754 | 1 | -1.170230301 |
| 25834 | | 1.784493964 | 1 | -1.0055996 |
| 114614 | | -2.122396631 | 1 | 1.048477073 |
| 22823 | | 1.462565869 | 1 | -1.0055996 |
| 123263 | mitochondria | 2.04752837 | 1 | 1.002962414 |
| 4760 | | -1.537434131 | 1 | -1.212050477 |
| 4931 | | 1.199531464 | 1 | -1.179628999 |
| 133015 | | 1.632490871 | 1 | -1.330694974 |
| 24145 | synaptic plasticity | -1.344789053 | 1 | -1.212050477 |
| 202781 | | 1.269920791 | 1 | -1.627087976 |
| 5089 | | 1.462565869 | 1 | -1.112514804 |
| 5279 | | -1.537434131 | 1 | 1.109877618 |
| 10714 | | -1.537434131 | 1 | -1.627087976 |
| 5533 | | 1.269920791 | 1 | 1.209413291 |
| 439949 | | 1.199531464 | 1 | 1.399384235 |
| 8643 | | -1.022860958 | 1 | 1.247381142 |
| 51209 | | -1.759826552 | 1 | 1.109877618 |
| 10266 | | -1.537434131 | 1 | -1.112514804 |
| 5939 | | -2.537434131 | 1 | -1.153156788 |
| 85004 | | -1.537434131 | 1 | -1.697477304 |
| 23586 | | 1.04752837 | 1 | -1.212050477 |
| 83732 | | -2.344789053 | 1 | 1.109877618 |
| 200312 | | 1.784493964 | 1 | -1.212050477 |
| 22934 | | 2.269920791 | 1 | 1.016768213 |
| 79680 | | -1.122396631 | 1 | -1.252692462 |
| 10603 | | -2.344789053 | 1 | -2.212050477 |
| 123041 | | 1.269920791 | 1 | -1.475084883 |
| 79677 | | 1.04752837 | 1 | 1.91723254 |
| 147670 | | -2.122396631 | 1 | -1.375549209 |
| 25992 | | -1.052007303 | 1 | -1.112514804 |
| 10615 | | 2.921997488 | 1 | 1.109877618 |
| 54961 | | -1.859362226 | 1 | 1.524915117 |
| 121256 | | -2.537434131 | 1 | -1.08276746 |
| 169200 | | 1.462565869 | 1 | 1.753733808 |
| 7139 | | -1.95247163 | 1 | -1.627087976 |
| 7105 | immune response | 1.269920791 | 1 | 1.027415458 |

| | | | | |
|---|---|---|---|---|
| 100130890 | | 1.04752837 | 1 | 1.332270039 |
| 7443 | | -2.122396631 | 1 | -1.027625906 |
| 55062 | | -3.344789053 | 1 | 2.279802619 |
| 10413 | cellular differentiation | 1.199531464 | 1 | 1.060968017 |
| 7541 | | -1.537434131 | 1 | 2.109877618 |
| 10773 | | -1.274399725 | 1 | -1.060047384 |
| 7545 | | 1.462565869 | 1 | -1.475084883 |
| 58500 | | 1.04752837 | 1 | 1.694840118 |
| 10793 | | -2.122396631 | 1 | -1.0055996 |
| 84874 | | 1.269920791 | 1 | 1.302522696 |
| 150472 | | -1.537434131 | 1 | -1.330694974 |

ıor 4

| p.adj |
| --- |
| 1 |
| 1 |
| 1 |
| 1 |
| 1 |
| 1 |
| 1 |
| 1 |
| 1 |
| 1 |
| 1 |
| 1 |
| 1 |
| 1 |
| 1 |
| 1 |
| 1 |
| 1 |
| 1 |
| 1 |
| 1 |
| 1 |
| 1 |
| 1 |
| 1 |
| 1 |
| 1 |
| 1 |
| 1 |
| 1 |
| 1 |
| 1 |
| 1 |
| 1 |
| 1 |
| 1 |
| 1 |
| 1 |
| 1 |

| |
|---|
| 1 |
| 1 |
| 1 |
| 0.318784191 |
| 1 |
| 1 |
| 1 |
| 1 |
| 1 |
| 1 |
| 1 |
| 1 |
| 1 |
| 1 |
| 1 |
| 1 |
| 1 |
| 1 |
| 1 |
| 1 |
| 1 |
| 1 |
| 1 |
| 1 |
| 1 |
| 1 |
| 1 |
| 1 |
| 1 |
| 1 |
| 1 |
| 1 |
| 1 |
| 1 |
| 1 |
| 1 |
| 1 |
| 1 |
| 1 |
| 1 |
| 1 |
| 1 |
| 1 |

1
1
1
1
1
1
1
1
1
1
1

**Data S13. Differentially expressed genes (DEGs) in astrocytes following 10Hz.**

| Gene symbol | Gene name | EntrezID |
|---|---|---|
| ABHD14B | abhydrolase domain containing 14B | 84836 |
| ADAM19 | ADAM metallopeptidase domain 19 | 8728 |
| ADAMTS8 | ADAM metallopeptidase with thrombospondin type 1 motif 8 | 11095 |
| AFAP1L2 | actin filament associated protein 1 like 2 | 84632 |
| ALMS1 | ALMS1 centrosome and basal body associated protein | 7840 |
| AS3MT | arsenite methyltransferase | 57412 |
| CEPT1 | choline/ethanolamine phosphotransferase 1 | 10390 |
| CFAP210 | cilia and flagella associated protein 210 | 129881 |
| CSF3 | colony stimulating factor 3 | 1440 |
| ECD | ecdysoneless cell cycle regulator | 11319 |
| ENGASE | endo-beta-N-acetylglucosaminidase | 64772 |
| FAM222B | family with sequence similarity 222 member B | 55731 |
| GABPA | GA binding protein transcription factor subunit alpha | 2551 |
| GDAP2 | ganglioside induced differentiation associated protein 2 | 54834 |
| GLIS3 | GLIS family zinc finger 3 | 169792 |
| GRM7 | glutamate metabotropic receptor 7 | 2917 |
| HAS1 | hyaluronan synthase 1 | 3036 |
| HECTD3 | HECT domain E3 ubiquitin protein ligase 3 | 79654 |
| INTS2 | integrator complex subunit 2 | 57508 |
| KATNA1 | katanin catalytic subunit A1 | 11104 |
| KCNQ1OT1 | KCNQ1 opposite strand/antisense transcript 1 | 10984 |
| KIF26B | kinesin family member 26B | 55083 |
| KLF11 | KLF transcription factor 11 | 8462 |
| LOH12CR2 | loss of heterozygosity on chromosome 12, region 2 | 503693 |
| MEX3D | mex-3 RNA binding family member D | 399664 |
| MINDY1 | MINDY lysine 48 deubiquitinase 1 | 55793 |
| MSANTD2 | Myb/SANT DNA binding domain containing 2 | 79684 |
| MTIF2 | mitochondrial translational initiation factor 2 | 4528 |
| NEPRO | nucleolus and neural progenitor protein | 25871 |
| NSUN6 | NOP2/Sun RNA methyltransferase 6 | 221078 |
| NUP54 | nucleoporin 54 | 53371 |
| PACRG | parkin coregulated | 135138 |
| PLEKHF2 | pleckstrin homology and FYVE domain containing 2 | 79666 |
| PLGLB1 | plasminogen like B1 | 5343 |
| PSMB8 | proteasome 20S subunit beta 8 | 5696 |
| REXO4 | REX4 homolog, 3'-5' exonuclease | 57109 |
| RPGR | retinitis pigmentosa GTPase regulator | 6103 |
| SAPCD2 | suppressor APC domain containing 2 | 89958 |
| SHROOM1 | shroom family member 1 | 134549 |

| | | |
|---|---|---|
| SSTR3 | somatostatin receptor 3 | 6753 |
| SYT6 | synaptotagmin 6 | 148281 |
| TMEM260 | transmembrane protein 260 | 54916 |
| TUT1 | terminal uridylyl transferase 1, U6 snRNA-specific | 64852 |
| UFL1 | UFM1 specific ligase 1 | 23376 |
| XRCC1 | X-ray repair cross complementing 1 | 7515 |
| ZFY | zinc finger protein Y-linked | 7544 |
| ZNF615 | zinc finger protein 615 | 284370 |
| ZNF621 | zinc finger protein 621 | 285268 |
| ZNF865 | zinc finger protein 865 | 100507290 |

| Gene category | Donor 1 | | Don |
|---|---|---|---|
| | log2fc | p.adj | log2fc |
| | -2.197939378 | 1 | 1.077390033 |
| immune/ inflammatory response | 1.971985624 | 1 | -1.244538062 |
| | -1.487445995 | 1 | 1.133973561 |
| | 1.387023123 | 1 | -1.566466157 |
| | 2.387023123 | 1 | 1.755461938 |
| | -1.349942471 | 1 | 1.077390033 |
| | -1.291048782 | 1 | 1.077390033 |
| | 1.387023123 | 1 | -1.092534969 |
| | 1.609415544 | 1 | -1.829500563 |
| | -1.12755005 | 1 | -1.382041586 |
| | -1.612976877 | 1 | 1.077390033 |
| | 1.194378045 | 1 | 1.56281686 |
| | -1.612976877 | 1 | 1.340424439 |
| | -1.612976877 | 1 | 2.340424439 |
| | -3.197939378 | 1 | 1.755461938 |
| | 1.609415544 | 1 | -3.051892984 |
| immune/ inflammatory response | -1.028014376 | 1 | 1.340424439 |
| | 1.123988717 | 1 | -2.051892984 |
| | 1.194378045 | 1 | 1.077390033 |
| | 1.194378045 | 1 | 1.755461938 |
| | 1.708951218 | 1 | -1.566466157 |
| | 1.708951218 | 1 | -1.244538062 |
| immune/ inflammatory response | -1.098403704 | 1 | -1.244538062 |
| | 1.387023123 | 1 | 1.340424439 |
| | 1.194378045 | 1 | 1.340424439 |
| | -1.028014376 | 1 | 1.340424439 |
| | 1.387023123 | 1 | -1.829500563 |
| mitochondria | 1.556948125 | 1 | -1.566466157 |
| | 1.293913719 | 1 | 1.018496344 |
| | -1.028014376 | 1 | -1.244538062 |
| | -1.098403704 | 1 | -1.566466157 |
| | -1.934904972 | 1 | 1.755461938 |
| | 1.123988717 | 1 | -1.092534969 |
| | -1.612976877 | 1 | 1.340424439 |
| | -1.028014376 | 1 | 2.56281686 |
| | 2.556948125 | 1 | 1.170499437 |
| | 1.971985624 | 1 | -1.244538062 |
| | 1.387023123 | 1 | 1.077390033 |
| | 1.123988717 | 1 | -1.829500563 |

| | | | |
|---|---|---|---|
| immune/ inflammatory response | -1.612976877 | 1 | 1.340424439 |
| | -1.028014376 | 1 | 1.340424439 |
| | -1.197939378 | 1 | 2.077390033 |
| | -1.197939378 | 1 | -1.829500563 |
| immune/ inflammatory response | -1.291048782 | 1 | 1.077390033 |
| | -1.028014376 | 1 | -1.382041586 |
| | -1.028014376 | 1 | -1.829500563 |
| | -1.028014376 | 1 | 1.755461938 |
| | 1.194378045 | 1 | 1.340424439 |
| | -1.349942471 | 1 | -2.244538062 |

| ior 2 | Donor 4 | |
|---|---|---|
| p.adj | log2fc | p.adj |
| 1 | 1.241481943 | 1 |
| 1 | -1.896021581 | 1 |
| 1 | -1.398521921 | 1 |
| 1 | 1.519015918 | 1 |
| 1 | 2.425906514 | 1 |
| 1 | -1.770490699 | 1 |
| 1 | 1.103978419 | 1 |
| 1 | 1.103978419 | 1 |
| 1 | -1.612228615 | 1 |
| 1 | -1.033525105 | 1 |
| 1 | -1.480984082 | 1 |
| 1 | -1.3554532 | 1 |
| 1 | 1.103978419 | 1 |
| 1 | -1.3554532 | 1 |
| 1 | -1.065946582 | 1 |
| 1 | -1.596461299 | 1 |
| 1 | -1.703376503 | 1 |
| 1 | -1.3554532 | 1 |
| 1 | 1.273903421 | 1 |
| 1 | 1.425906514 | 1 |
| 1 | -1.480984082 | 1 |
| 1 | -1.217949676 | 1 |
| 1 | 1.68894092 | 1 |
| 1 | 1.219455637 | 1 |
| 1 | 1.103978419 | 1 |
| 1 | -1.31105908 | 1 |
| 1 | 2.103978419 | 1 |
| 1 | 1.482490042 | 1 |
| 1 | -1.011498798 | 1 |
| 1 | -1.480984082 | 1 |
| 1 | 1.021516259 | 1 |
| 1 | 1.273903421 | 1 |
| 1 | -1.480984082 | 1 |
| 1 | -1.011498798 | 1 |
| 1 | 1.103978419 | 1 |
| 1 | -1.632987175 | 1 |
| 1 | -1.118414002 | 1 |
| 1 | -1.065946582 | 1 |
| 1 | -1.480984082 | 1 |

| | | |
|---|---|---|
| 1 | -1.480984082 | 1 |
| 1 | 1.32637084 | 1 |
| 1 | -1.3554532 | 1 |
| 1 | -1.896021581 | 1 |
| 1 | -2.118414002 | 1 |
| 1 | 1.296623497 | 1 |
| 1 | 2.103978419 | 1 |
| 1 | 1.103978419 | 1 |
| 1 | -2.802912177 | 1 |
| 1 | 1.32637084 | 1 |

**Data S14. Raw patch-clamp electrophysiology data of intrinsinc neuronal properies of hu**

| Cell ID | Donor ID | Donor # | Treatment | Input Resistance (MΩ) | RMP (mV) |
|---|---|---|---|---|---|
| 5422002_C1 | 5422002 | Donor 1 | iTBS | 193.09 | -54 |
| 5422002_C4 | 5422002 | Donor 1 | iTBS | 99.08 | -59 |
| 5422002_C7 | 5422002 | Donor 1 | iTBS | 52.91 | -74 |
| 5422003_C3 | 5422003 | Donor 2 | iTBS sham | 61.35 | -64.2 |
| 5422005_C4 | 5422005 | Donor 3 | iTBS | 176.33 | -69.7 |
| 5422005_C2 | 5422005 | Donor 3 | iTBS sham | 71.15 | -69.7 |
| 5422005_C3 | 5422005 | Donor 3 | iTBS sham | 86.56 | -74.7 |
| 5422010_C1 | 5422010 | Donor 4 | iTBS sham | 68.6 | -61.6 |
| 5422010_C4 | 5422010 | Donor 4 | iTBS | 153.03 | -66.8 |
| 5422010_C3 | 5422010 | Donor 4 | iTBS sham | 58.54 | -65.6 |

**ıman layer 2/3 pyramidal neurons recorded 3 h following iTBS or iTBS sham stimulation.**

| Rheobase (pA) | AP threshold (mV) | AP half-width (ms) | Max firing frequency (Hz) |
|---|---|---|---|
| 50 | -51.8 | 0.453 | 46 |
| 50 | -38.8 | 0.784 | 37 |
| 175 | -37.5 | 0.486 | 56 |
| 100 | -45.3 | 1.36 | 24 |
| 50 | -46.15 | 0.933 | 37 |
| 100 | -34.7 | 0.783 | 22 |
| 125 | -46.28 | 1.095 | 21 |
| - | - | - | - |
| 125 | -33.4 | 1.17 | 21 |
| 75 | -42.9 | 0.8484 | 34 |

**Data S15. Summary of human spatial transcriptomics samples (n = 16).**

| Donor ID | Donor # | Age | Gender | Cortical region | Treatment |
|---|---|---|---|---|---|
| 5422002 | Donor 1 | 20 | Male | Right Temporal | iTBS<br>iTBS sham<br>10Hz<br>10Hz sham |
| 5422003 | Donor 2 | 44 | Male | Right Frontal | iTBS<br>iTBS sham<br>10Hz<br>10Hz sham |
| 5422005 | Donor 3 | 68 | Male | Right Occipital | iTBS<br>iTBS sham<br>10Hz<br>10Hz sham |
| 5422010 | Donor 4 | 40 | Male | Left Parietal | iTBS<br>iTBS sham<br>10Hz<br>10Hz sham |

| Visium slide number | Visium area number | Number of spots under tissue |
|---|---|---|
| V13Y10-013 | A1 | 1327 |
| V13Y10-013 | B1 | 376 |
| V13Y10-013 | C1 | 1439 |
| V13Y10-013 | D1 | 1711 |
| V13Y10-015 | A1 | 1445 |
| V13Y10-015 | B1 | 254 |
| V13Y10-015 | C1 | 1212 |
| V13Y10-015 | D1 | 935 |
| V13Y10-012 | A1 | 1286 |
| V13Y10-012 | B1 | 1087 |
| V13Y10-012 | C1 | 512 |
| V13Y10-012 | D1 | 569 |
| V13Y10-014 | A1 | 1541 |
| V13Y10-014 | B1 | 1743 |
| V13Y10-014 | C1 | 1768 |
| V13Y10-014 | D1 | 1929 |

| Total number of reads | Mean reads per spot | Median UMIs per spot |
|---|---|---|
| 432290334 | 325765 | 2949 |
| 316758537 | 842443 | 3779 |
| 353004929 | 245313 | 1778 |
| 361871580 | 211497 | 1622 |
| 380458355 | 263293 | 1030 |
| 289625380 | 1140257 | 1530 |
| 286856187 | 236680 | 1111 |
| 305013554 | 326218 | 1146 |
| 357038663 | 277635 | 1323 |
| 294685734 | 271100 | 484 |
| 346796148 | 677336 | 586 |
| 401156556 | 705020 | 740 |
| 694989459 | 450999 | 2608 |
| 712790050 | 408944 | 2255 |
| 679788366 | 384496 | 2533 |
| 626391387 | 324723 | 2055 |

| Median genes per spot | Number of spots under tissue (after QC filtering) |
|---|---|
| 1406 | 1319 |
| 1761 | 372 |
| 852 | 1414 |
| 860 | 1710 |
| 679 | 1443 |
| 868 | 251 |
| 710 | 1208 |
| 742 | 934 |
| 818 | 1286 |
| 345 | 1087 |
| 388 | 511 |
| 443 | 569 |
| 1322 | 1509 |
| 1216 | 1686 |
| 1328 | 1761 |
| 1105 | 1925 |
